# A Macro-scale Comparison Algorithm for Analysis of TCR Repertoire Completeness

**DOI:** 10.1101/2021.03.07.434284

**Authors:** Fernando Esponda, Petr Šulc, Joseph Blattman, Stephanie Forrest

**Affiliations:** Department of Computer Science, ITAM, Mexico City, Mexico, 01080; Biodesign Center for Biocomputation, Security and Society, Arizona State University, Tempe, AZ, 85287; Biodesign Center for Molecular Design and Biomimetics, Arizona State University, Tempe, AZ, 85287; Biodesign Center for Immunotherapy, Vaccines and Virotherapy, Arizona State University, Tempe, AZ, 85287

## Abstract

Recent advances in biotechnology are beginning to generate whole *immunome* datasets, which will enable the comparison of immune repertoires between individuals, e.g., to assess immunocompetence. Existing algorithms cluster cell types based on the relative expression abundance of about 20 000 genes, but such algorithms have limited utility when comparing immunome datasets with many higher orders of magnitude (>10^12^) of variation, such as occurs in immunoreceptor sequences in highly polyclonal naive repertoires.

In this paper we present a method for comparing immune repertoires by identifying macro-level features that are conserved between similar individuals. Our method allows us to detect some blind spots in naive populations and to assess whether a repertoire is likely complete by examining only a sample of its sequences.

**Author Summary:** In this paper we present a method for comparing the immune repertoire of different individuals. Repertoires are represented by a sample of genetic sequences. Our technique coarse grains each individual’s data into groups, matches groups between individual’s and finds significant differences.

## Introduction

Comparison of large data sets is a mainstay of modern biology, and it has improved our understanding of the relationships among molecules, cells, and systems. As tools for acquiring information on biological systems have scaled up, however, new analysis methods are required to draw conclusions of biological significance. Here we consider the problem of analyzing highly diverse immunoreceptor repertoires at the macro-scale, showing how to analyze repertoire datasets to gain insight about the completeness, or immunocompetence, of an individual during immune reconstitution (e.g., that occurs during bone marrow reconstitution) or immune decline (e.g., during aging). The prospect of whole immunome studies, which can shed light on the repertoire’s role in immunotherapy treatments and on how the repertoire changes over time, has many important implicatins. For example, such studies could improve our ability to predict a patient’s response to therapy [1], and they open up new possibilities for personalized medicine and novel vaccination strategies.

In this paper, we focus on the following questions: How to determine if a repertoire is complete based on an incomplete sample of its sequences? How to compare two repertoires based on incomplete sampling of both? We address these questions by identifying the macro-level structure of repertoires, using clustering methods to group related sequences, and then comparing and contrasting clusters between repertoires.

Whole-repertoire comparisons are challenging using existing methods because of the high heterogeneity of such systems. Diversity in TCR is generated via a multi-step semi-random process of selection of variable (V), diversity (D) and joining (J) gene segments, random cleavage of DNA hairpins to generate palindromic sequences (p-nucleotides), and addition of non-templated nucleotides (n-nucleotides).

Estimates of total TCR diversity in a single individual are many orders of magnitude greater than the total number of cells in any individual [2–4]. Thus, even in genetically identical clones with identical life histories, no two individuals express the same naive T cell receptor repertoire; moreover, since the repertoire changes over time, even a single individual does not express exactly the same sequences throughout its lifetime. This issue has new urgency relevant to our SARS-CoV-2 infections, where improved understanding of a repertoire’s macro-scale structure and how it changes over time could inform which individuals would most benefit from a given vaccine. In other diseases, vaccine longevity is highly variable, although the reasons are not well understood [5], and improved understanding of repertoire dynamics could provide insight. These complexities point to the need for a robust framework for analyzing highly diverse polyclonal BCR and TCR repertoires, which allows comparison among individuals in a population and conclusions regarding immunocompetence of individuals.

Historically, researchers have assessed immune diversity either by analyzing TCR-Vb usage distributions [3], i.e. immunospectratyping, or by studying the Gaussian distribution of CDR3 lengths within a given V gene segment population [6, 7]. Such analyses can provide information about how skewed a distribution is from a normal state, but they do not provide information about the total diversity within a given immune population. More recently, both TCR and BCR repertoires have been studied quantitatively by calculating the probability of particular sequences being generated by the V(D)J recombination [8–11]. Other research has developed metrics for comparing the “proximity” of TCRs based on their similarity using a weighted sum of mismatches between aligned TCR sequences and used this approach to assign sequences to different clusters [12]. Other work has used clustering methods to detect new TCR sequences that emerge in patients after vaccination by looking for new TCR sequences that are more enriched in the cluster than what would be expected with a null model [13]. Finally, clustering methods have been applied to TCR sequences to characterize human and mouse repertoires and found several *public* TCR sequences that are shared by different individuals [8]. This earlier work, however, focuses on individual sequences to compare repertoires. Here, we focus on clusters of sequences which lets us better compare repertoires by abstracting away particular details from individual sequences.

In this paper we propose the Repertoire Comparison Algorithm (RCA) for macro-scale comparisons of TCR repertoire samples between individuals. Because only a few sequences are shared between any two individuals, exact sequence comparisons provide limited information. Our approach enables comparison of entire repertoires between different individuals (or possibly repertoire of same individual at different times) that does not rely on identifying shared sequences. Instead, RCA first identifies important clusters in each individual and then finds their correspondences. Such an approach allows us to identify missing clusters between two sequence datasets.

Our clustering algorithm generates a coarse-grained representation of a repertoire by grouping similar sequences into clusters. Because the clusters are derived from a sample of sequences, several factors can affect which clusters are identified, including: the sampling technique, the size of the sample, and the individual’s repertoire. We hypothesize that meaningful similarities and differences between individual repertoires can be discovered by comparing their clusters and controlling for the size of the sample. In particular, we look for common clusters across individuals, which we refer to as *public clusters*, illustrated in Fig.1. We examine differences in the distributions of these clusters to determine similarities between repertoires and to detect anomalies, such as the existence of unexpected groups of sequences or the absence of anticipated clusters.

**Fig. 1.**
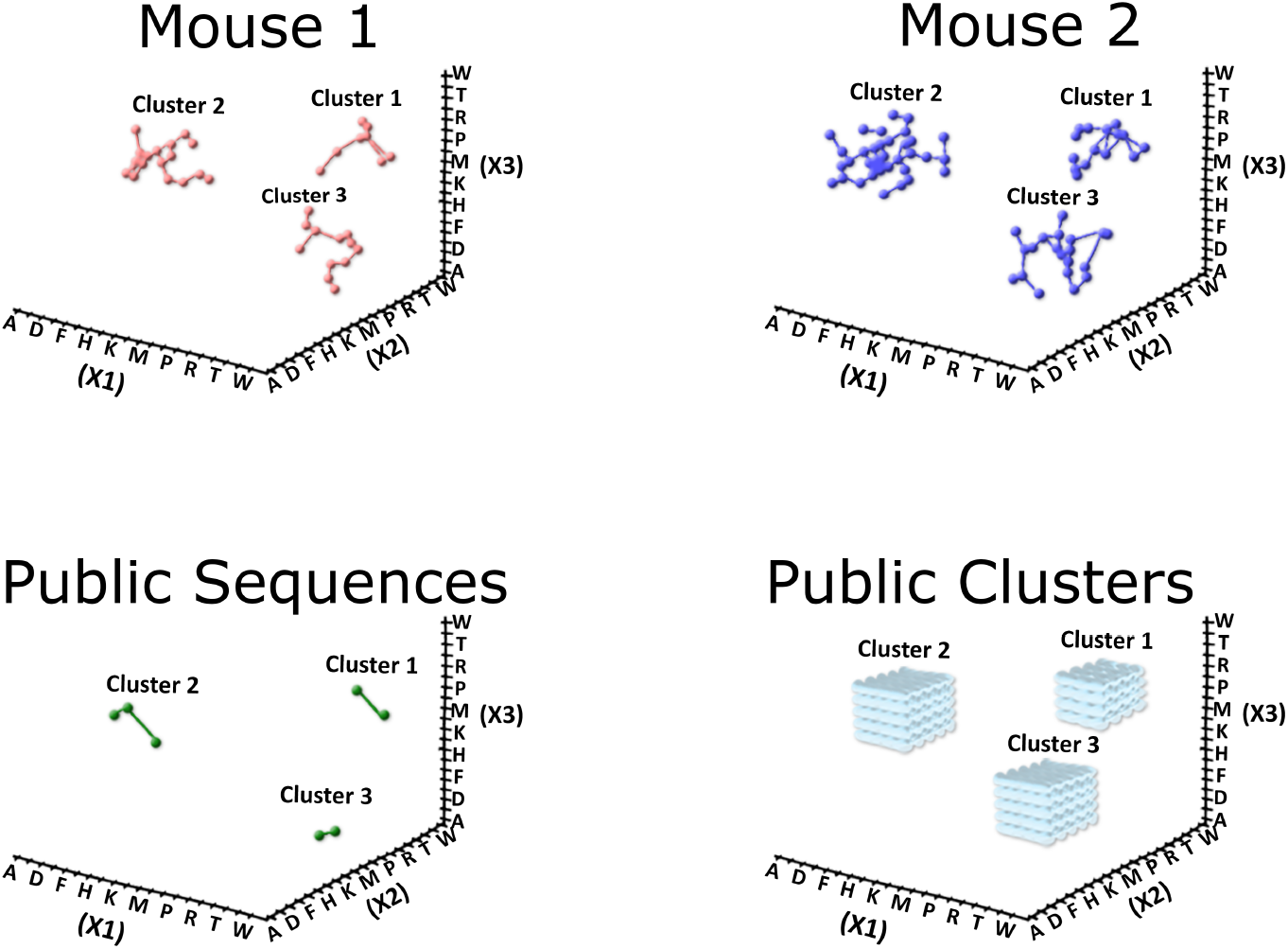
Schematic illustration of hypothetical macro-structure generated by RCA, illustrated for three sequence positions *X*_1_, *X*_2_, and *X*_3_. Each axis corresponds to one sequence, and each position on each axis denotes a possible amino acid. Panels (a) and (b) show sequences for two hypothetical repertoires and how they are grouped into three clusters. Panel (c) highlights the sequences that a hypothetical Mouse 1 and Mouse 2 have in common. Panel (d) shows the public clusters—all the possible sequences in the regions from which different individuals draw their repertoires.

Applying RCA to mice TCR repertoires suggests that for similar individuals the distribution of clusters (in terms of number of clusters and number of sequences in each cluster) is remarkably consistent. We also generated synthetic repertoires that are significantly larger than the mouse data we have, finding that clusters from the mouse data appear in the large synthetic repertoires with similar relative sizes but that some clusters in the synthetic repertoires are not found in the mice datasets. A potential application of our method is detecting interesting differences, or anomalies, between datasets. We test this by systematically deleting sequences from a repertoire and observing how it affects the clusters and their distributions.

## Results

We developed the RCA software *(github link will be included here for the publication)* and used it to study the macro-structure of a single repertoire under different similarity matching rules for quantifying the similarity of two sequences. Next, we compared pairs of repertoires, showing how to identify corresponding clusters to identify high-level commonalities between repertoires. Finally, we used the clustering approach for comparing repertoires to determine if a repertoire is complete, showing how our method can be leveraged to detect anomalies in repertoire.

### Clustering the repertoire

We implemented our software using the DBSCAN clustering algorithm (see Methods Sect. Clustering) and tested it on both synthetic and experimental data (Sect. Data Description). The synthetic dataset was obtained by simulating the V(D)J recombination process using the OLGA software tool [9], and the experimental dataset was obtained from CD8 sorted cells from C57BL/6 mouse spleens

We considered two metrics for assessing the difference between sequences, Levenshtein and Hamming distance, and found that Hamming distance is more appropriate for comparing repertoires. Although Levenshtein distance is a common choice for generating mutation networks, we found that its restricted version, Hamming distance, provides a better solution for our course-grained approach because it leads to a larger number of clusters for a given sample size, which in turn allows us to more accurately detect differences between repertoire samples (Fig. 2).

**Fig. 2.**
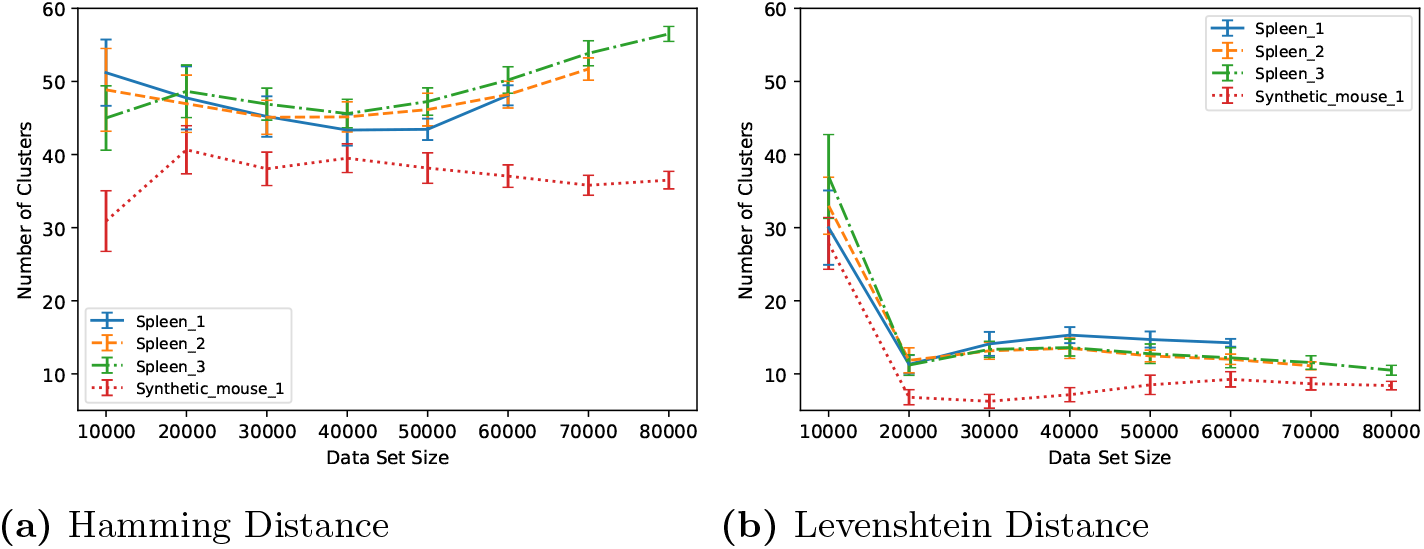
Comparison of distance metrics. Panels (a) and (b) show results for Hamming Distance and Levenshtein Distance respectively. The *y* axis reports the number of clusters identified by RCA using the distance metric on a dataset sample of size *x*. Dataset size refers to the number of distinct amino acid sequences in the dataset. Each data point is the average of 20 runs of RCA on a random sample of *x* sequences selected from the dataset indicated in the legend.

To study the effect of dataset size, we sampled subsets of the murine datasets. Figure 2 shows how the number of clusters changes as the size of the data set increases for Hamming and Levenshtein distances. Clusters that contain fewer than 0.1% of the total sampled sequences are not reported (Fig. 2). Thus, not all sequences in a sample are assigned to a cluster. The exact threshold value is not crucial for our results, which are not sensitive to the exact threshold used, although the running time of the algorithm is strongly affected by the threshold value.

Figure 2 shows that Hamming Distance is more sensitive than Levenshtein and can identify more clusters. This discrepancy is more pronounced for larger datasets. Hamming Distance identifies nearly twice as many clusters for small datasets and three times as many for larger datasets. This is expected because Hamming Distance is more sensitive than Levenshtein, in the sense that the number of possible sequences that can match a given sequence is much smaller for Hamming than for Levenshtein. The figure also shows that the number of clusters found with Hamming Distance generally increases with dataset size in the murine examples, while the number of clusters levels off quickly with Levenshtein Distance.

We also find that RCA produces fewer clusters in the synthetic datasets compared to murine data for the same sample sizes (Fig. 2). However as shown in the next section larger synthetic data sizes do exhibit all the clusters of smaller size murine data (Fig. 3b). Synthetic data were produced from naive repertoire sequences using OLGA software tool [9], which accounts for V(D)J recombination probabilities and does not account for thymic selection that occurs thereafter. We conjecture that a larger synthetic dataset is required for it to exhibit a similar number of clusters than a smaller murine sample because the murine data is the result a larger pool from which some sequences were removed by thymic selection. It is in the larger naive sample where there are enough similar sequences for a cluster identification.

**Fig. 3.**
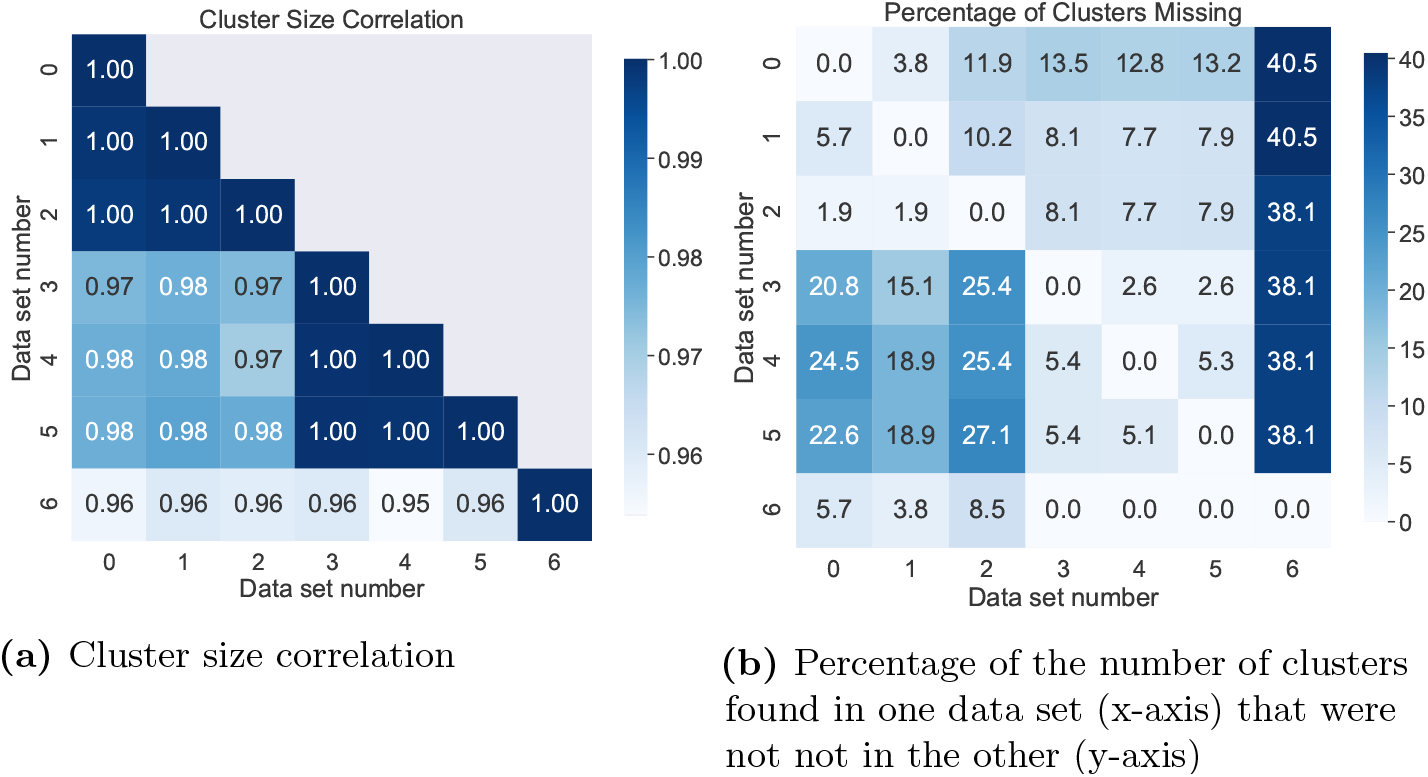
Comparing datasets. Panel (a) reports the correlation between repertoire samples, and panel (b) reports the percentage of missing clusters.

**Fig. 4.**
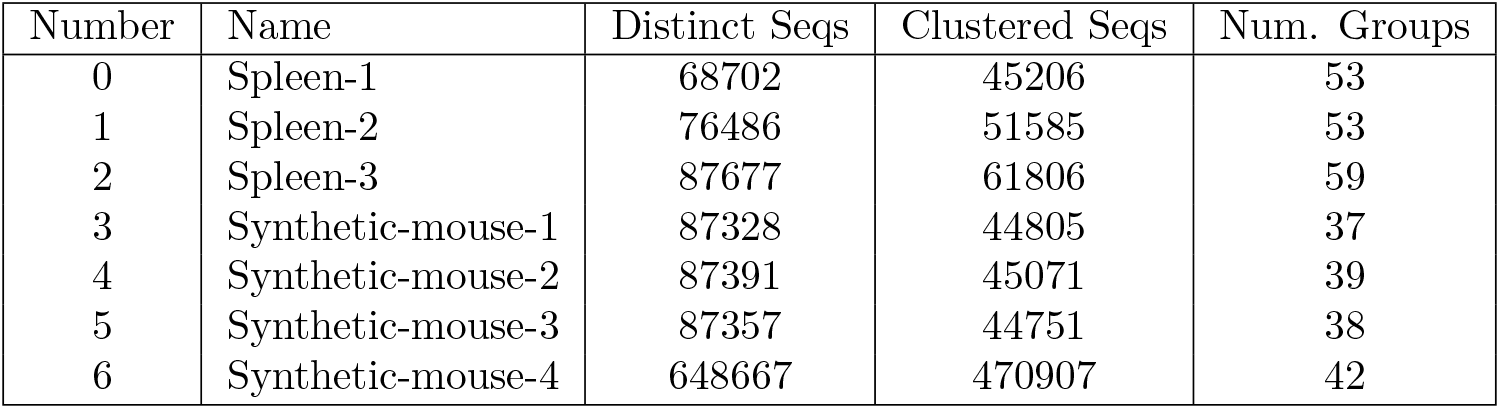
Repertoire datasets and clustering results. The minimum cluster size for all data sets is either 0.1% of the number of distinct sequences in a data set size or at least 30 sequences in a cluster, whichever is larger. The first two columns report an ID number and a descriptive name of the dataset. ‘Distinct Seqs’ indicates the total number of sequences in the dataset; ‘Clustered Seqs’ refers to the number of distinct amino-acid sequences that the algorithm assigned to a cluster; ‘Num. Groups’ reports the number of clusters above the threshold size that were found for each dataset. Datasets 0-2 were generated from the ImmunoSeq of TCR from mouse spleen, and datasets 3-6 were generated synthetically using OLGA.

Based on these results we adopted Hamming Distance as the similarity metric Because Hamming Distance generates more clusters for a given sample size, which allows more detailed comparisons between repertoires.

It is important to note that as long as the clustering algorithm and the distance metric capture fine grained similarities between sequences our procedure will be able to leverage the resulting groups to compare macro-level properties between repertoires.

### Finding corresponding clusters in two different repertoires

We used RCA’s matching algorithm (Sec. Cluster Matching) to identify corresponding clusters in different repertoires and found a large number of clusters that appear in each of three murine datasets. The distribution of relative sizes of these clusters is highly conserved across the datasets. We also found similarities between the natural and synthetic data but only when the size of the synthetic sample was significantly larger than the natural samples.

We applied our method to seven data sets (Sect. Data Description). The experimental data were obtained from naive (CD44lo) CD8+ T cells, magnetically sorted from 8 wk old C57Bl/6 splenocytes through ImmunoSeq TCR profiling, and the synthetic data were generated with OLGA tool. Table 4 summarizes the data sets and the clustering results. Clusters from one dataset that were not found in the other are considered to be *missing*. We used these data to produce Figs. 3a and 3b, which show the pairwise macro-level similarities and differences between datasets. Figure 3a shows that the three experimental data sets have highly correlated cluster sizes (0.99 Pearson correlation). This means that matching clusters have similar sizes relative to their dataset size. Further, the same correlation is found between synthetic datasets. There is also high correlation (> 0.95) between synthetic and natural datasets. These results suggest that RCA identifies macro-structure that is conserved among repertoires.

Figure 3b shows the percentage of clusters that are present in one data set but missing from another. Because we used a single clustering threshold across all datasets, it is unsurprising that the larger datasets generated more clusters than the smaller ones. With more sequences in a larger dataset, some small clusters are consolidated and some new clusters are found because they have the minimum number of sequences required (Algorithm 2). We found more missing clusters when comparing repertoires from mouse spleens than when we compared clusters between synthetic data sets. We interpret this result as evidence that there is a consistent structure captured by the algorithm in each dataset despite differences among individual sequences. The the murine datasets, we find missing clusters in part because of the difference in data set sizes. We account for varying dataset size in the following section. A larger number of clusters is missing from the synthetic repertoires when compared to real repertoires, but these differences disappear when the number of unique synthetic sequences is increased to around 10 times larger than the mice datasets. This reinforces our earlier point about the possible role of selection process in determining the probability of sequences ultimately belonging to a repertoire.

Furthermore, we studied an alternative cluster matching algorithm, based on the direct contact analysis (DCA) model [14], to match clusters from different samples. The DCA method has previously been used successfully on sequence ensembles to identify not not in the other (y-axis) structural contacts in proteins and RNA molecules for structure prediction [14, 15]. It was also shown that the trained model can generate new sequences that have the same function as those on which it was trained [16]. Although DCA is less accurate than our current cluster matching algorithm, once it is trained, it is computationally more efficient than ours.

Using the pseudo-likelihood method [17] (see Methods for details), we parametrized the DCA model on the aligned TCR sequences that belong to an identified cluster in one murine sample. We then assigned the score (Eq. (1)) to all the sequences in the respective clusters in the other sample. The DCA approach indeed matched 90% of the clusters identically to the matchings found by RCA.

However, when we omitted the term in the scoring function that depends on pairwise correlation of different aminoacids in the TCR, prediction accuracy did not change significantly, suggesting that identities of aminoacid residues for a given position in the TCR are sufficient for discriminating between the clusters (Supporting Information). This result suggests that the clusters are determined by conserved aminoacids at a set of positions.

### Comparing the macro-structure of different repertoires

Having established that our clustering and matching procedures capture macro properties of a repertoire that are shared between individuals, we now use these findings to determine how similar two repertoires are to each other. For example, if we have a dataset from one individual with a complete repertoire, we can use RCA to determine if another repertoire is complete or has significant differences. Such an analysis could be useful in assessing why some individuals are susceptible to a particular disease and others are not, either because of genetic predisposition or because of their life histories. It might also allow us to study how an individual’s repertoire changes as he or she ages, compare pre- and post-infection repertoires, and identify anomalous functional or phenotypic characteristics among individuals.

We compare two datasets: a *Reference* repertoire and a *Target* repertoire, and focus on the number of clusters in the Reference repertoire that are missing in the Target repertoire as the signal to analyze (see Repertoire Comparison for details).

We found that unaltered repertoire samples from different individuals are more similar to each other than they are when compared to their own original repertoire after sequences have been deleted.

We generated anomalous datasets by simulating the changes a superantigen would induce in mice TCR repertoires by deleting some of its sequences (Tables 9a, 9b) and ran RCA on these ablated datasets and the corresponding nonablated data. Our aim is to explore if our method finds more differences between altered and unaltered data sets than between unaltered datasets or datasets affected by the same superantigen.

Figure 5 shows the mean and standard deviation of the number of missing clusters using the original mice spleen data as Reference and the superantigen manipulated dataset as Target. Values in bold report the expected number of missing clusters when the Reference and Target are created from the same original data set; this is the expected number of missing clusters that can be attributed to differences in sample size, e.g., the entry for Spleen 1-Spleen 1 reports that on average there are 15 missing clusters between same size random samples of Spleen 1. In most cases the difference in means between unaltered data sets is smaller than between these and the superantigen manipulated samples, e.g., Spleen 1-Spleen 1 vs. Spleen 1-Spleen 2 (15.0 vs 12.6) and Spleen 1-Spleen 1 vs. Spleen 1-Spleen 2SEB (15.0 vs. 4.6). This difference shows that RCA is sensitive enough to distinguish the effects of a superantigen on a repertoire from the inherent differences of two normal repertoires.

**Fig. 5.**
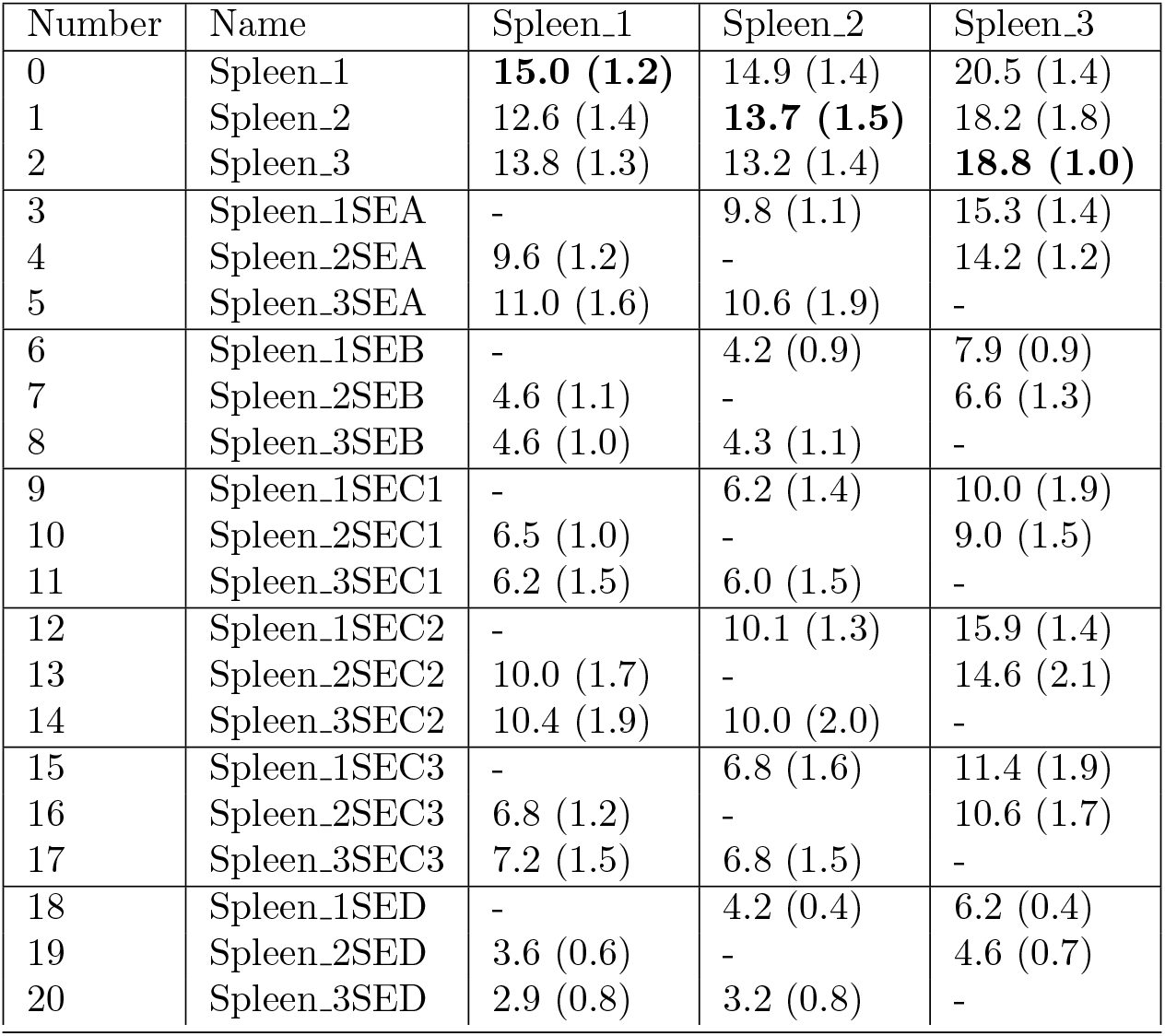
Identifying missing clusters. The original data sets are used as the Reference (indicated in column Name). The Target data sets (rows) include the unaltered murine samples and those samples manipulated to simulate the effects of the different superantigens. The mean and (standard deviation) are reported for the number of clusters that occur in the Reference data set but are missing from the Target repertoire. The expected number of missing sequences based only on sample size differences is shown in bold.

We also conducted the complementary experiment where the Reference data set is affected by the superantigen and the Targets are either the unaltered samples or the altered data sets from the other mice. Figure 6 shows the results for two super pathogens (SEA and SEC1) as illustration. Here we observe again that most datasets affected by the same superantigen have similar means e.g. Spleen 1SEA-Spleen 1SEA vs Spleen 1SEA-Spleen 2SEA while the means differ more markedly when compared to an unaltered data set e.g., Spleen 1SEA-Spleen 1SEA vs Spleen 1SEA-Spleen 2.

**Fig. 6.**
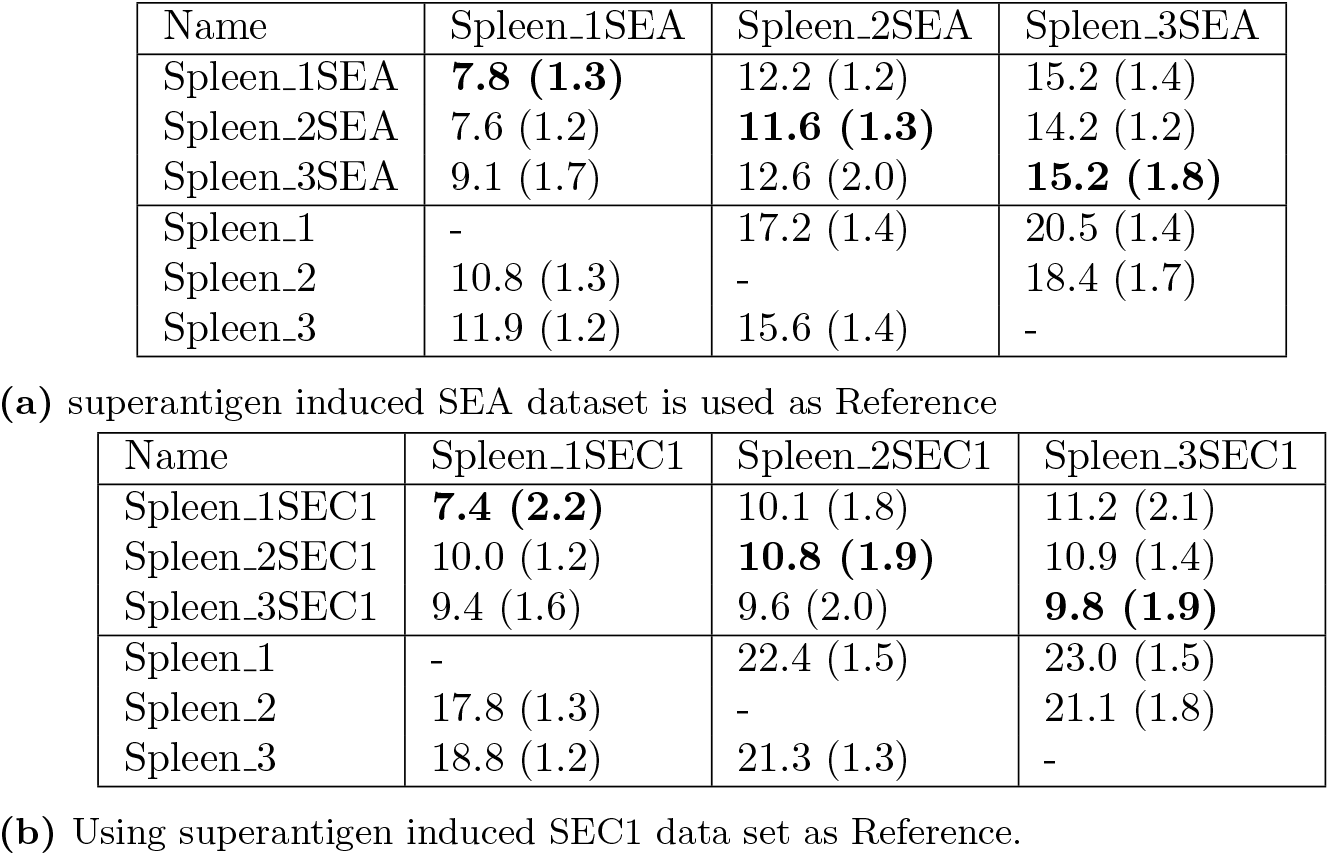
superantigens: Missing clusters for altered datasets (columns) as Reference and original data sets as Target (rows). The normalcy standard is shown in bold.

We next measured the distance between the missing cluster distributions of the Reference-Reference samples and the Reference-Target samples using the Earth Mover’s Distance (EMD) [18]. Table 7 and Fig. 7 show the distance between all pairs of distributions of altered and unaltered data sets. We expect low numbers between unmanipulated datasets and between datasets with the same manipulations, i.e., when two repertoires were affected by the same superantigen. We expect high numbers when comparing unmanipulated and manipulated datasets and between datasets affected by a different superantigen. The procedure detects similarities between all of the original data sets, finding low distances between them, while reporting higher distance values between Reference and Target when one of them is affected by a superantigen.

**Fig. 7.**
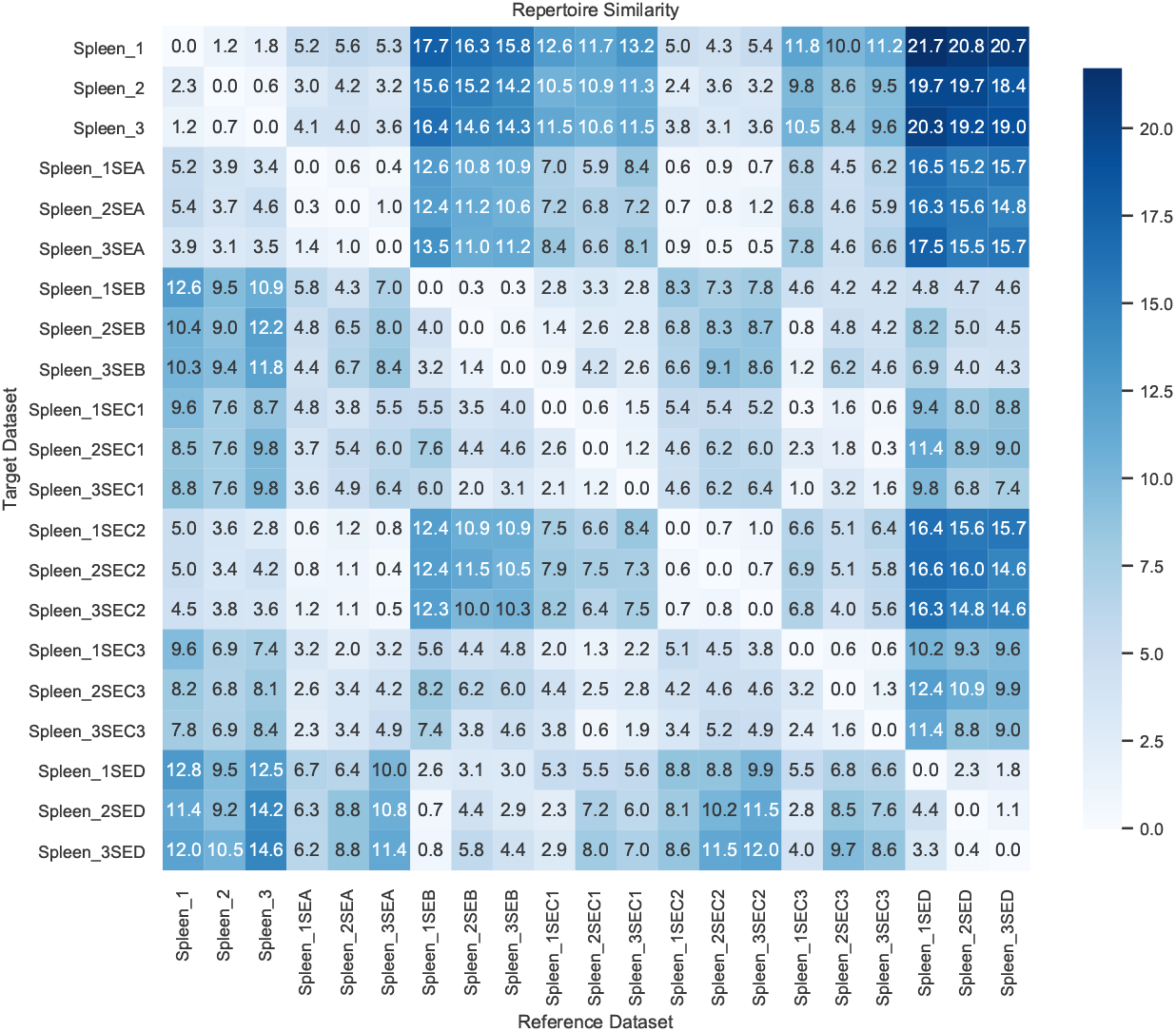
Earth Mover’s Distance (EMD) between repertoire samples. Reference datasets are set as columns and Target datasets as rows. Datasets number to name correspondence can be seen in Table 5.

Finally, we illustrate in Fig. 8 the effectiveness of our procedure for detecting systematic differences between repertoires. Here we selected the detection threshold to be the maximum Earth Mover’s Distance (EMD) between the unaltered datasets Fig. 7 and flag two datasets as different if the maximum between the Reference-Target and the Targer-Reference EMD’s is above it. This procedure could be applied in an experimental setting where the Reference data set, say from a healthy mouse, is used to assess which of a series of repertoire samples from other mice are the most different from it and warrant further investigation.

**Fig. 8.**
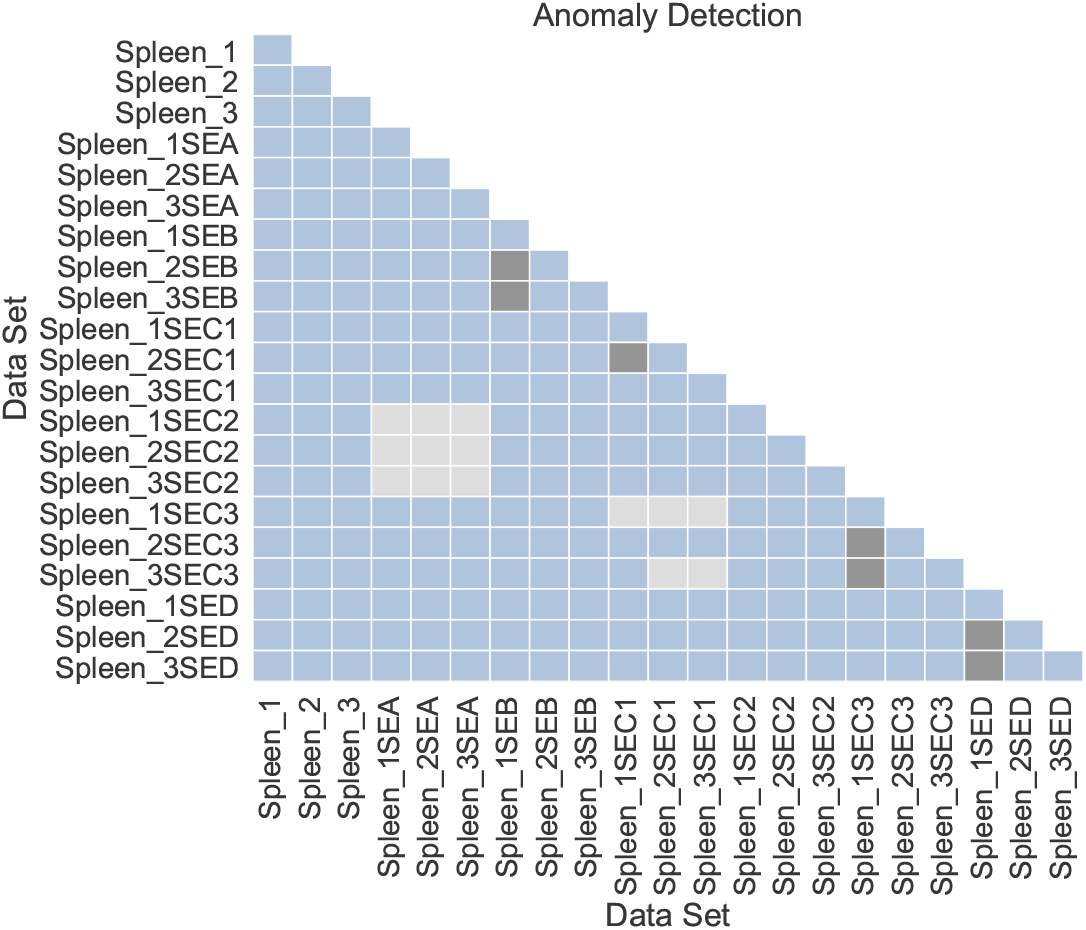
Anomaly detection using Earth Mover’s Distance (EMD). The similarity between two data sets is the maximum EMD of the Reference-Target and Target-Reference EMDs. The similarity threshold was set as the maximum EMD between unaltered repertoires (2.3) Fig.7. Dark squares are false negatives, grey squares false positives and the blue area represents correct classifications as either similar or dissimilar. For this example we have 175 true positives, 14 true negatives, 7 false positives and 14 false negatives with a precision, recall and accuracy of 0.96, 0.93, 0.90 respectively.

**Fig. 9.**
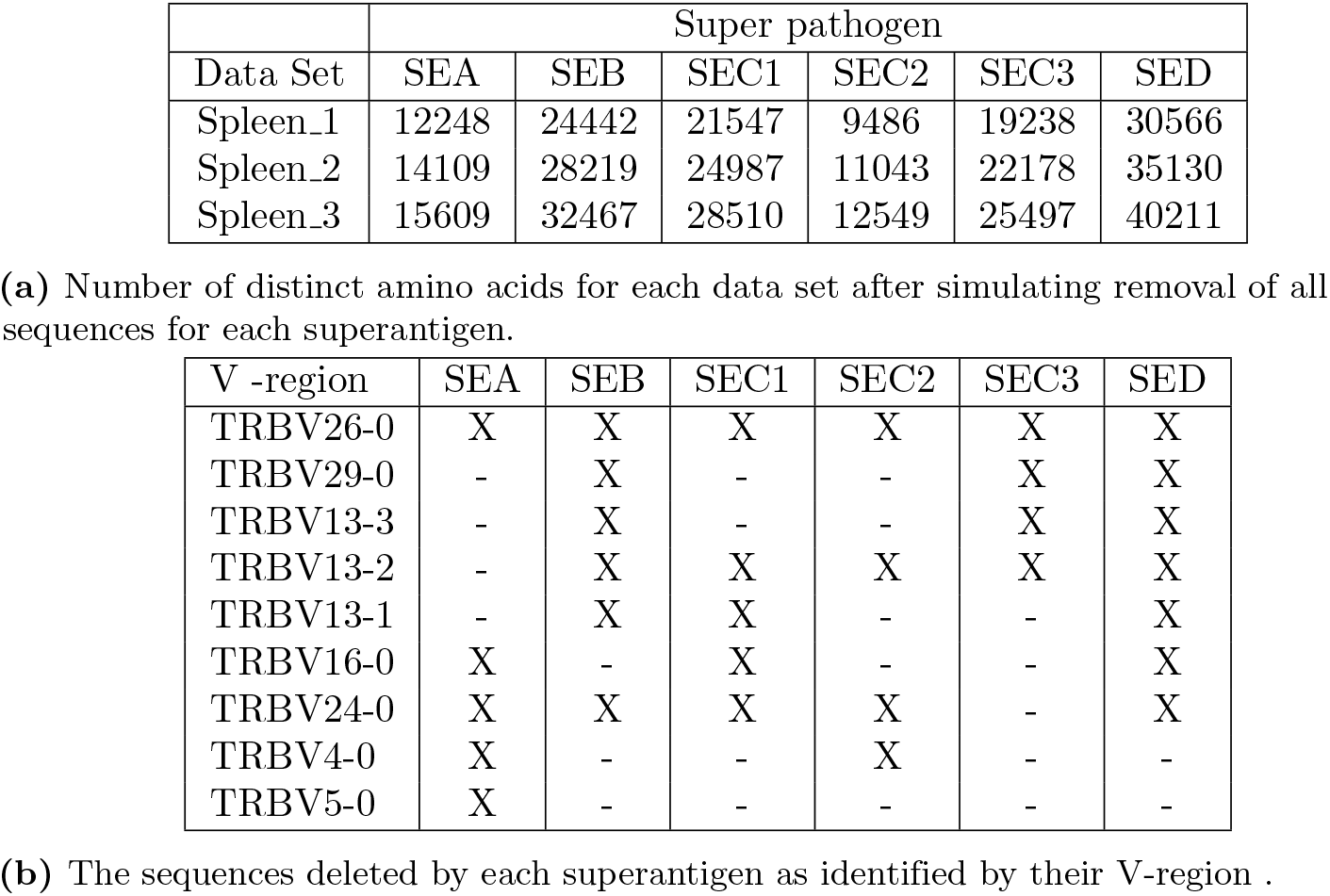
Data from mouse spleen reperotoire’s TCR after computational manipulation of the data to simulate the effect of a superantigen.

## Discussion

With whole immune repertoires becoming available, there is a need for methods to assess repertoire completeness, compare the repertoires of different individuals, and detect the presence of anomalies. Because most sequences are rare, comparing sequences directly is unlikely to be informative, and we instead opted to consider the macro-level properties of each repertoire using cluster analysis.

Extending the idea of a public sequence, we propose the concept of a public cluster: a set containing all sequences that are similar under some criterion and from which each individual repertoire draws its specific instances.

RCA identifies sequences across repertoire samples that belong to the same public cluster and records the sequences that belong to each public cluster in each individual. Based on experimental data from three naive mice from the same laboratory, which were raised under similar conditions, our results show that the number and size of individual clusters are conserved across these individuals, even when their individual sequences vary greatly. These results suggest that RCA captures intrinsic macro-level properties of their repertoires. To a lesser extent, the experiments show that this structure occurs in synthetically-generated (using OLGA tool [9]) repertoires that simulate V(D)J recombination to generate TCR sequences before they undergo positive and negative selection in thymus. We also showed (Supporting Information) that RCA matches clusters correctly between different repertoires even when using a different scoring function. The alternative scoring function accounts for homologous positions and pairwise-correlations between amino acids in the TCR receptor.

When we simulated the effect of superantigens on our datasets (by deleting sequences from the samples), we found that unaltered repertoire samples from different individuals are more similar to each other than they are when compared to their own original repertoire after sequences have been deleted. These results suggest that RCA detects when two repertoire samples have similar macro-structure and when they have significant discrepancies.

Because our primary objective was to investigate macro structures in repertoires, our analysis focused on how many clusters match and don’t match between individuals and on their relative sizes. We leave for future work many other avenues that can potentially reveal significant similarities and differences between individuals, for example, cluster properties such as size, compactness and sequence probability distributions. We could also study the role of certain sequences within a cluster such as their centrality and their role as public sequences. In the future we plan to test RCA on murine samples from different labs and assess RCA’s ability to detect discrepancies arising from different laboratory procedures to extract genetic samples.

## Methods

### Data Description

#### Mice and Cell Sorting

All mice were maintained under specific-pathogen free conditions at The Biodesign Institute, and experiments were performed in compliance with institutional guidelines as approved by Institutional Animal Care and Use Committee of Arizona State University. *CD*8^+^ T cells were purified by positive immunomagnetic cell sorting (>95% *CD*8^+^, >95% *CD*44^*lo*^; Miltenyi Biotec) as previously described [19] from spleens of 6-8 week-old donor C57BL/6 mice. Single cell suspensions were prepared from splenocytes as previously described [20]. Briefly, erythrocytes were lysed with ammonium chloride lysis (ACK) buffer purchased from Lonza (Allendale, NJ). Aliquots of all samples were analyzed by FACS staining as previously described in 96 well plates with fluorochrome-labeled monoclonal antibodies: anti-CD8 (clone 53-6.7), anti-CD44 (clone IM7), anti-CD4 (clone GK1.5) [21]. Samples were then fixed in 1% paraformaldehyde solution and immediately acquired on a BD LSR II Fortessa flow cytometer (San Jose, CA) and analyzed using FlowJo Software (Tree-Star, Ashland, OR). All monoclonal antibodies were purchased from BD Pharmigen (San Diego, CA) or eBiosciences (San Diego, CA).

#### TCR CDR3 Sequencing

Three samples consisting of between 1.16×10^6^ – 1.48×10^6^ CD8 sorted cells from C57BL/6 mouse spleens were transferred to Adaptive Biotechnologies (Seattle, WA) for standard ImmunoSEQ TCR*β* profiling [22]. Generation of the library of TCR CDR3 encoding sequencing amplicons using the ImmunoSEQ technology requires the use of multiple comprehensive multiplex primer sets. Briefly, equimolar pools of 45 Vb forward primers and 13 Jb reverse primers, each specific to the known functional murine TCR*β* V and J gene segments are used to generate 60nt reads of TCR*β* CDR3 amplicons. Additionally, each primer contains at the 5’ end the universal forward and reverse primer sequences compatible with the base Illumina sequencing technology. Sequencing was performed on the MiSeq analyzer using 2×100 paired end reads.

Read data were processed by Adaptive Biotechnologies for correction of PCR biases informed by synthetic repertoires and identification of germline gene segments, the number of insertions and deletions, and functional status using standardized definitions from the international ImMunoGeneTics information system [23].

By using the ImmunoSEQ high-throughput sequencing approach on genomic DNA extracted from purified CD8+ T cells, we were able to analyze the CDR3 sequence repertoire from over 1 10^6^ T cells from each of the spleens of three C57BL/6 mice. Samples consisted of 1.16×10^6^, 1.34×10^6^, and 1.48×10^6^ CD8 sorted T cells from each of the respective mice. Each of the three samples yielded roughly 10^5^ unique CDR3 sequence reads with an average CDR3 length of around 42 nucleotides for both total reads and productive reads.

#### Synthetic Data

We used the OLGA software tool [9] to generate synthetic mouse TCR beta sequence data sets. This software produces n-frame nucleotide CDR3 sequences from a generative model of V(D)J recombination. Sequences are generated independently of one another with no errors.

#### Simulated Data

We simulated the superantigens in the following way, as shown in Table 9:

### Algorithms

RCA assumes that the sequences found in a repertoire sample are a function of the sampling technique and the number of such sequences present in the full repertoire (reflecting their probability of generation, selection, etc). The larger the sample, the more likely it is to include sequences that are less frequent in the full repertoire. Further, we assume that similar sequences have similar probabilities of being generated and that individuals with similar life histories FIXME: is this the right term? are likely to have many similar sequences in their repertoire.

#### Clustering

We started with the DBSCAN algorithm [24] (Algorithm 2) and made modifications to improve efficiency (Algorithm 3). We combined DBSCAN with the distance metric to find a repertoire’s cluster set.

A cluster under DBSCAN is a set of sequences for which there exists a mutation path between any two of its elements; two sequences *S*_1_ and *S_n_* belong to the same cluster if there is a path < *S*_1_*, S*_2_*, S*_3_*,…, S_n_* > between them for which the distance between *S_i_* and *S_i_*_+1_ is at most one mutation for all *i* and all *S_i_* are in the set.

For each sequence in a dataset, the algorithm identifies all similar sequences in the dataset—sequences of distance one or less. To improve efficiency, we created a new procedure (Fig. 3) which generates, for each sequence, all sequences at distance one from it and then checks for their presence in the repertoire. The distance once sequences are stored as a Hash table, allowing constant time for each comparison. This produces a significant time efficiency, reducing the operation from *O*(*l·n*^2^) to *O*(*l·n*), where *l* is the sequence length and *n* the dataset size, provided that the number of possible sequences at distance one from a given sequence is smaller than the repertoire sizem, which is almost always the case.

**Algorithm 2:**
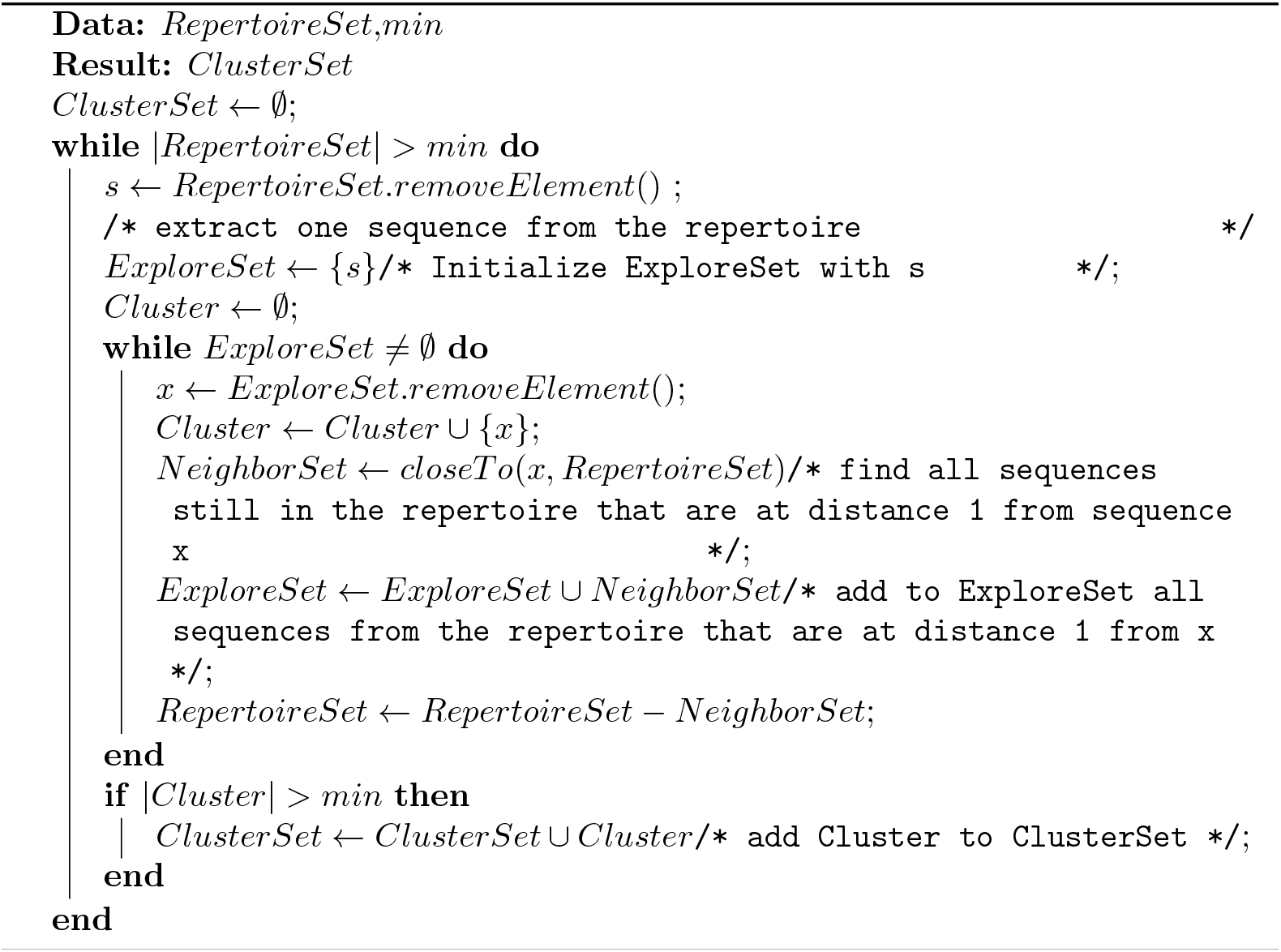
DBscan

**Algorithm 3:**
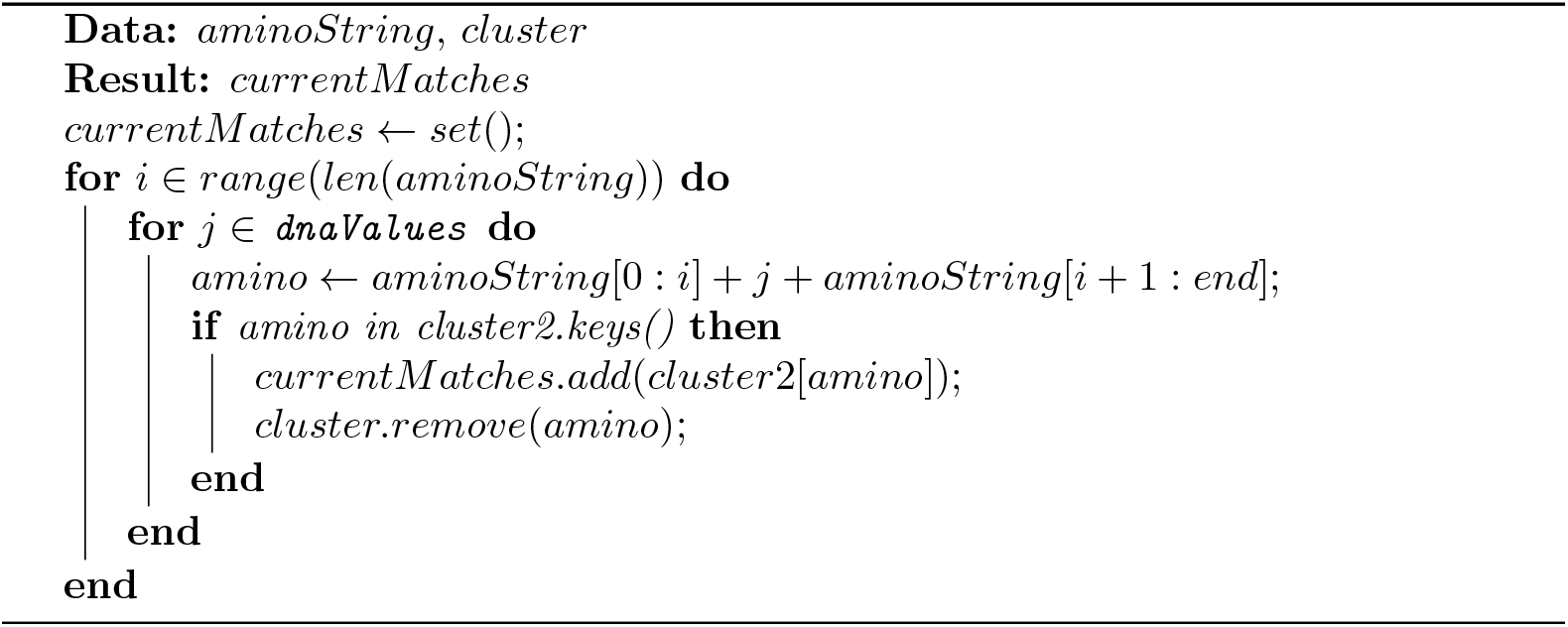
closeTo (with Hamming Distance)

#### Cluster Matching

Repertoire samples are compared by matching their respective clusters. Two clusters match if the distance between any two of their sequences is at most one. that is, two clusters match if their respective sequences would belong to the same cluster if those sequences were in the same sample. This is not a one-to-one relationship. A cluster in one data set, say cluster 1, might match several clusters in another data set, clusters 4, 7 and 8 for example. In such a case, clusters 4, 7 and 8, are counted as a single cluster for the purpose of comparing those repertoires (see Algorithm 4).

**Algorithm 4:**
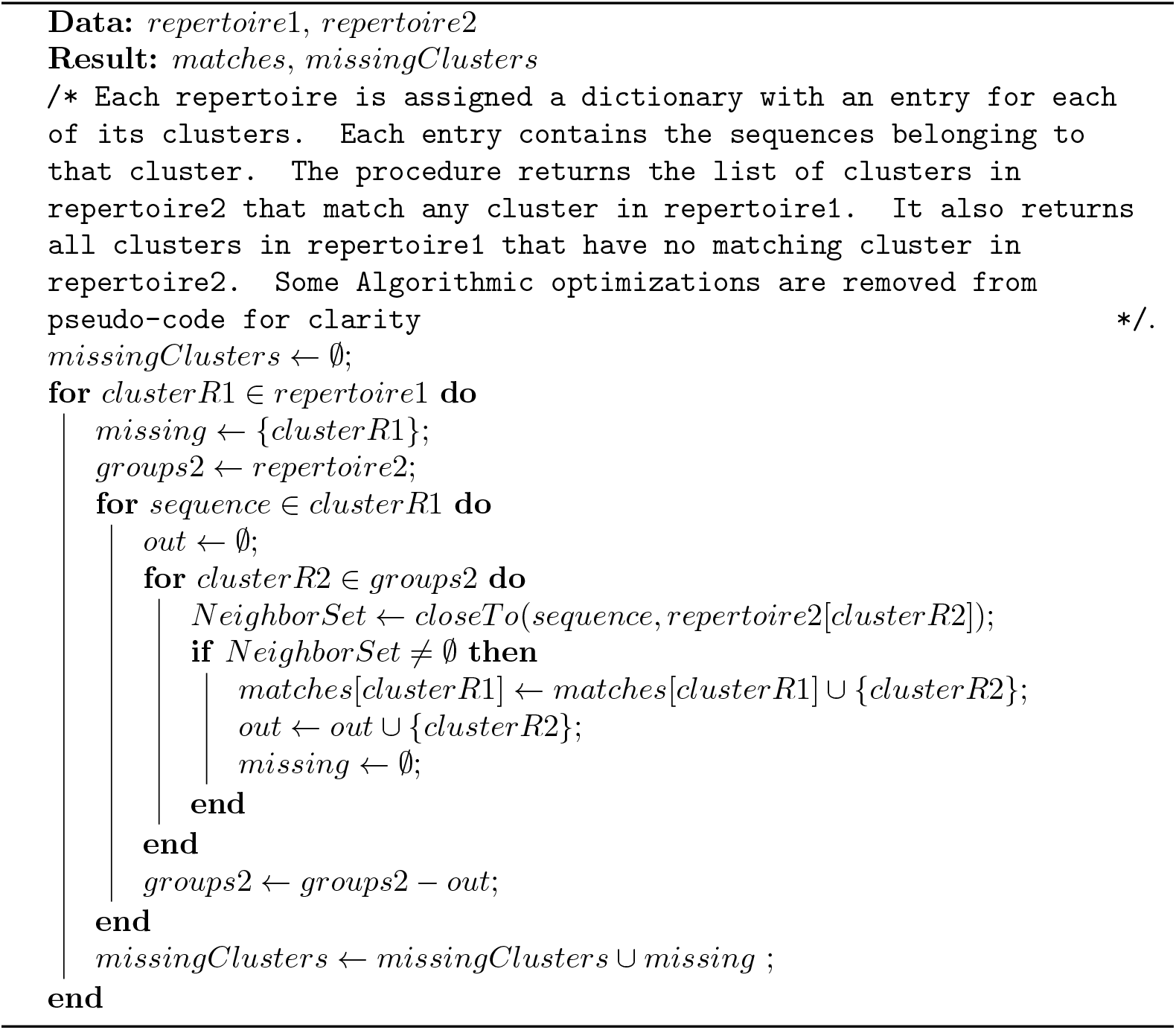
Repertoire Matching

#### Matching clusters with the DCA Model

The DCA method parametrizes the two-point tensor *J_ij_*(*A, B*) and an array *h_i_*(*A*) so that the probability of aminoacid sequence consisting of nucleotides *A*_1_*, A*_2_*,…, A_n_* is given by

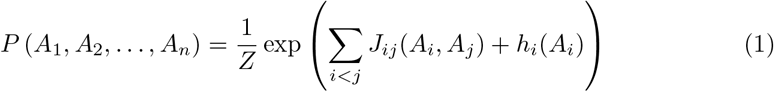

The tensors *J_ij_* and *h_i_* are parametrized on a set of aligned aminoacid sequences which correspond to all TCR sequences assigned to a single cluster, so that they reproduce frequencies *f_i_* of occurrence of aminoacids on position *i* and pairwise frequency *f_ij_* on positions *i* and *j*:

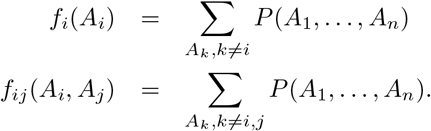

The score that assigns if TCR sequence *s* belongs to cluster *c* is given by Eq. (1). We trained the model on clusters from one chosen dataset and used it to assign TCR sequences to clusters from the other sample. We then compared the fraction of TCR sequences that were assigned by the DCA score to the same cluster as with the clustering algorithm, and it was between 90 to 94% for all sequences considered. The correct assignment rate was only weakly dependent on the two-site correlation (tensor *J_i_j*), and using the Eq. (1) alone to assign the sequences to clusters with only *h_i_* parameter produces nearly the same assignment. The sequence logos for TCR seqeunces belonging to the same cluster as well as the scores assigned to sequences belonging to respective clusters with the DCA method are shown in detail in the Supp. Information.

#### Repertoire Comparison

Comparing repertoires requires more than simply matching clusters between two datasets: On the one hand, the number of clusters found is a function of the size of the repertoire sample (Sect. *Clustering the repertoire*) and on the other, different samples of the same size are likely to contain some differences from random sampling, i.e., by chance, different samples of the same size are likely to contain some unmatched clusters even in the absence of anomalies, because our datasets are a sample of all the sequences in the full repertoire.

To identify missing clusters, we account for the effect of different size datasets and for the fact that datasets of the same size from the same exact individual are expected to have some mismatched clusters. We account for this by first generating uniform random samples that are the same size as the dataset without replacement from both the Reference and Target (the results presented here use a sample size of 41k which is the size of our smallest super-antigen dataset). Next, we compute the number of clusters that are missing from the Reference in each of the Reference samples and compute the number of missing clusters from the Reference set in each of the Target samples. Note that swapping the Reference and Target datasets is likely to produce different distributions. Finally we compare the missing cluster distributions from n repetitions of the Reference-Reference and the Reference-Target trials using the Earth Mover’s Distance (EMD) or Wasserstein metric (Fig. 7 shows the results of 20 repetitions). This metric quantifies the minimum amount of values that need to be “moved” between categories to transform one discrete distribution into another. The algorithm is described in Algorithm 5.

**Algorithm 5:**
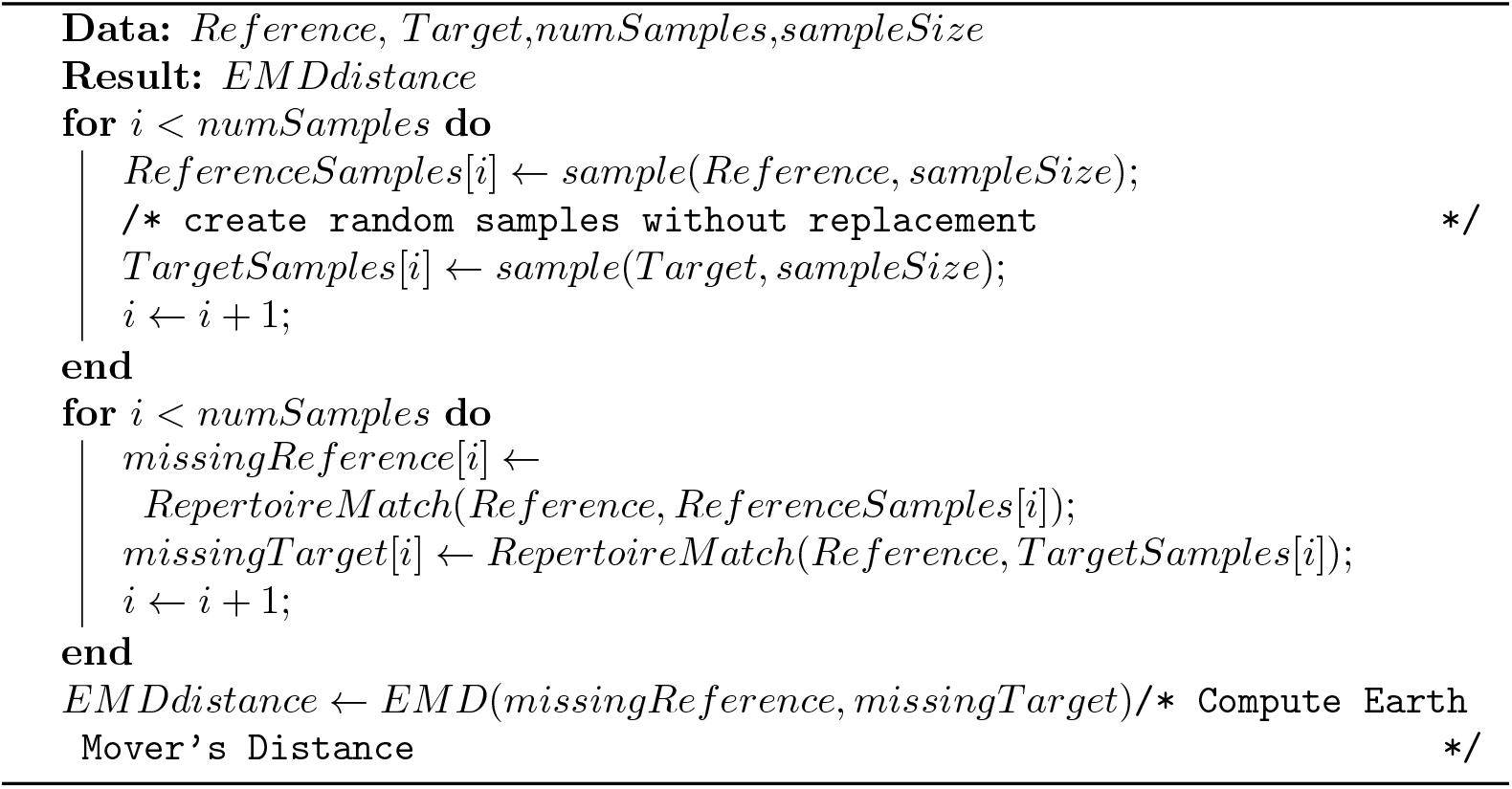
Repertoire Comparison

## Acknowledgements

We gratefully acknowledge the adaptive lab for their comments on the manuscript.

## References

1. Balachandran VP, Łuksza M, Zhao JN, Makarov V, Moral JA, Remark R, et al. Identification of unique neoantigen qualities in long-term survivors of pancreatic cancer. Nature. 2017;551(7681):512–516.

2. Casrouge A, Beaudoing E, Dalle S, Pannetier C, Kanellopoulos J, Kourilsky P. Size estimate of the *αβ* TCR repertoire of naive mouse splenocytes. The Journal of Immunology. 2000;164(11):5782–5787.

3. Arstila TP, Casrouge A, Baron V, Even J, Kanellopoulos J, Kourilsky P. A direct estimate of the human *αβ* T cell receptor diversity. Science. 1999;286(5441):958–961.

4. Schwab R, Szabo P, Manavalan JS, Weksler ME, Posnett DN, Pannetier C, et al. Expanded CD4+ and CD8+ T cell clones in elderly humans. The Journal of Immunology. 1997;158(9):4493–4499.

5. Cohen J. How long do vaccines last? The surprising answer may help protect people longer. Science. 2019;10.

6. Zajac AJ, Blattman JN, Murali-Krishna K, Sourdive DJ, Suresh M, Altman JD, et al. Viral immune evasion due to persistence of activated T cells without effector function. The Journal of experimental medicine. 1998;188(12):2205–2213.

7. Blattman JN, Sourdive DJ, Murali-Krishna K, Ahmed R, Altman JD. Evolution of the T cell repertoire during primary, memory, and recall responses to viral infection. The Journal of Immunology. 2000;165(11):6081–6090.

8. Madi A, Poran A, Shifrut E, Reich-Zeliger S, Greenstein E, Zaretsky I, et al. T cell receptor repertoires of mice and humans are clustered in similarity networks around conserved public CDR3 sequences. Elife. 2017;6:e22057.

9. Sethna Z, Elhanati Y, Callan Jr CG, Walczak AM, Mora T. OLGA: fast computation of generation probabilities of B-and T-cell receptor amino acid sequences and motifs. Bioinformatics. 2019;35(17):2974–2981.

10. Murugan A, Mora T, Walczak AM, Callan CG. Statistical inference of the generation probability of T-cell receptors from sequence repertoires. Proceedings of the National Academy of Sciences. 2012;109(40):16161–16166.

11. Tanyi JL, Bobisse S, Ophir E, Tuyaerts S, Roberti A, Genolet R, et al. Personalized cancer vaccine effectively mobilizes antitumor T cell immunity in ovarian cancer. Science Translational Medicine. 2018;10(436).

12. Dash P, Fiore-Gartland AJ, Hertz T, Wang GC, Sharma S, Souquette A, et al. Quantifiable predictive features define epitope-specific T cell receptor repertoires. Nature. 2017;547(7661):89–93.

13. Pogorelyy MV, Minervina AA, Shugay M, Chudakov DM, Lebedev YB, Mora T, et al. Detecting T cell receptors involved in immune responses from single repertoire snapshots. PLoS Biology. 2019;17(6):e3000314.

14. Morcos F, Pagnani A, Lunt B, Bertolino A, Marks DS, Sander C, et al. Direct-coupling analysis of residue coevolution captures native contacts across many protein families. Proceedings of the National Academy of Sciences. 2011;108(49):E1293–E1301.

15. De Leonardis E, Lutz B, Ratz S, Cocco S, Monasson R, Schug A, et al. Direct-Coupling Analysis of nucleotide coevolution facilitates RNA secondary and tertiary structure prediction. Nucleic acids research. 2015;43(21):10444–10455.

16. Russ WP, Figliuzzi M, Stocker C, Barrat-Charlaix P, Socolich M, Kast P, et al. An evolution-based model for designing chorismate mutase enzymes. Science. 2020;369(6502):440–445.

17. Ekeberg M, Hartonen T, Aurell E. Fast pseudolikelihood maximization for direct-coupling analysis of protein structure from many homologous amino-acid sequences. Journal of Computational Physics. 2014;276:341–356.

18. Pele O, Werman M. Fast and robust earth mover’s distances. In: 2009 IEEE 12th International Conference on Computer Vision. IEEE; 2009. p. 460–467.

19. Mora JR, Bono MR, Manjunath N, Weninger W, Cavanagh LL, Rosemblatt M, et al. Selective imprinting of gut-homing T cells by Peyer’s patch dendritic cells. Nature. 2003;424(6944):88–93.

20. Murali-Krishna K, Altman JD, Suresh M, Sourdive DJ, Zajac AJ, Miller JD, et al. Counting antigen-specific CD8 T cells: a reevaluation of bystander activation during viral infection. Immunity. 1998;8(2):177–187.

21. Altman JD, Moss PA, Goulder PJ, Barouch DH, McHeyzer-Williams MG, Bell JI, et al. Phenotypic analysis of antigen-specific T lymphocytes. Science. 1996;274(5284):94–96.

22. Robins HS, Campregher PV, Srivastava SK, Wacher A, Turtle CJ, Kahsai O, et al. Comprehensive assessment of T-cell receptor *β*-chain diversity in *αβ* T cells. Blood, The Journal of the American Society of Hematology. 2009;114(19):4099–4107.

23. Monod MY, Giudicelli V, Chaume D, Lefranc MP. IMGT/JunctionAnalysis: the first tool for the analysis of the immunoglobulin and T cell receptor complex V–J and V–D–J JUNCTIONs. Bioinformatics. 2004;20(suppl 1):i379–i385.

24. Ester M, Kriegel HP, Sander J, Xu X, et al. A density-based algorithm for discovering clusters in large spatial databases with noise. In: Kdd. vol. 96-34; 1996. p. 226–231.

