## Supplementary Information for "A Macro-scale Comparison Algorithm for Analysis of TCR Repertoire Completeness"

We provide here further details about training the DCA model for assigning sequences to clusters as well as provide sequence logos for individual TCR sequence clusters identified by our clustering algorithm.

#### Methods

Each cluster of the the three datasets (Spleen-1, Spleen-2, Spleen-3) underwent plmDCA analysis using no reweighting coefficient on similar sequences. From this analysis, single position and pairwise amino acid identity probabilities were determined and stored in the H (single position) and J (pairwise) matrix generated in each run.

Subsequently, each individual cluster of each dataset was compared to all clusters in the three datasets. To accomplish the comparison, the H and J matrix of the individual cluster of a single dataset was energetically scored against all other clusters according to the Hamiltonian of the plmDCA method. An additional comparison using just the single position parameters of a cluster was also performed.

#### Results

The derived parameters can be viewed as both residue and position specific probabilities of sequence identity unique to each individual cluster. Another cluster scoring closely to the target cluster would indicate a high probability of the clusters being related in sequence space. Unlike other methods, a higher energy indicates a higher probability of the scored sequence belonging to the scoring cluster. Using this method we verified the results provided in the given file (matches.pickle) albeit with a very small number of differences in each cluster comparison. The differences are available in txt file ('spleen_results.txt') I have put here on Overleaf.

Notably, scoring of sequences using just single position parameters (no pairwise) provided nearly identical results and matched the overall relative separation between clusters compared to the inclusion of pairwise parameters. No notable increase or decrease in performance was observed possibly indicating the absence of evolutionary pairwise correlations in the clustering algorithm used to generate Spleen-1, Spleen-2, Spleen-3.

#### Drawbacks/Deficiencies

Unexplored was the relationship between cluster sequence number and prediction results. Many clusters had low sequence number (<5000) but the regularization terms were not tuned to account for this deficiency. Thus training parameters of these clusters may be less precise than clusters with sufficient numbers (>5000). Despite this, the agreement of the plmDCA and the provided results was quite good.

### Data Table of Contents

**Page No.**

**1 Dataset 1 (Spleen-1) 1**

1.1 Dataset 1's Sequence Logos and Pairwise Parameter (J) Matrices . . . . . . . . . . . . . . . . . . . . . . . . 1-18
1.2 Energy Comparison using Dataset 1's Single Position (H) and Pairwise (J) Parameters . . . . . . . . . . . . . 19-36
1.3 Energy Comparison using Dataset 1's Single Position Parameters (H) exclusively . . . . . . . . . . . . . . . 37-54

**2 Dataset 2 (Spleen-2) 55**

2.1 Dataset 2's Sequence Logos and Pairwise Parameter (J) Matrices . . . . . . . . . . . . . . . . . . . . . . . . 55-72
2.2 Energy Comparison using Dataset 2's Single Position (H) and Pairwise (J) Parameters . . . . . . . . . . . . . 73-90
2.3 Energy Comparison using Dataset 2's Single Position Parameters (H) exclusively . . . . . . . . . . . . . . . 91-108

**3 Dataset 3 (Spleen-3) 109**

3.1 Dataset 3's Sequence Logos and Pairwise Parameter (J) Matrices . . . . . . . . . . . . . . . . . . . . . . . . 55-72
3.2 Energy Comparison using Dataset 3's Single Position (H) and Pairwise (J) Parameters . . . . . . . . . . . . . 73-90
3.3 Energy Comparison using Dataset 3's Single Position Parameters (H) exclusively . . . . . . . . . . . . . . . 91-108

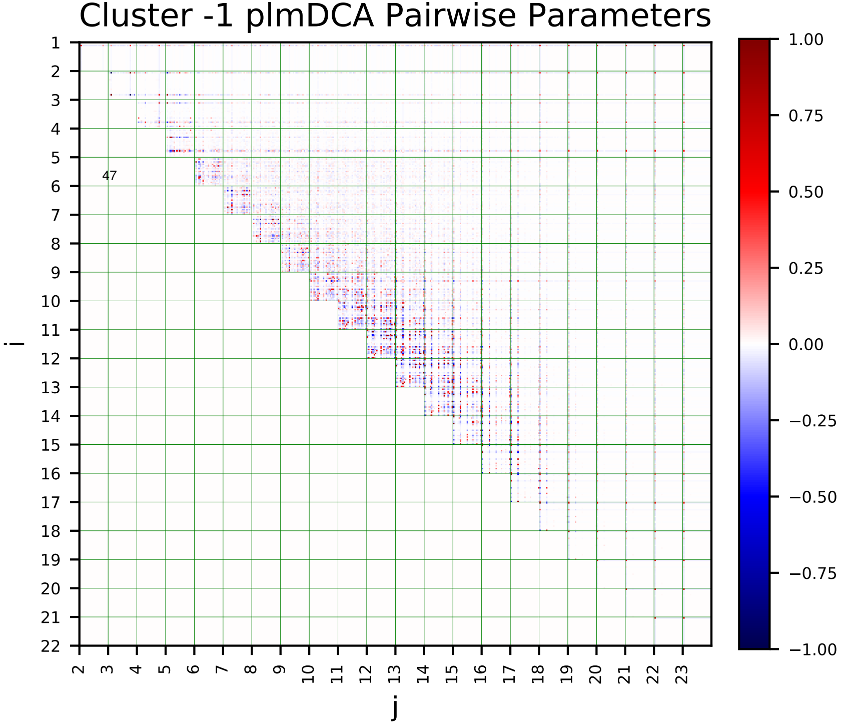
Cluster -1 plmDCA Pairwise Parameters
47
i
j

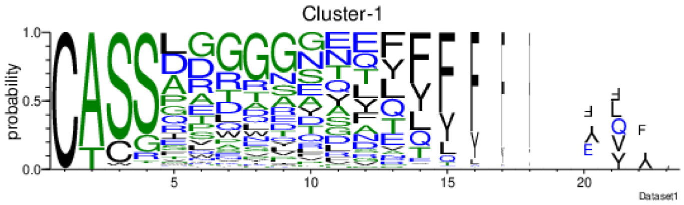
Cluster-1
probability
Dataset1

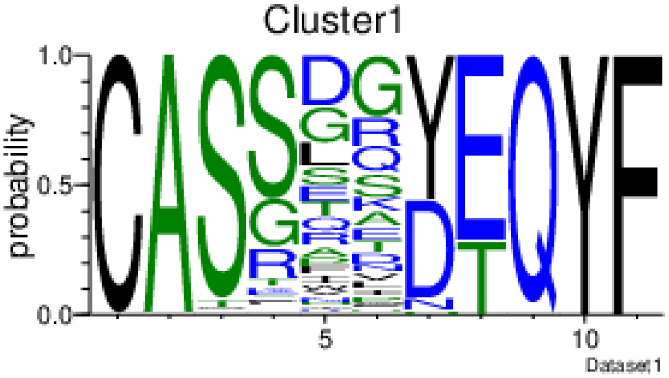
Cluster1
probability
Dataset1

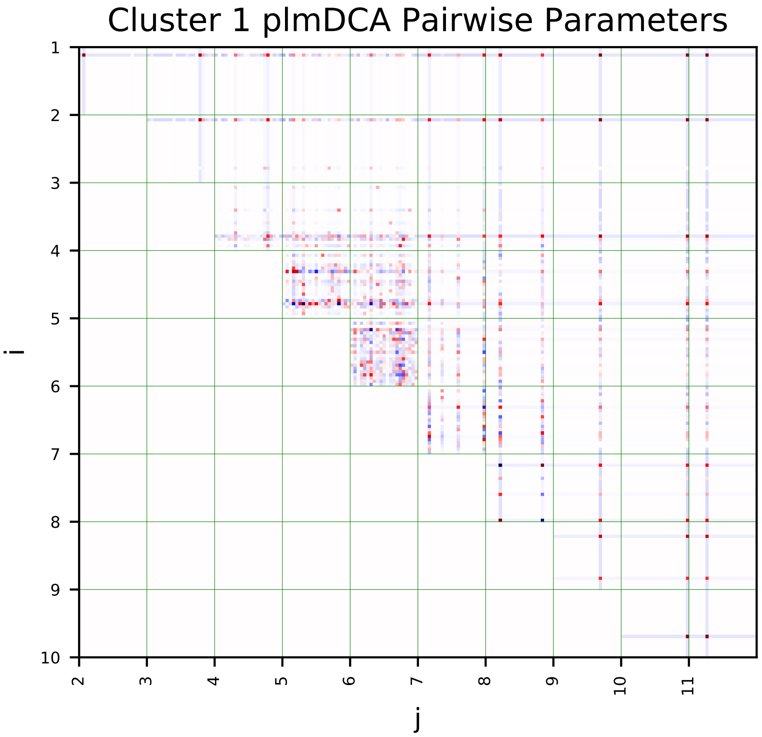
Cluster 1 plmDCA Pairwise Parameters
i
j

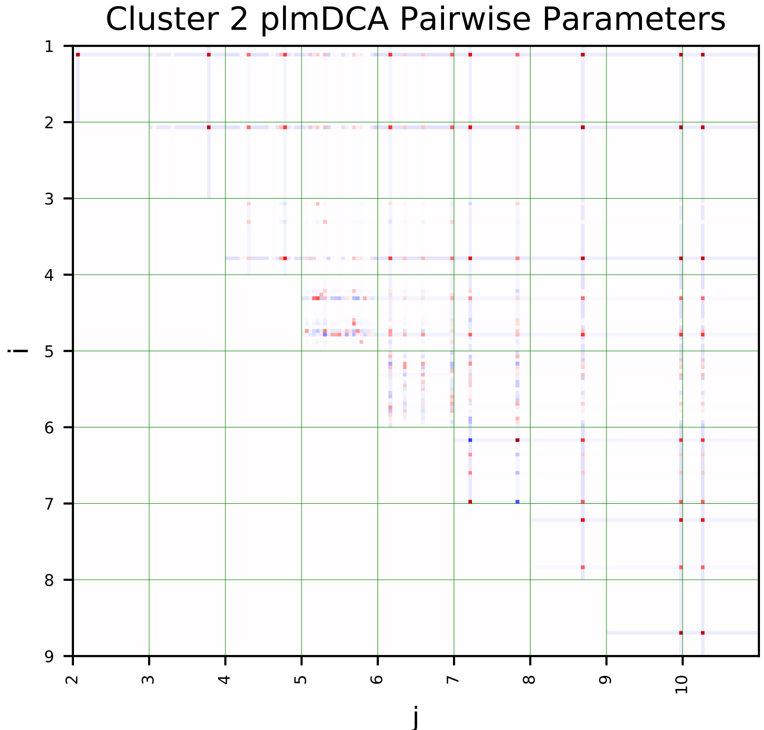
Cluster 2 plmDCA Pairwise Parameters
i
j

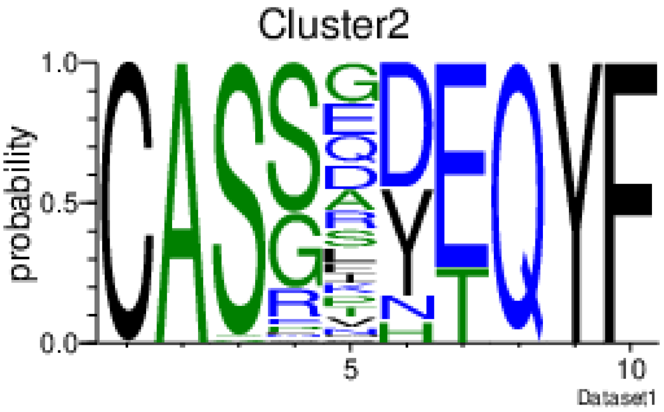
Cluster2
probability
Dataset1

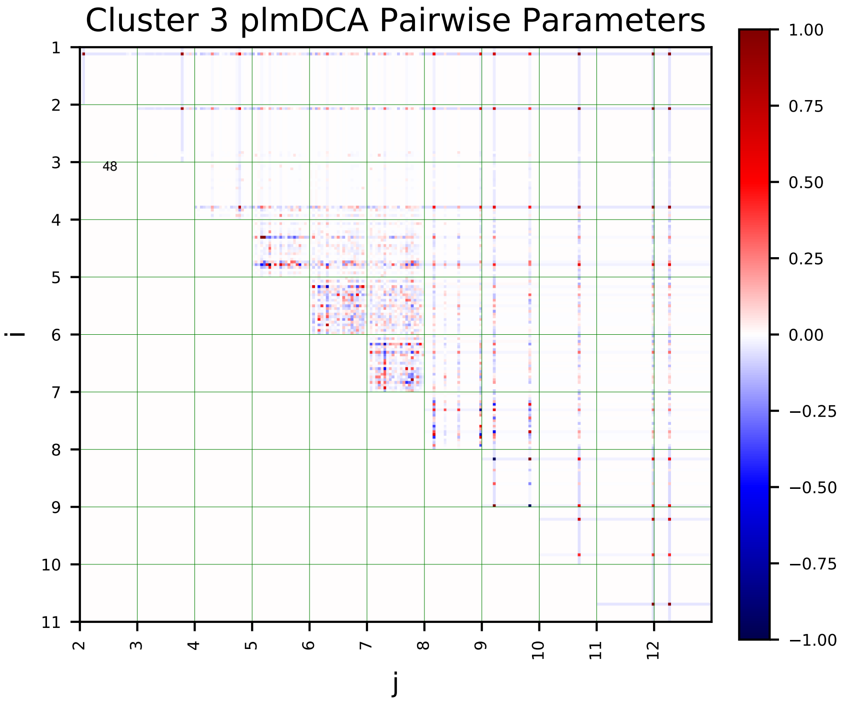
Cluster 3 plmDCA Pairwise Parameters
i
j
48
1.00
0.75
0.50
0.25
0.00
−0.25
−0.50
−0.75
−1.00

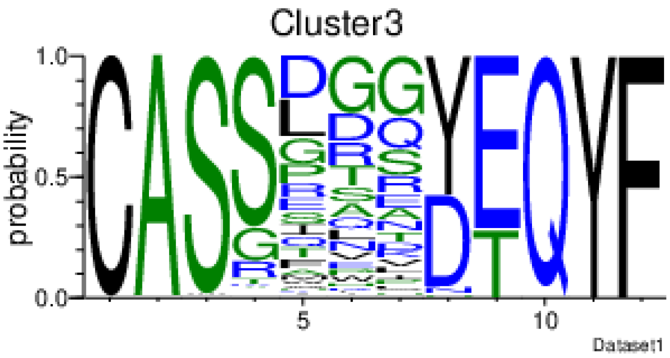
Cluster3
probability
Dataset1

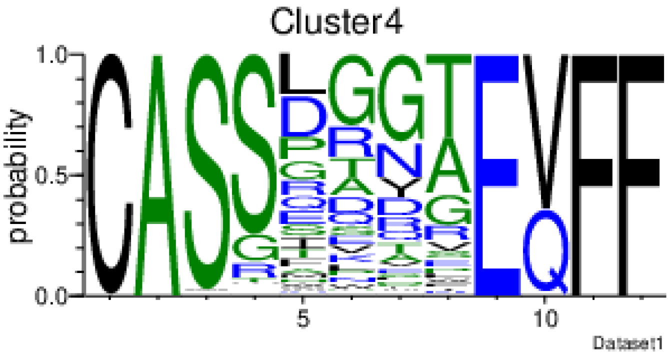
Cluster4
probability
Dataset1

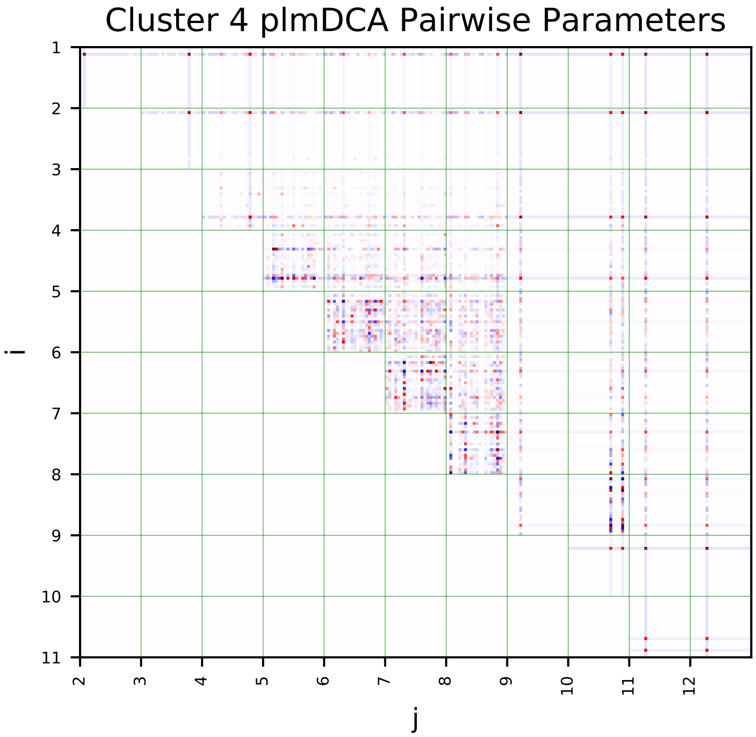
Cluster 4 plmDCA Pairwise Parameters
i
j

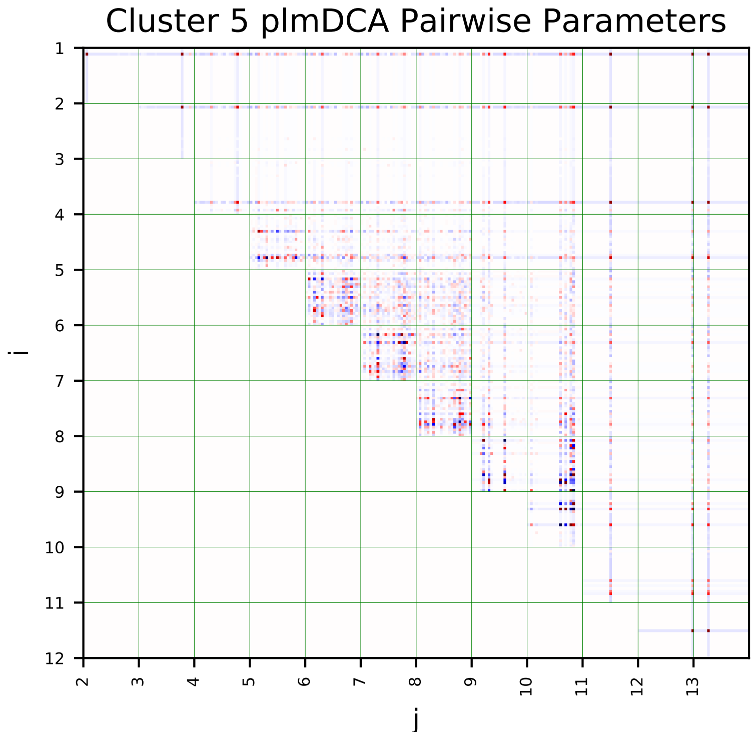
Cluster 5 plmDCA Pairwise Parameters
i
j

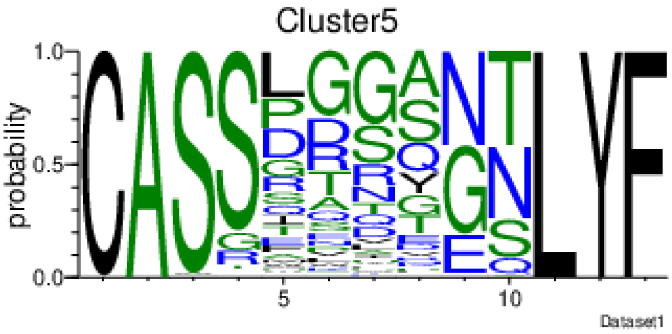
Cluster5
probability
Dataset1

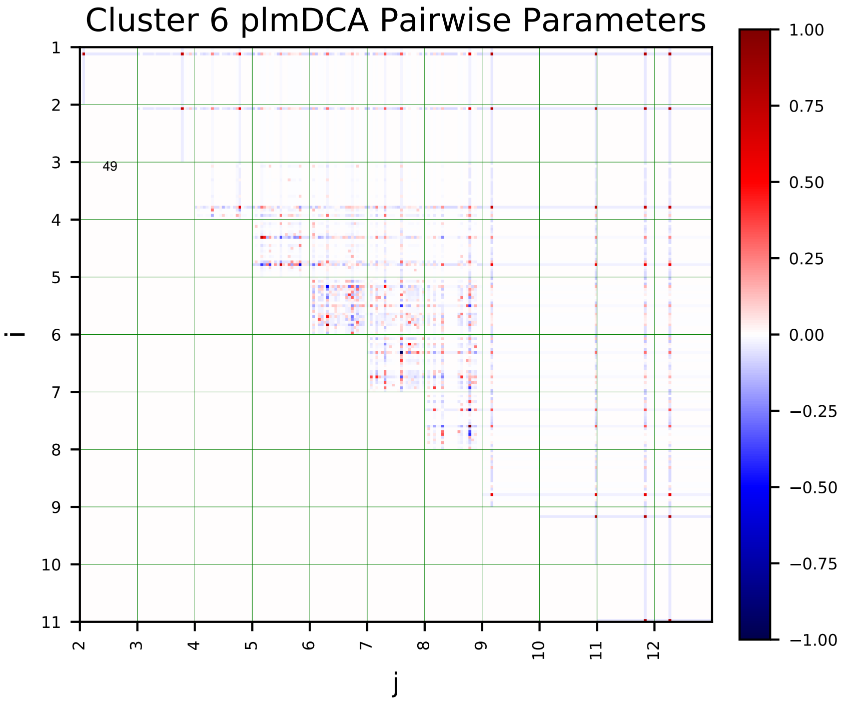
Cluster 6 plmDCA Pairwise Parameters
i
j
49
1.00
0.75
0.50
0.25
0.00
−0.25
−0.50
−0.75
−1.00

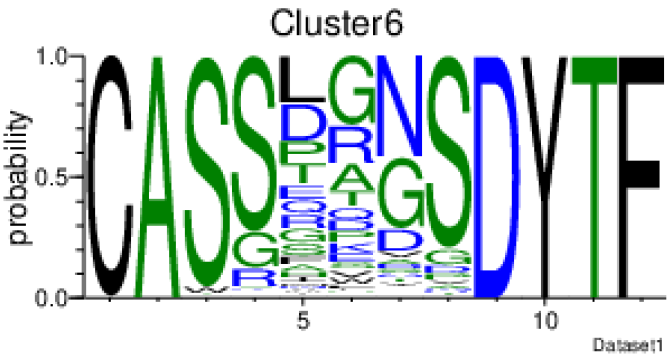
Cluster6
probability
Dataset1

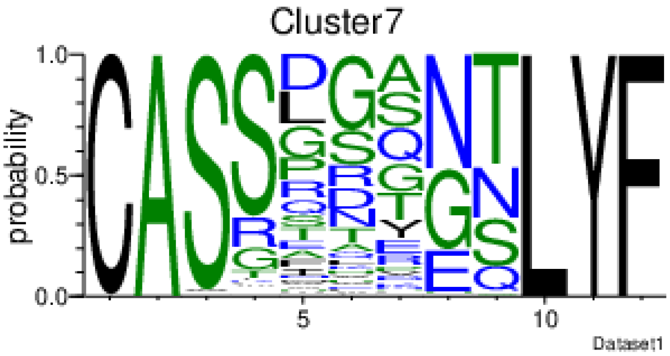
Cluster7
probability
Dataset1

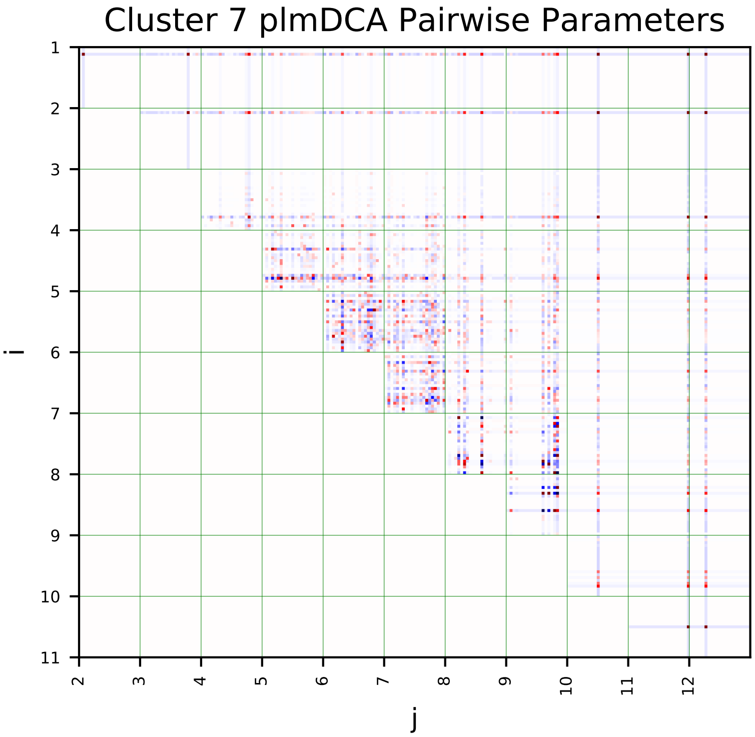
Cluster 7 plmDCA Pairwise Parameters
i
j

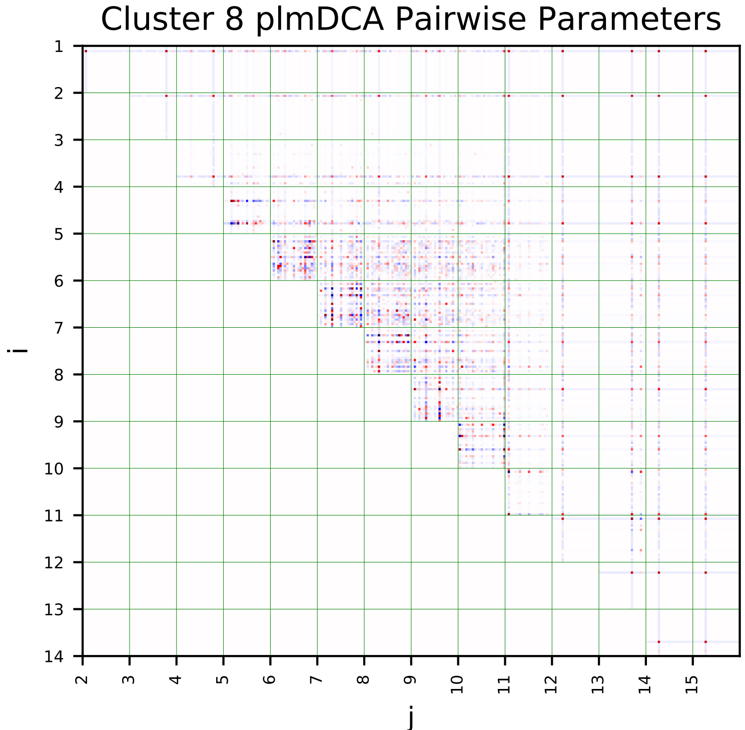
Cluster 8 plmDCA Pairwise Parameters
i
j

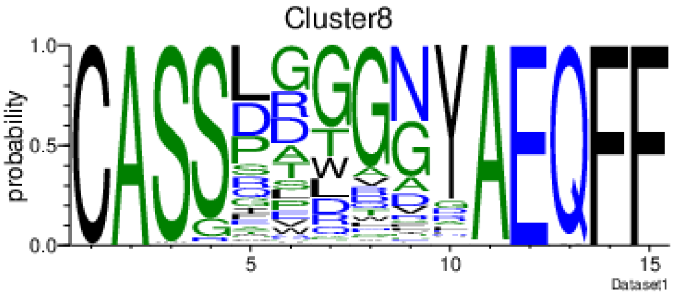
Cluster8
probability
Dataset1

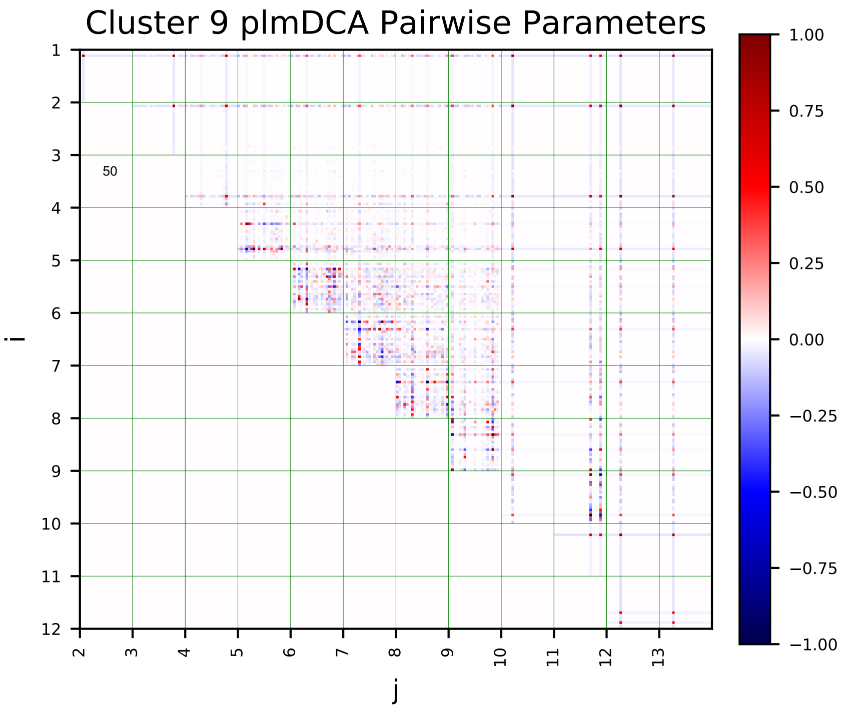
Cluster 9 plmDCA Pairwise Parameters
i
j
50

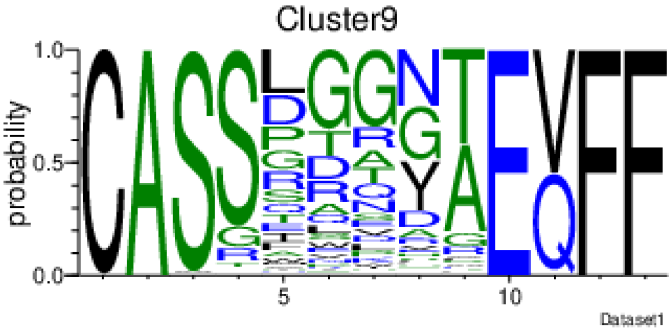
Cluster9
probability
Dataset1

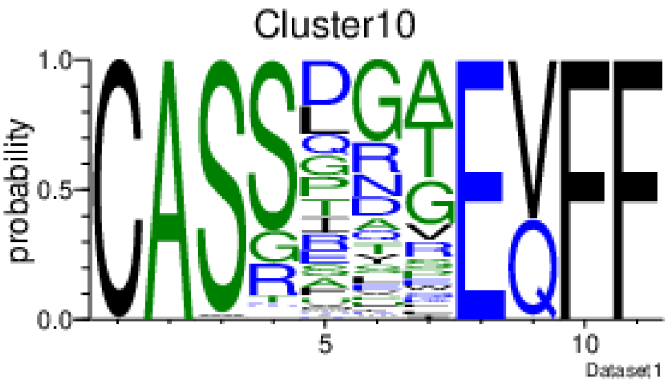
Cluster10
probability
Dataset1

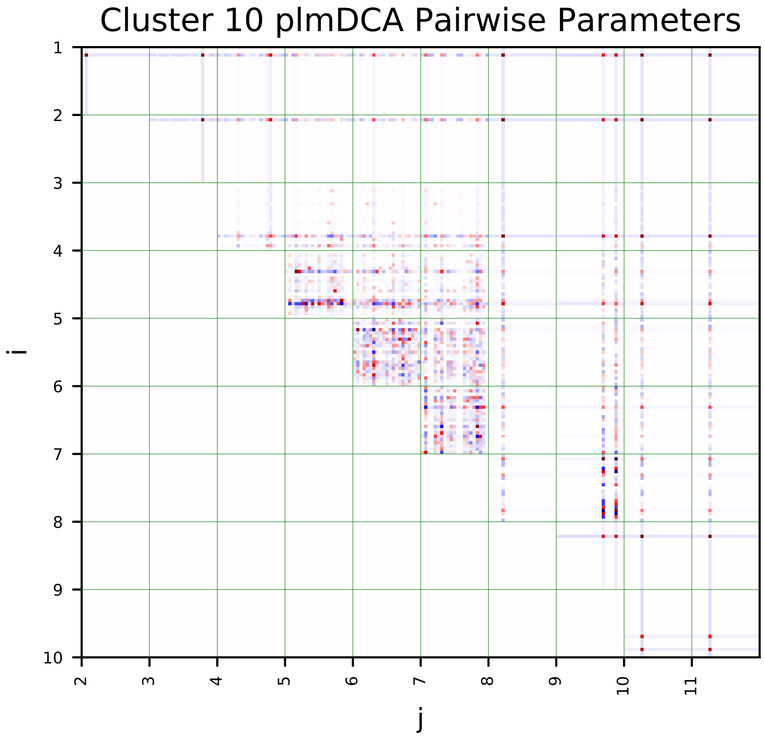
Cluster 10 plmDCA Pairwise Parameters
i
j

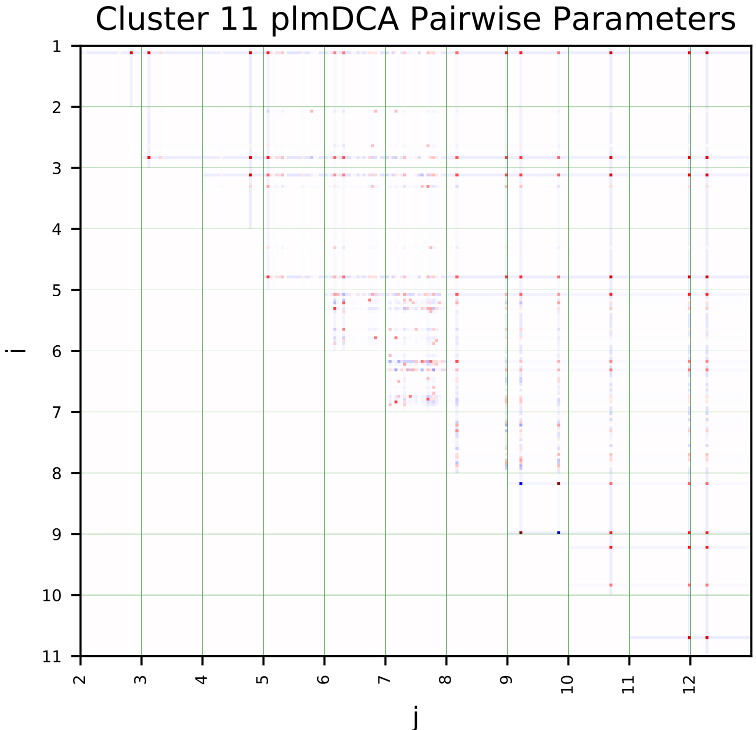
Cluster 11 plmDCA Pairwise Parameters
i
j

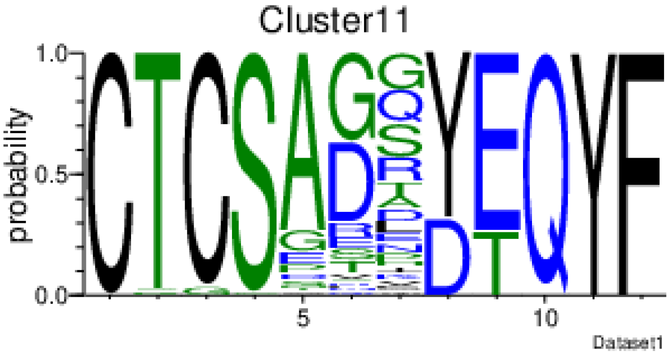
Cluster11
probability
Dataset1

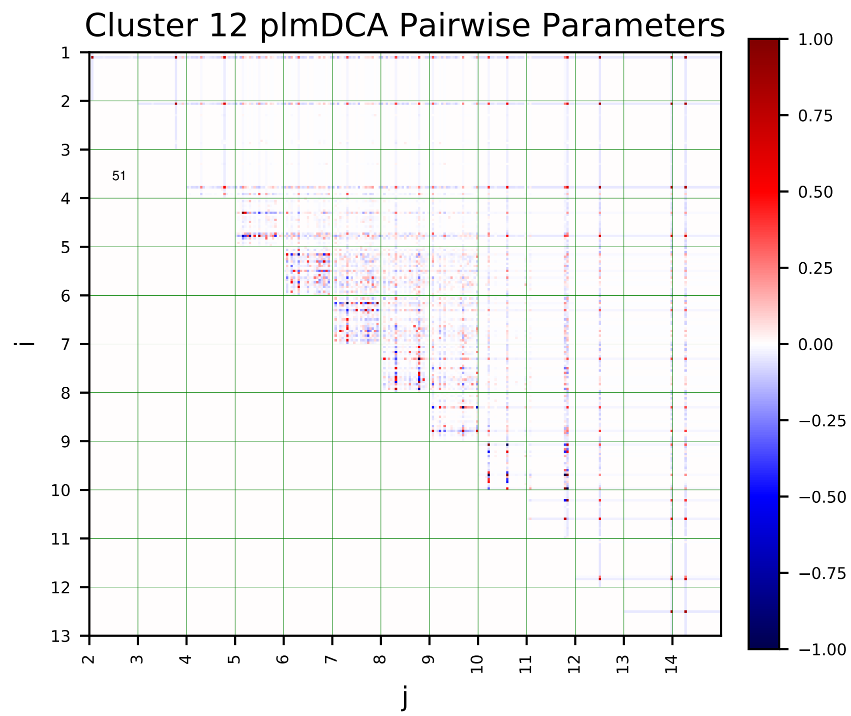
Cluster 12 plmDCA Pairwise Parameters
i
j
1.00
0.75
0.50
0.25
0.00
−0.25
−0.50
−0.75
−1.00

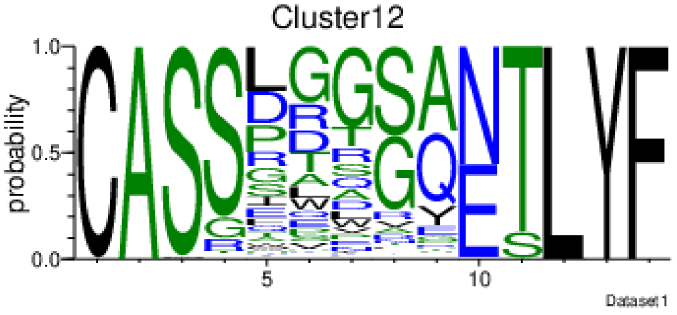
Cluster12
probability
Dataset1

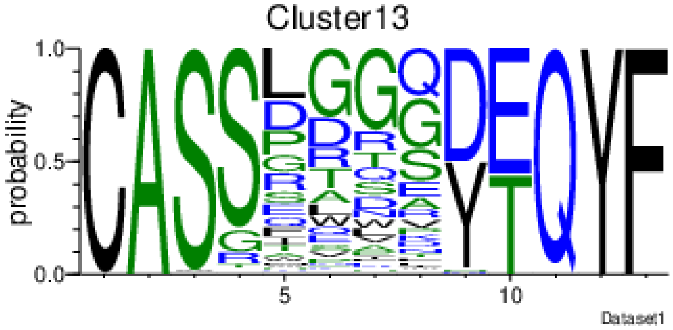
Cluster13
probability
Dataset1

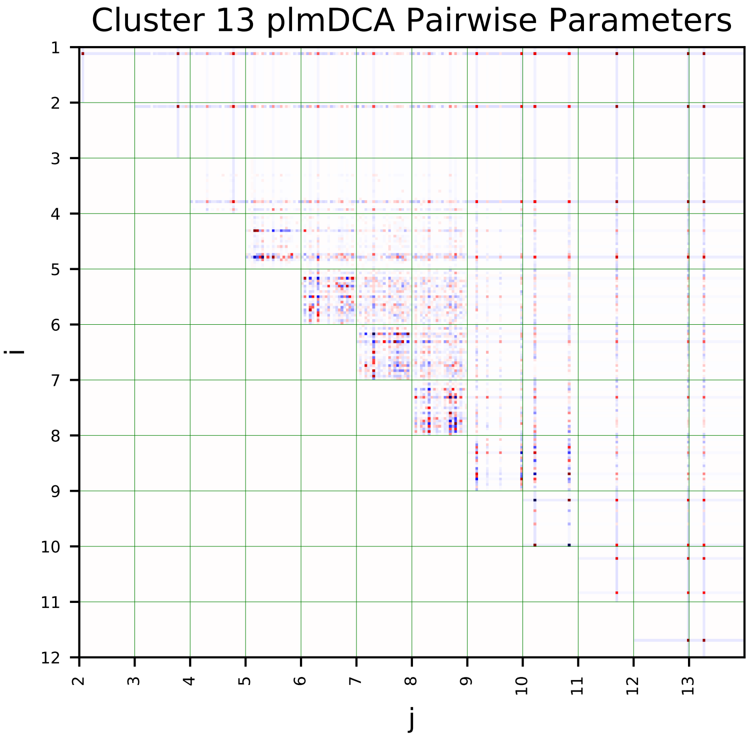
Cluster 13 plmDCA Pairwise Parameters
i
j

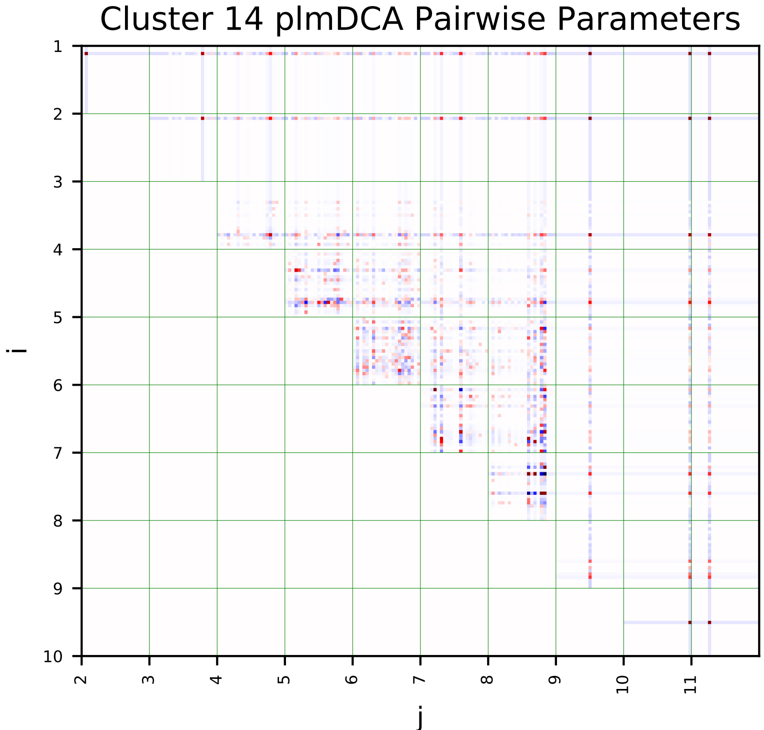
Cluster 14 plmDCA Pairwise Parameters
i
j

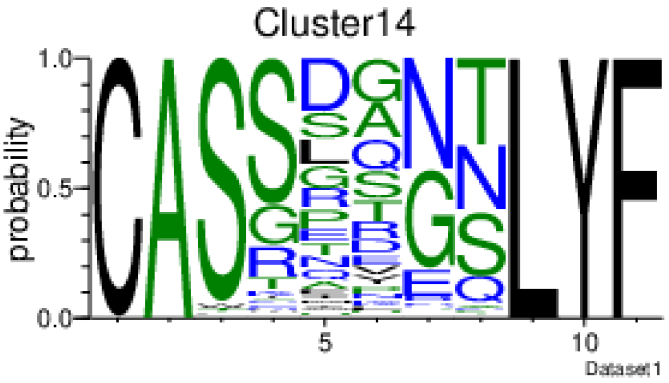
Cluster14
probability
Dataset1

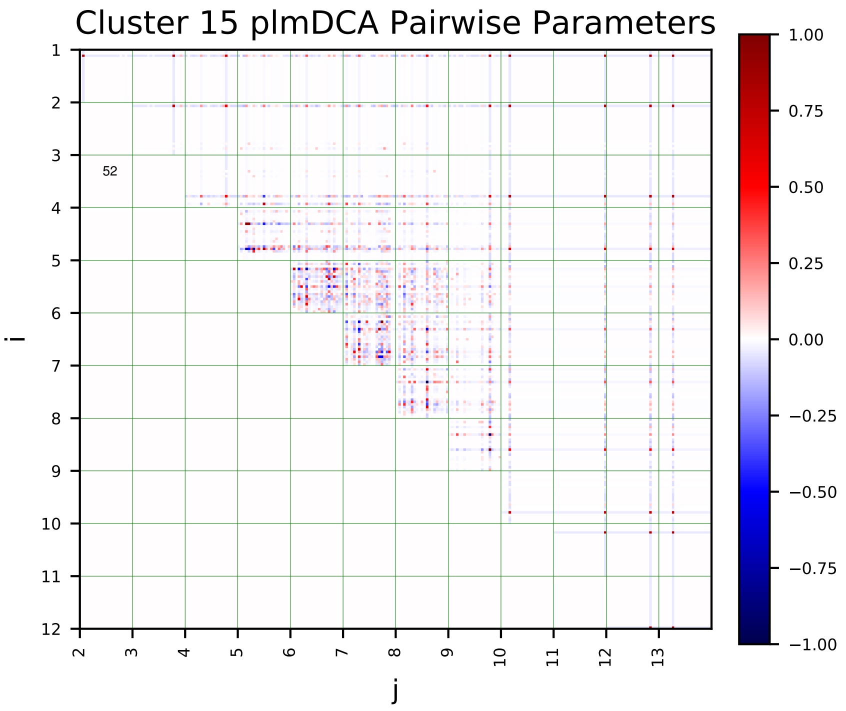
Cluster 15 plmDCA Pairwise Parameters
i
j
52

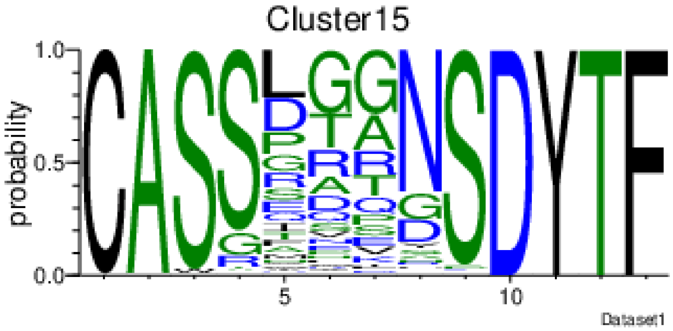
Cluster15
probability
Dataset1

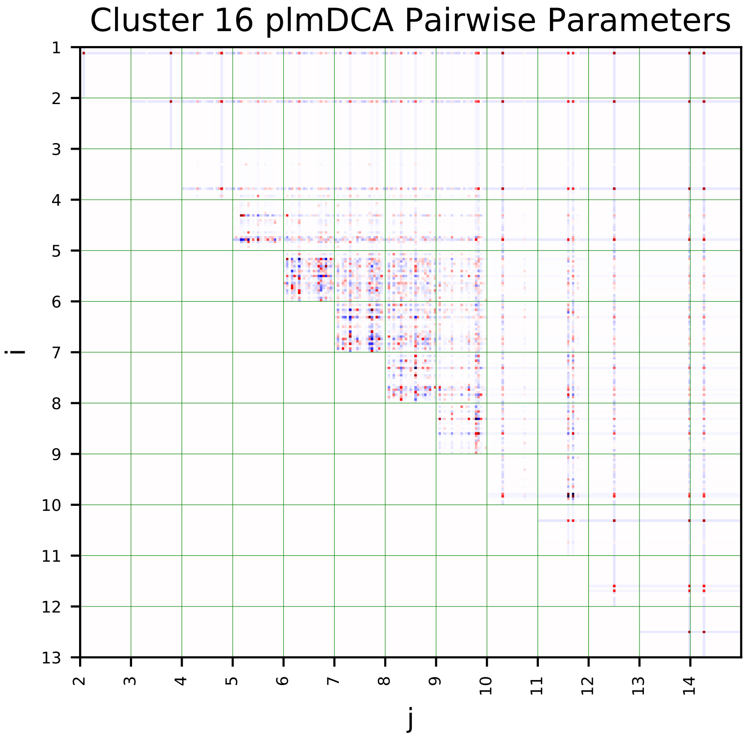
Cluster 16 plmDCA Pairwise Parameters
i
j

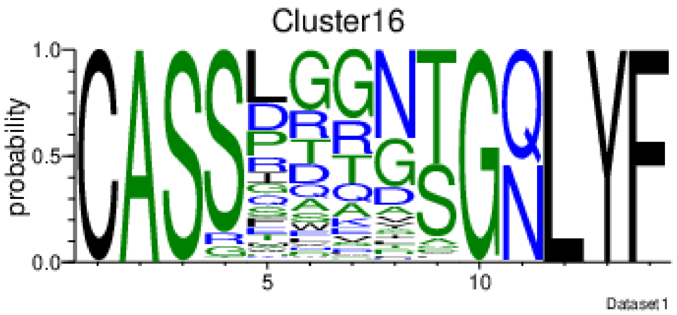
Cluster16
probability
Dataset1

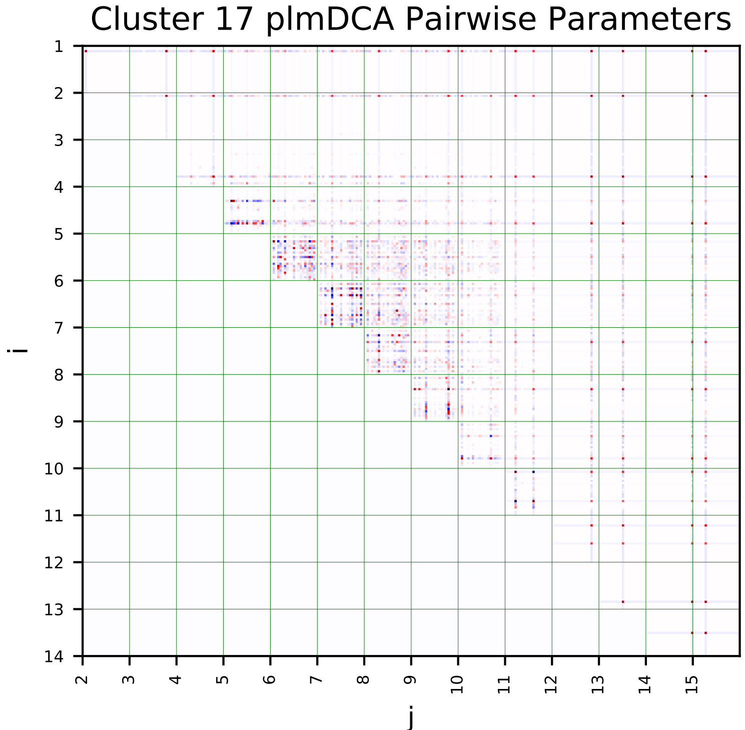
Cluster 17 plmDCA Pairwise Parameters
i
j

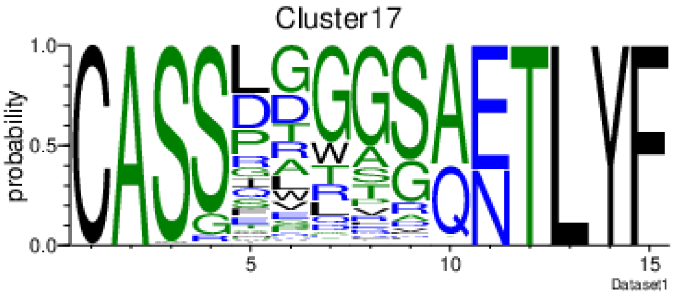
Cluster17
probability
Dataset1

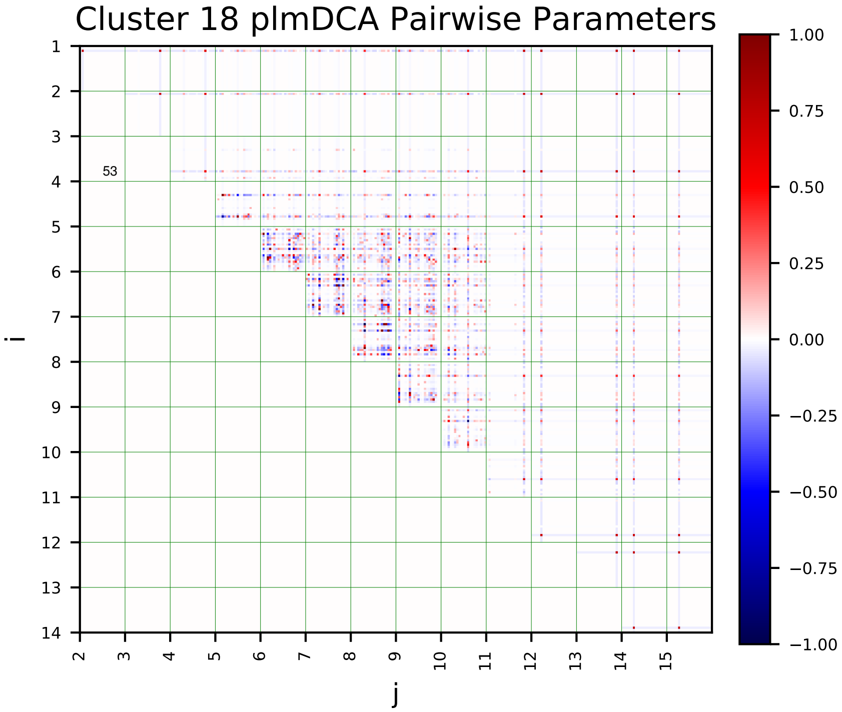
Cluster 18 plmDCA Pairwise Parameters
i
j
53
1.00
0.75
0.50
0.25
0.00
−0.25
−0.50
−0.75
−1.00

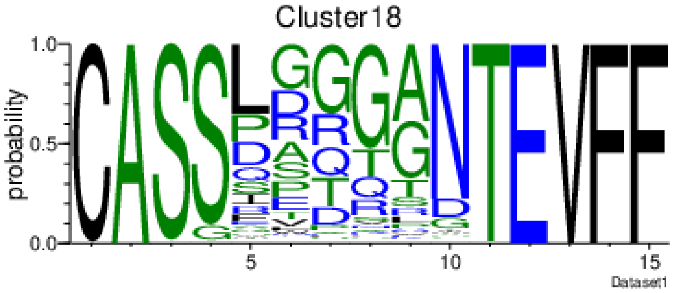
Cluster18
probability
Dataset1

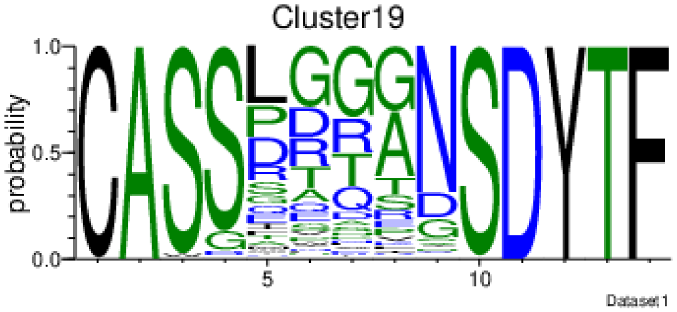
Cluster19
probability
Dataset1

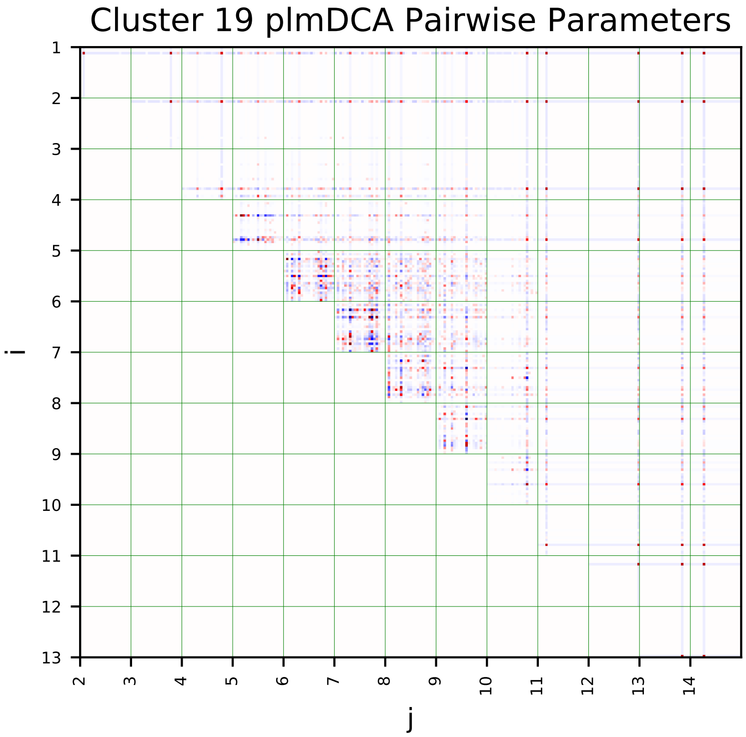
Cluster 19 plmDCA Pairwise Parameters
i
j

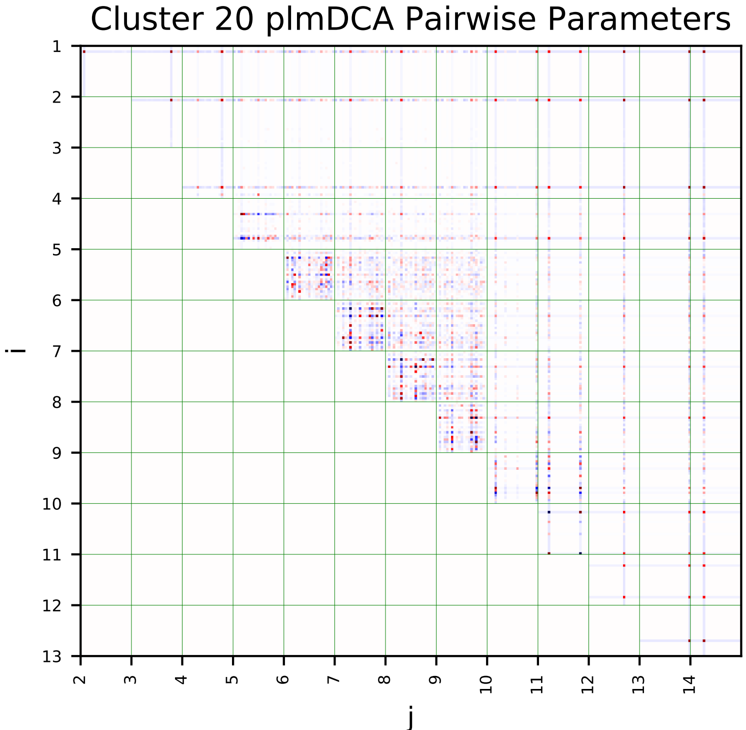
Cluster 20 plmDCA Pairwise Parameters
i
j

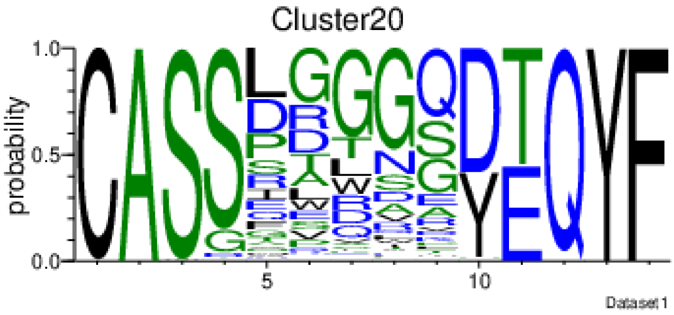
Cluster20
probability
Dataset1

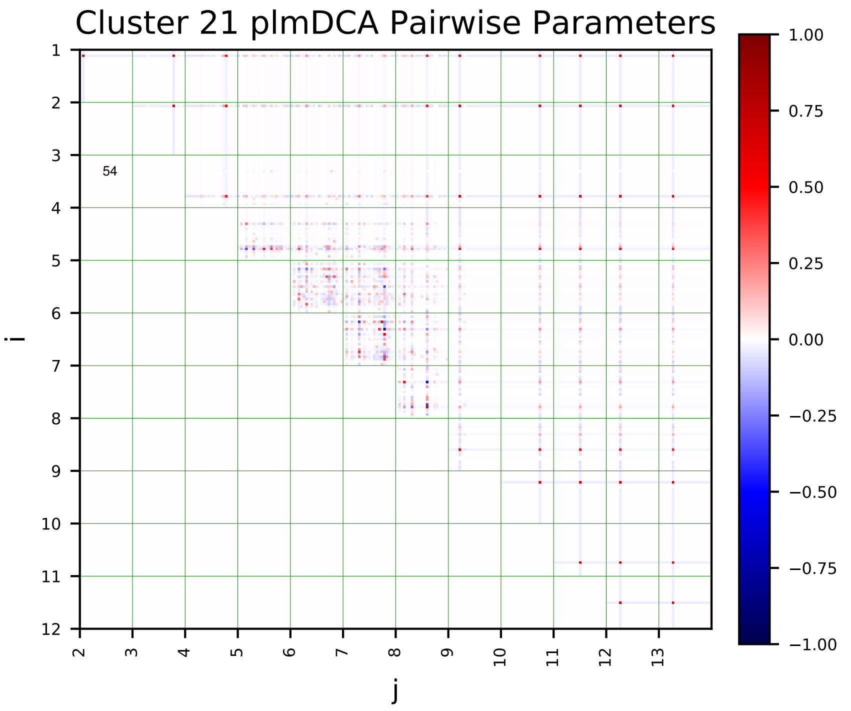
Cluster 21 plmDCA Pairwise Parameters
i
j
54
1.00
0.75
0.50
0.25
0.00
−0.25
−0.50
−0.75
−1.00

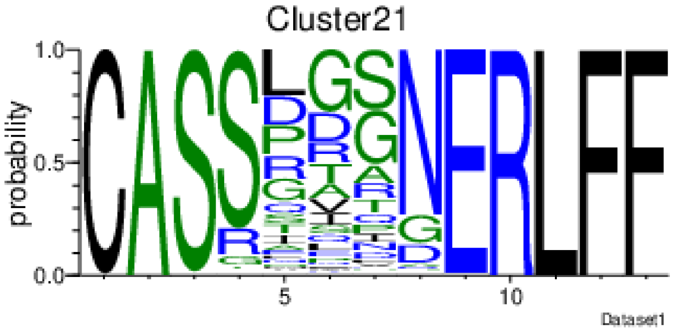
Cluster21
probability
Dataset1

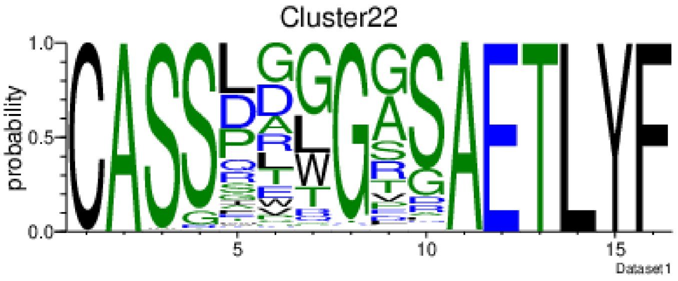
Cluster22
probability
Dataset1

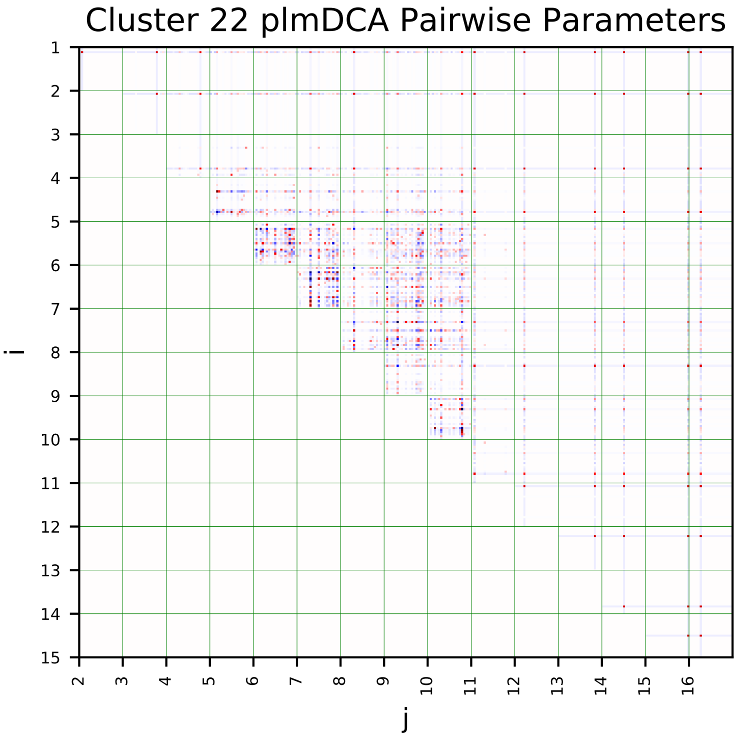
Cluster 22 plmDCA Pairwise Parameters
i
j

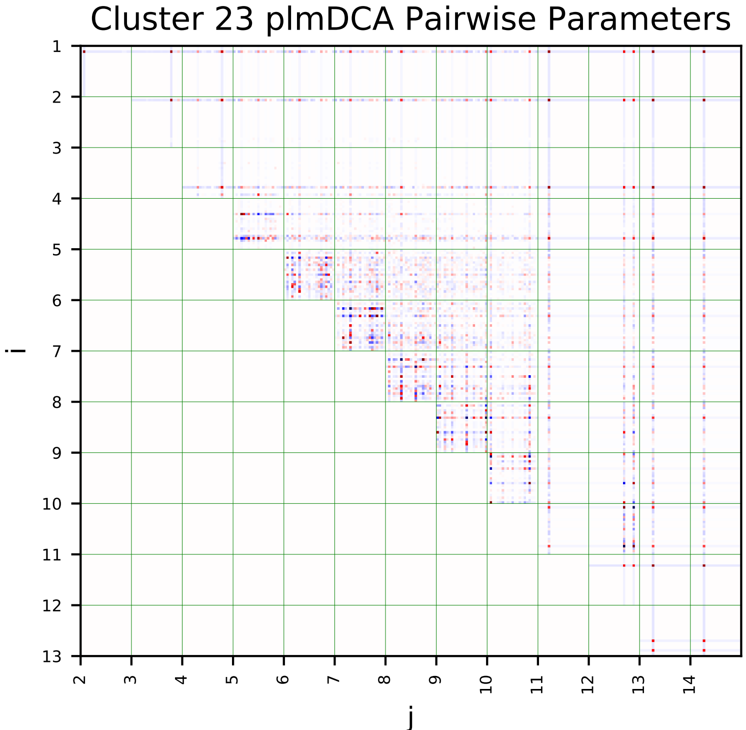
Cluster 23 plmDCA Pairwise Parameters
i
j

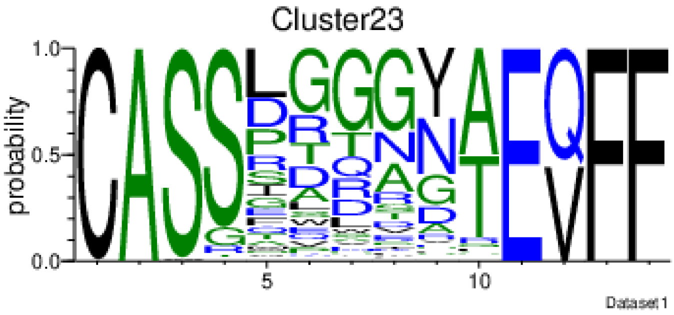
Cluster23
probability
Dataset1

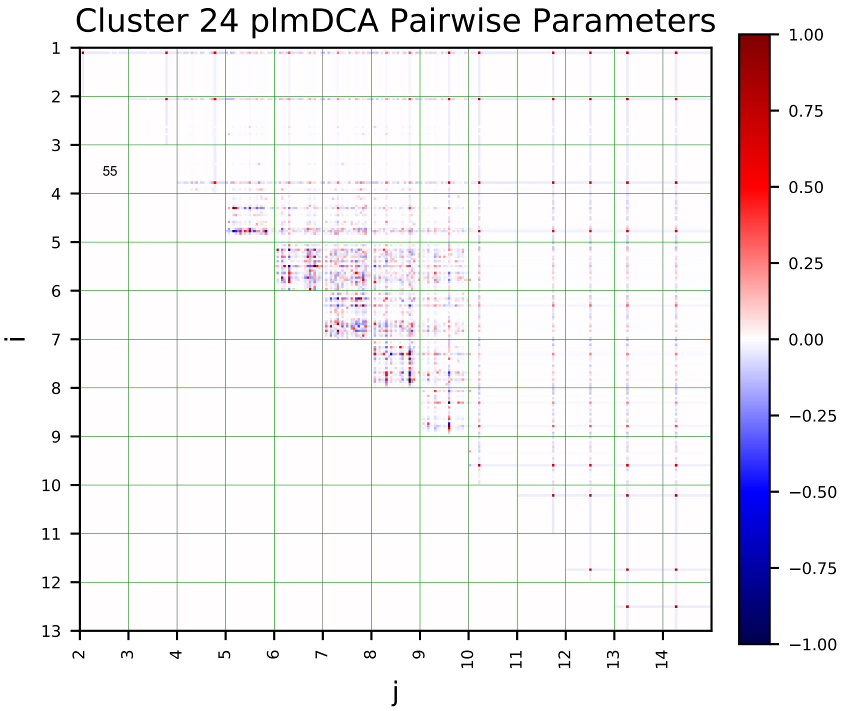
Cluster 24 plmDCA Pairwise Parameters
i
j

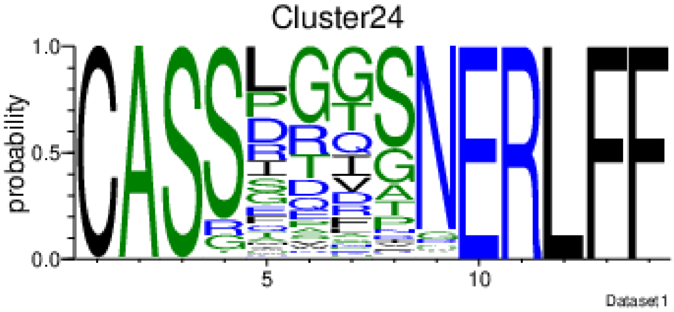
Cluster24
probability
Dataset1

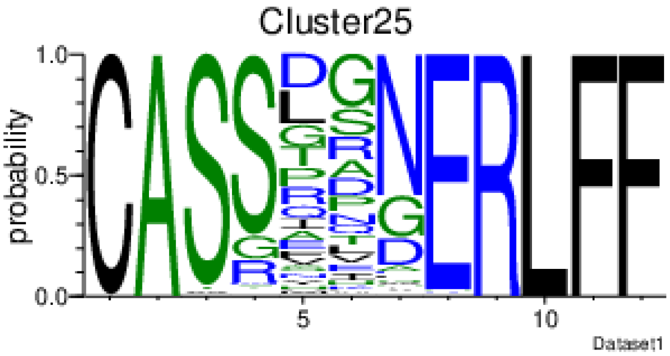
Cluster25
probability
Dataset1

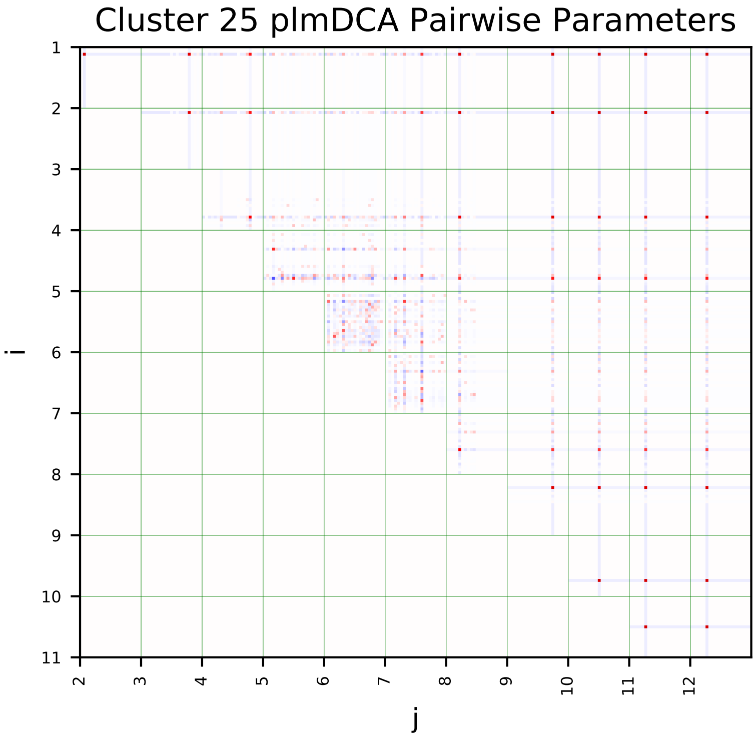
Cluster 25 plmDCA Pairwise Parameters
i
j

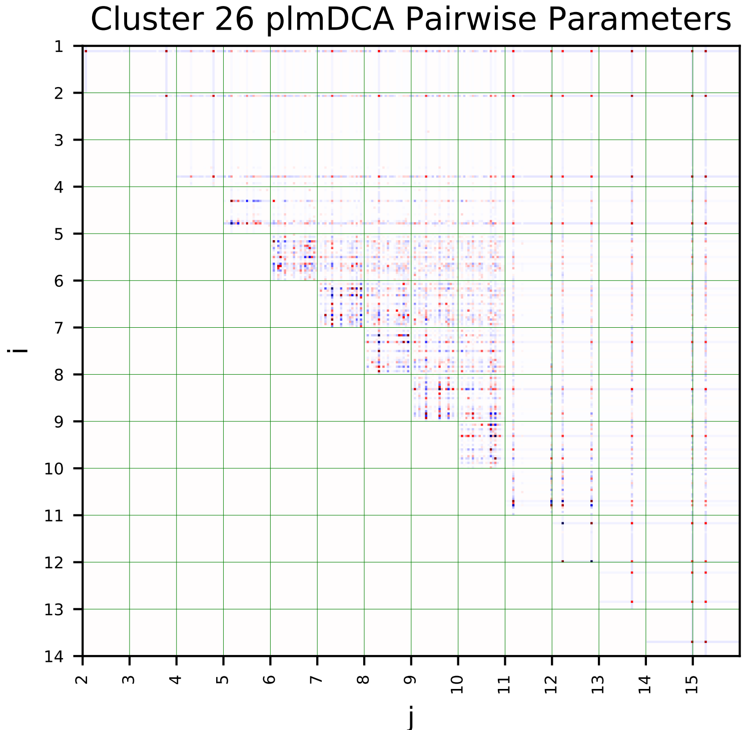
Cluster 26 plmDCA Pairwise Parameters
i
j

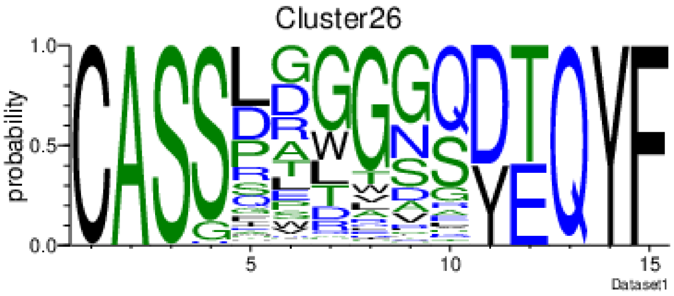
Cluster26
probability
Dataset1

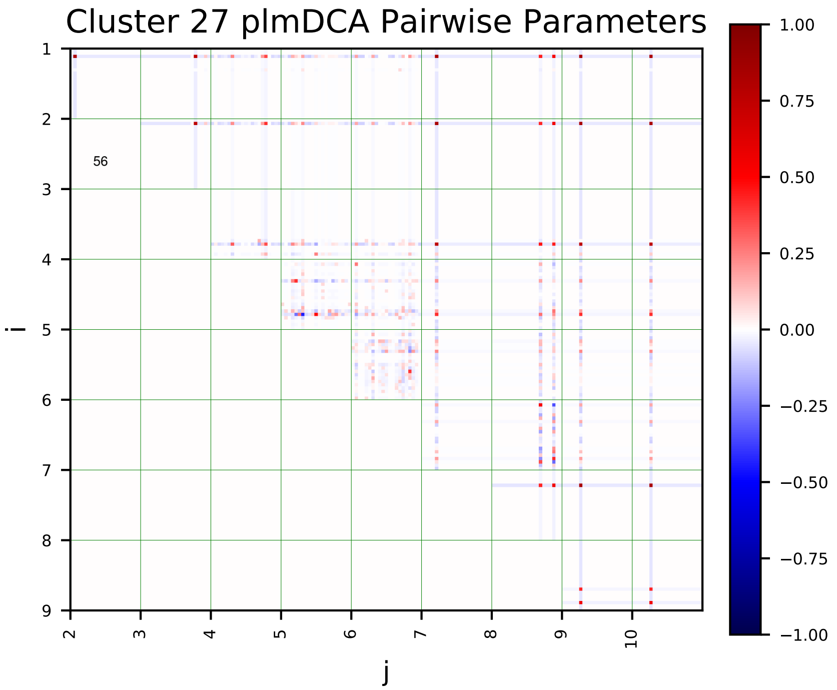
Cluster 27 plmDCA Pairwise Parameters
i
j

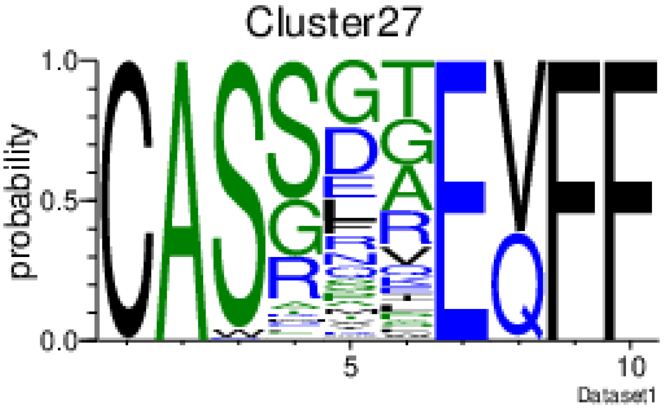
Cluster27
probability
Dataset1

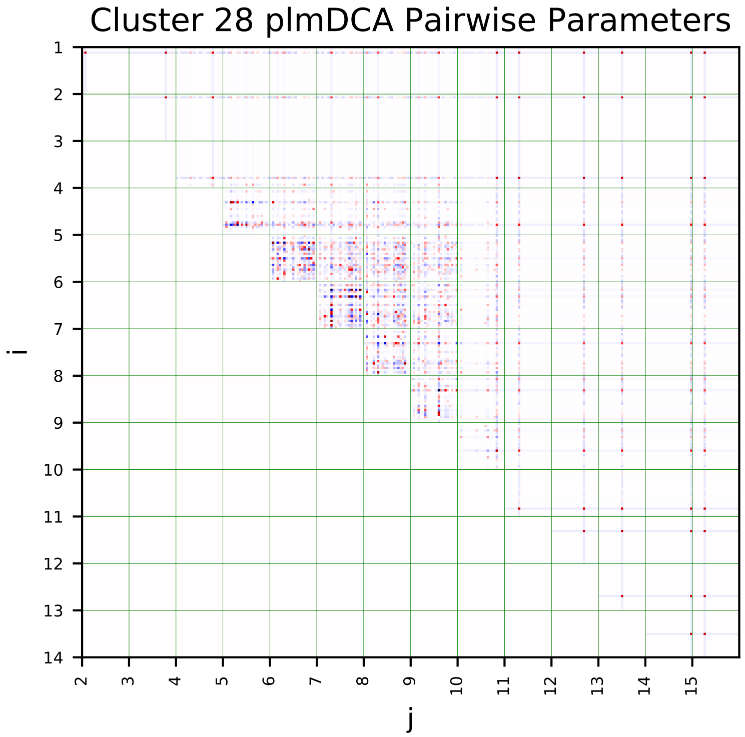
Cluster 28 plmDCA Pairwise Parameters
i
j

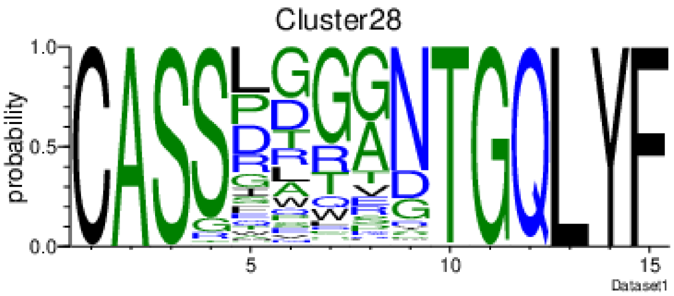
Cluster28
probability
Dataset1

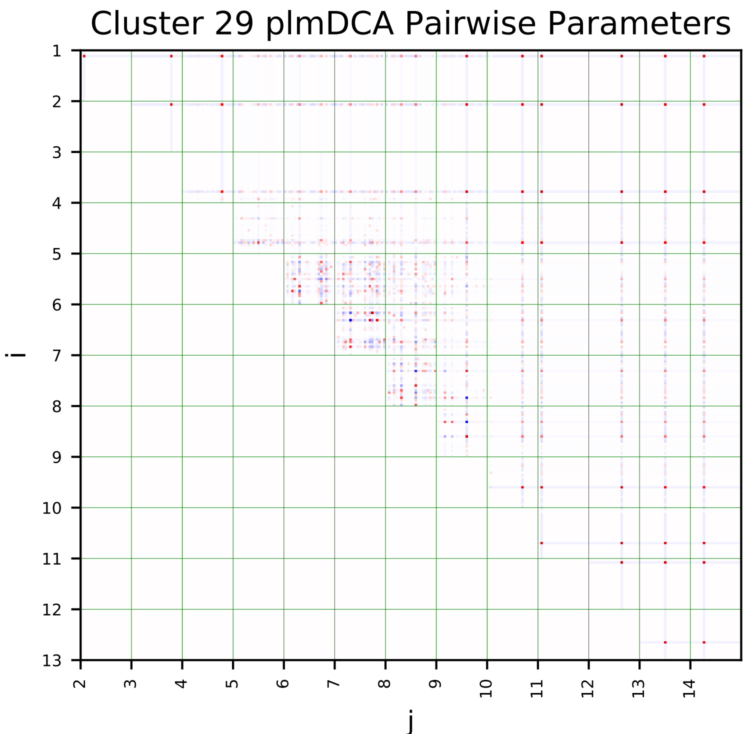
Cluster 29 plmDCA Pairwise Parameters
i
j

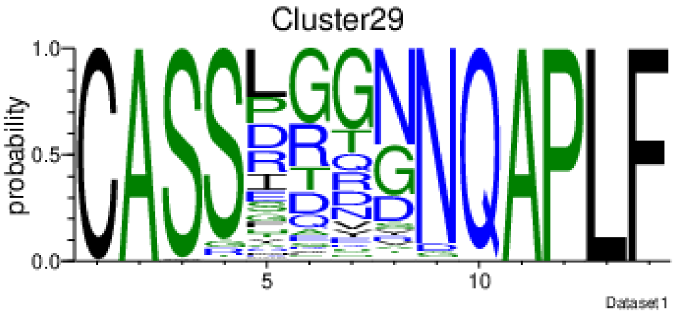
Cluster29
probability
Dataset1

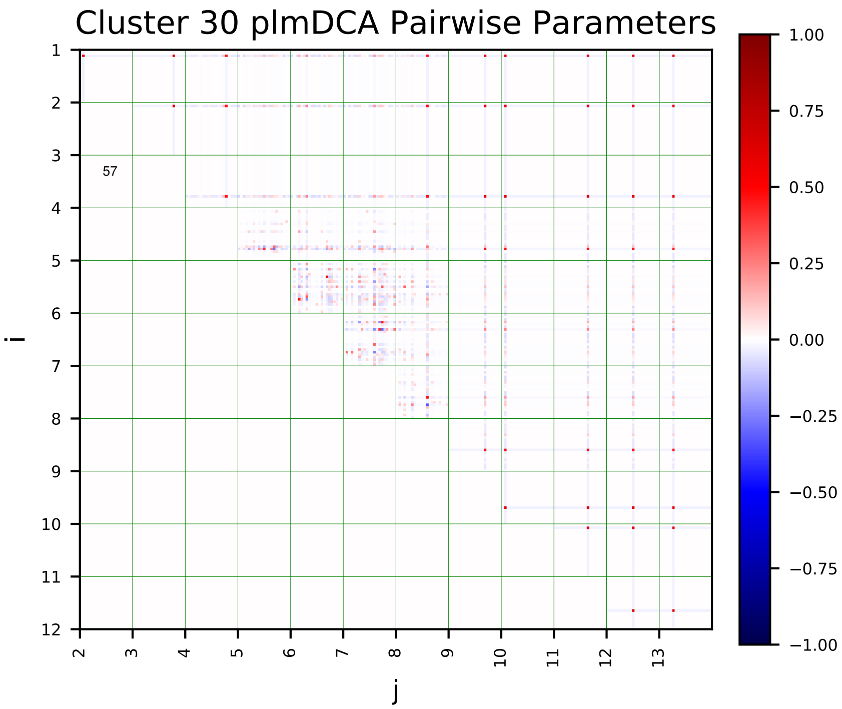
Cluster 30 plmDCA Pairwise Parameters
i
j
57

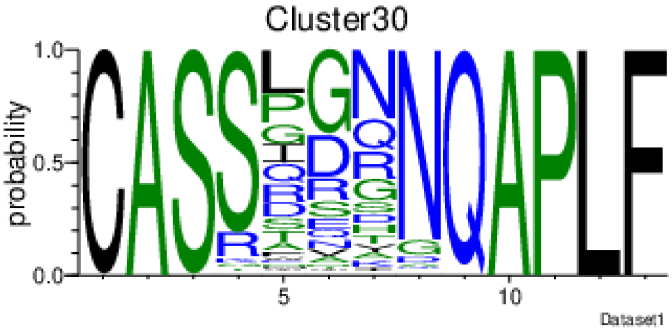
Cluster30
probability
Dataset1

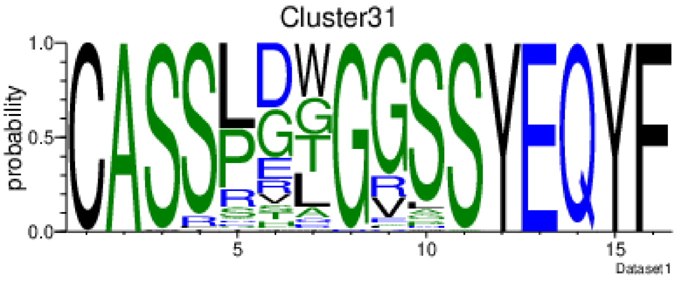
Cluster31
probability
Dataset1

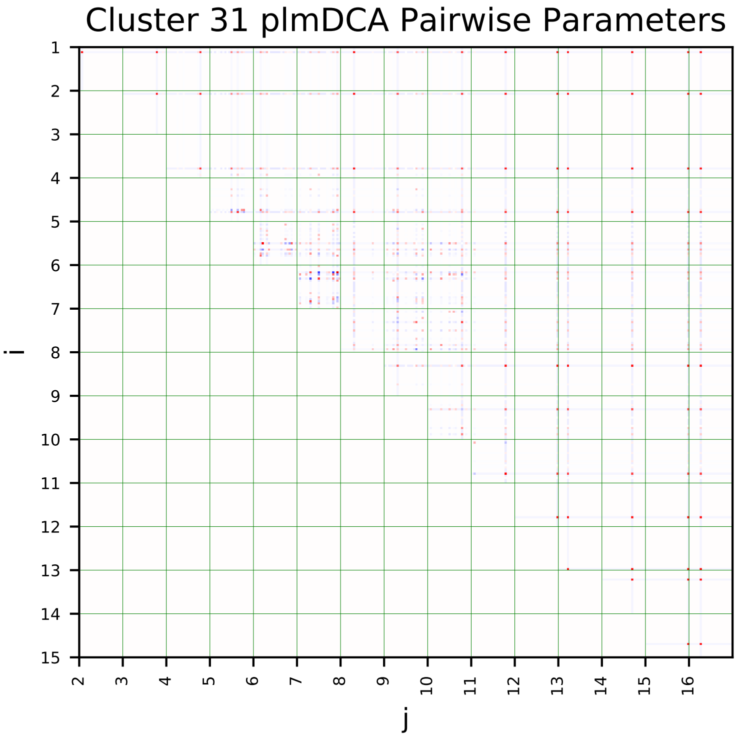
Cluster 31 plmDCA Pairwise Parameters
i
j

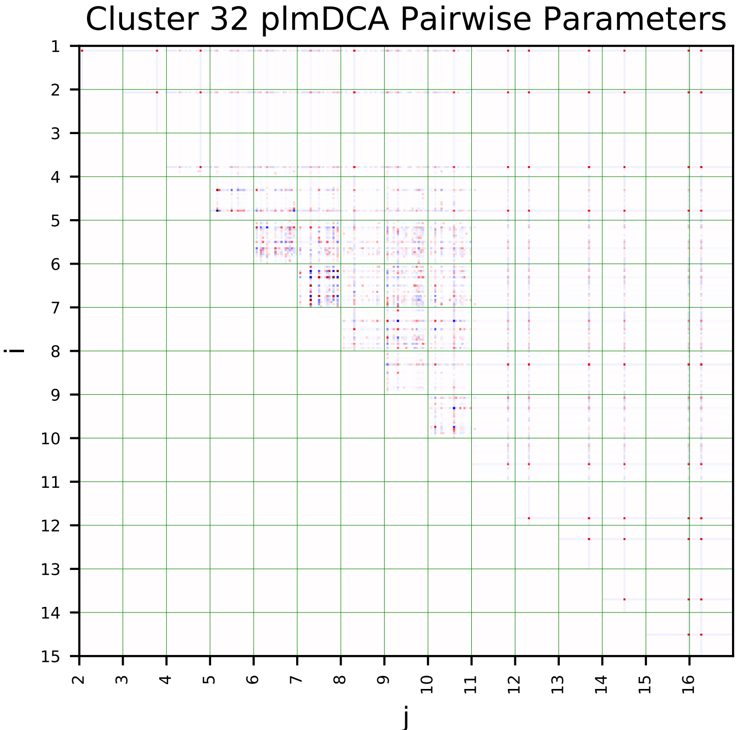
Cluster 32 plmDCA Pairwise Parameters
i
j

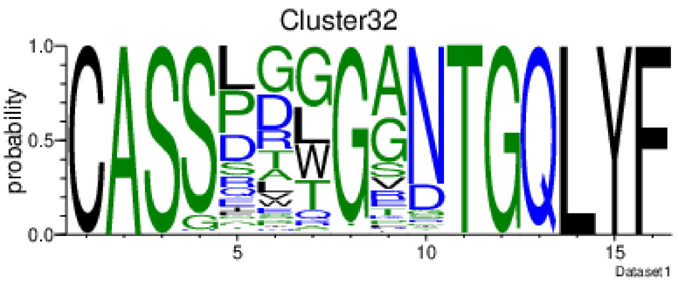
Cluster32
probability
Dataset1

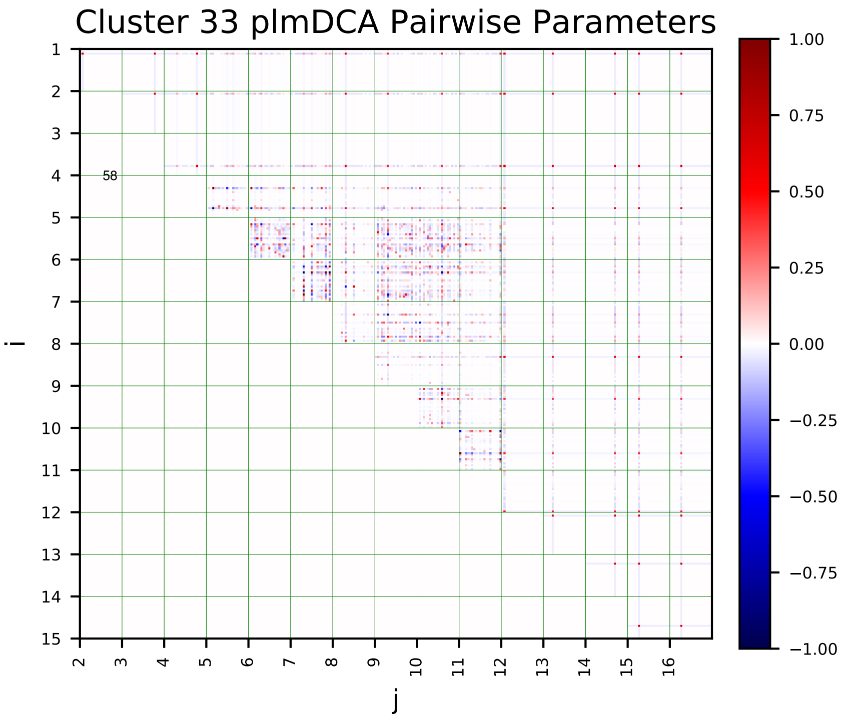
Cluster 33 plmDCA Pairwise Parameters
58
i
j

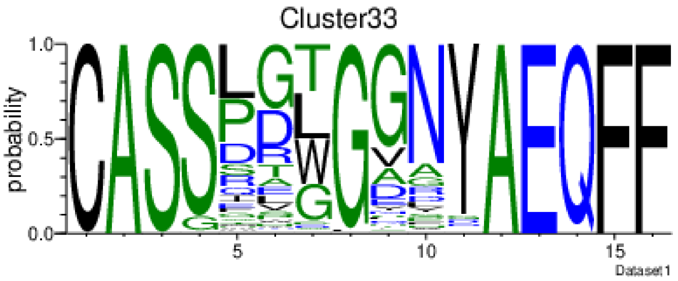
Cluster33
probability
Dataset1

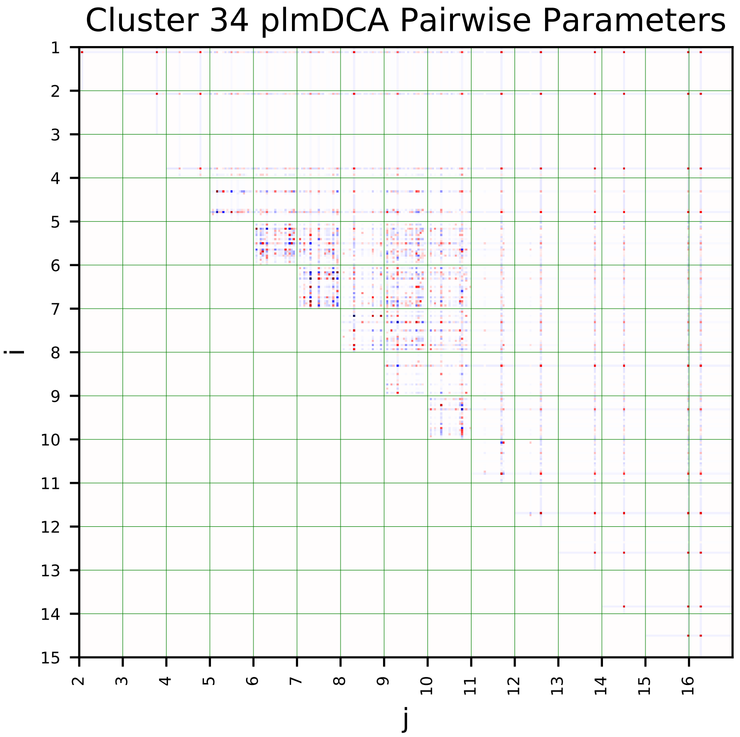
Cluster 34 plmDCA Pairwise Parameters
i
j

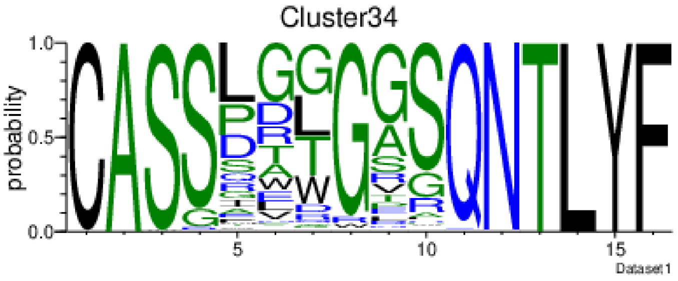
Cluster34
probability
Dataset1

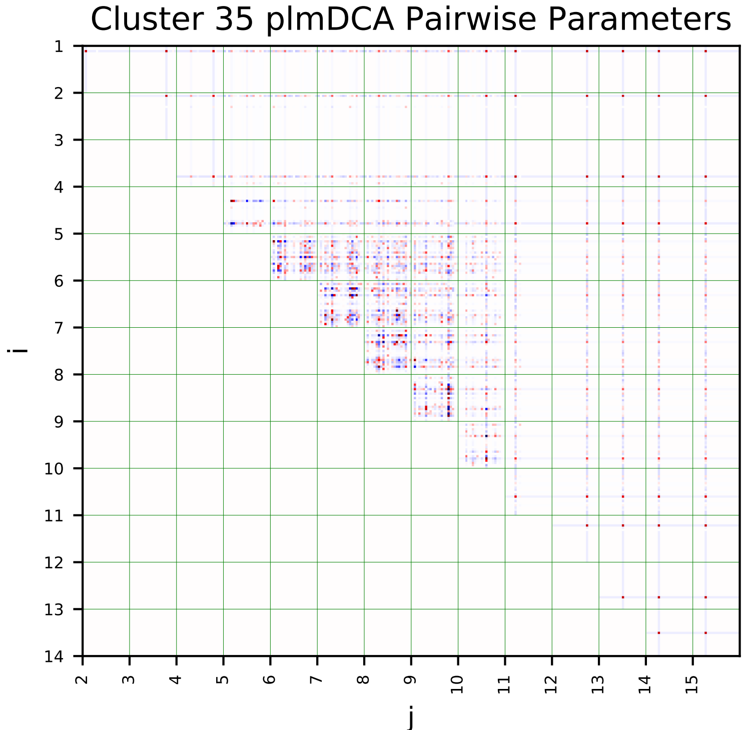
Cluster 35 plmDCA Pairwise Parameters
i
j

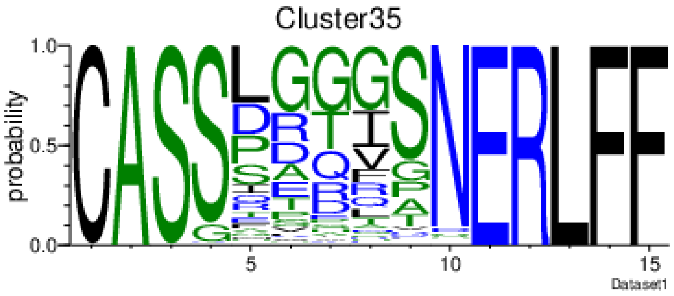
Cluster35
probability
Dataset1

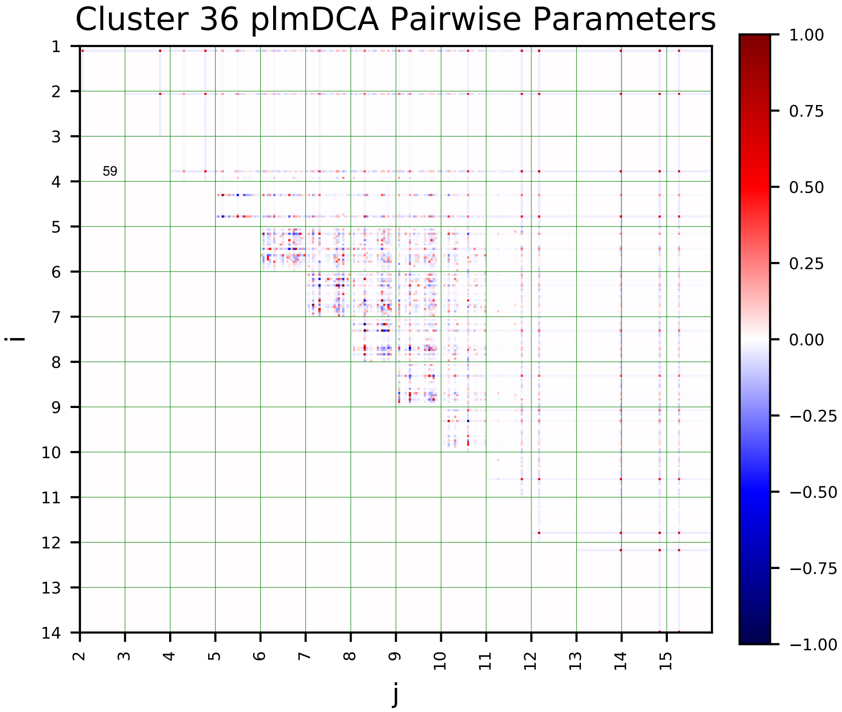
Cluster 36 plmDCA Pairwise Parameters
i
j
59
1.00
0.75
0.50
0.25
0.00
−0.25
−0.50
−0.75
−1.00

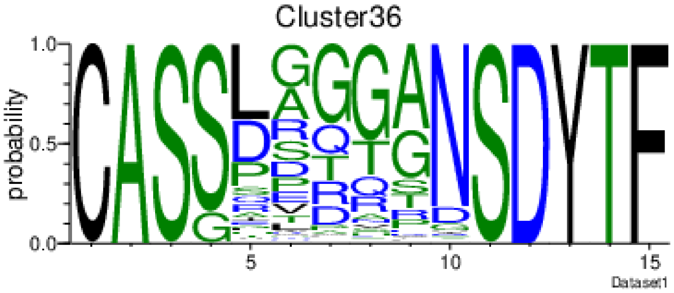
Cluster36
probability
Dataset1

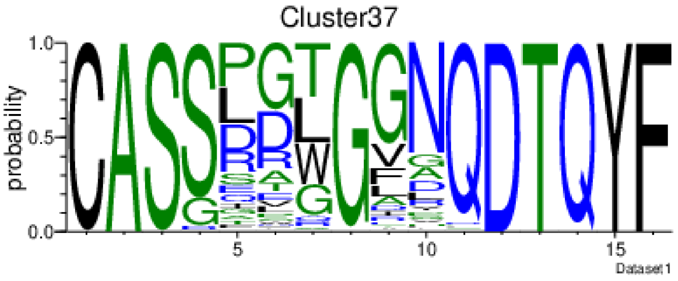
Cluster37
probability
Dataset1

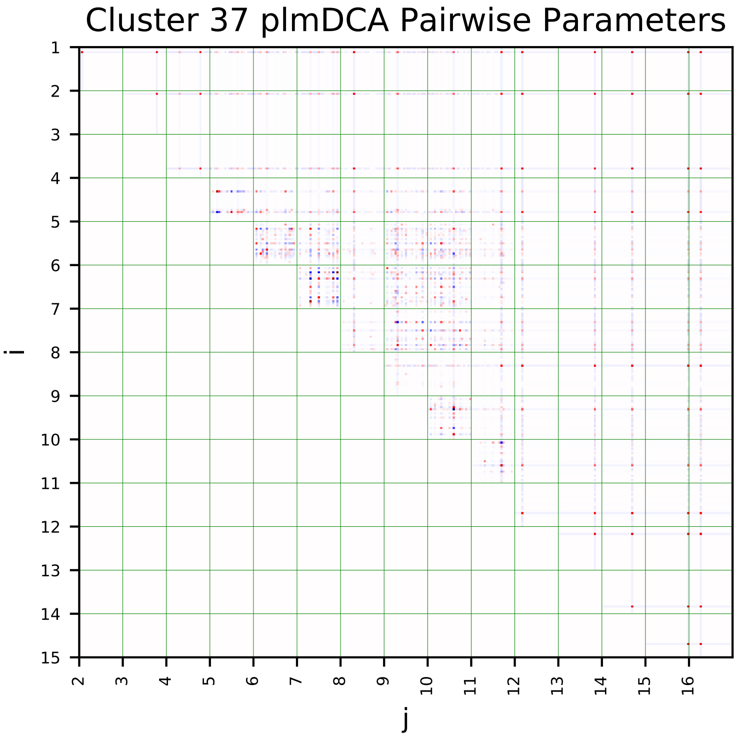
Cluster 37 plmDCA Pairwise Parameters
i
j

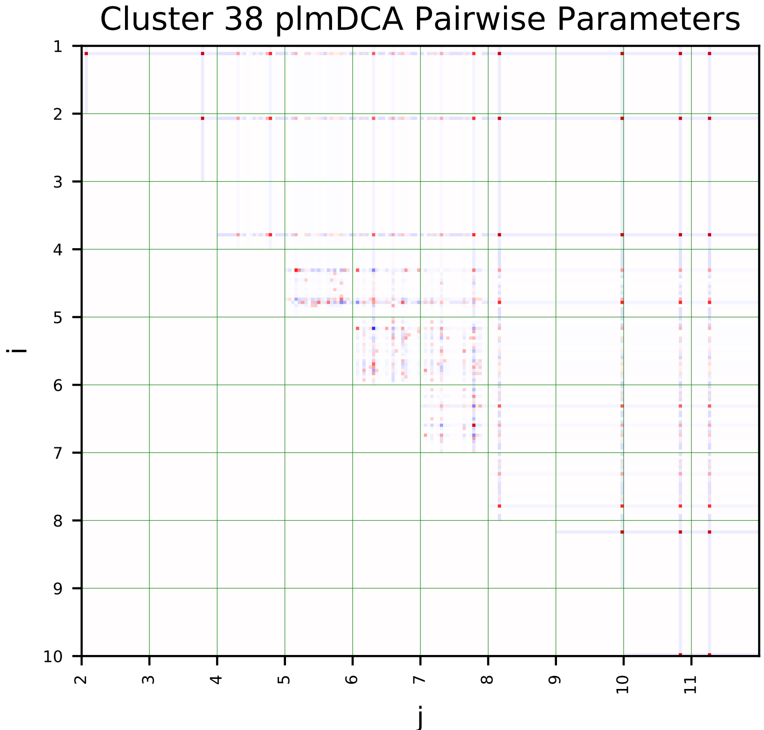
Cluster 38 plmDCA Pairwise Parameters
i
j

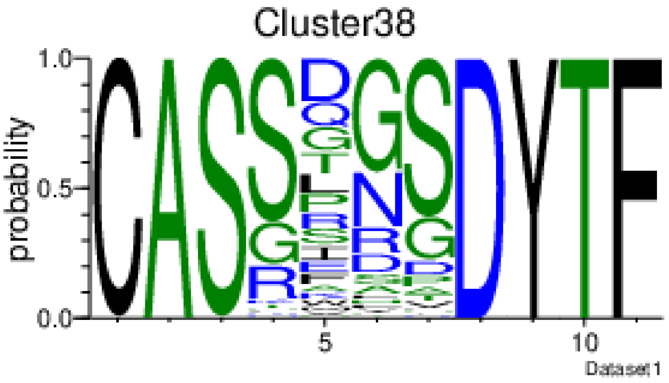
Cluster38
probability
Dataset1

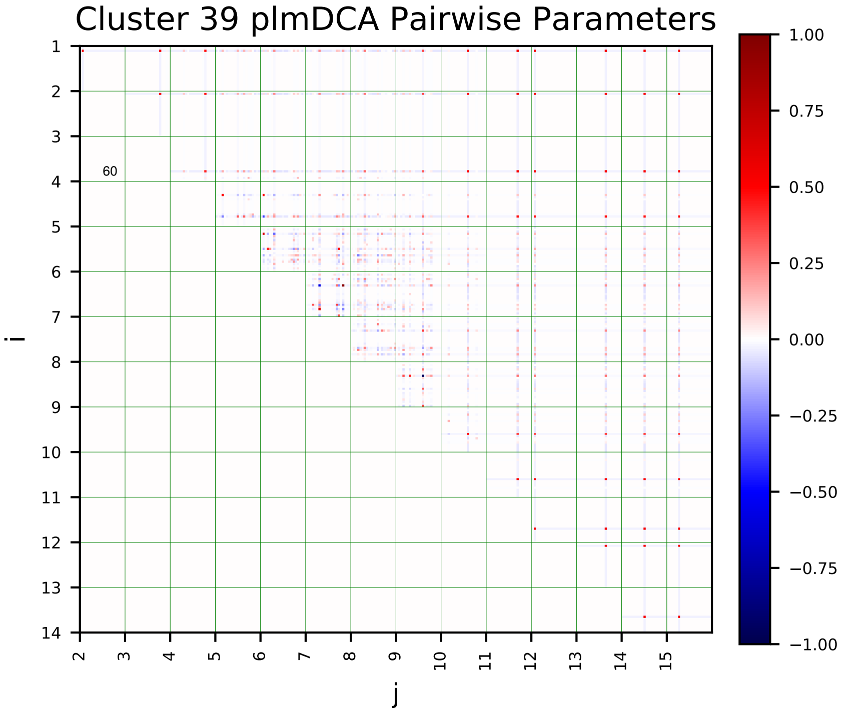
Cluster 39 plmDCA Pairwise Parameters
i
j
60
1.00
0.75
0.50
0.25
0.00
−0.25
−0.50
−0.75
−1.00

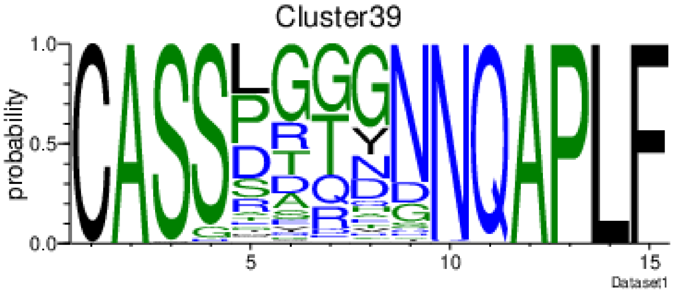
Cluster39
probability
Dataset1

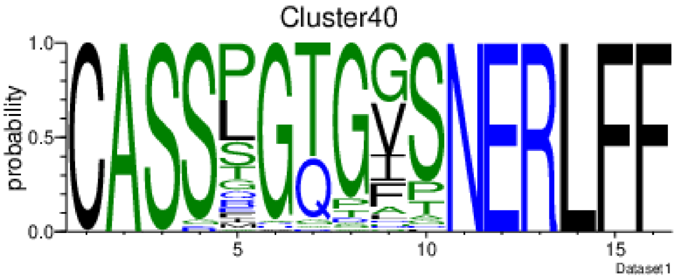
Cluster40
probability
Dataset1

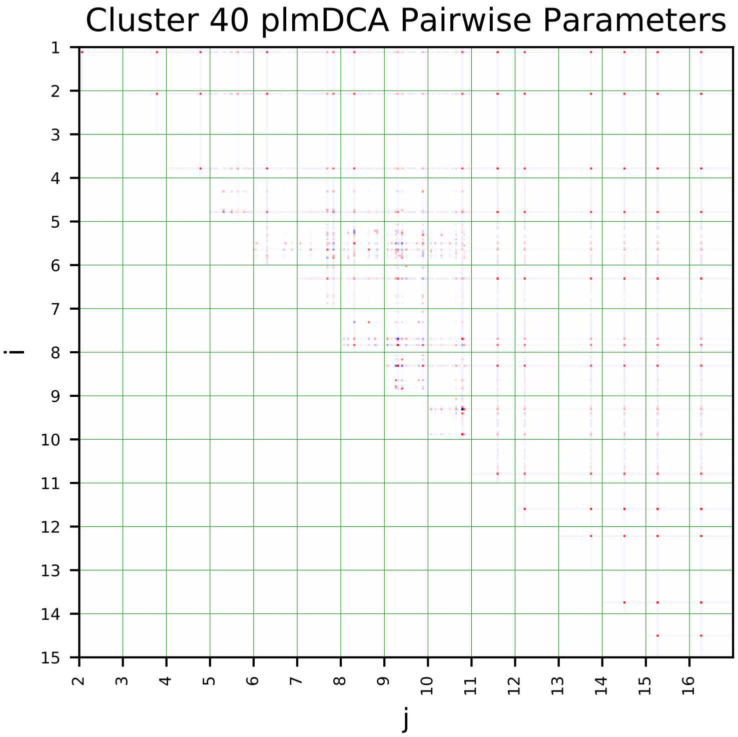
Cluster 40 plmDCA Pairwise Parameters
i
j

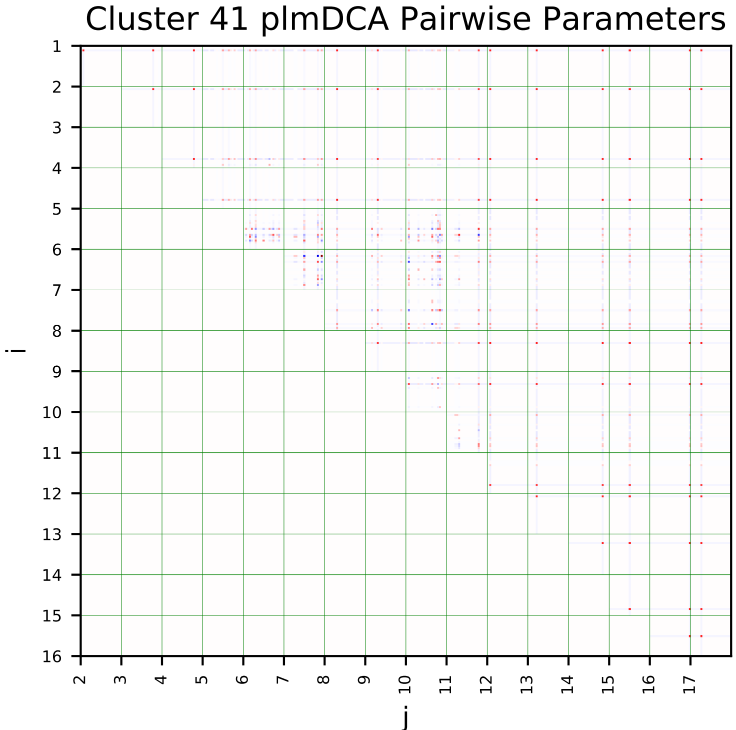
Cluster 41 plmDCA Pairwise Parameters
i
j

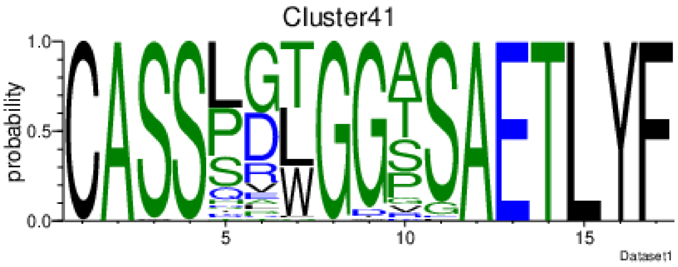
Cluster41
probability
Dataset1

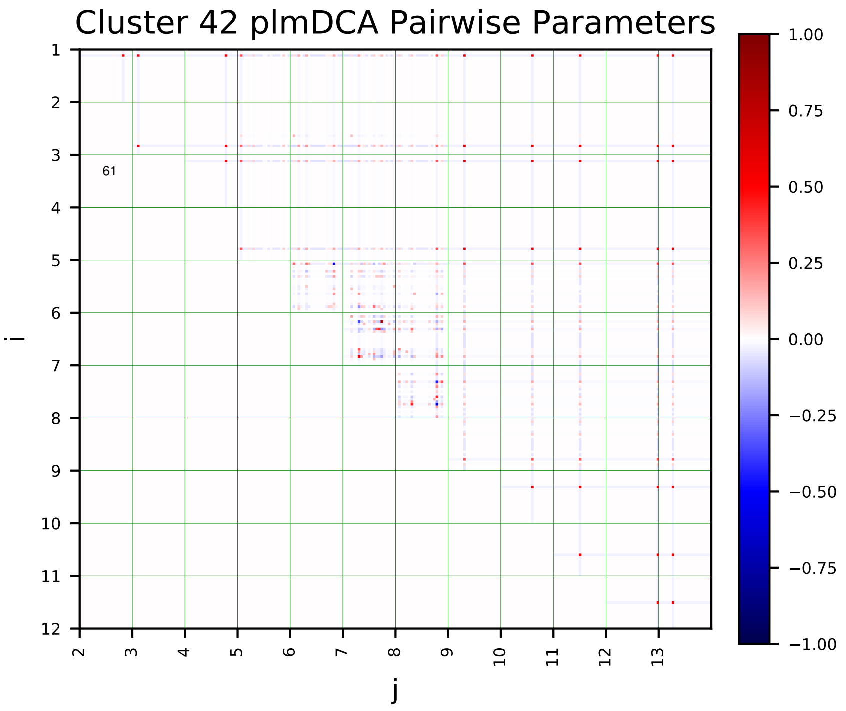
Cluster 42 plmDCA Pairwise Parameters
i
j
61
1.00
0.75
0.50
0.25
0.00
−0.25
−0.50
−0.75
−1.00

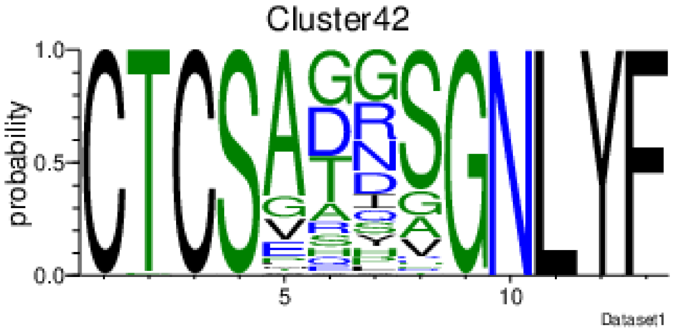
Cluster42
probability
1.0
0.5
0.0
5
10
Dataset1

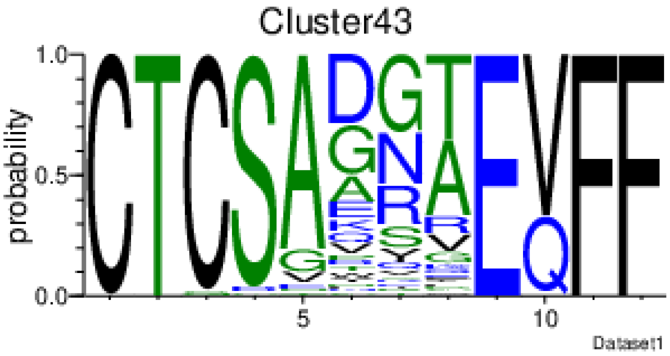
Cluster43
probability
1.0
0.5
0.0
5
10
Dataset1

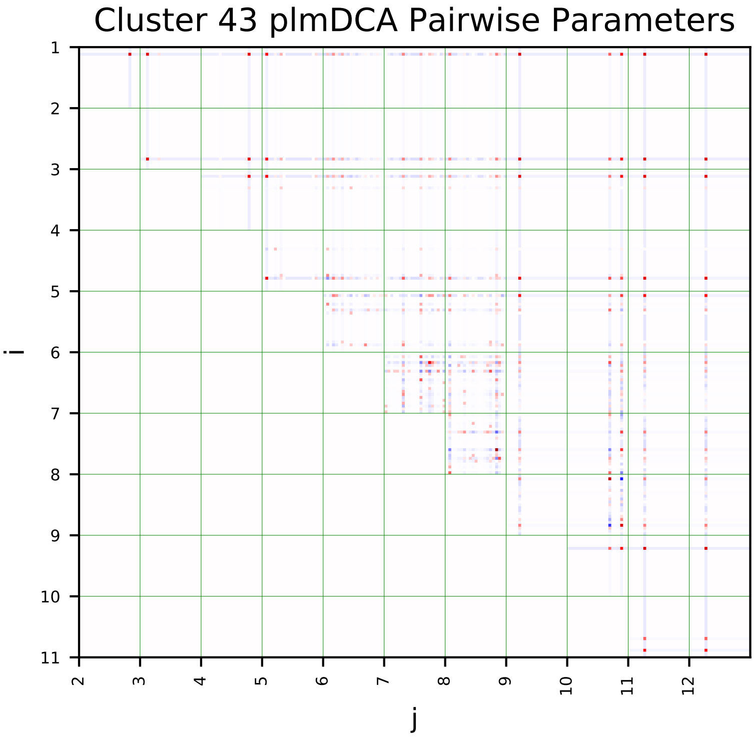
Cluster 43 plmDCA Pairwise Parameters
i
j

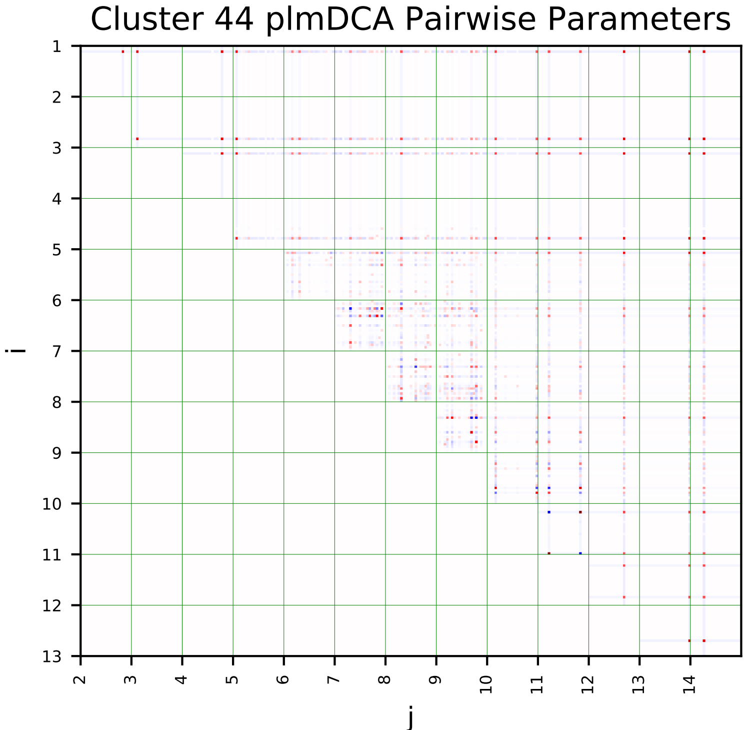
Cluster 44 plmDCA Pairwise Parameters
i
j

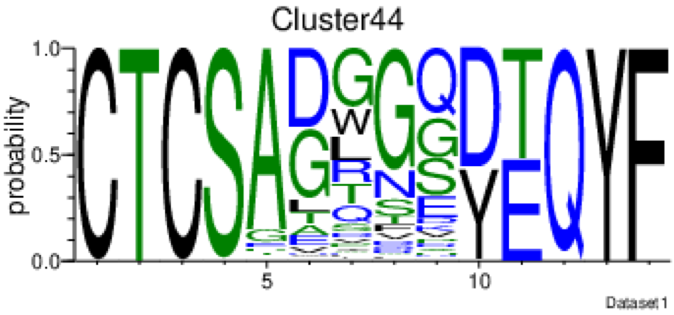
Cluster44
probability
1.0
0.5
0.0
5
10
Dataset1

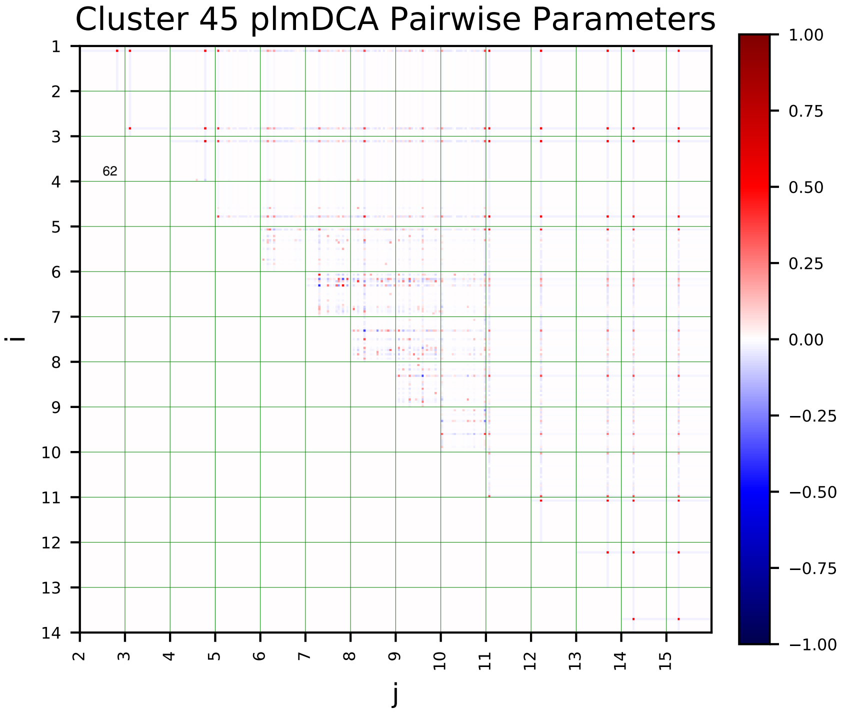
Cluster 45 plmDCA Pairwise Parameters
i
j
62

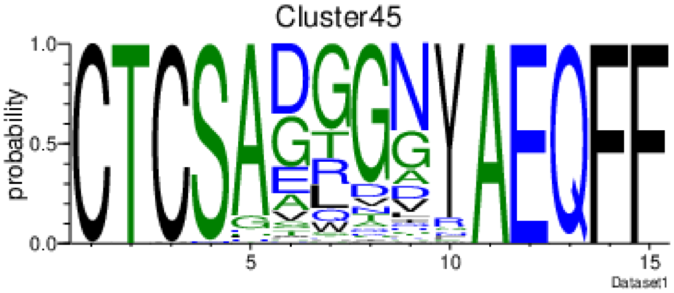
Cluster45
probability
Dataset1

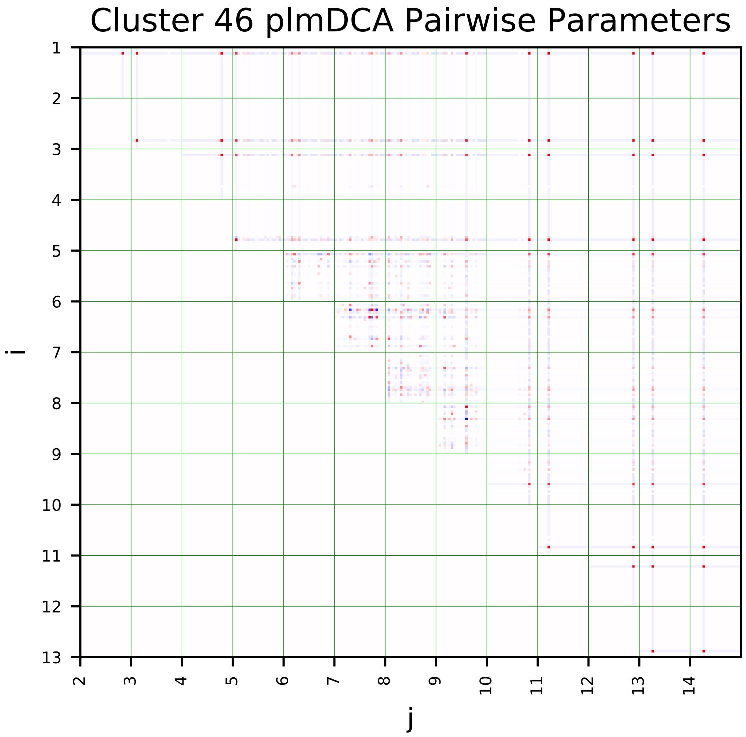
Cluster 46 plmDCA Pairwise Parameters
i
j

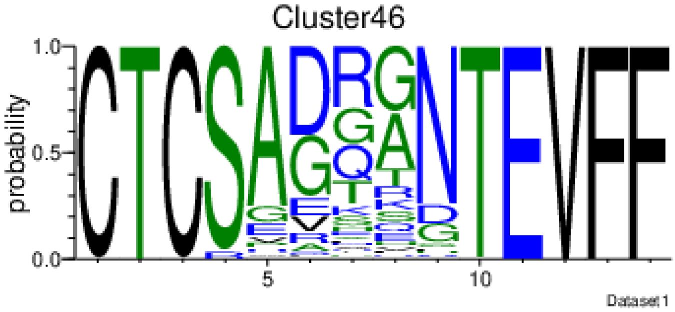
Cluster46
probability
Dataset1

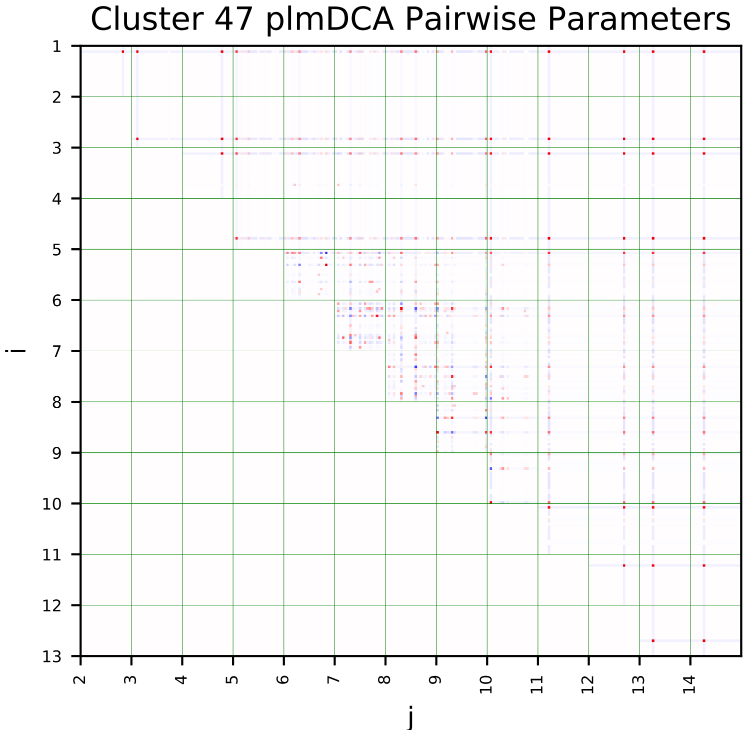
Cluster 47 plmDCA Pairwise Parameters
i
j

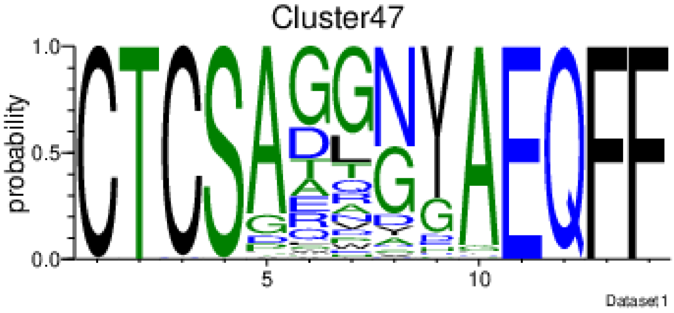
Cluster47
probability
Dataset1

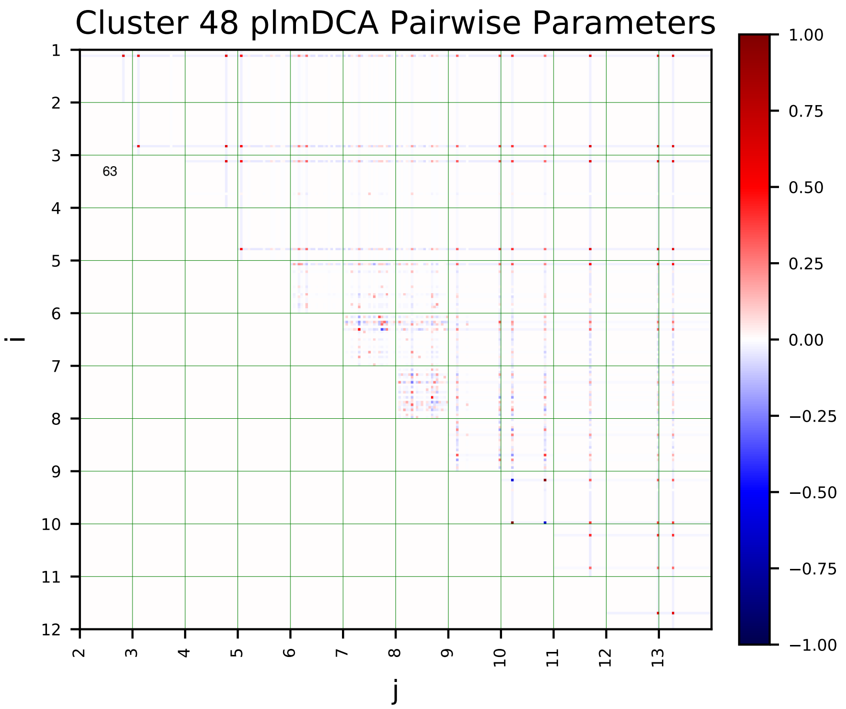
Cluster 48 plmDCA Pairwise Parameters
i
j
63

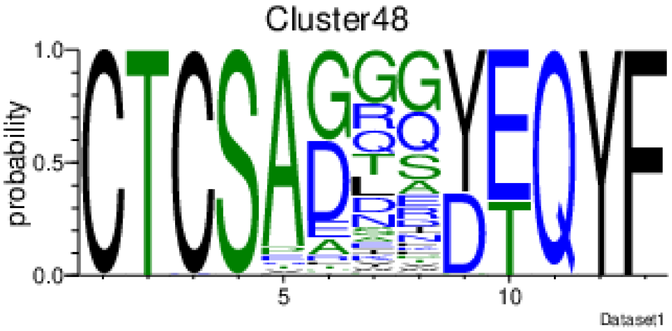
Cluster48
probability
Dataset1

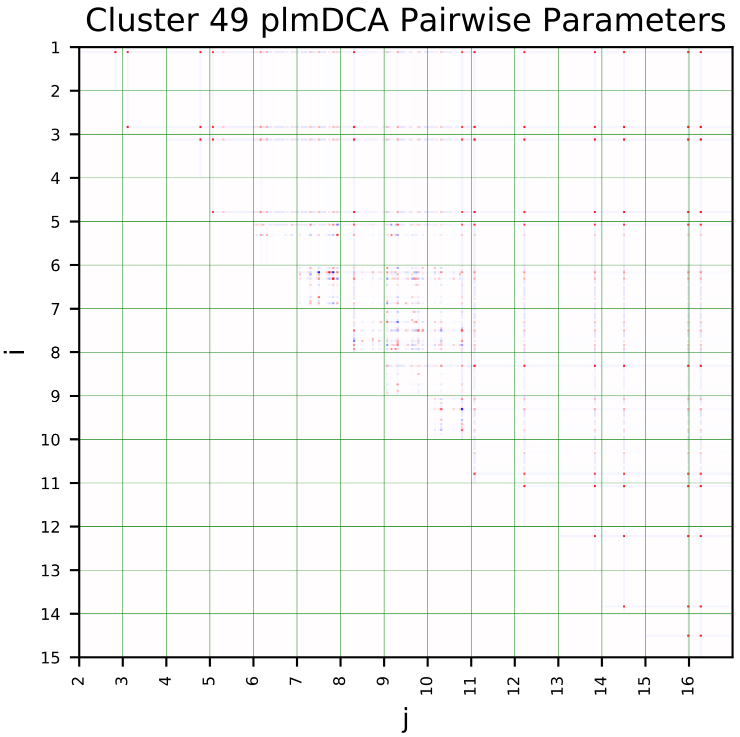
Cluster 49 plmDCA Pairwise Parameters
i
j

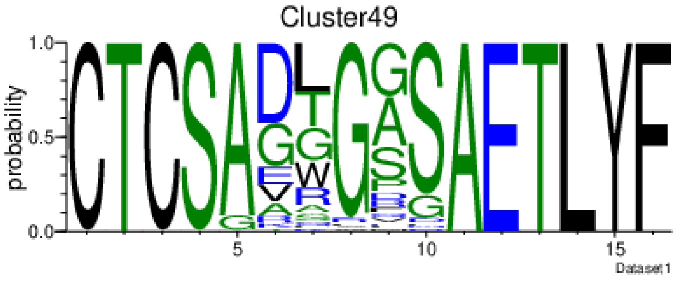
Cluster49
probability
Dataset1

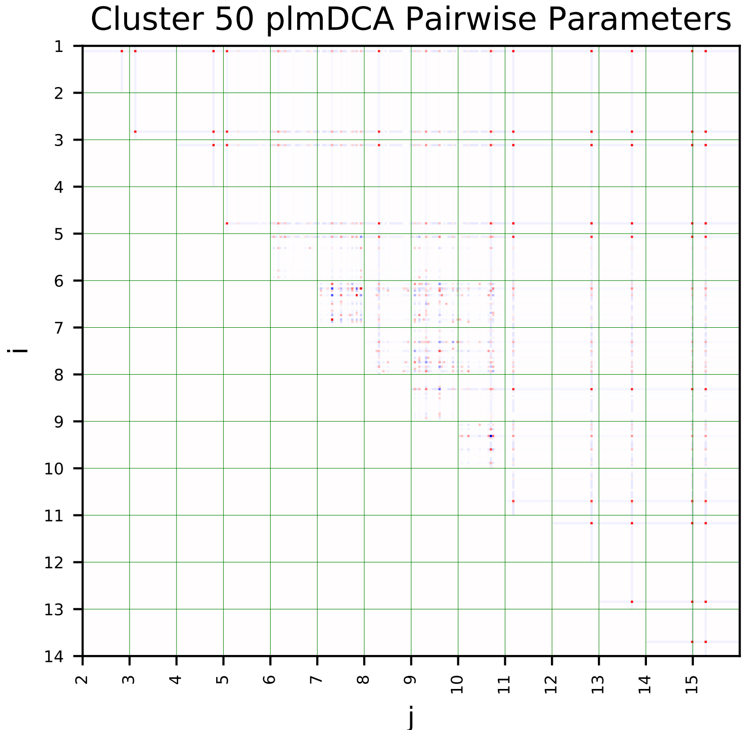
Cluster 50 plmDCA Pairwise Parameters
i
j

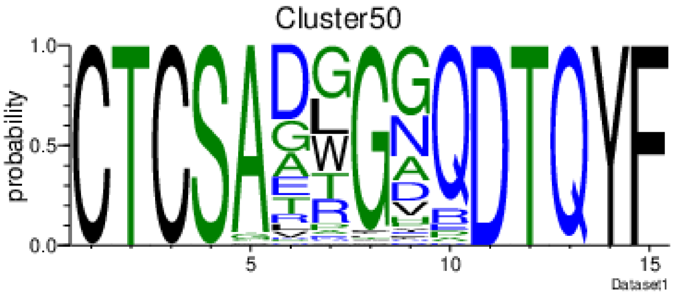
Cluster50
probability
Dataset1

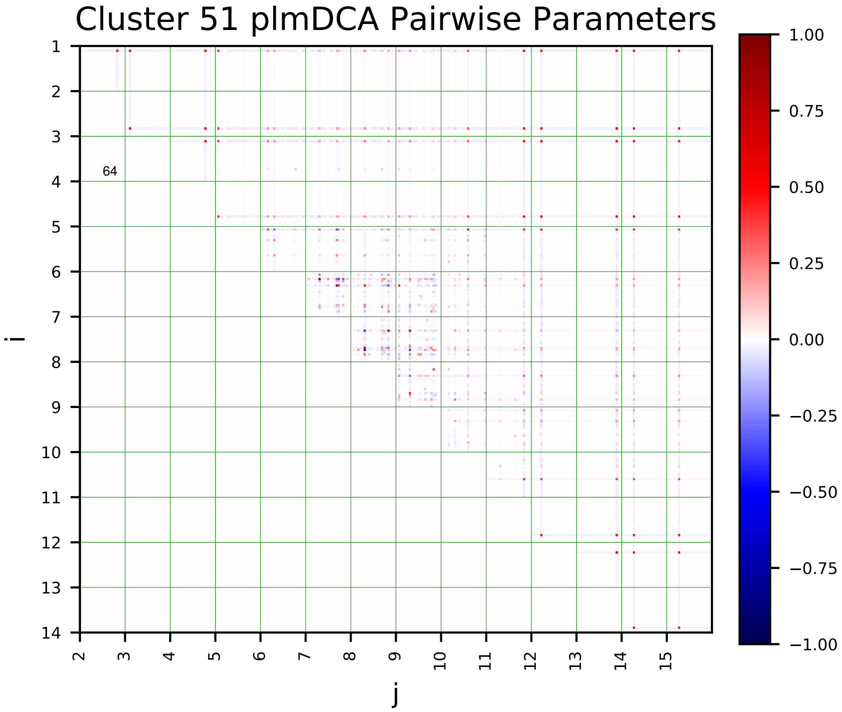
Cluster 51 plmDCA Pairwise Parameters
i
j
64

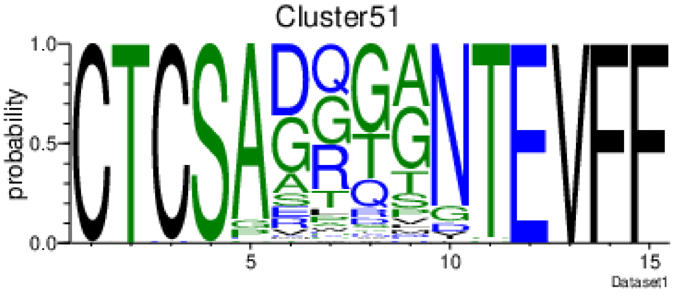
Cluster51
probability
Dataset1

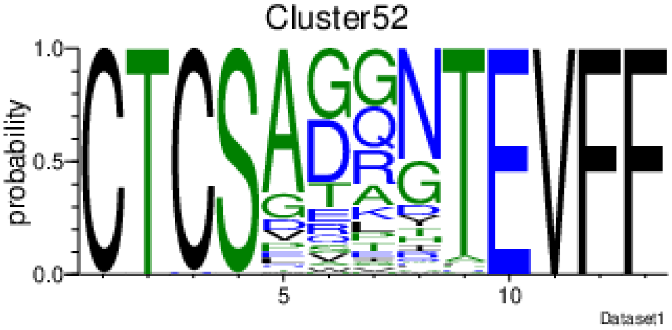
Cluster52
probability
Dataset1

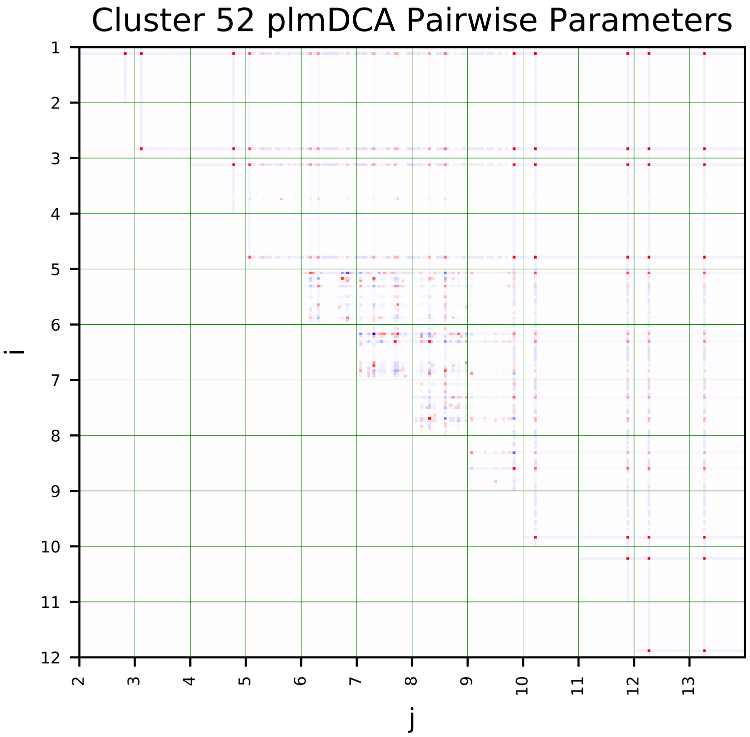
Cluster 52 plmDCA Pairwise Parameters
i
j

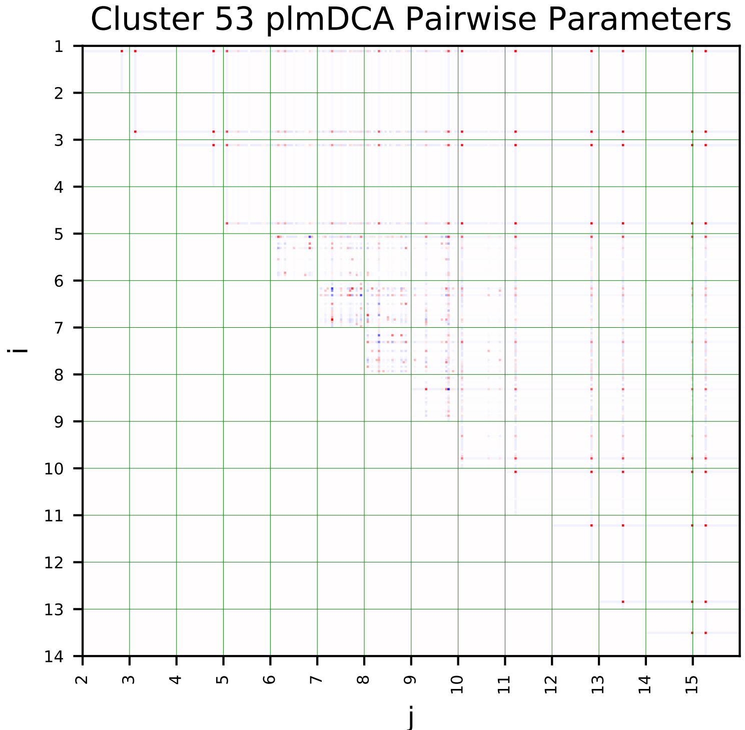
Cluster 53 plmDCA Pairwise Parameters
i
j

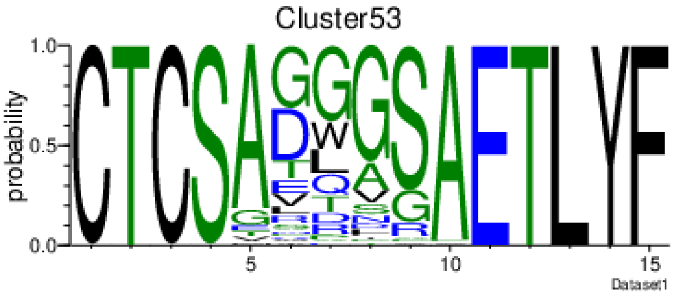
Cluster53
probability
Dataset1

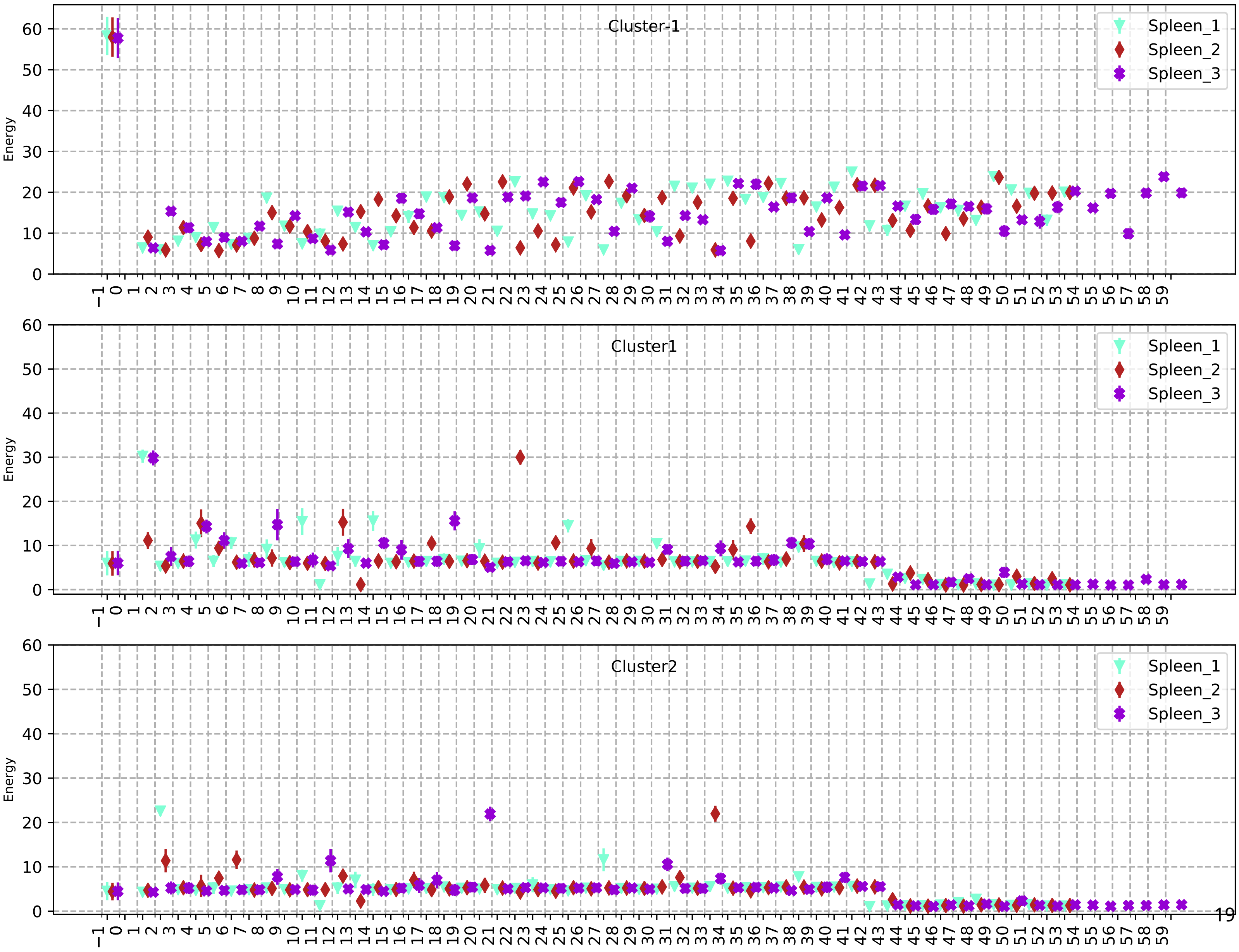

Cluster-1
Cluster1
Cluster2
Spleen_1
Spleen_2
Spleen_3
Energy

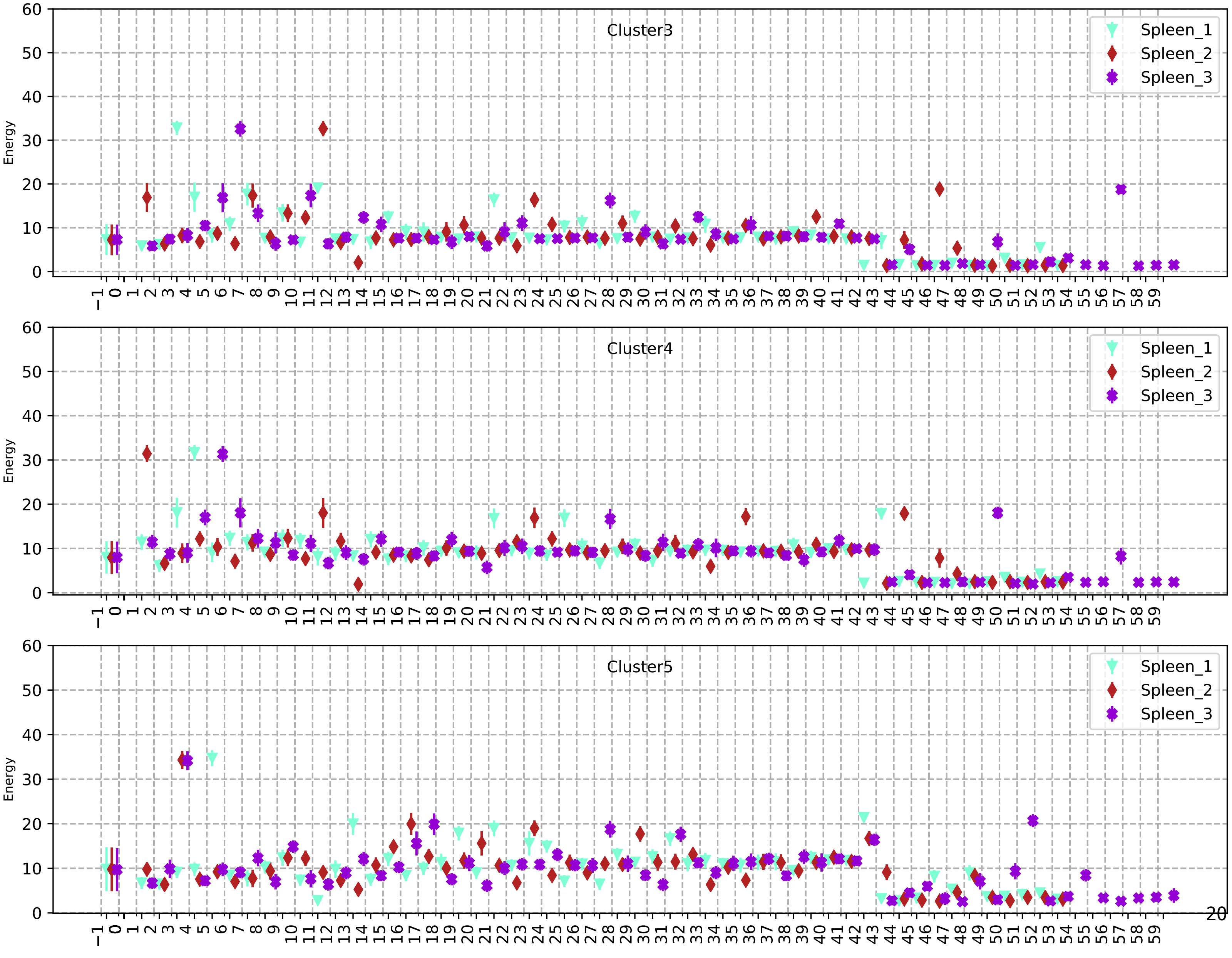
Cluster3
Spleen_1
Spleen_2
Spleen_3
Energy
Cluster4
Spleen_1
Spleen_2
Spleen_3
Energy
Cluster5
Spleen_1
Spleen_2
Spleen_3
Energy

67

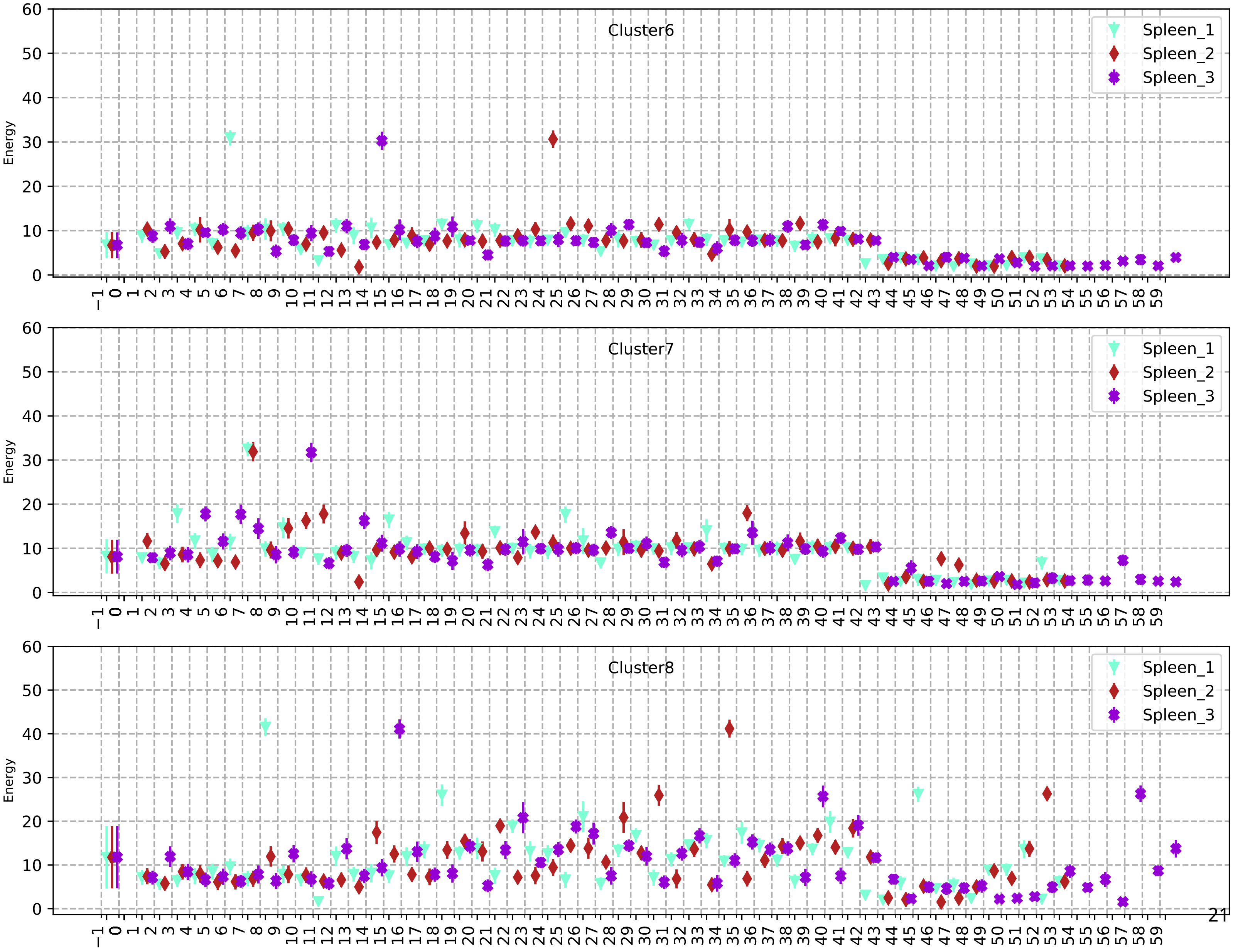

Cluster6
Cluster7
Cluster8
Spleen_1
Spleen_2
Spleen_3
Energy

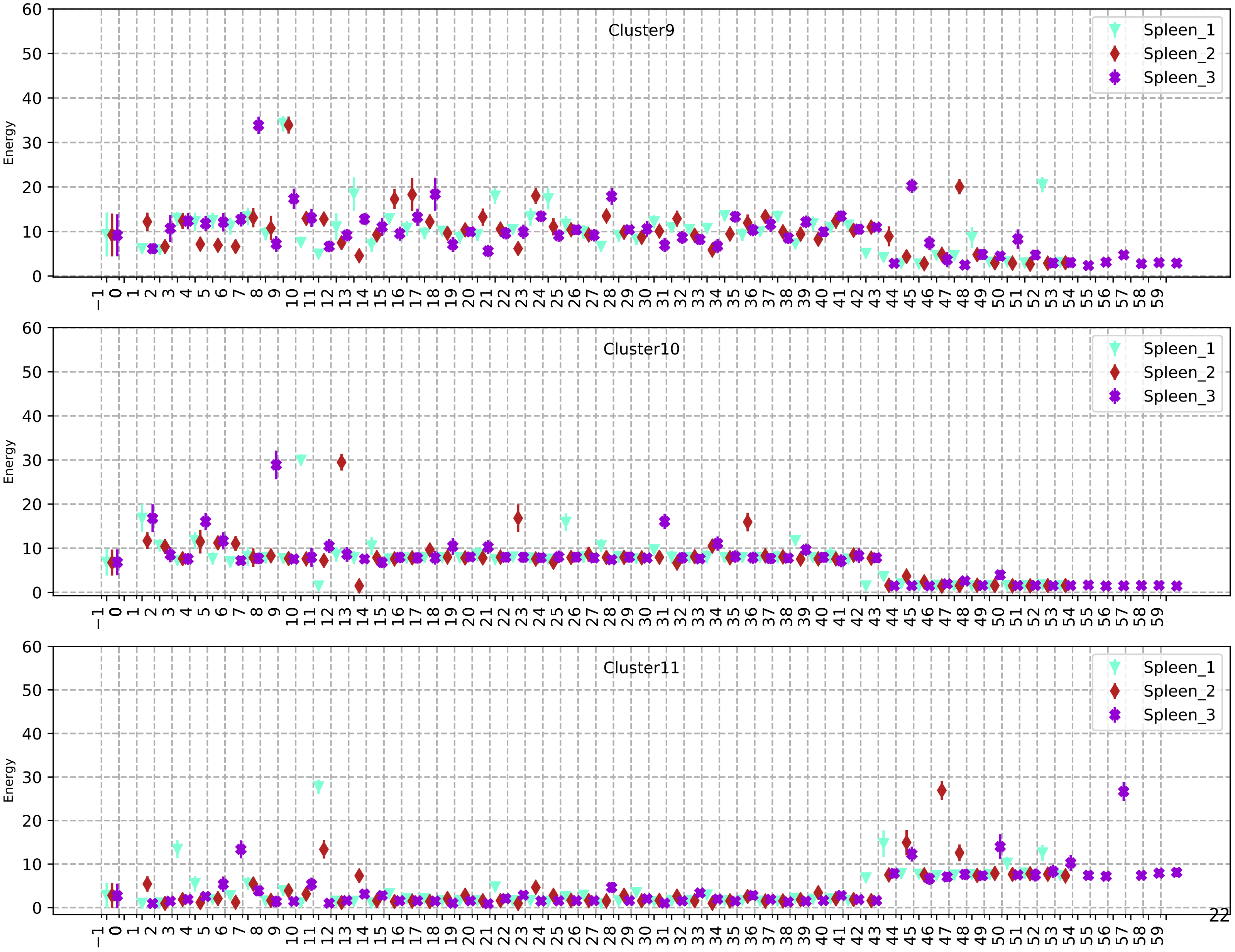
Cluster9
Cluster10
Cluster11
Spleen_1
Spleen_2
Spleen_3
Energy

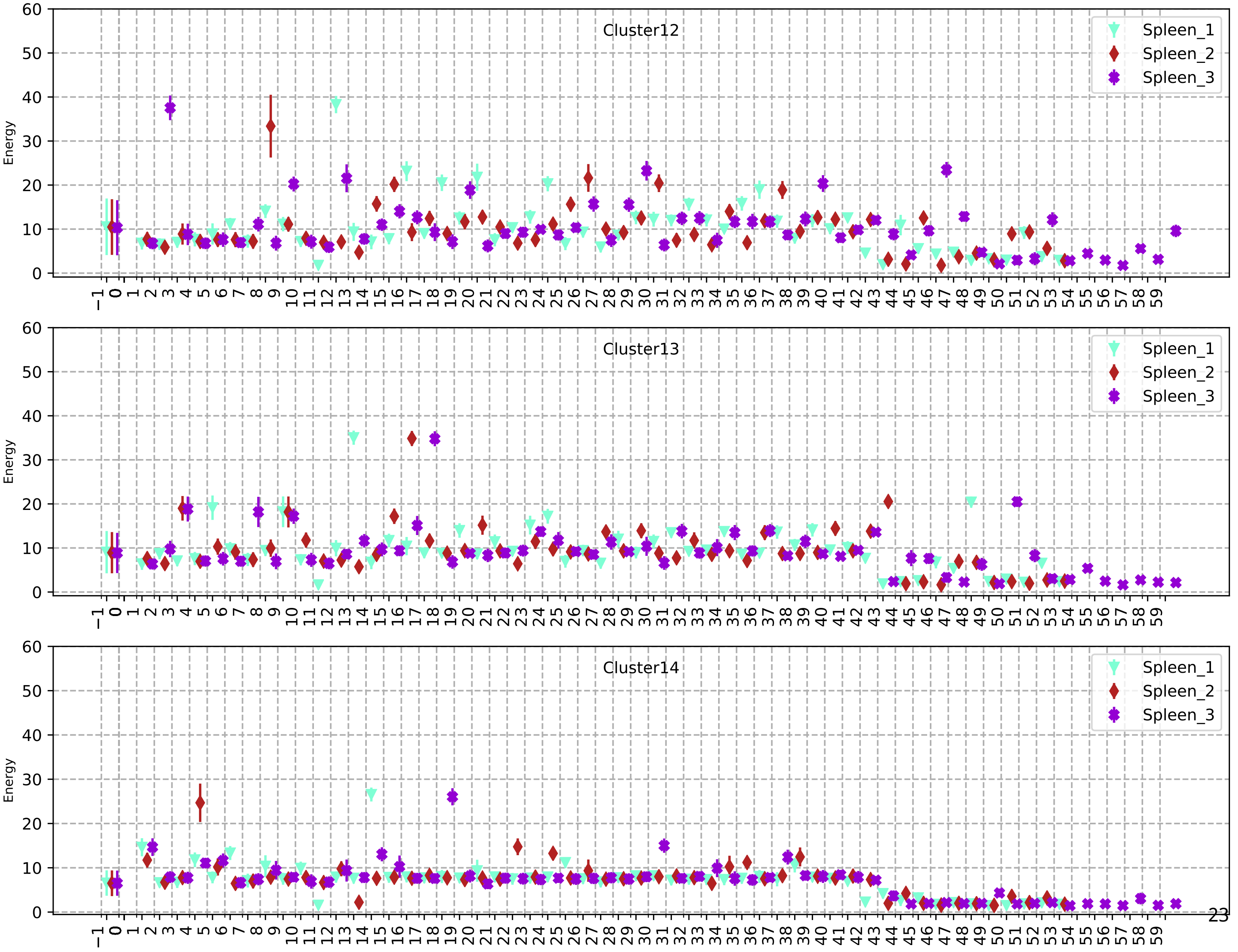
Cluster12
Spleen_1
Spleen_2
Spleen_3
Energy
Cluster13
Spleen_1
Spleen_2
Spleen_3
Energy
Cluster14
Spleen_1
Spleen_2
Spleen_3
Energy

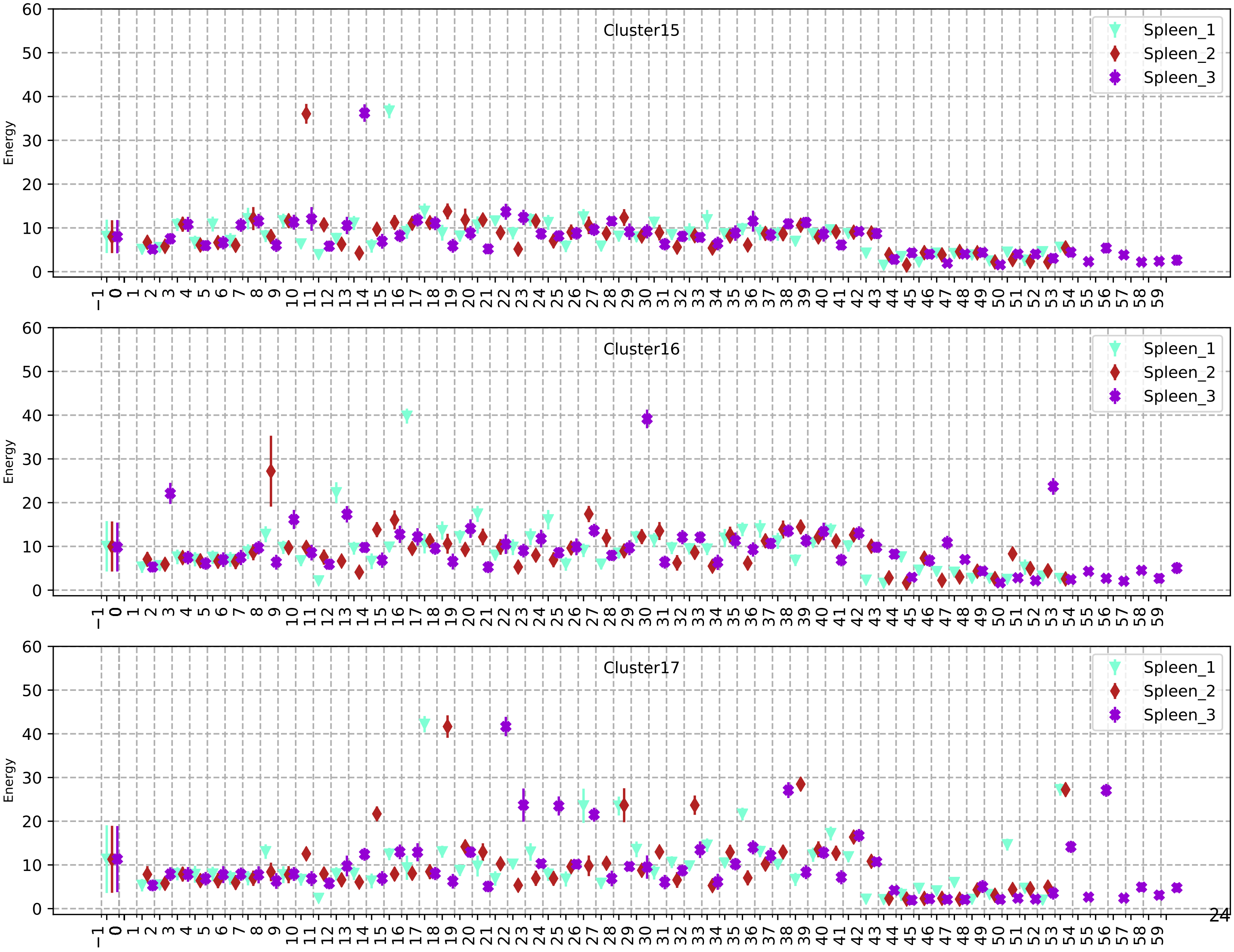
Cluster15
Spleen_1
Spleen_2
Spleen_3
Energy
Cluster16
Spleen_1
Spleen_2
Spleen_3
Energy
Cluster17
Spleen_1
Spleen_2
Spleen_3
Energy

71

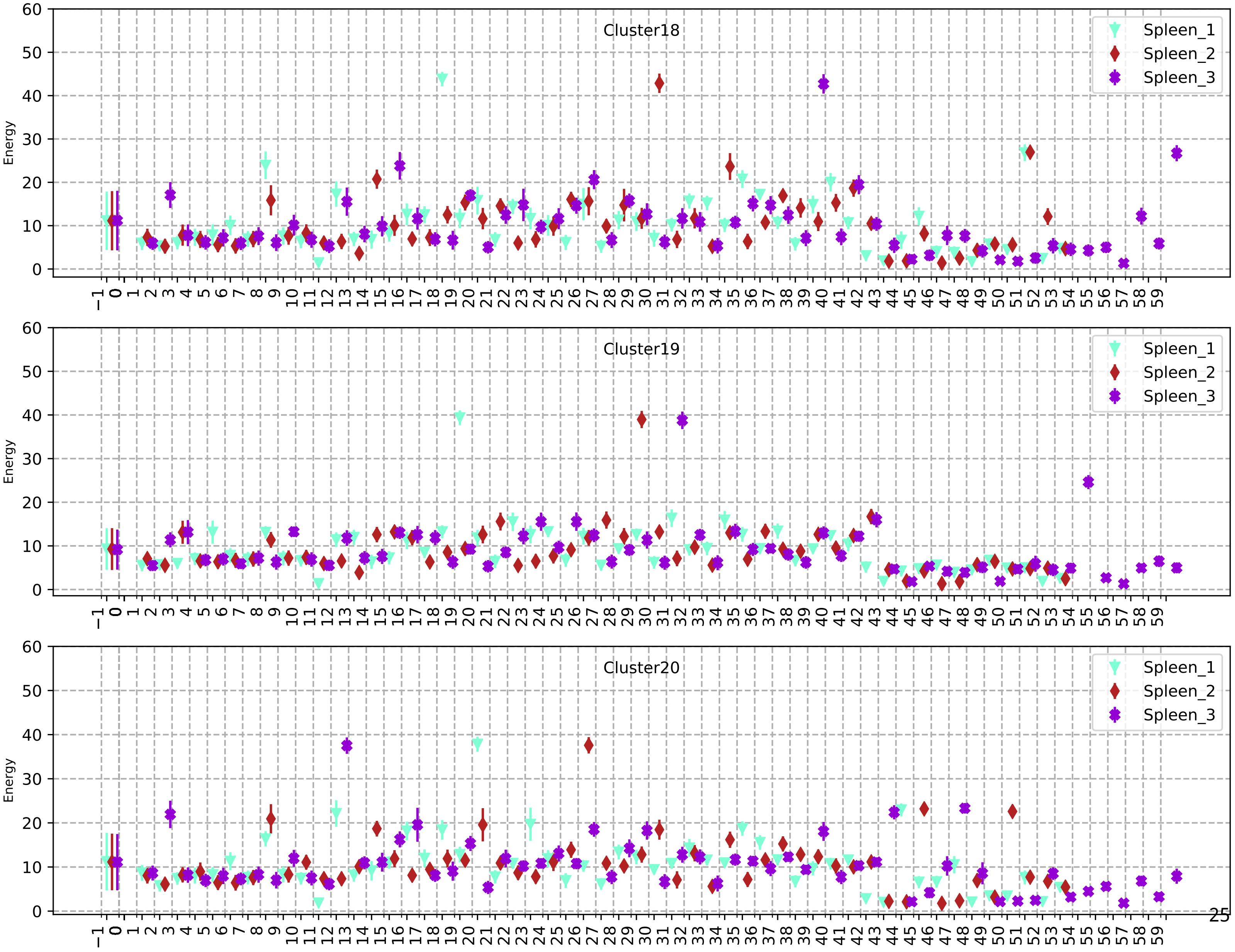
Cluster18
Cluster19
Cluster20
Spleen_1
Spleen_2
Spleen_3
Energy

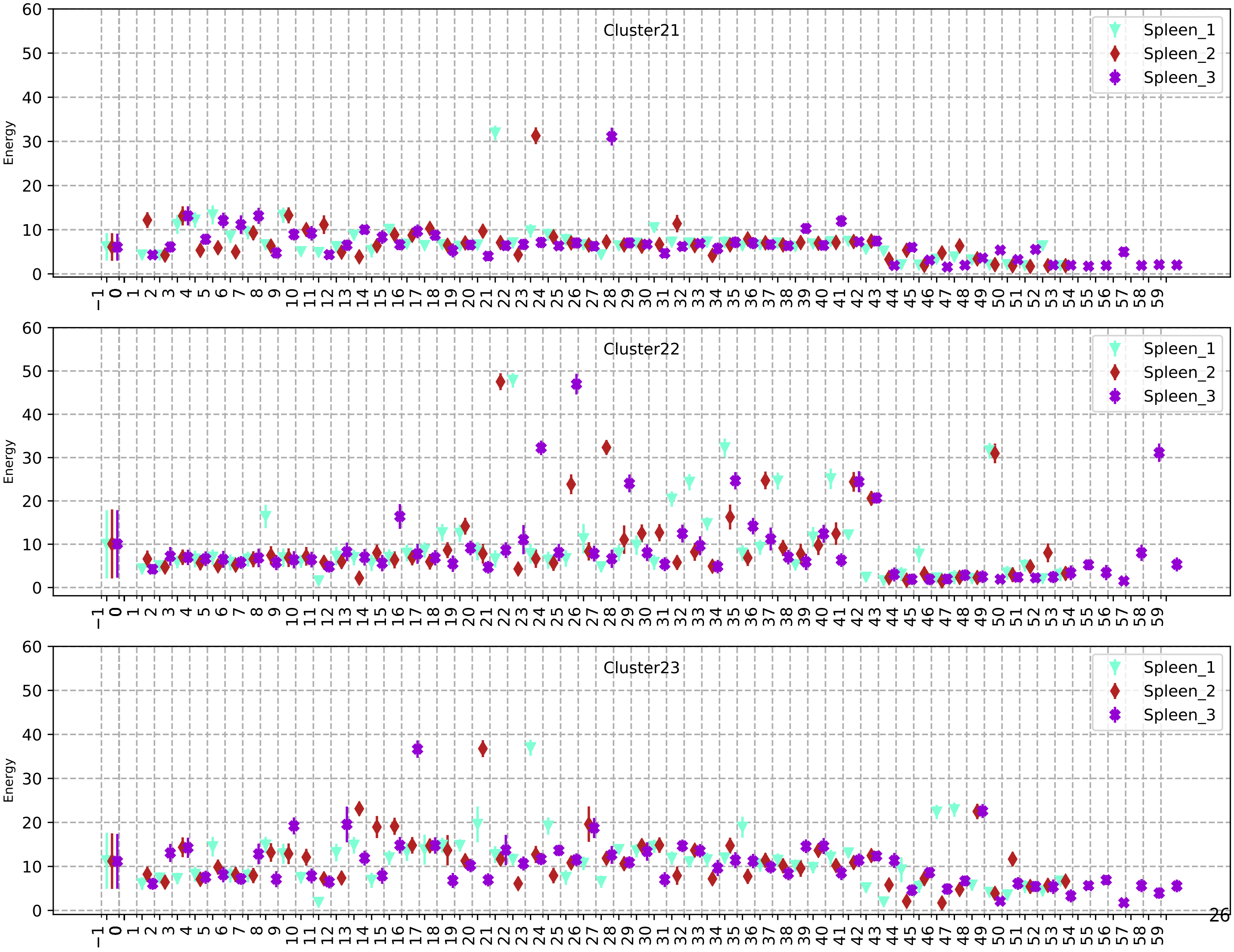
Cluster21
Cluster22
Cluster23
Spleen_1
Spleen_2
Spleen_3
Energy

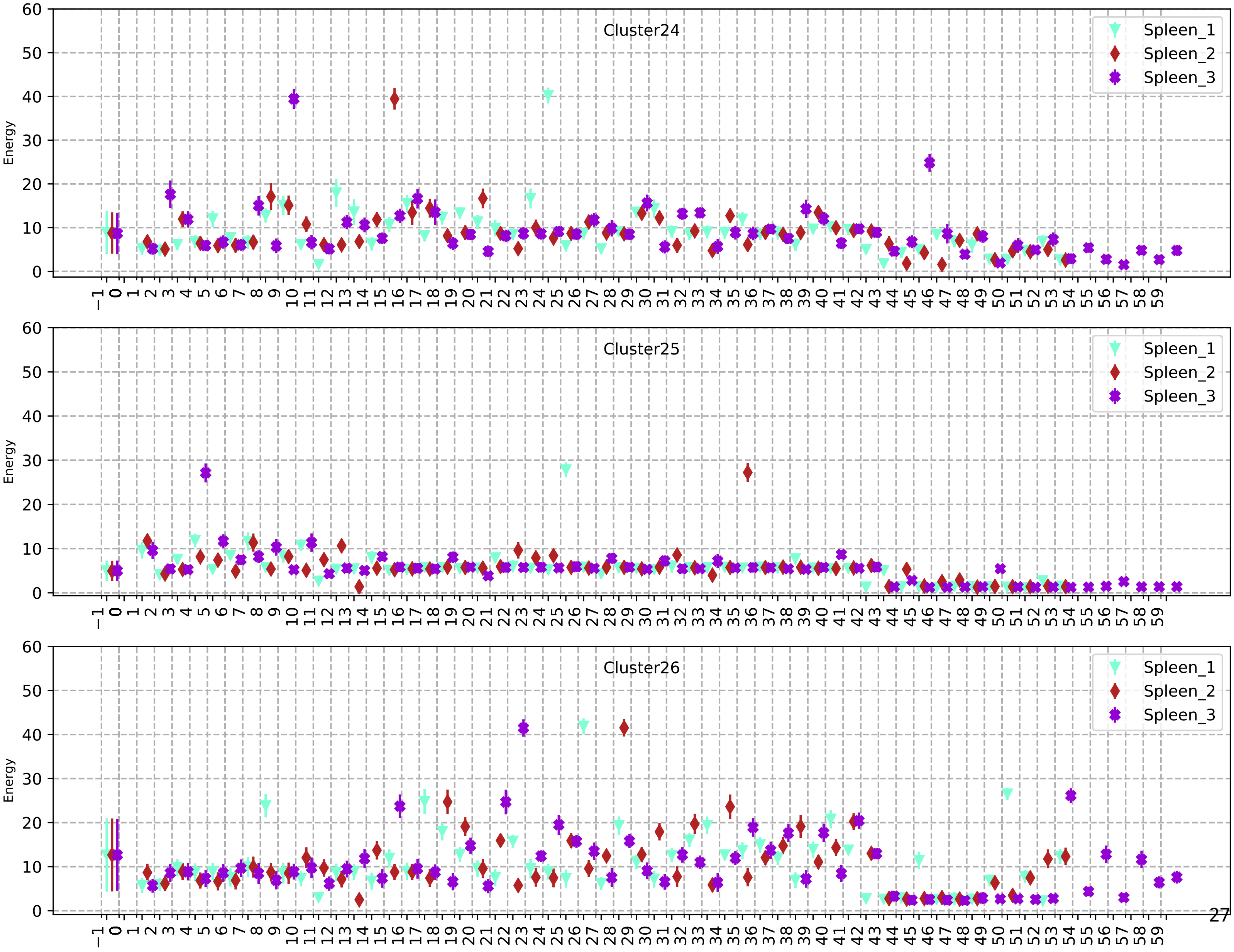
Cluster24
Cluster25
Cluster26
Spleen_1
Spleen_2
Spleen_3
Energy

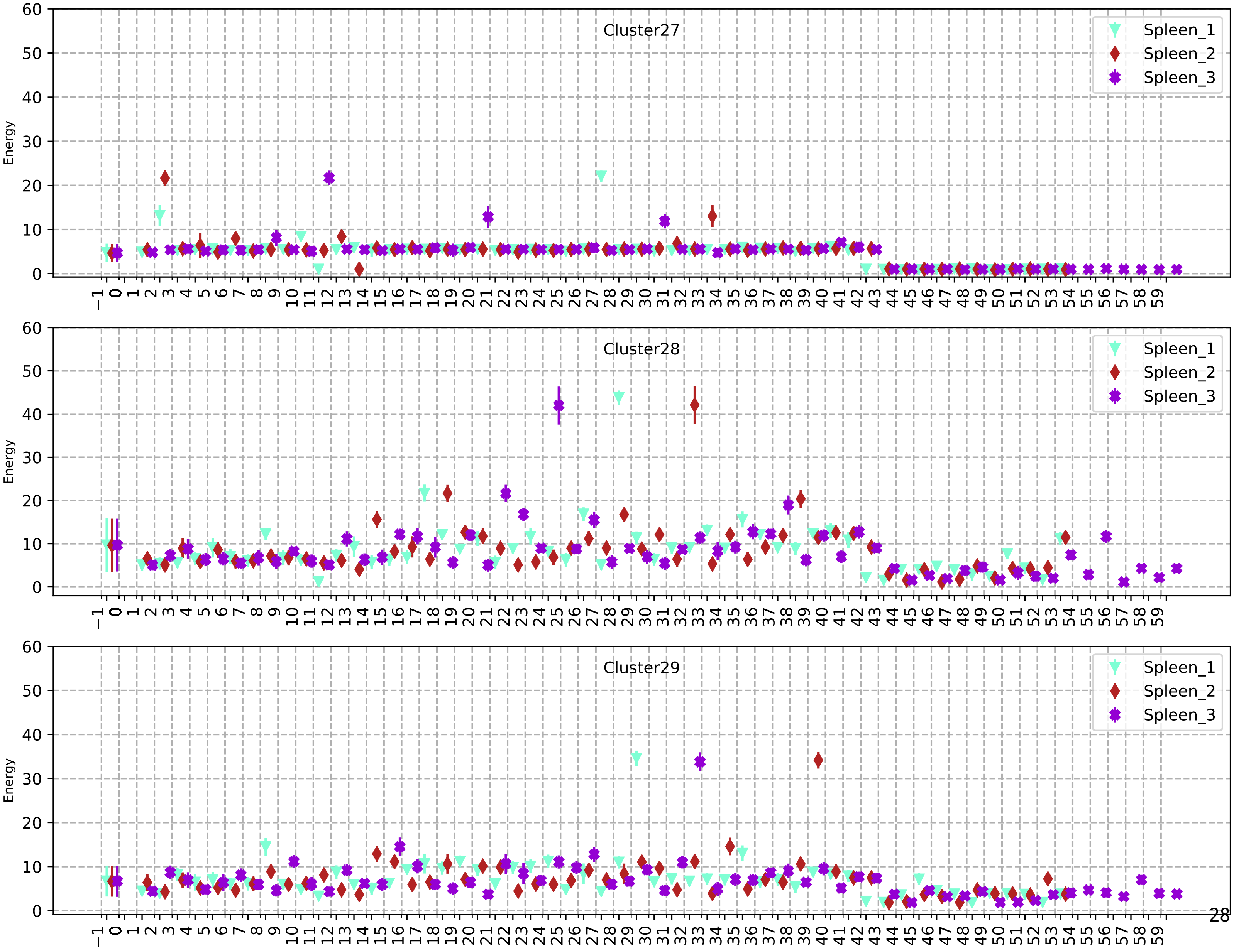
Cluster27
Cluster28
Cluster29
Spleen_1
Spleen_2
Spleen_3
Energy

75

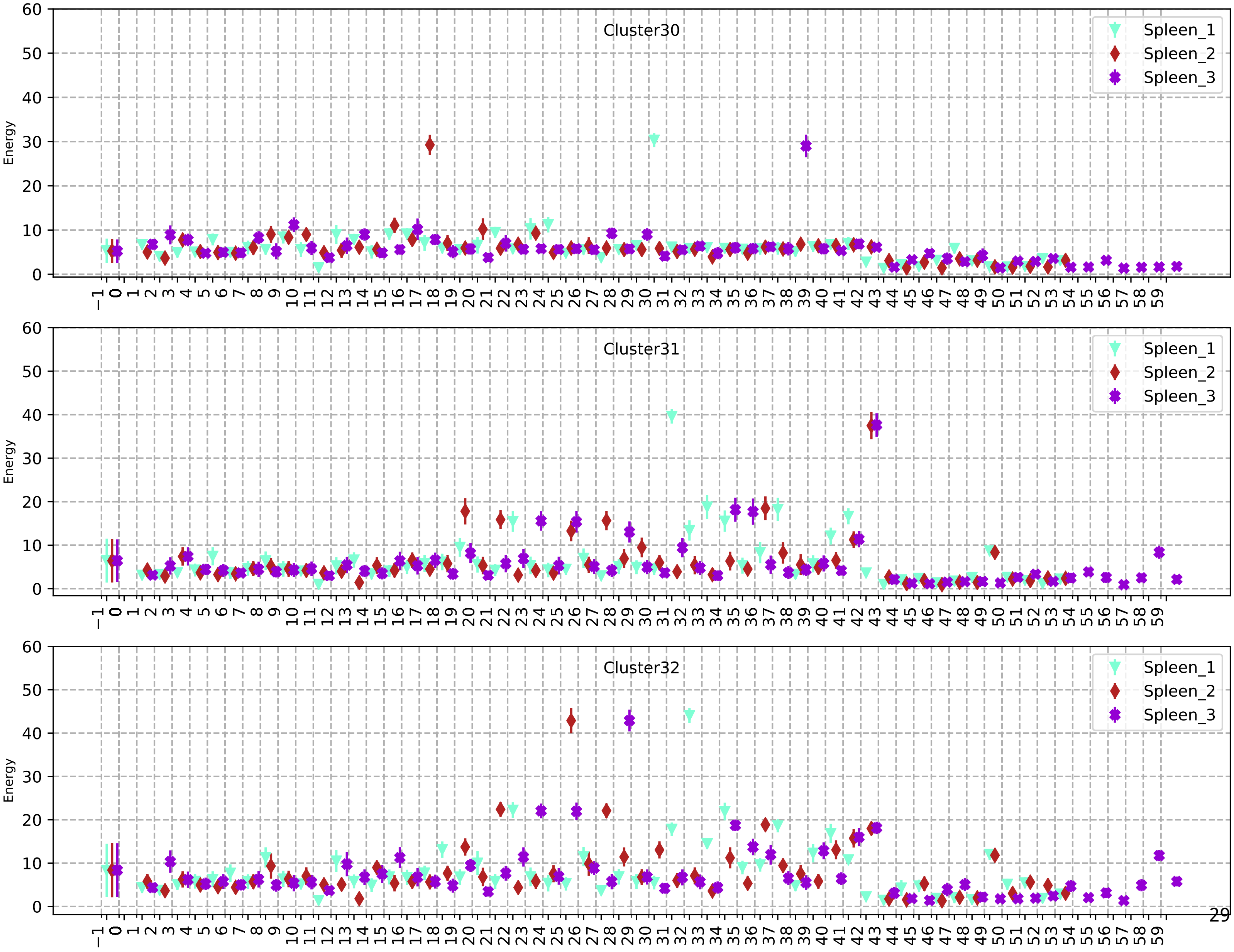

Cluster30
Cluster31
Cluster32
Spleen_1
Spleen_2
Spleen_3
Energy

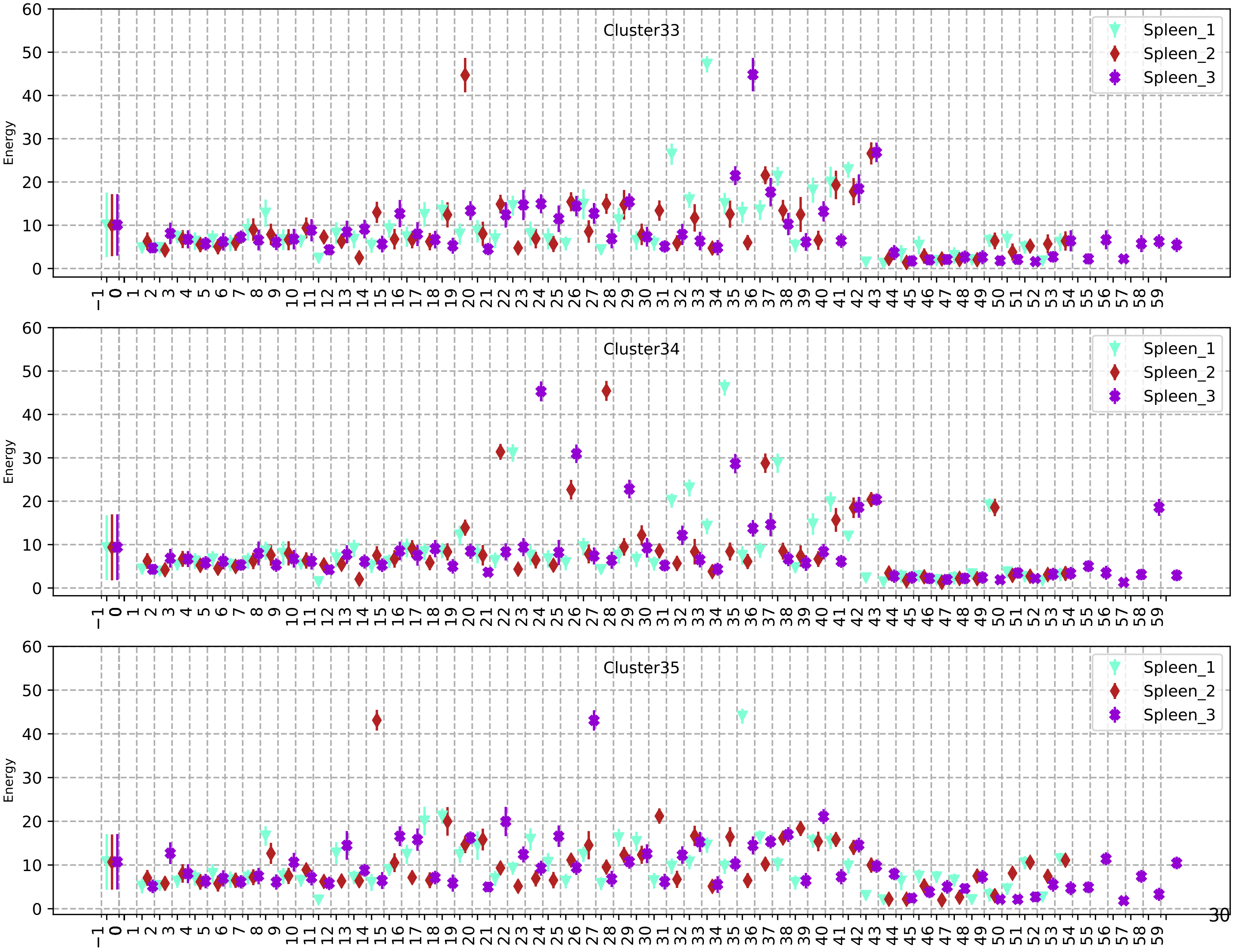
Cluster33
Cluster34
Cluster35
Spleen_1
Spleen_2
Spleen_3
Energy

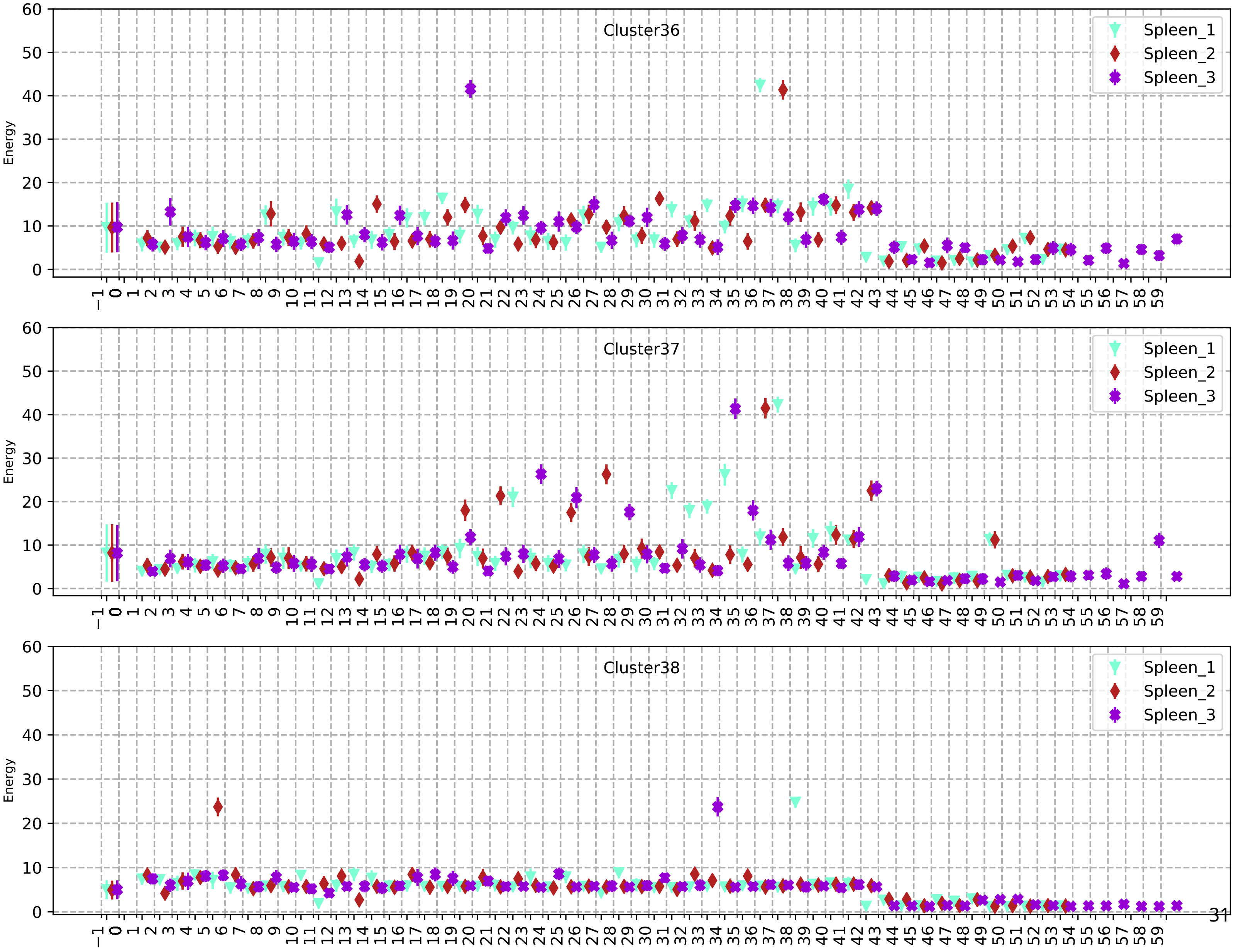
Cluster36
Cluster37
Cluster38
Spleen_1
Spleen_2
Spleen_3
Energy

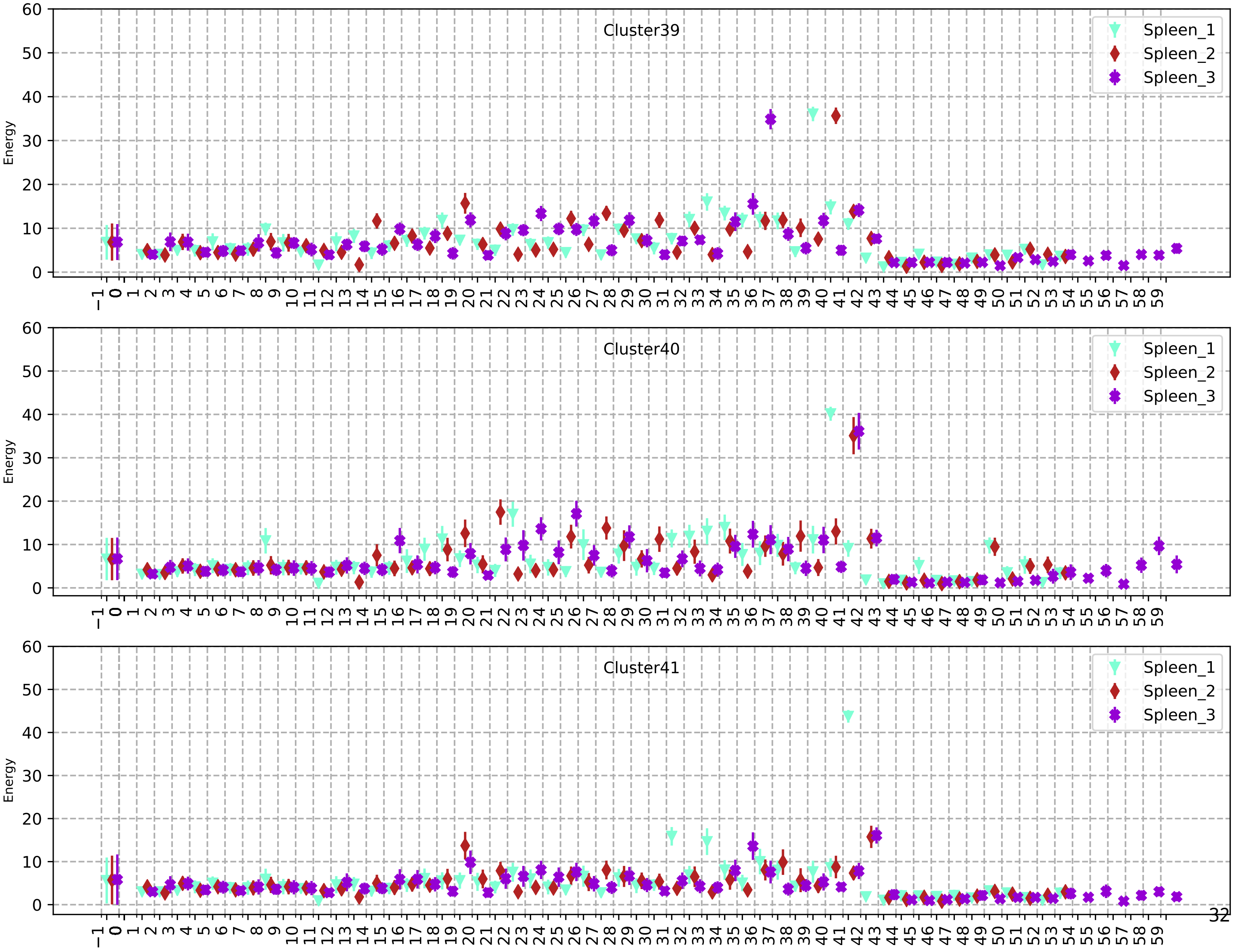
Cluster39
Cluster40
Cluster41
Spleen_1
Spleen_2
Spleen_3
Energy
0
10
20
30
40
50
60
−1 0 1 2 3 4 5 6 7 8 9 10 11 12 13 14 15 16 17 18 19 20 21 22 23 24 25 26 27 28 29 30 31 32 33 34 35 36 37 38 39 40 41 42 43 44 45 46 47 48 49 50 51 52 53 54 55 56 57 58 59

79

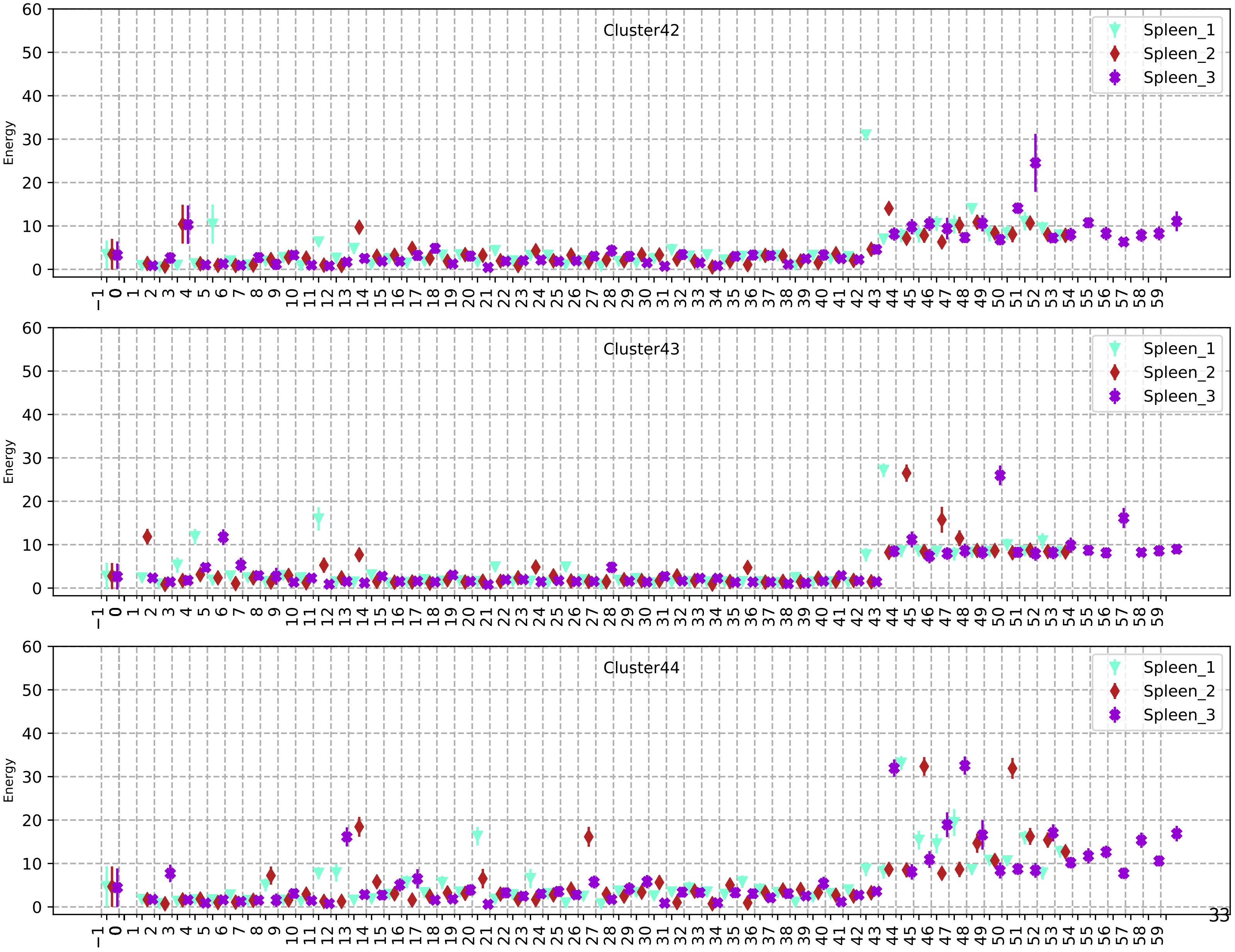

Cluster42
Cluster43
Cluster44
Spleen_1
Spleen_2
Spleen_3
Energy


33

80

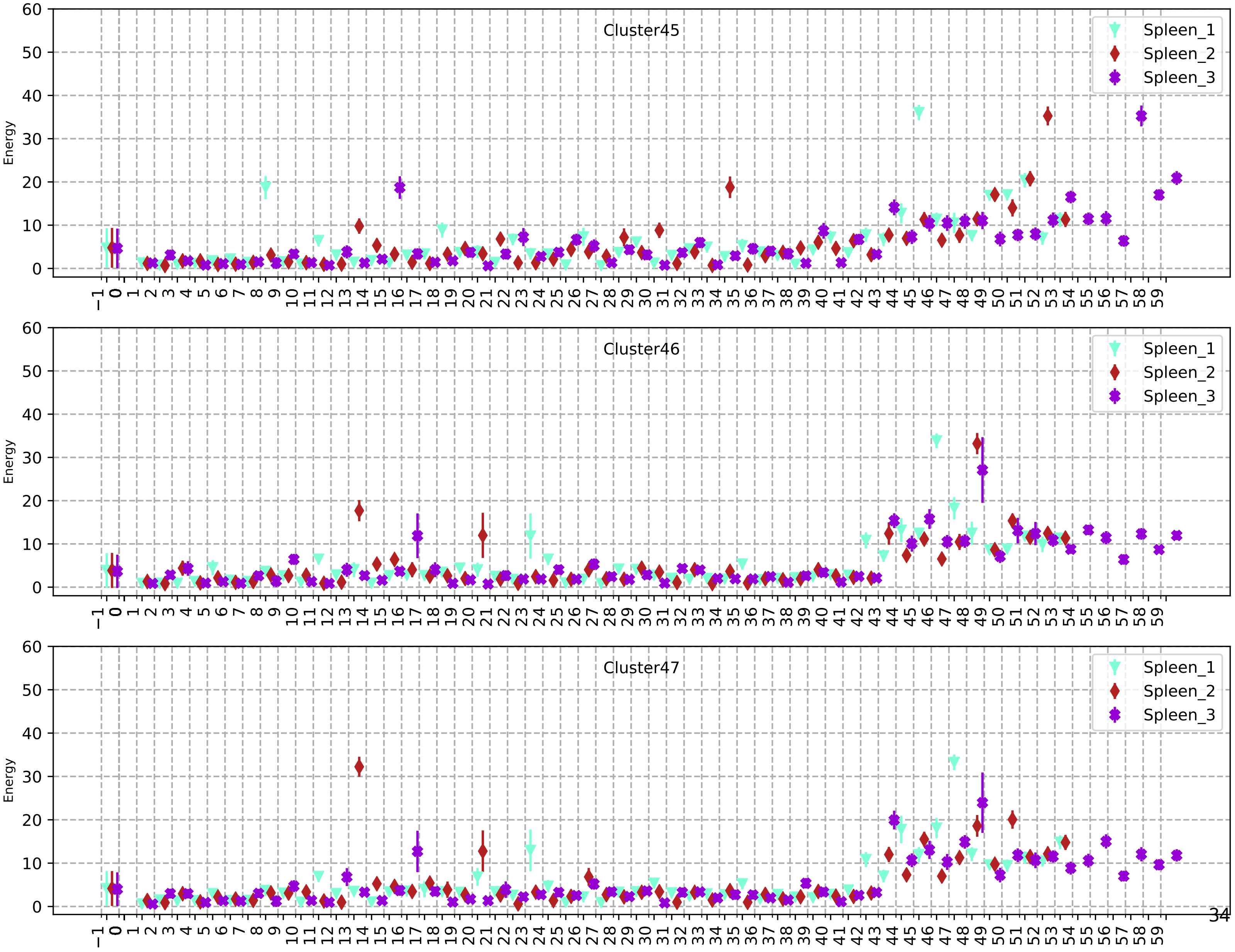

Cluster45
Spleen_1
Spleen_2
Spleen_3
Energy
Cluster46
Spleen_1
Spleen_2
Spleen_3
Energy
Cluster47
Spleen_1
Spleen_2
Spleen_3
Energy

81

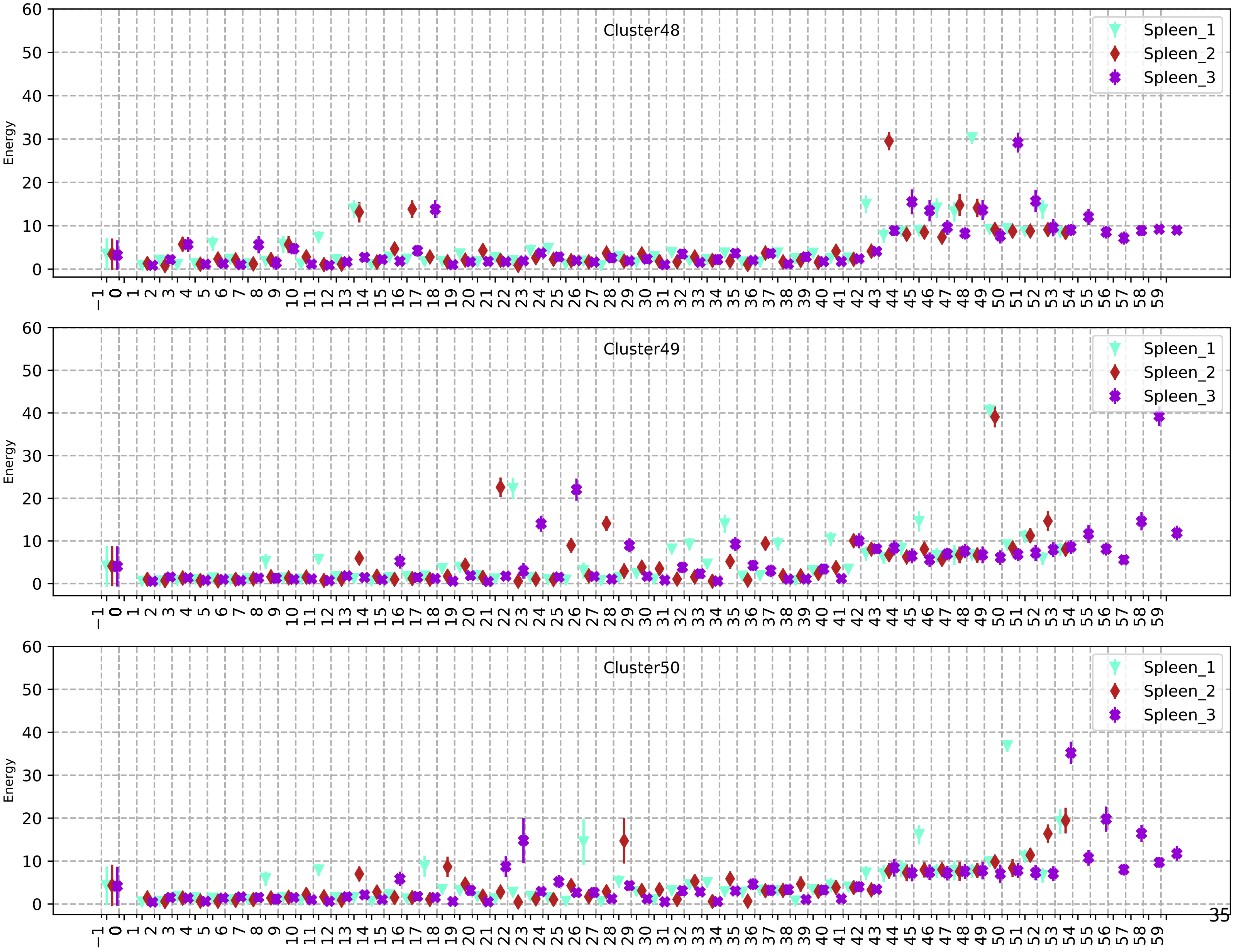

Cluster48
Cluster49
Cluster50
Energy
Spleen_1
Spleen_2
Spleen_3

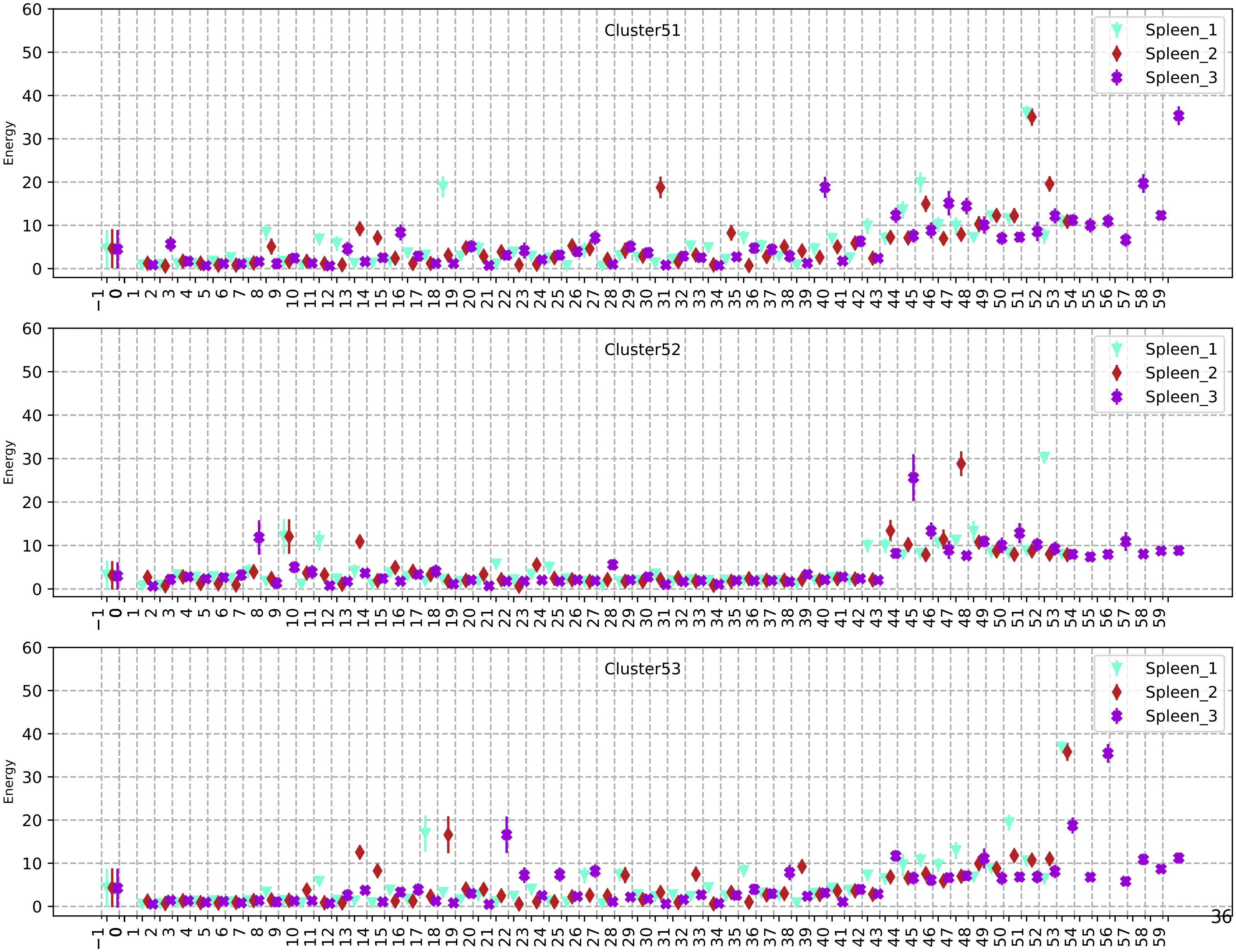
Cluster51
Cluster52
Cluster53
Spleen_1
Spleen_2
Spleen_3
Energy

83

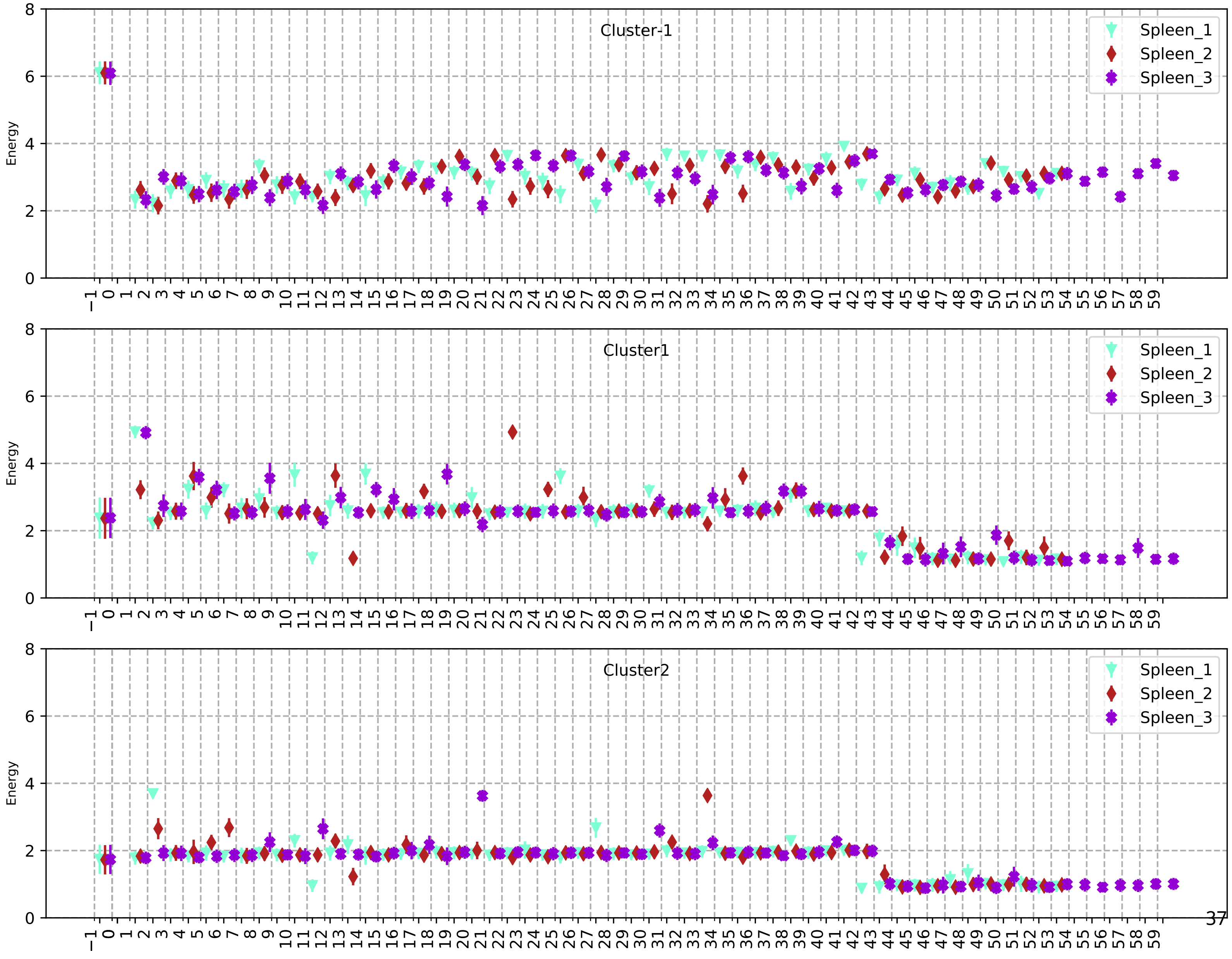

Cluster-1
Cluster1
Cluster2
Spleen_1
Spleen_2
Spleen_3
Energy

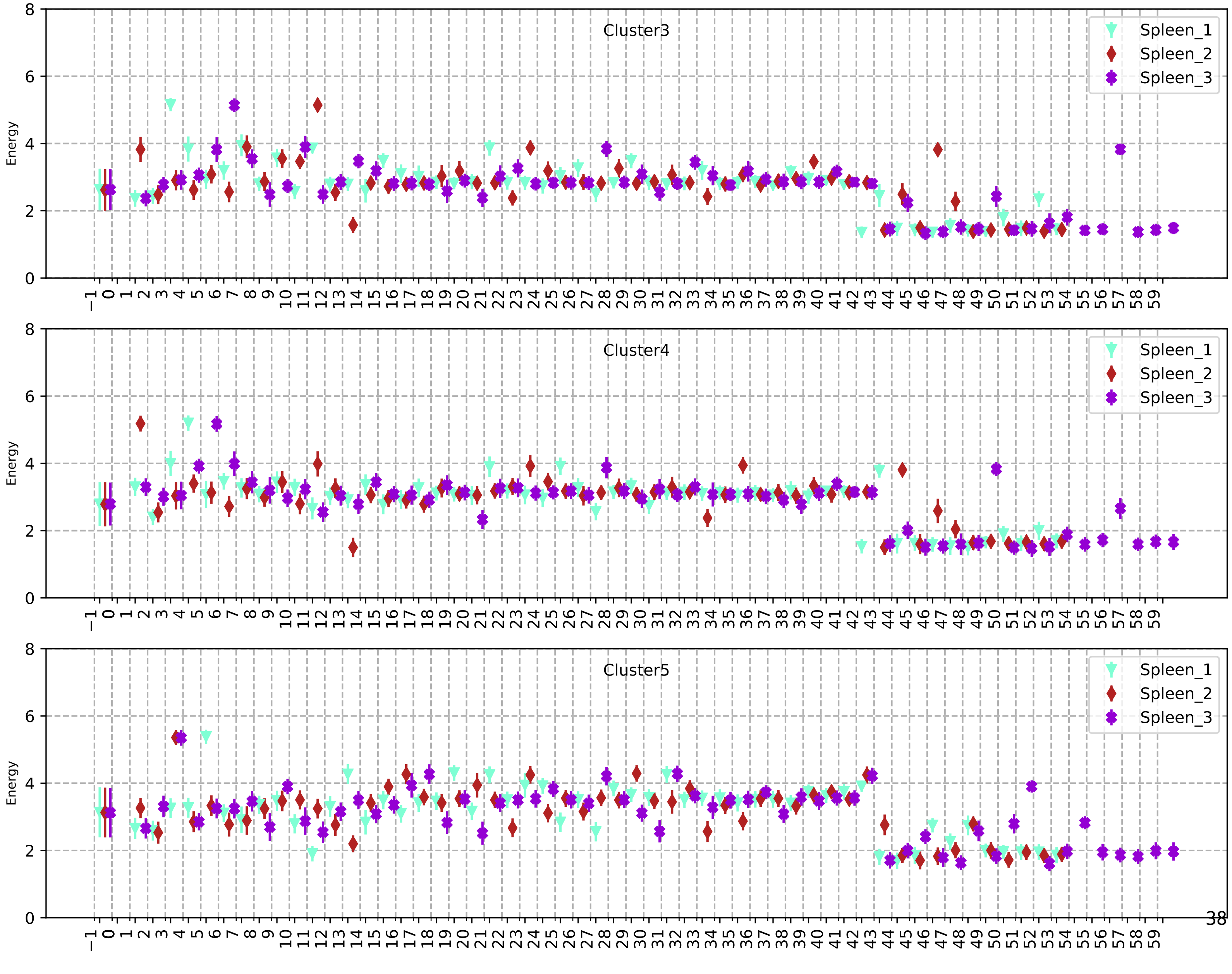
Cluster3
Cluster4
Cluster5
Spleen_1
Spleen_2
Spleen_3
Energy

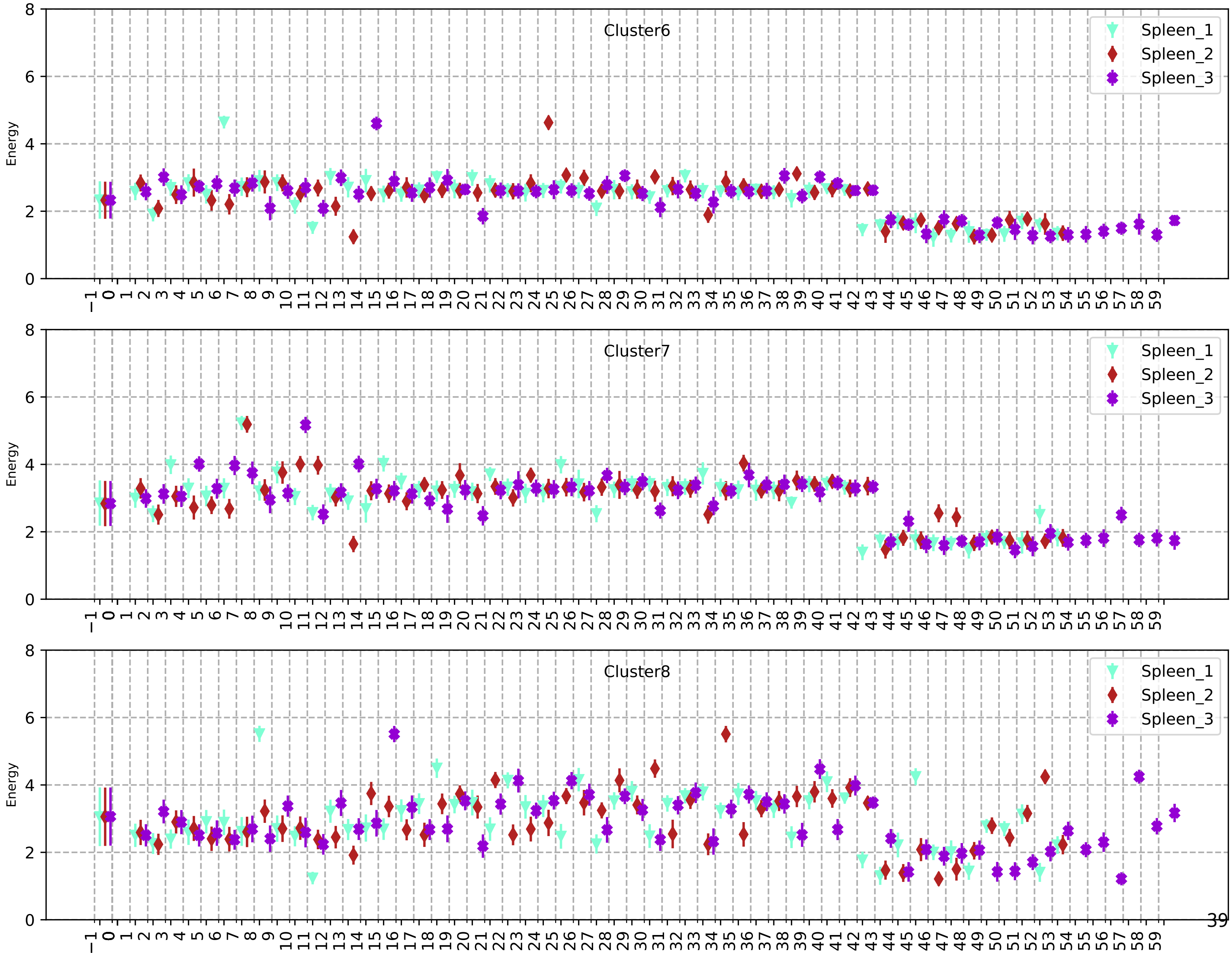
Cluster6
Cluster7
Cluster8
Spleen_1
Spleen_2
Spleen_3
Energy

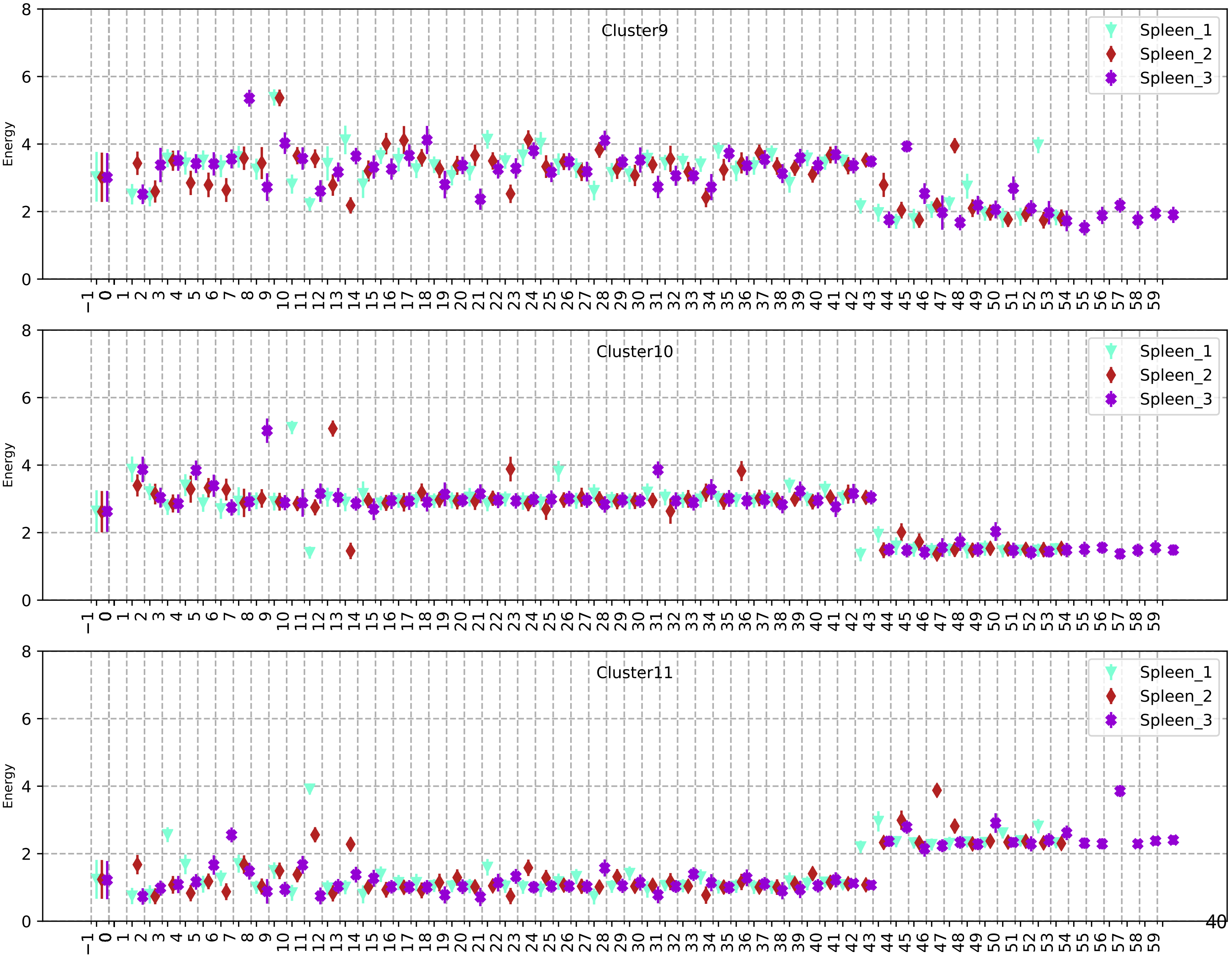
Cluster9
Cluster10
Cluster11
Spleen_1
Spleen_2
Spleen_3
Energy

87

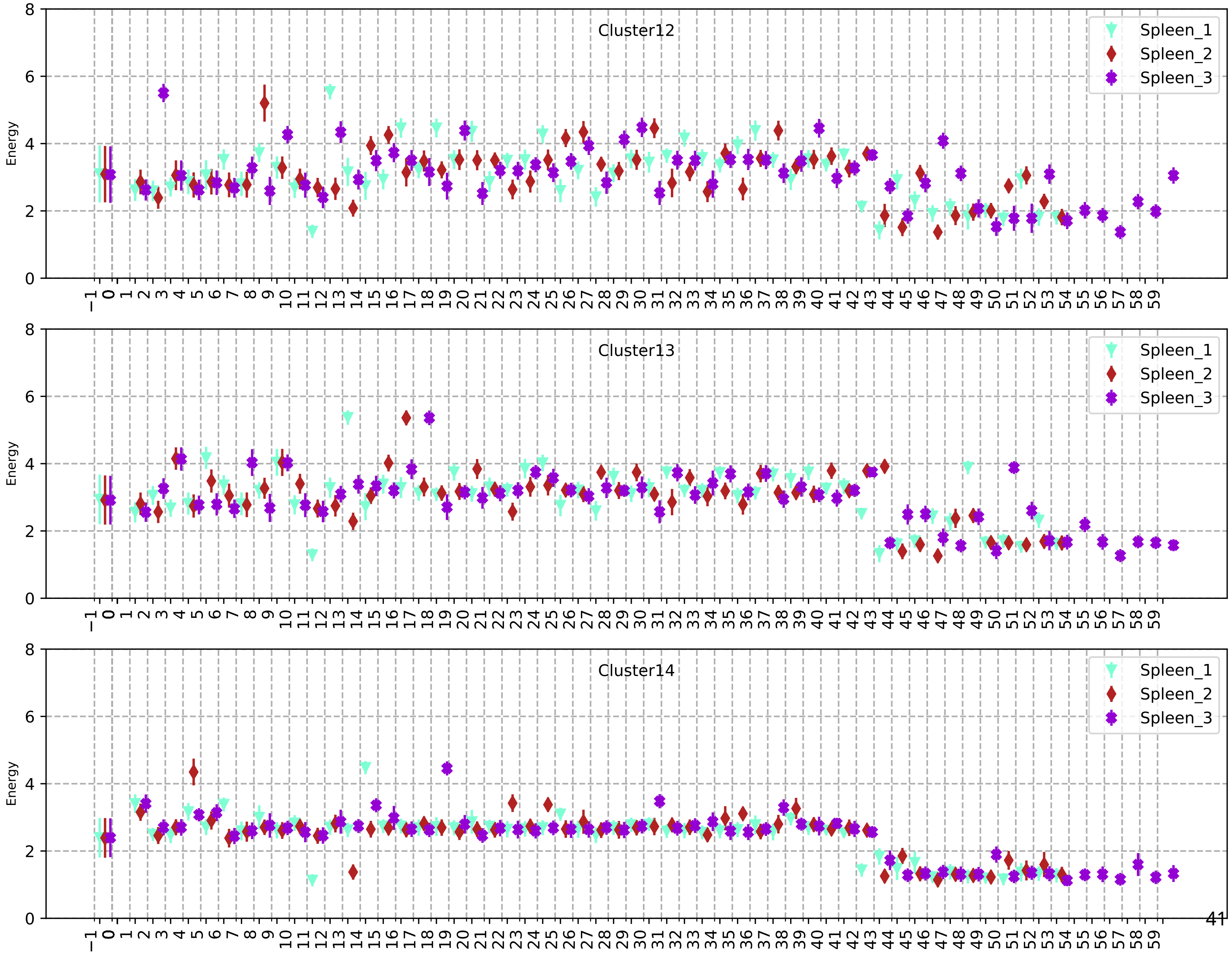
Cluster12
Cluster13
Cluster14
Spleen_1
Spleen_2
Spleen_3
Energy
0
2
4
6
8
−1 0 1 2 3 4 5 6 7 8 9 10 11 12 13 14 15 16 17 18 19 20 21 22 23 24 25 26 27 28 29 30 31 32 33 34 35 36 37 38 39 40 41 42 43 44 45 46 47 48 49 50 51 52 53 54 55 56 57 58 59

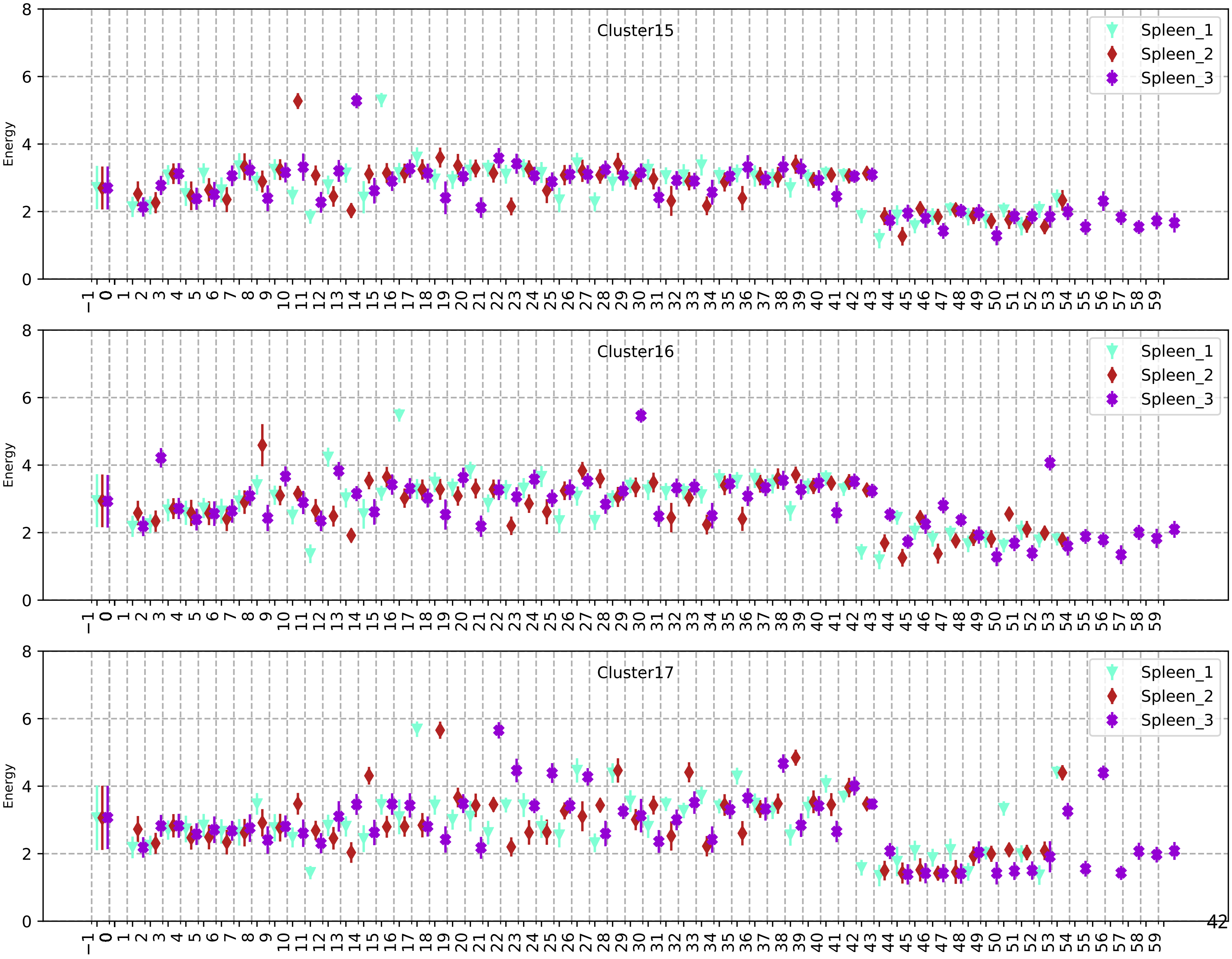
Cluster15
Spleen_1
Spleen_2
Spleen_3
Energy
Cluster16
Spleen_1
Spleen_2
Spleen_3
Energy
Cluster17
Spleen_1
Spleen_2
Spleen_3
Energy

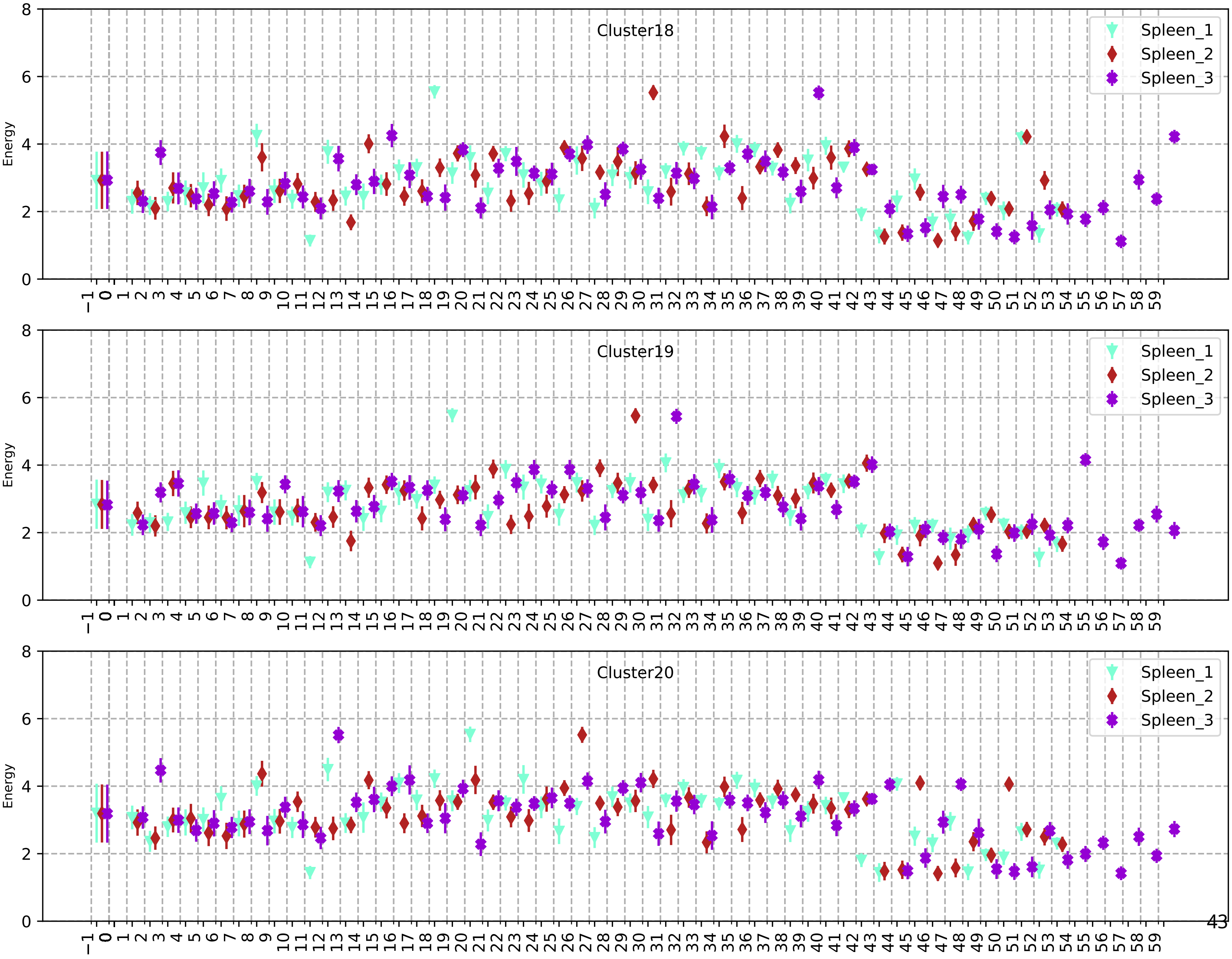
Cluster18
Cluster19
Cluster20
Spleen_1
Spleen_2
Spleen_3
Energy

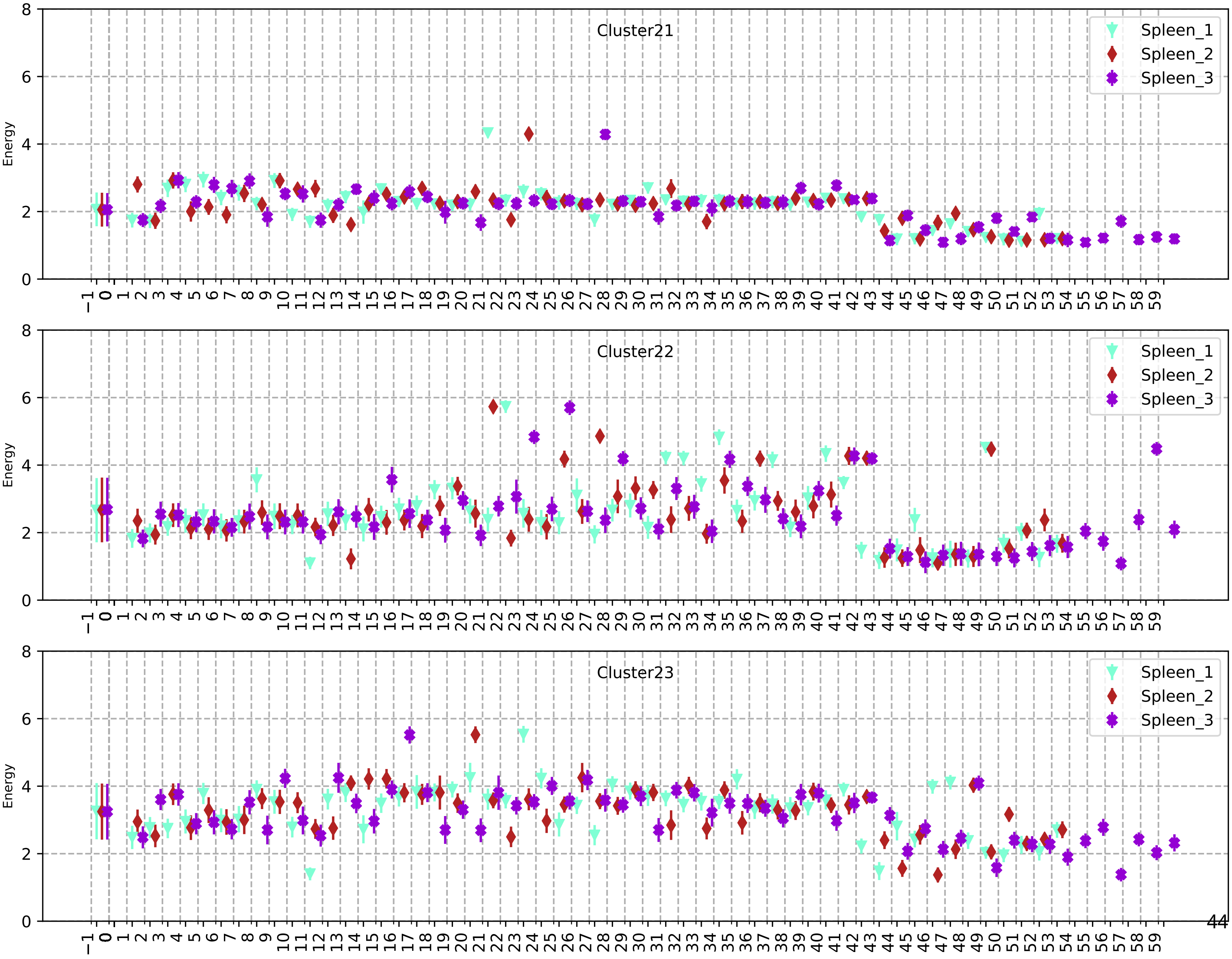
Cluster21
Cluster22
Cluster23
Energy
Spleen_1
Spleen_2
Spleen_3

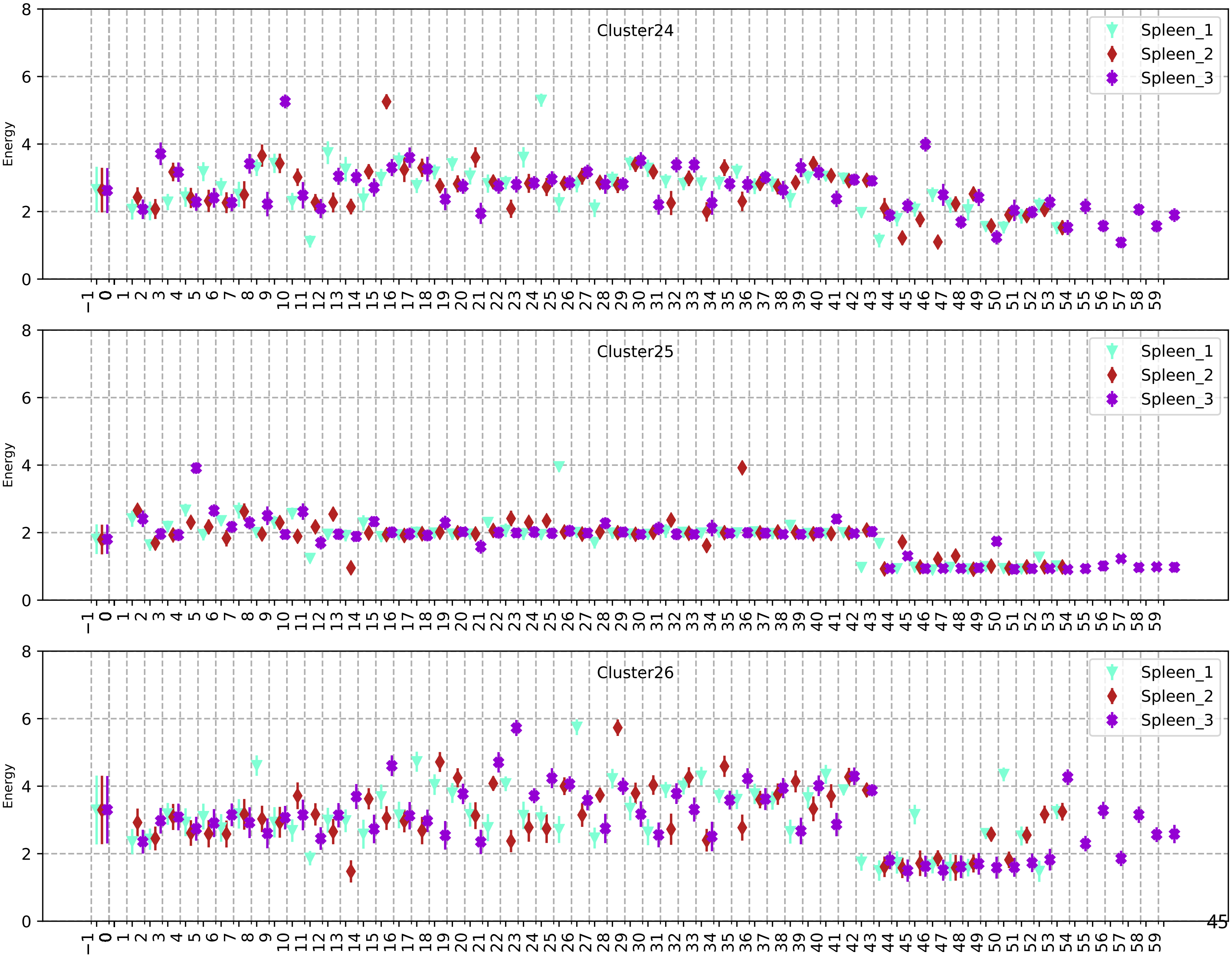
Cluster24
Cluster25
Cluster26
Spleen_1
Spleen_2
Spleen_3
Energy
0
2
4
6
8
−1 0 1 2 3 4 5 6 7 8 9 10 11 12 13 14 15 16 17 18 19 20 21 22 23 24 25 26 27 28 29 30 31 32 33 34 35 36 37 38 39 40 41 42 43 44 45 46 47 48 49 50 51 52 53 54 55 56 57 58 59

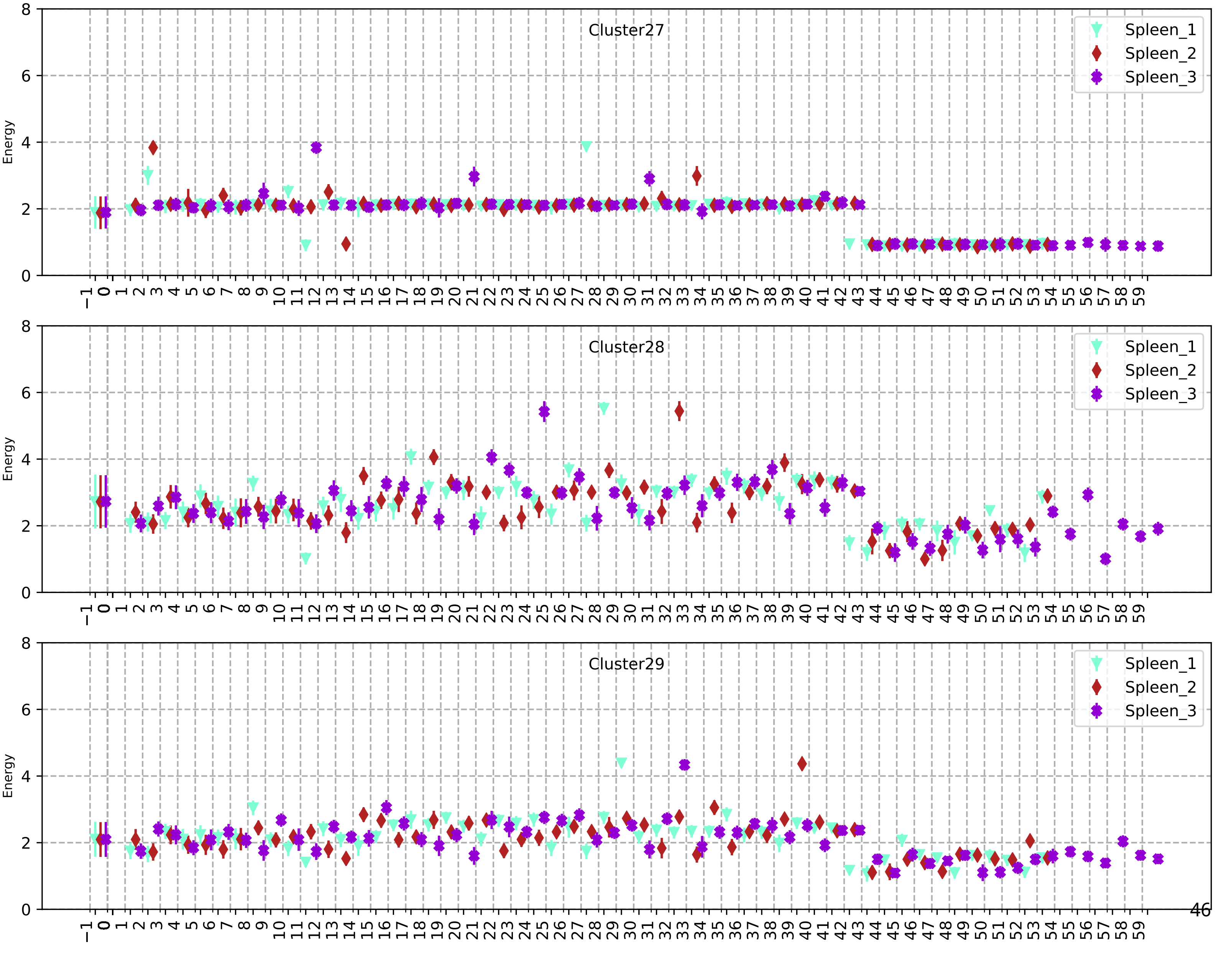

Cluster27
Cluster28
Cluster29
Spleen_1
Spleen_2
Spleen_3
Energy

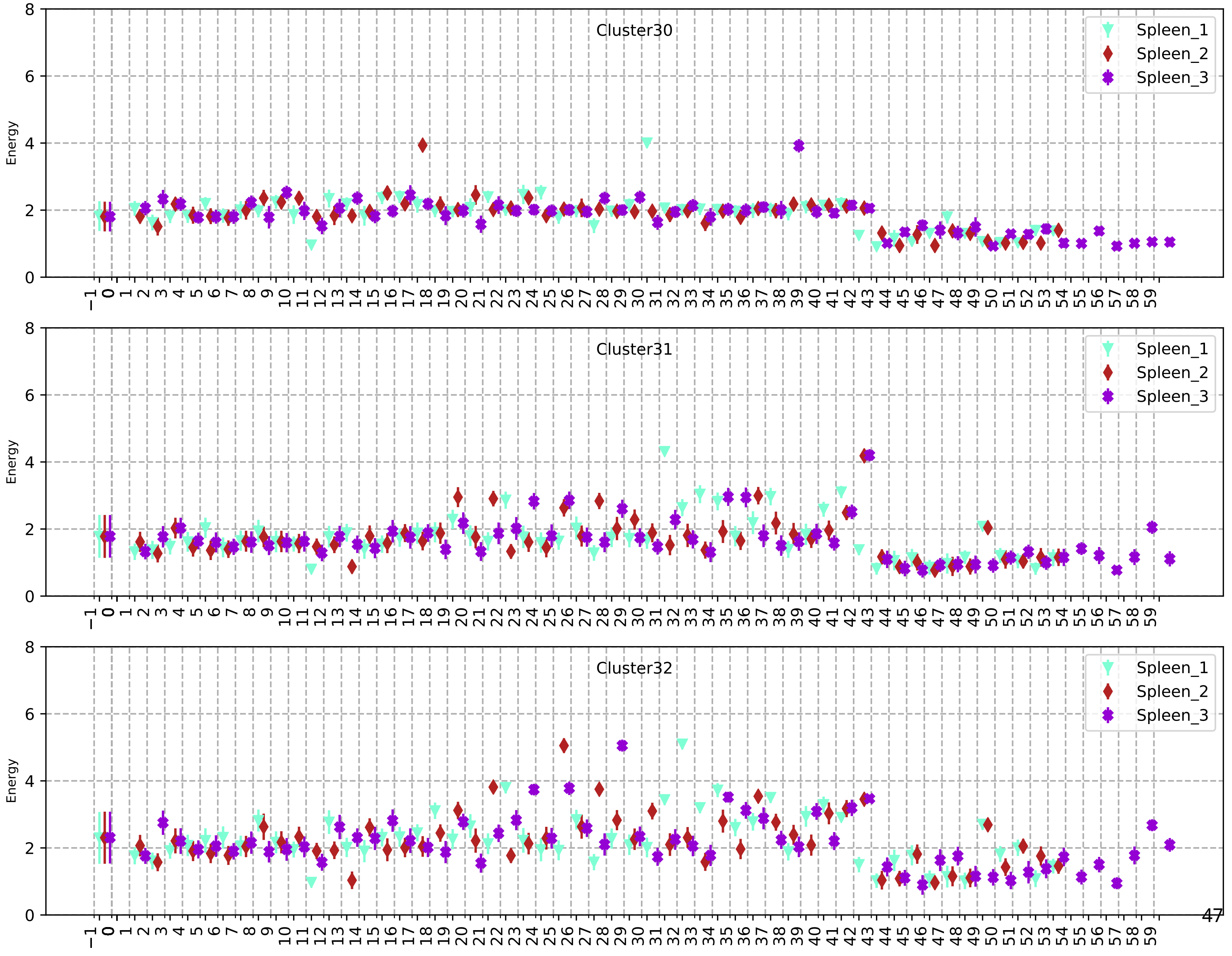
Cluster30
Spleen_1
Spleen_2
Spleen_3
Energy
Cluster31
Spleen_1
Spleen_2
Spleen_3
Energy
Cluster32
Spleen_1
Spleen_2
Spleen_3
Energy

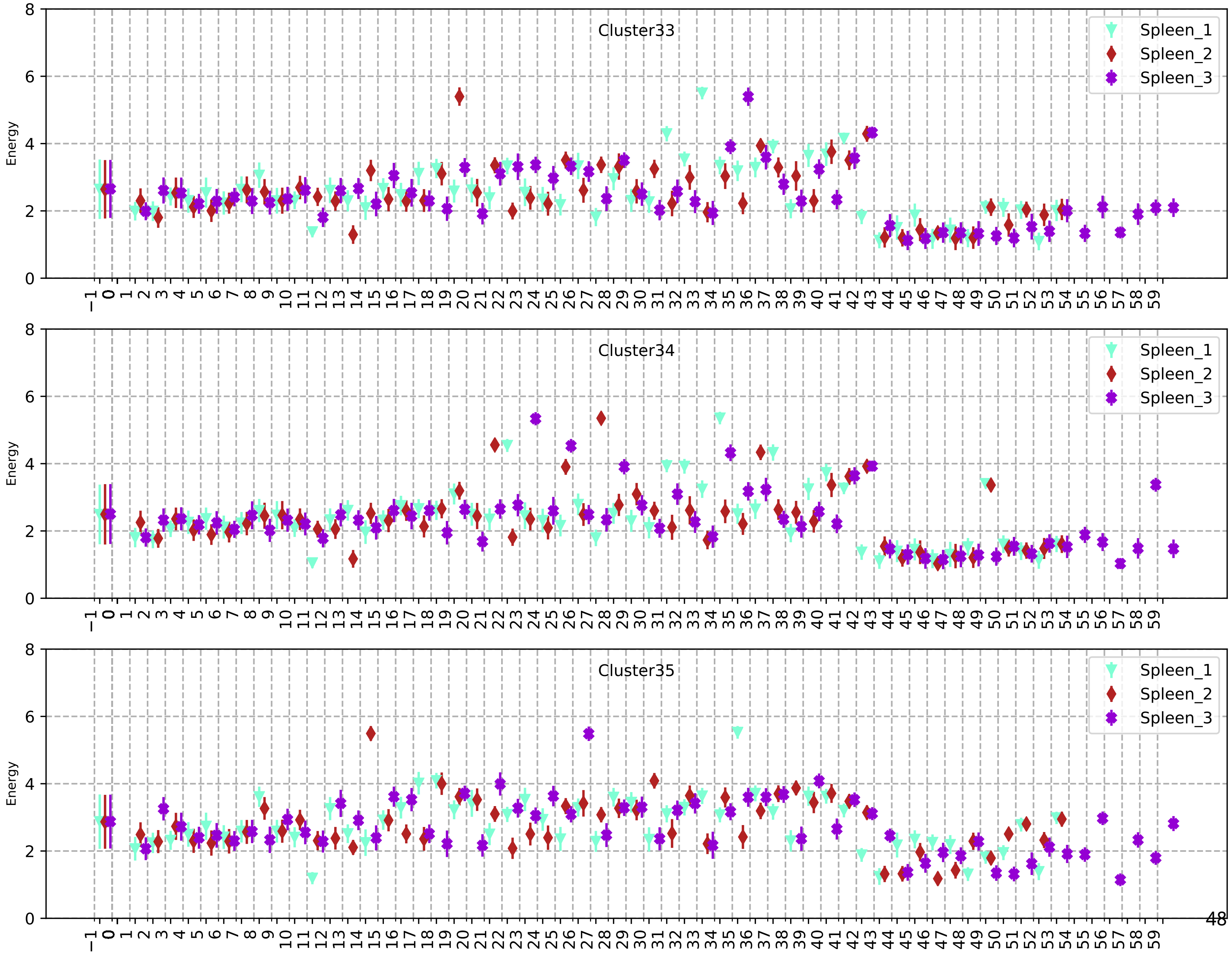
Cluster33
Cluster34
Cluster35
Energy
Spleen_1
Spleen_2
Spleen_3

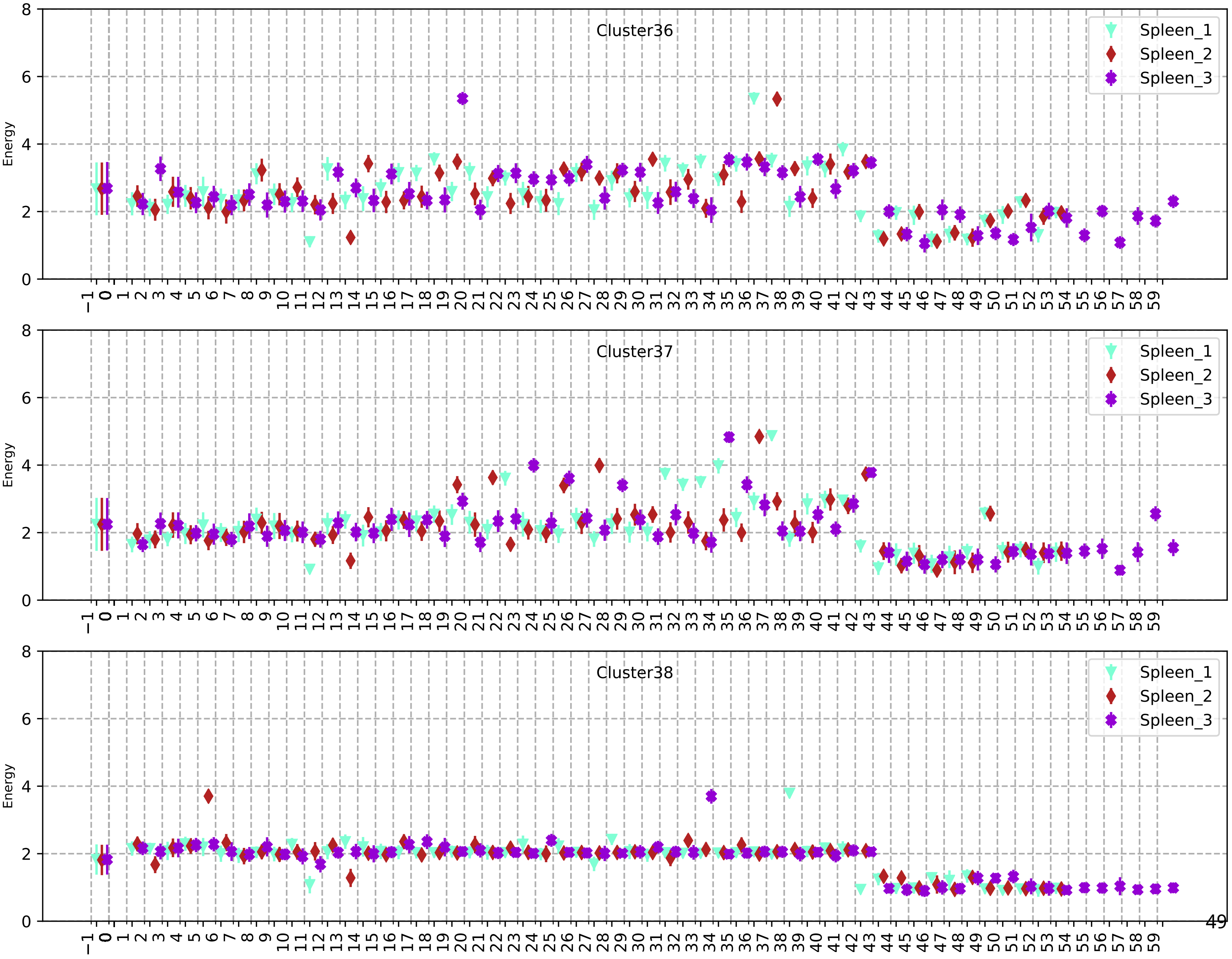
Cluster36
Cluster37
Cluster38
Spleen_1
Spleen_2
Spleen_3
Energy
0 2 4 6 8
−1 0 1 2 3 4 5 6 7 8 9 10 11 12 13 14 15 16 17 18 19 20 21 22 23 24 25 26 27 28 29 30 31 32 33 34 35 36 37 38 39 40 41 42 43 44 45 46 47 48 49 50 51 52 53 54 55 56 57 58 59

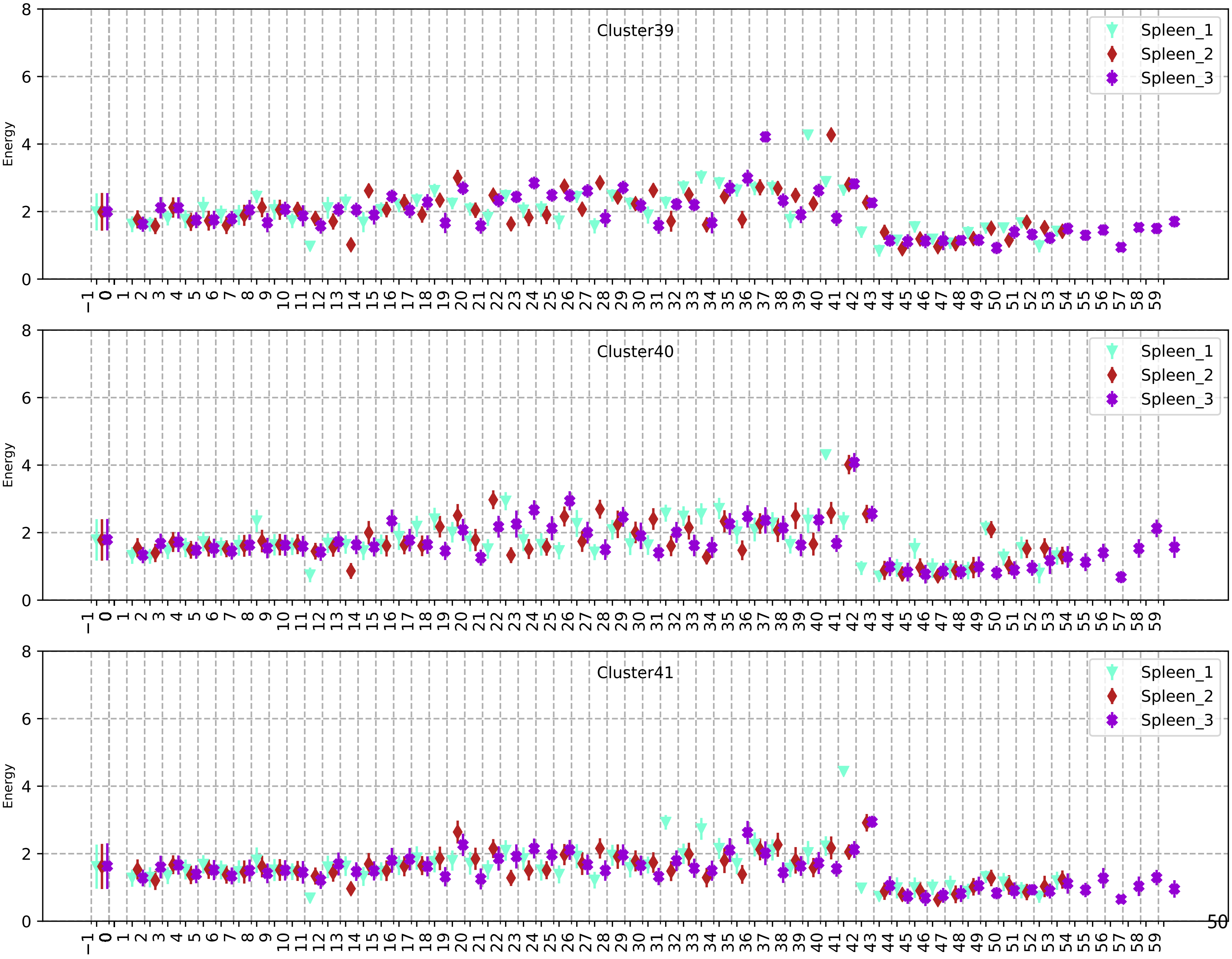
Cluster39
Spleen_1
Spleen_2
Spleen_3
Energy
Cluster40
Spleen_1
Spleen_2
Spleen_3
Energy
Cluster41
Spleen_1
Spleen_2
Spleen_3
Energy

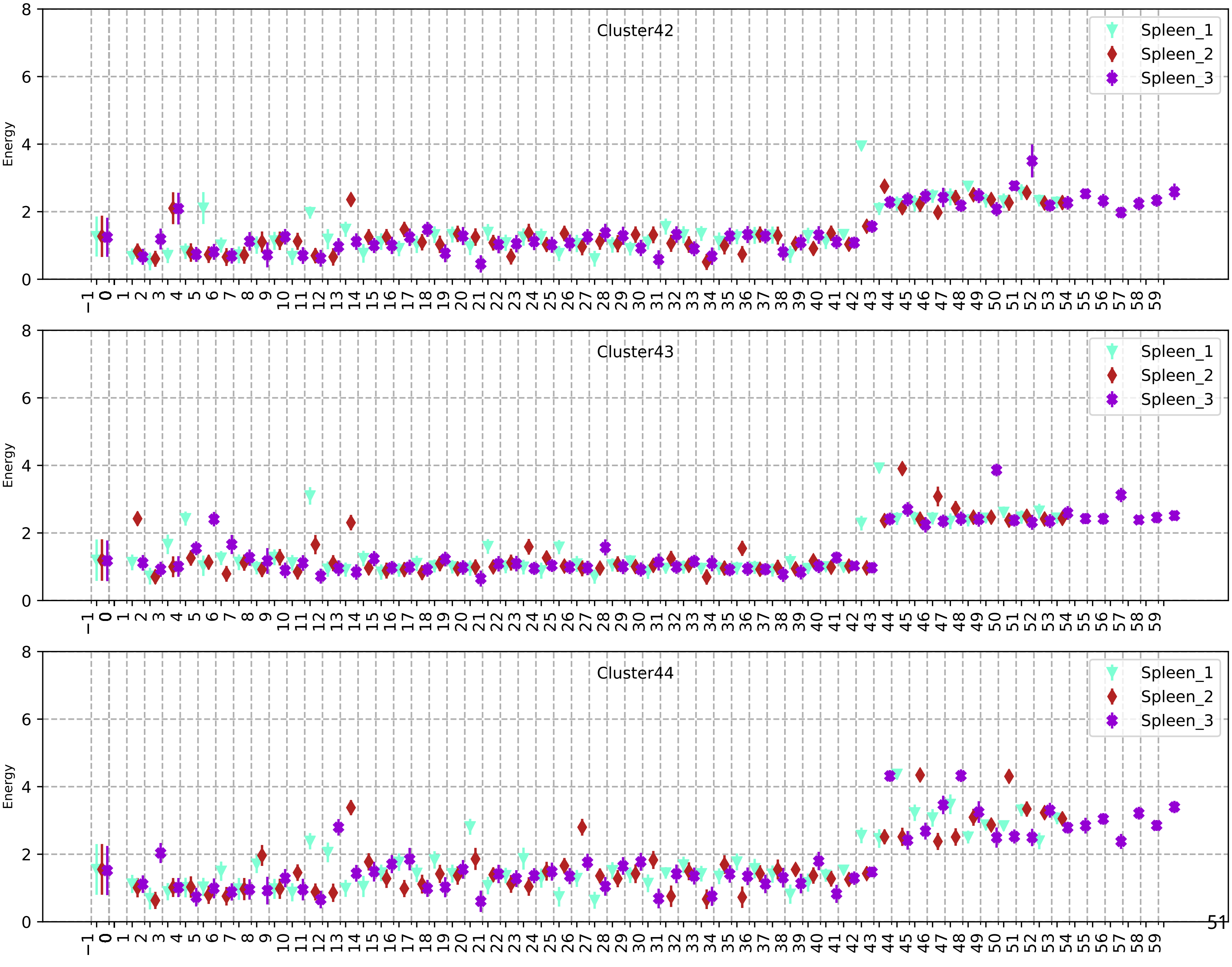
Cluster42
Cluster43
Cluster44
Spleen_1
Spleen_2
Spleen_3
Energy

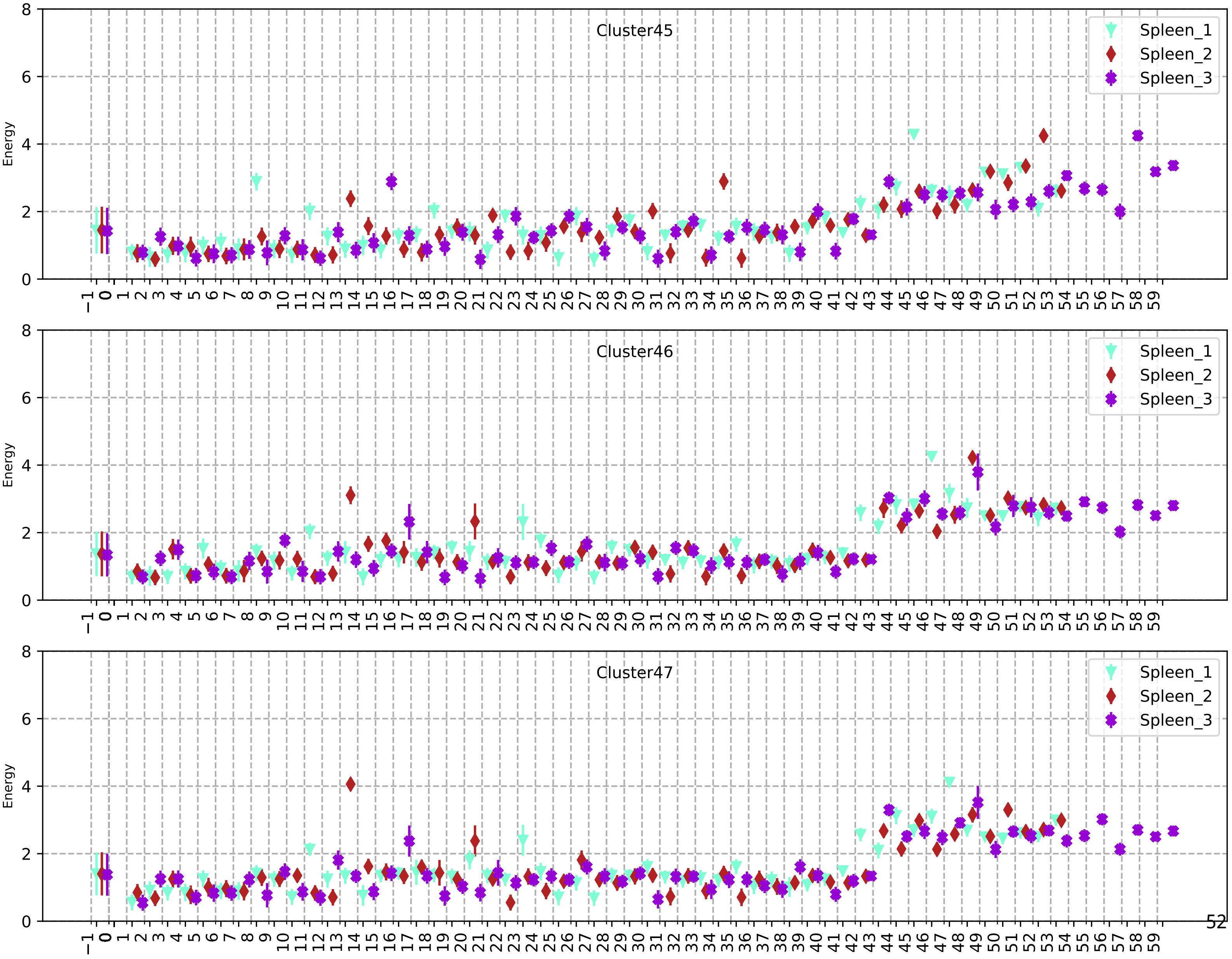
Cluster45
Cluster46
Cluster47
Spleen_1
Spleen_2
Spleen_3
Energy
0
2
4
6
8
−1 0 1 2 3 4 5 6 7 8 9 10 11 12 13 14 15 16 17 18 19 20 21 22 23 24 25 26 27 28 29 30 31 32 33 34 35 36 37 38 39 40 41 42 43 44 45 46 47 48 49 50 51 52 53 54 55 56 57 58 59

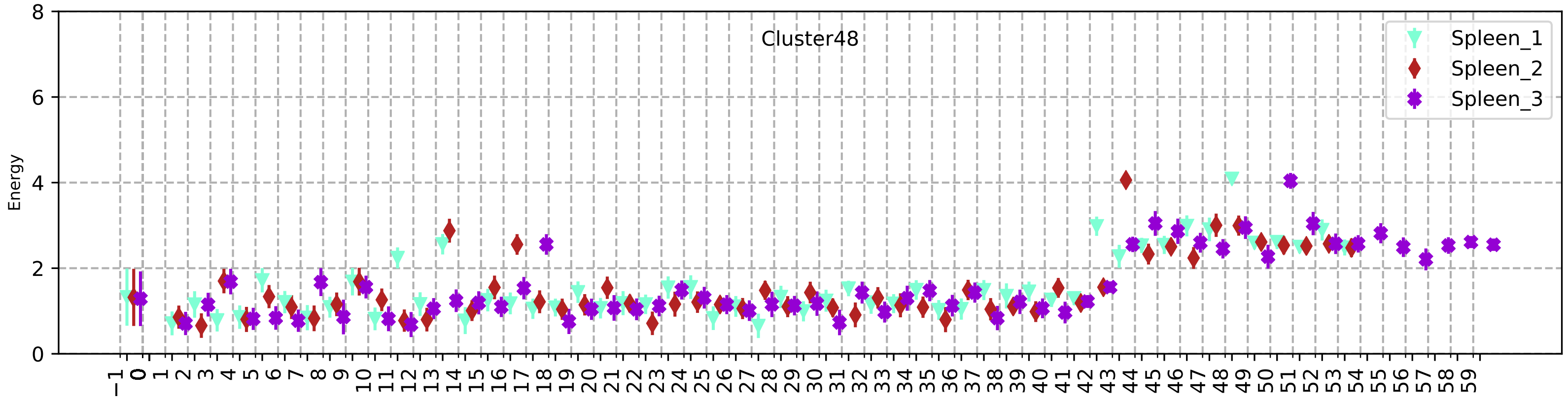
Cluster48
Spleen_1
Spleen_2
Spleen_3
Energy

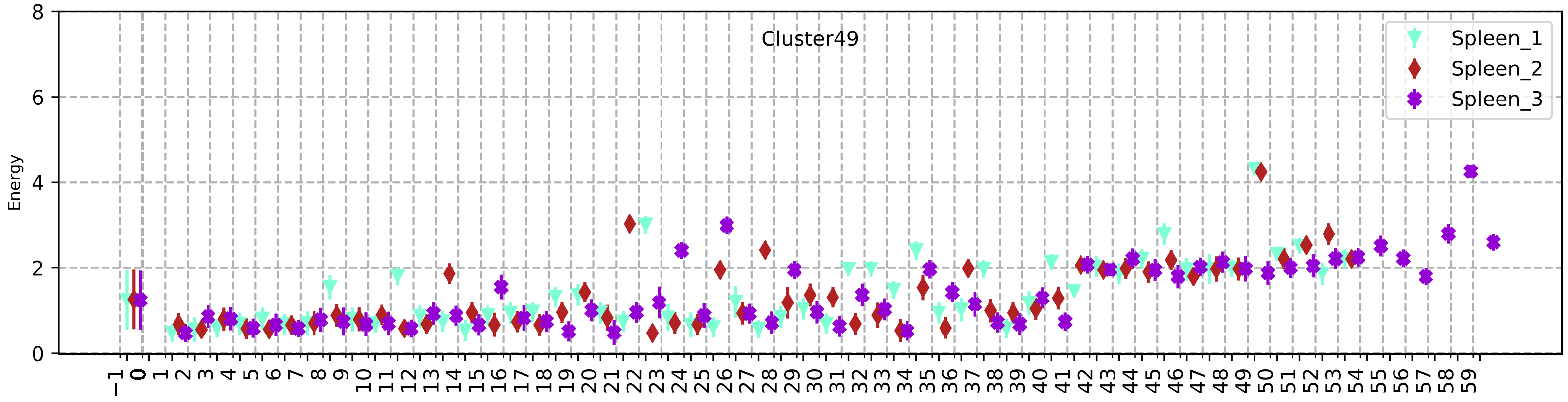
Cluster49
Spleen_1
Spleen_2
Spleen_3
Energy

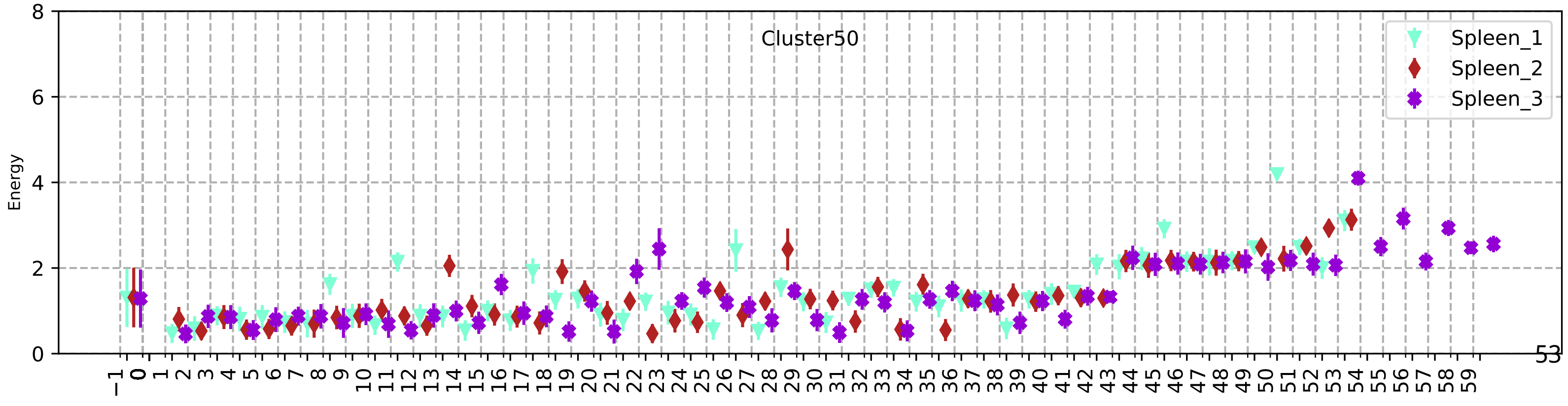
Cluster50
Spleen_1
Spleen_2
Spleen_3
Energy

100

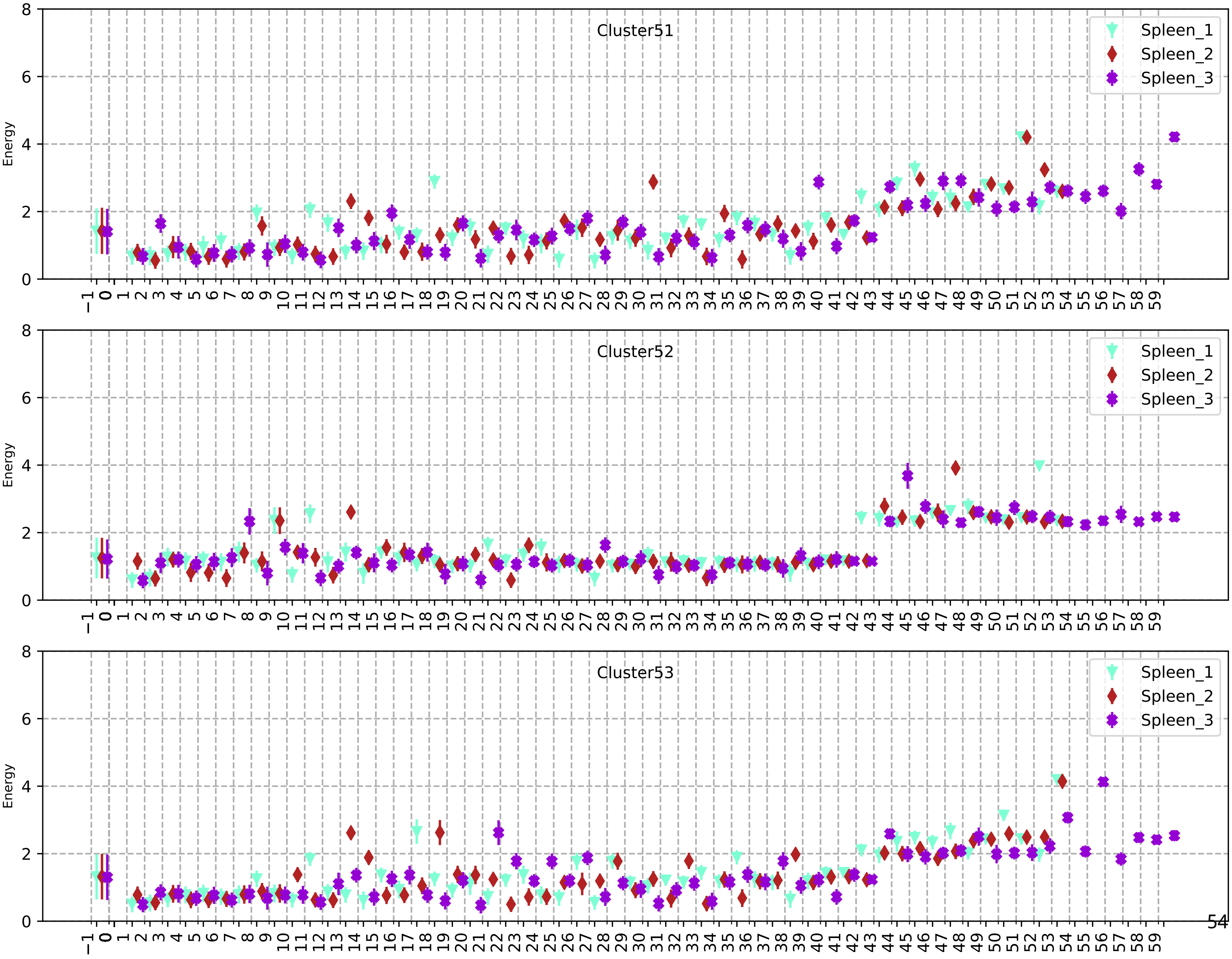
Cluster51
Cluster52
Cluster53
Spleen_1
Spleen_2
Spleen_3
Energy

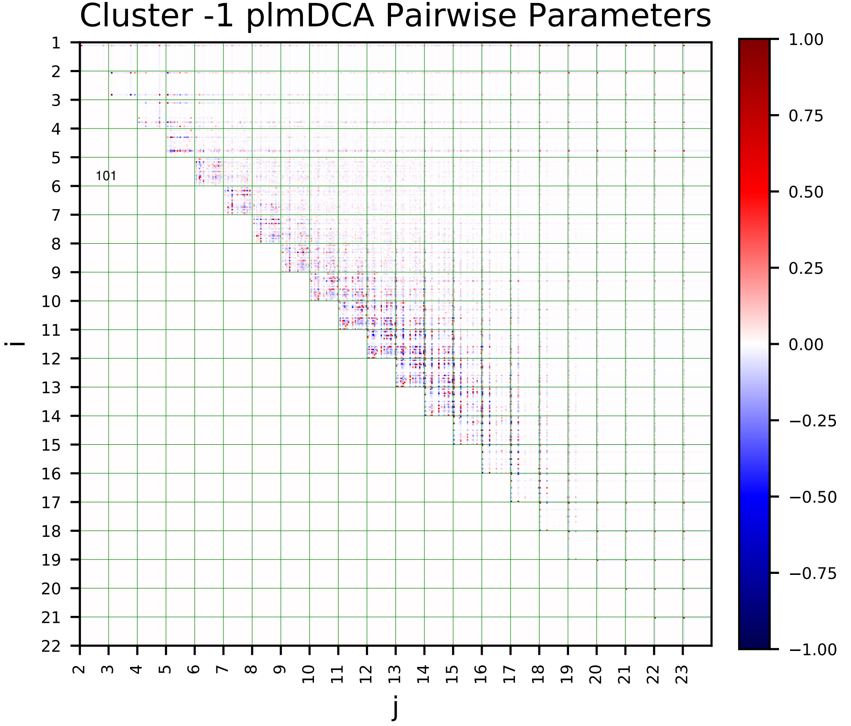
Cluster -1 plmDCA Pairwise Parameters
i
j
101

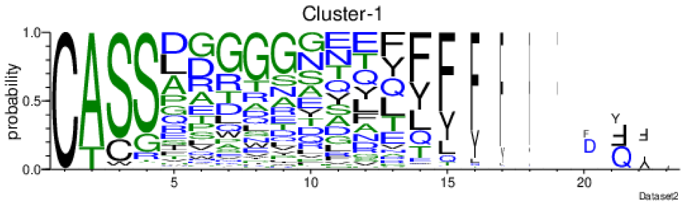
Cluster-1
probability
Dataset2

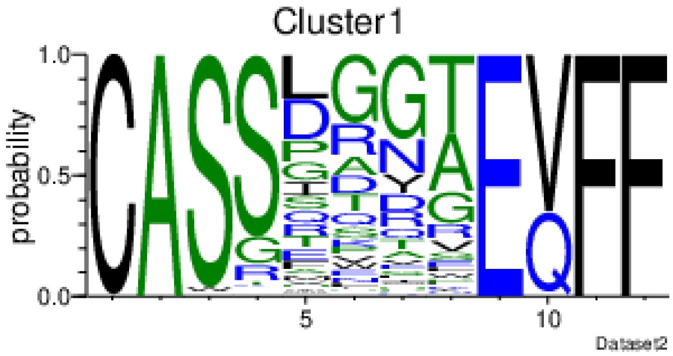
Cluster1
probability
Dataset2

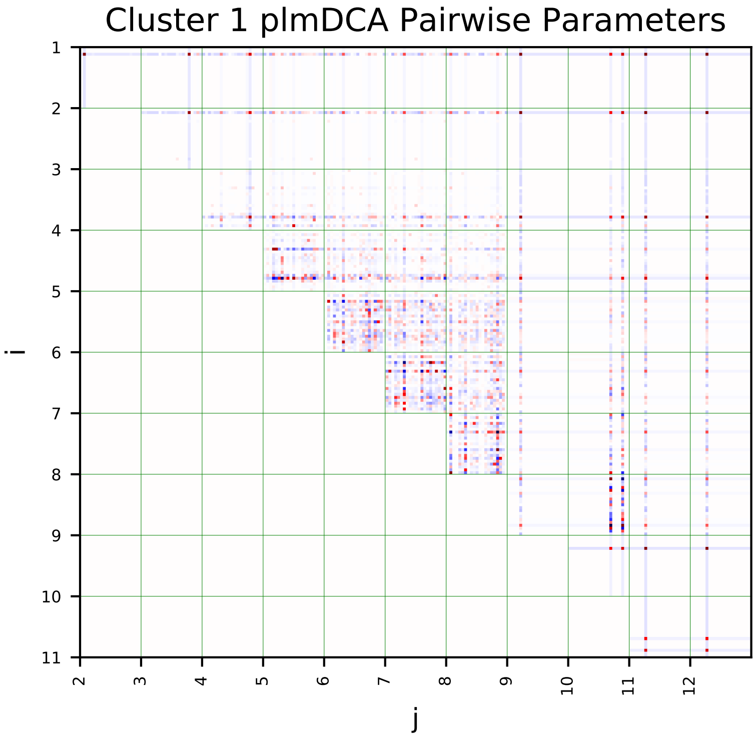
Cluster 1 plmDCA Pairwise Parameters
i
j

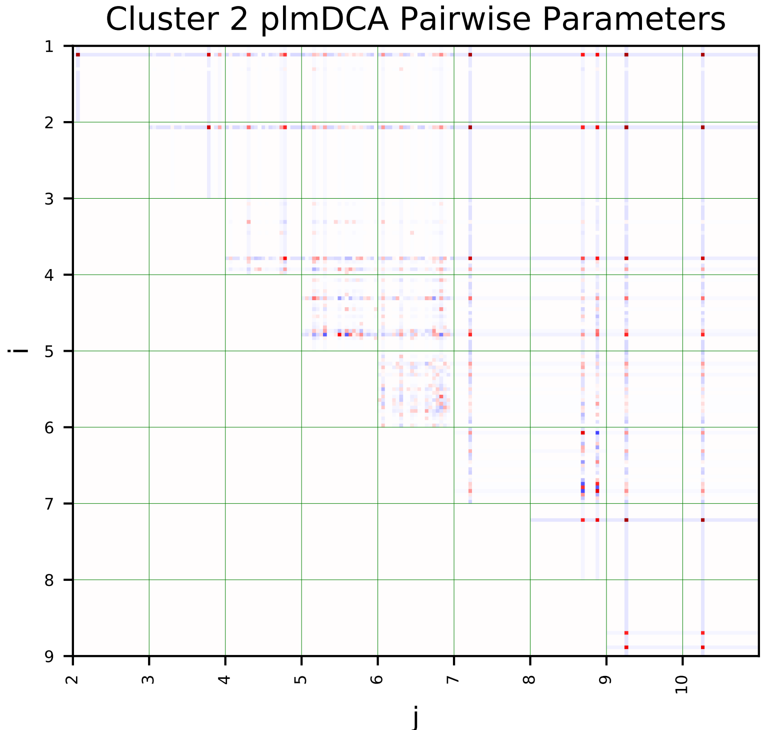
Cluster 2 plmDCA Pairwise Parameters
i
j

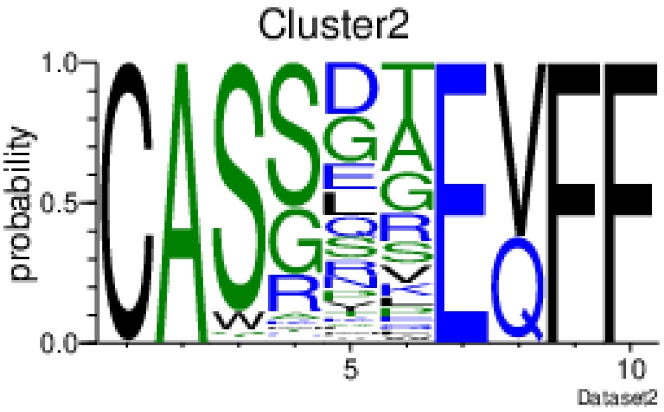
Cluster2
probability
Dataset2

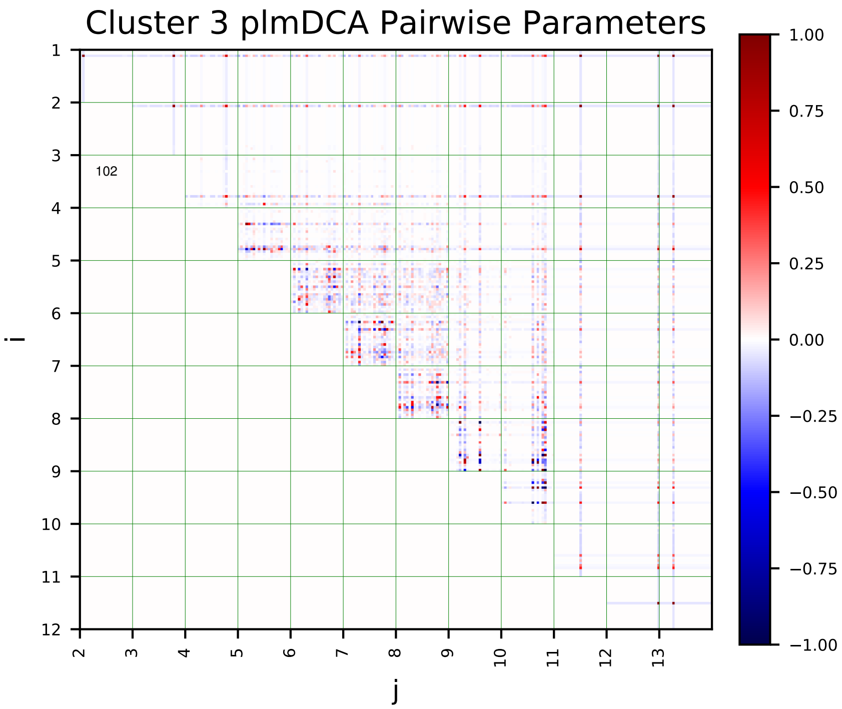
Cluster 3 plmDCA Pairwise Parameters
i
j
102

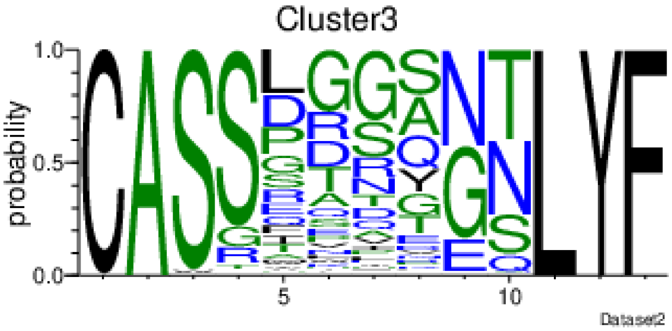
Cluster3
probability
Dataset2

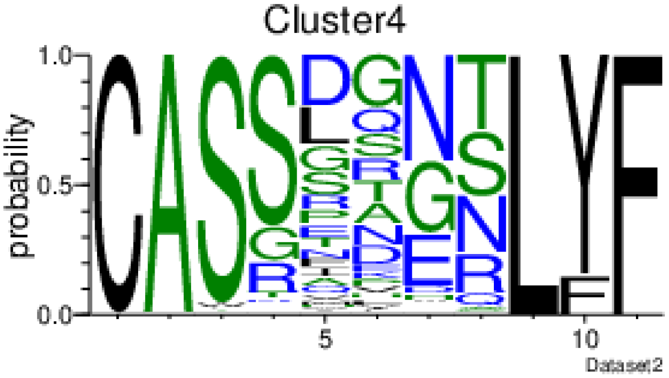
Cluster4
probability
Dataset2

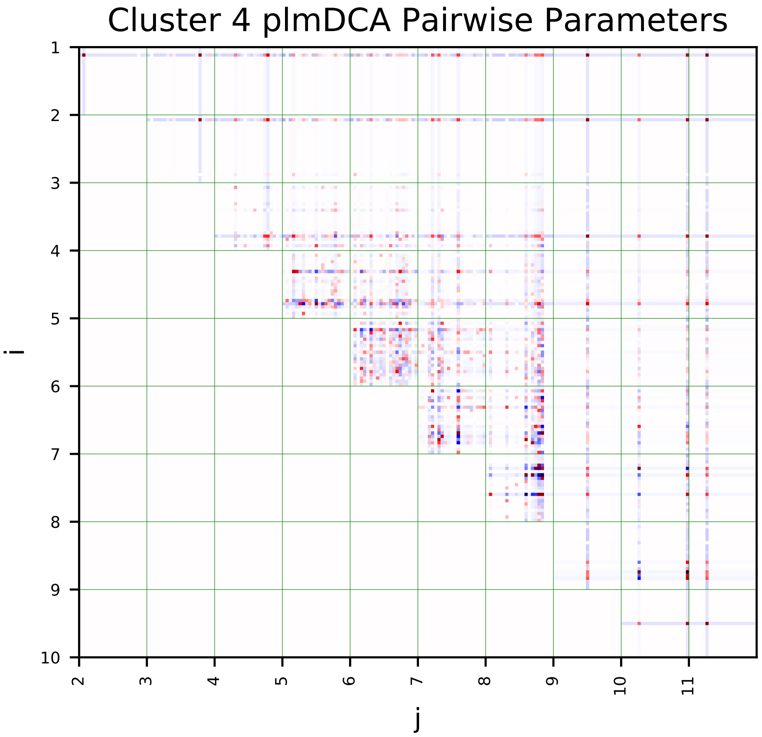
Cluster 4 plmDCA Pairwise Parameters
i
j

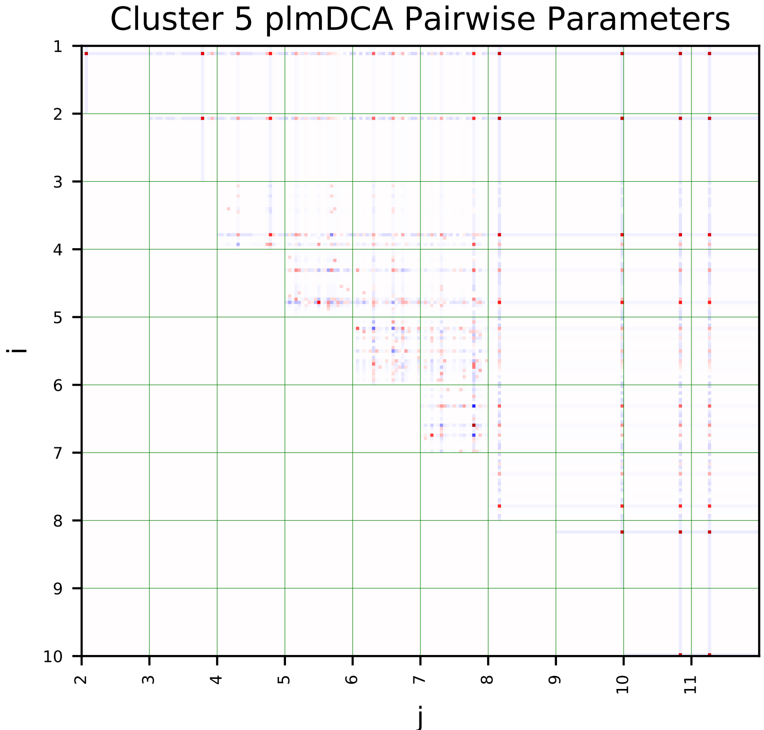
Cluster 5 plmDCA Pairwise Parameters
i
j

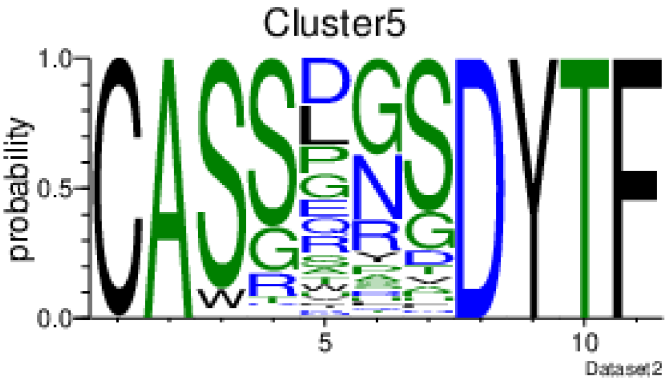
Cluster5
probability
Dataset2

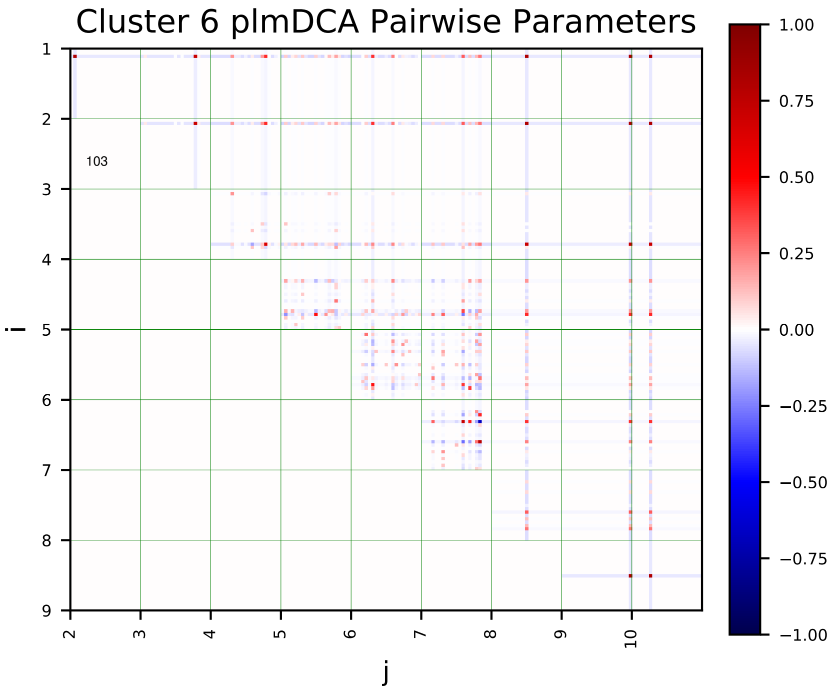
Cluster 6 plmDCA Pairwise Parameters
103
i
j

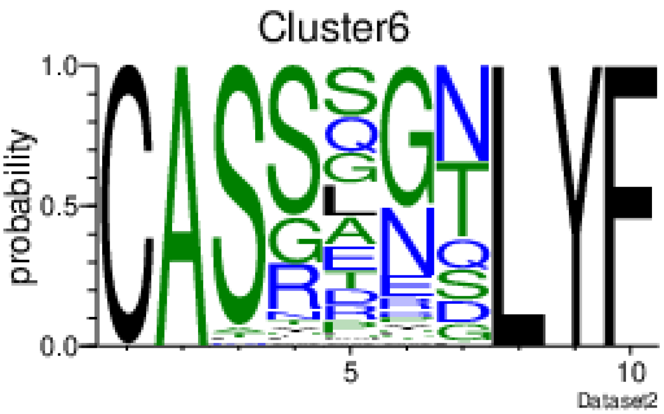
Cluster6
probability
Dataset2

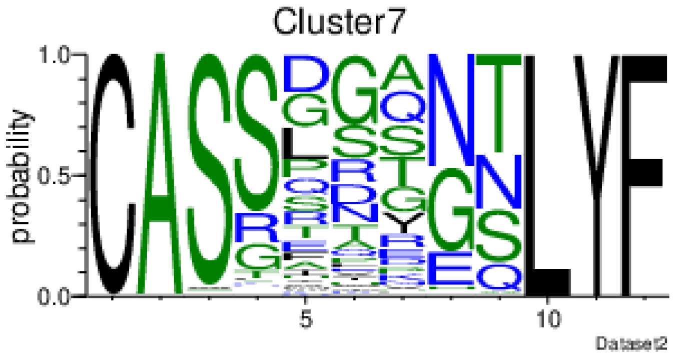
Cluster7
probability
Dataset2

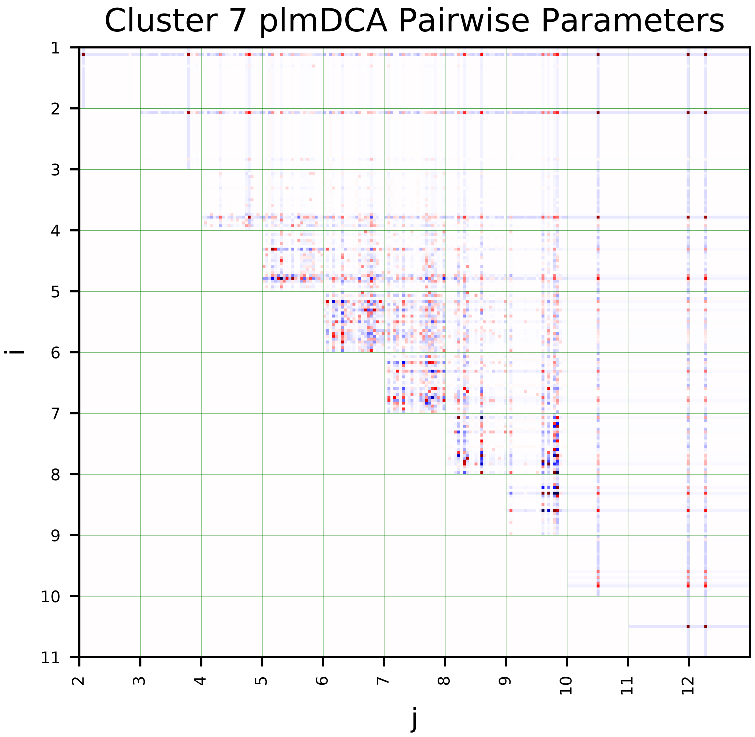
Cluster 7 plmDCA Pairwise Parameters
i
j

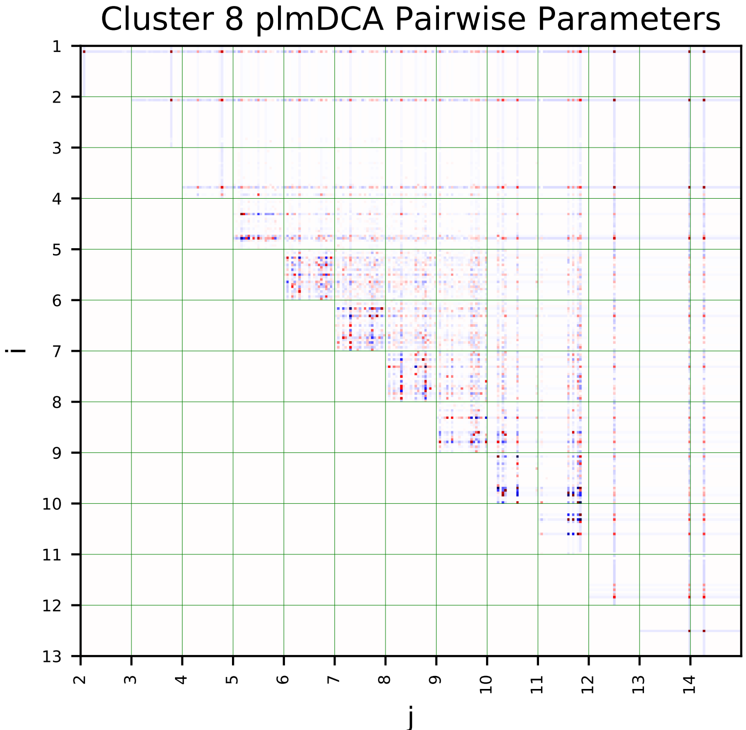
Cluster 8 plmDCA Pairwise Parameters
i
j

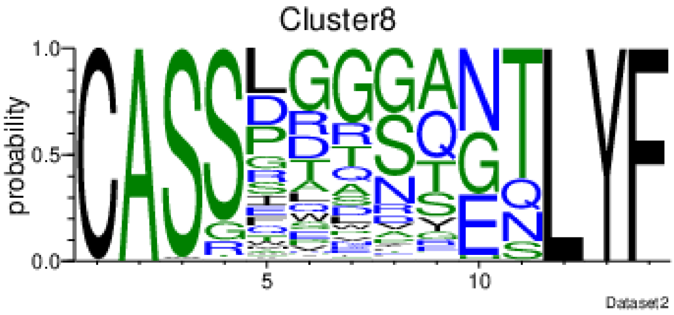
Cluster8
probability
Dataset2

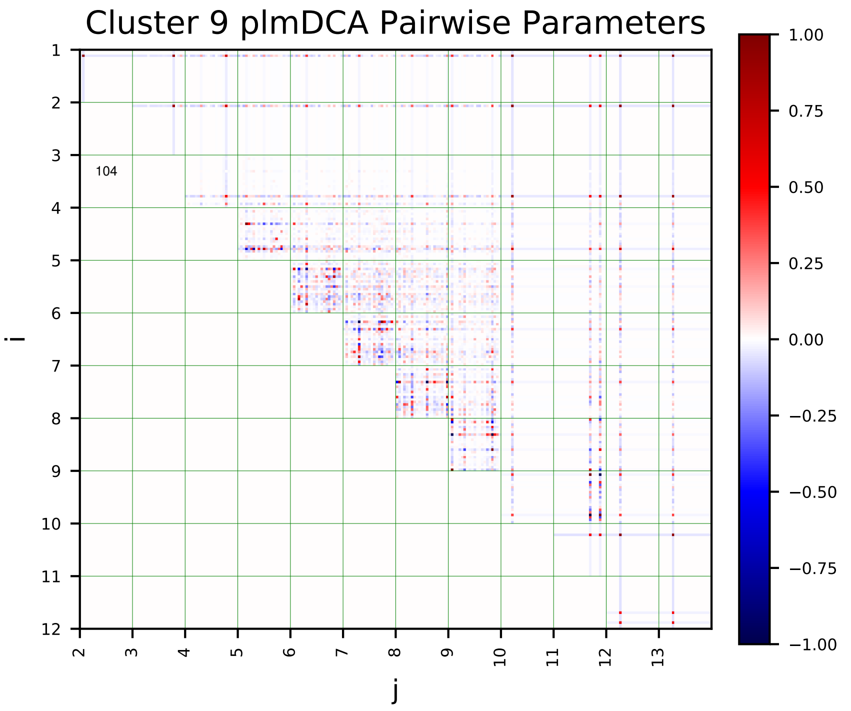
Cluster 9 plmDCA Pairwise Parameters
i
j
104

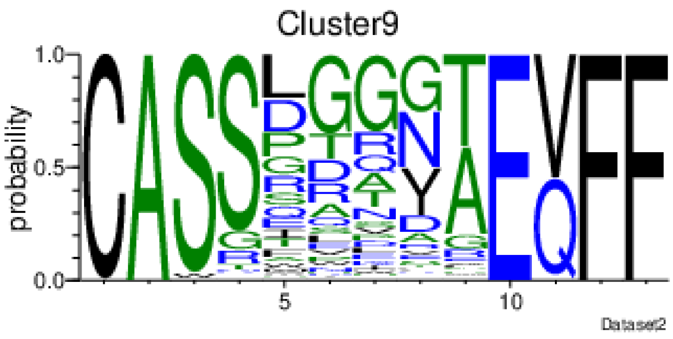
Cluster9
probability
Dataset2

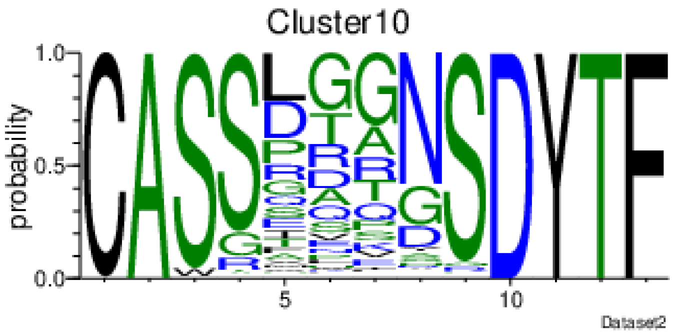
Cluster10
probability
Dataset2

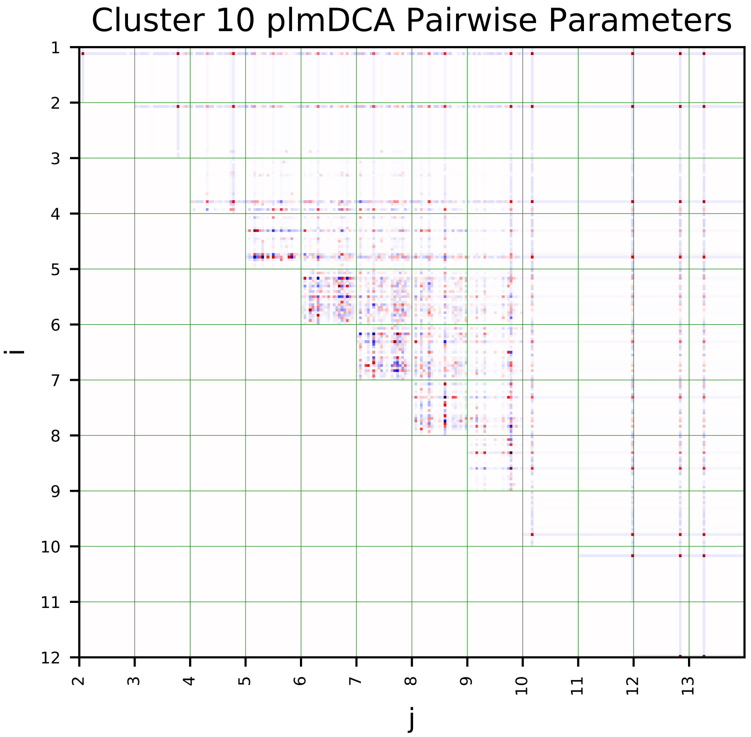
Cluster 10 plmDCA Pairwise Parameters
i
j

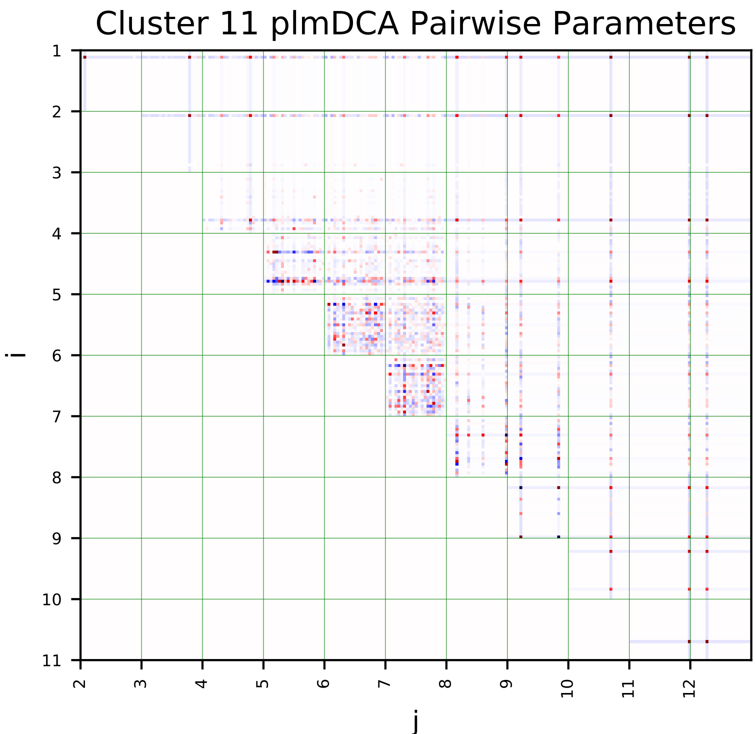
Cluster 11 plmDCA Pairwise Parameters
i
j

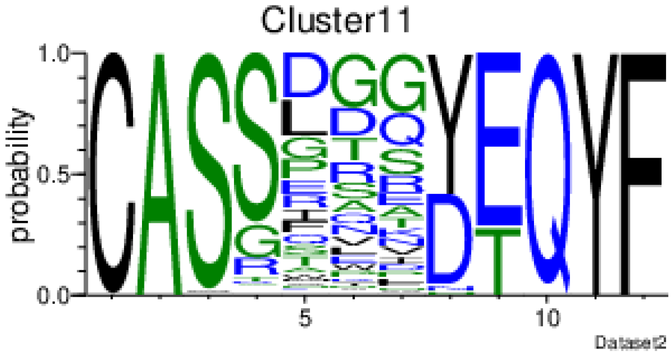
Cluster11
probability
Dataset2

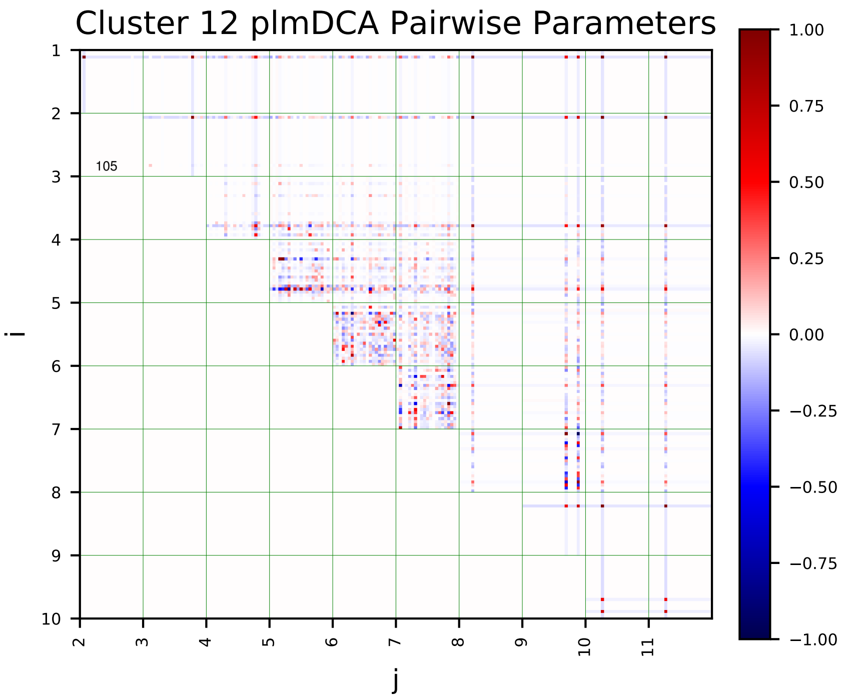
Cluster 12 plmDCA Pairwise Parameters
i
j
105
1.00
0.75
0.50
0.25
0.00
−0.25
−0.50
−0.75
−1.00

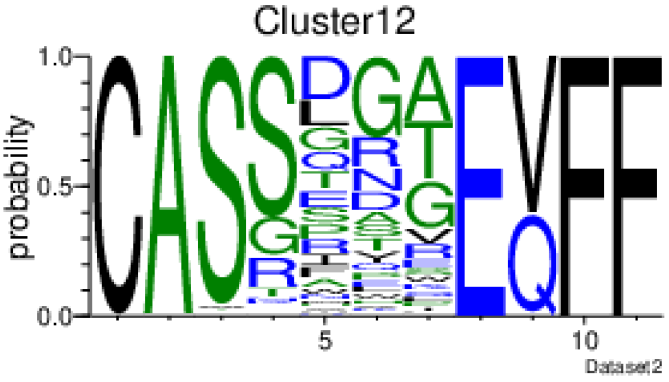
Cluster12
probability
Dataset2

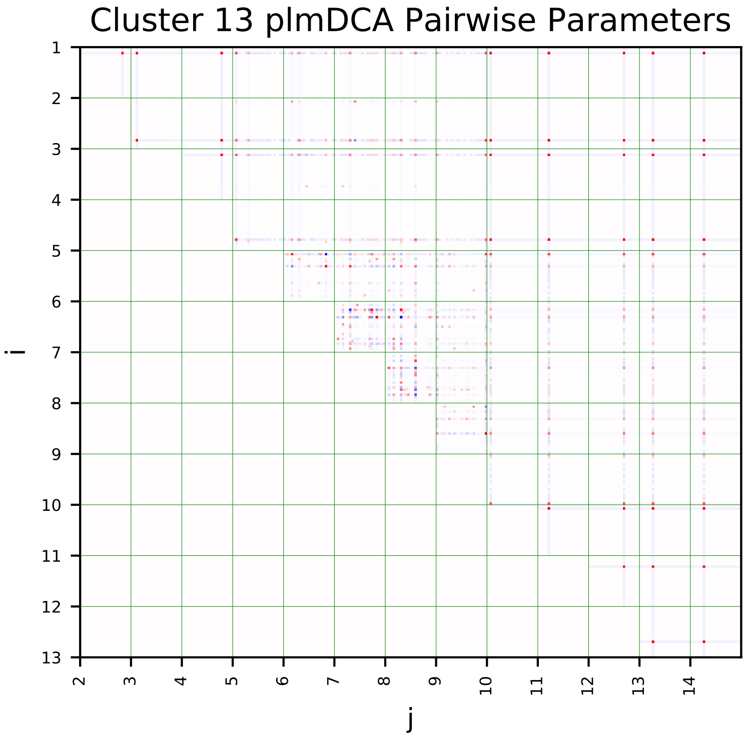
Cluster 13 plmDCA Pairwise Parameters
i
j

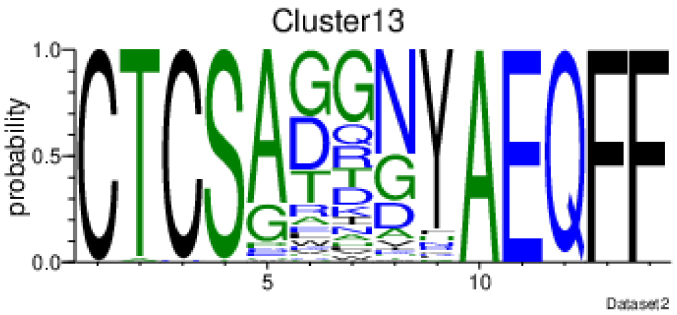
Cluster13
probability
Dataset2

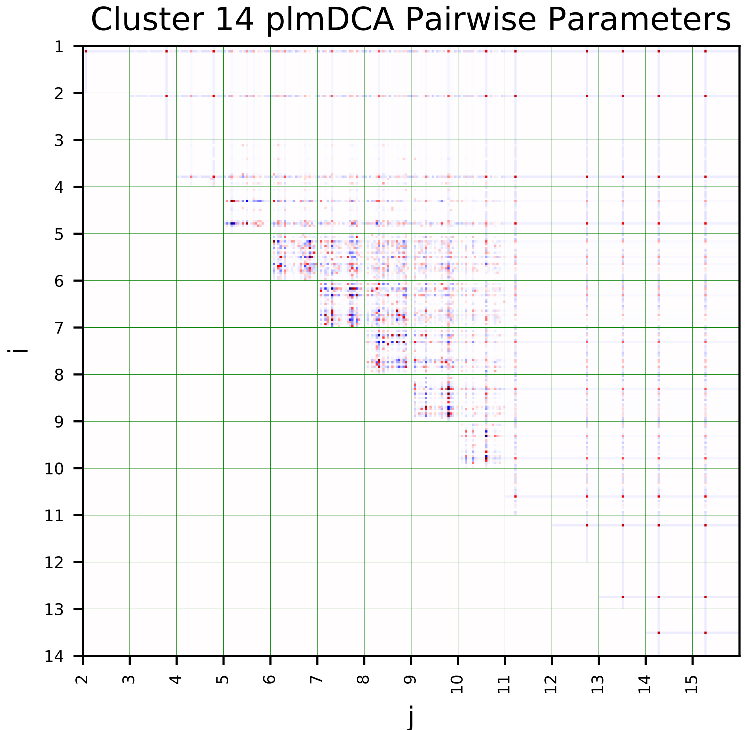
Cluster 14 plmDCA Pairwise Parameters
i
j

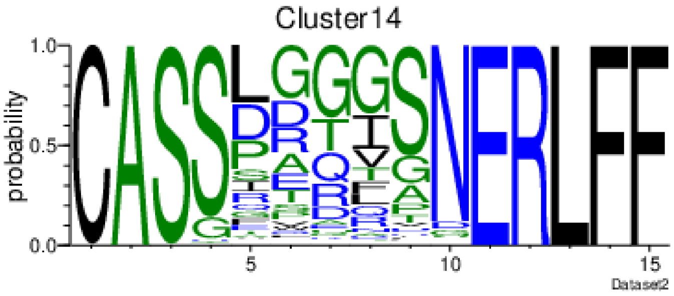
Cluster14
probability
Dataset2

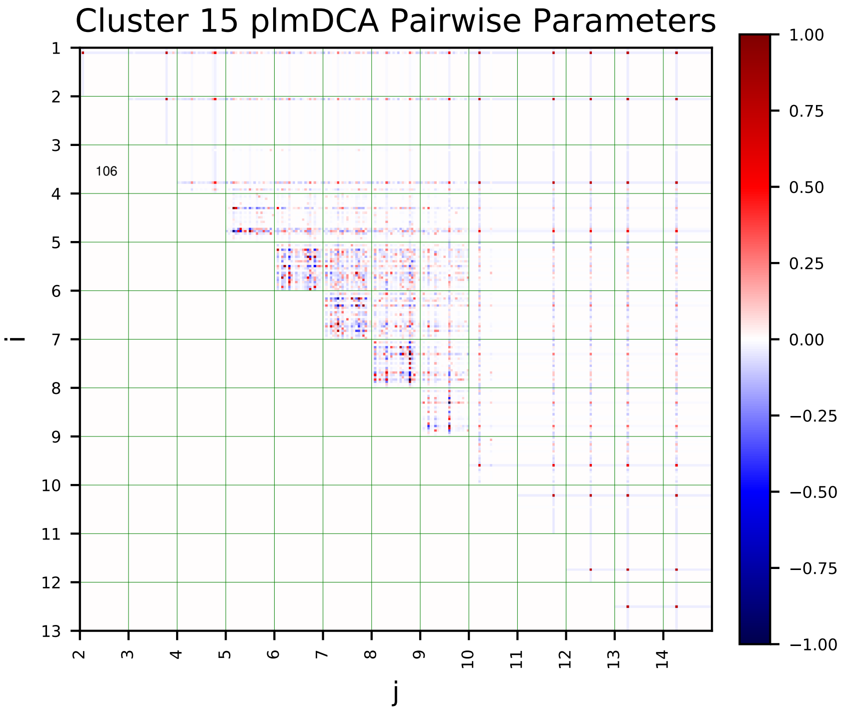
Cluster 15 plmDCA Pairwise Parameters
i
j
106
1.00
0.75
0.50
0.25
0.00
−0.25
−0.50
−0.75
−1.00

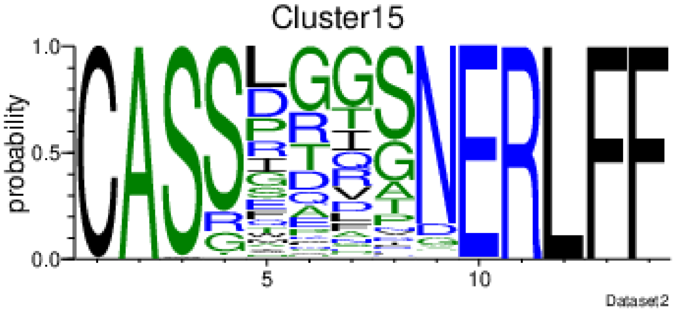
Cluster15
probability
Dataset2

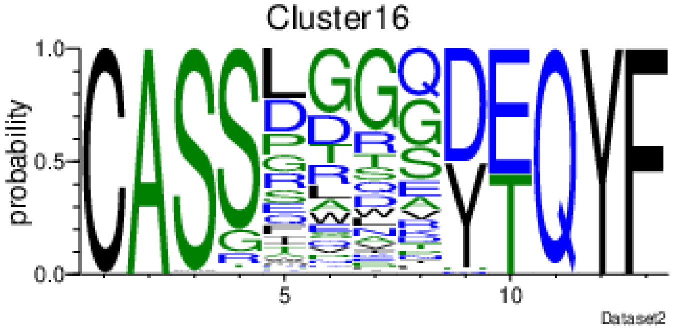
Cluster16
probability
Dataset2

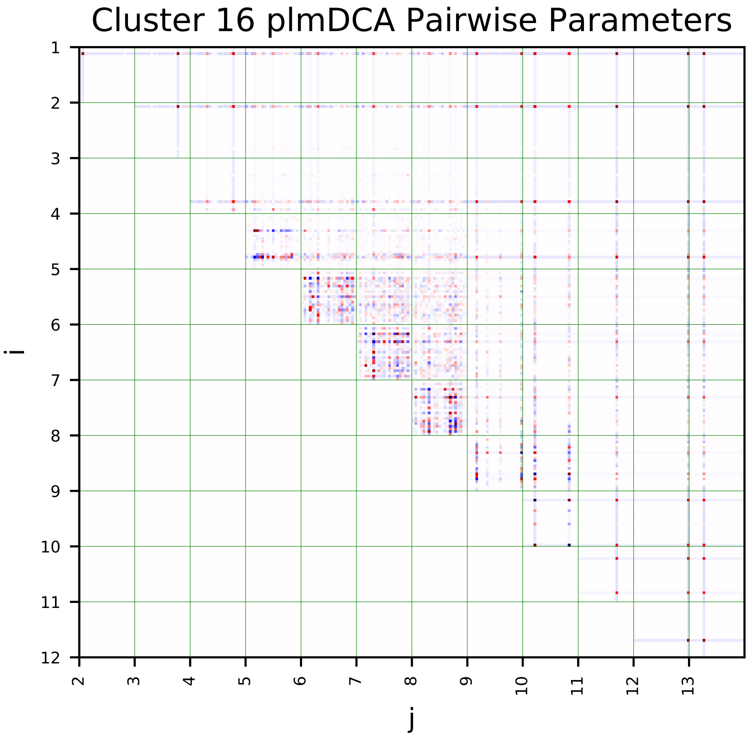
Cluster 16 plmDCA Pairwise Parameters
i
j

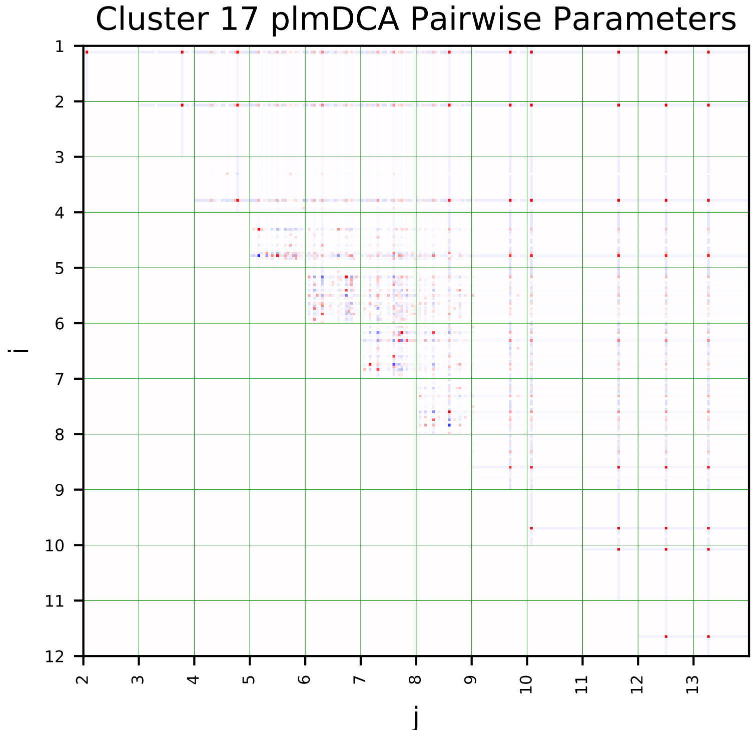
Cluster 17 plmDCA Pairwise Parameters
i
j

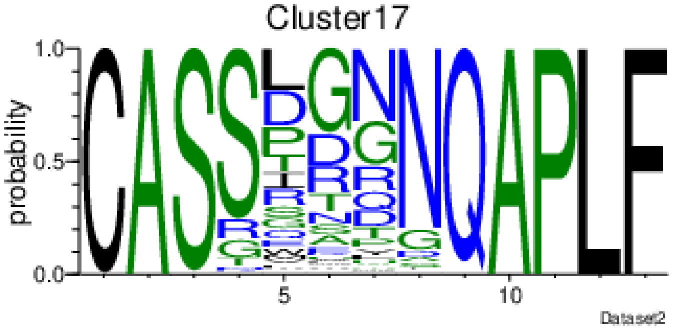
Cluster17
probability
Dataset2

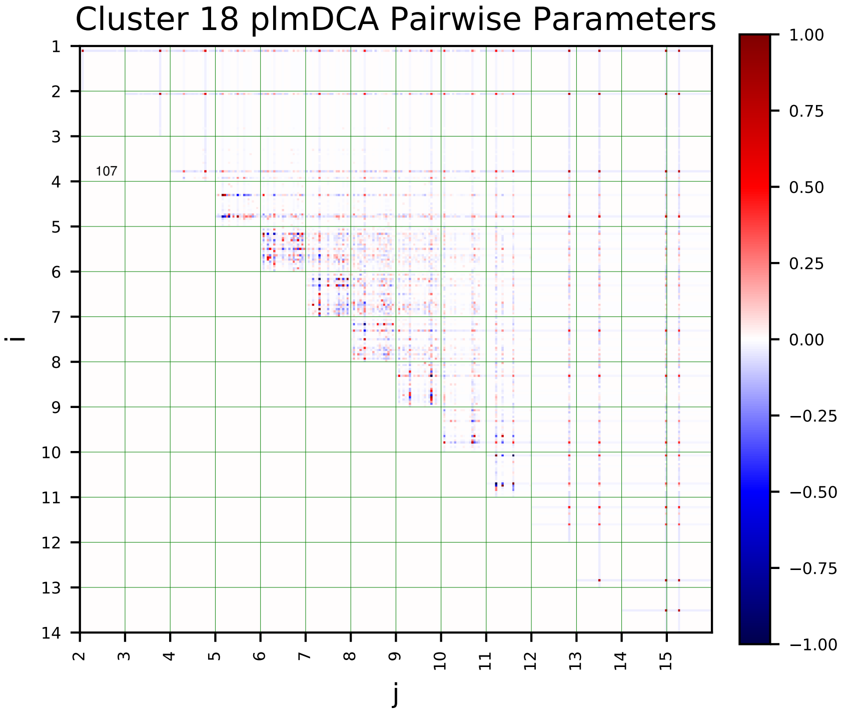
Cluster 18 plmDCA Pairwise Parameters
i
j
107

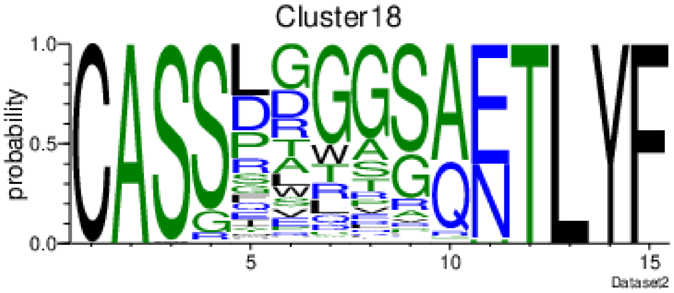
Cluster18
probability
Dataset2

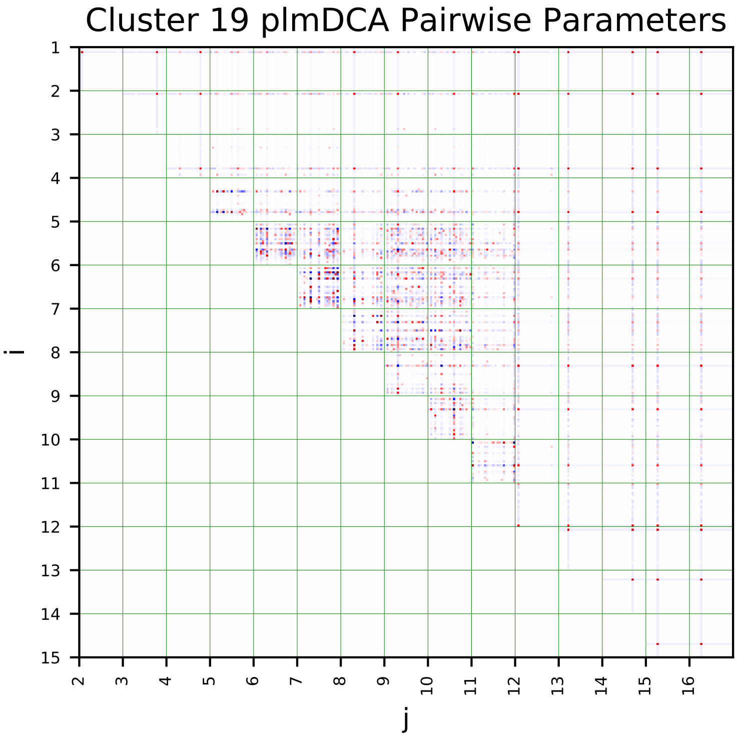
Cluster 19 plmDCA Pairwise Parameters
i
j

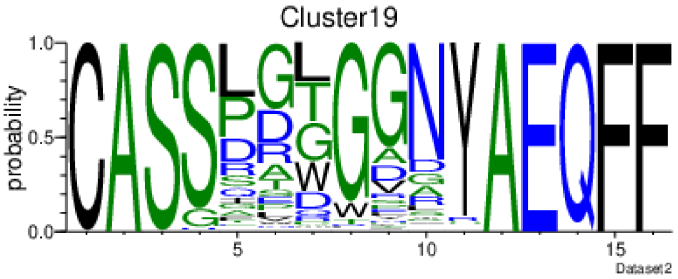
Cluster19
probability
Dataset2

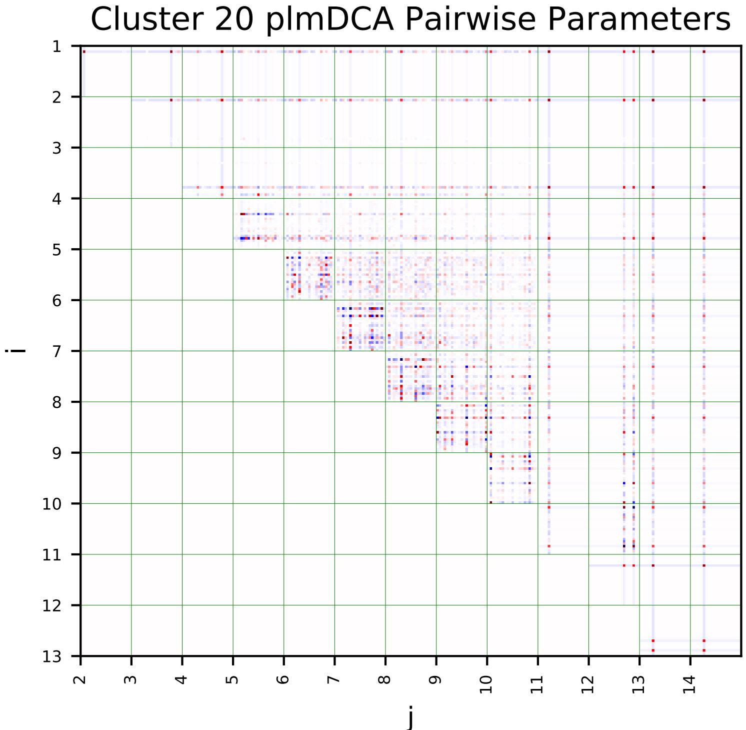
Cluster 20 plmDCA Pairwise Parameters
i
j

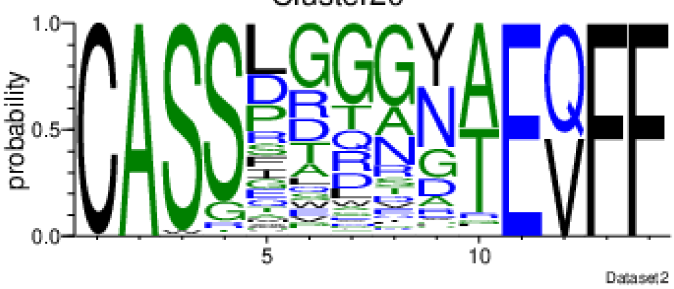
Cluster20
probability
Dataset2

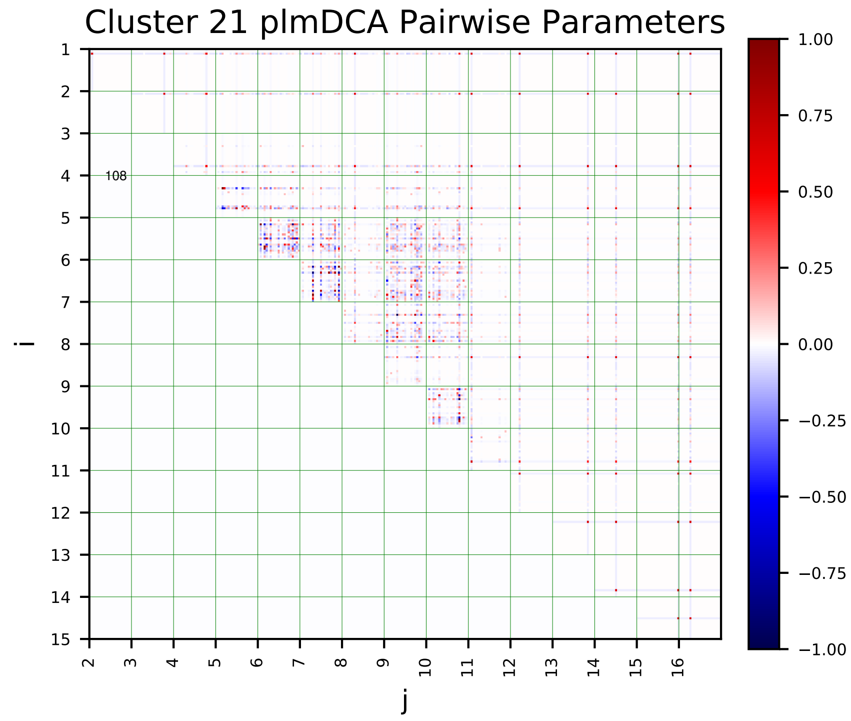
Cluster 21 plmDCA Pairwise Parameters
i
j

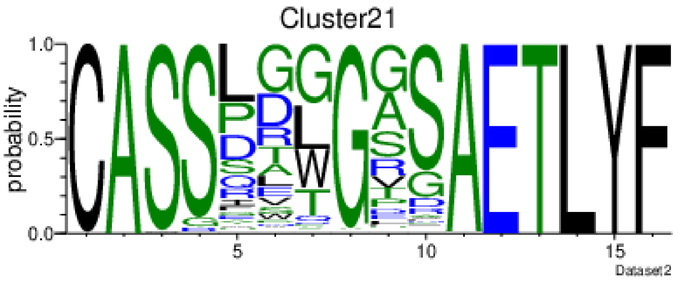
Cluster21
probability
Dataset2

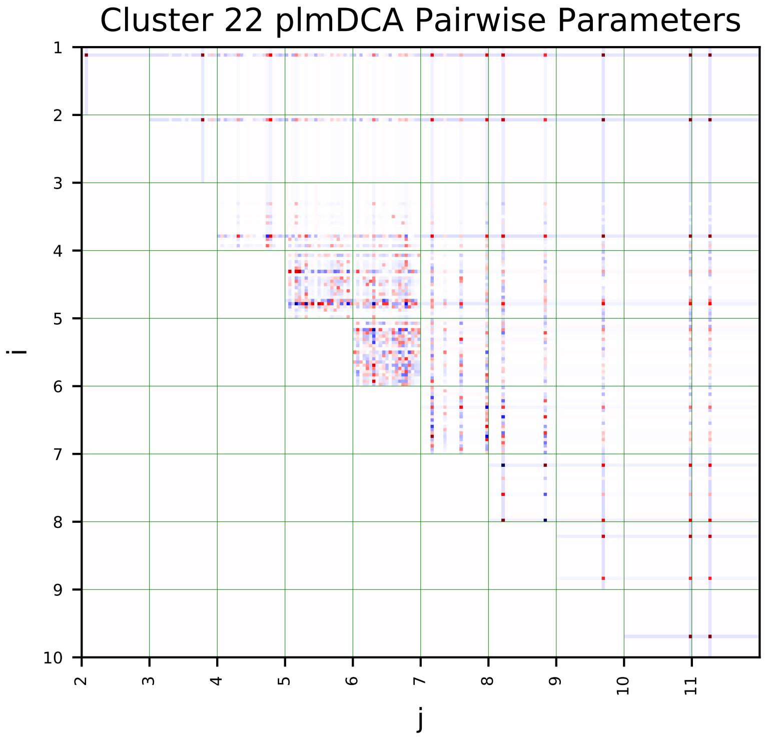
Cluster 22 plmDCA Pairwise Parameters
i
j

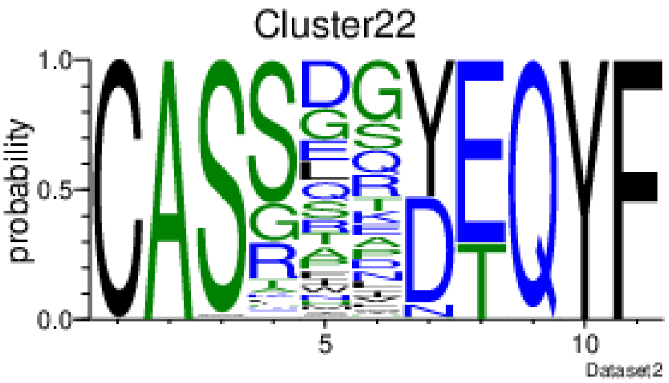
Cluster22
probability
Dataset2

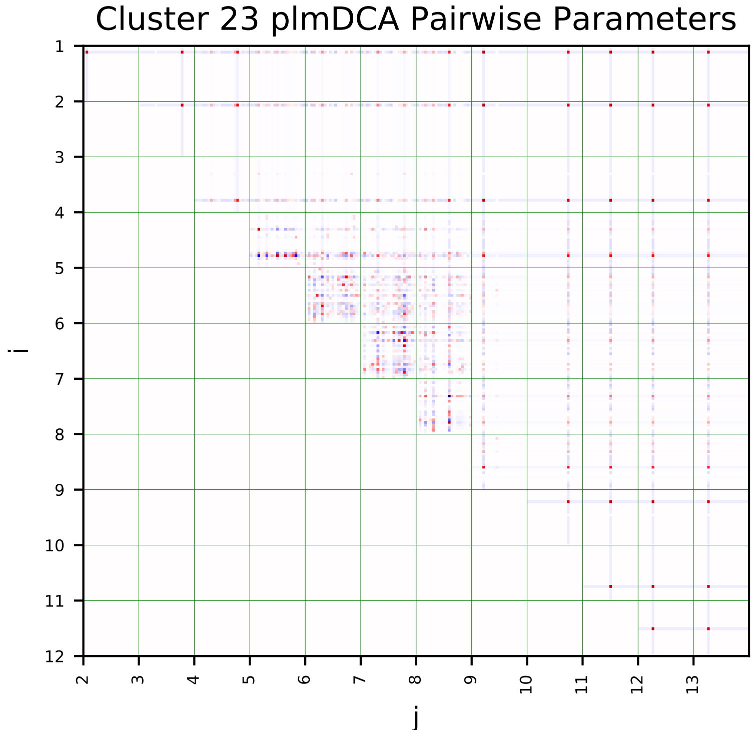
Cluster 23 plmDCA Pairwise Parameters
i
j

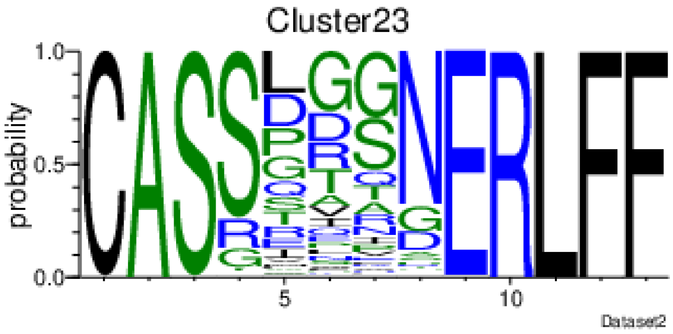
Cluster23
probability
Dataset2

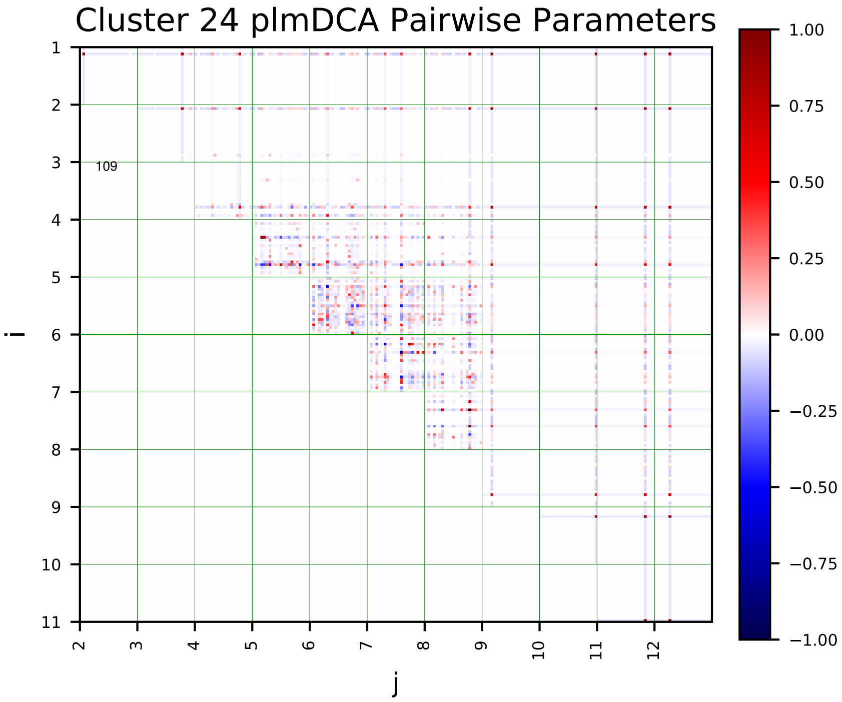
Cluster 24 plmDCA Pairwise Parameters
109
i
j

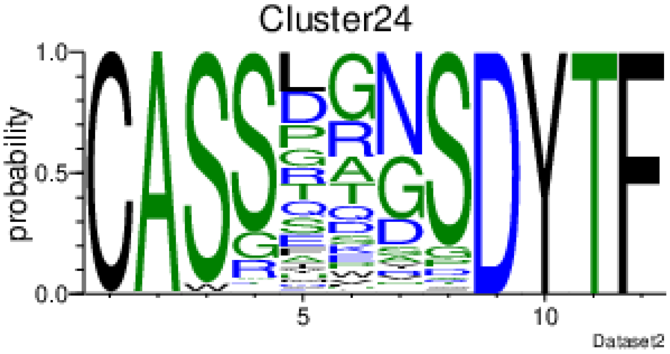
Cluster24
probability
Dataset2

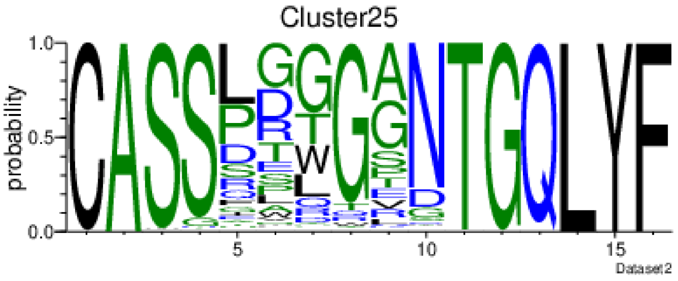
Cluster25
probability
Dataset2

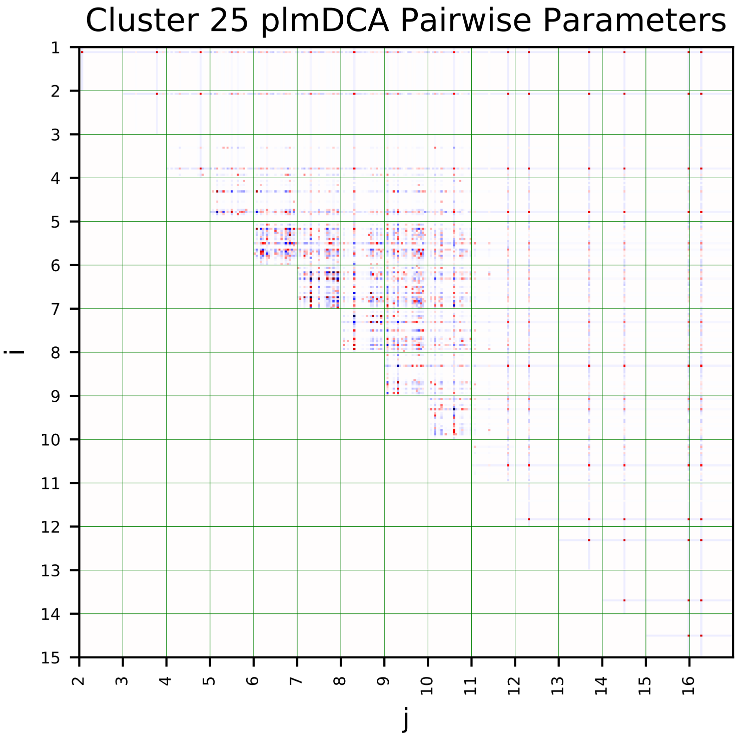
Cluster 25 plmDCA Pairwise Parameters
i
j

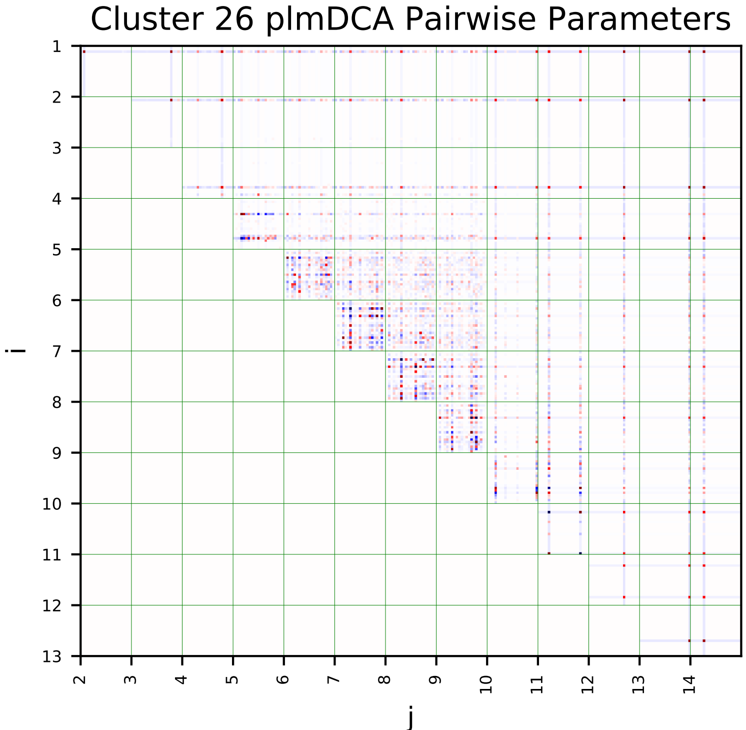
Cluster 26 plmDCA Pairwise Parameters
i
j

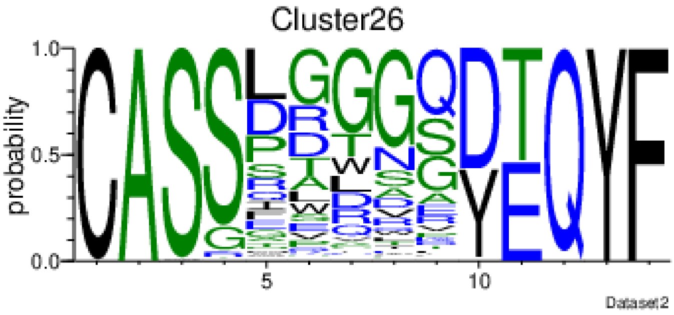
Cluster26
probability
Dataset2

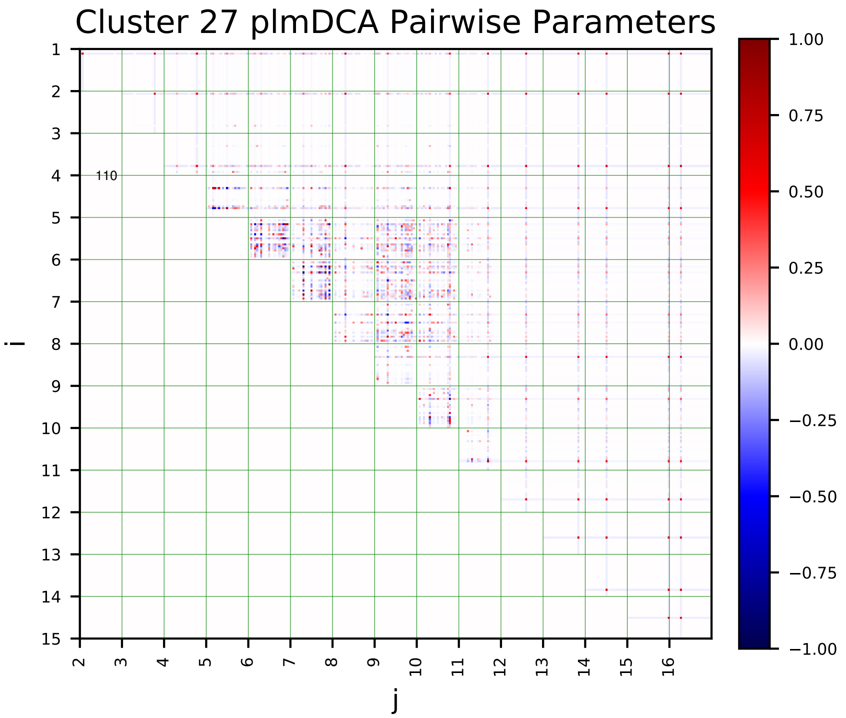
Cluster 27 plmDCA Pairwise Parameters
i
j
110

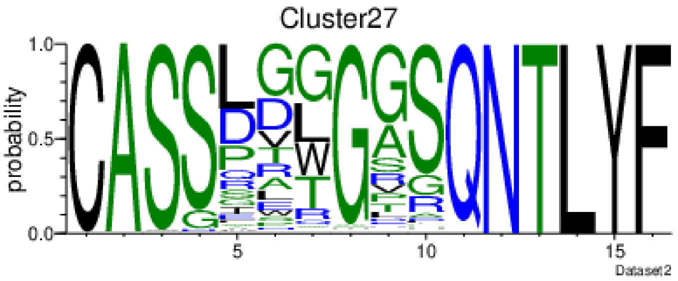
Cluster27
probability
Dataset2

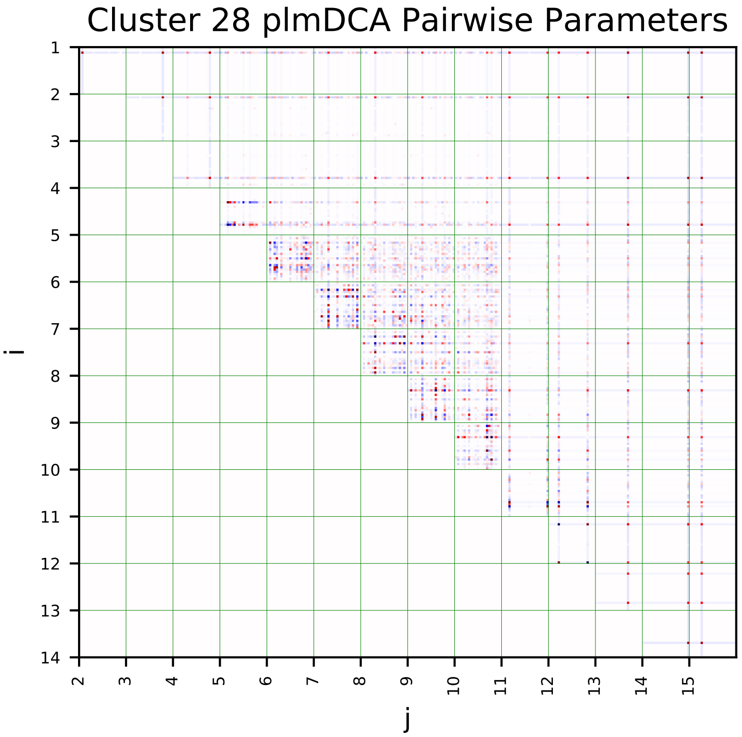
Cluster 28 plmDCA Pairwise Parameters
i
j

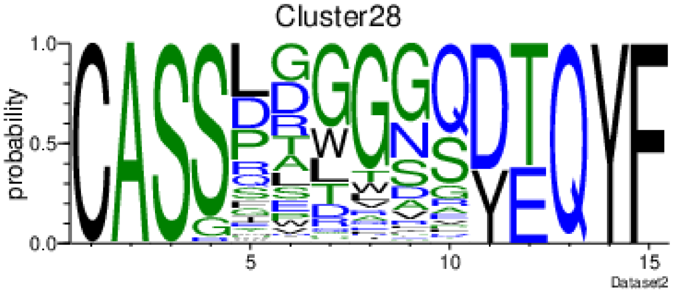
Cluster28
probability
Dataset2

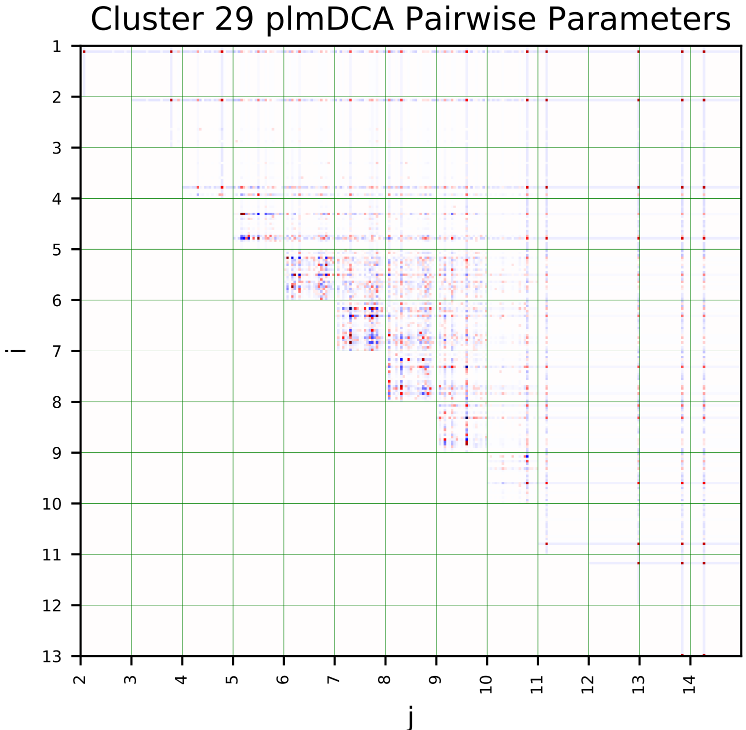
Cluster 29 plmDCA Pairwise Parameters
i
j

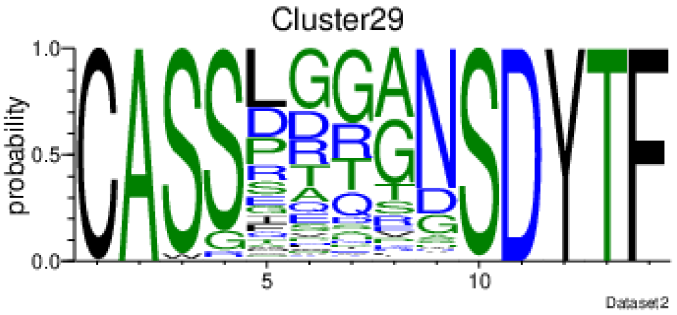
Cluster29
probability
Dataset2

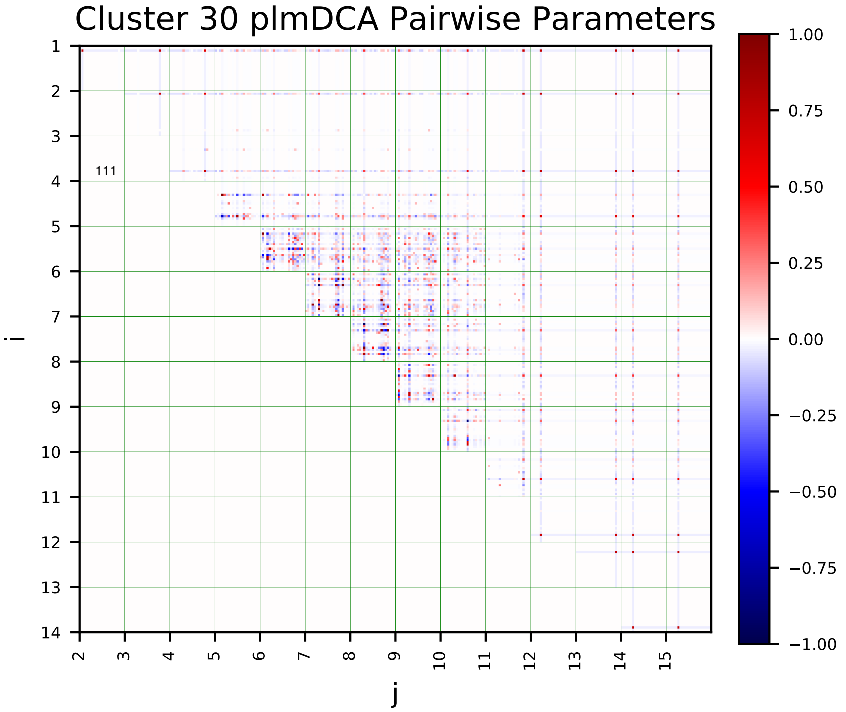
Cluster 30 plmDCA Pairwise Parameters
i
j

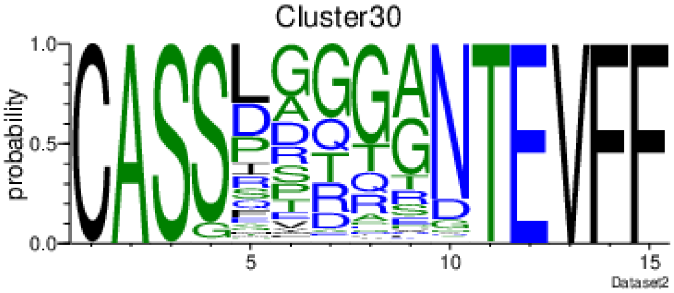
Cluster30
probability
Dataset2

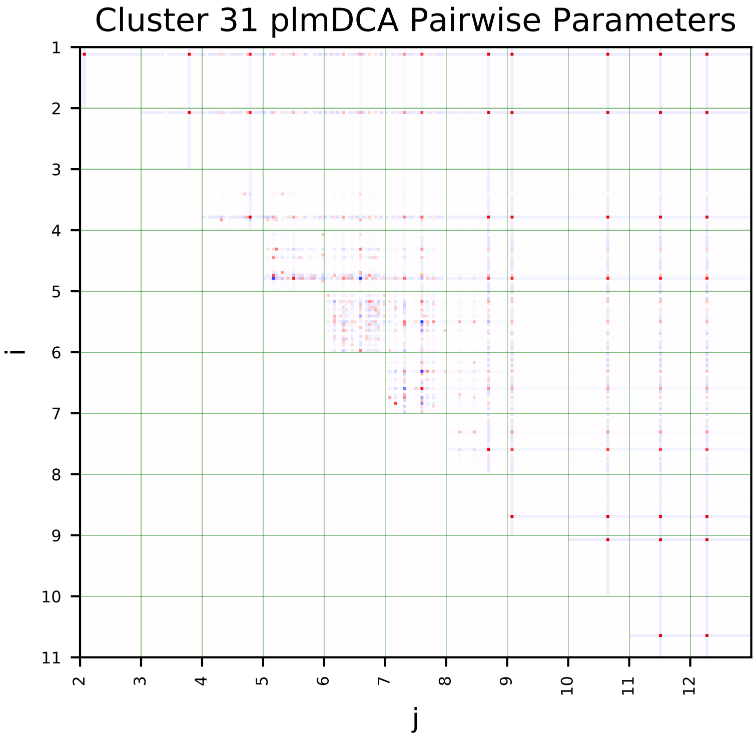
Cluster 31 plmDCA Pairwise Parameters
i
j

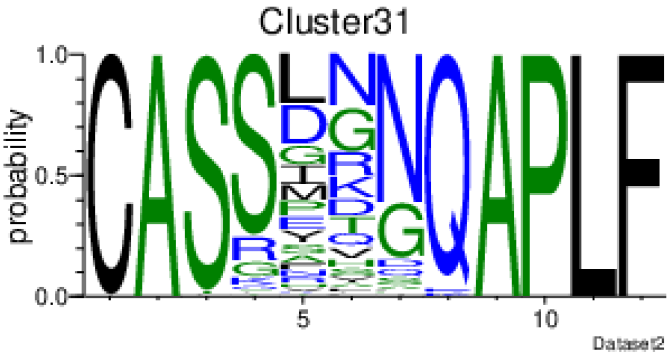
Cluster31
probability
Dataset2

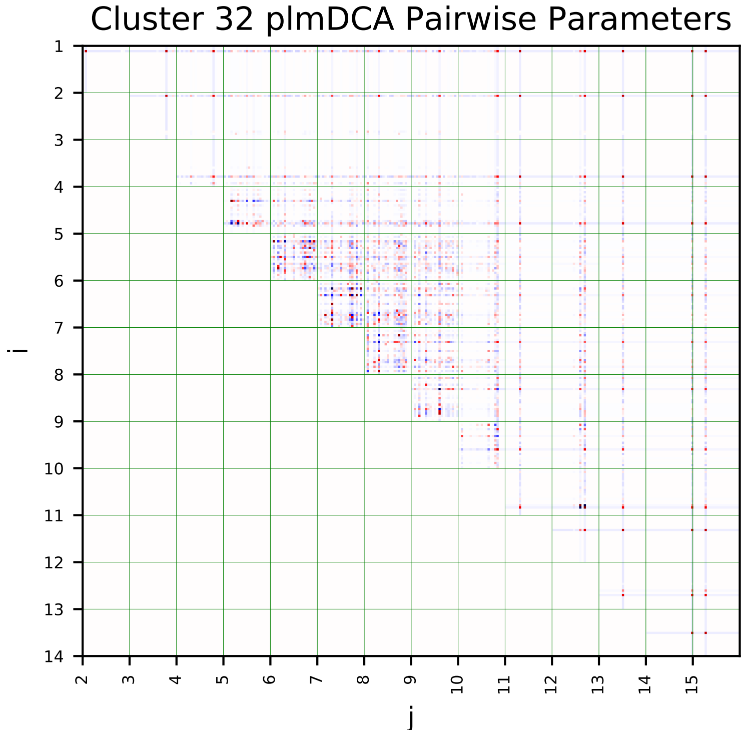
Cluster 32 plmDCA Pairwise Parameters
i
j

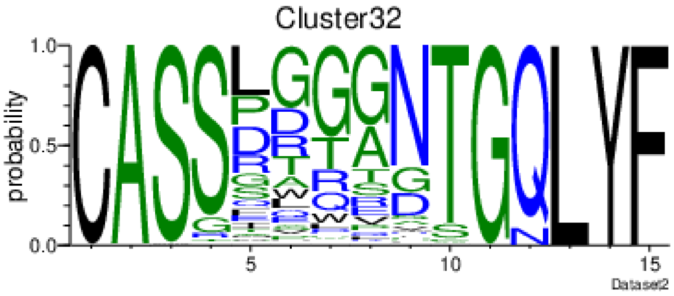
Cluster32
probability
Dataset2

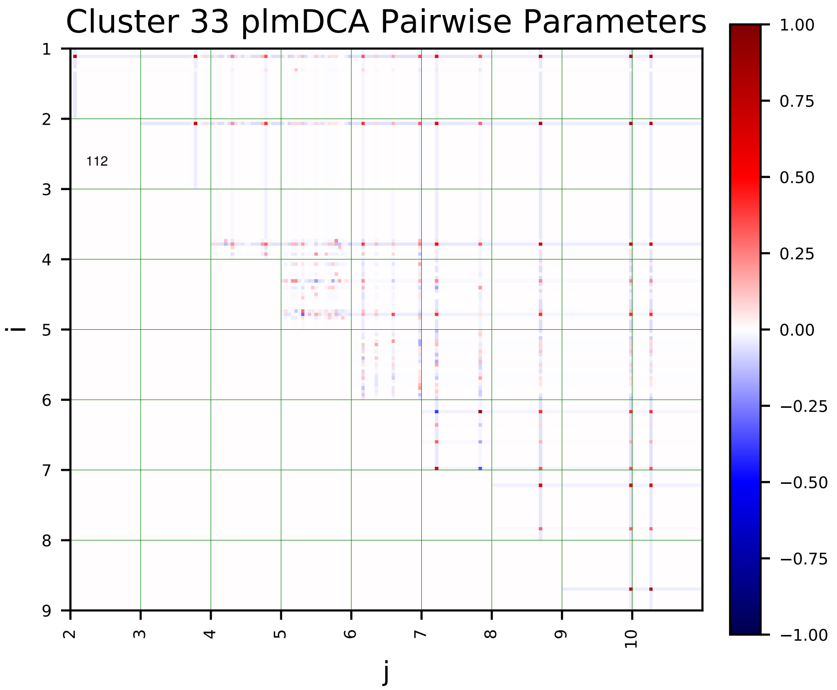
Cluster 33 plmDCA Pairwise Parameters
112
i
j
1.00
0.75
0.50
0.25
0.00
−0.25
−0.50
−0.75
−1.00

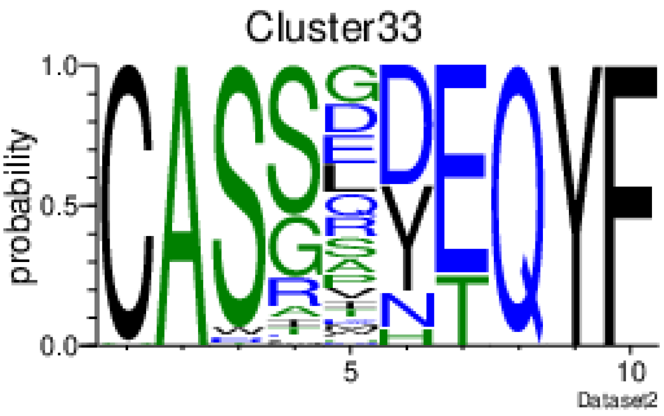
Cluster33
probability
Dataset2

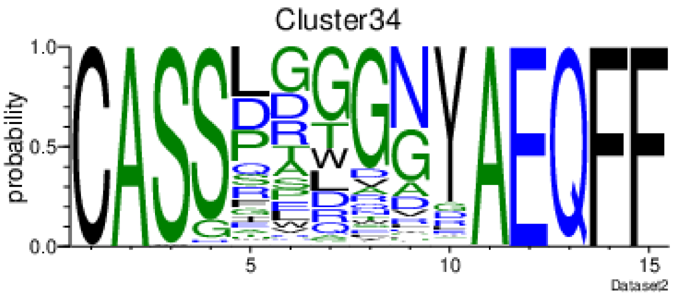
Cluster34
probability
Dataset2

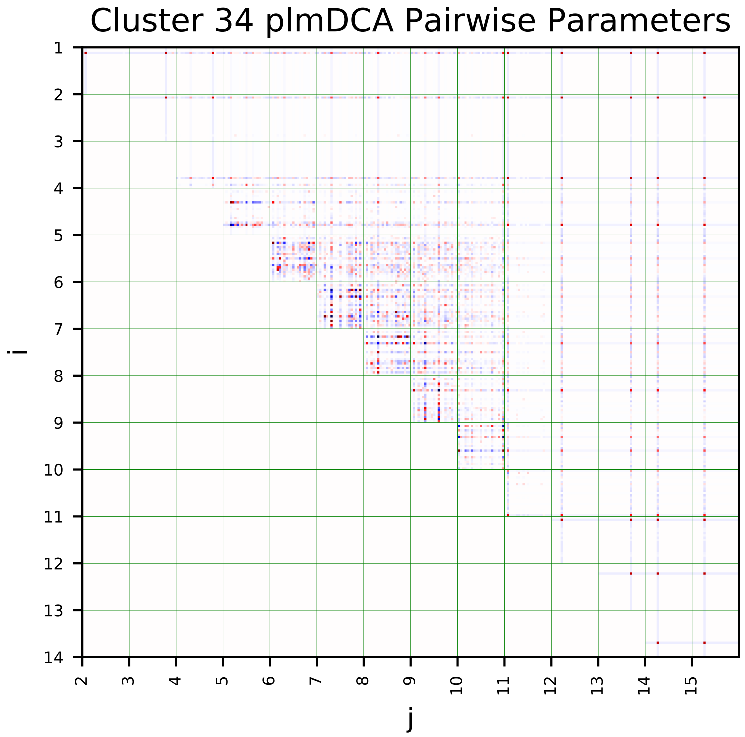
Cluster 34 plmDCA Pairwise Parameters
i
j

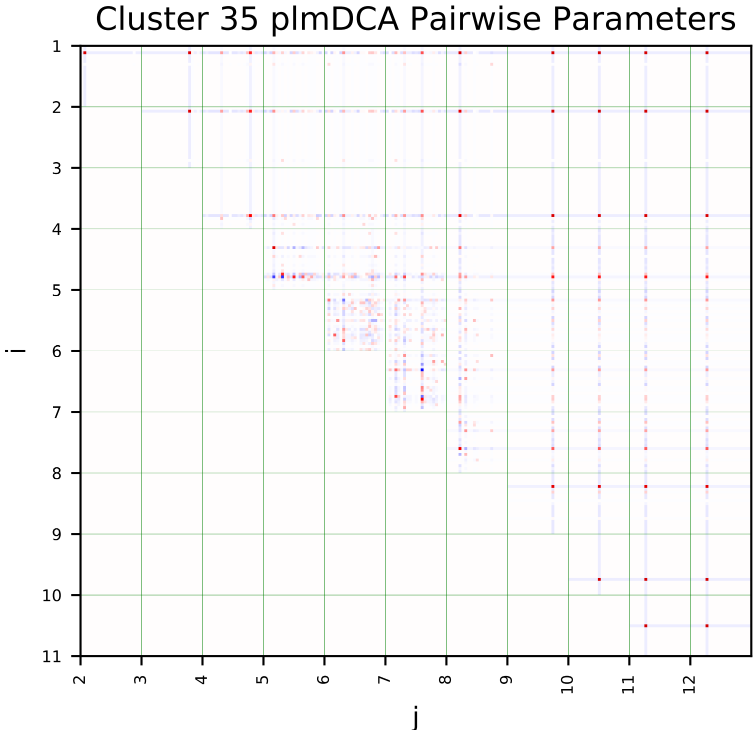
Cluster 35 plmDCA Pairwise Parameters
i
j

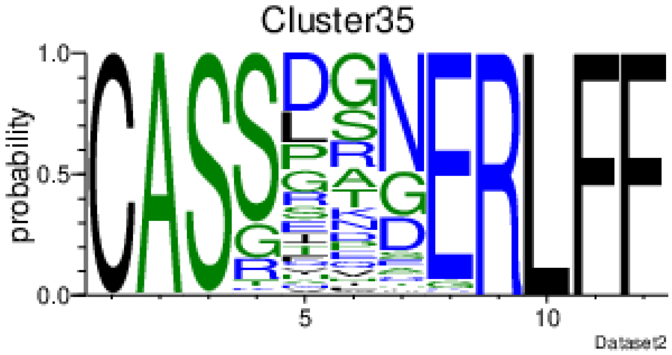
Cluster35
probability
Dataset2

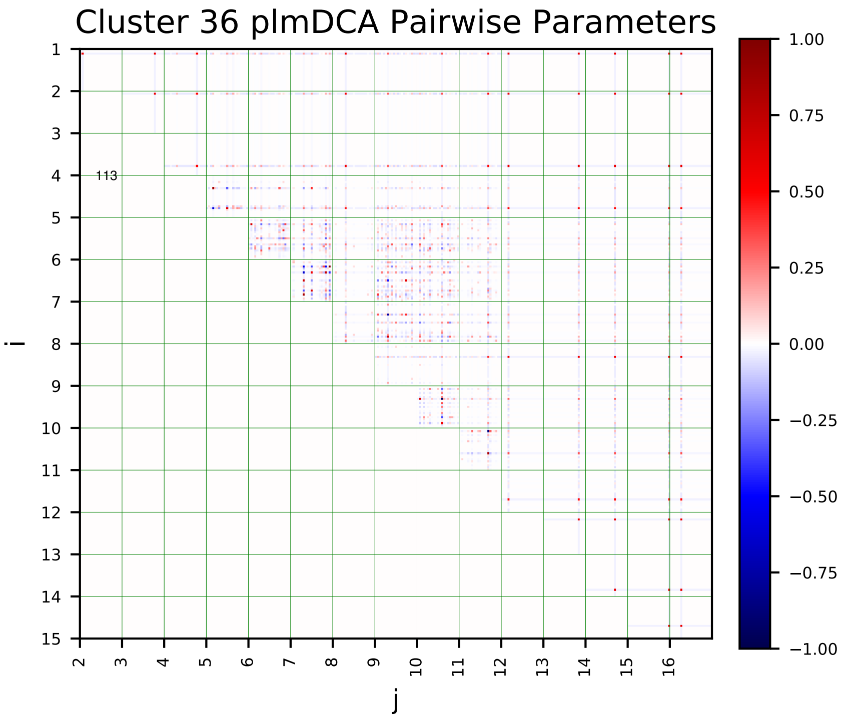
Cluster 36 plmDCA Pairwise Parameters
i
j
1.00
0.75
0.50
0.25
0.00
−0.25
−0.50
−0.75
−1.00

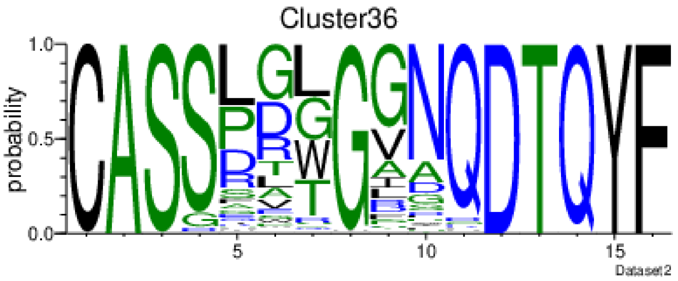
Cluster36
probability
Dataset2

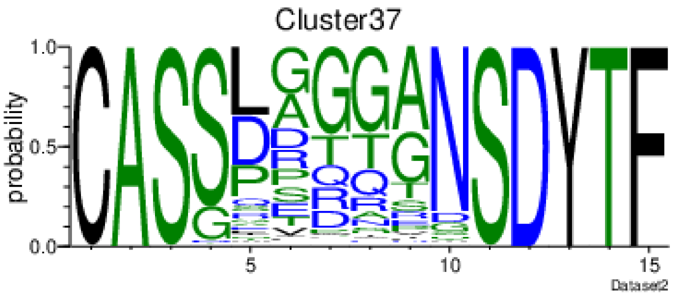
Cluster37
probability
Dataset2

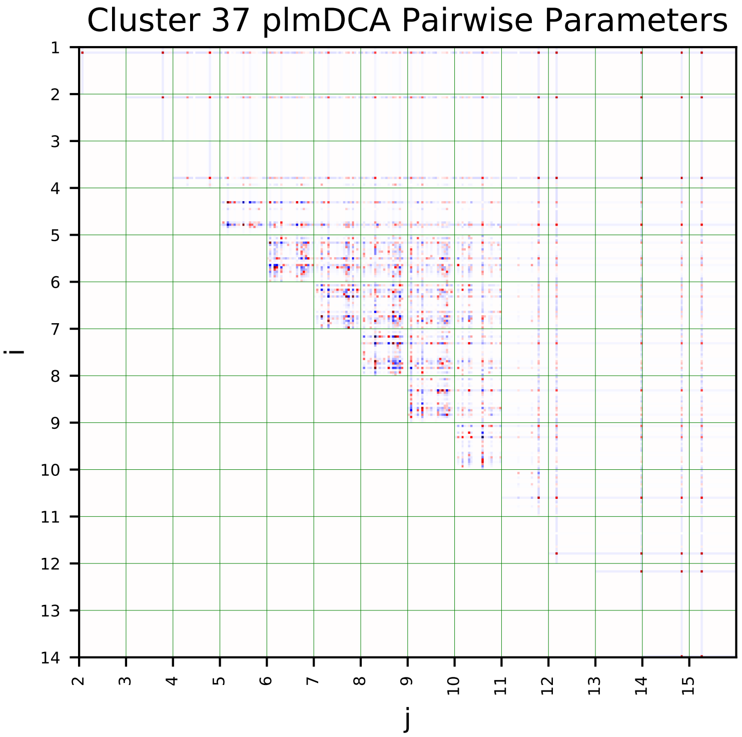
Cluster 37 plmDCA Pairwise Parameters
i
j

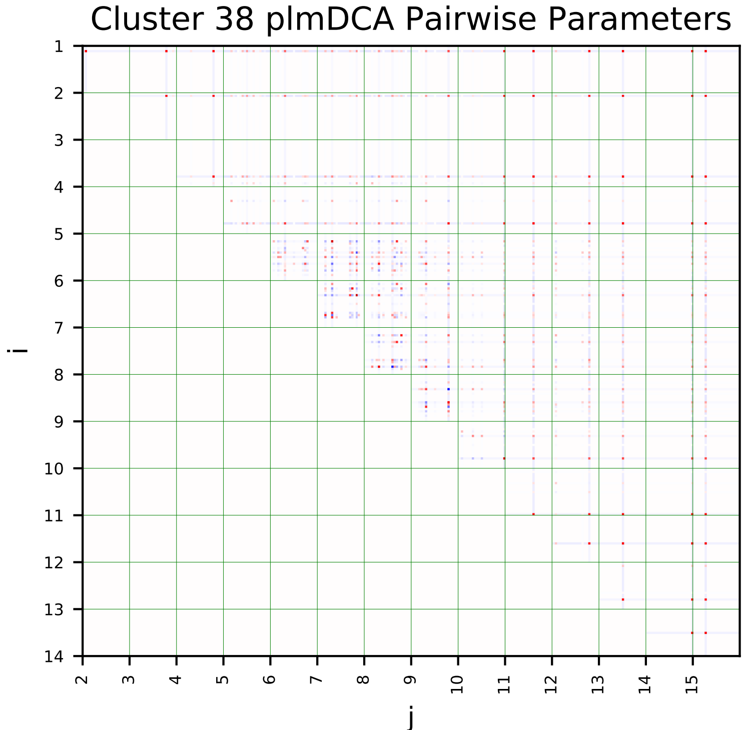
Cluster 38 plmDCA Pairwise Parameters
i
j

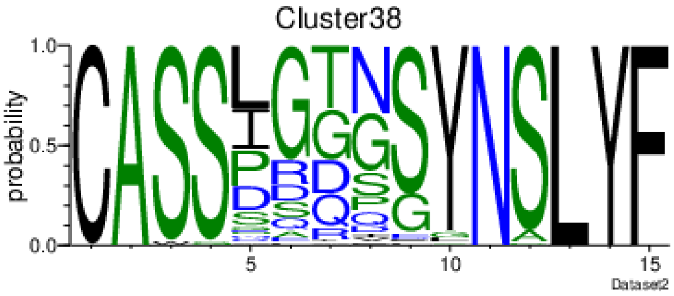
Cluster38
probability
Dataset2

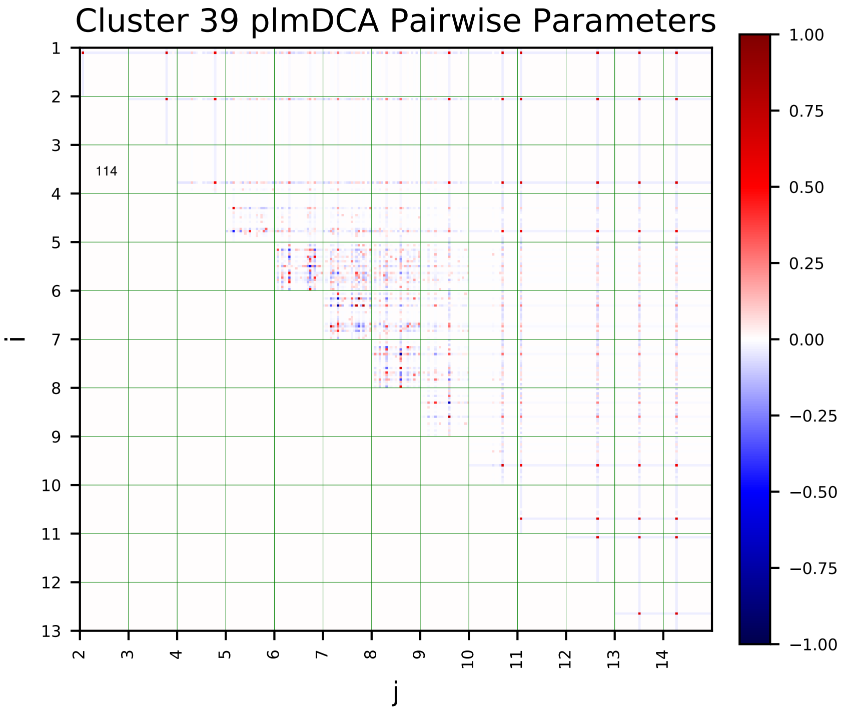
Cluster 39 plmDCA Pairwise Parameters
i
j
114

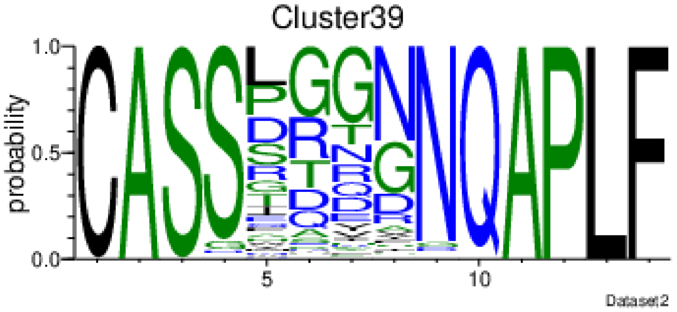
Cluster39
probability
Dataset2

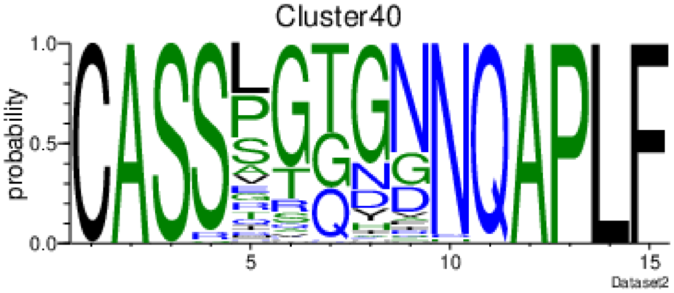
Cluster40
probability
Dataset2

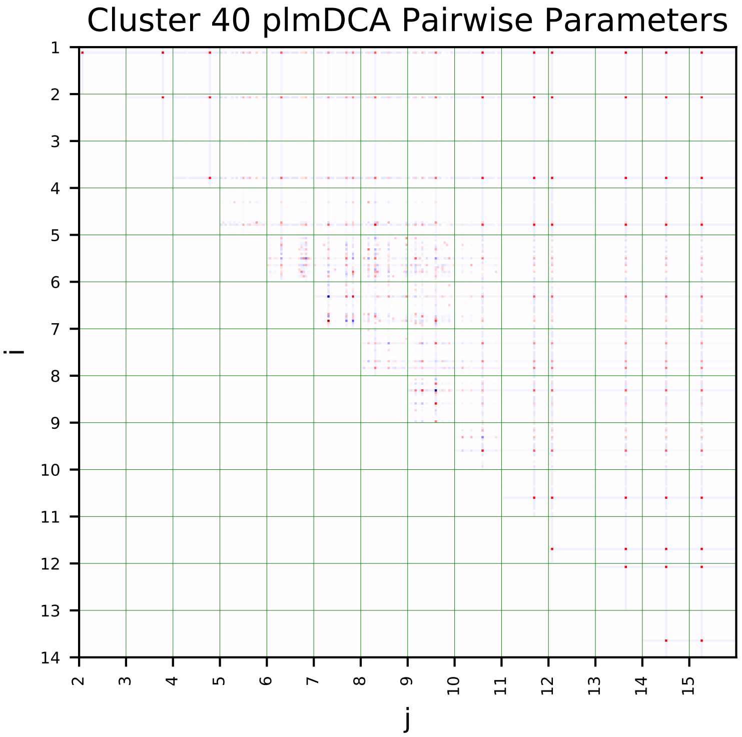
Cluster 40 plmDCA Pairwise Parameters
i
j

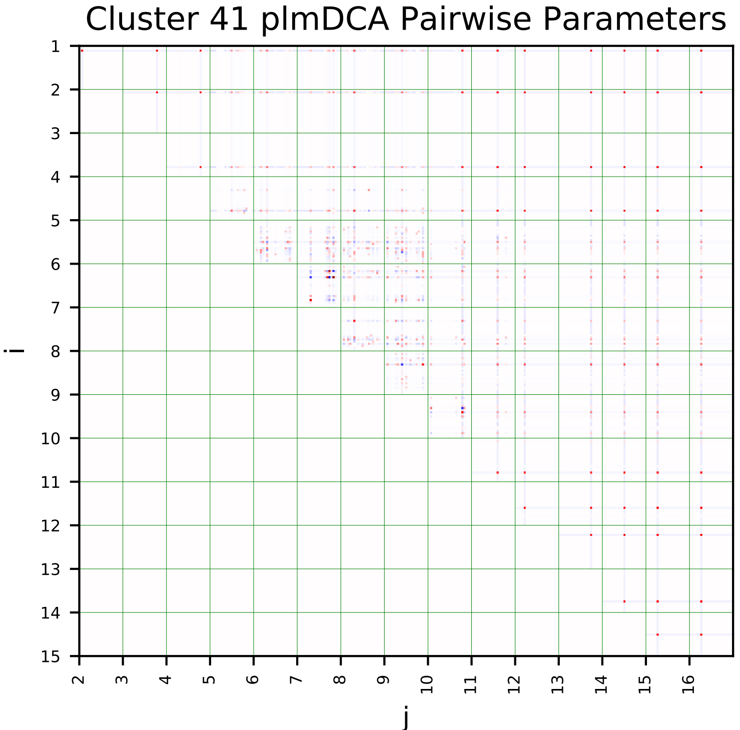
Cluster 41 plmDCA Pairwise Parameters
i
j

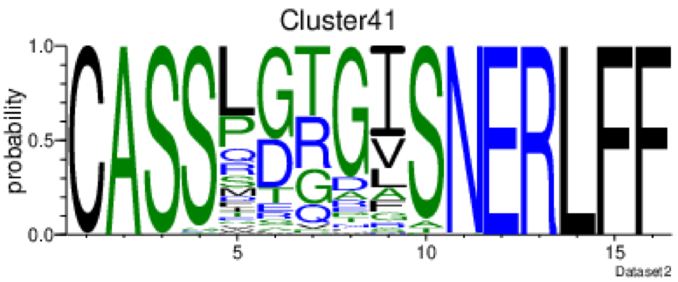
Cluster41
probability
Dataset2

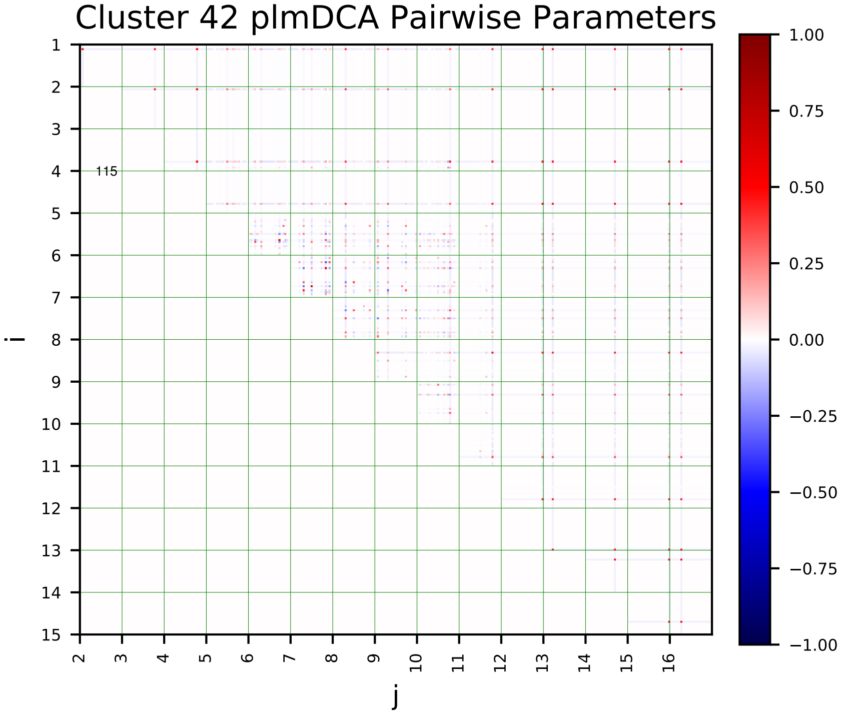
Cluster 42 plmDCA Pairwise Parameters
i
j
115
1.00
0.75
0.50
0.25
0.00
−0.25
−0.50
−0.75
−1.00

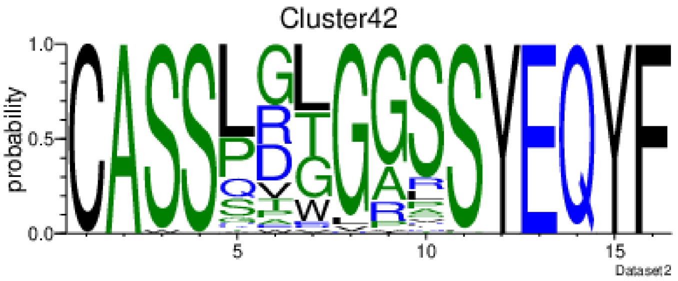
Cluster42
probability
Dataset2

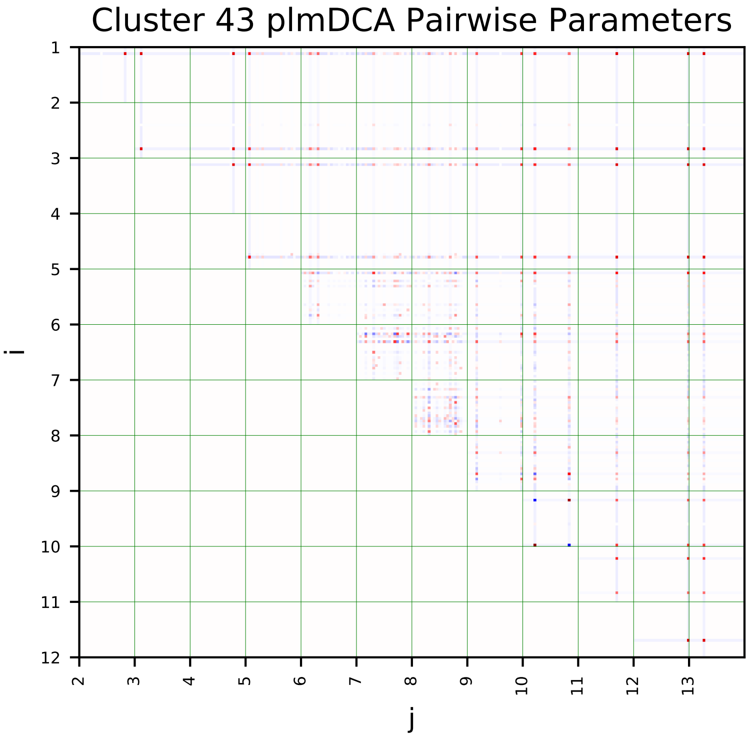
Cluster 43 plmDCA Pairwise Parameters
i
j

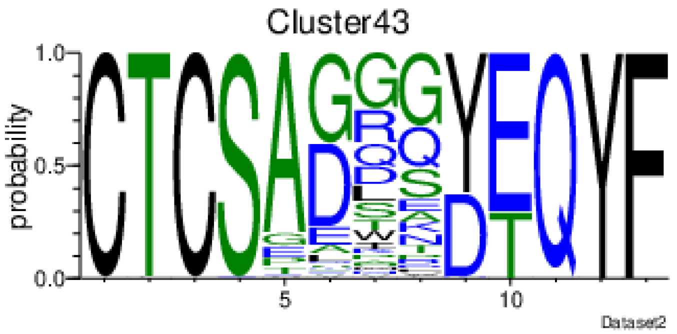
Cluster43
probability
Dataset2

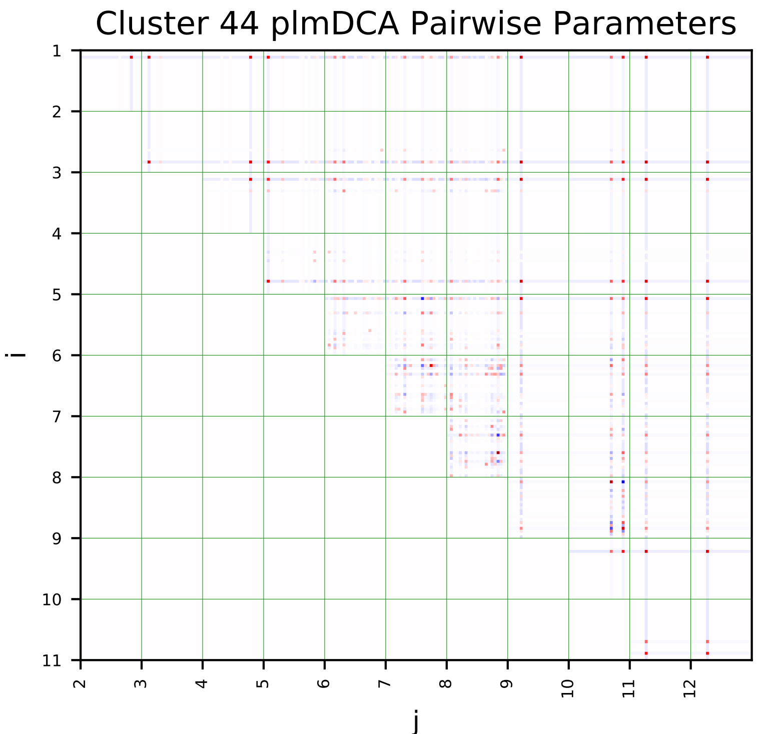
Cluster 44 plmDCA Pairwise Parameters
i
j

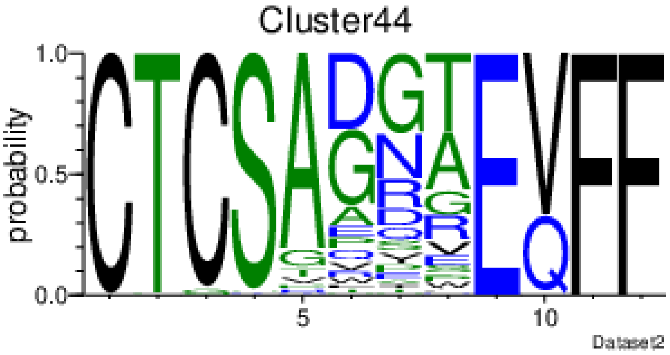
Cluster44
probability
Dataset2

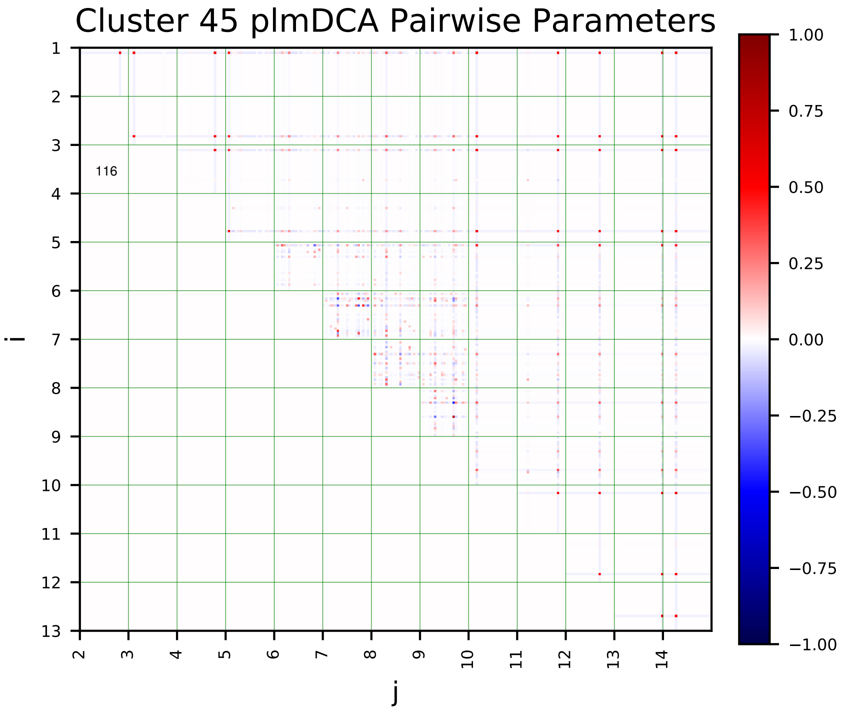
Cluster 45 plmDCA Pairwise Parameters
i
j
116
1.00
0.75
0.50
0.25
0.00
−0.25
−0.50
−0.75
−1.00

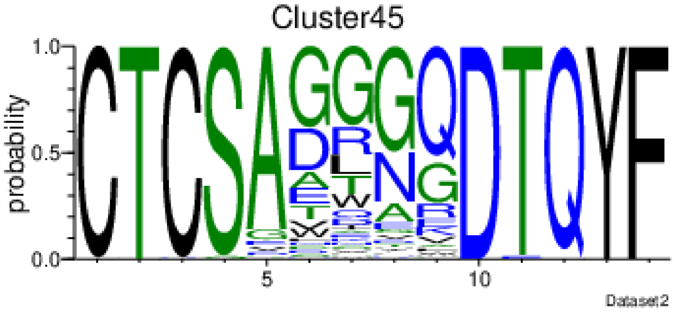
Cluster45
probability
Dataset2

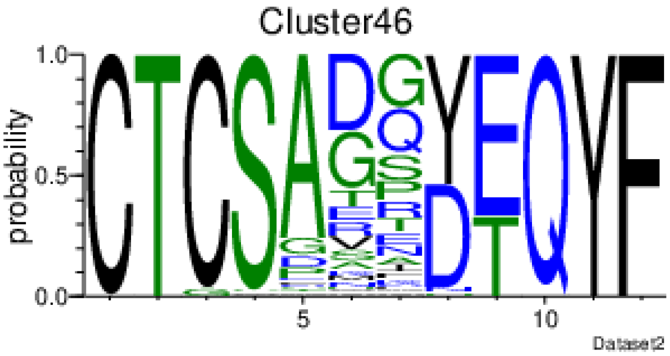
Cluster46
probability
Dataset2

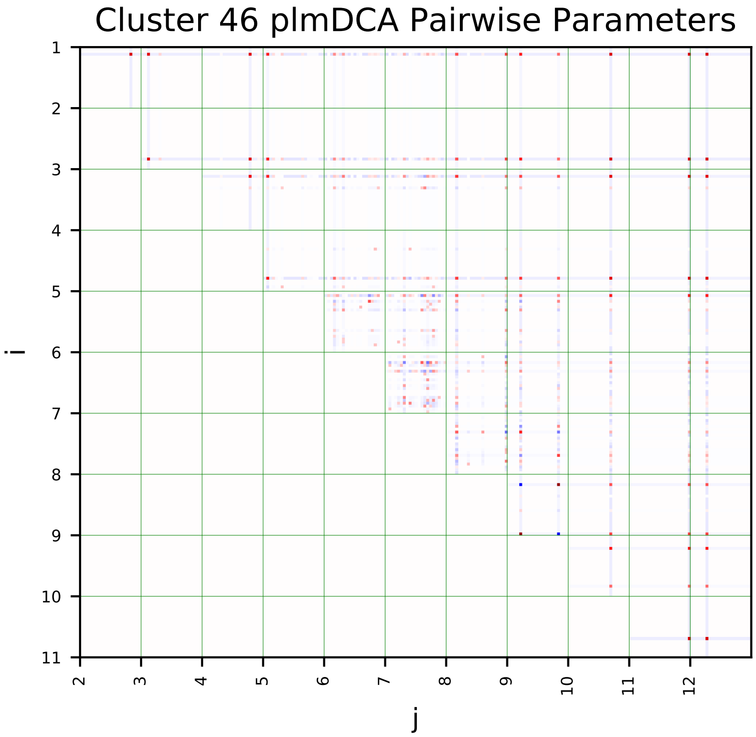
Cluster 46 plmDCA Pairwise Parameters
i
j

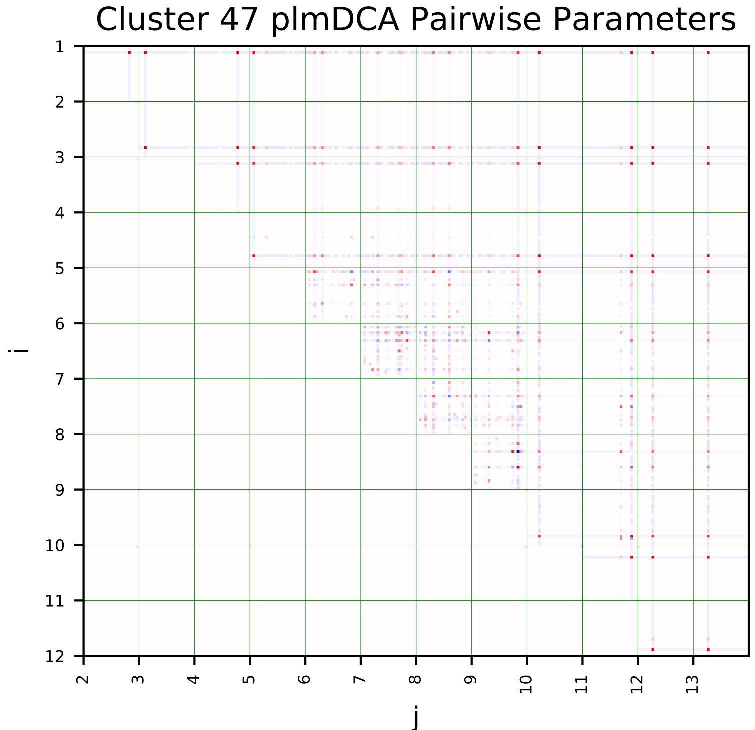
Cluster 47 plmDCA Pairwise Parameters
i
j

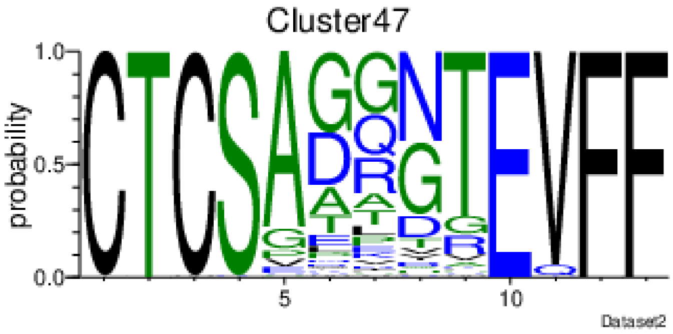
Cluster47
probability
Dataset2

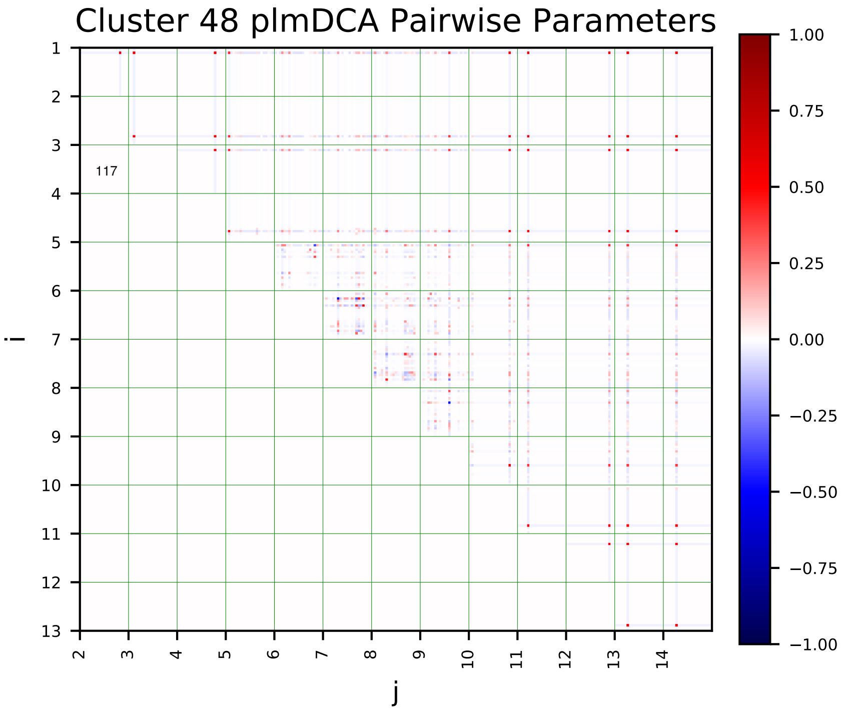
Cluster 48 plmDCA Pairwise Parameters
i
j

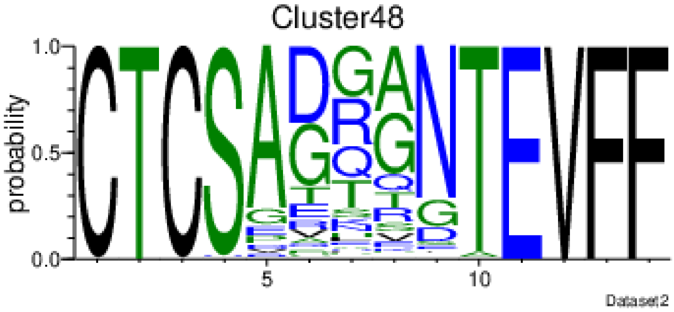
Cluster48
probability
Dataset2

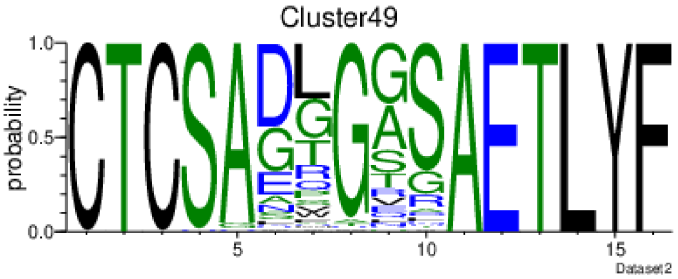
Cluster49
probability
Dataset2

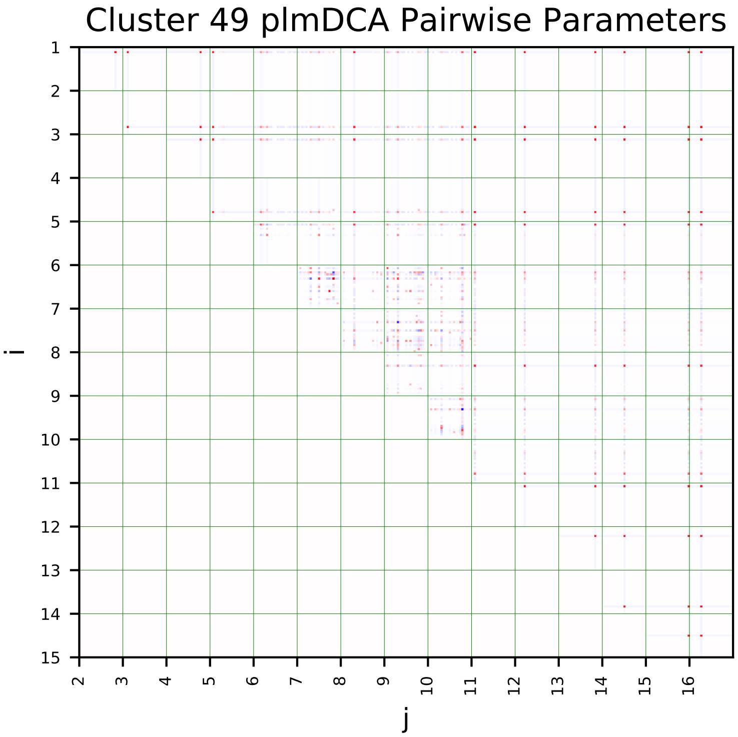
Cluster 49 plmDCA Pairwise Parameters
i
j

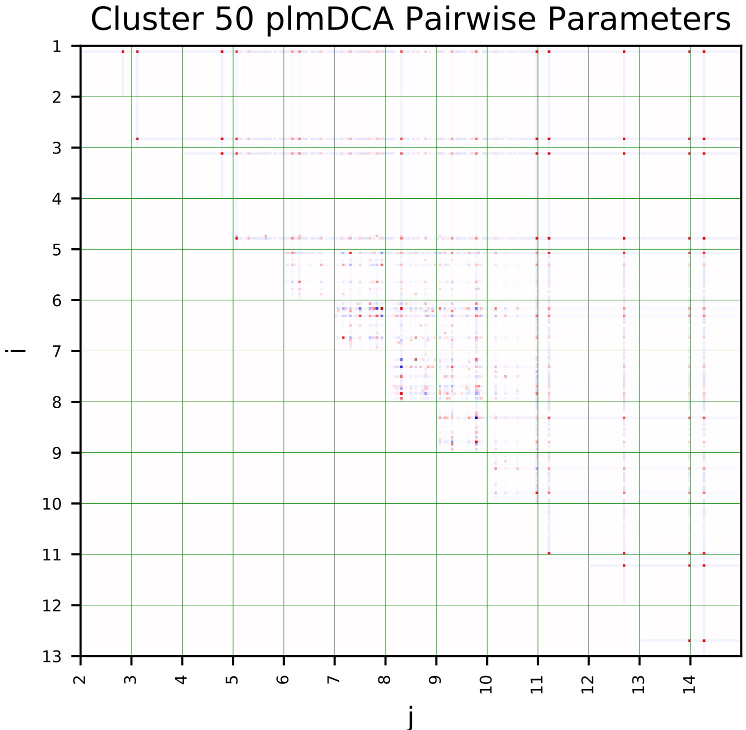
Cluster 50 plmDCA Pairwise Parameters
i
j

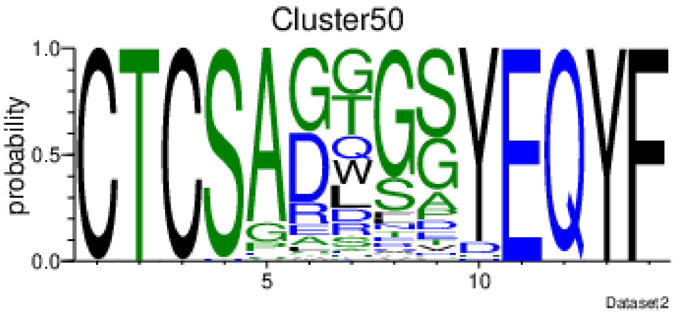
Cluster50
probability
Dataset2

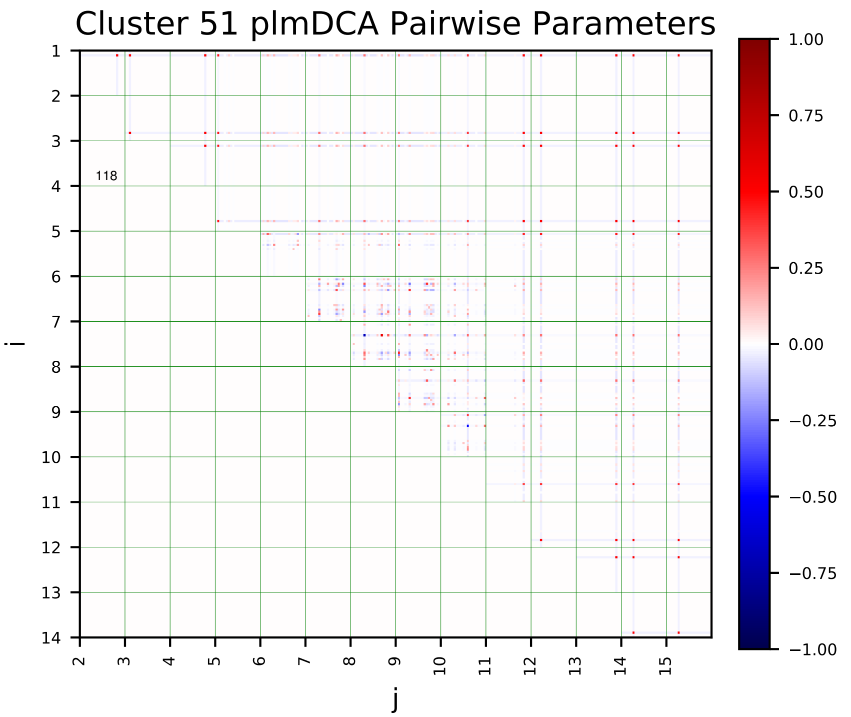
Cluster 51 plmDCA Pairwise Parameters
118
i
j

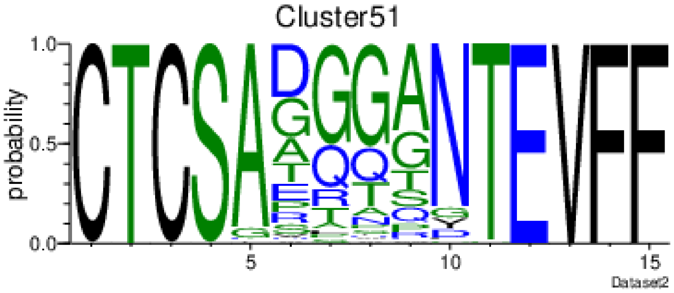
Cluster51
probability
Dataset2

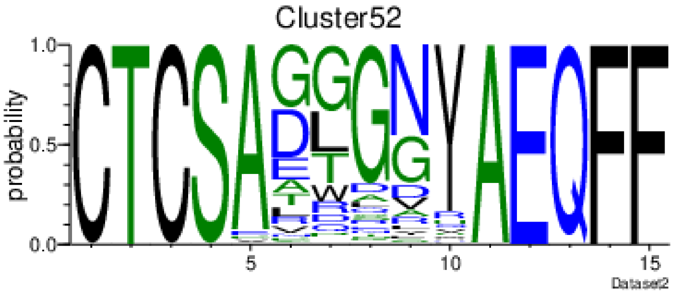
Cluster52
probability
Dataset2

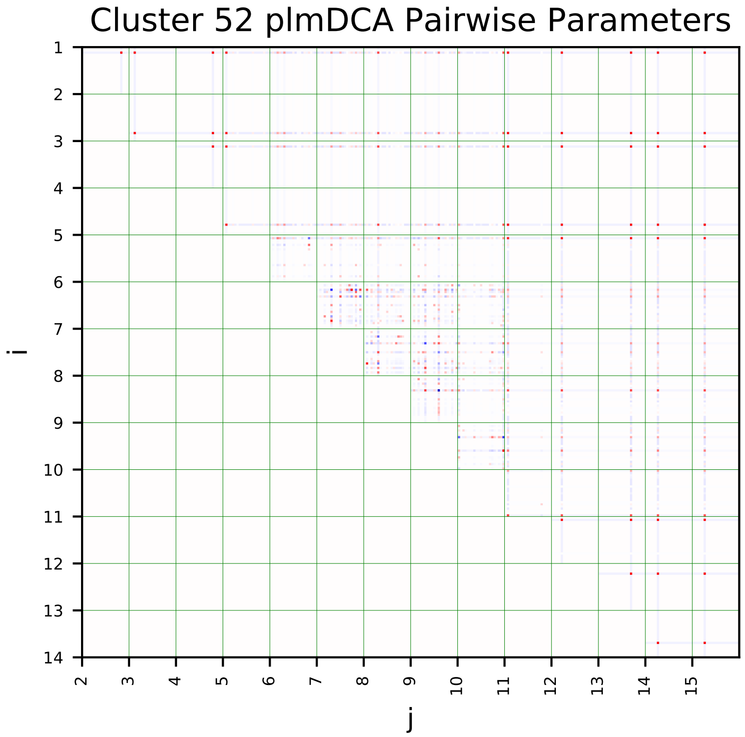
Cluster 52 plmDCA Pairwise Parameters
i
j

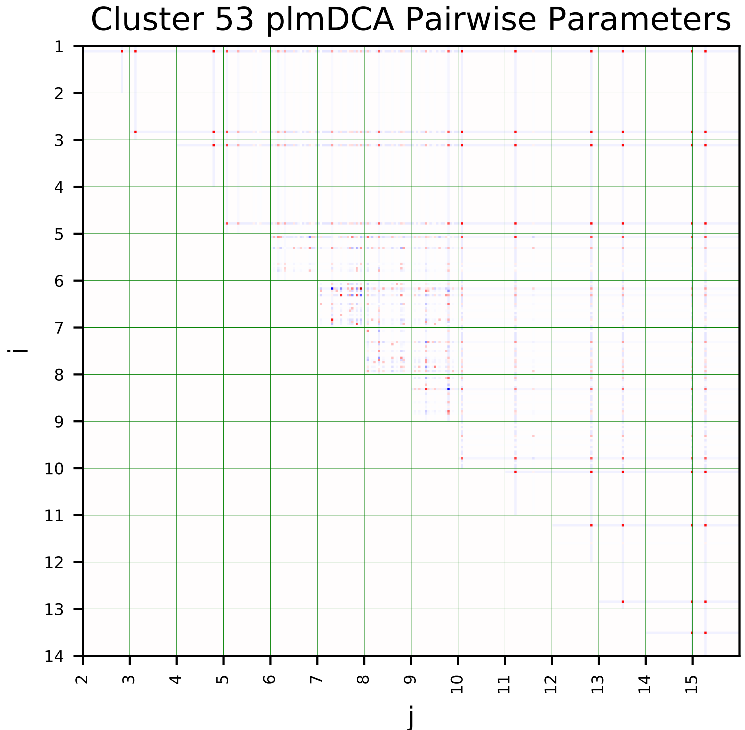
Cluster 53 plmDCA Pairwise Parameters
i
j

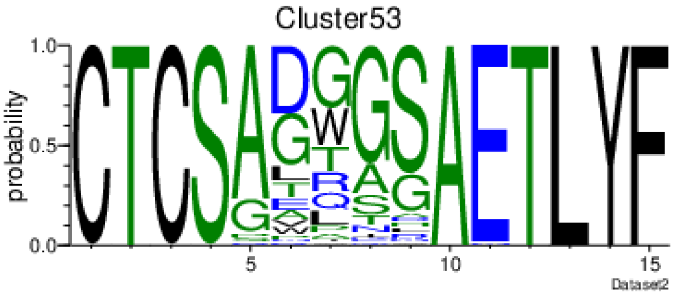
Cluster53
probability
Dataset2

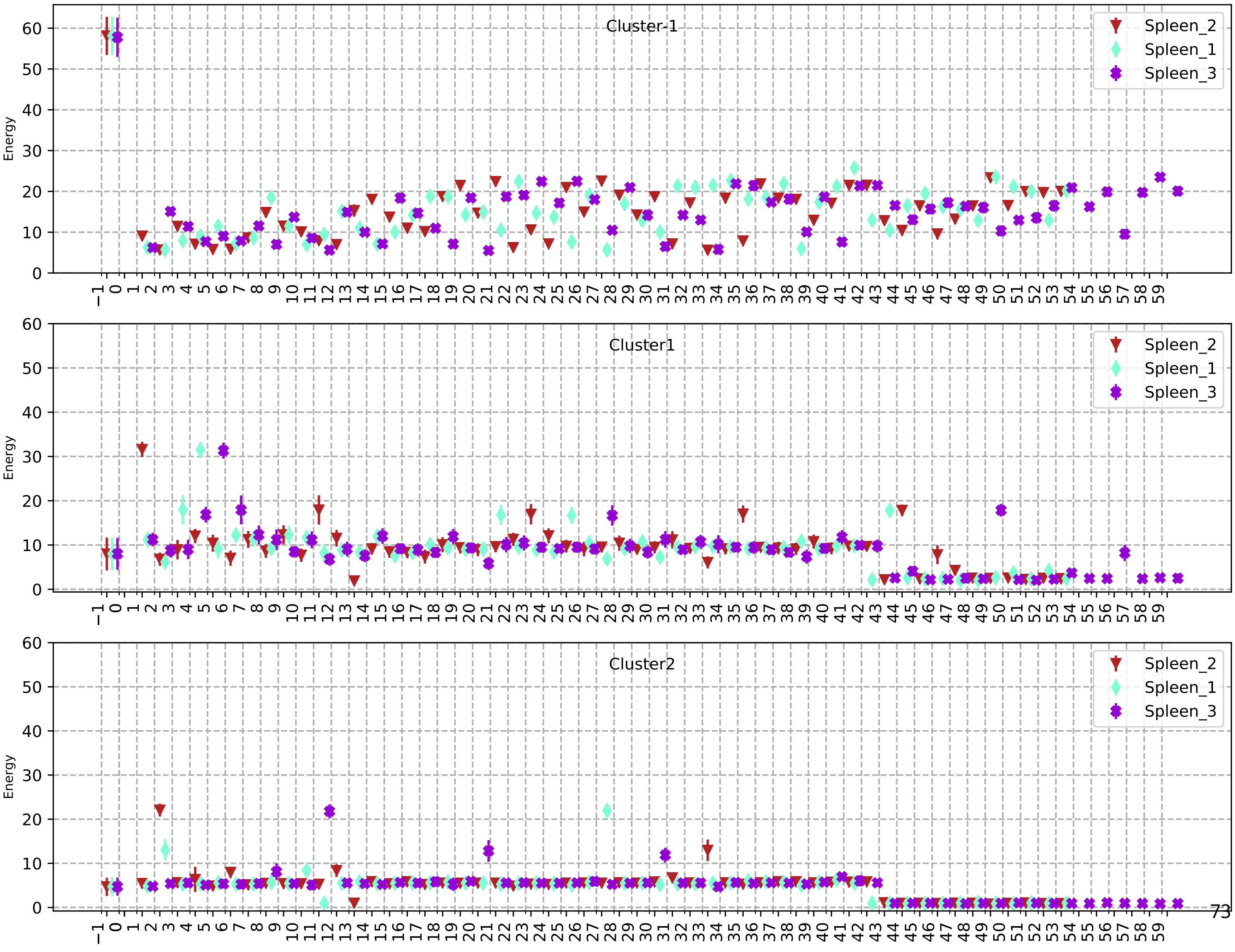
Cluster-1
Cluster1
Cluster2
Spleen_2
Spleen_1
Spleen_3
Energy

120

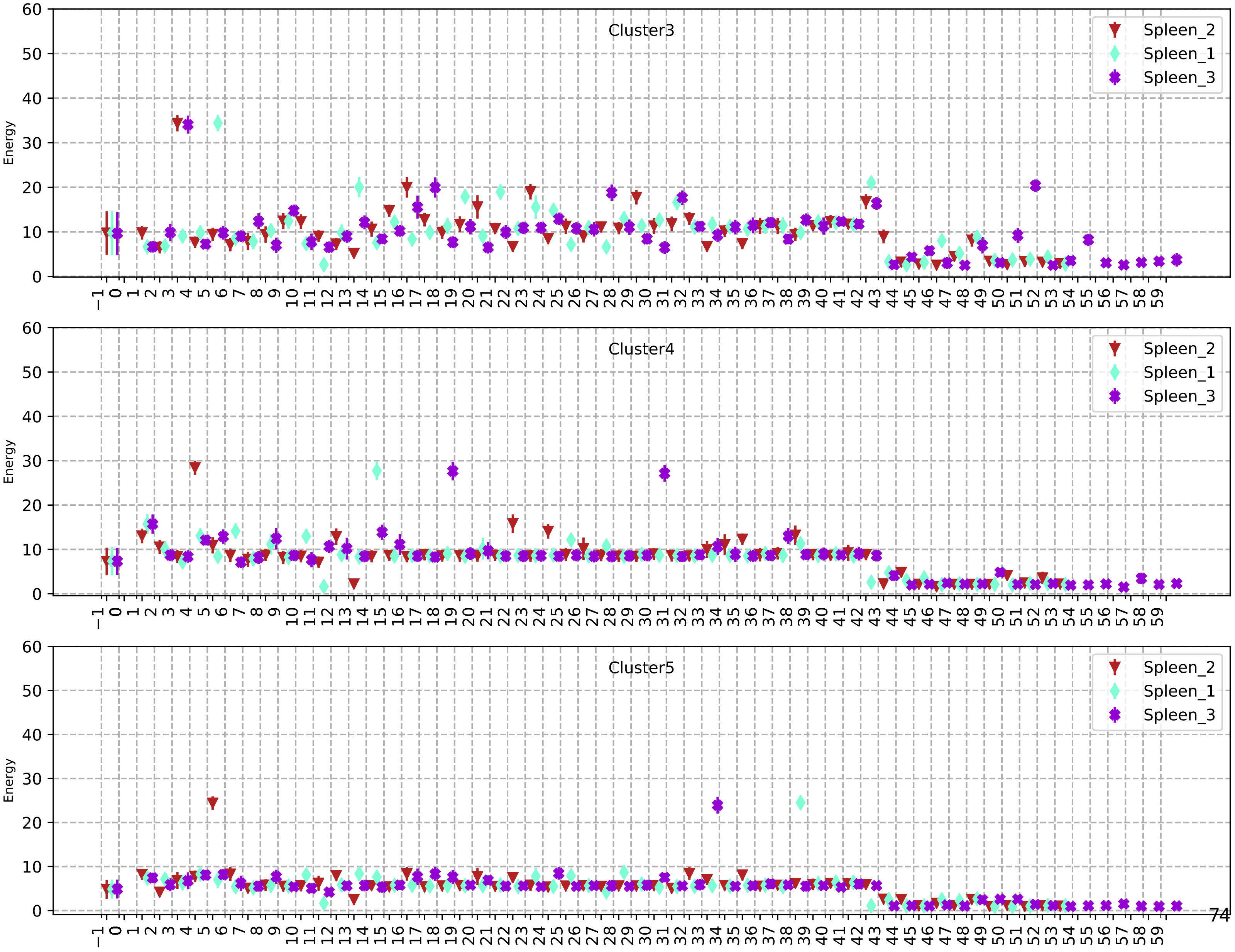

Cluster3
Cluster4
Cluster5
Spleen_2
Spleen_1
Spleen_3
Energy

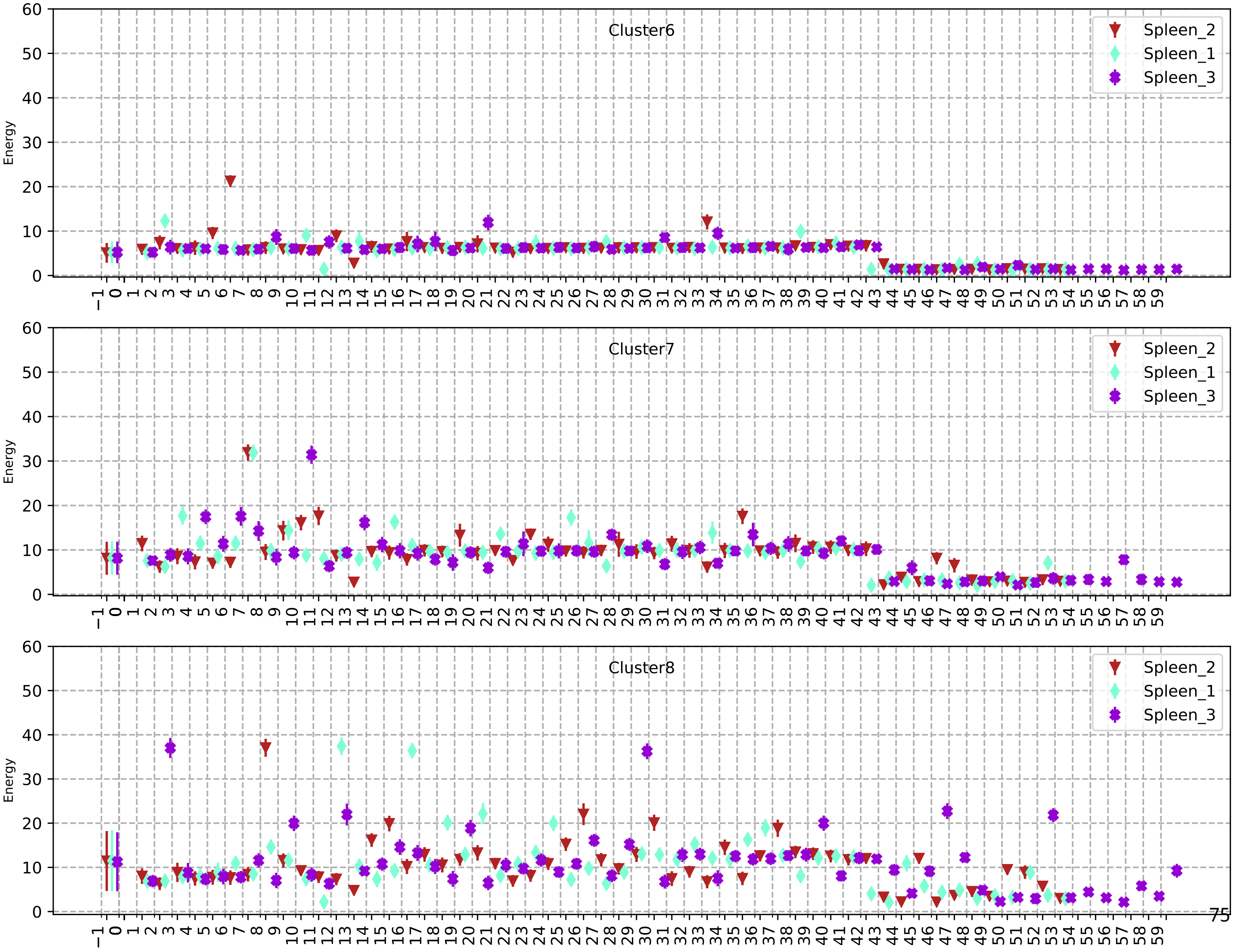
Cluster6
Spleen_2
Spleen_1
Spleen_3
Energy
Cluster7
Spleen_2
Spleen_1
Spleen_3
Energy
Cluster8
Spleen_2
Spleen_1
Spleen_3
Energy

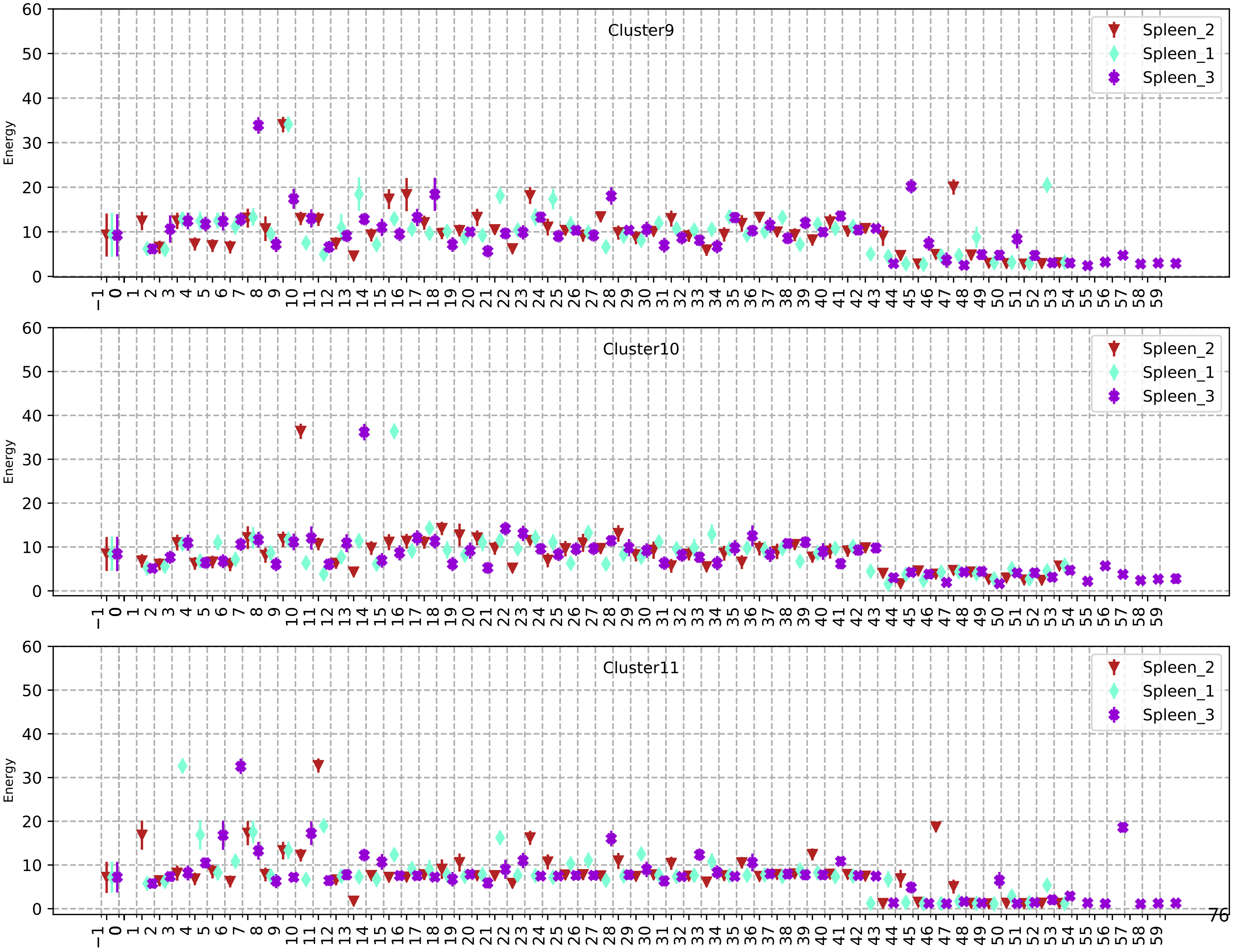
Cluster9
Spleen_2
Spleen_1
Spleen_3
Energy
Cluster10
Spleen_2
Spleen_1
Spleen_3
Energy
Cluster11
Spleen_2
Spleen_1
Spleen_3
Energy

123

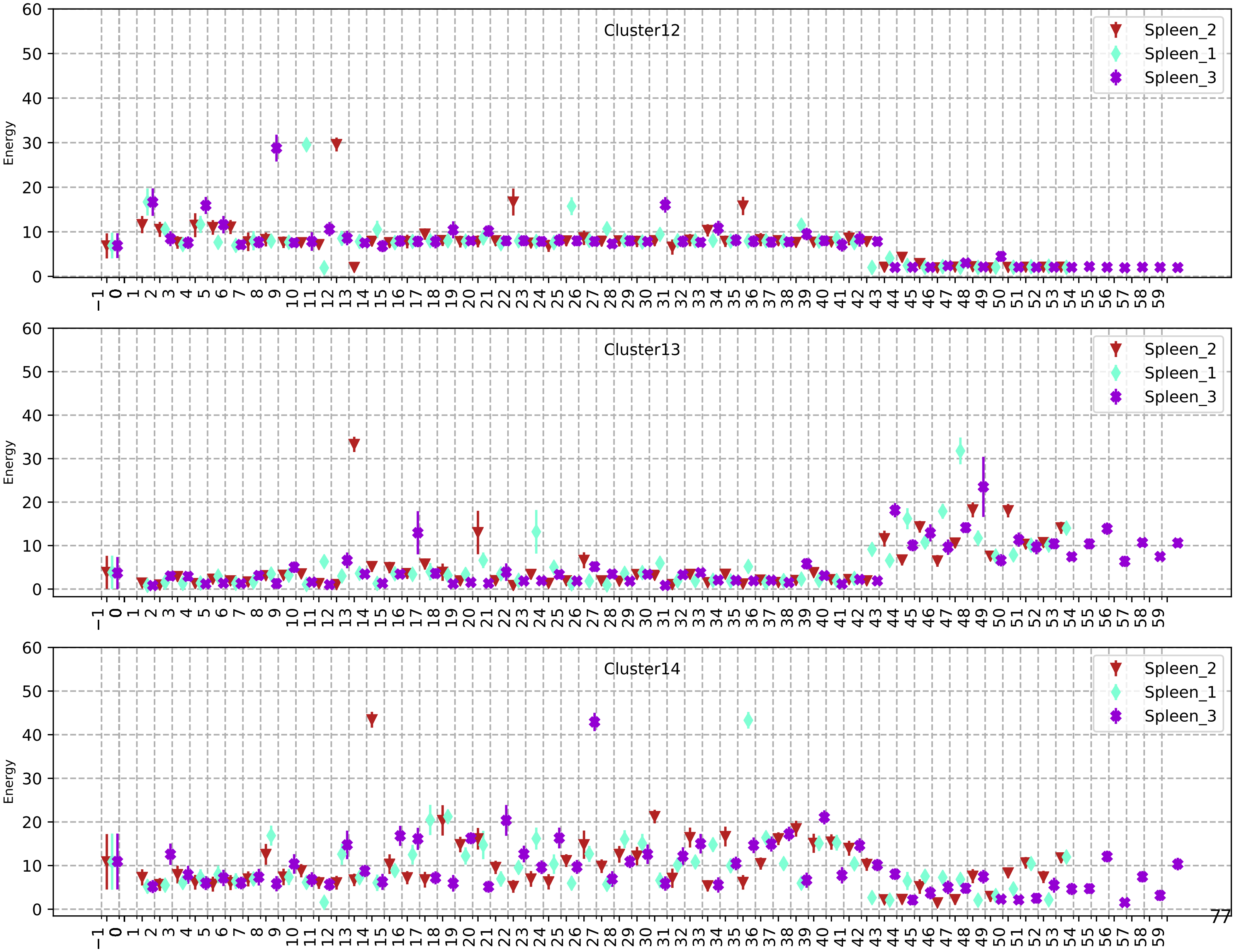
Cluster12
Cluster13
Cluster14
Spleen_2
Spleen_1
Spleen_3
Energy
0
10
20
30
40
50
60

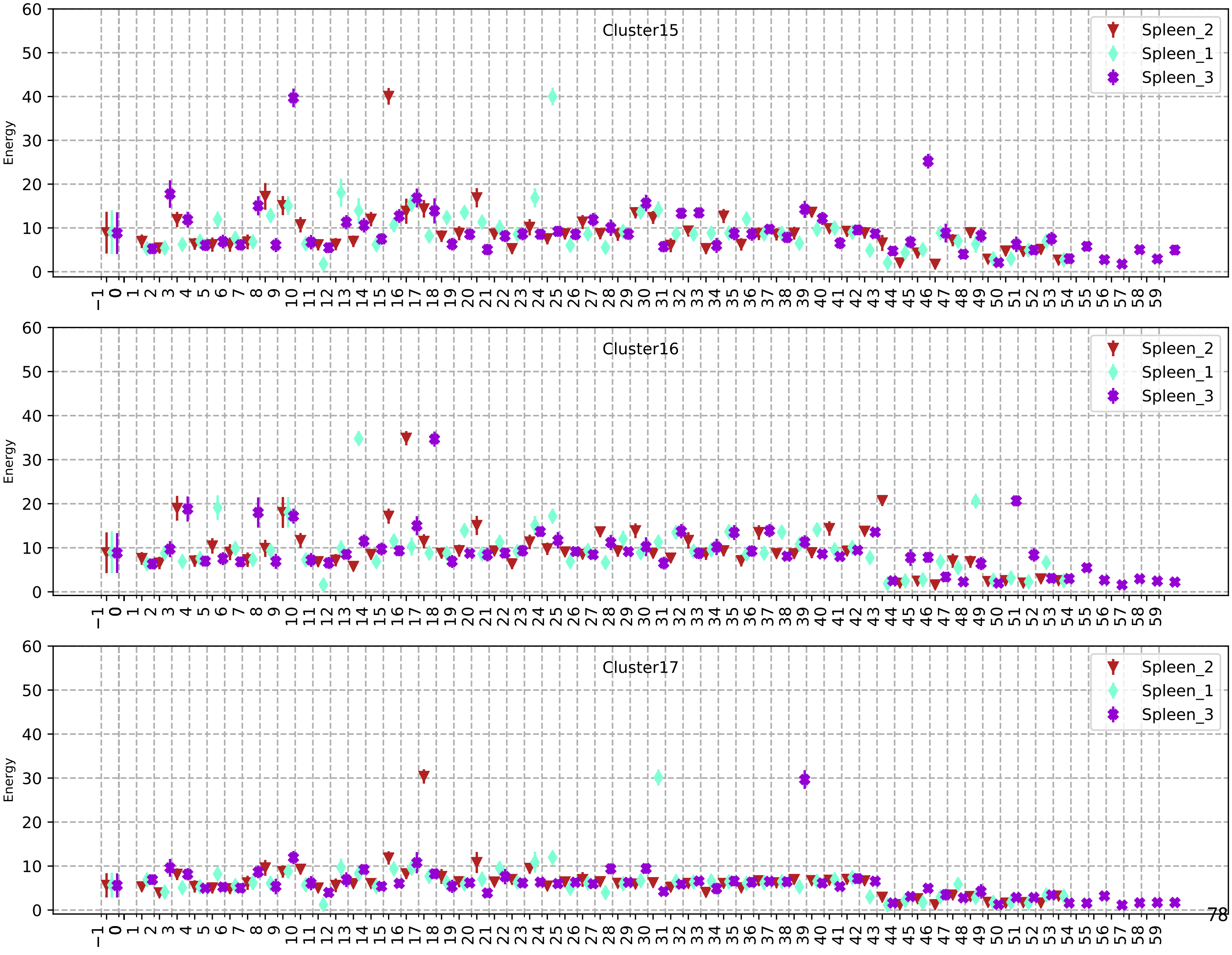

Cluster15
Cluster16
Cluster17
Spleen_2
Spleen_1
Spleen_3
Energy

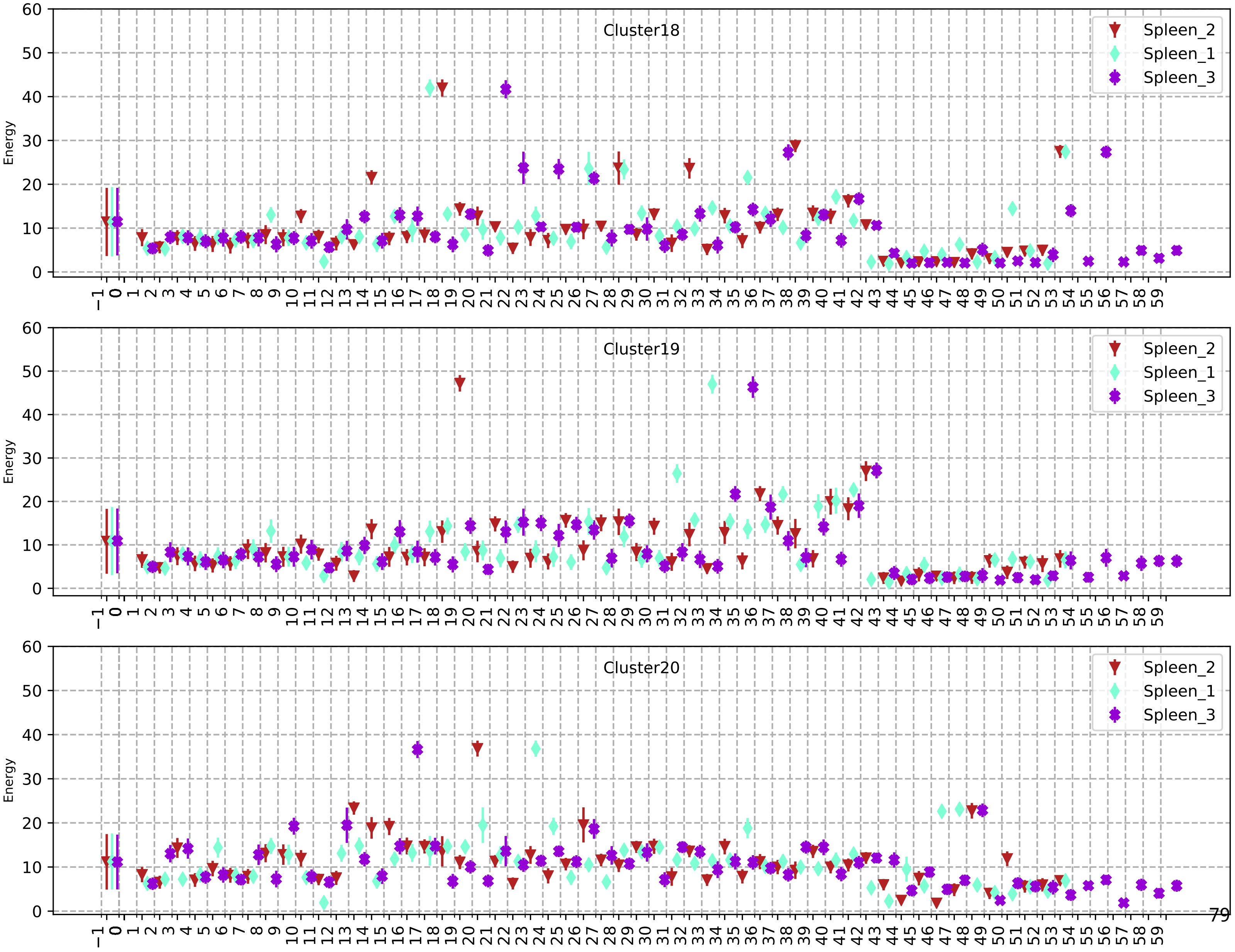
Cluster18
Spleen_2
Spleen_1
Spleen_3
Energy
Cluster19
Spleen_2
Spleen_1
Spleen_3
Energy
Cluster20
Spleen_2
Spleen_1
Spleen_3
Energy

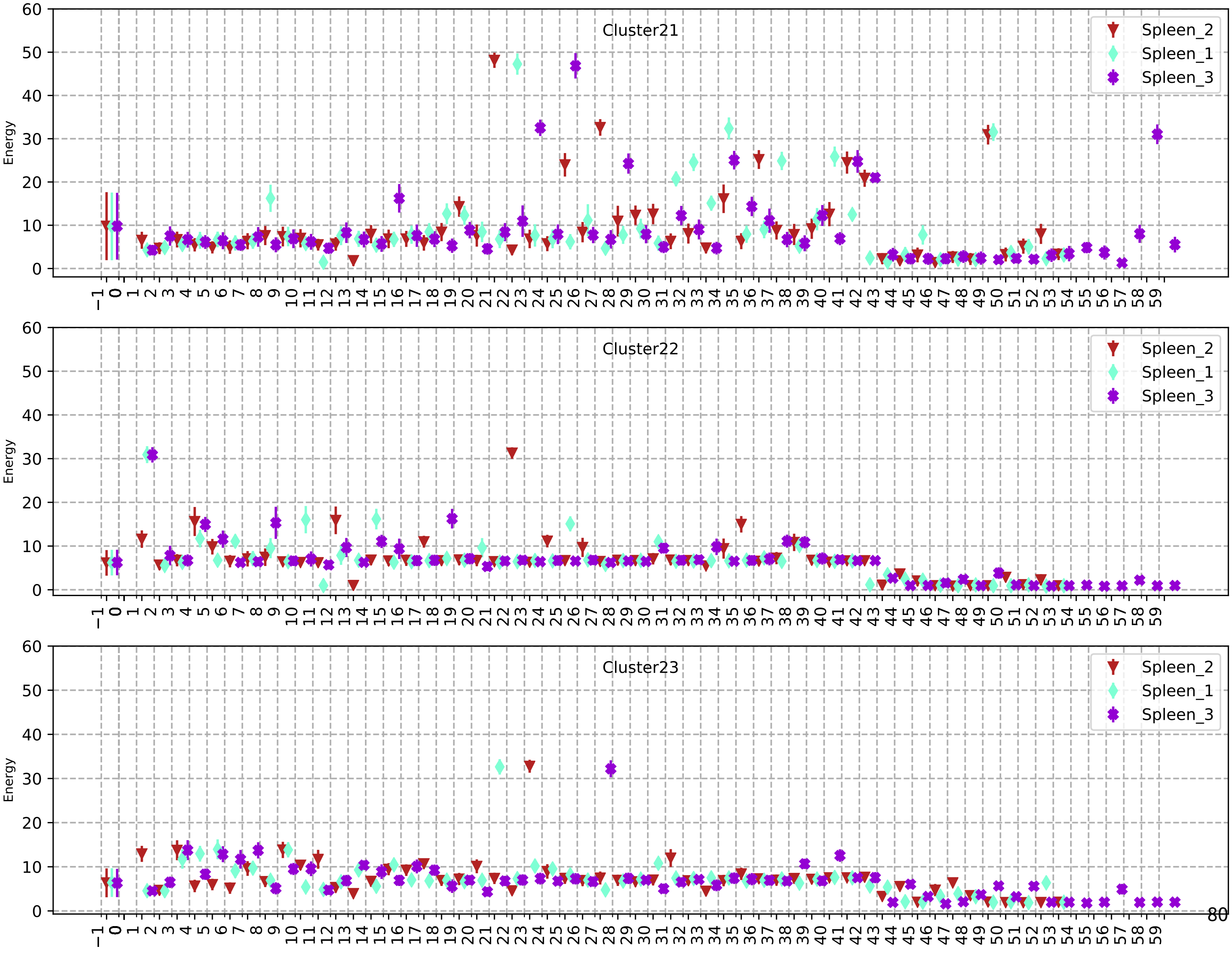
Cluster21
Cluster22
Cluster23
Spleen_2
Spleen_1
Spleen_3
Energy
0 10 20 30 40 50 60
−1 0 1 2 3 4 5 6 7 8 9 10 11 12 13 14 15 16 17 18 19 20 21 22 23 24 25 26 27 28 29 30 31 32 33 34 35 36 37 38 39 40 41 42 43 44 45 46 47 48 49 50 51 52 53 54 55 56 57 58 59

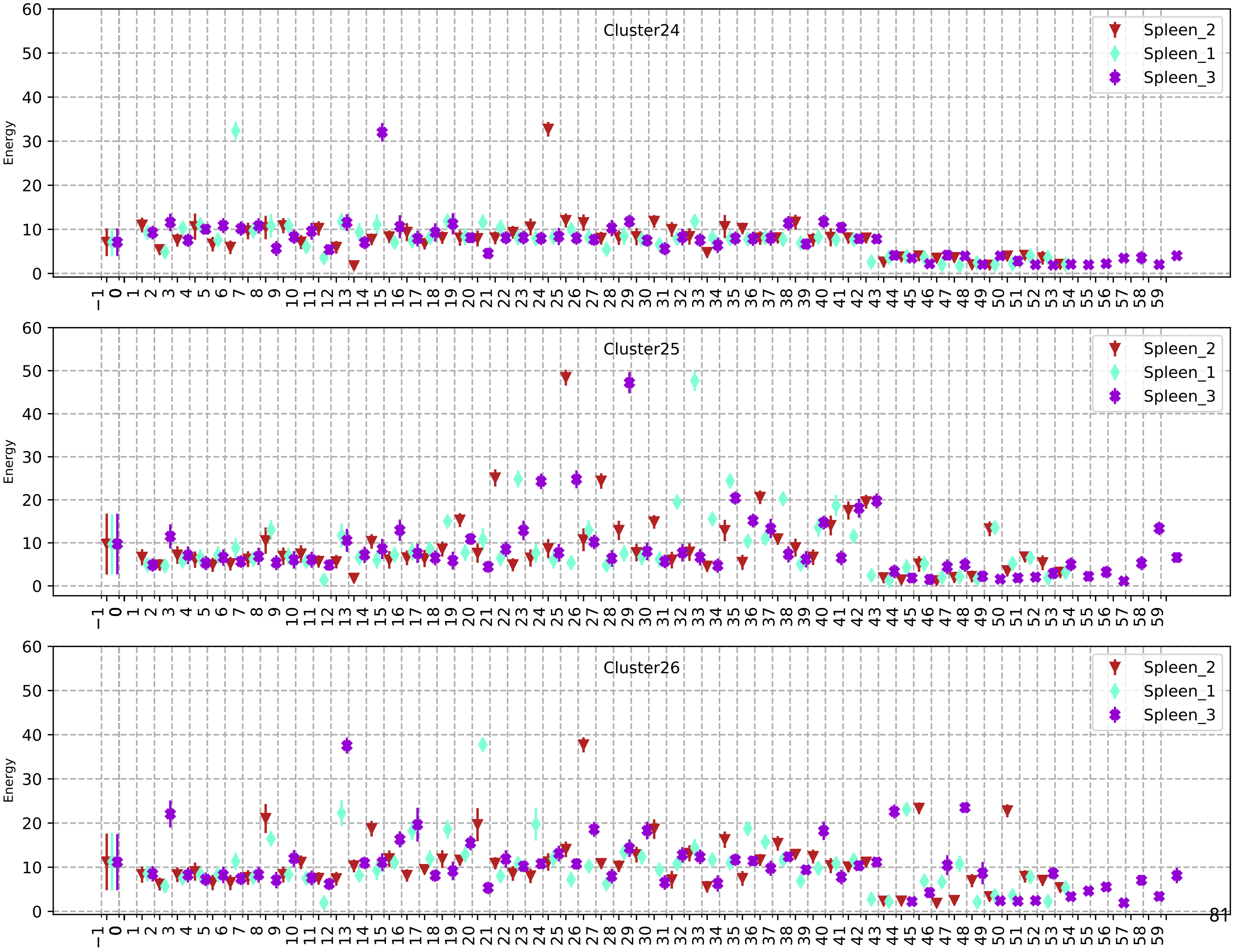
Cluster24
Cluster25
Cluster26
Spleen_2
Spleen_1
Spleen_3
Energy

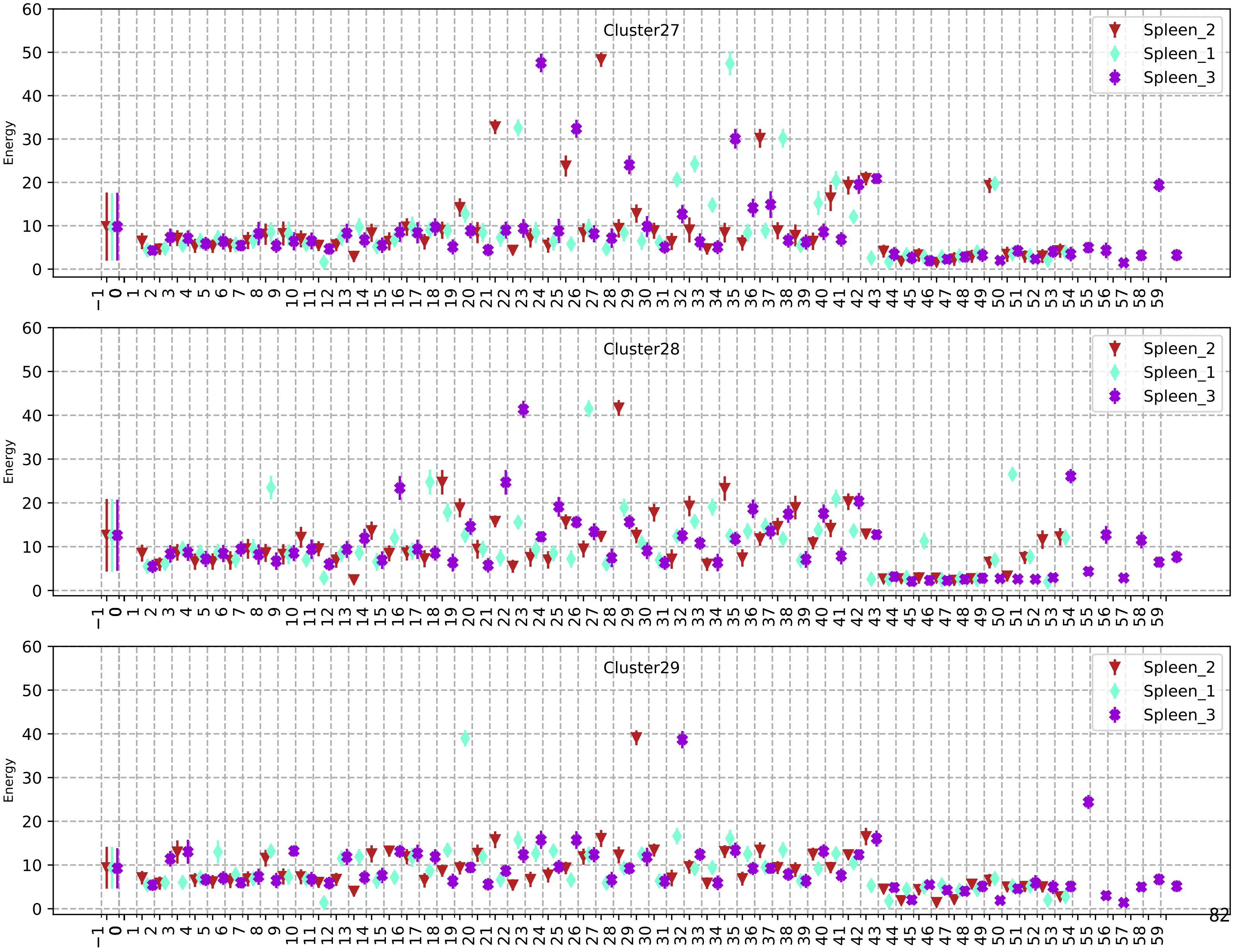
Cluster27
Cluster28
Cluster29
Spleen_2
Spleen_1
Spleen_3
Energy

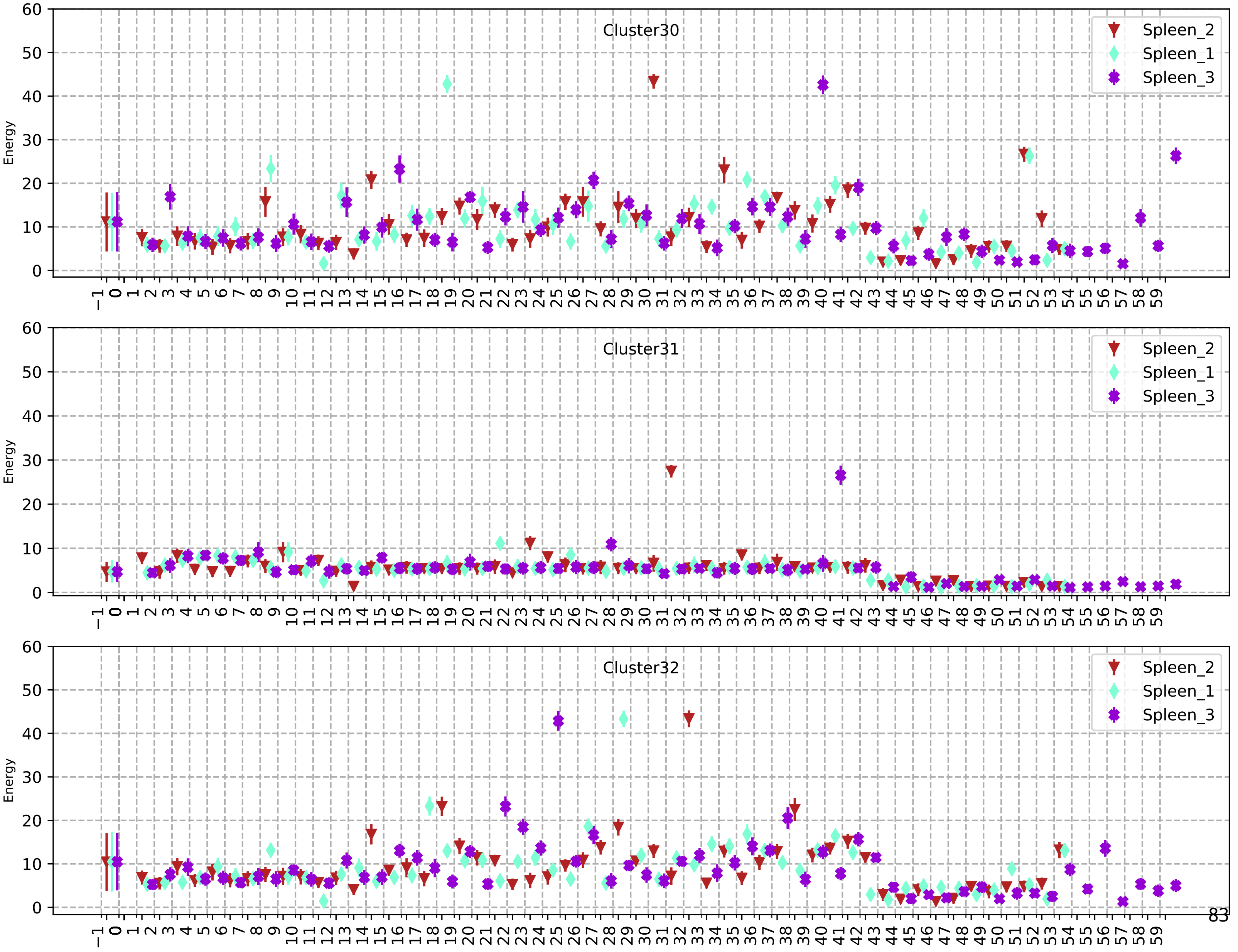
Cluster30
Cluster31
Cluster32
Spleen_2
Spleen_1
Spleen_3
Energy

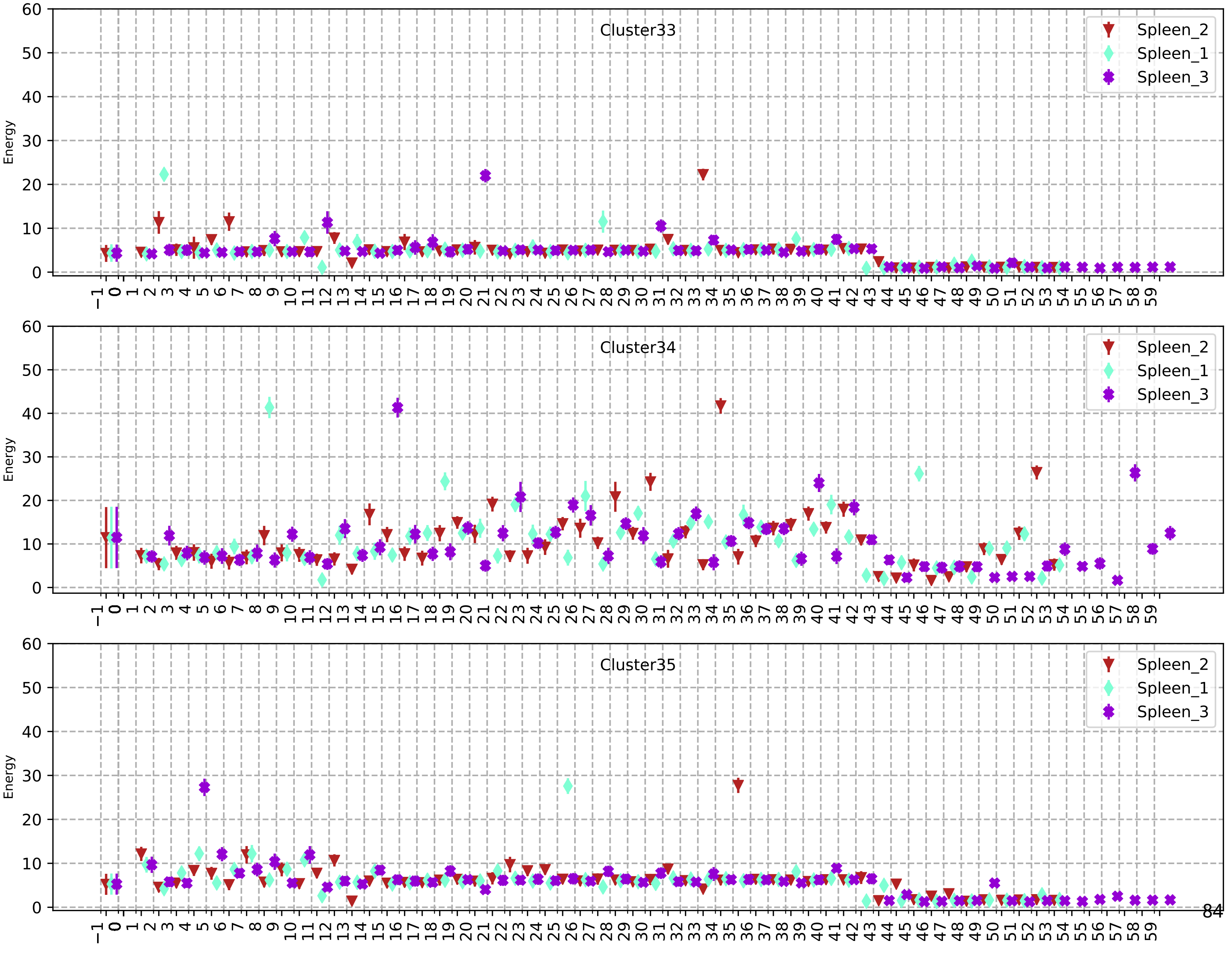

Cluster33
Cluster34
Cluster35
Spleen_2
Spleen_1
Spleen_3
Energy

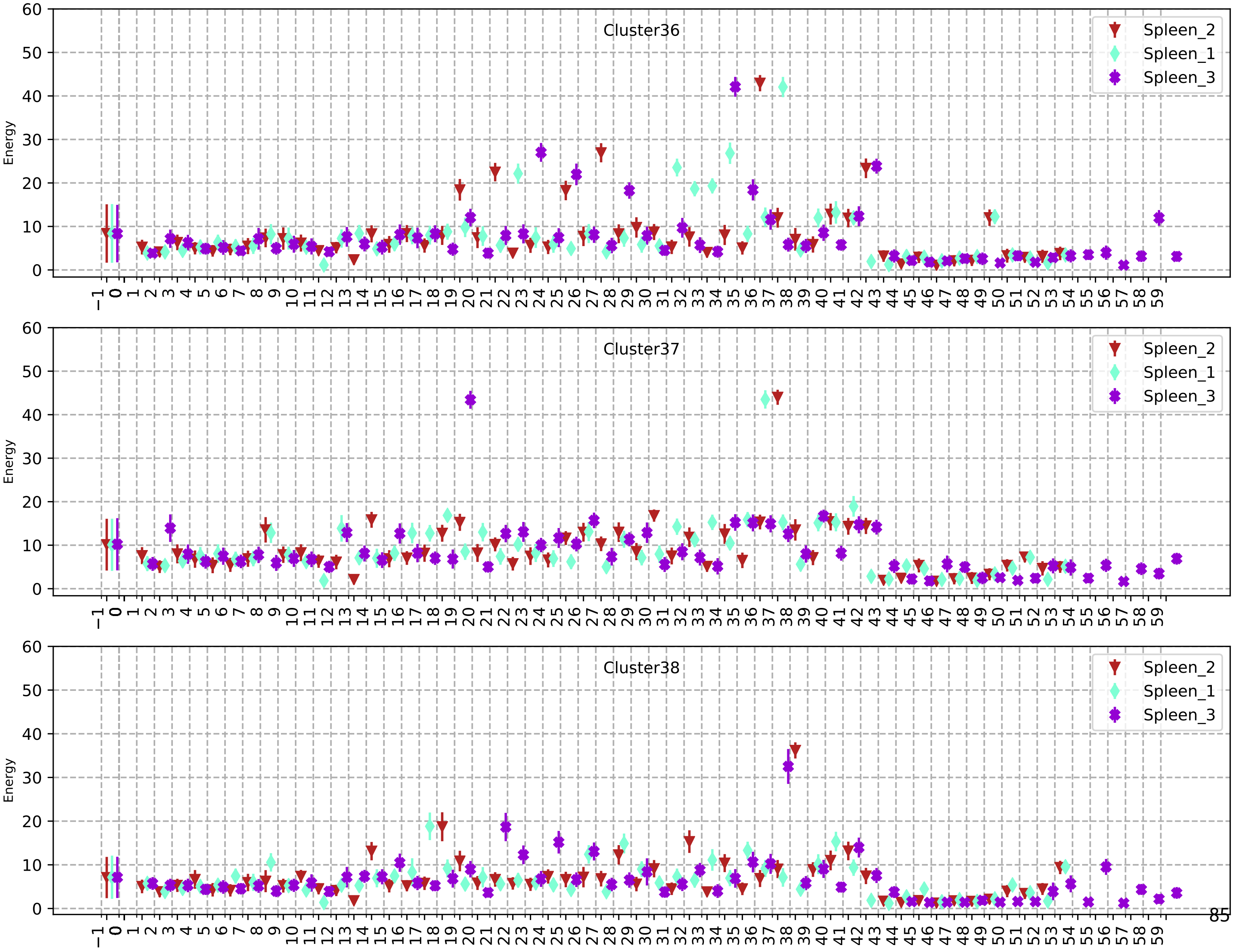
Cluster36
Cluster37
Cluster38
Spleen_2
Spleen_1
Spleen_3
Energy

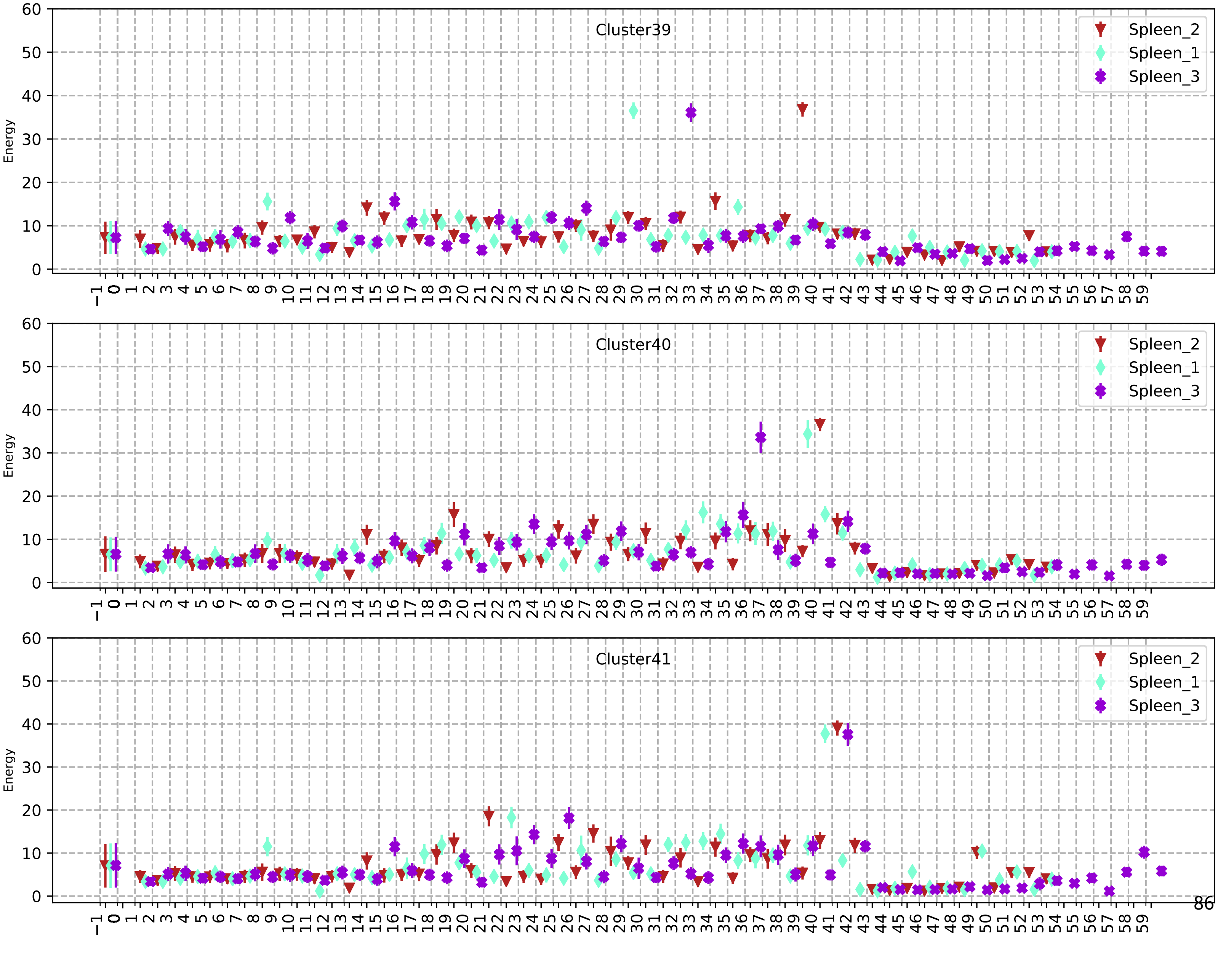
Cluster39
Cluster40
Cluster41
Spleen_2
Spleen_1
Spleen_3
Energy

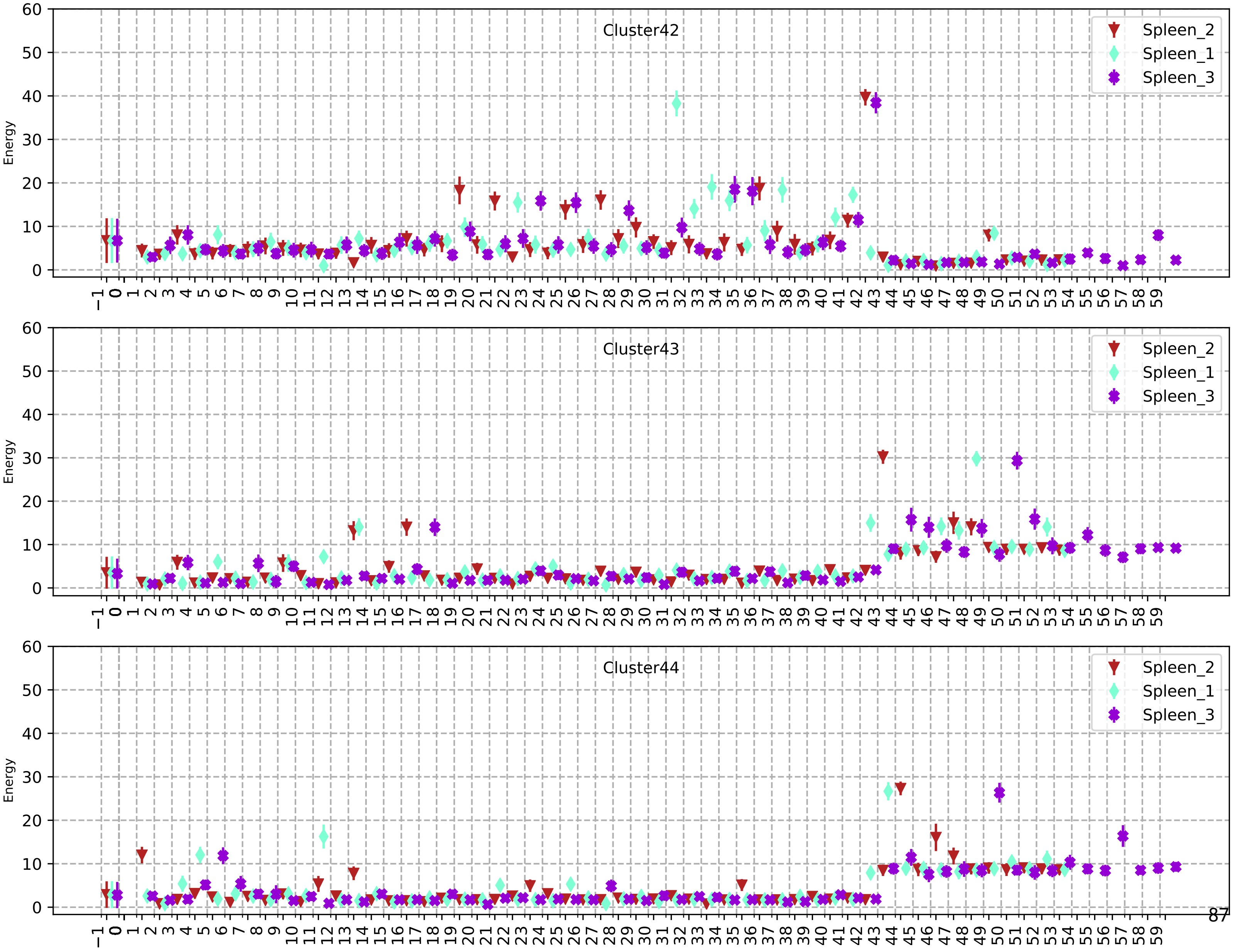
Cluster42
Cluster43
Cluster44
Spleen_2
Spleen_1
Spleen_3
Energy

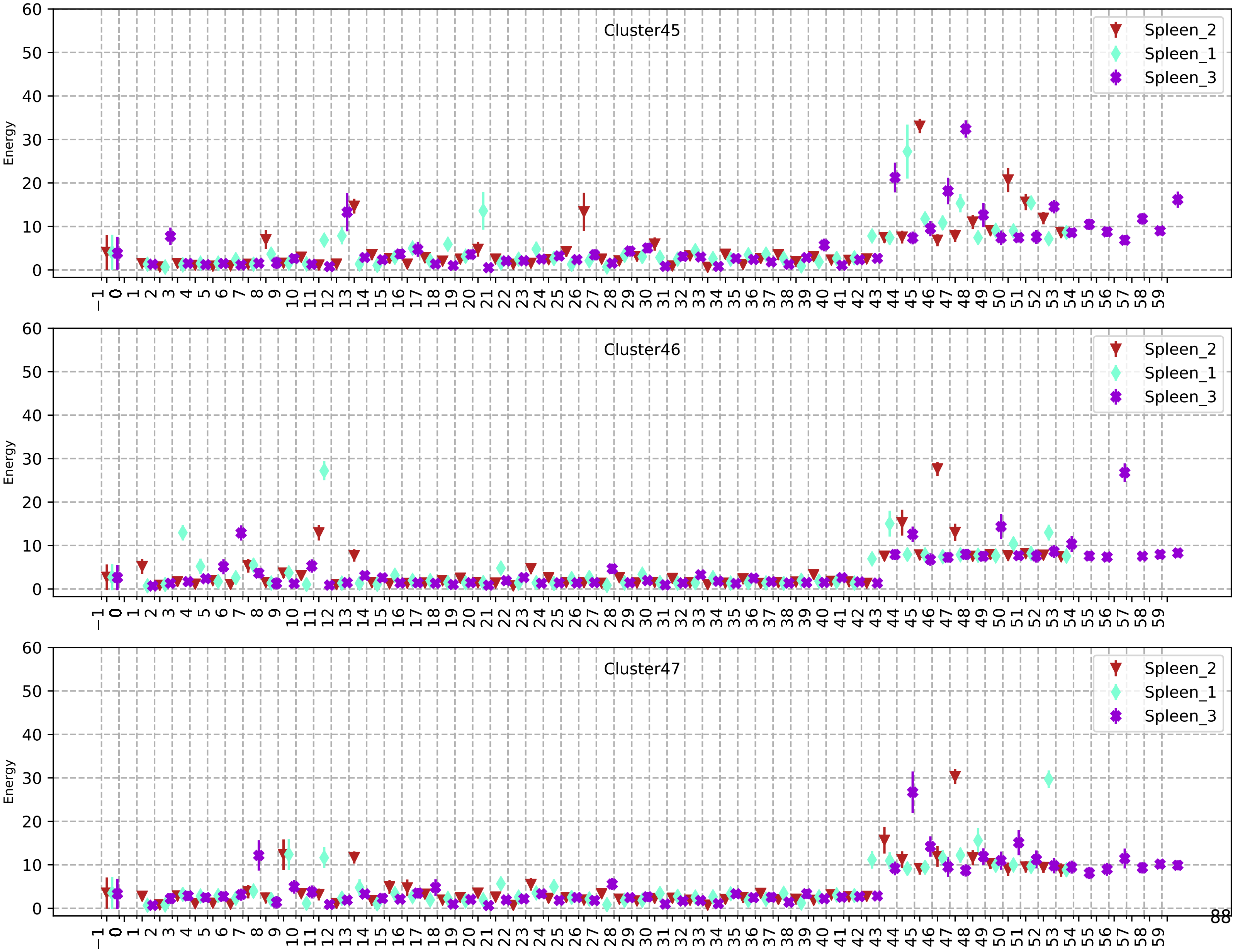
Cluster45
Cluster46
Cluster47
Spleen_2
Spleen_1
Spleen_3
Energy
0
10
20
30
40
50
60
−1 0 1 2 3 4 5 6 7 8 9 10 11 12 13 14 15 16 17 18 19 20 21 22 23 24 25 26 27 28 29 30 31 32 33 34 35 36 37 38 39 40 41 42 43 44 45 46 47 48 49 50 51 52 53 54 55 56 57 58 59

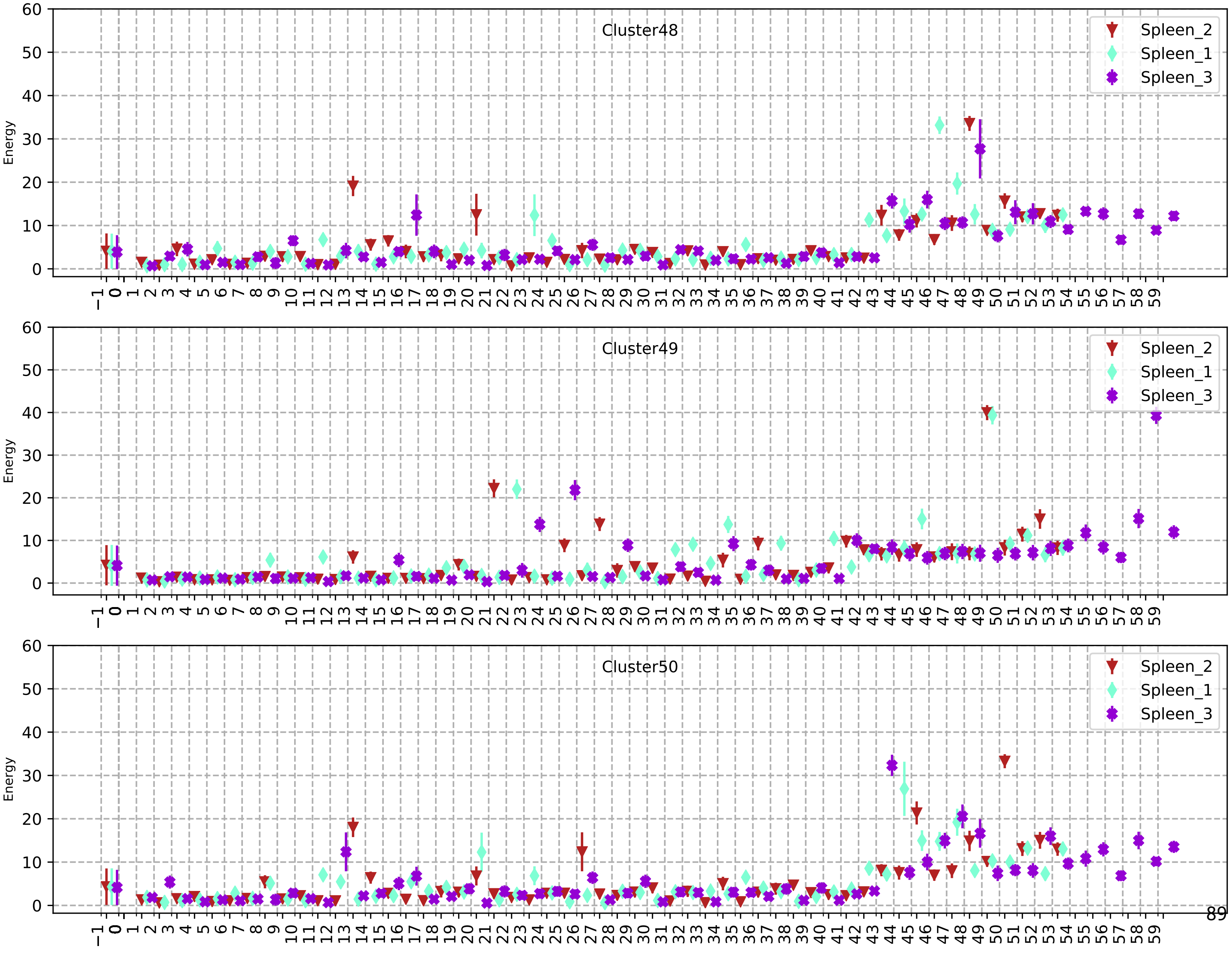
Cluster48
Energy
Spleen_2
Spleen_1
Spleen_3
Cluster49
Energy
Spleen_2
Spleen_1
Spleen_3
Cluster50
Energy
Spleen_2
Spleen_1
Spleen_3

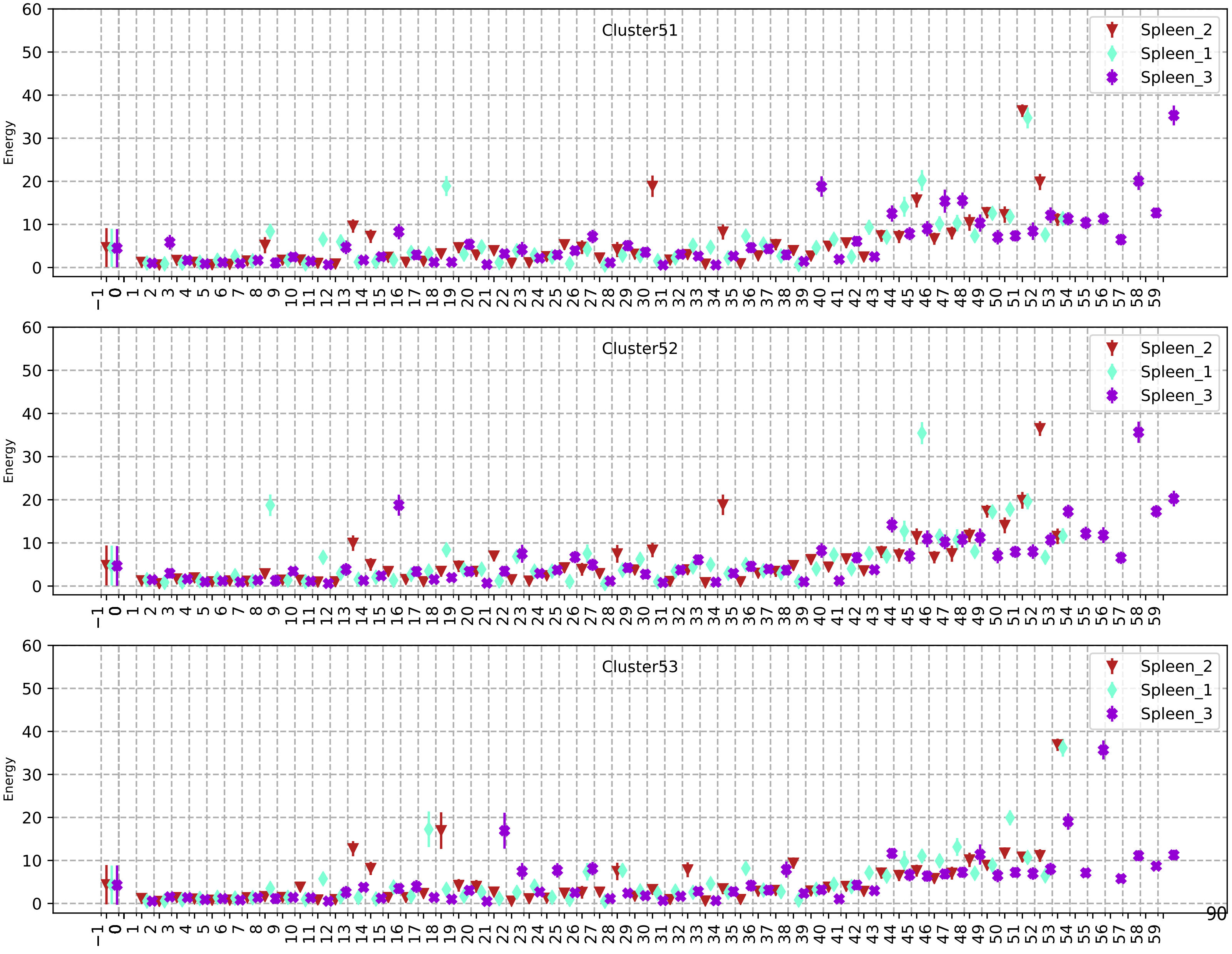
Cluster51
Cluster52
Cluster53
Spleen_2
Spleen_1
Spleen_3
Energy

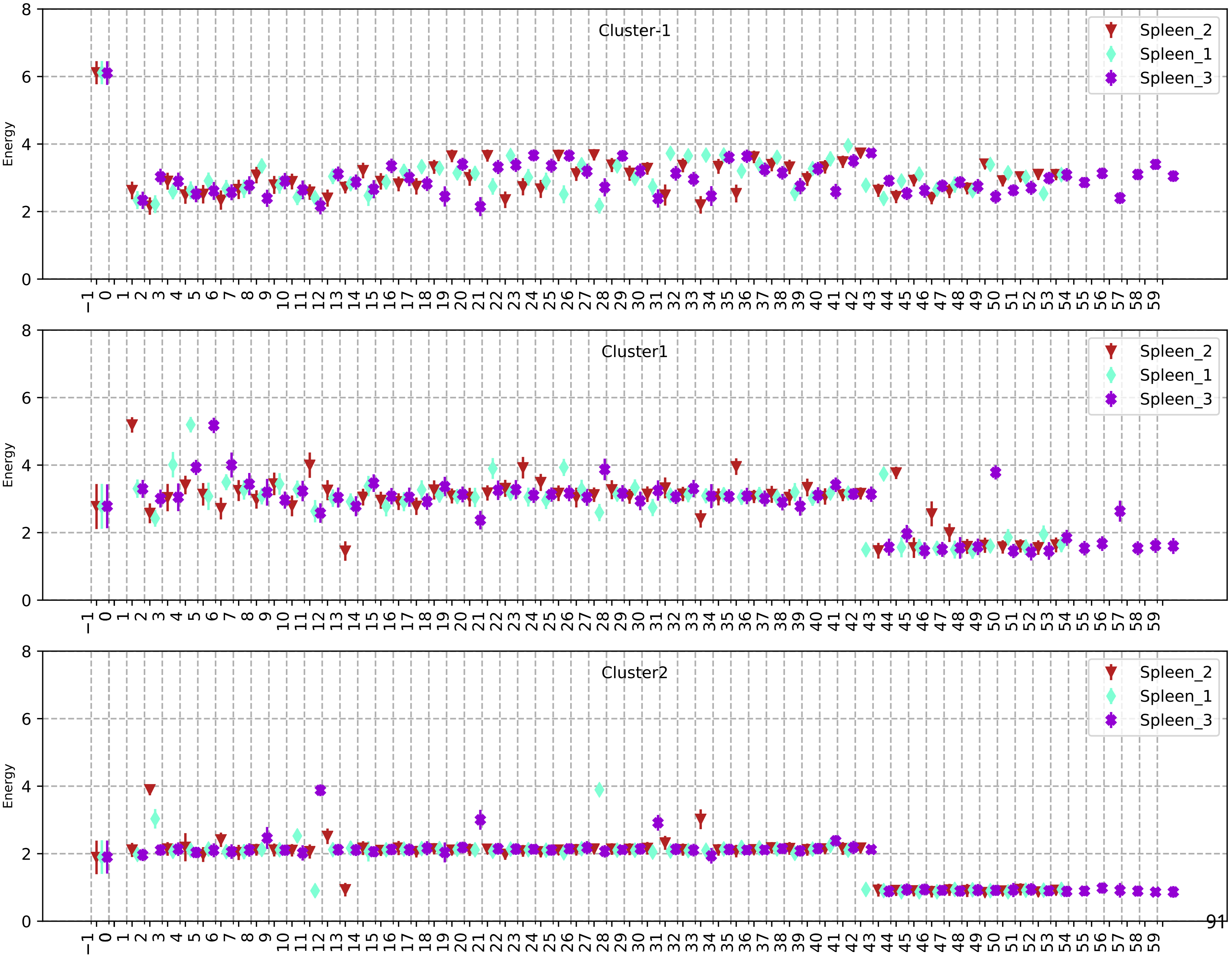
Cluster-1
Spleen_2
Spleen_1
Spleen_3
Energy
Cluster1
Spleen_2
Spleen_1
Spleen_3
Energy
Cluster2
Spleen_2
Spleen_1
Spleen_3
Energy

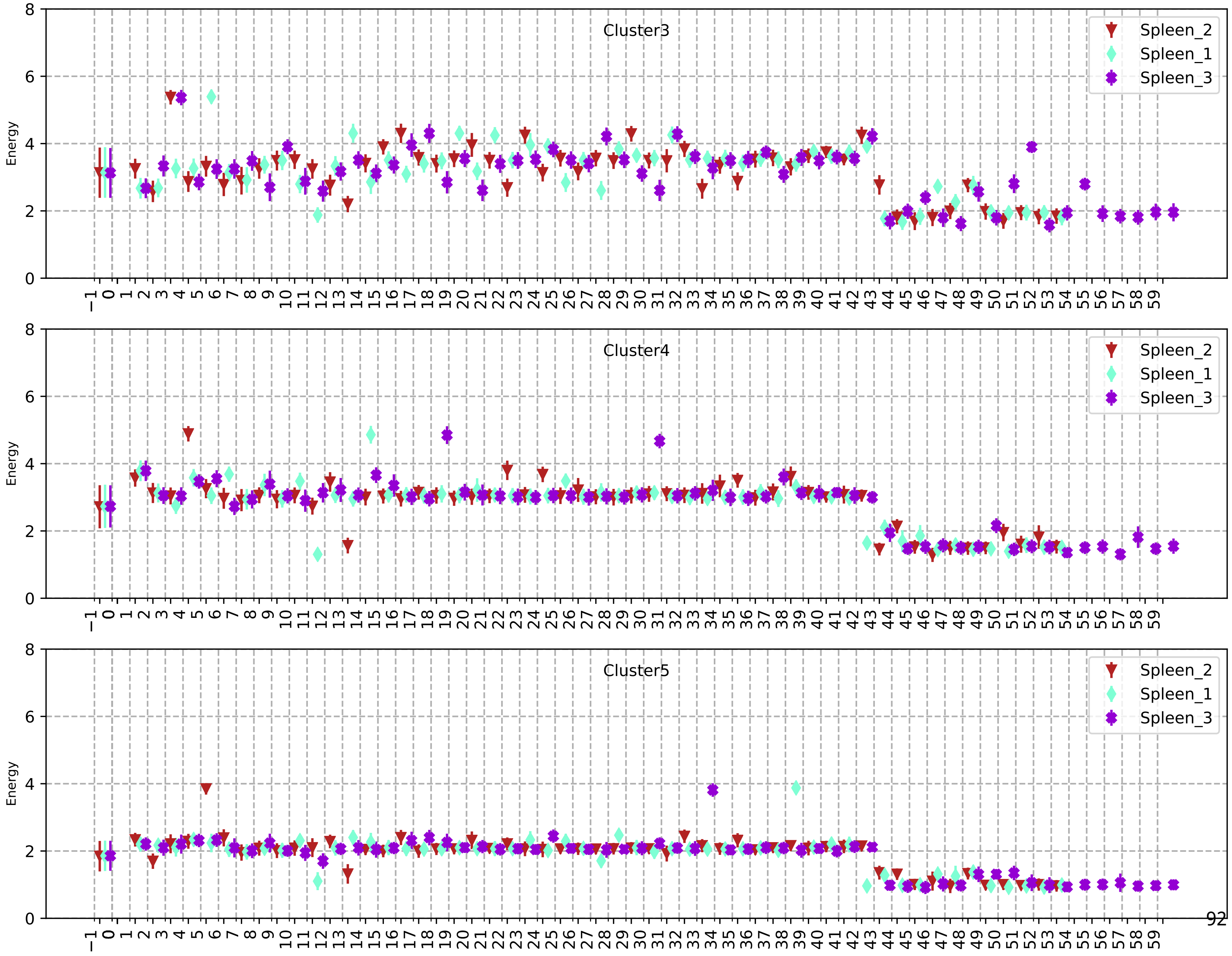

Cluster3
Cluster4
Cluster5
Spleen_2
Spleen_1
Spleen_3
Energy

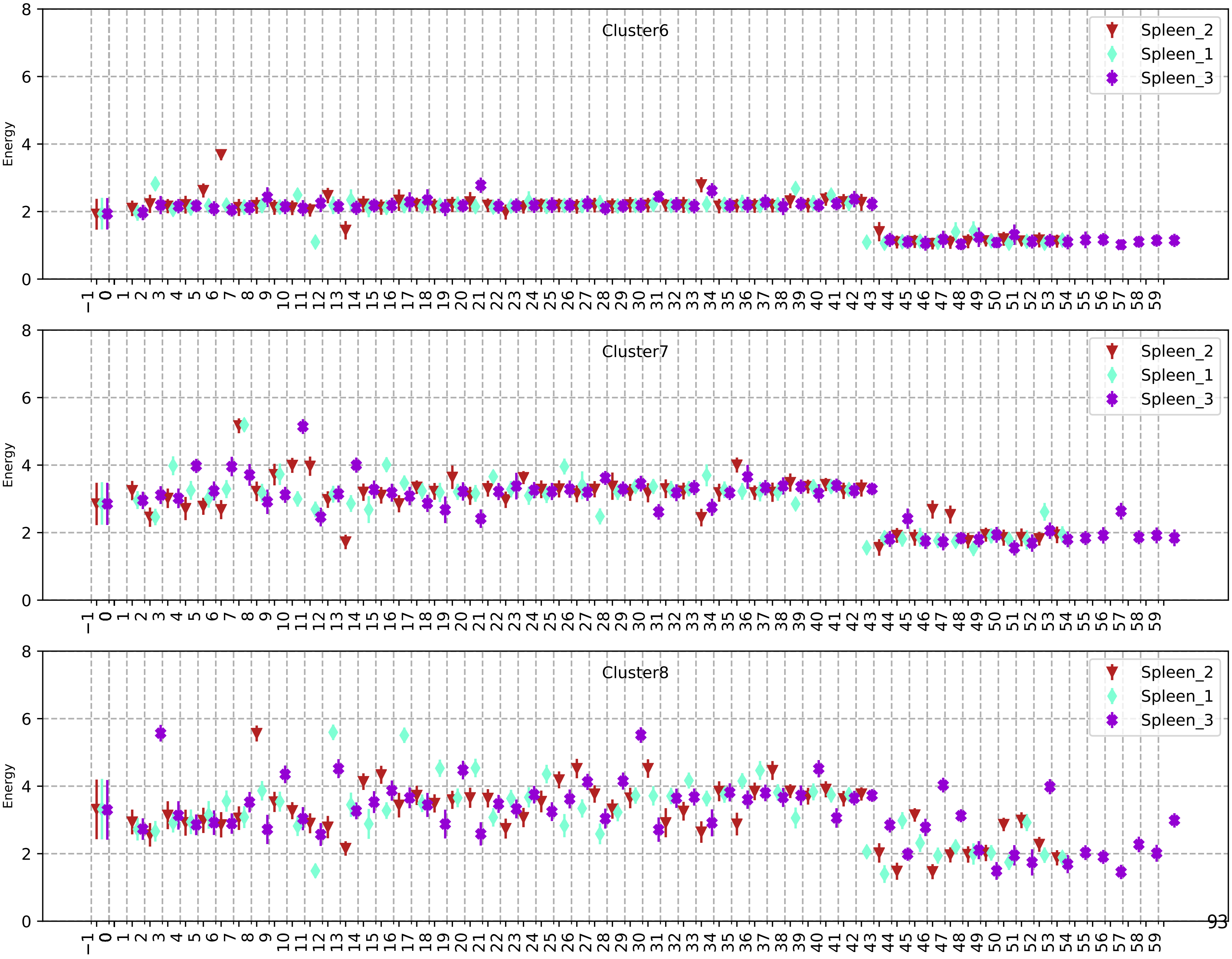
Cluster6
Cluster7
Cluster8
Spleen_2
Spleen_1
Spleen_3
Energy

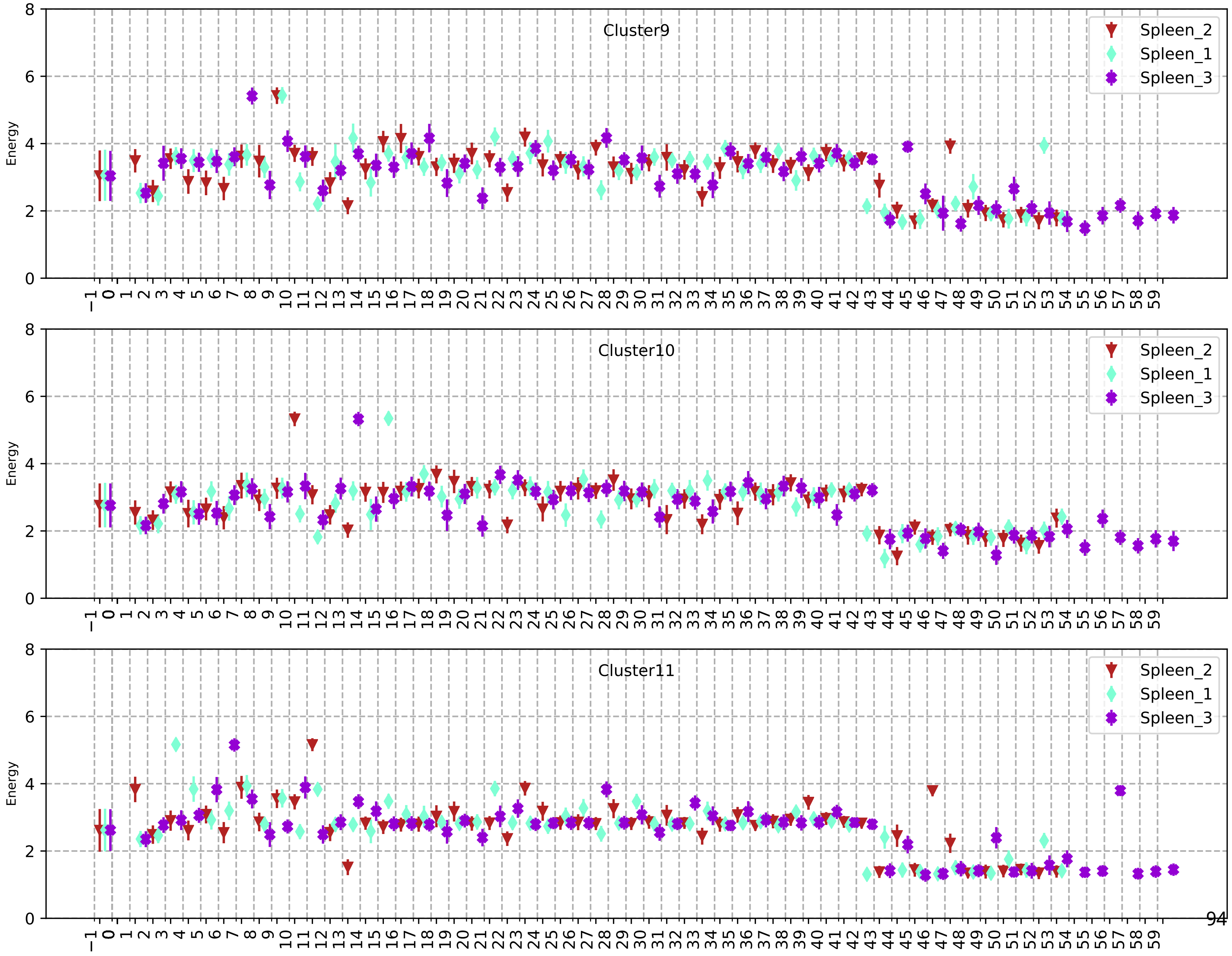
Cluster9
Cluster10
Cluster11
Energy
Spleen_2
Spleen_1
Spleen_3

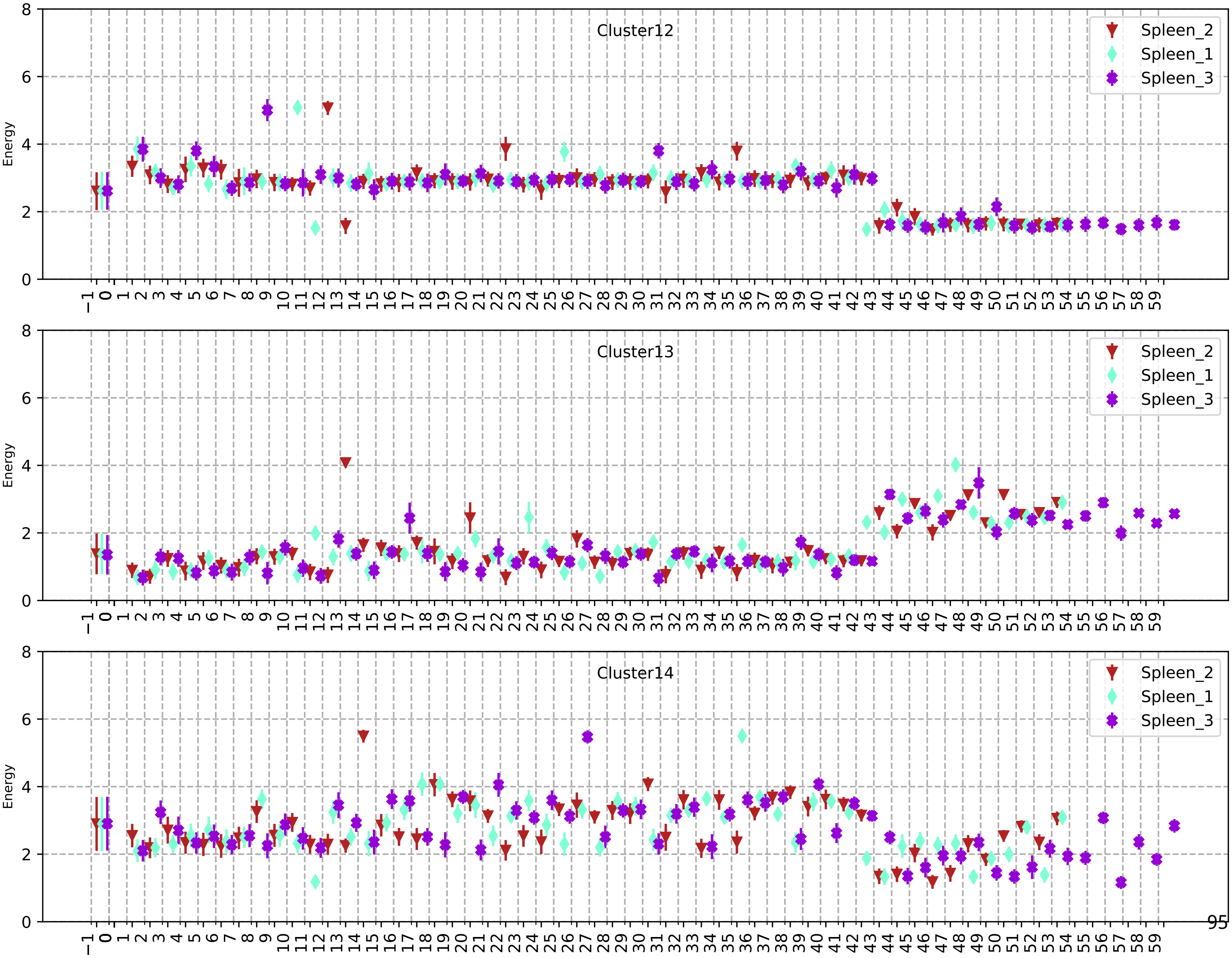
Cluster12
Spleen_2
Spleen_1
Spleen_3
Energy
Cluster13
Spleen_2
Spleen_1
Spleen_3
Energy
Cluster14
Spleen_2
Spleen_1
Spleen_3
Energy

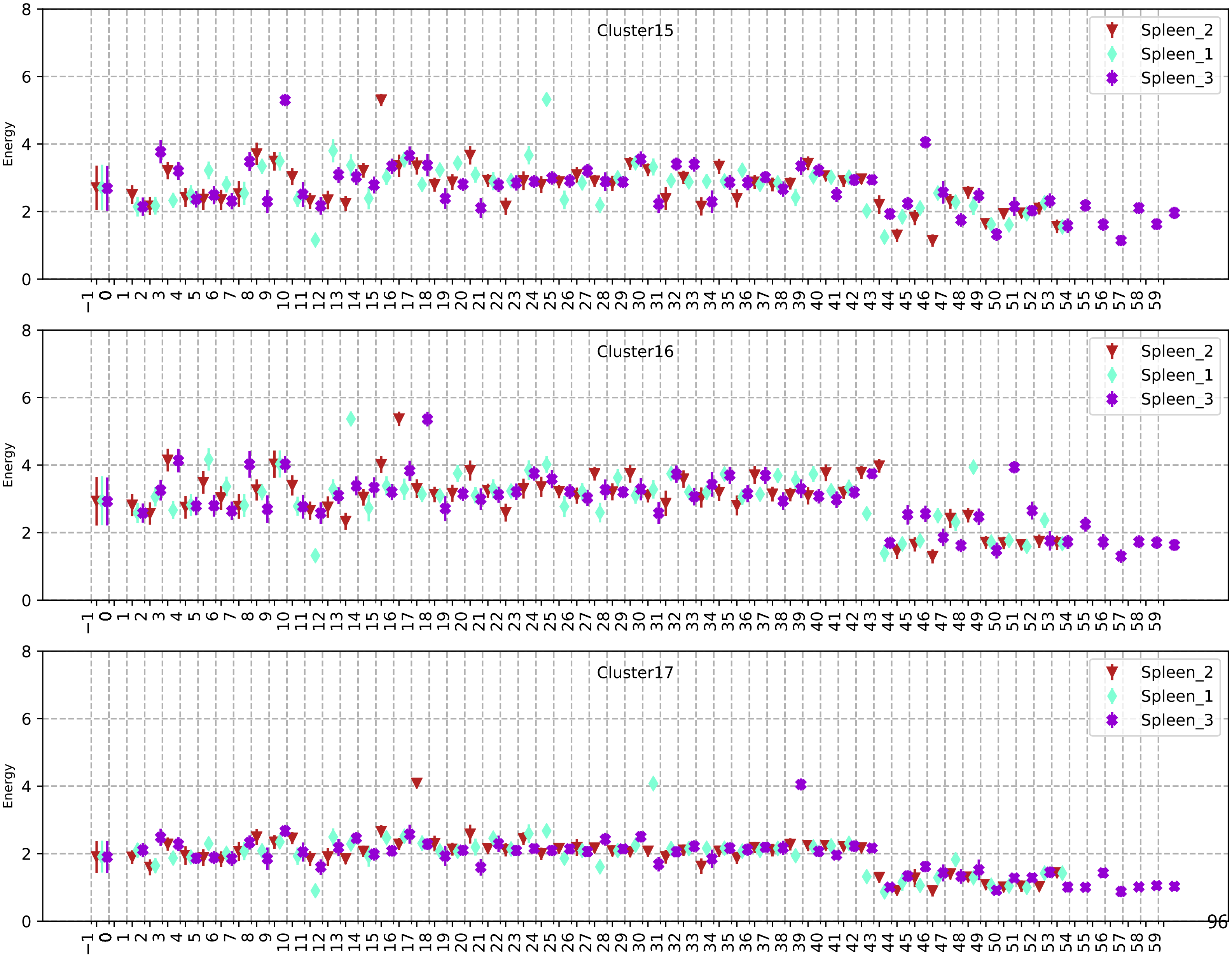
Cluster15
Cluster16
Cluster17
Energy
Spleen_2
Spleen_1
Spleen_3
0
2
4
6
8
−1 0 1 2 3 4 5 6 7 8 9 10 11 12 13 14 15 16 17 18 19 20 21 22 23 24 25 26 27 28 29 30 31 32 33 34 35 36 37 38 39 40 41 42 43 44 45 46 47 48 49 50 51 52 53 54 55 56 57 58 59

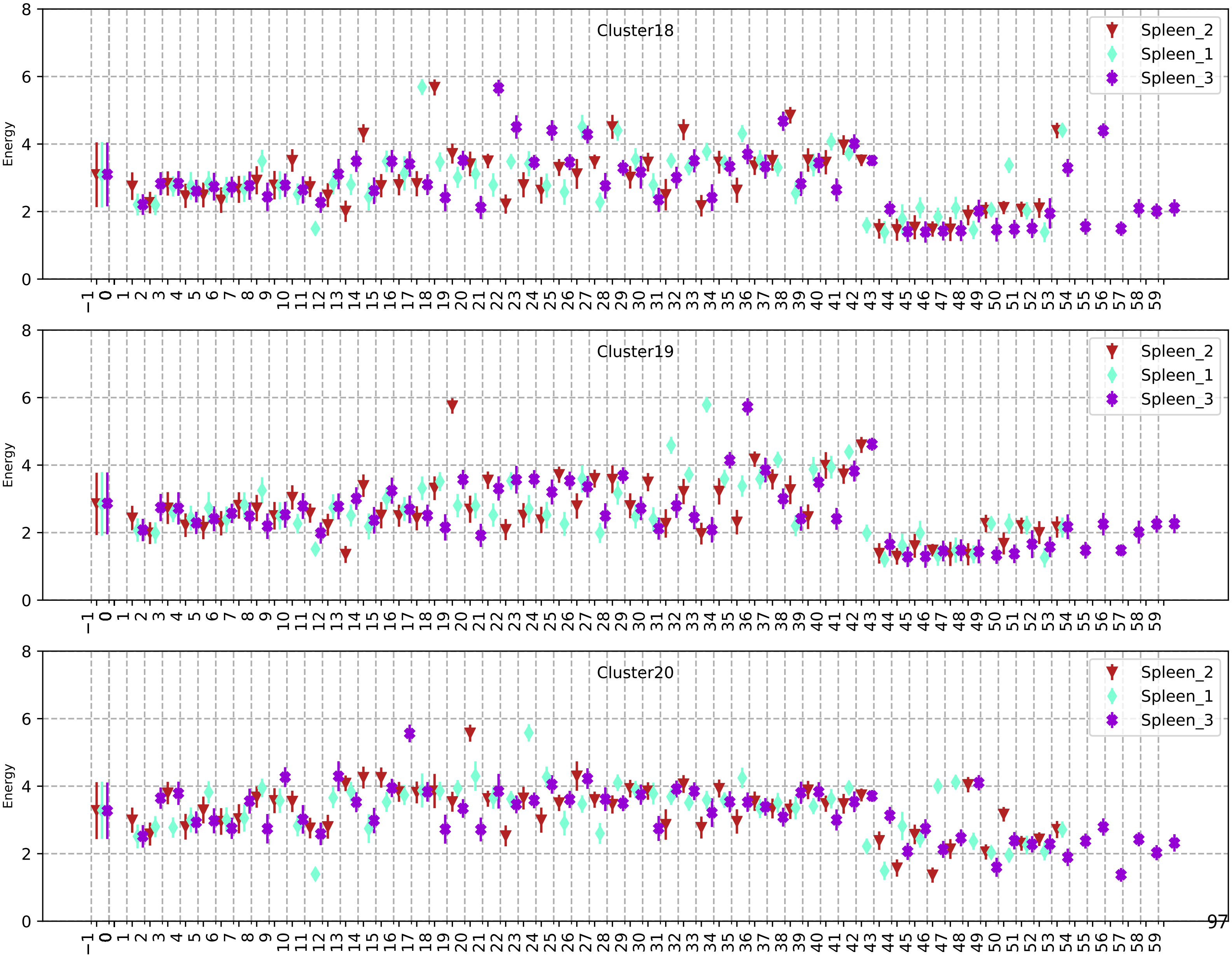
Cluster18
Cluster19
Cluster20
Energy
Spleen_2
Spleen_1
Spleen_3

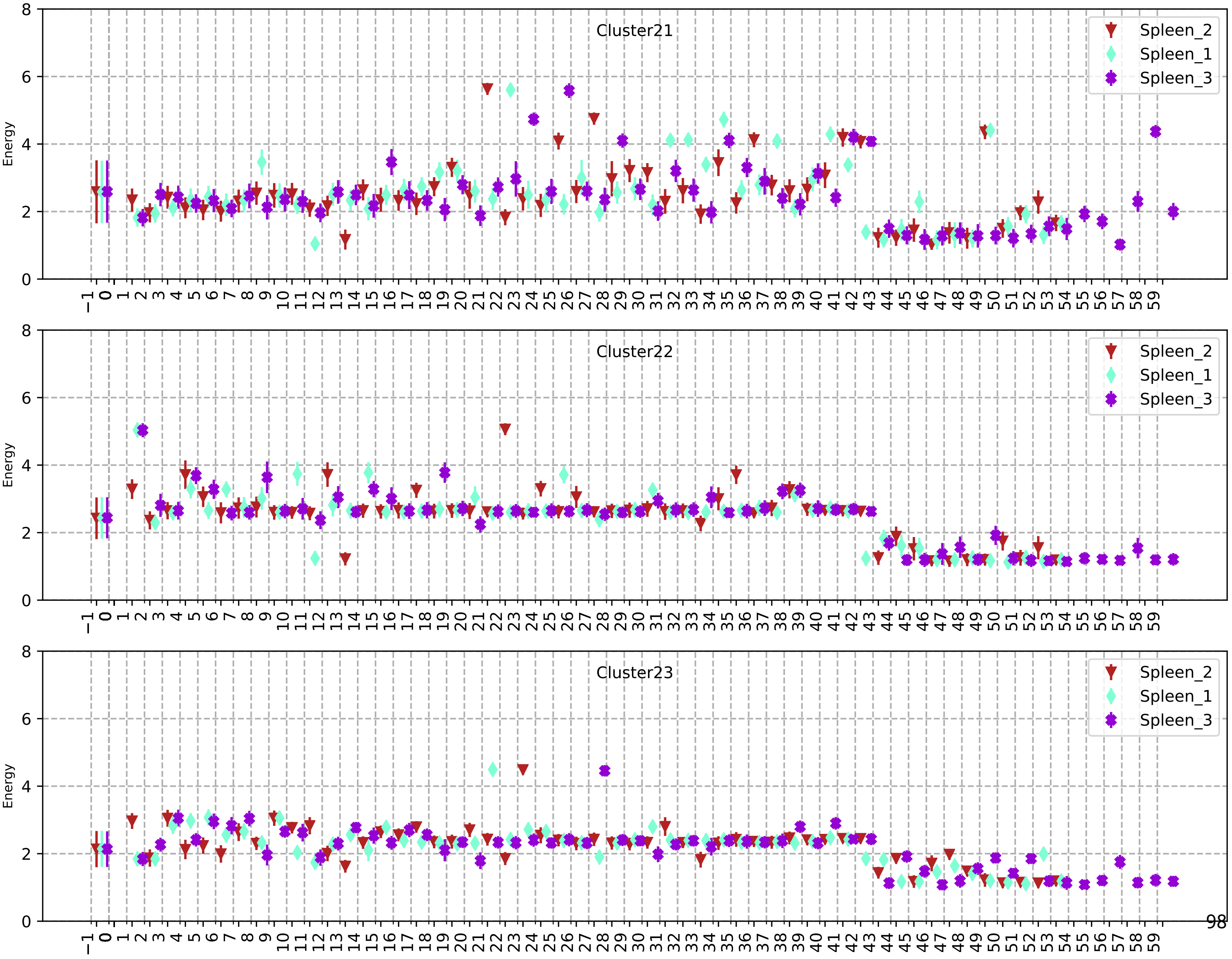
Cluster21
Cluster22
Cluster23
Spleen_2
Spleen_1
Spleen_3
Energy

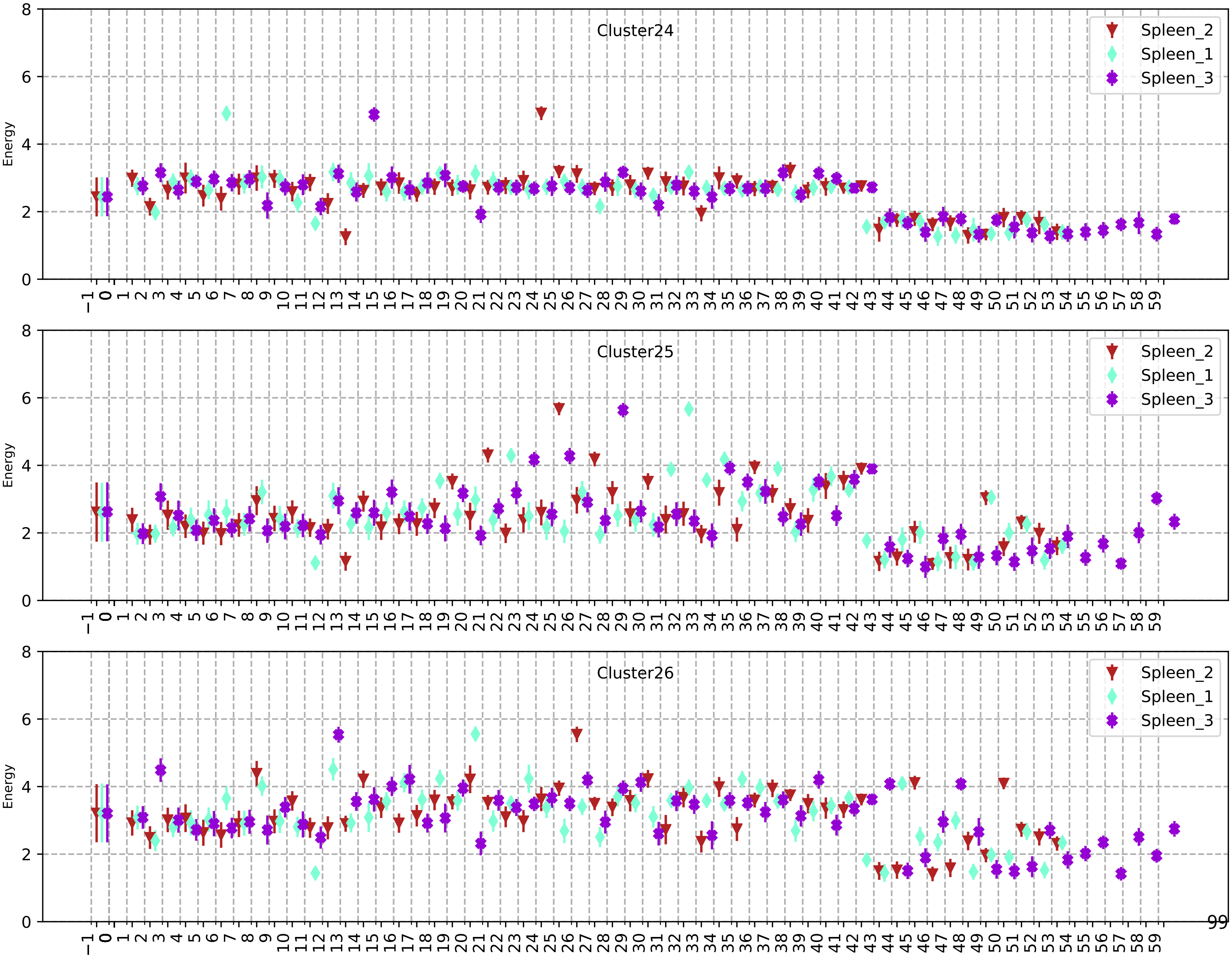
Cluster24
Cluster25
Cluster26
Energy
Spleen_2
Spleen_1
Spleen_3

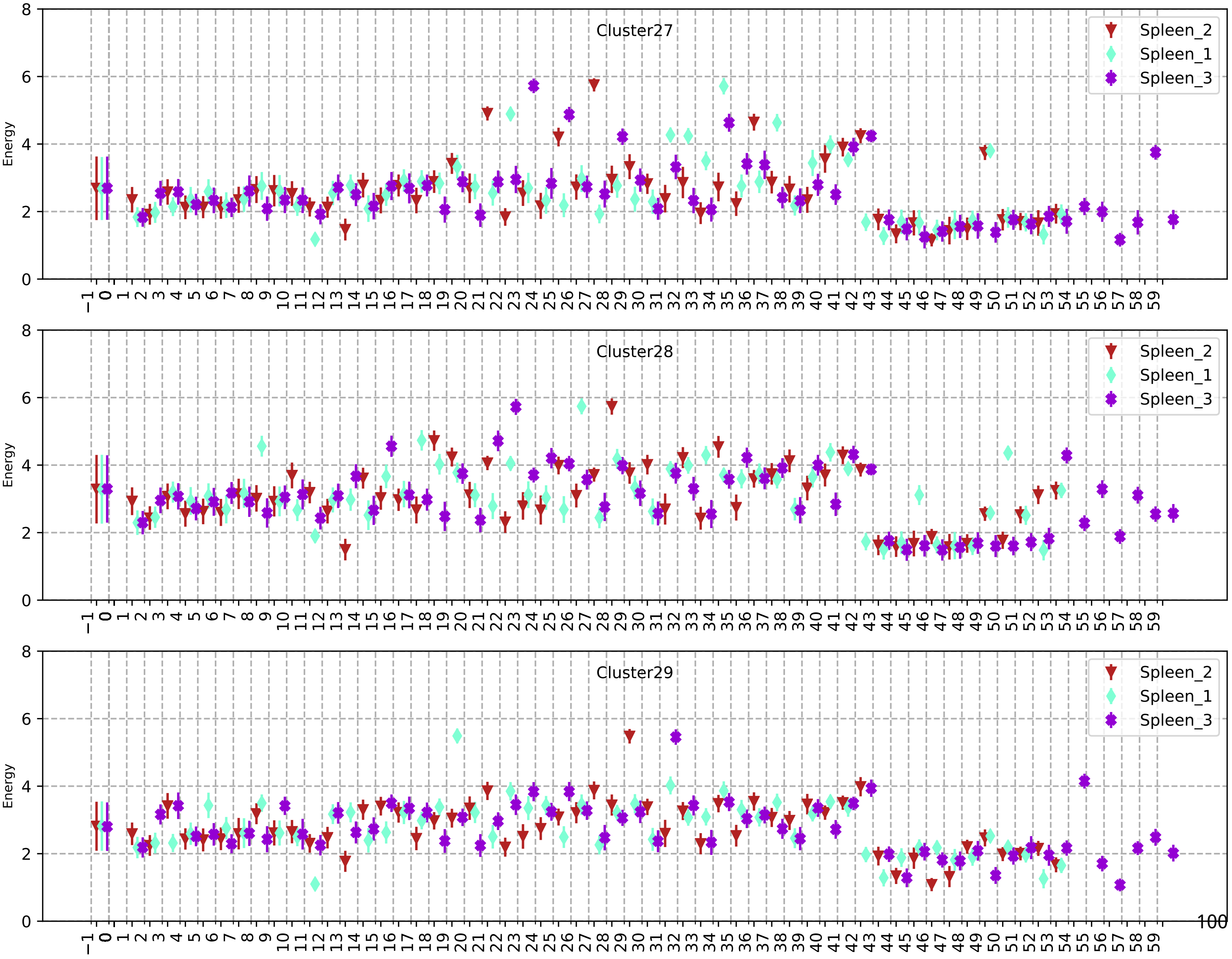
Cluster27
Cluster28
Cluster29
Energy
Spleen_2
Spleen_1
Spleen_3
100

147

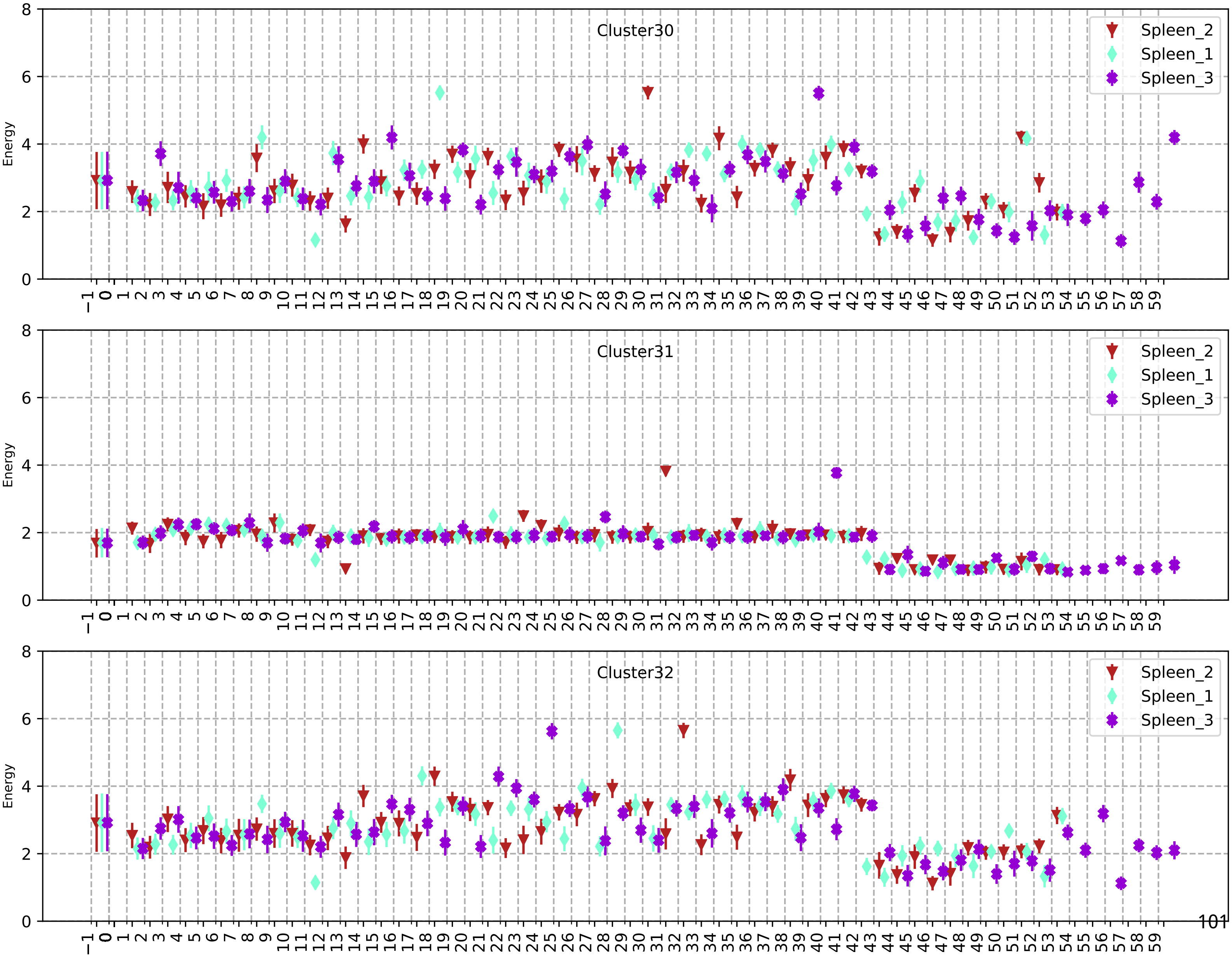

Cluster30
Spleen_2
Spleen_1
Spleen_3
Energy
Cluster31
Spleen_2
Spleen_1
Spleen_3
Energy
Cluster32
Spleen_2
Spleen_1
Spleen_3
Energy

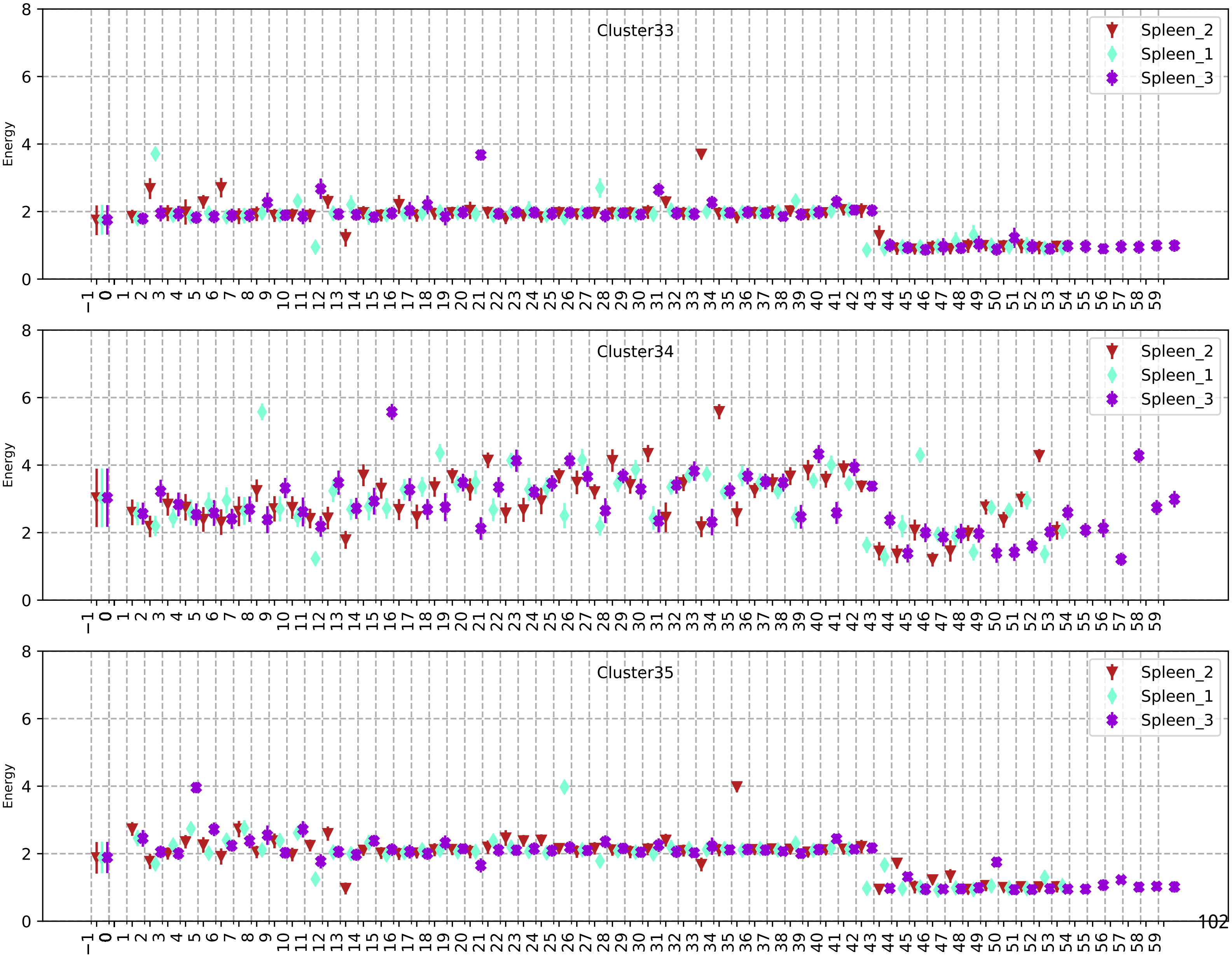
Cluster33
Spleen_2
Spleen_1
Spleen_3
Energy
Cluster34
Spleen_2
Spleen_1
Spleen_3
Energy
Cluster35
Spleen_2
Spleen_1
Spleen_3
Energy

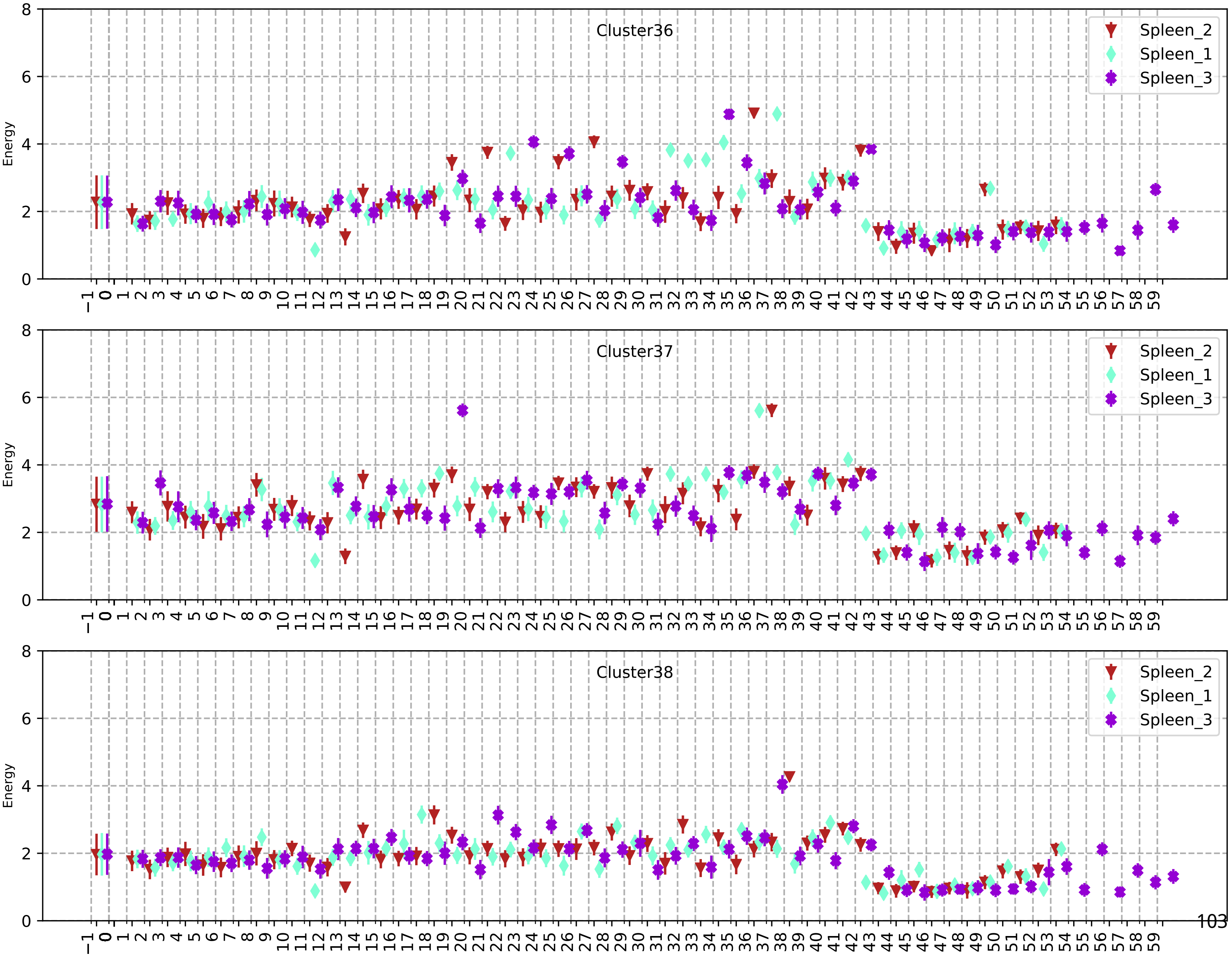
Cluster36
Spleen_2
Spleen_1
Spleen_3
Energy
Cluster37
Spleen_2
Spleen_1
Spleen_3
Energy
Cluster38
Spleen_2
Spleen_1
Spleen_3
Energy

150

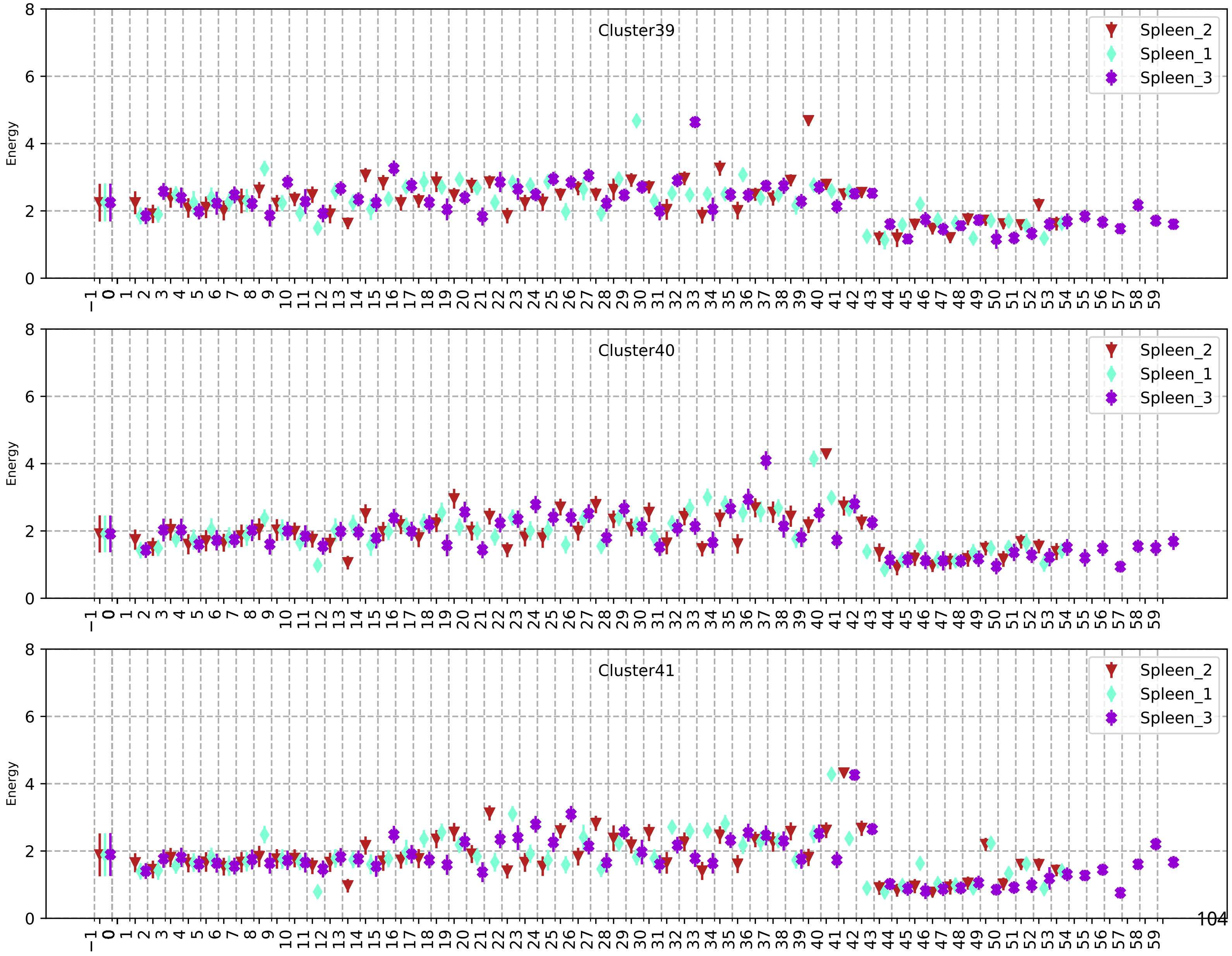
Cluster39
Cluster40
Cluster41
Spleen_2
Spleen_1
Spleen_3
Energy
0
2
4
6
8
−1 0 1 2 3 4 5 6 7 8 9 10 11 12 13 14 15 16 17 18 19 20 21 22 23 24 25 26 27 28 29 30 31 32 33 34 35 36 37 38 39 40 41 42 43 44 45 46 47 48 49 50 51 52 53 54 55 56 57 58 59

104

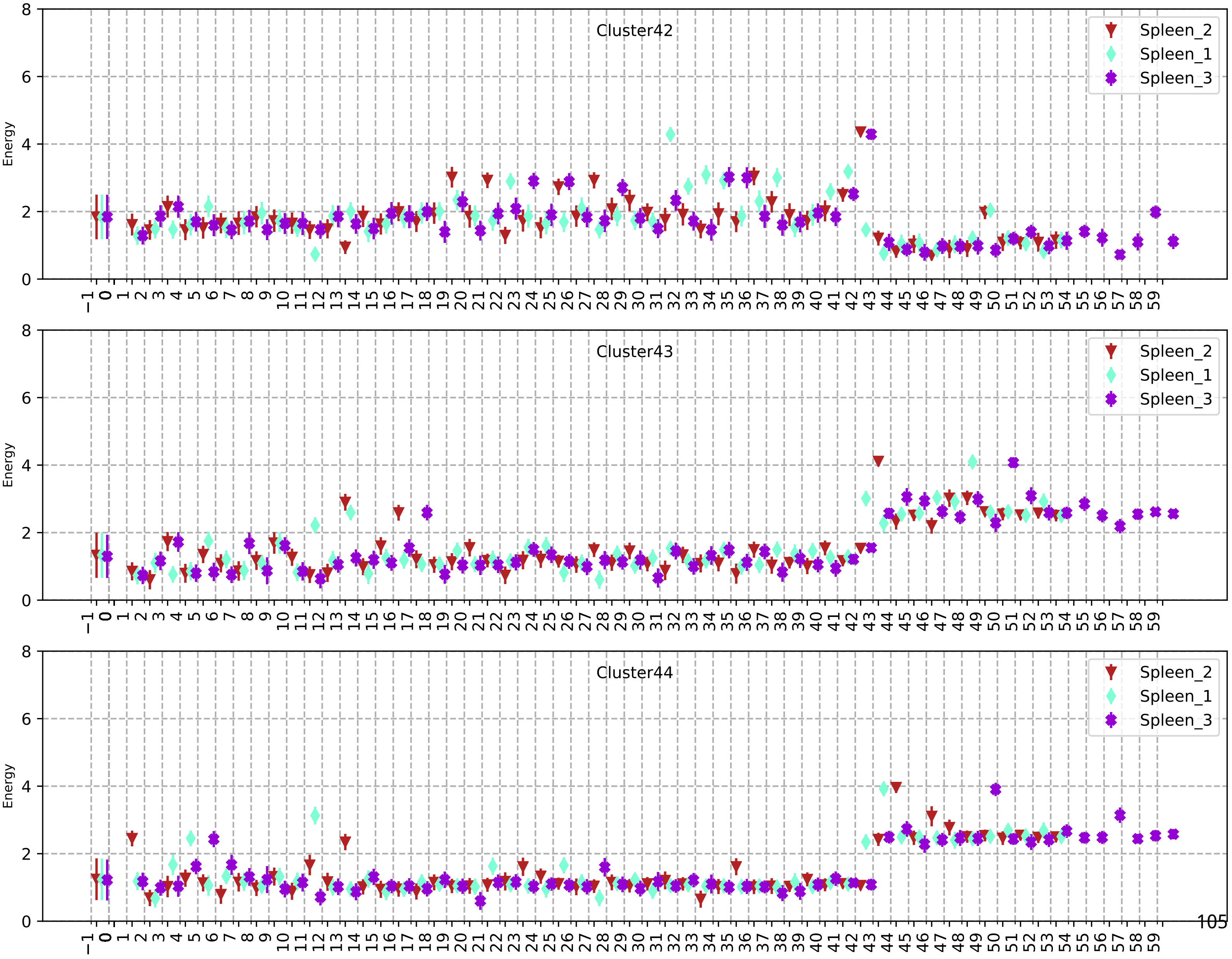
Cluster42
Cluster43
Cluster44
Energy
Spleen_2
Spleen_1
Spleen_3

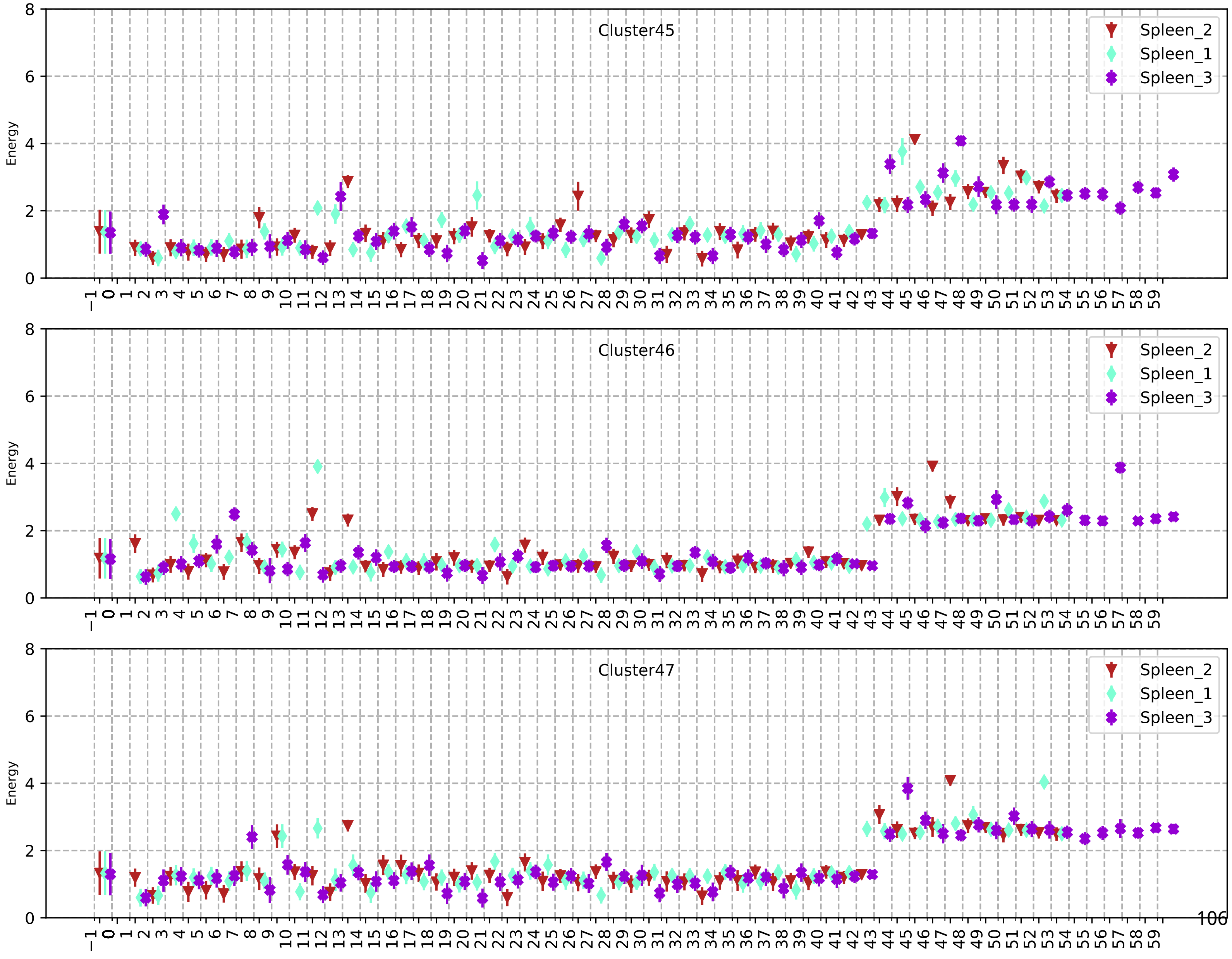
Cluster45
Cluster46
Cluster47
Spleen_2
Spleen_1
Spleen_3
Energy

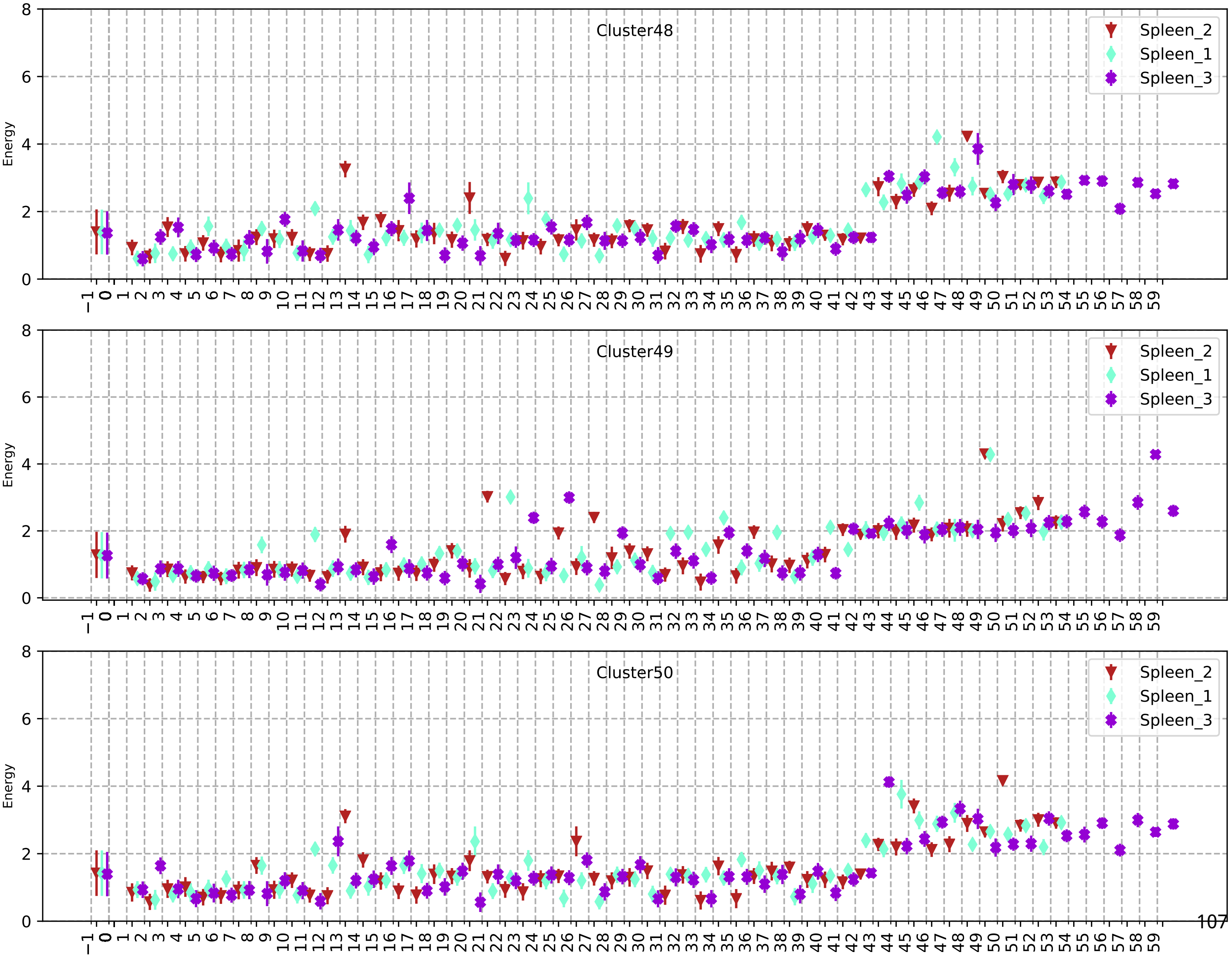
Cluster48
Cluster49
Cluster50
Spleen_2
Spleen_1
Spleen_3
Energy

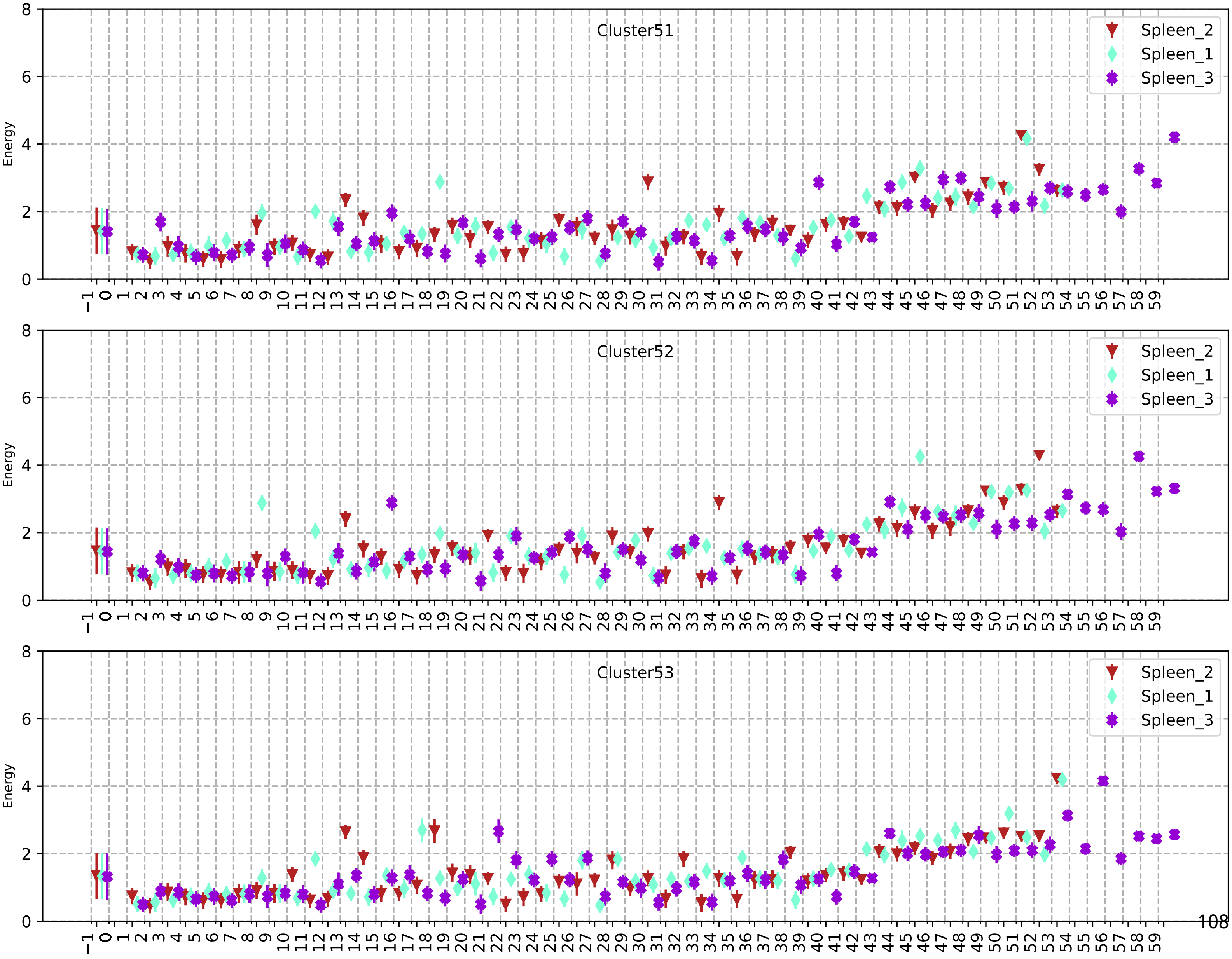
Cluster51
Cluster52
Cluster53
Energy
Spleen_2
Spleen_1
Spleen_3

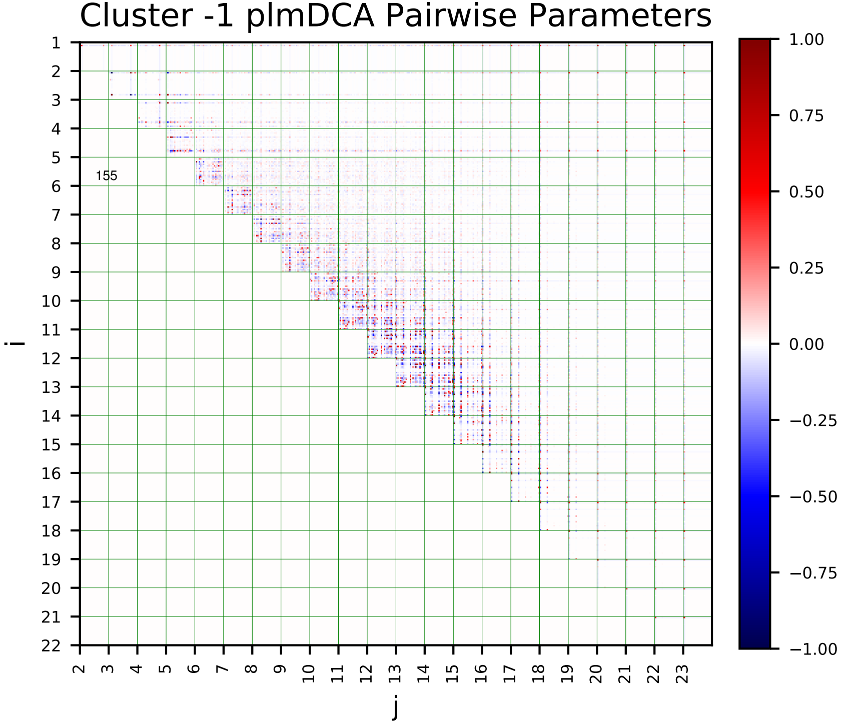
Cluster -1 plmDCA Pairwise Parameters
155
i
j
1.00
0.75
0.50
0.25
0.00
−0.25
−0.50
−0.75
−1.00

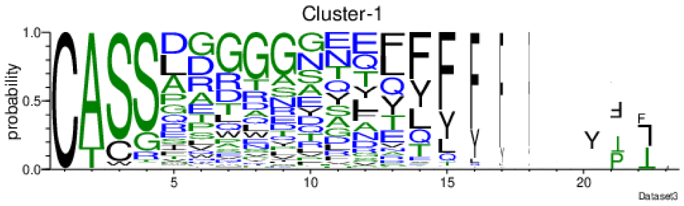
Cluster-1
probability
Dataset3

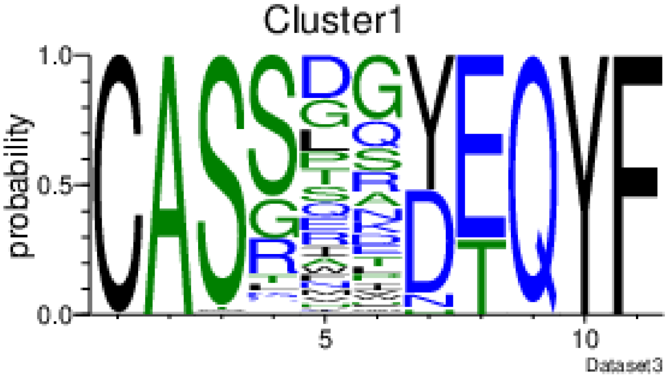
Cluster1
probability
Dataset3

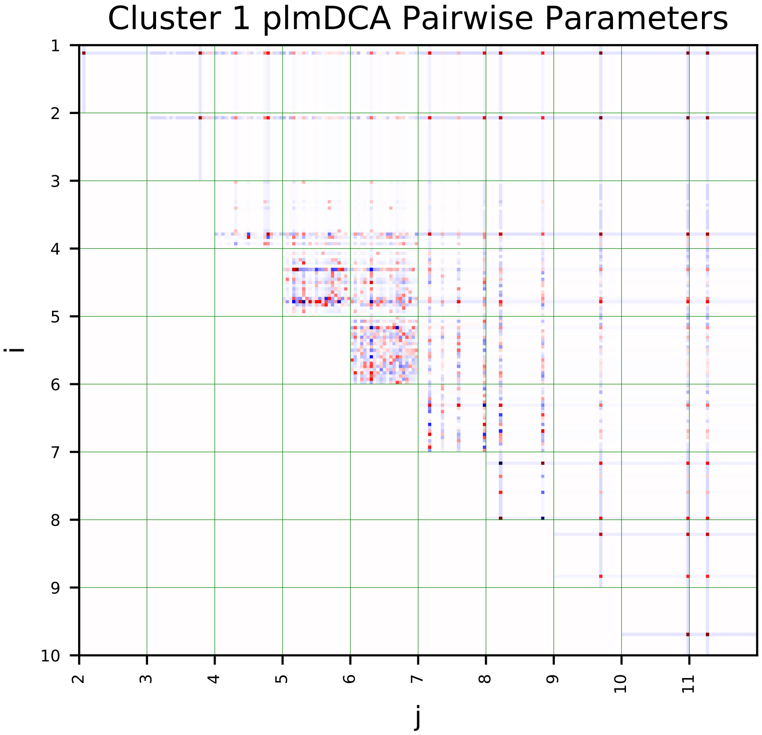
Cluster 1 plmDCA Pairwise Parameters
i
j

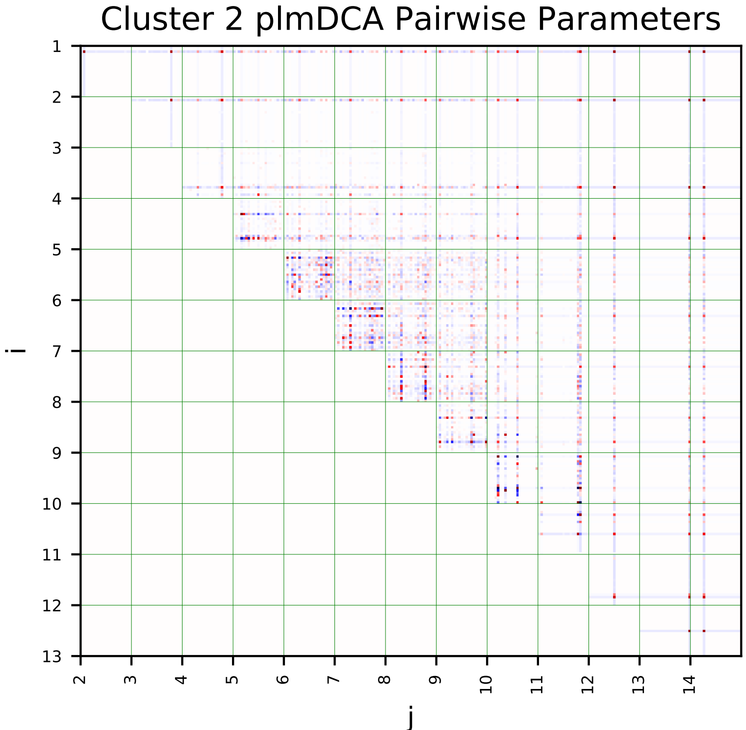
Cluster 2 plmDCA Pairwise Parameters
i
j

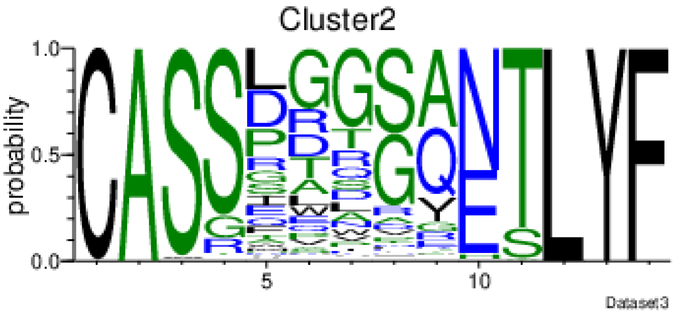
Cluster2
probability
Dataset3

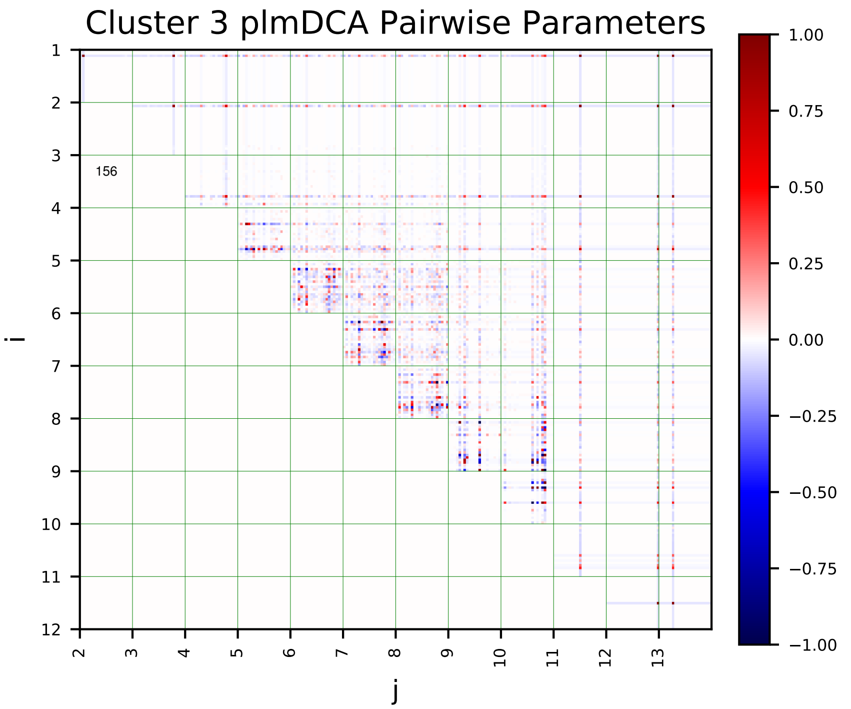
Cluster 3 plmDCA Pairwise Parameters
156
i
j

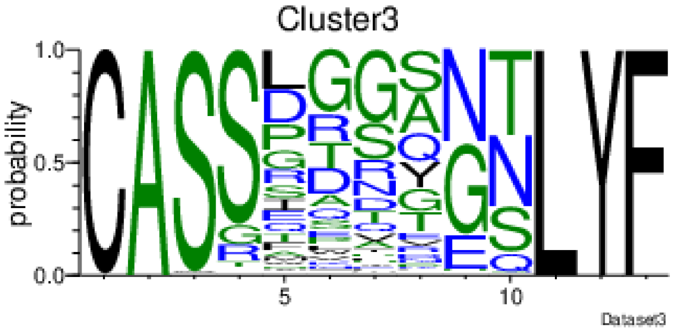
Cluster3
probability
Dataset3

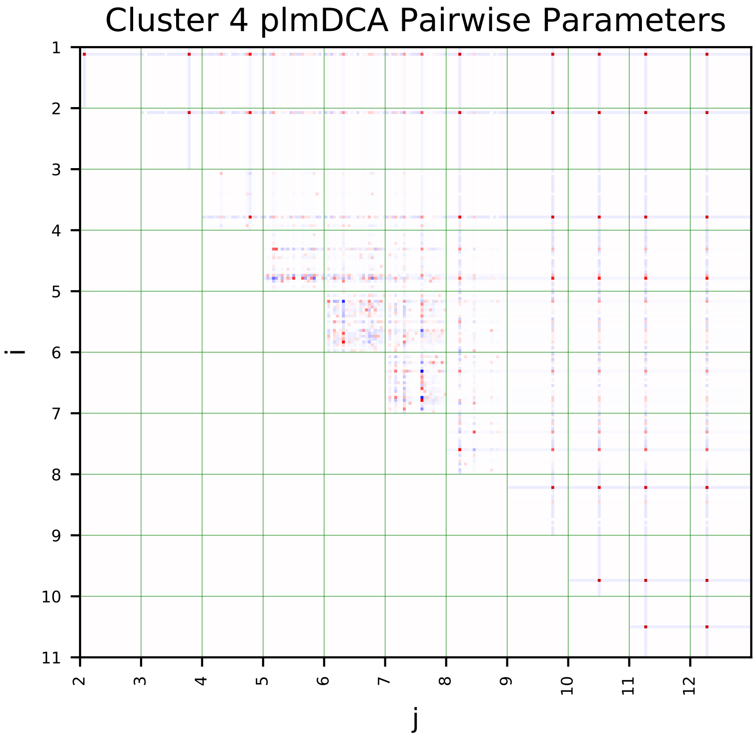
Cluster 4 plmDCA Pairwise Parameters
i
j

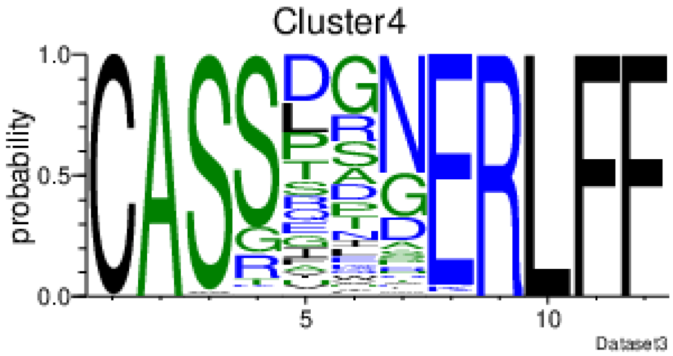
Cluster4
probability
Dataset3

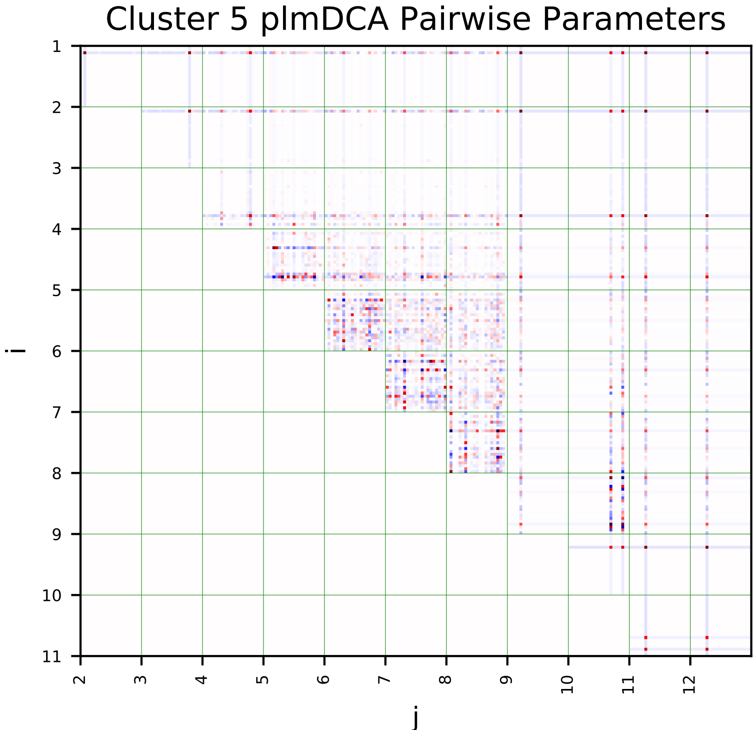
Cluster 5 plmDCA Pairwise Parameters
i
j

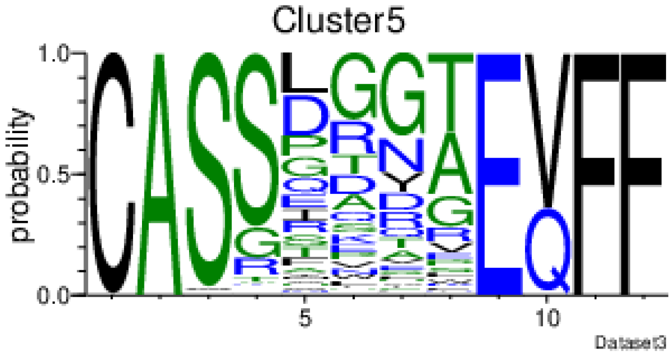
Cluster5
probability
Dataset3

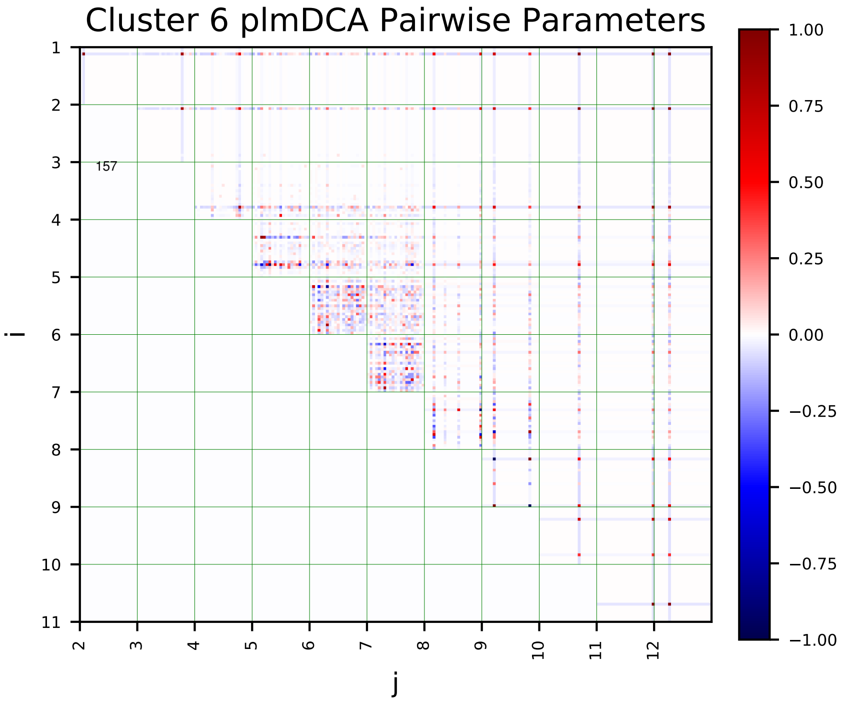
Cluster 6 plmDCA Pairwise Parameters
157
i
j

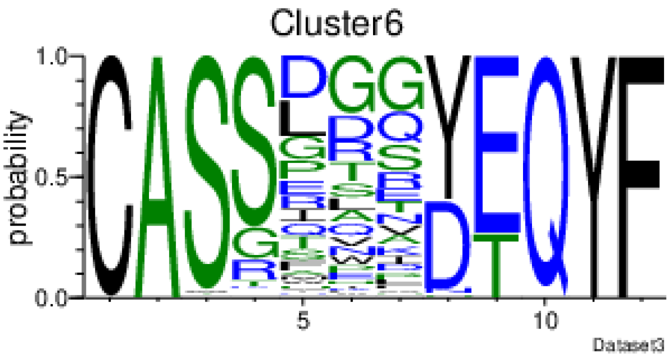
Cluster6
probability
Dataset3

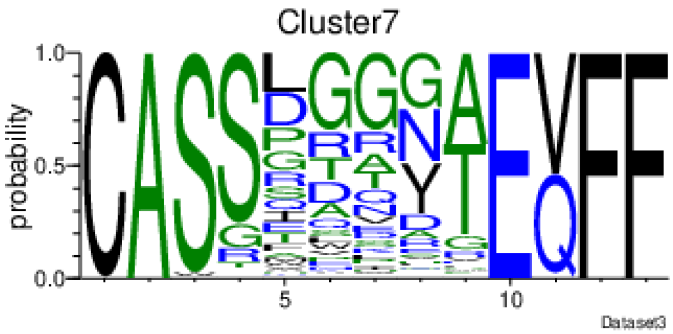
Cluster7
probability
Dataset3

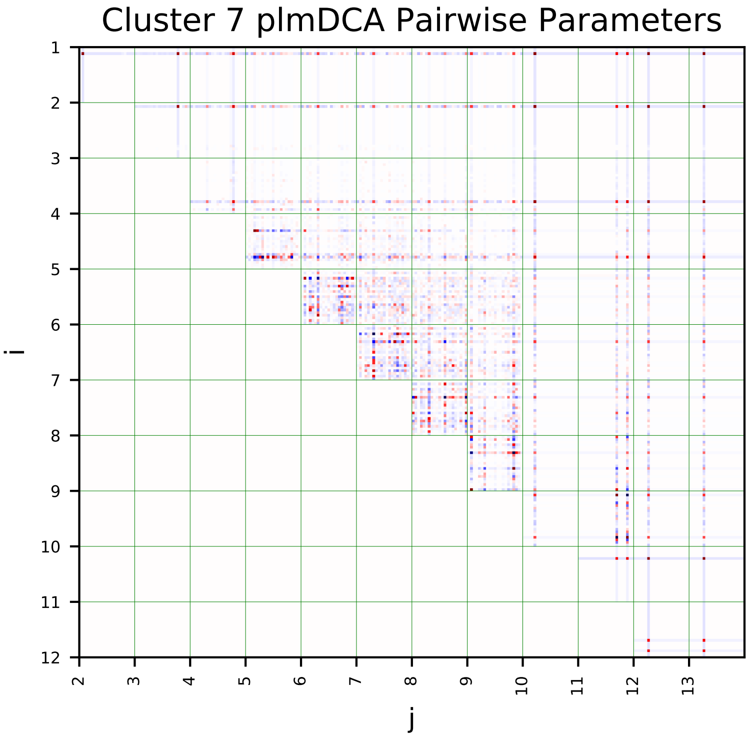
Cluster 7 plmDCA Pairwise Parameters
i
j

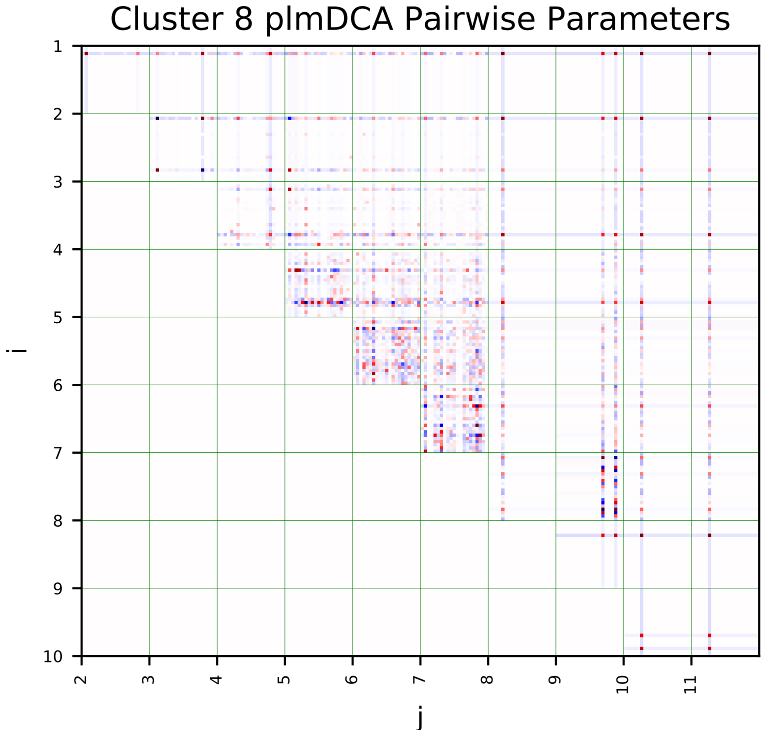
Cluster 8 plmDCA Pairwise Parameters
i
j

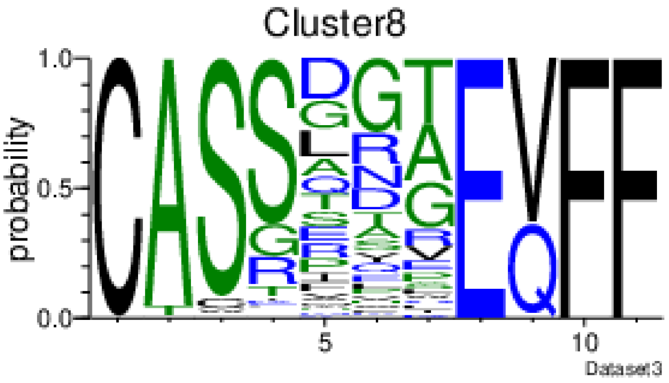
Cluster8
probability
Dataset3

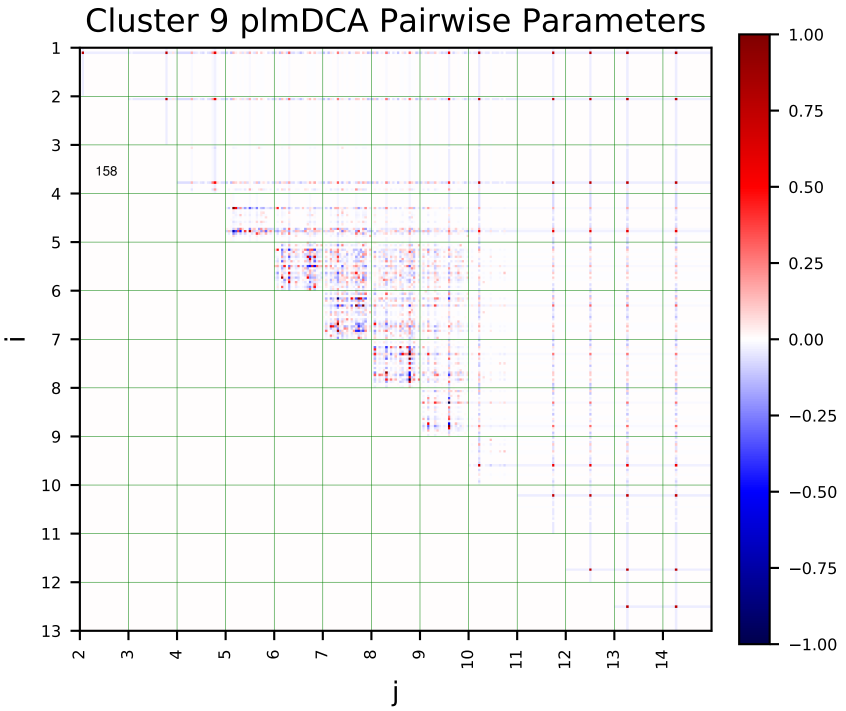
Cluster 9 plmDCA Pairwise Parameters
i
j
158
1.00
0.75
0.50
0.25
0.00
−0.25
−0.50
−0.75
−1.00

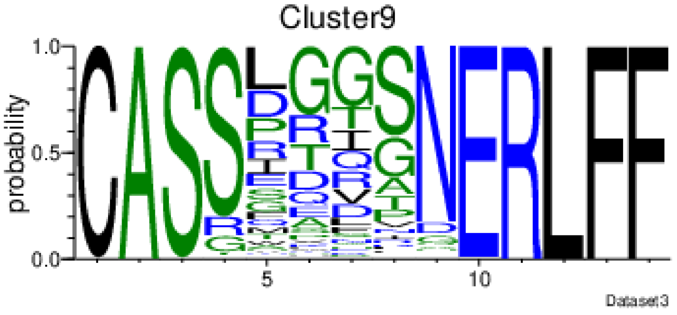
Cluster9
probability
Dataset3

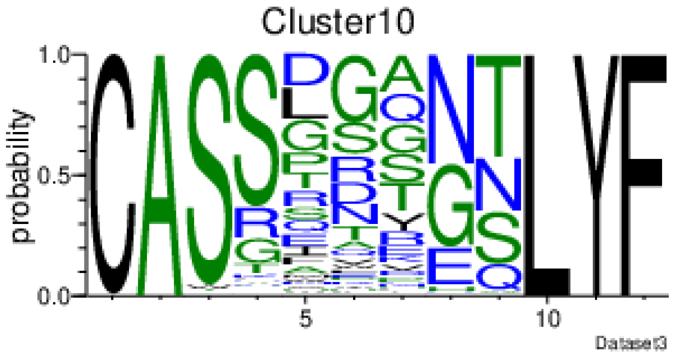
Cluster10
probability
Dataset3

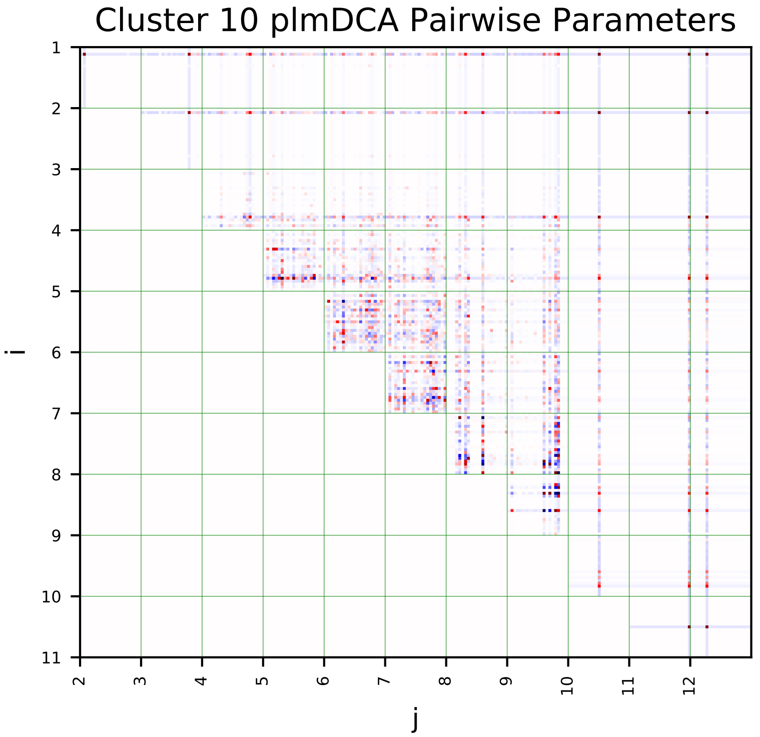
Cluster 10 plmDCA Pairwise Parameters
i
j

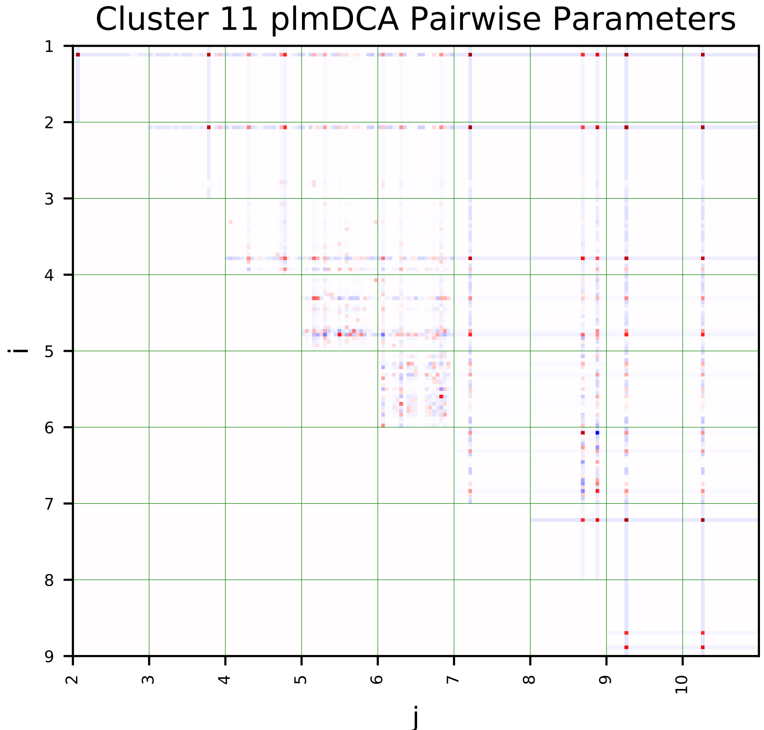
Cluster 11 plmDCA Pairwise Parameters
i
j

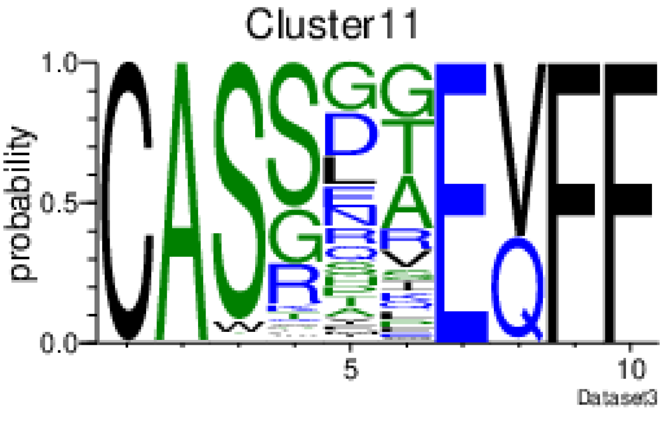
Cluster11
probability
Dataset3

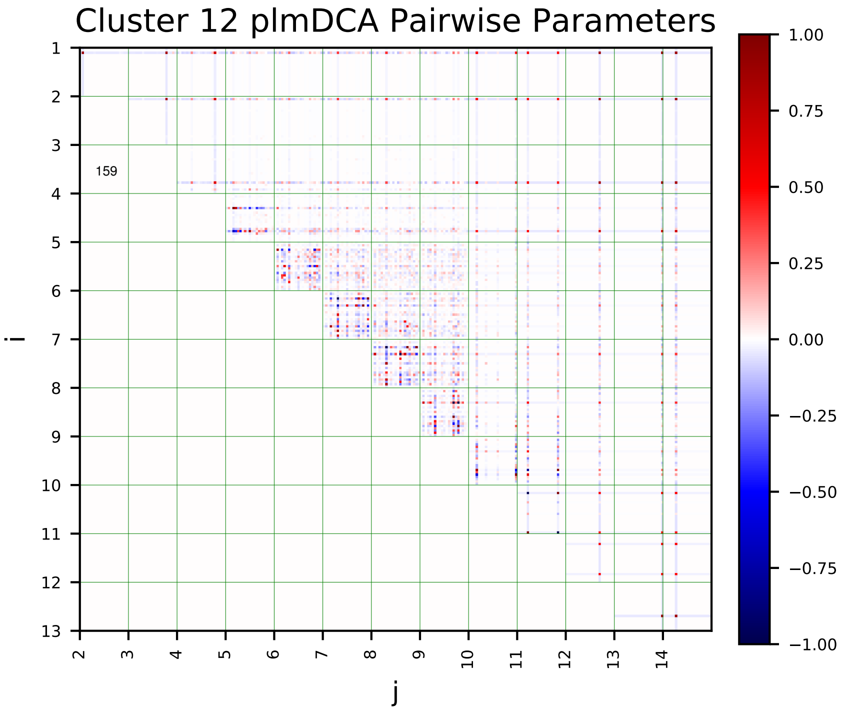
Cluster 12 plmDCA Pairwise Parameters
i
j
159

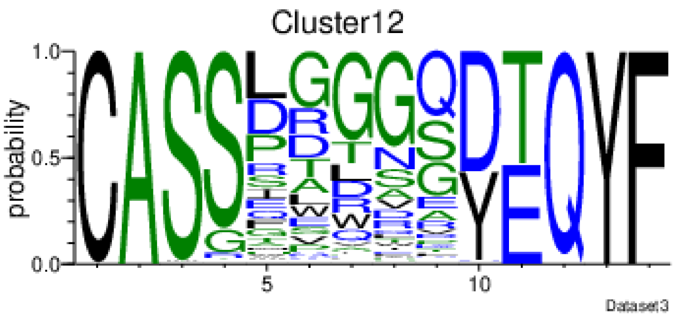
Cluster12
probability
Dataset3

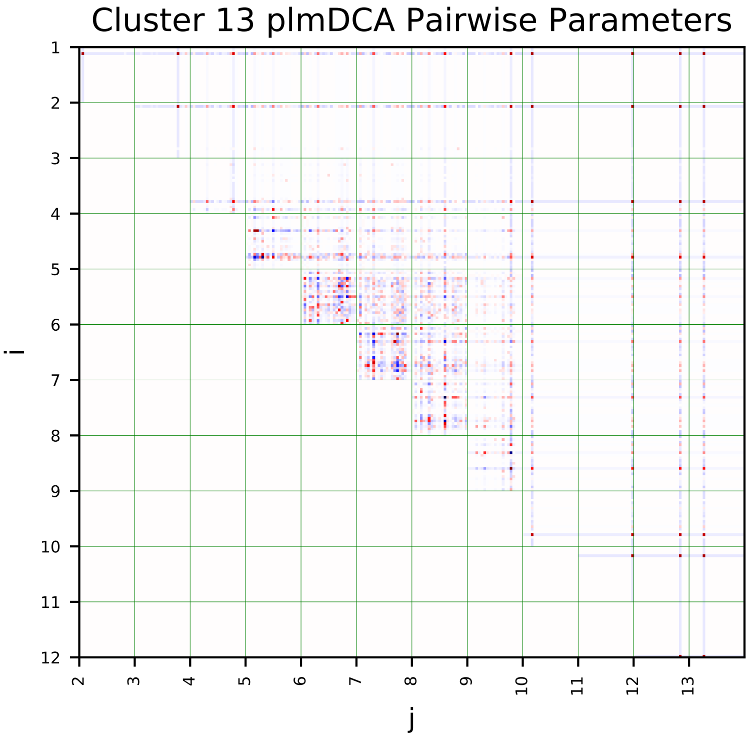
Cluster 13 plmDCA Pairwise Parameters
i
j

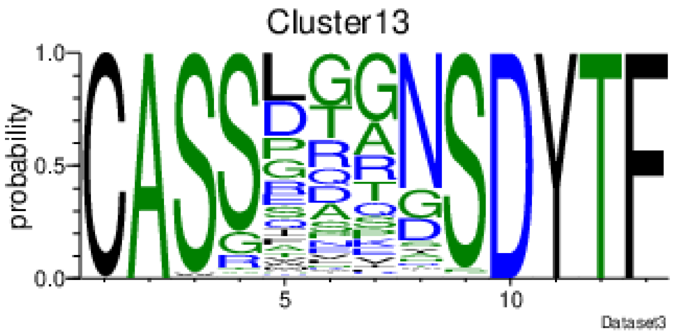
Cluster13
probability
Dataset3

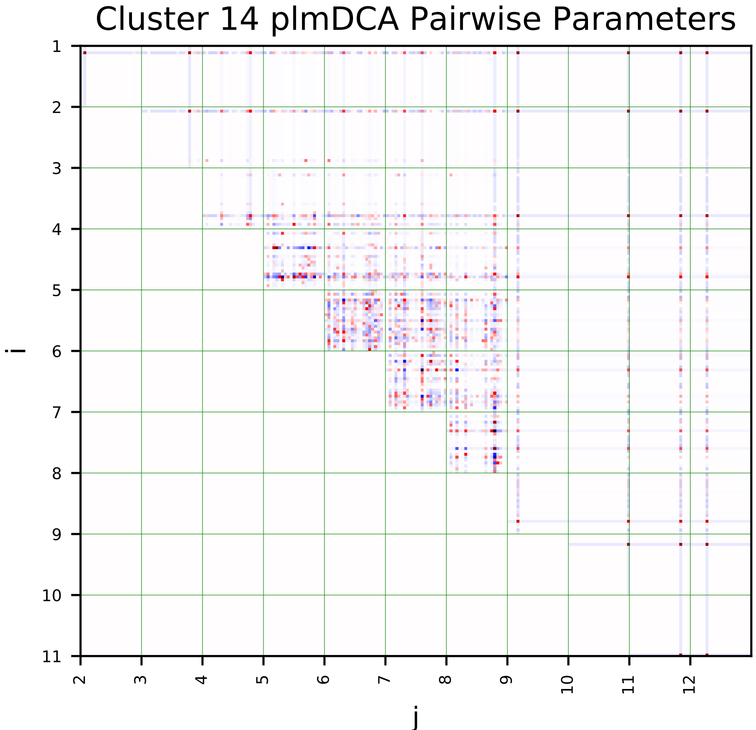
Cluster 14 plmDCA Pairwise Parameters
i
j

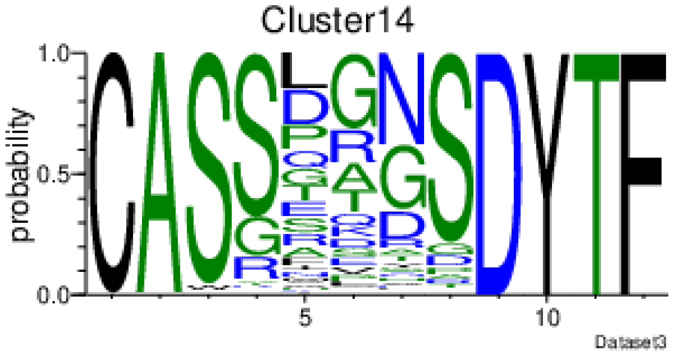
Cluster14
probability
Dataset3

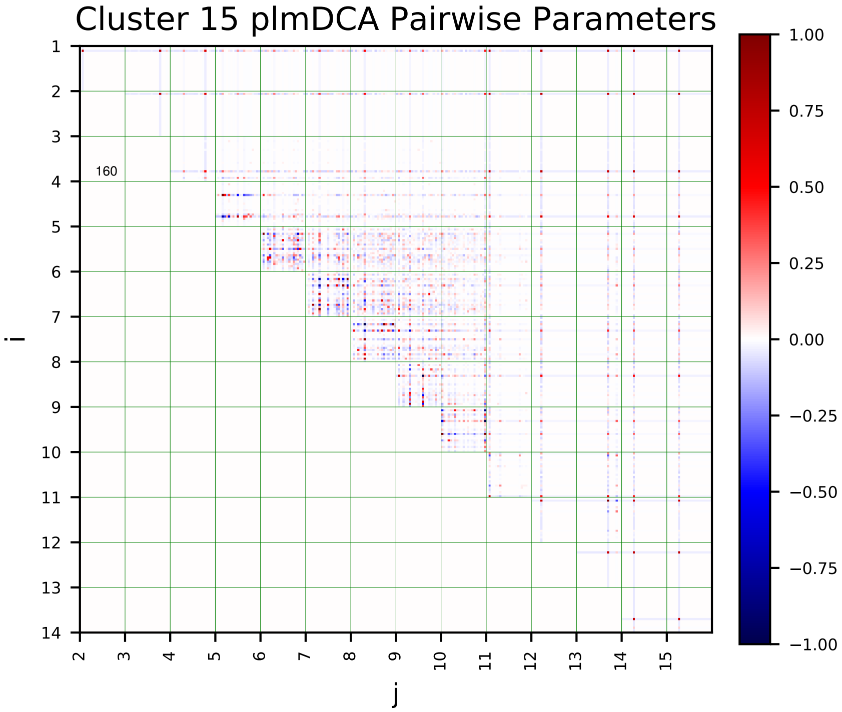
Cluster 15 plmDCA Pairwise Parameters
160
i
j

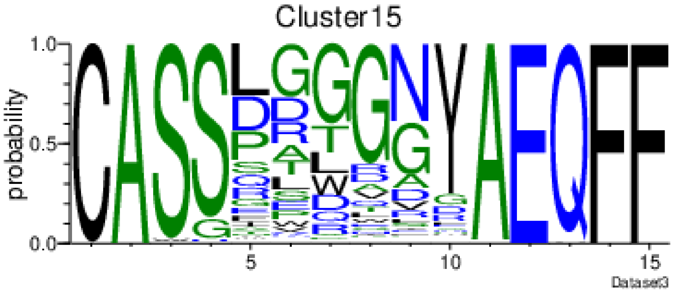
Cluster15
probability
Dataset3

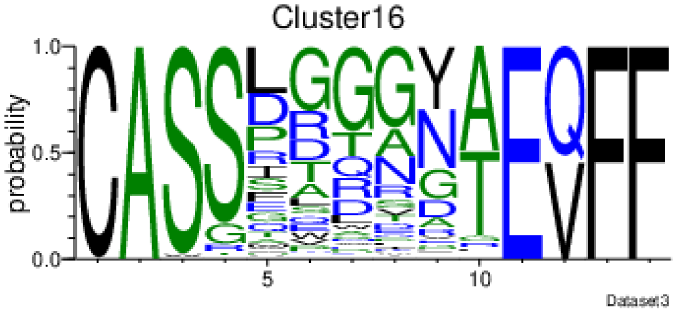
Cluster16
probability
Dataset3

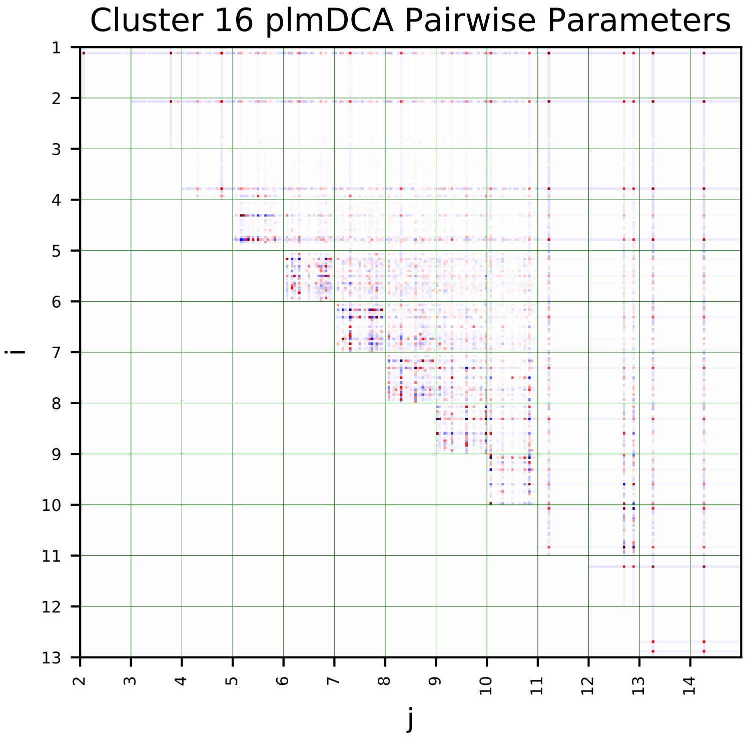
Cluster 16 plmDCA Pairwise Parameters
i
j

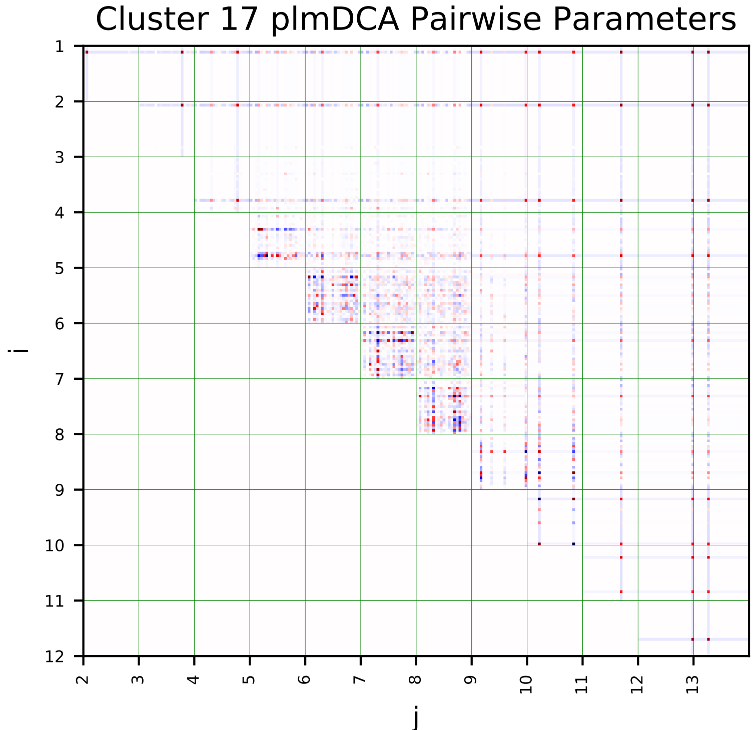
Cluster 17 plmDCA Pairwise Parameters
i
j

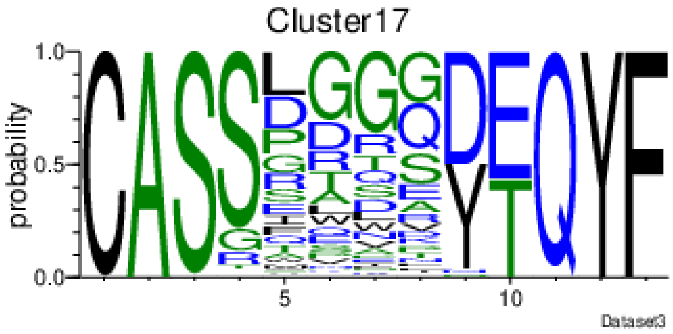
Cluster17
probability
Dataset3

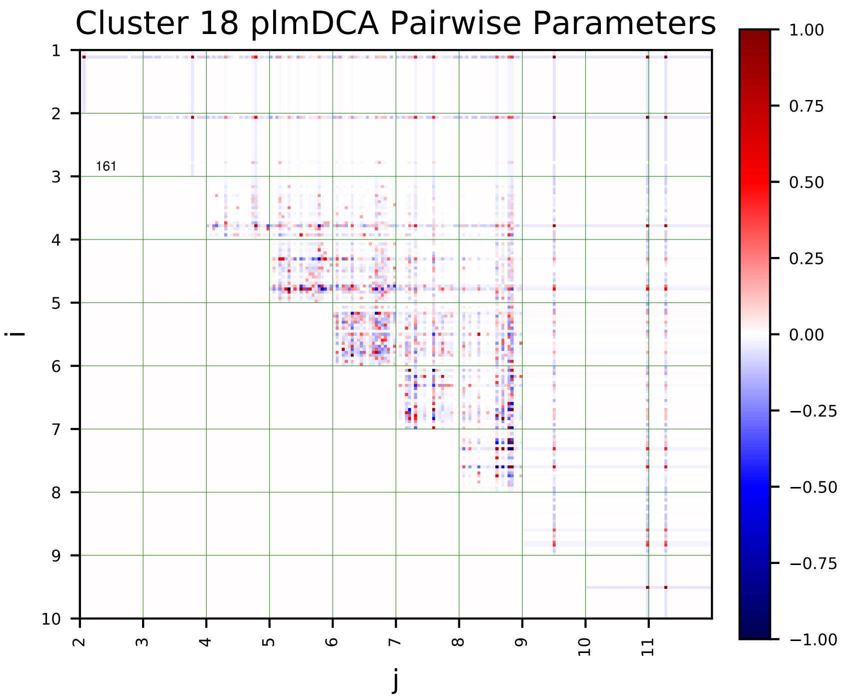
Cluster 18 plmDCA Pairwise Parameters
i
j
161
1.00
0.75
0.50
0.25
0.00
−0.25
−0.50
−0.75
−1.00

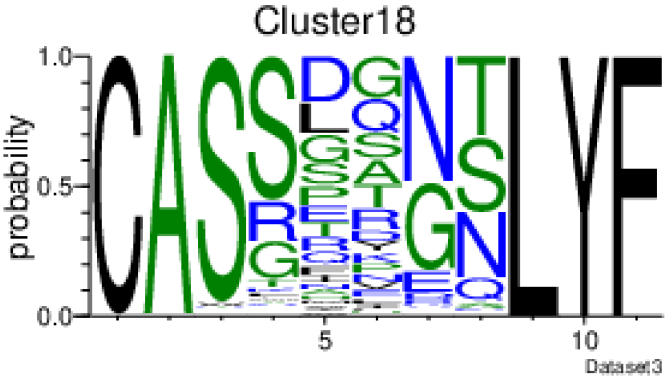
Cluster18
probability
Dataset3

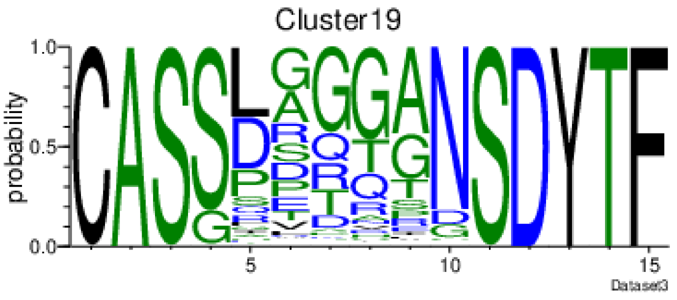
Cluster19
probability
Dataset3

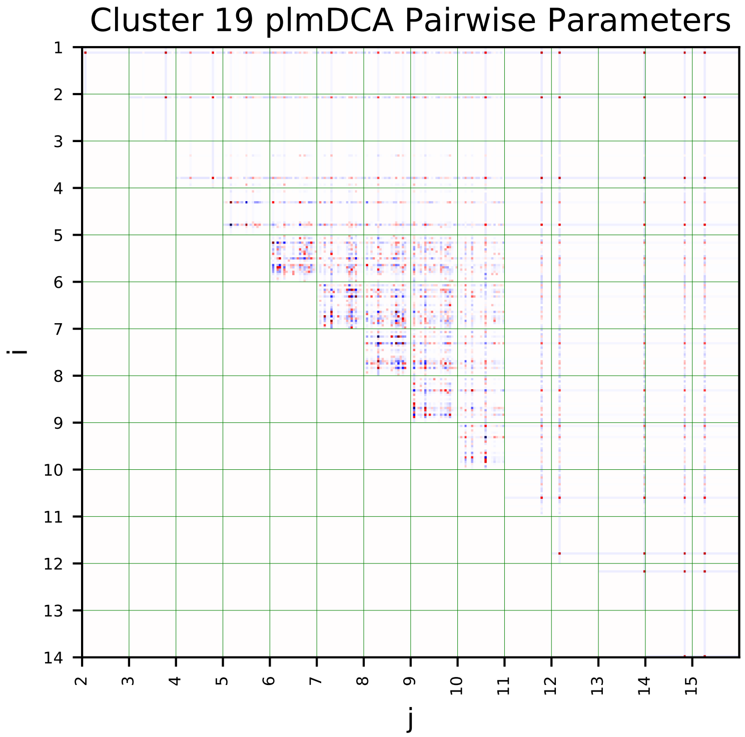
Cluster 19 plmDCA Pairwise Parameters
i
j

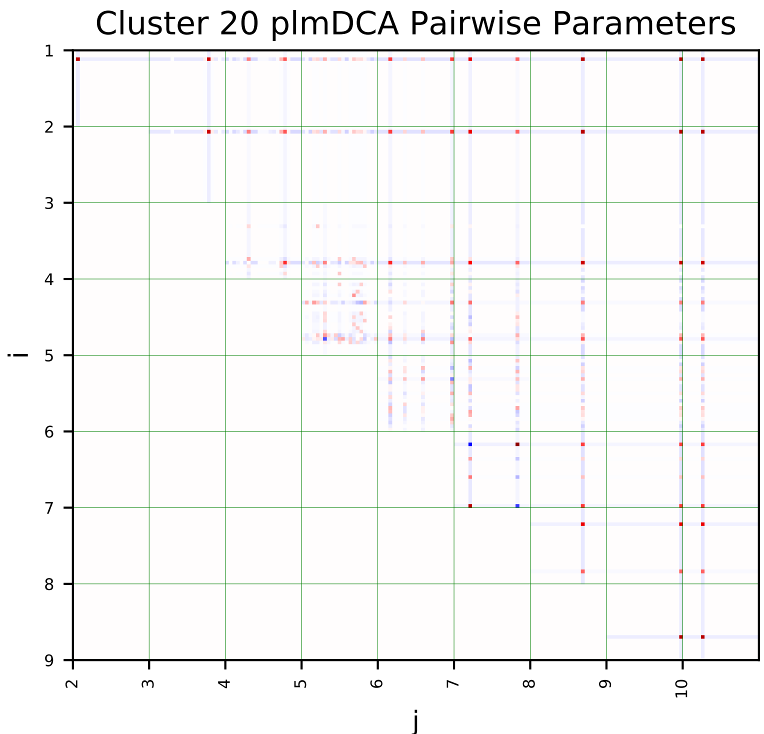
Cluster 20 plmDCA Pairwise Parameters
i
j

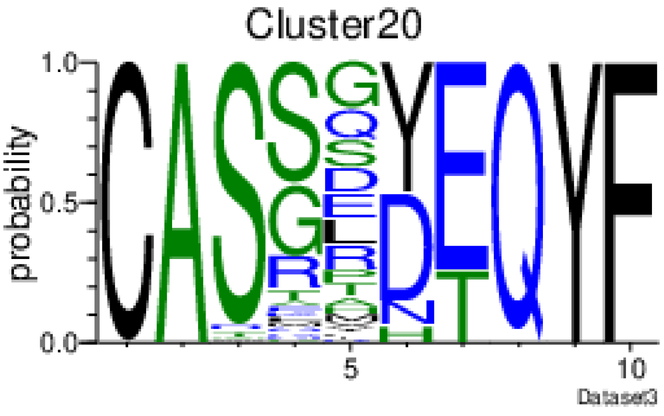
Cluster20
probability
Dataset3

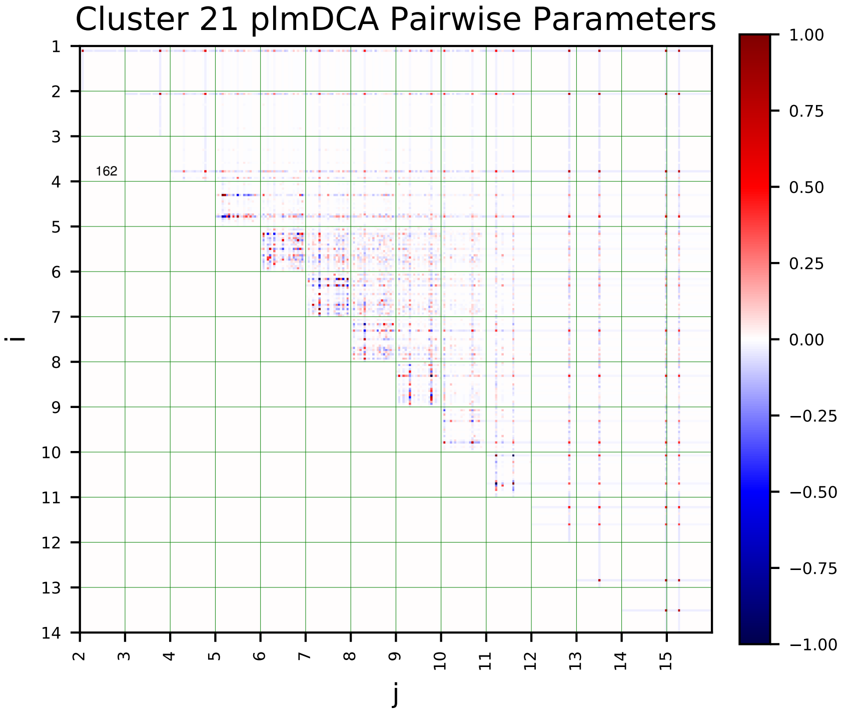
Cluster 21 plmDCA Pairwise Parameters
162
i
j
1.00
0.75
0.50
0.25
0.00
−0.25
−0.50
−0.75
−1.00

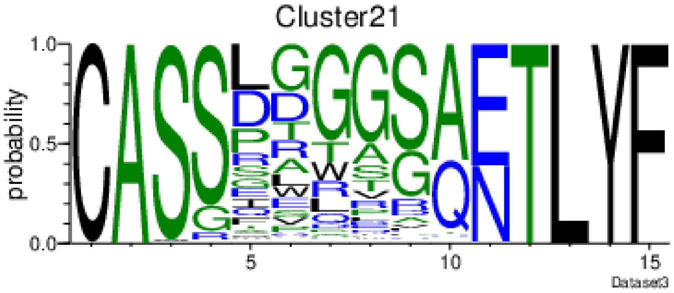
Cluster21
probability
Dataset3

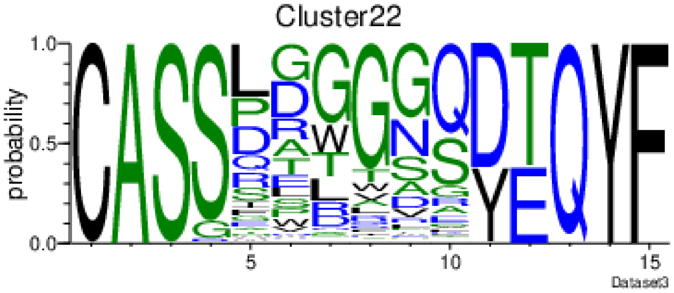
Cluster22
probability
Dataset3

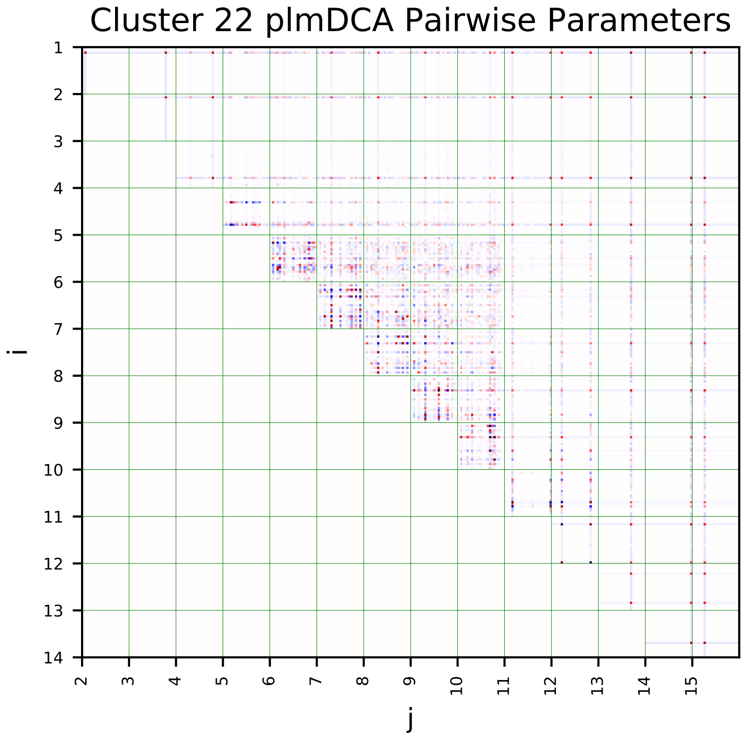
Cluster 22 plmDCA Pairwise Parameters
i
j

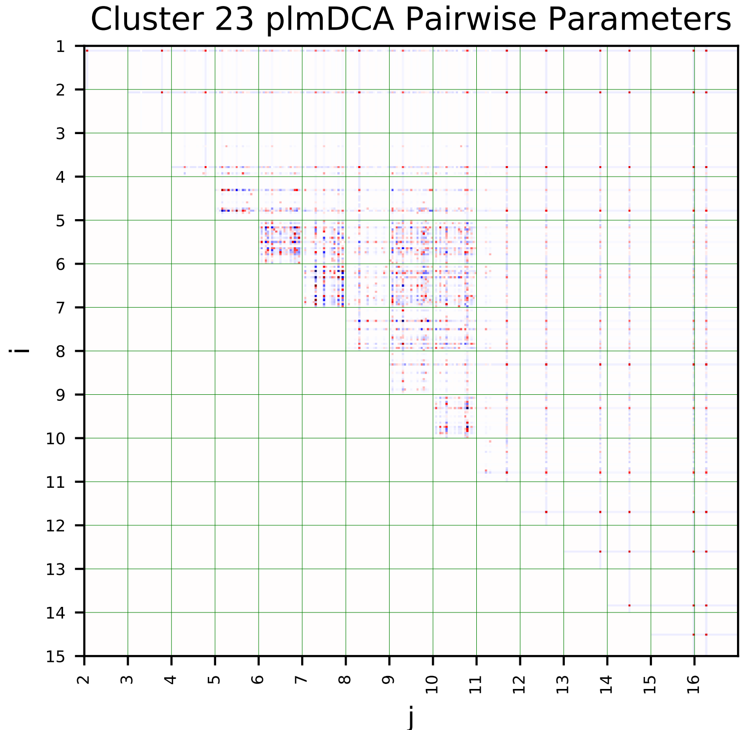
Cluster 23 plmDCA Pairwise Parameters
i
j

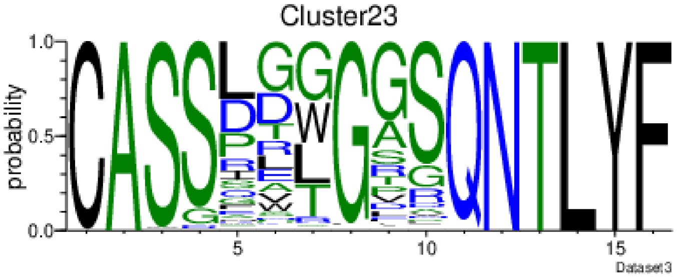
Cluster23
probability
Dataset3

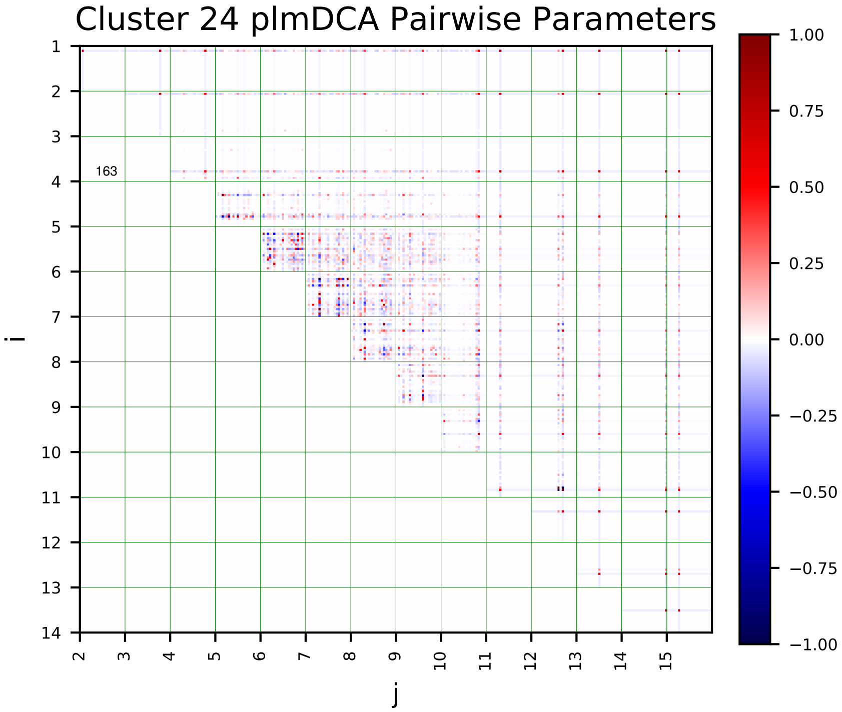
Cluster 24 plmDCA Pairwise Parameters
i
j
163

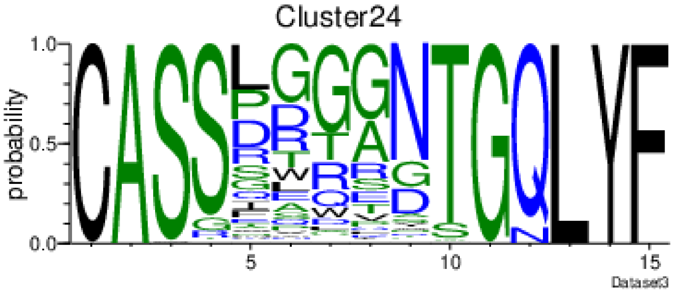
Cluster24
probability
Dataset3

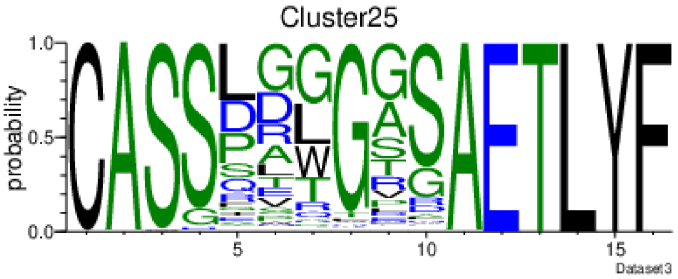
Cluster25
probability
Dataset3

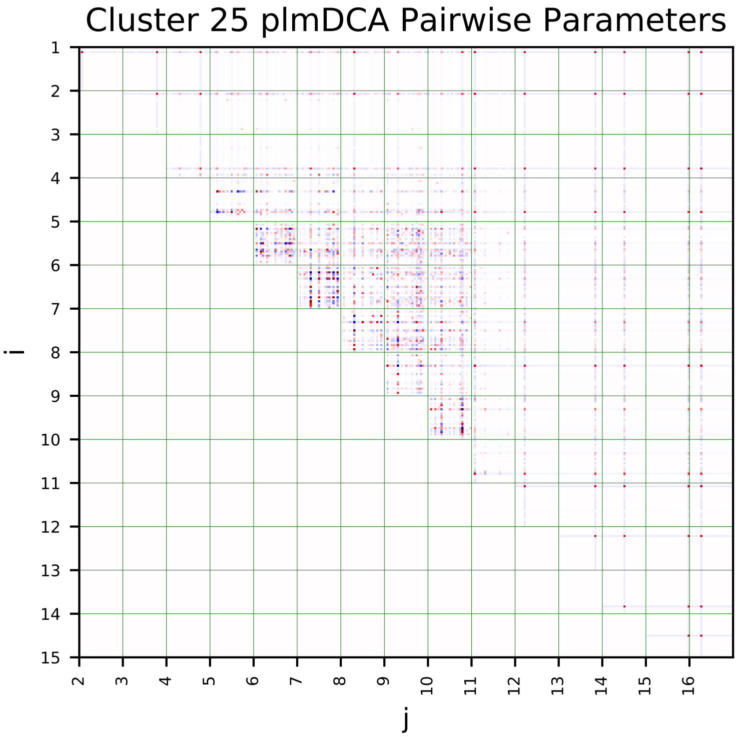
Cluster 25 plmDCA Pairwise Parameters
i
j

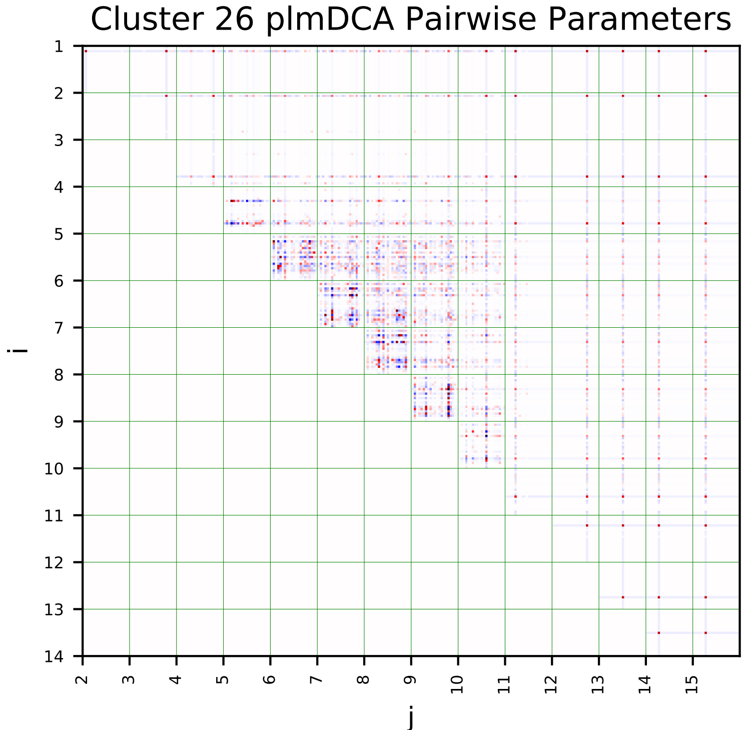
Cluster 26 plmDCA Pairwise Parameters
i
j

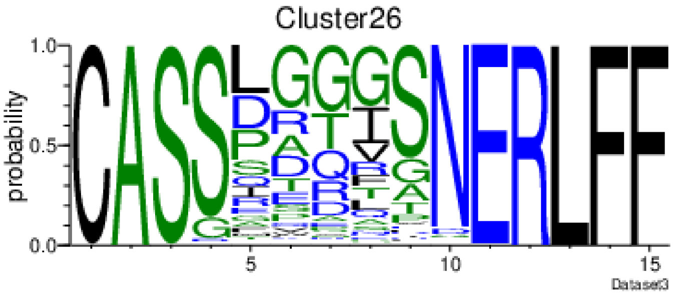
Cluster26
probability
Dataset3

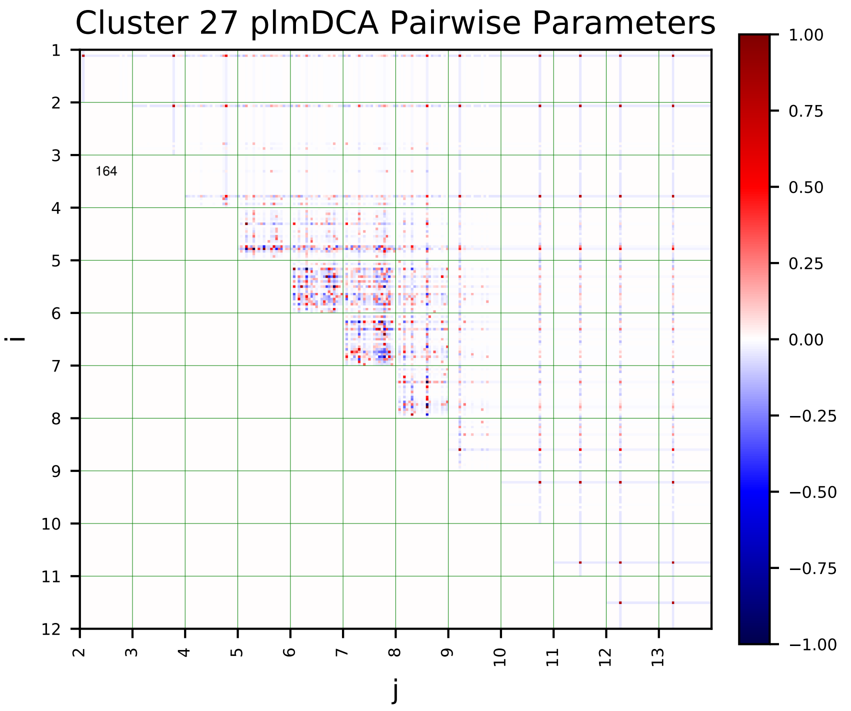
Cluster 27 plmDCA Pairwise Parameters
164
i
j
1.00
0.75
0.50
0.25
0.00
−0.25
−0.50
−0.75
−1.00

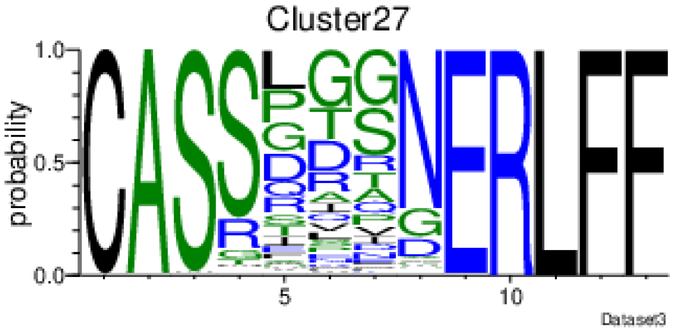
Cluster27
probability
Dataset3

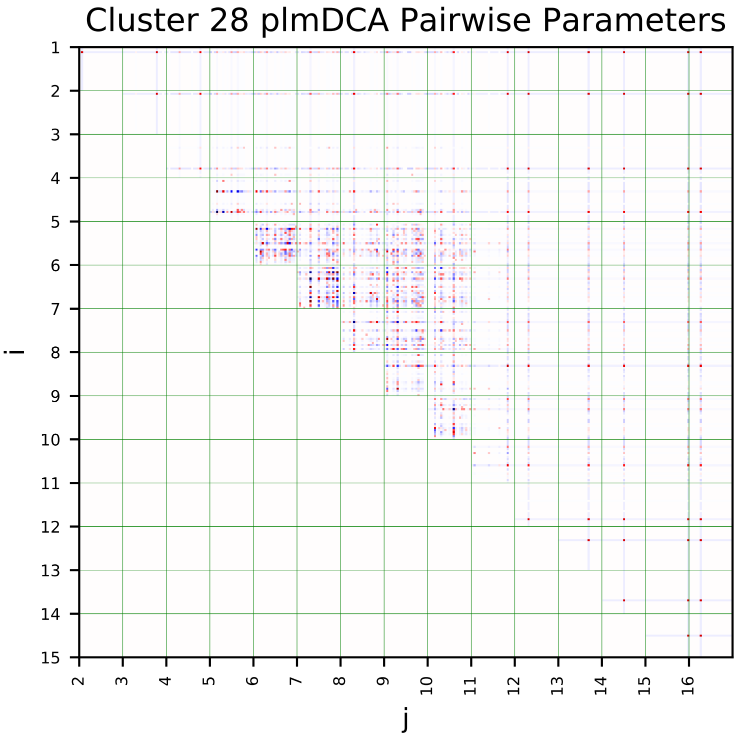
Cluster 28 plmDCA Pairwise Parameters
i
j

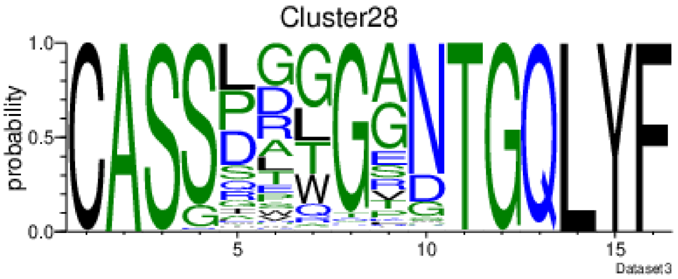
Cluster28
probability
Dataset3

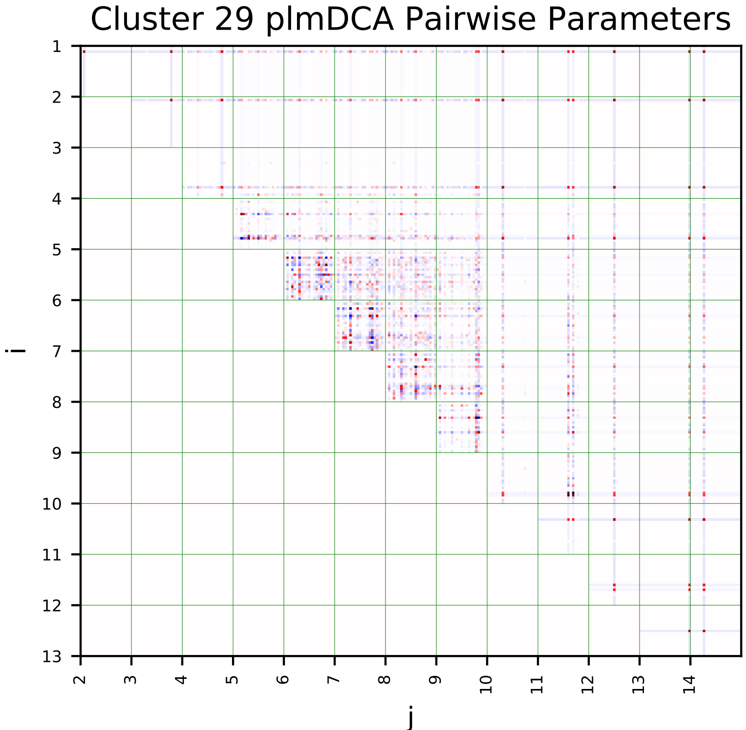
Cluster 29 plmDCA Pairwise Parameters
i
j

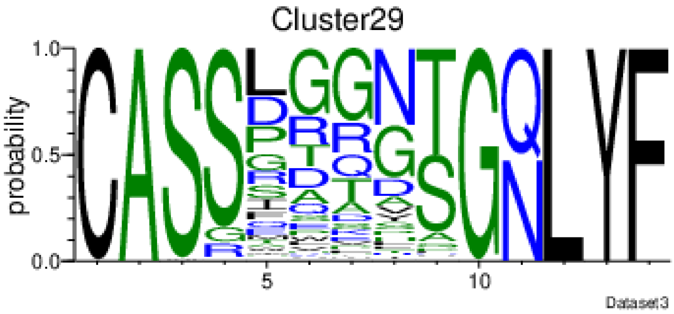
Cluster29
probability
Dataset3

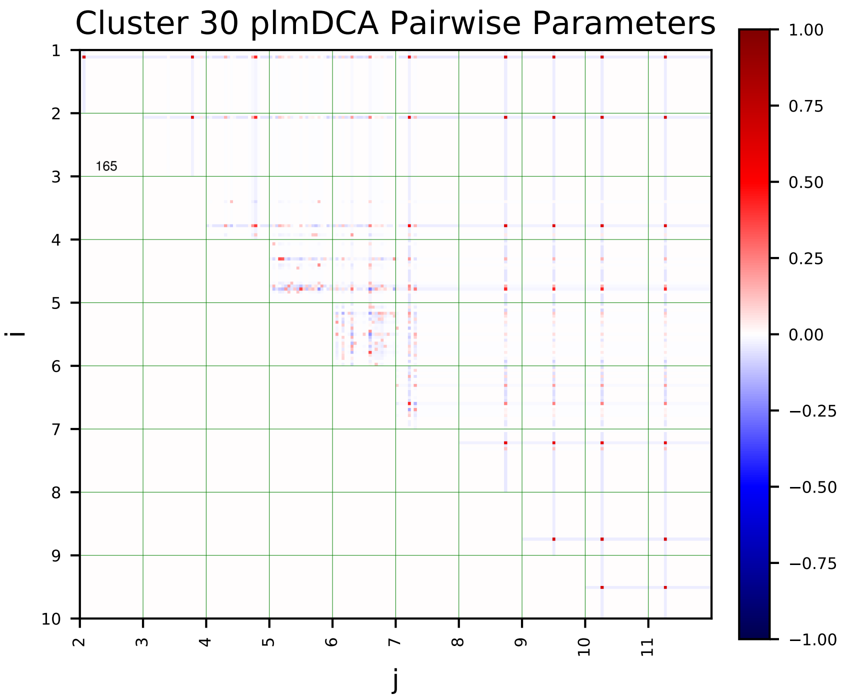
Cluster 30 plmDCA Pairwise Parameters
i
j
165
1.00
0.75
0.50
0.25
0.00
−0.25
−0.50
−0.75
−1.00

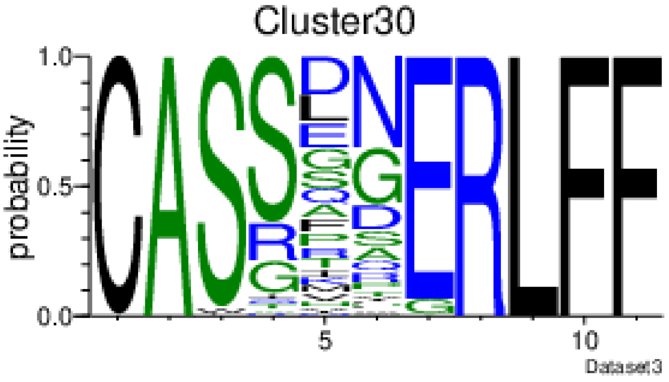
Cluster30
probability
Dataset3

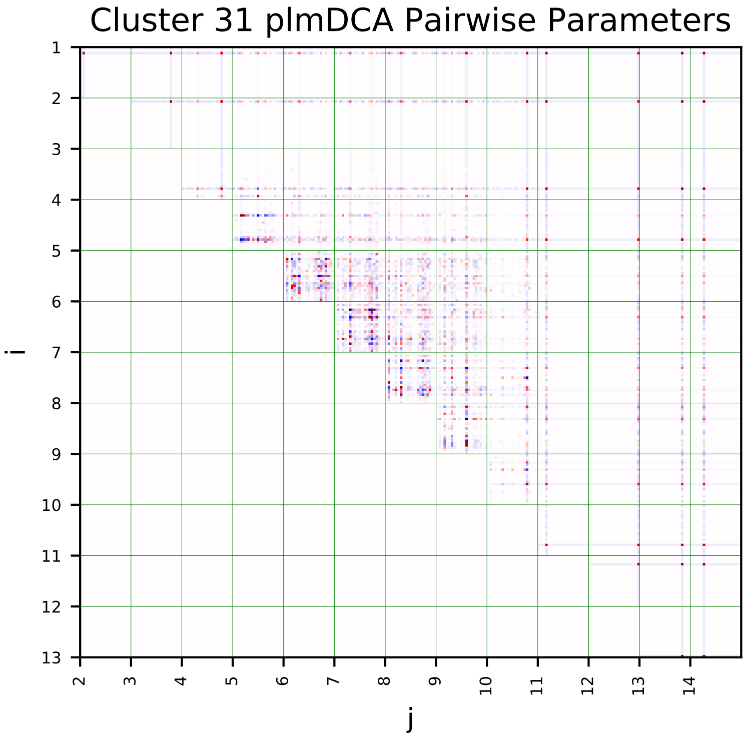
Cluster 31 plmDCA Pairwise Parameters
i
j

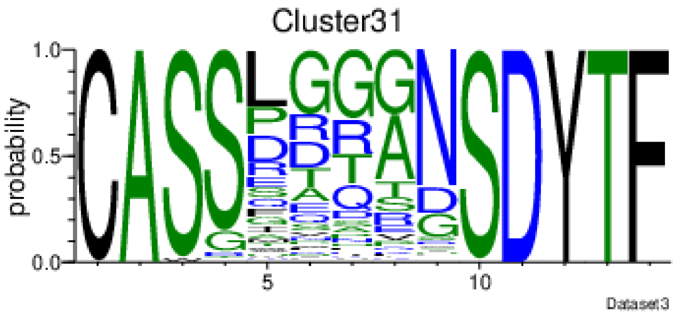
Cluster31
probability
Dataset3

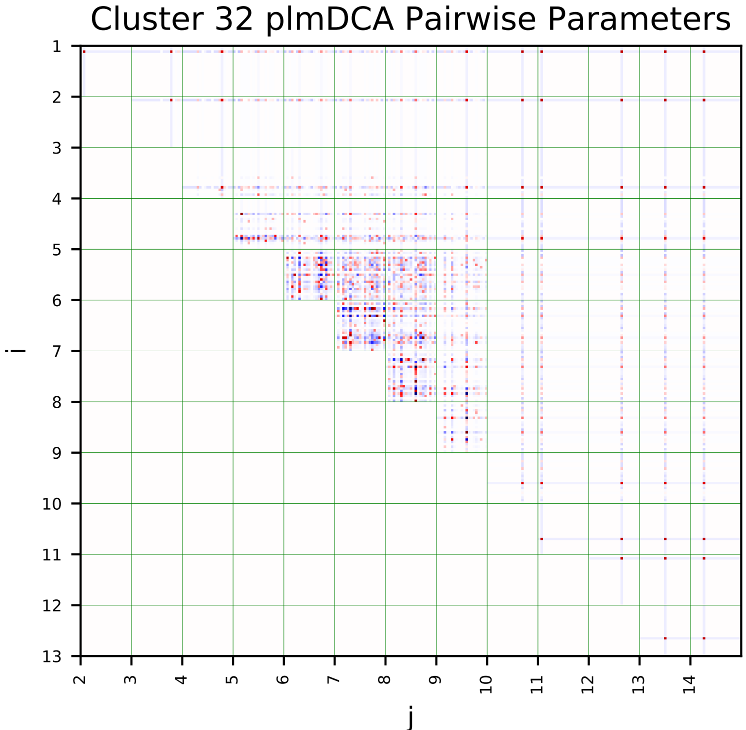
Cluster 32 plmDCA Pairwise Parameters
i
j

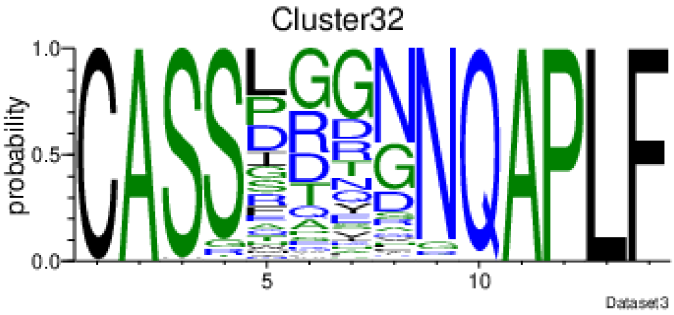
Cluster32
probability
Dataset3

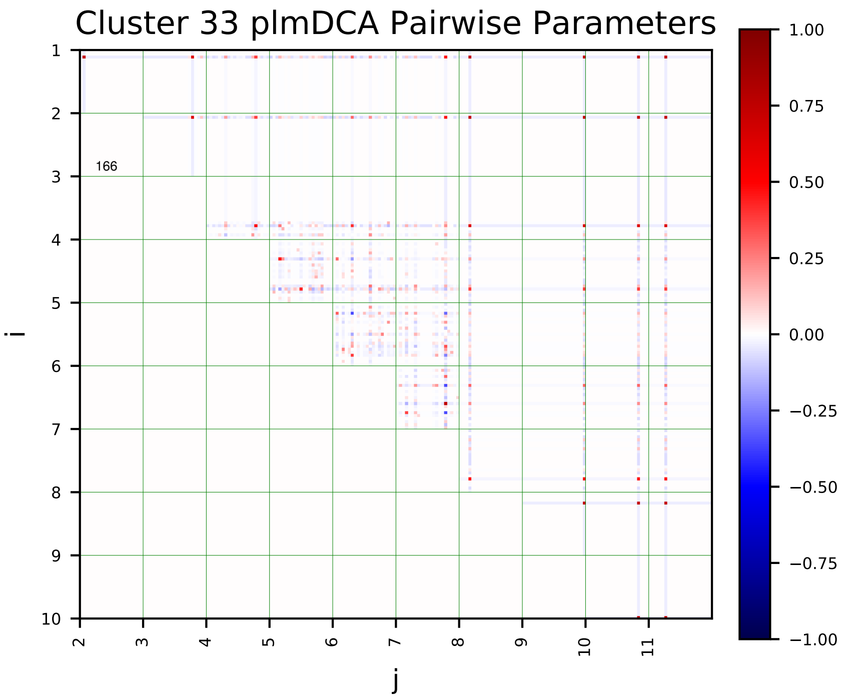
Cluster 33 plmDCA Pairwise Parameters
i
j
166

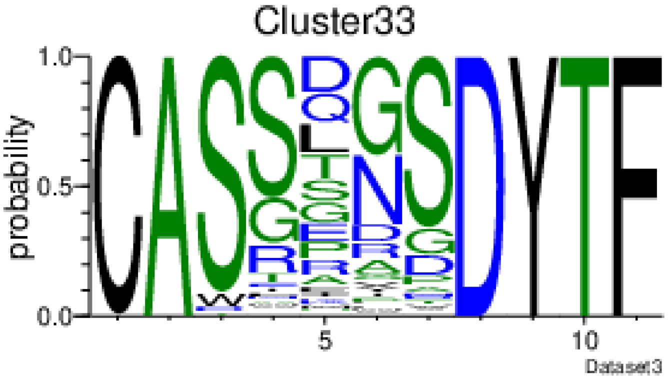
Cluster33
probability
Dataset3

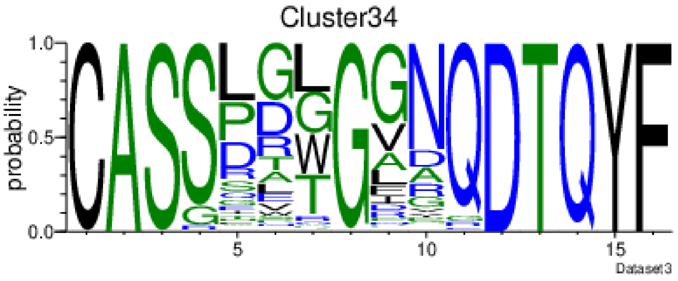
Cluster34
probability
Dataset3

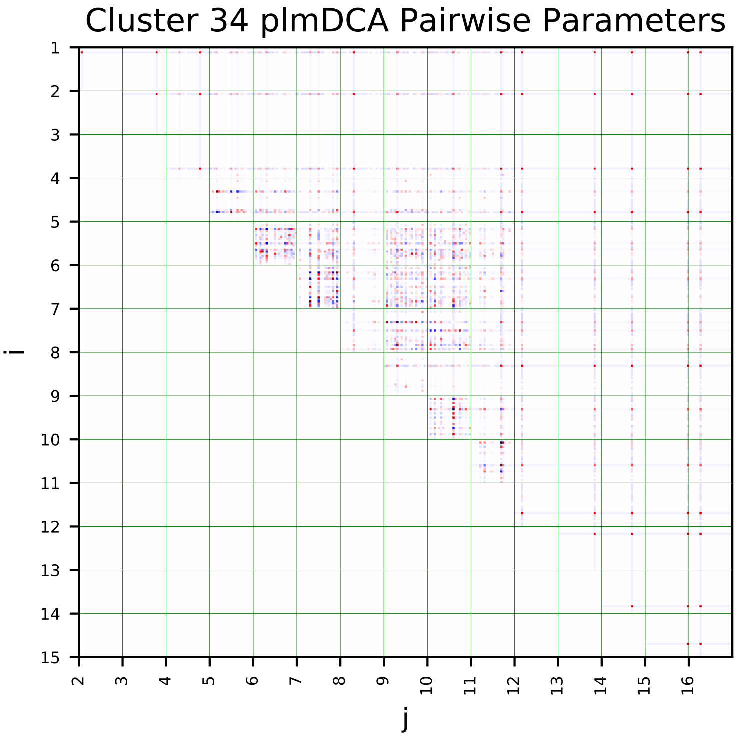
Cluster 34 plmDCA Pairwise Parameters
i
j

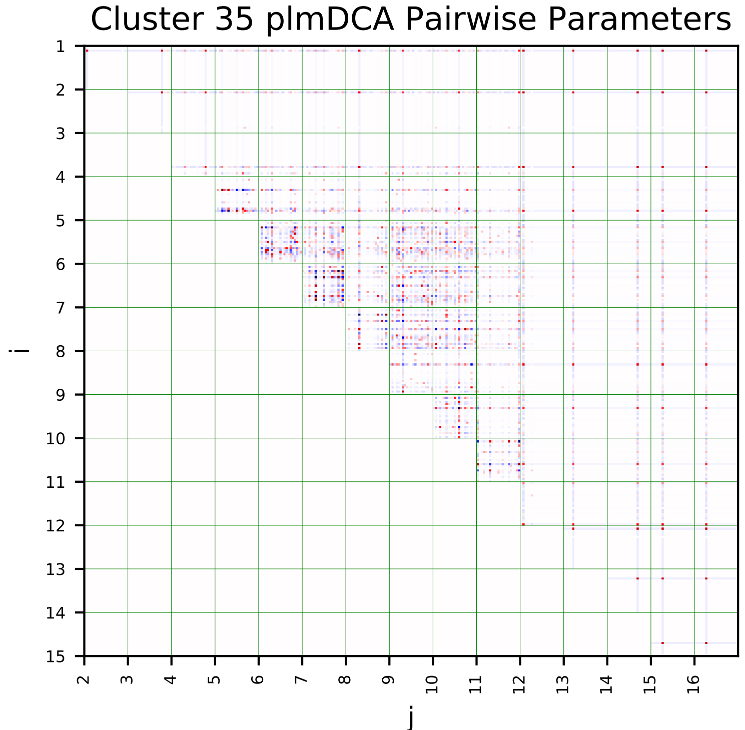
Cluster 35 plmDCA Pairwise Parameters
i
j

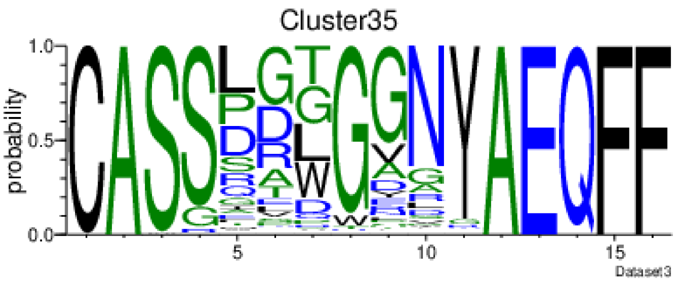
Cluster35
probability
Dataset3

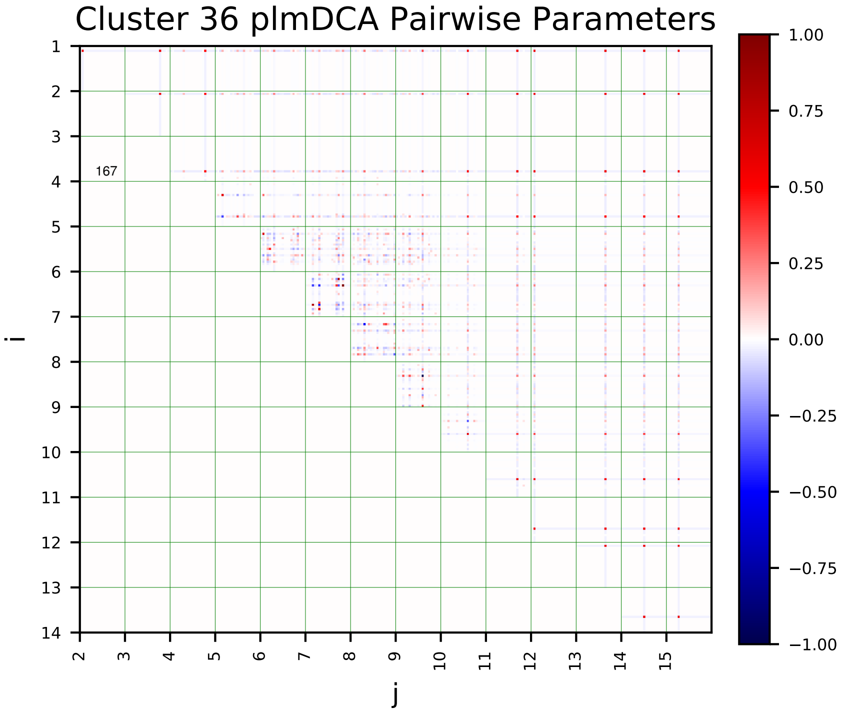
Cluster 36 plmDCA Pairwise Parameters
167
i
j

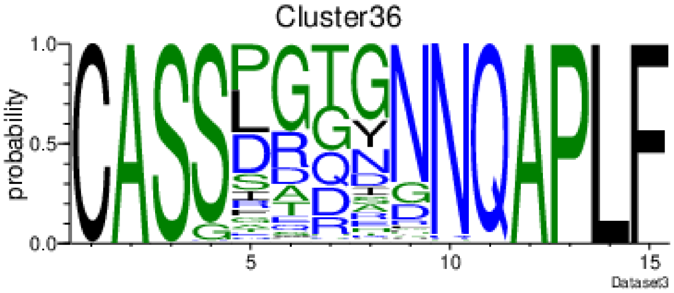
Cluster36
probability
Dataset3

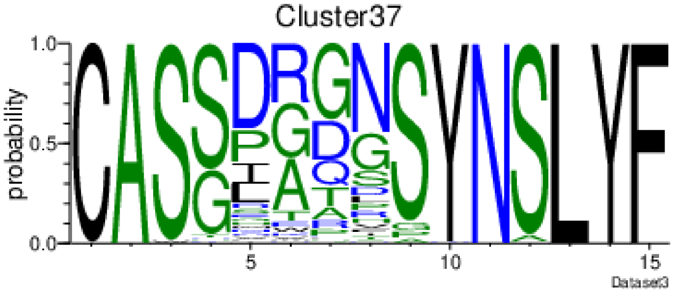
Cluster37
probability
Dataset3

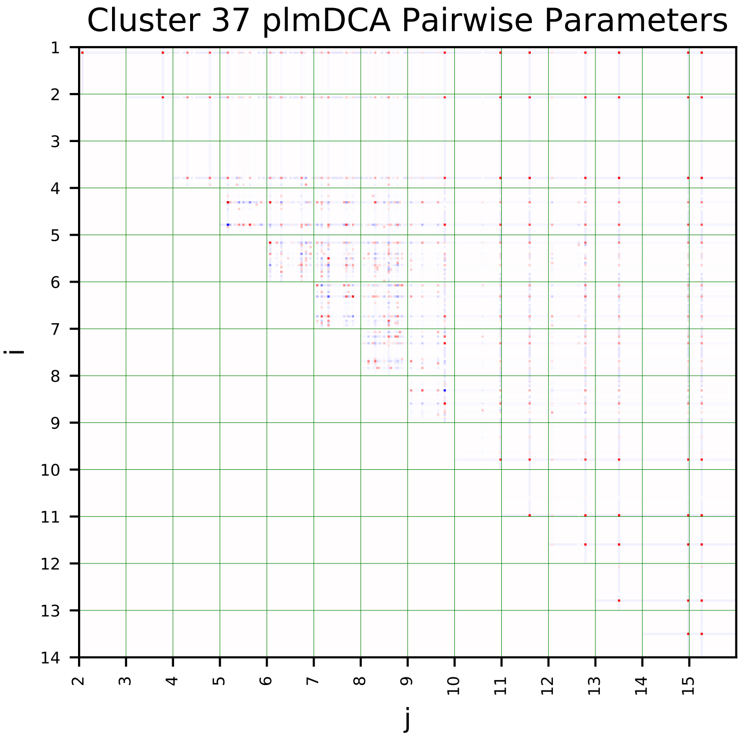
Cluster 37 plmDCA Pairwise Parameters
i
j

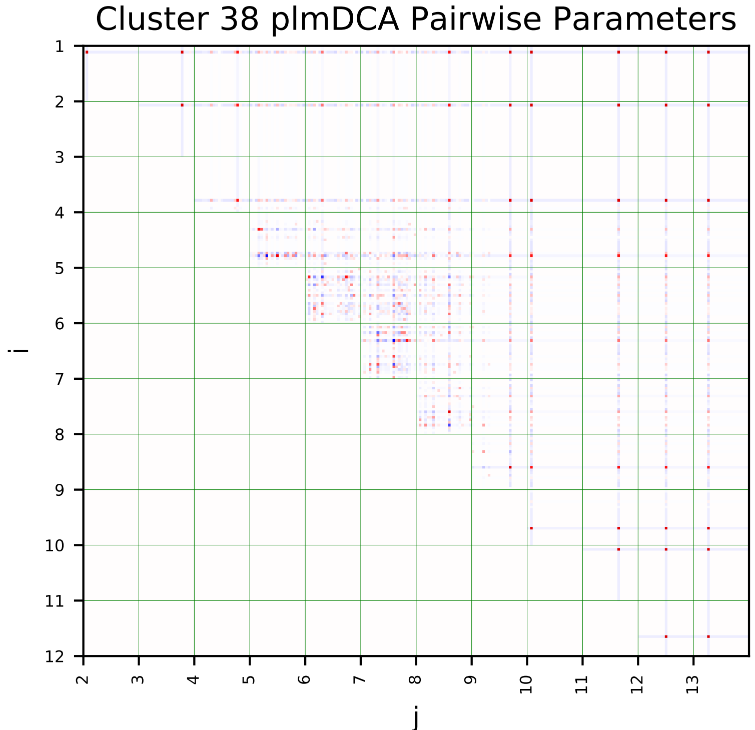
Cluster 38 plmDCA Pairwise Parameters
i
j

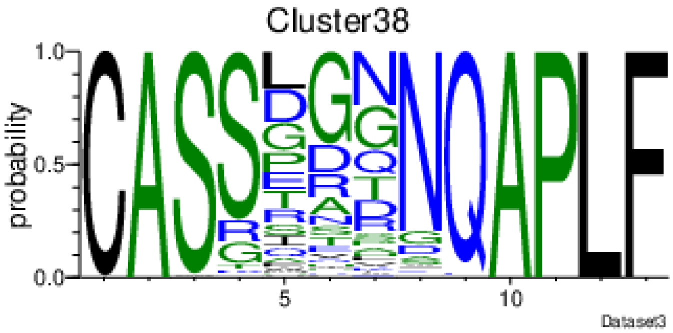
Cluster38
probability
Dataset3

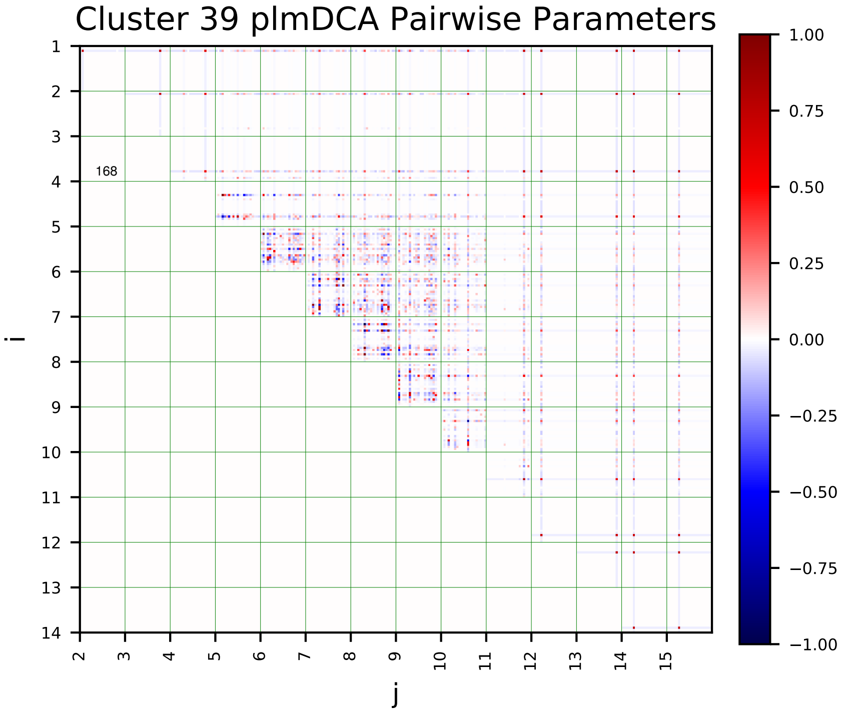
Cluster 39 plmDCA Pairwise Parameters
i
j
168

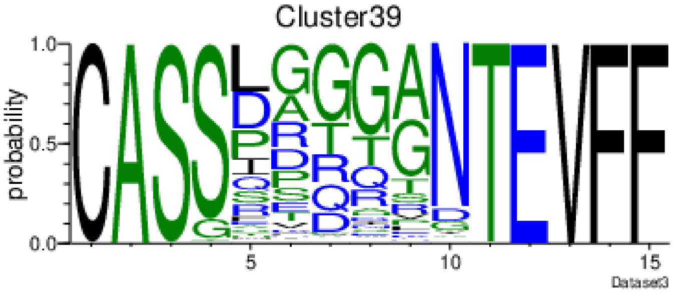
Cluster39
probability
Dataset3

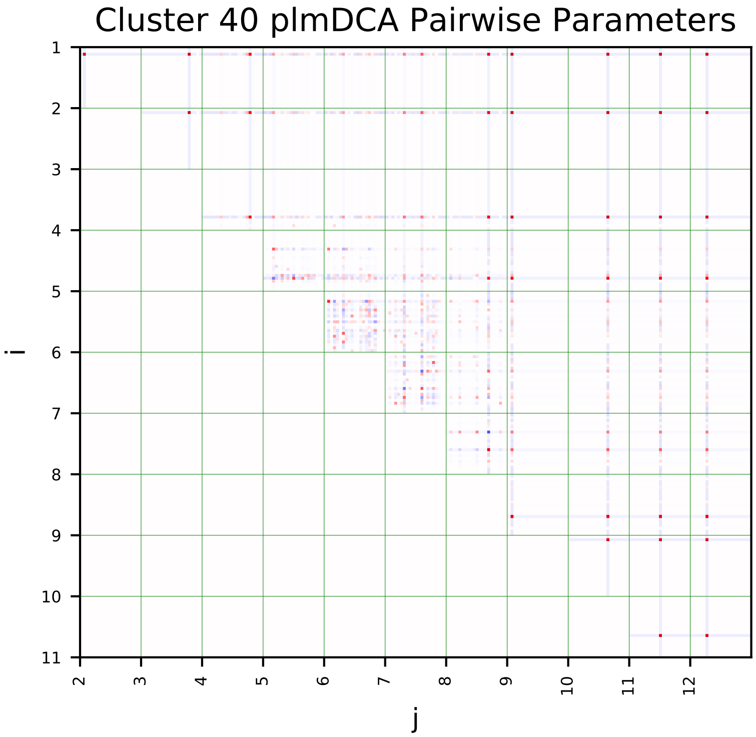
Cluster 40 plmDCA Pairwise Parameters
i
j

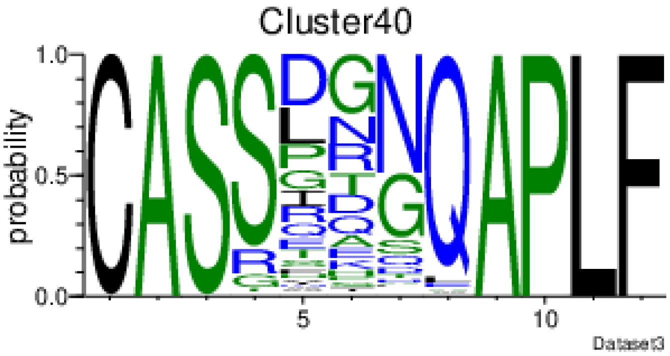
Cluster40
probability
Dataset3

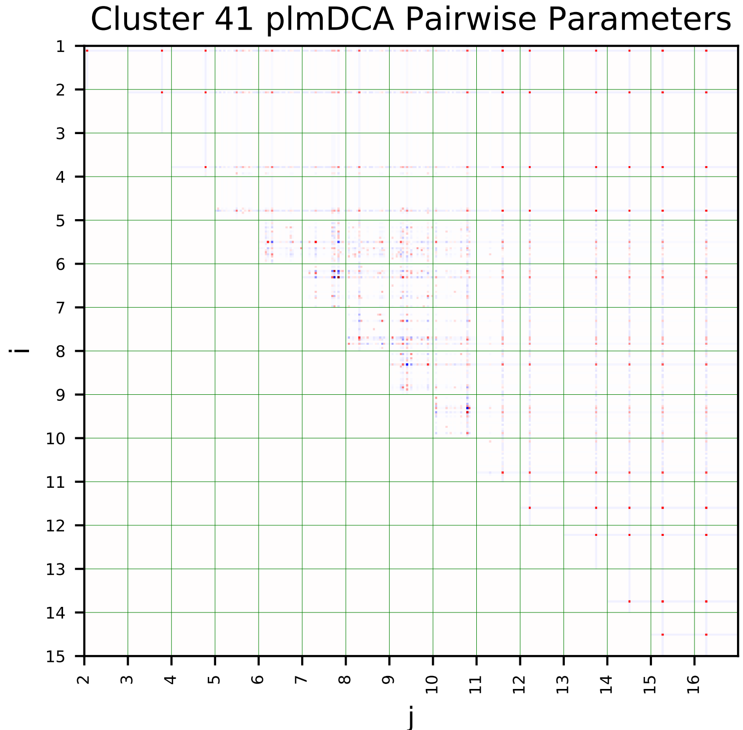
Cluster 41 plmDCA Pairwise Parameters
i
j

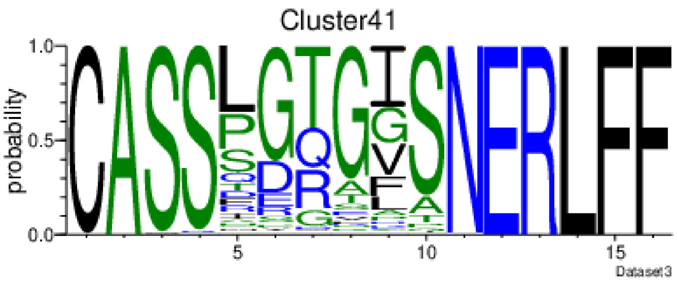
Cluster41
probability
Dataset3

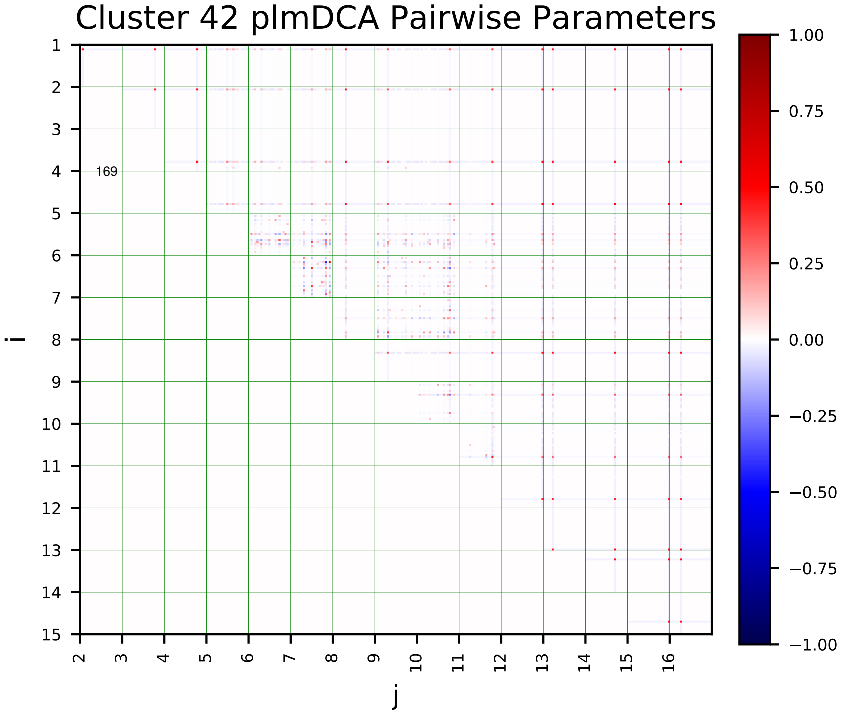
Cluster 42 plmDCA Pairwise Parameters
i
j
169

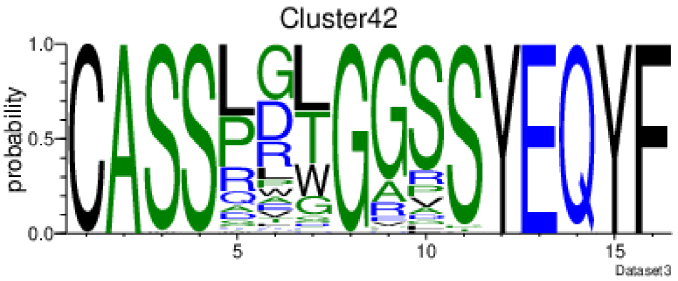
Cluster42
probability
Dataset3

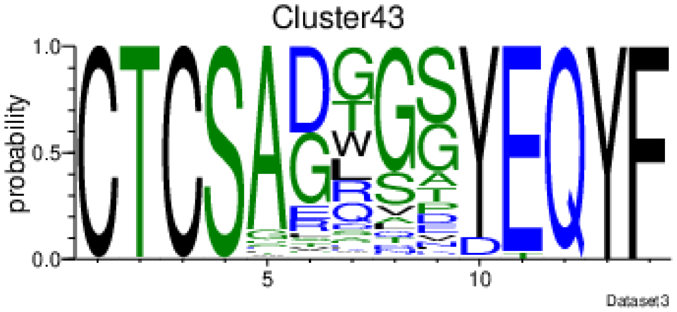
Cluster43
probability
Dataset3

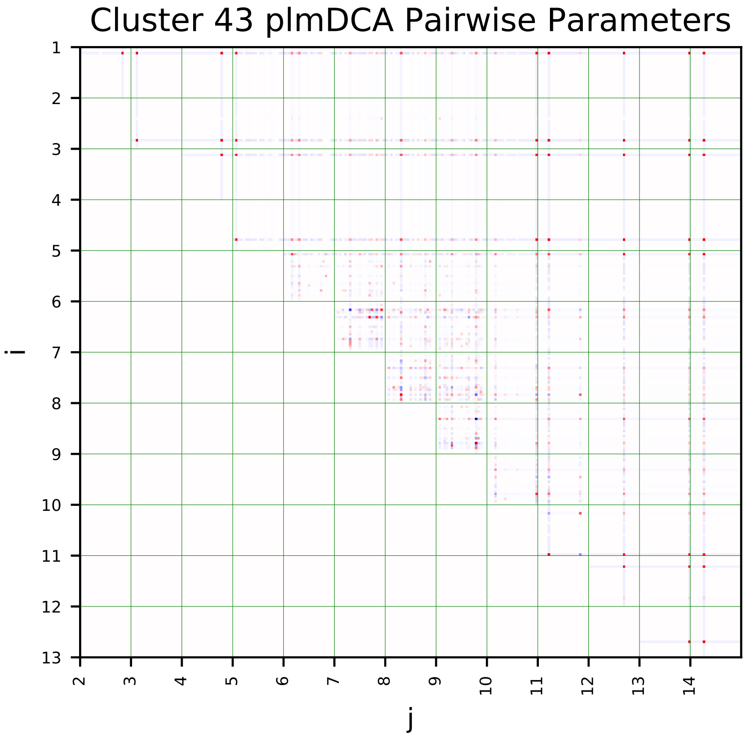
Cluster 43 plmDCA Pairwise Parameters
i
j

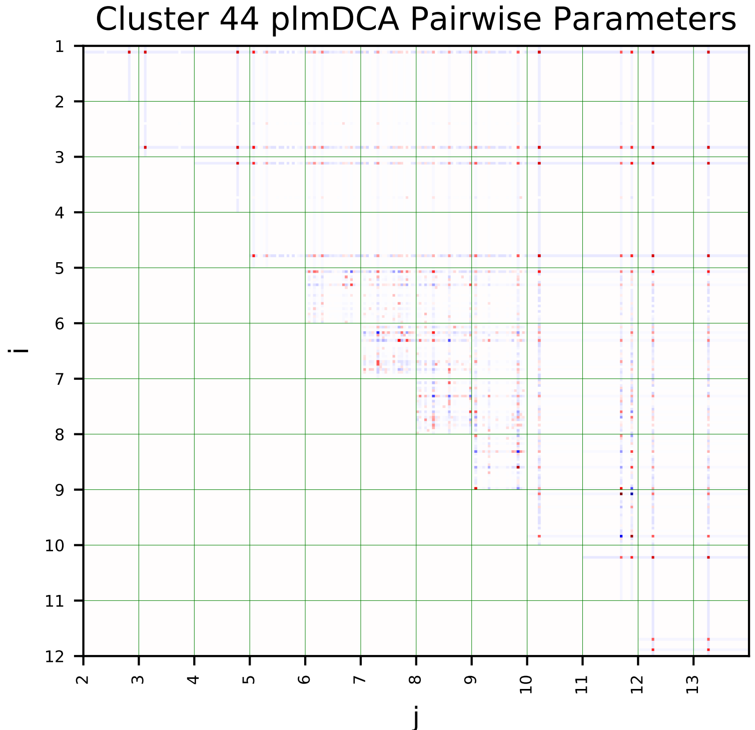
Cluster 44 plmDCA Pairwise Parameters
i
j

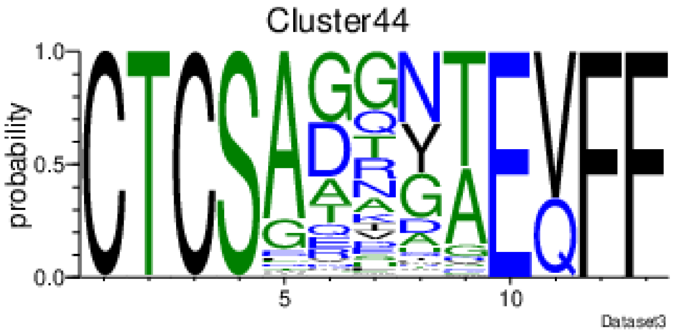
Cluster44
probability
Dataset3

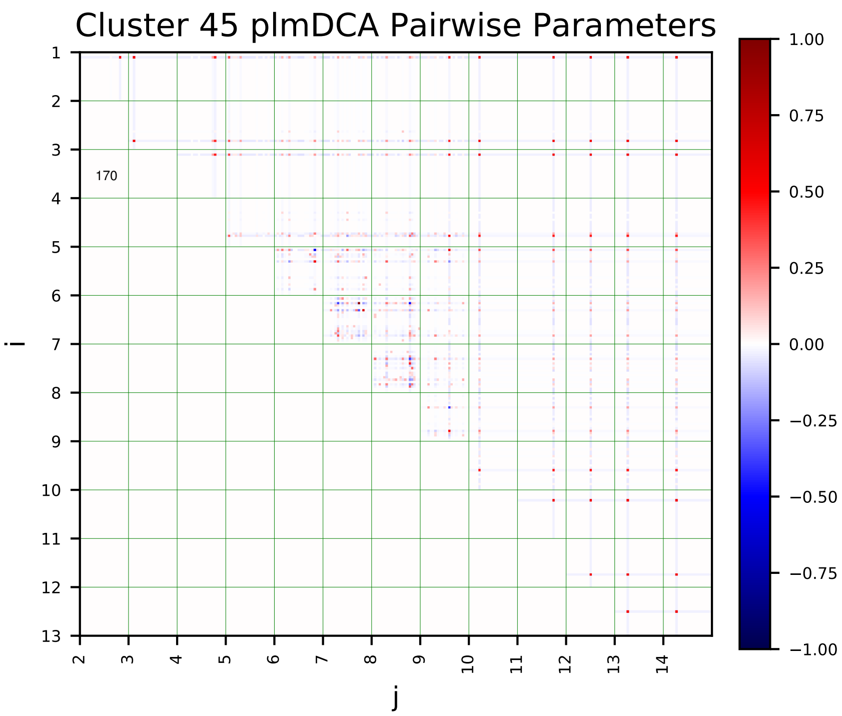
Cluster 45 plmDCA Pairwise Parameters
170
i
j

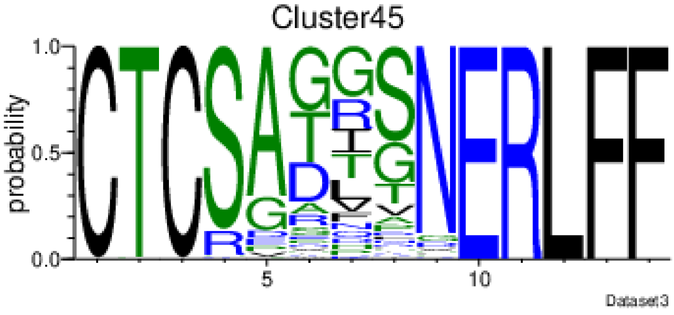
Cluster45
probability
Dataset3

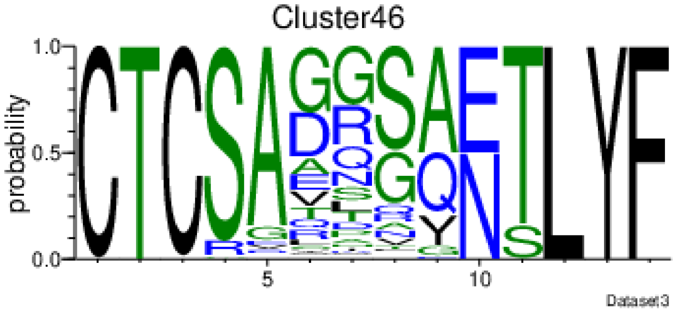
Cluster46
probability
Dataset3

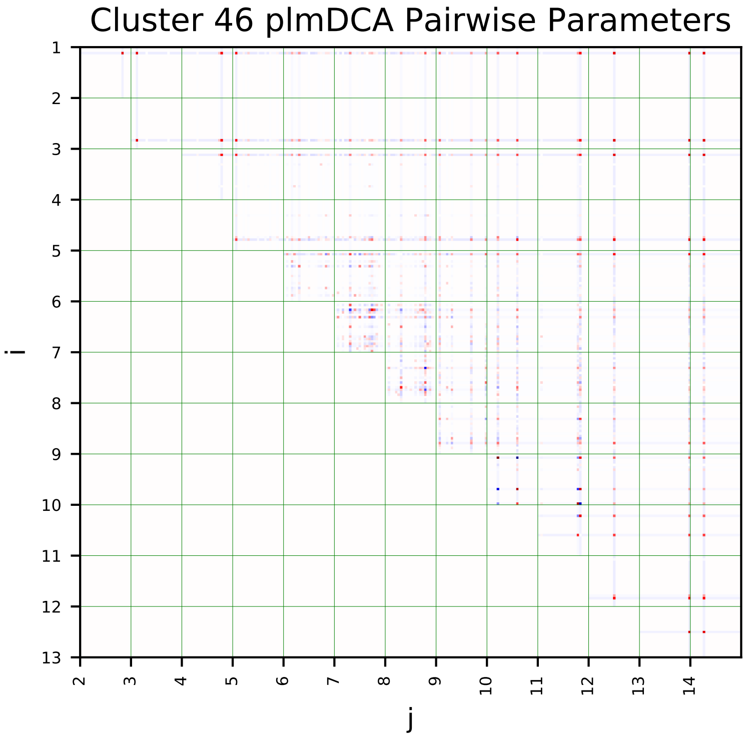
Cluster 46 plmDCA Pairwise Parameters
i
j

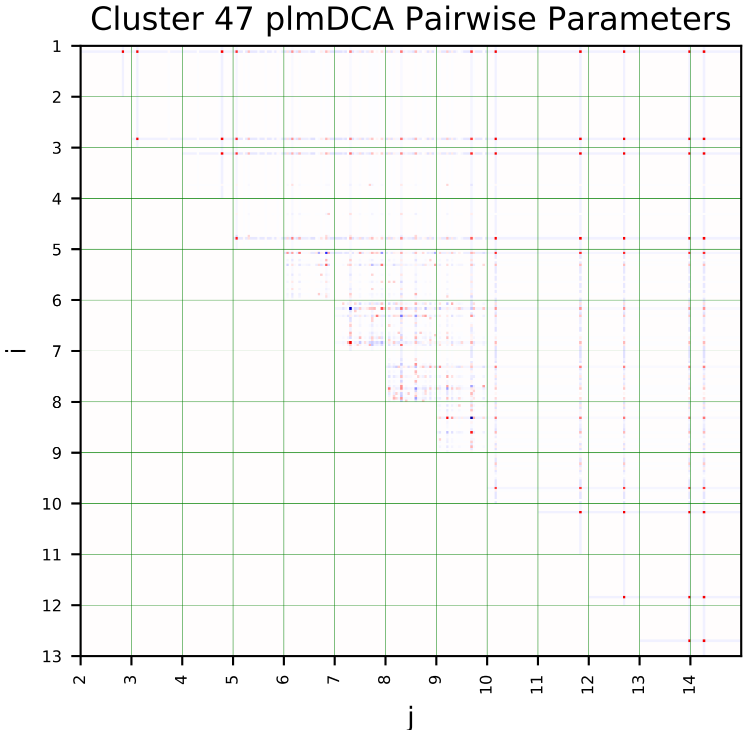
Cluster 47 plmDCA Pairwise Parameters
i
j

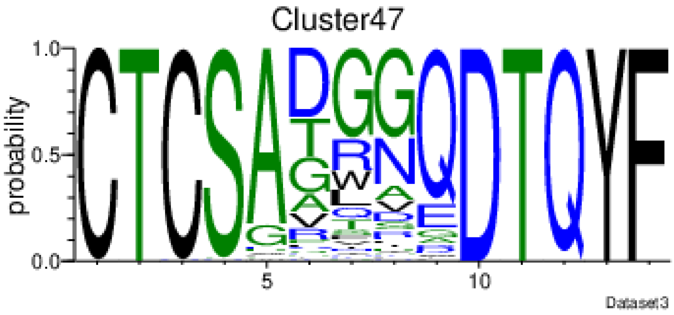
Cluster47
probability
Dataset3

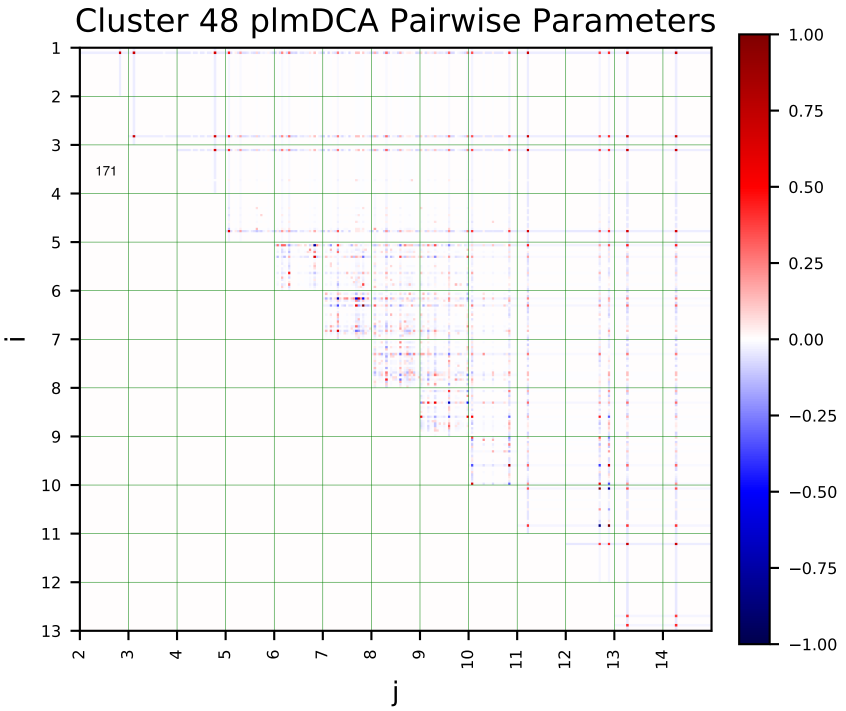
Cluster 48 plmDCA Pairwise Parameters
171
i
j
1.00
0.75
0.50
0.25
0.00
−0.25
−0.50
−0.75
−1.00

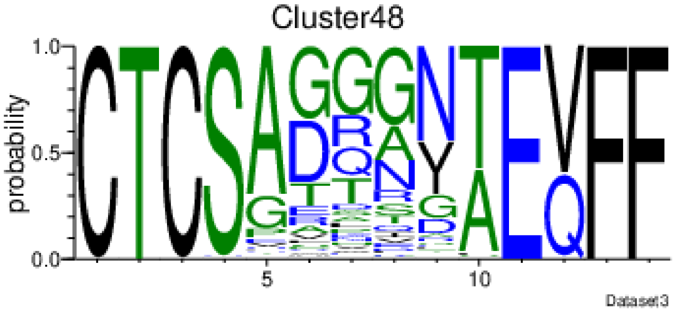
Cluster48
probability
Dataset3

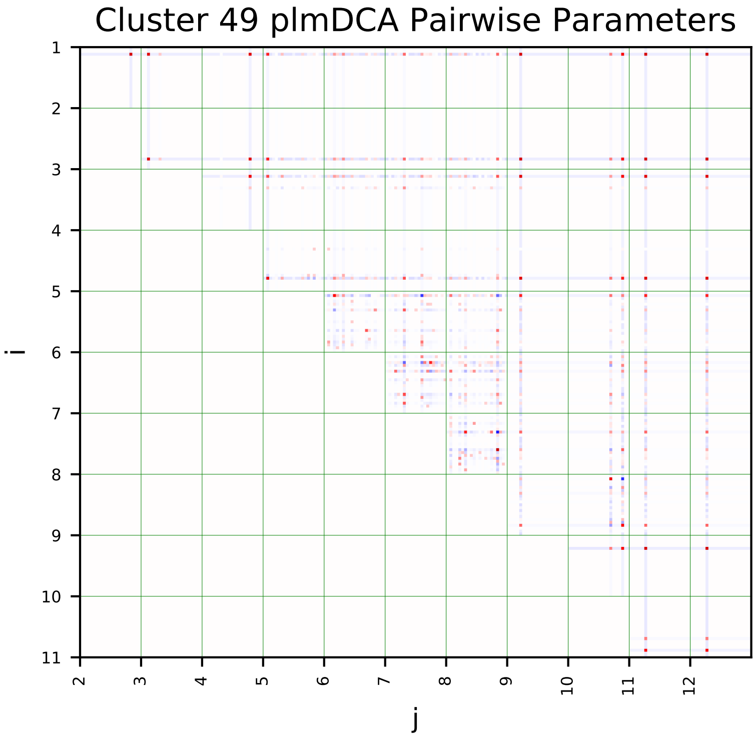
Cluster 49 plmDCA Pairwise Parameters
i
j

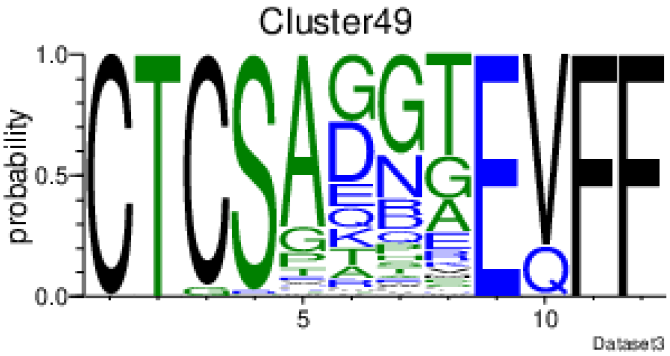
Cluster49
probability
Dataset3

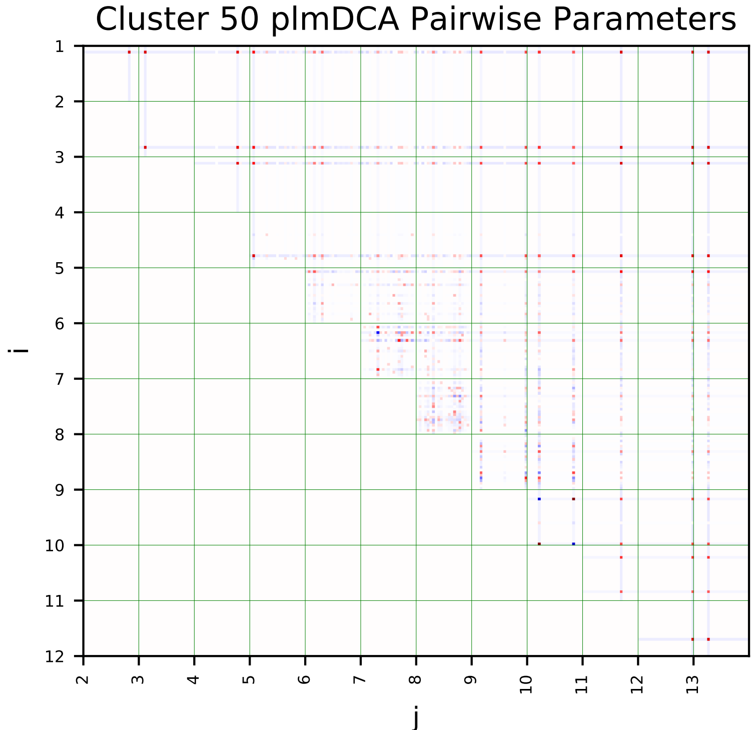
Cluster 50 plmDCA Pairwise Parameters
i
j

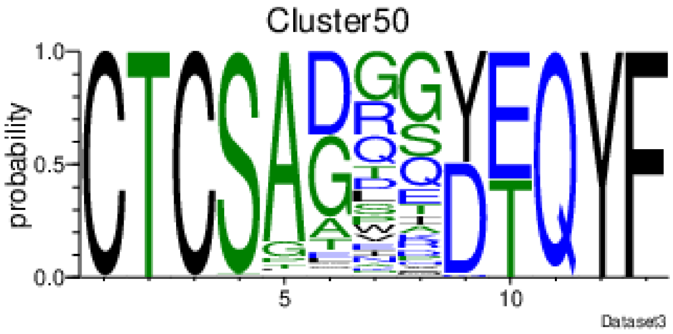
Cluster50
probability
Dataset3

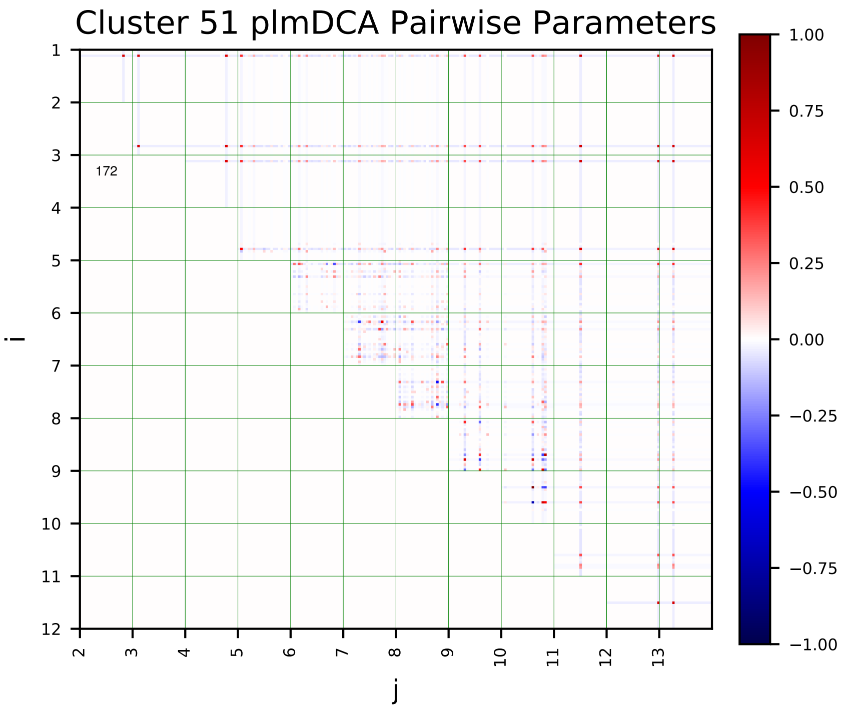
Cluster 51 plmDCA Pairwise Parameters
i
j
172

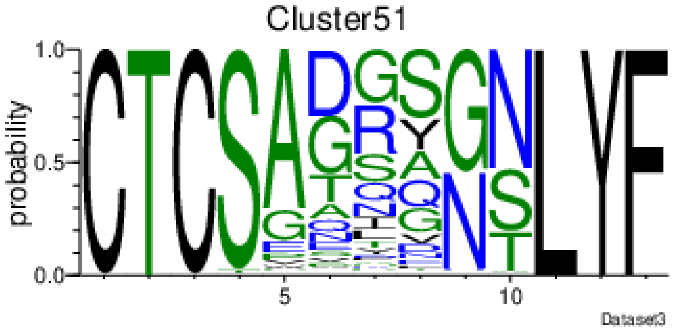
Cluster51
probability
Dataset3

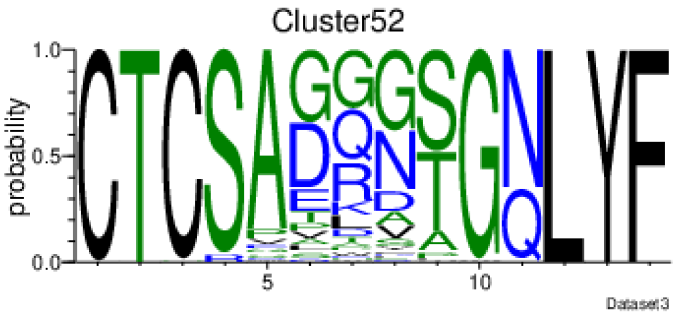
Cluster52
probability
Dataset3

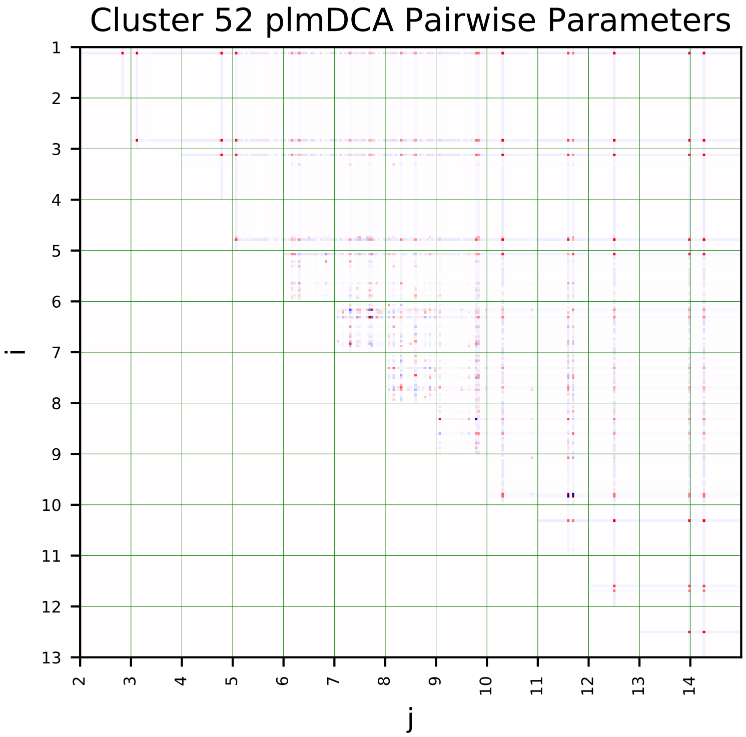
Cluster 52 plmDCA Pairwise Parameters
i
j

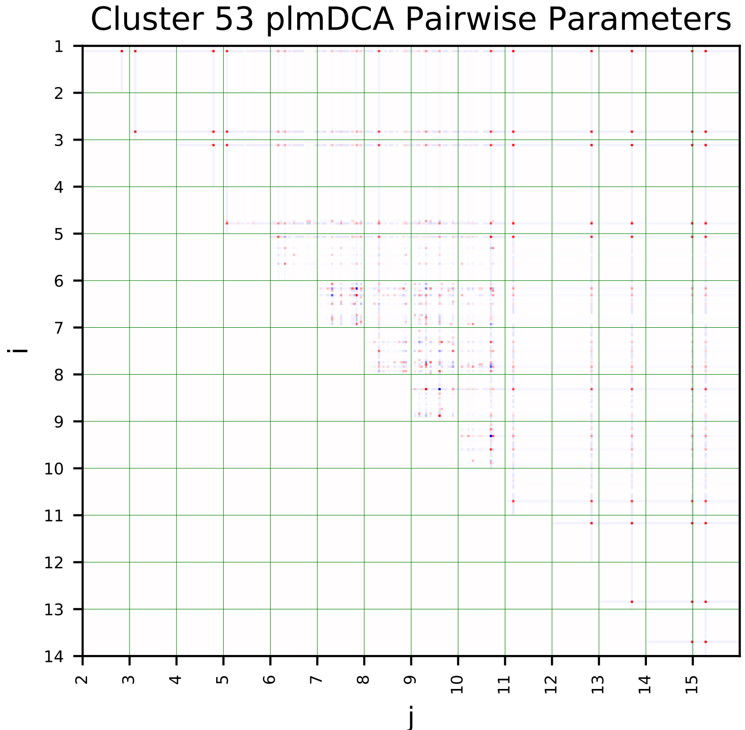
Cluster 53 plmDCA Pairwise Parameters
i
j

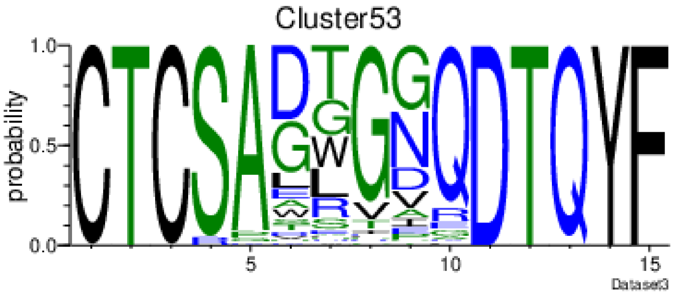
Cluster53
probability
Dataset3

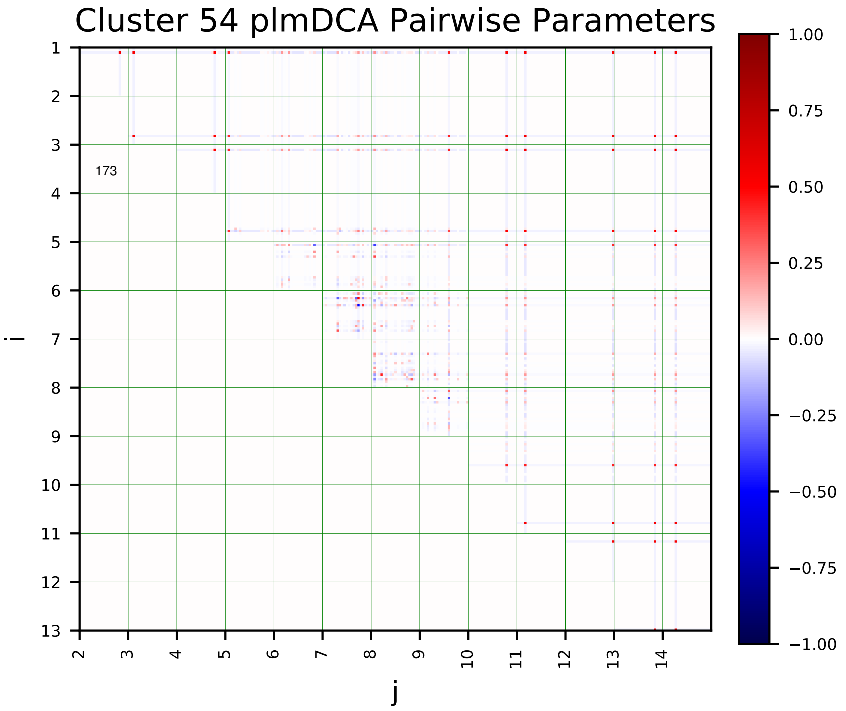
Cluster 54 plmDCA Pairwise Parameters
173
i
j
1.00
0.75
0.50
0.25
0.00
−0.25
−0.50
−0.75
−1.00

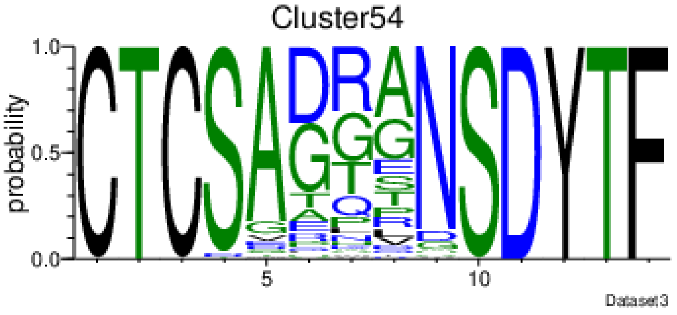
Cluster54
probability
Dataset3

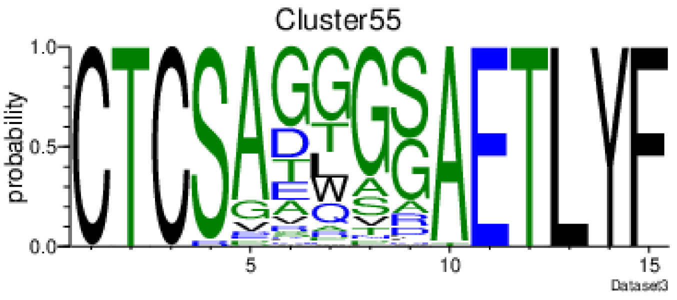
Cluster55
probability
Dataset3

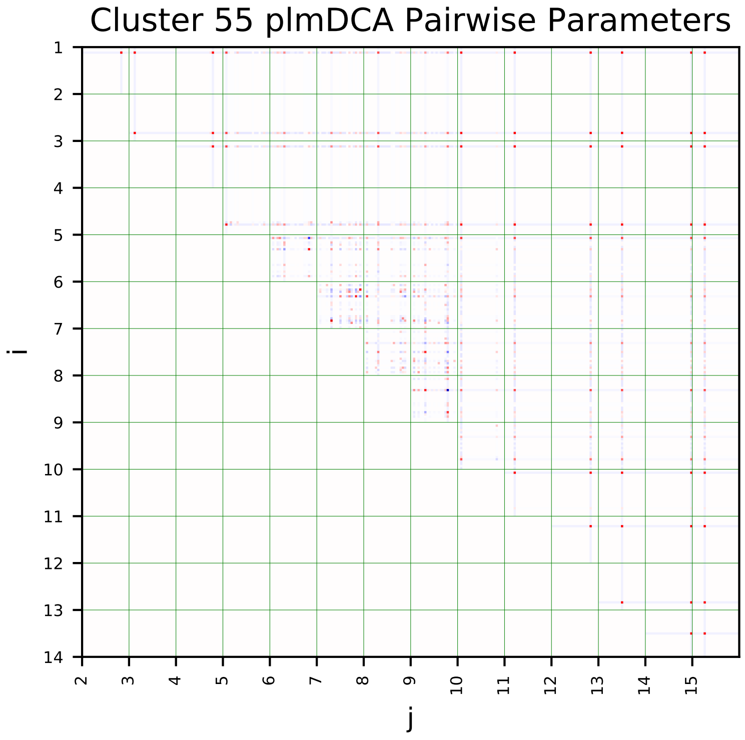
Cluster 55 plmDCA Pairwise Parameters
i
j

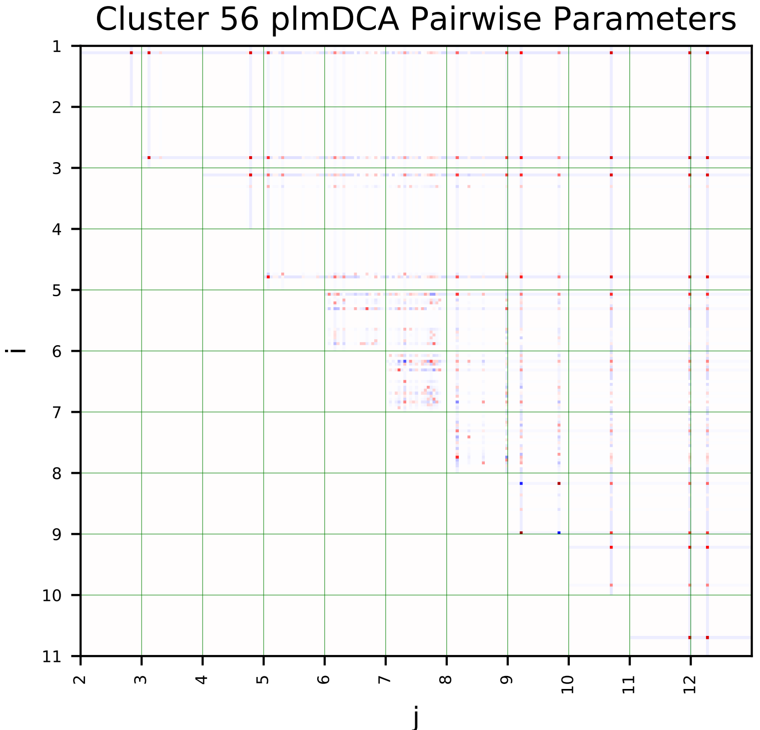
Cluster 56 plmDCA Pairwise Parameters
i
j

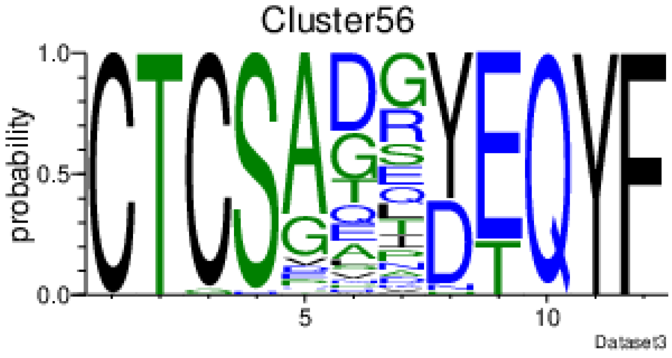
Cluster56
probability
Dataset3

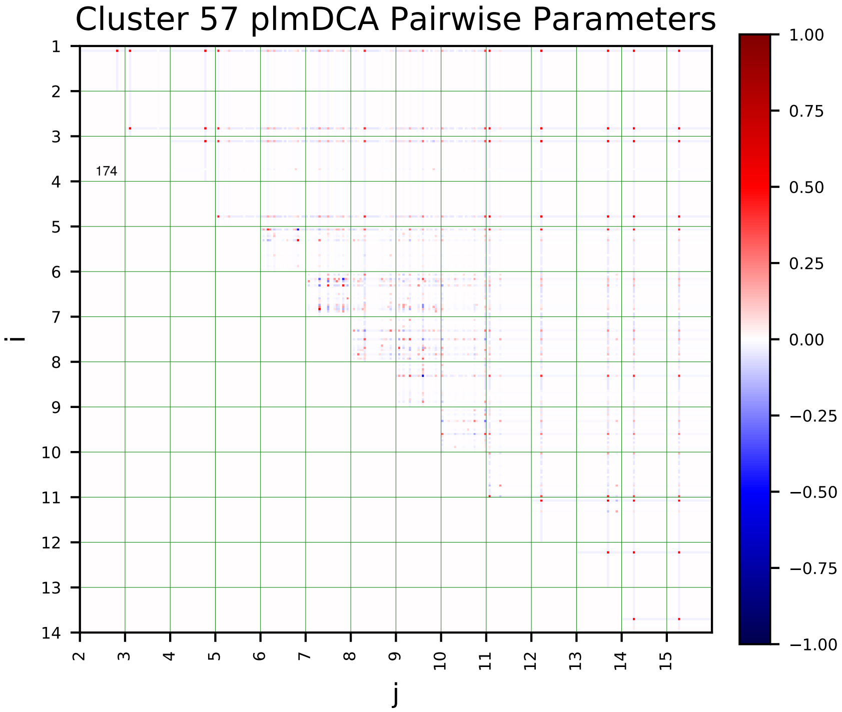
Cluster 57 plmDCA Pairwise Parameters
i
j
174
1.00
0.75
0.50
0.25
0.00
−0.25
−0.50
−0.75
−1.00

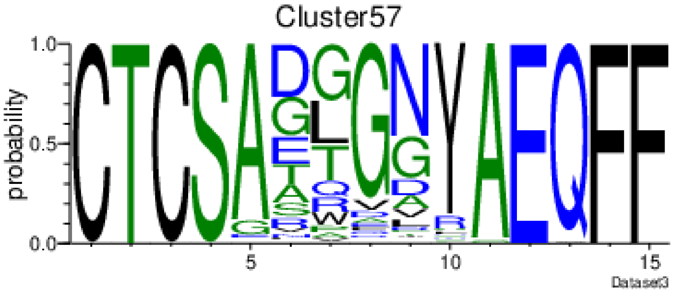
Cluster57
probability
Dataset3

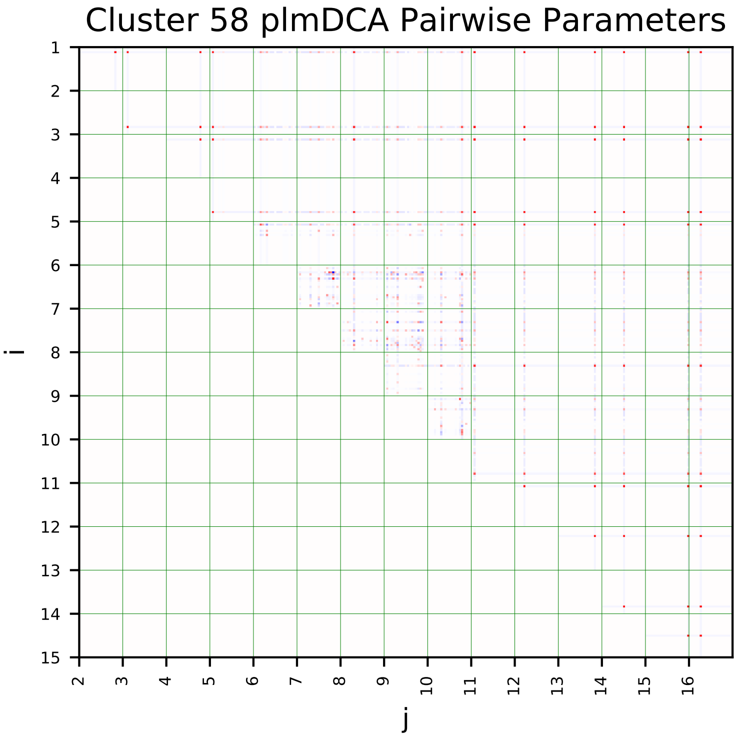
Cluster 58 plmDCA Pairwise Parameters
i
j

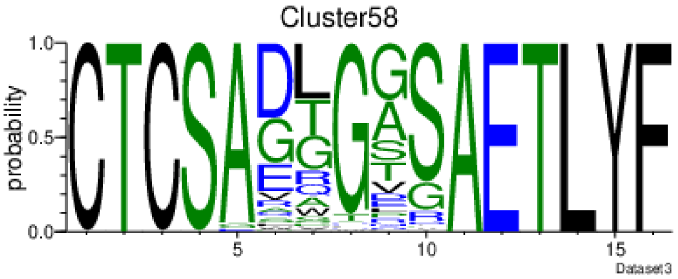
Cluster58
probability
Dataset3

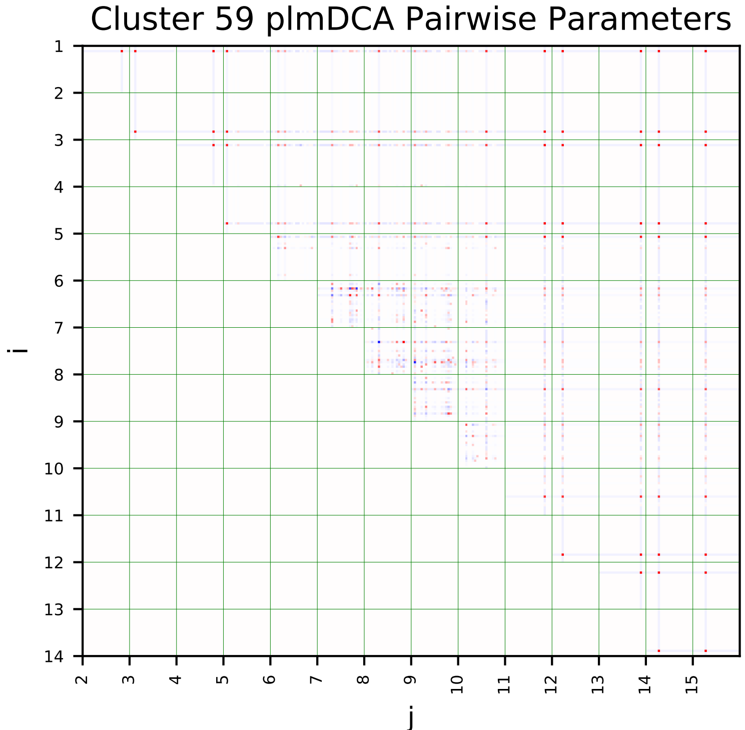
Cluster 59 plmDCA Pairwise Parameters
i
j

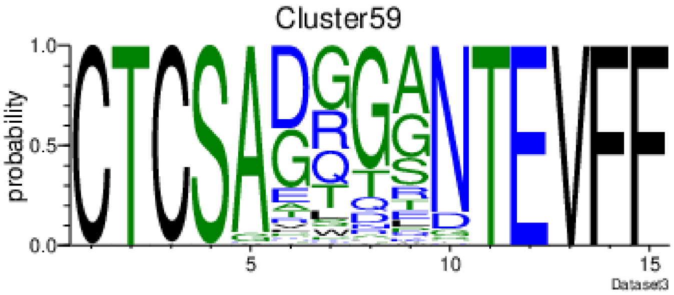
Cluster59
probability
Dataset3

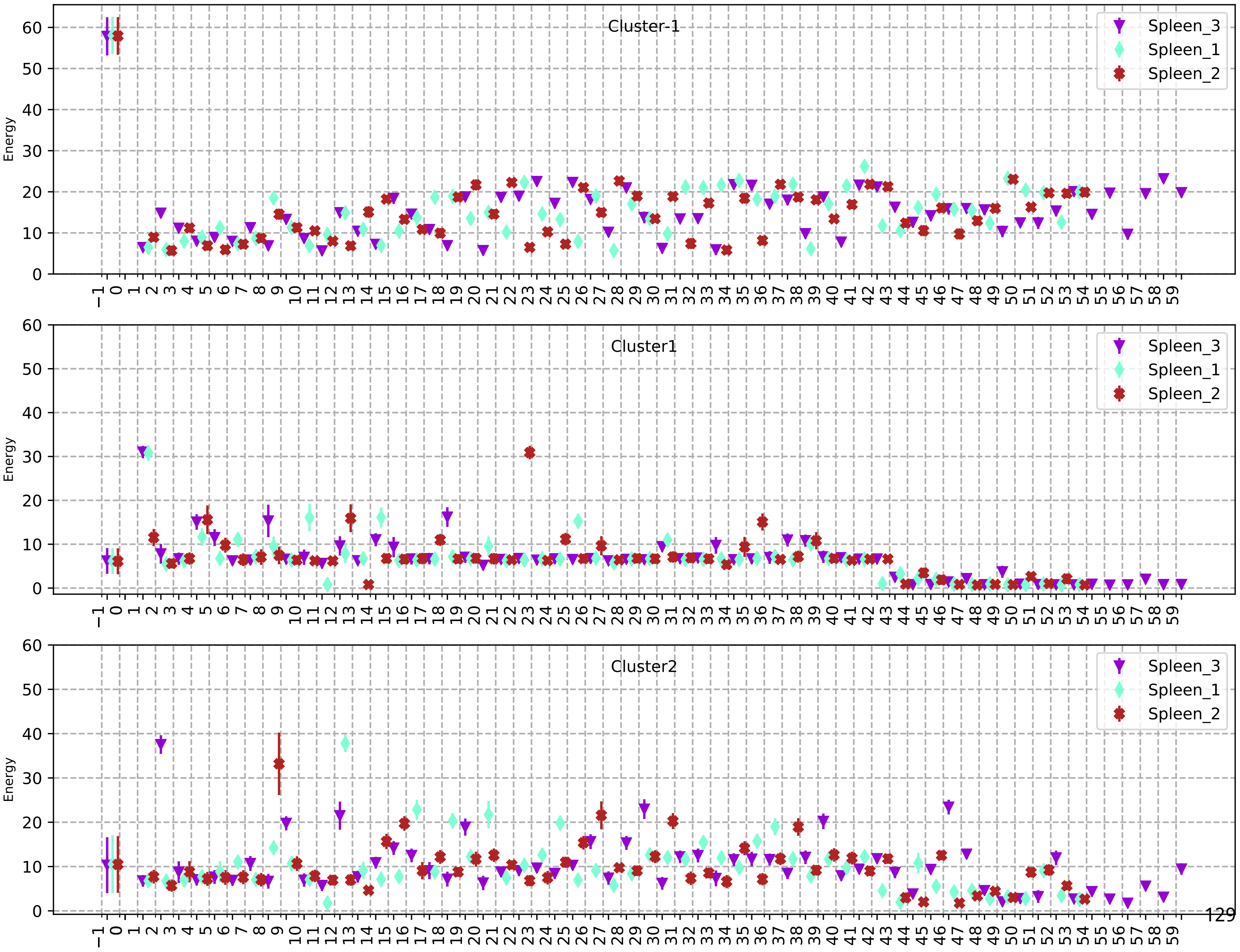

Cluster-1
Cluster1
Cluster2
Energy
Spleen_3
Spleen_1
Spleen_2

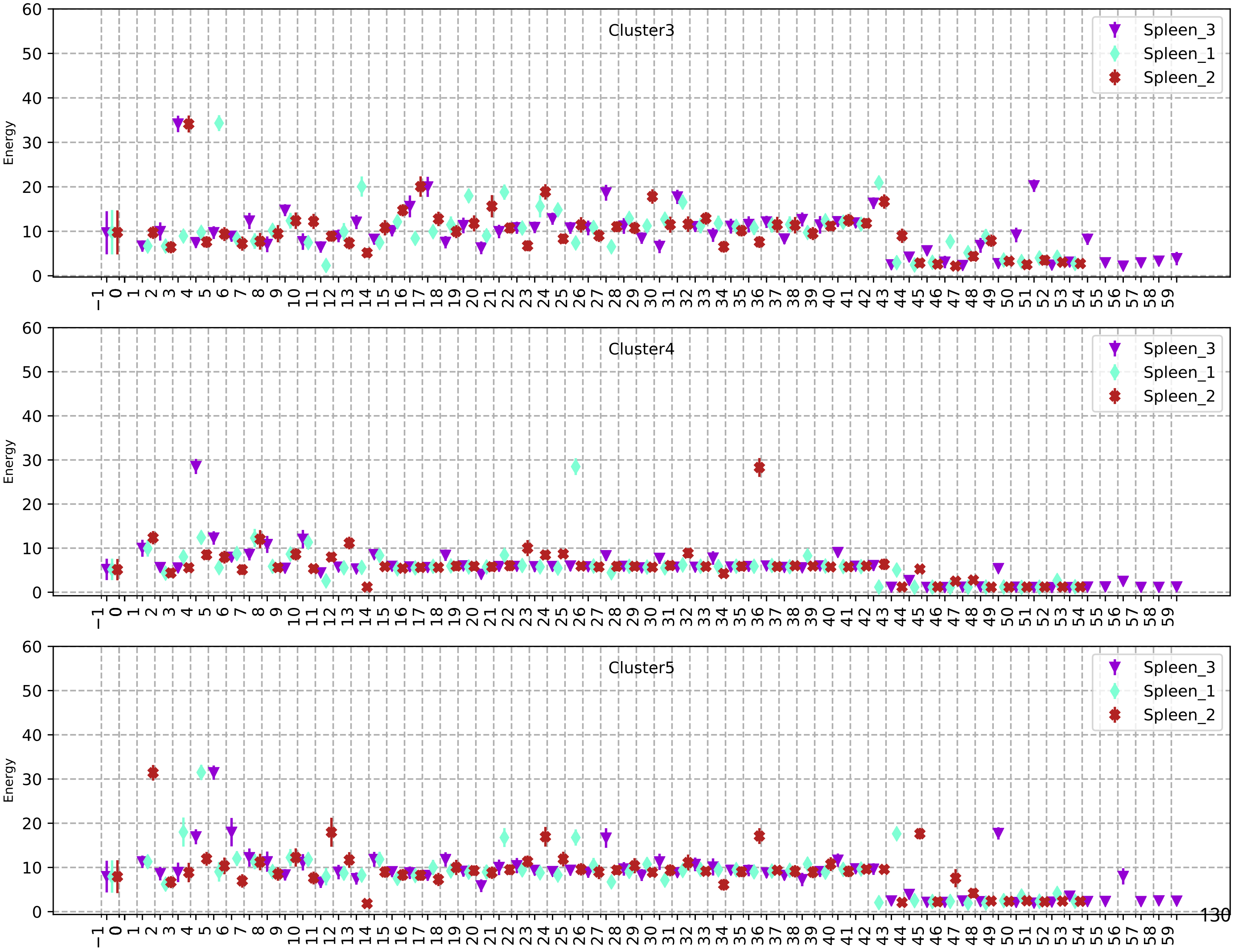
Cluster3
Cluster4
Cluster5
Spleen_3
Spleen_1
Spleen_2
Energy

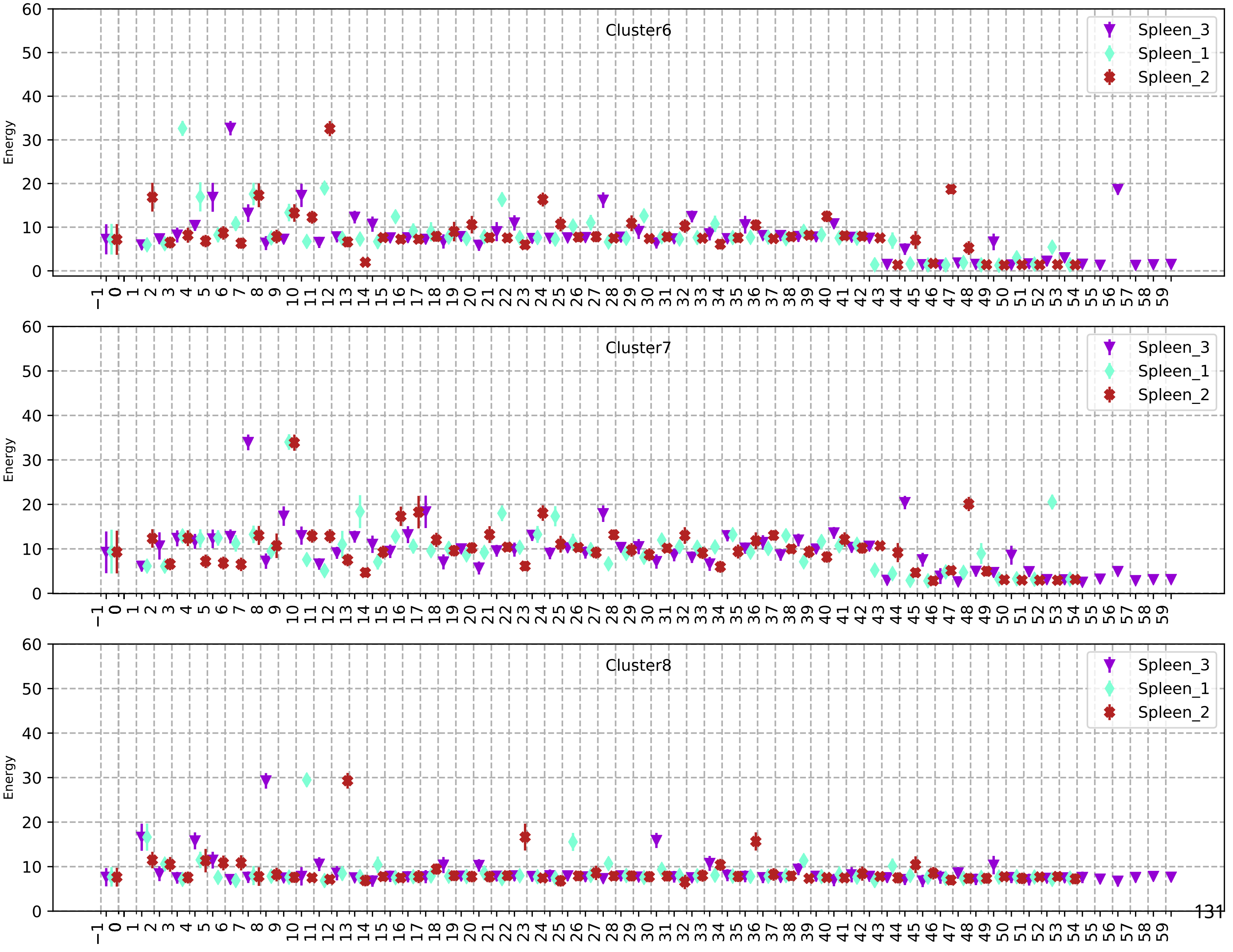
Cluster6
Cluster7
Cluster8
Spleen_3
Spleen_1
Spleen_2
Energy
0
10
20
30
40
50
60
−1 0 1 2 3 4 5 6 7 8 9 10 11 12 13 14 15 16 17 18 19 20 21 22 23 24 25 26 27 28 29 30 31 32 33 34 35 36 37 38 39 40 41 42 43 44 45 46 47 48 49 50 51 52 53 54 55 56 57 58 59

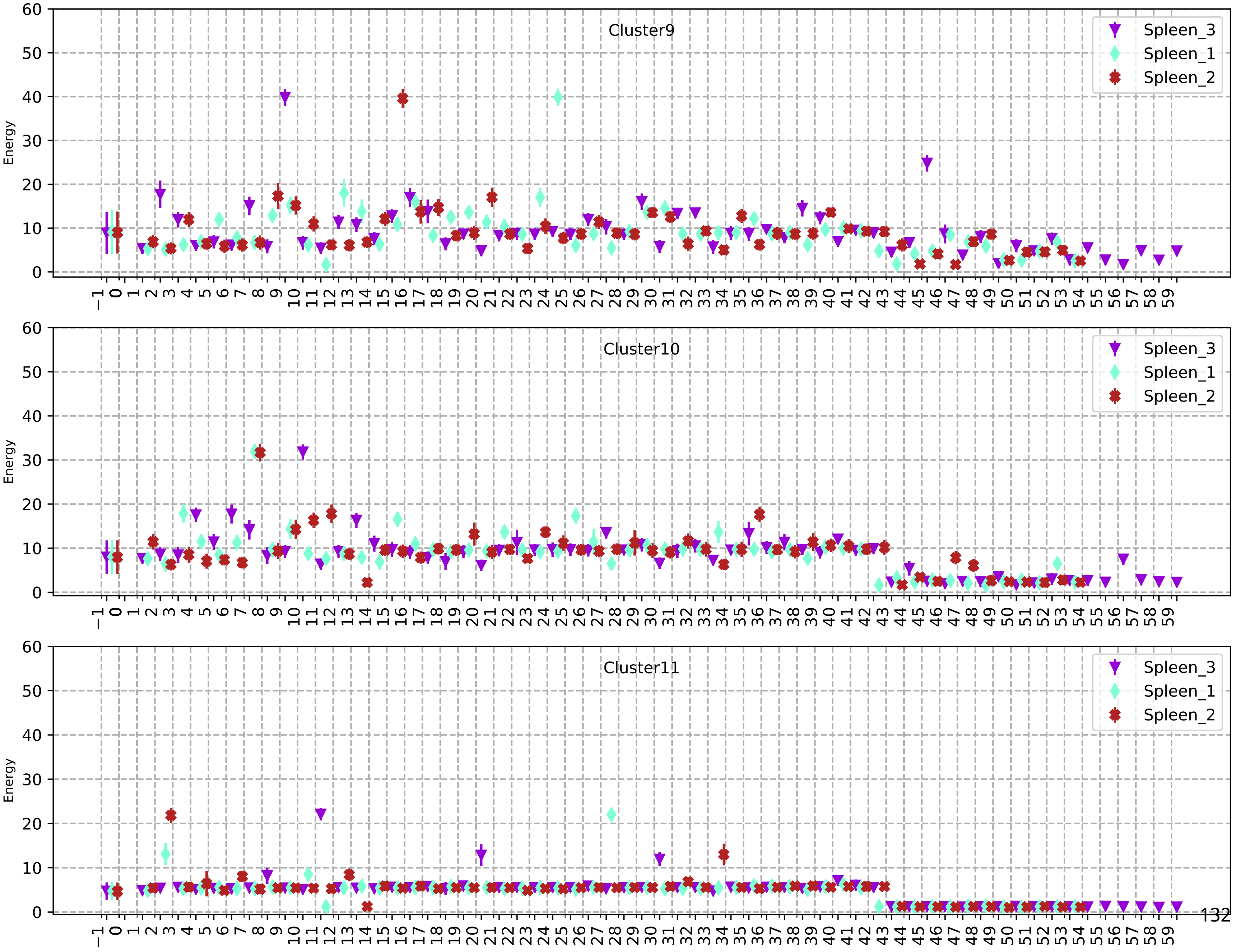
Cluster9
Cluster10
Cluster11
Spleen_3
Spleen_1
Spleen_2
Energy

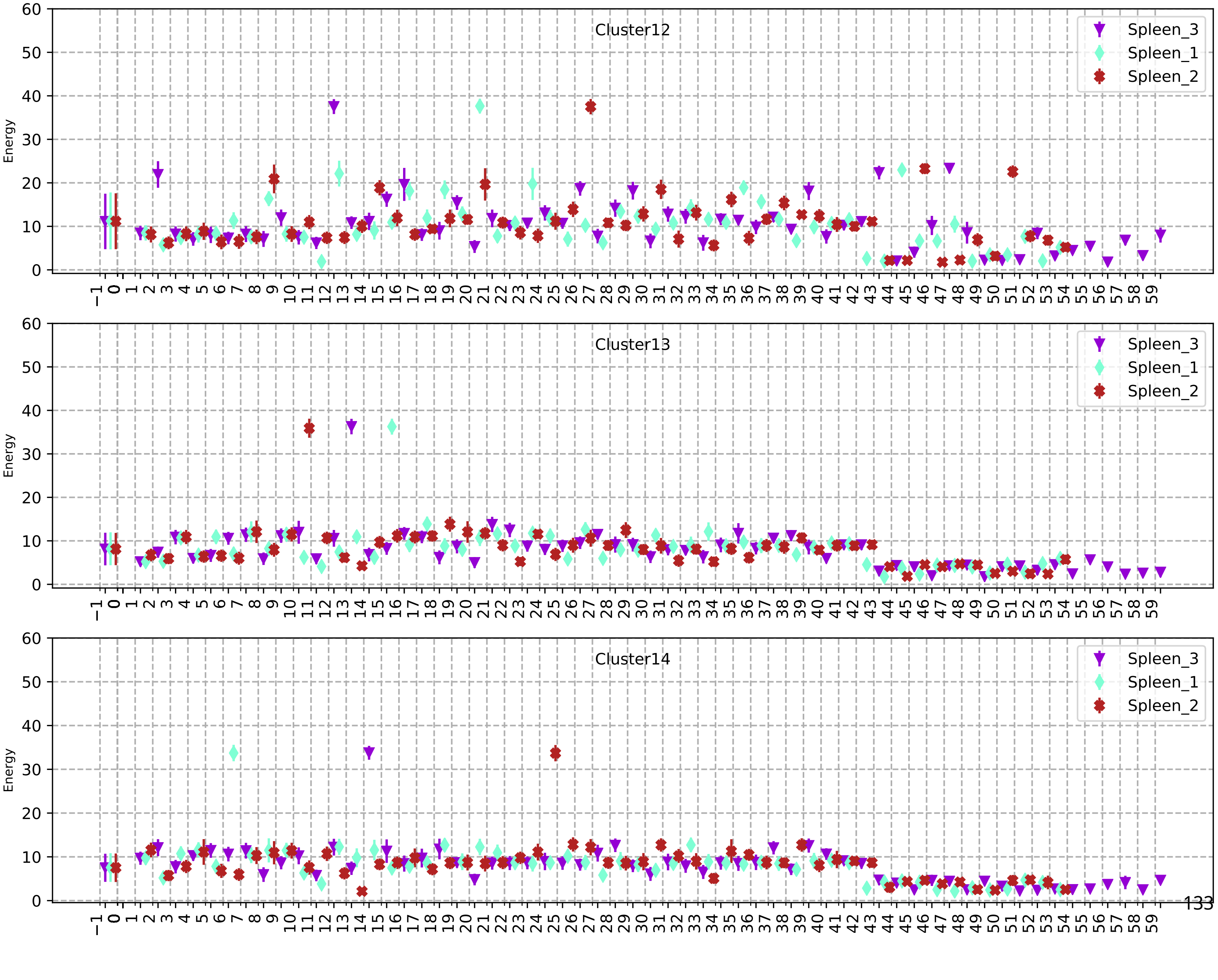
Cluster12
Cluster13
Cluster14
Energy
Spleen_3
Spleen_1
Spleen_2

180

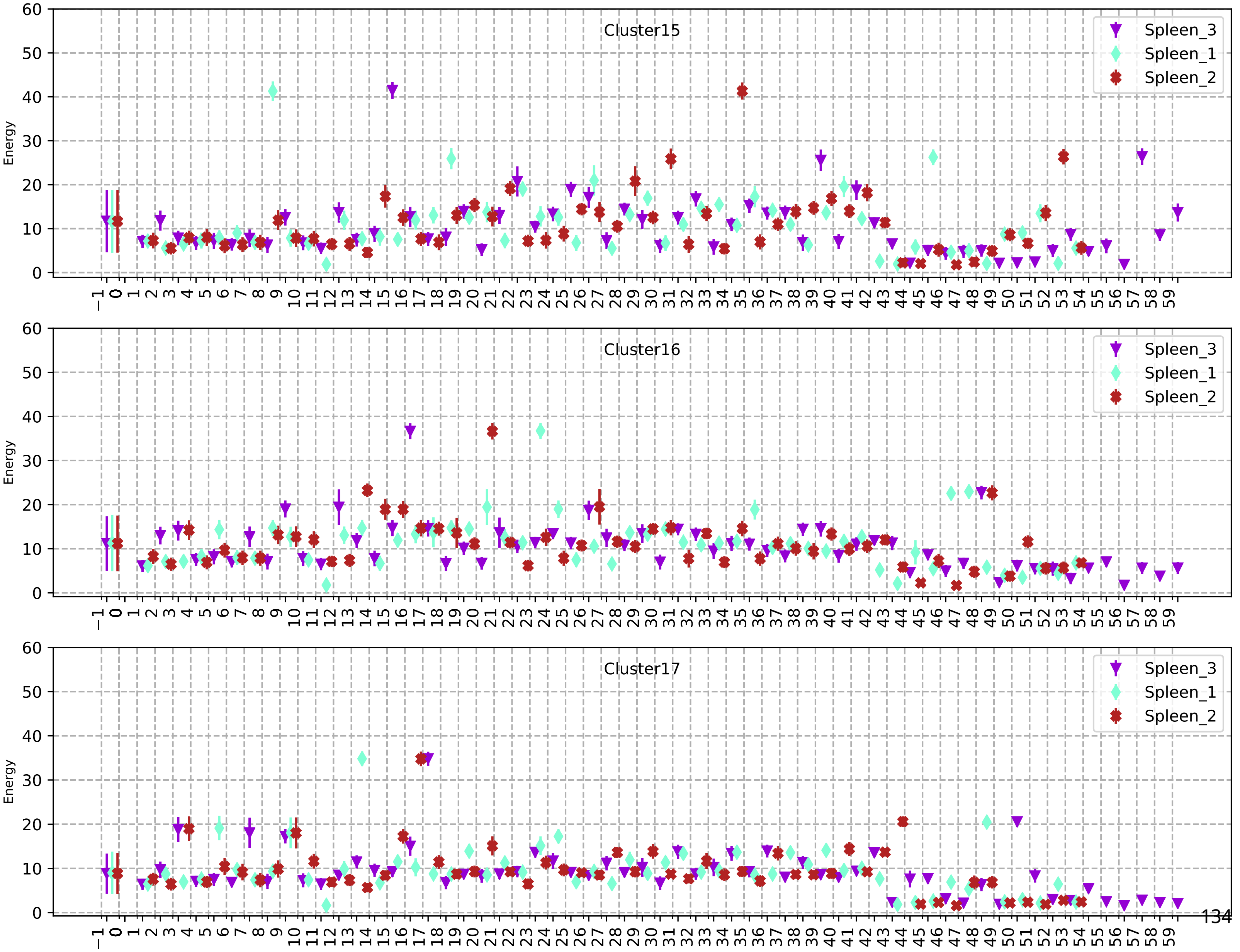
Cluster15
Cluster16
Cluster17
Energy
Spleen_3
Spleen_1
Spleen_2

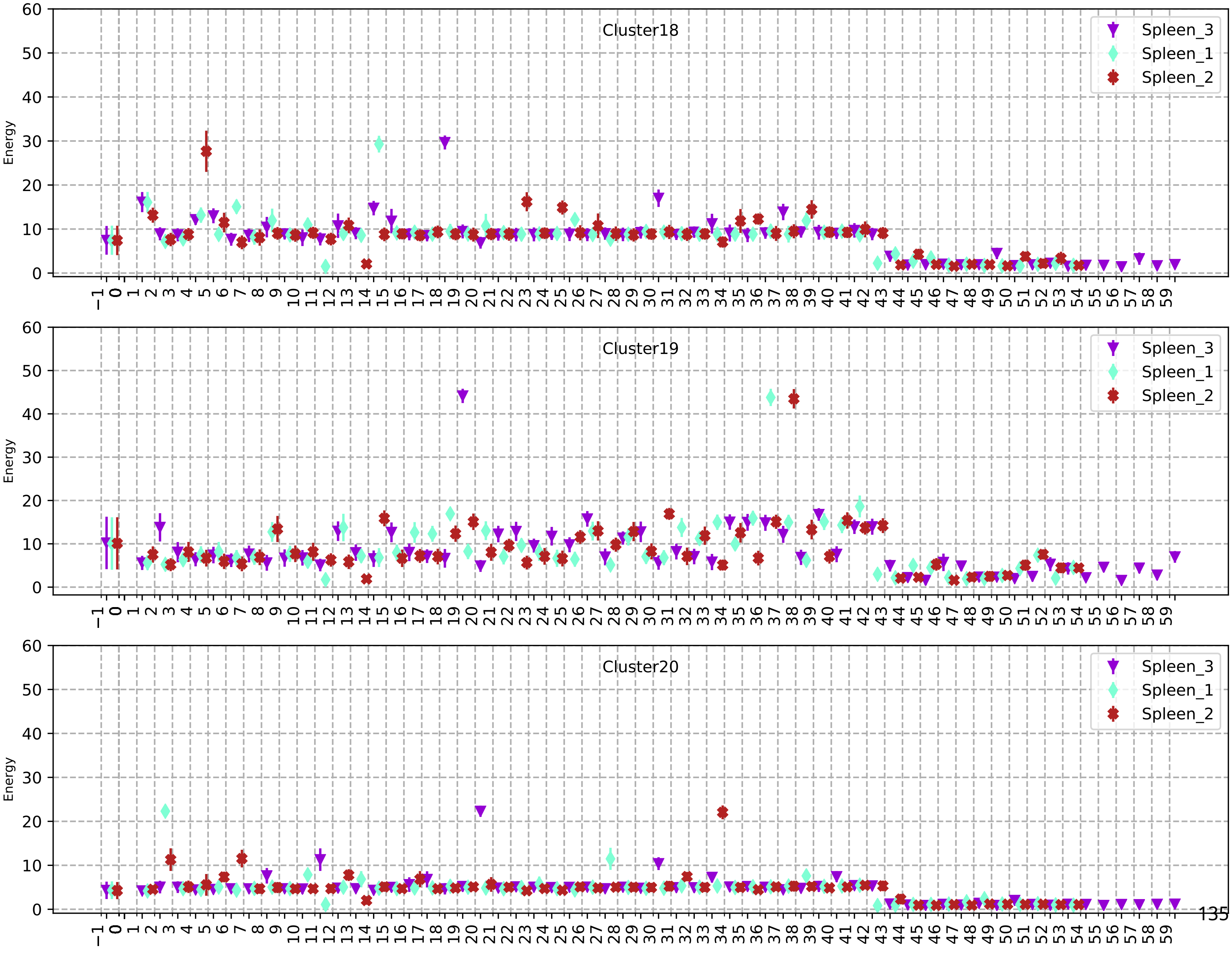
Cluster18
Spleen_3
Spleen_1
Spleen_2
Energy
Cluster19
Spleen_3
Spleen_1
Spleen_2
Energy
Cluster20
Spleen_3
Spleen_1
Spleen_2
Energy

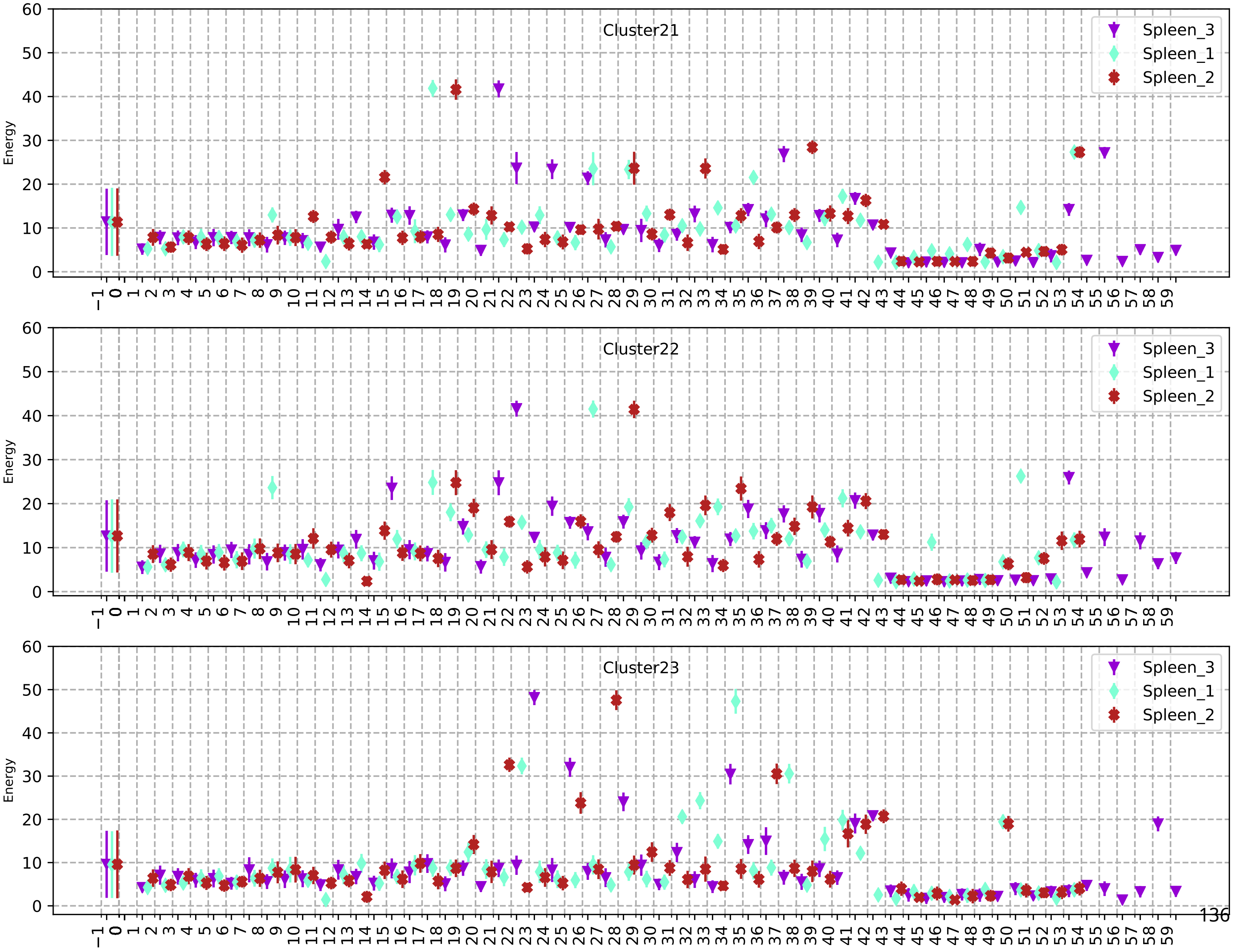
Cluster21
Cluster22
Cluster23
Energy
Spleen_3
Spleen_1
Spleen_2

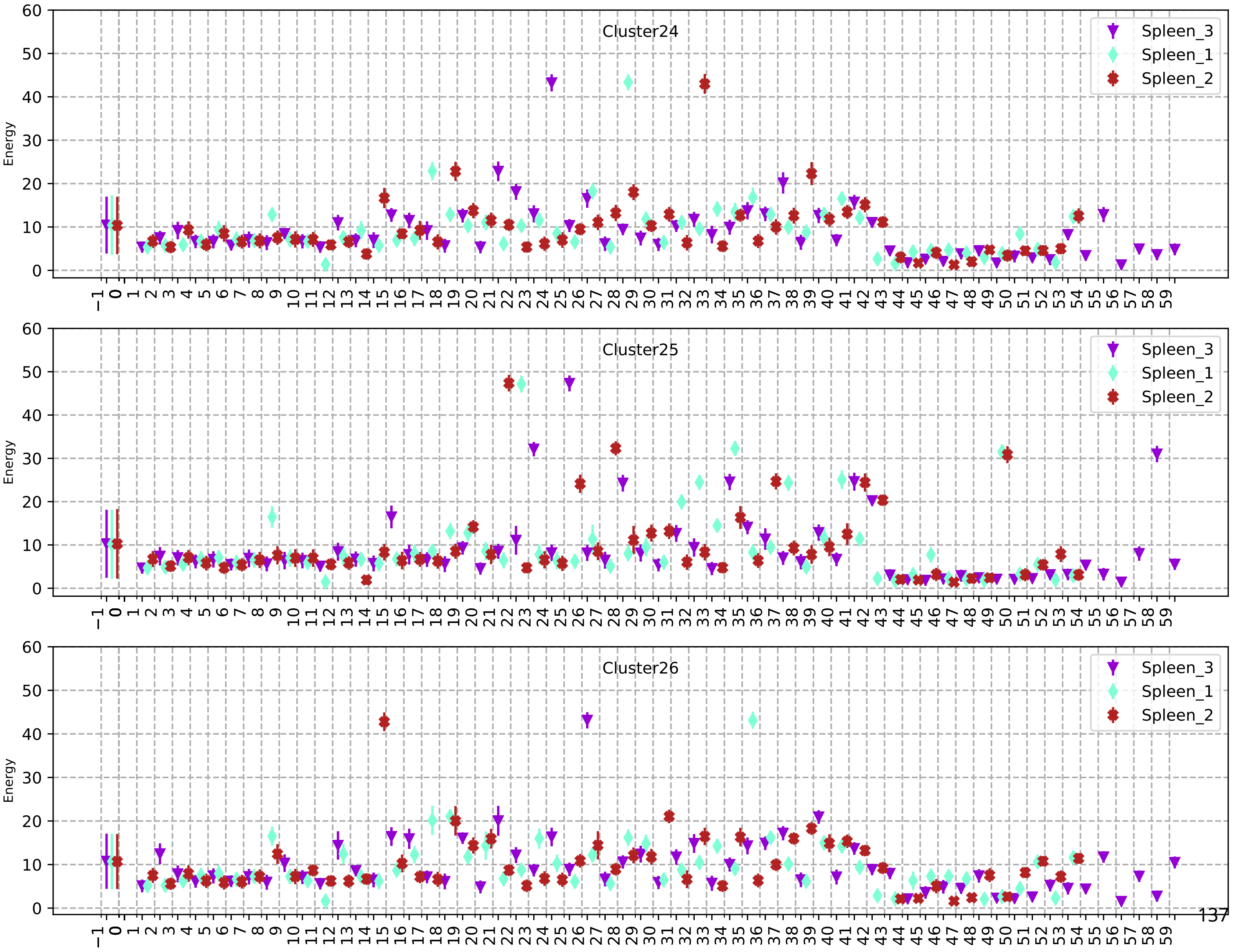
Cluster24
Cluster25
Cluster26
Spleen_3
Spleen_1
Spleen_2
Energy

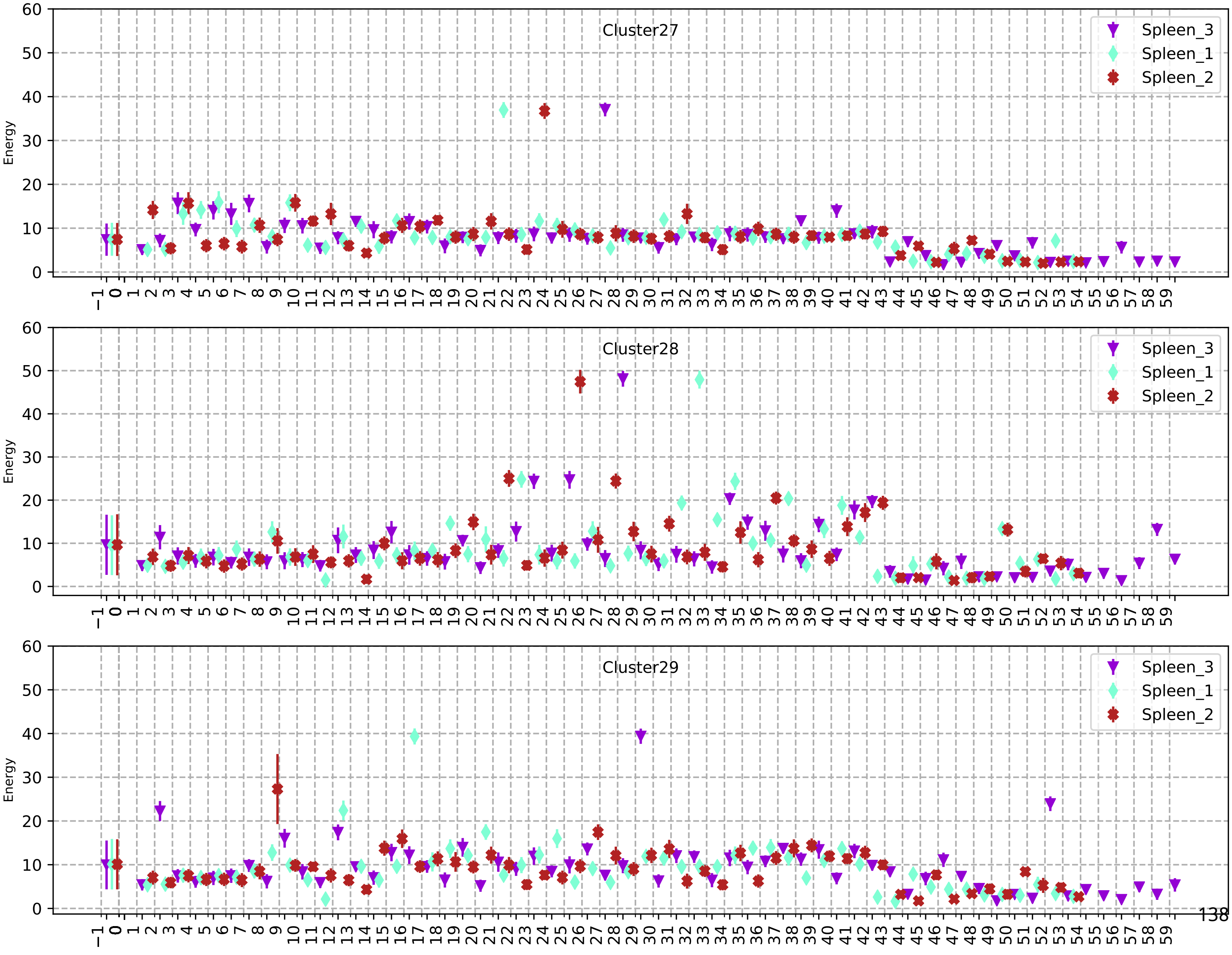
Cluster27
Spleen_3
Spleen_1
Spleen_2
Energy
Cluster28
Spleen_3
Spleen_1
Spleen_2
Energy
Cluster29
Spleen_3
Spleen_1
Spleen_2
Energy

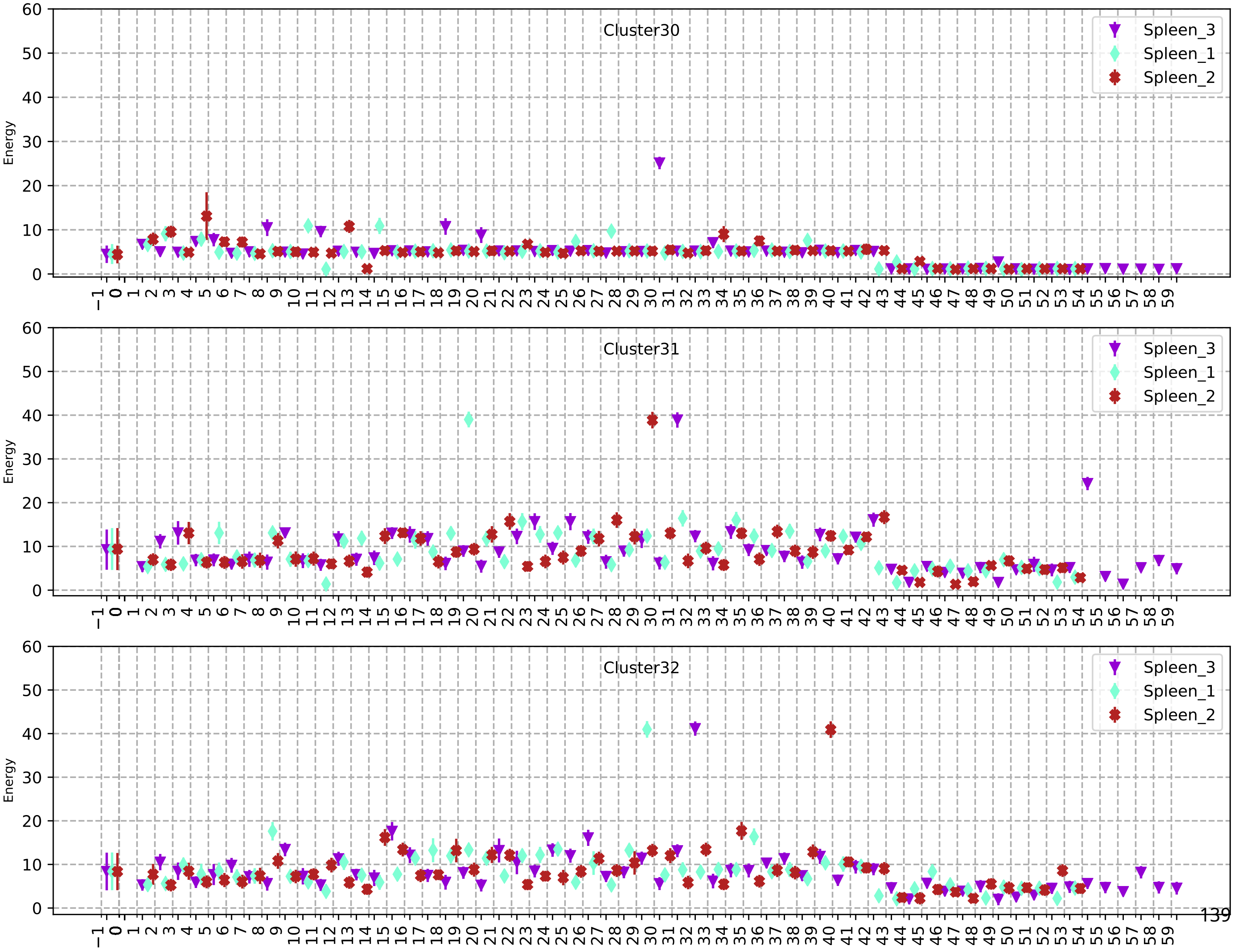
Cluster30
Spleen_3
Spleen_1
Spleen_2
Energy
Cluster31
Spleen_3
Spleen_1
Spleen_2
Energy
Cluster32
Spleen_3
Spleen_1
Spleen_2
Energy

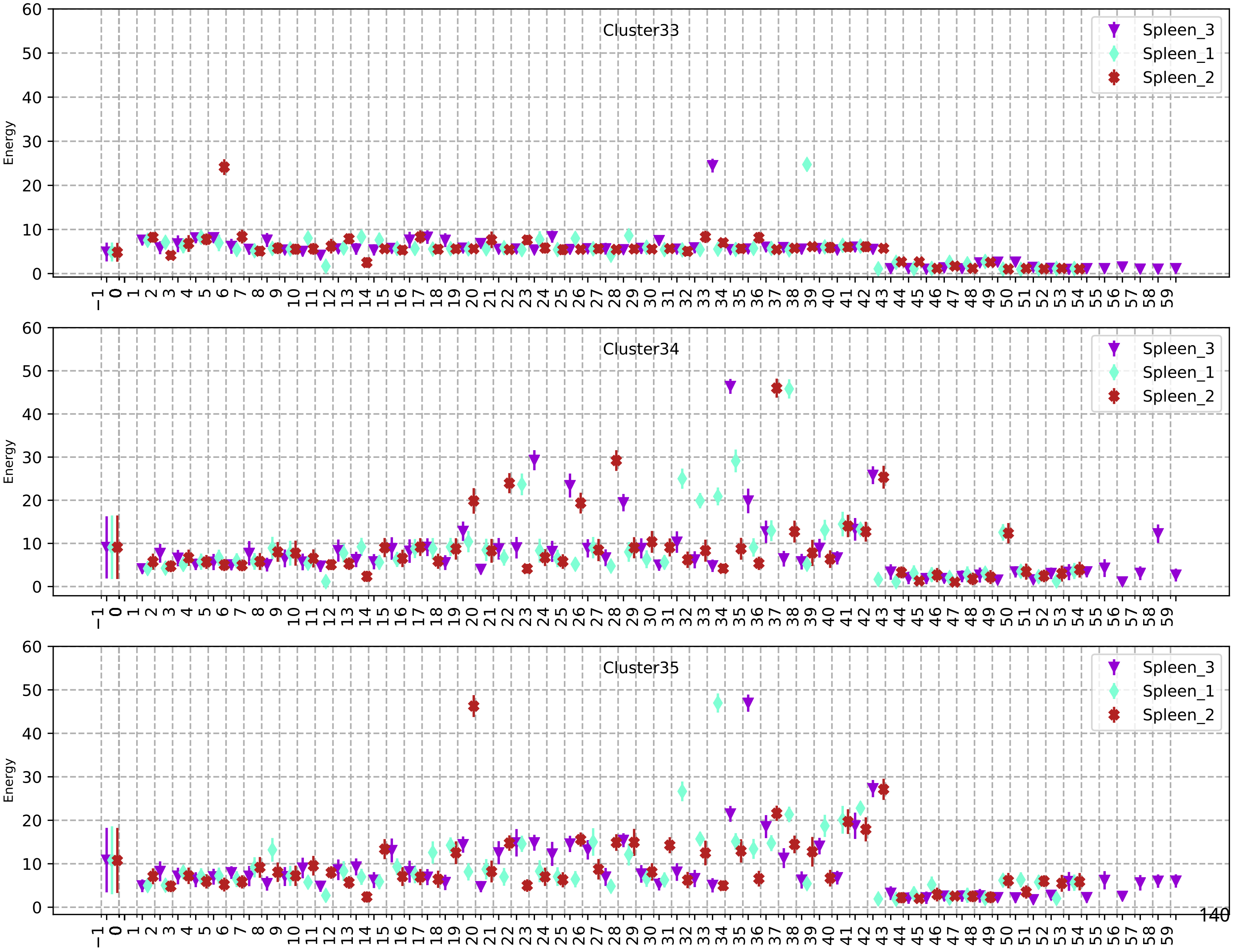
Cluster33
Cluster34
Cluster35
Spleen_3
Spleen_1
Spleen_2
Energy
−1 0 1 2 3 4 5 6 7 8 9 10 11 12 13 14 15 16 17 18 19 20 21 22 23 24 25 26 27 28 29 30 31 32 33 34 35 36 37 38 39 40 41 42 43 44 45 46 47 48 49 50 51 52 53 54 55 56 57 58 59
0 10 20 30 40 50 60

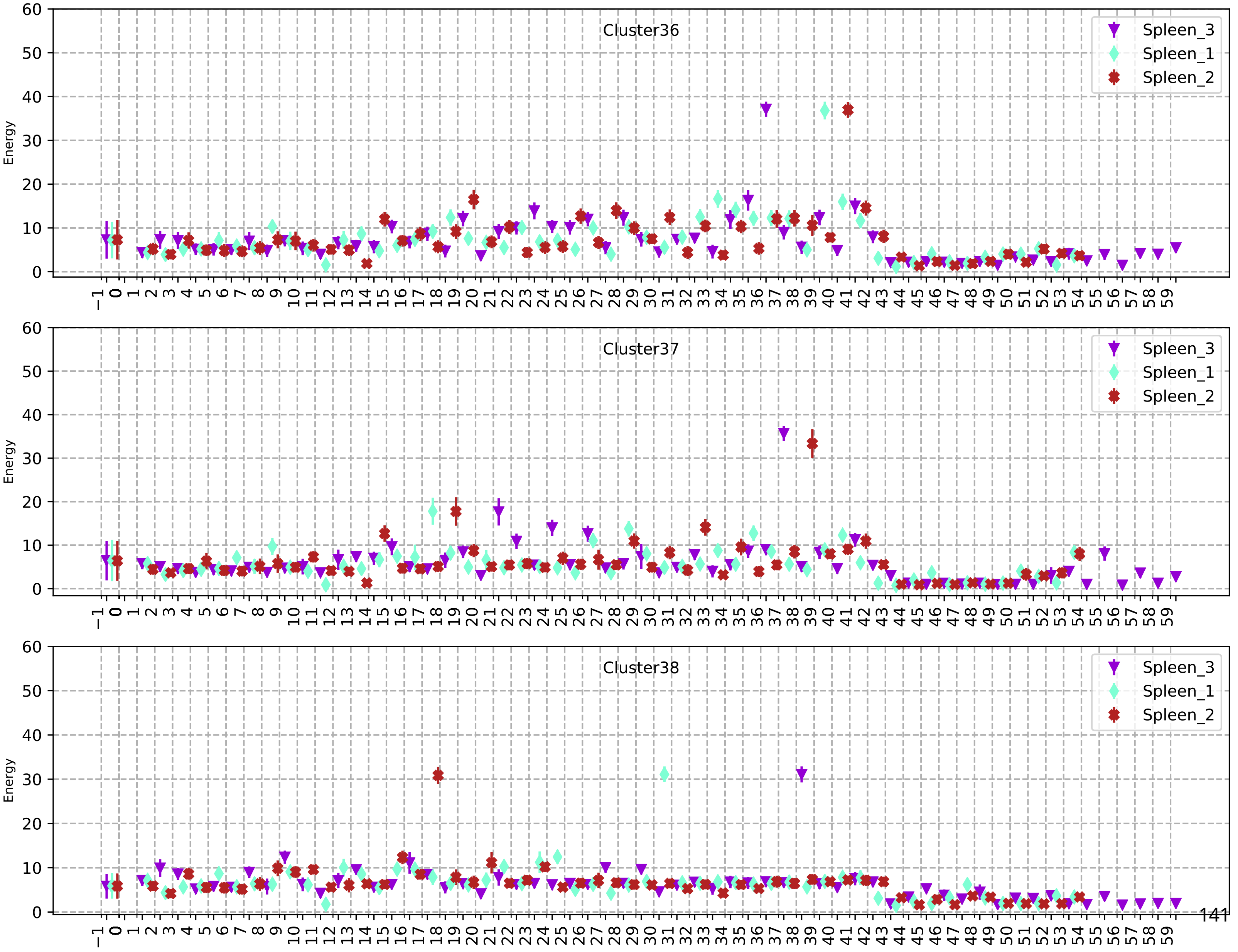
Cluster36
Spleen_3
Spleen_1
Spleen_2
Energy
Cluster37
Spleen_3
Spleen_1
Spleen_2
Energy
Cluster38
Spleen_3
Spleen_1
Spleen_2
Energy

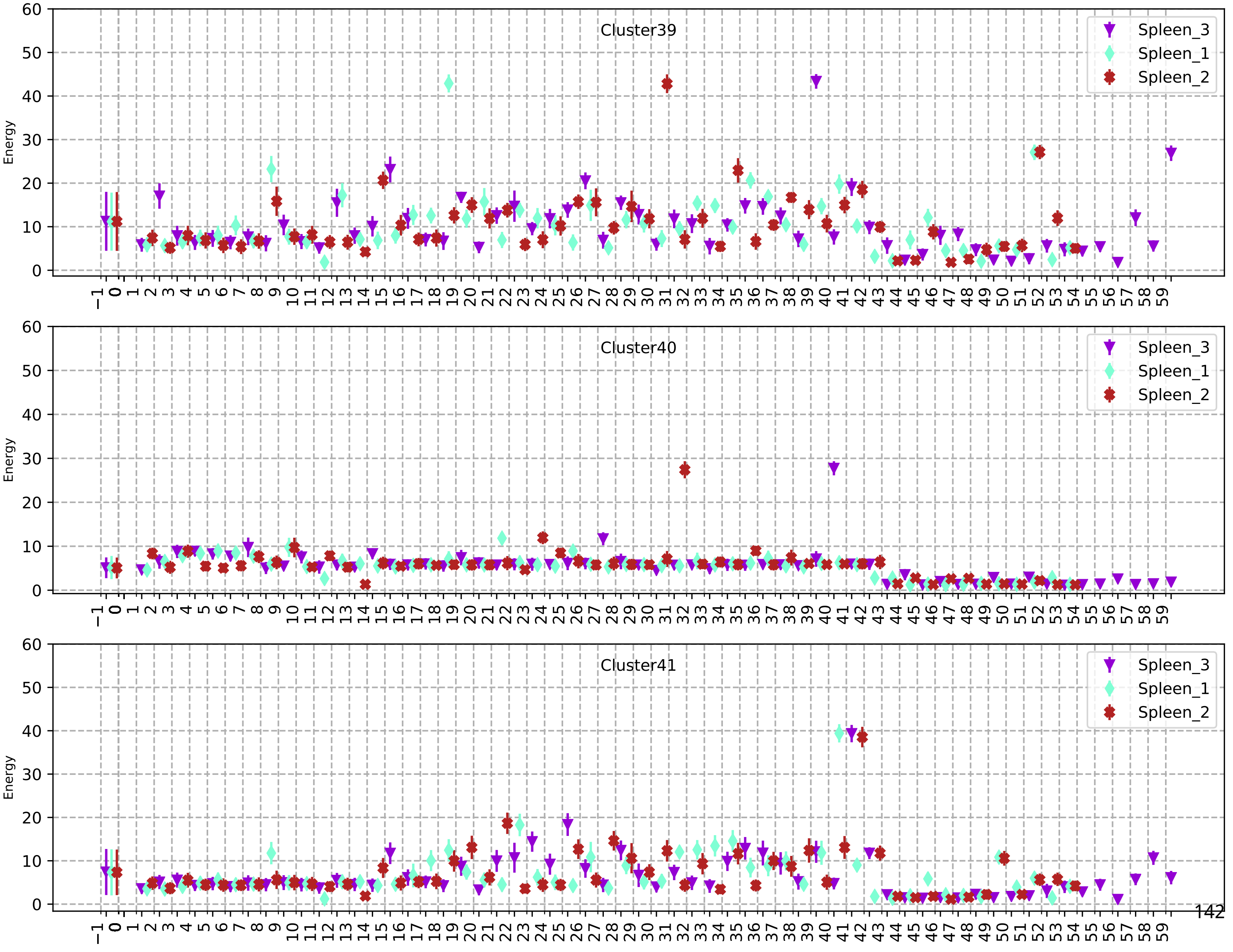
Cluster39
Cluster40
Cluster41
Spleen_3
Spleen_1
Spleen_2
Energy
0 10 20 30 40 50 60
−1 0 1 2 3 4 5 6 7 8 9 10 11 12 13 14 15 16 17 18 19 20 21 22 23 24 25 26 27 28 29 30 31 32 33 34 35 36 37 38 39 40 41 42 43 44 45 46 47 48 49 50 51 52 53 54 55 56 57 58 59

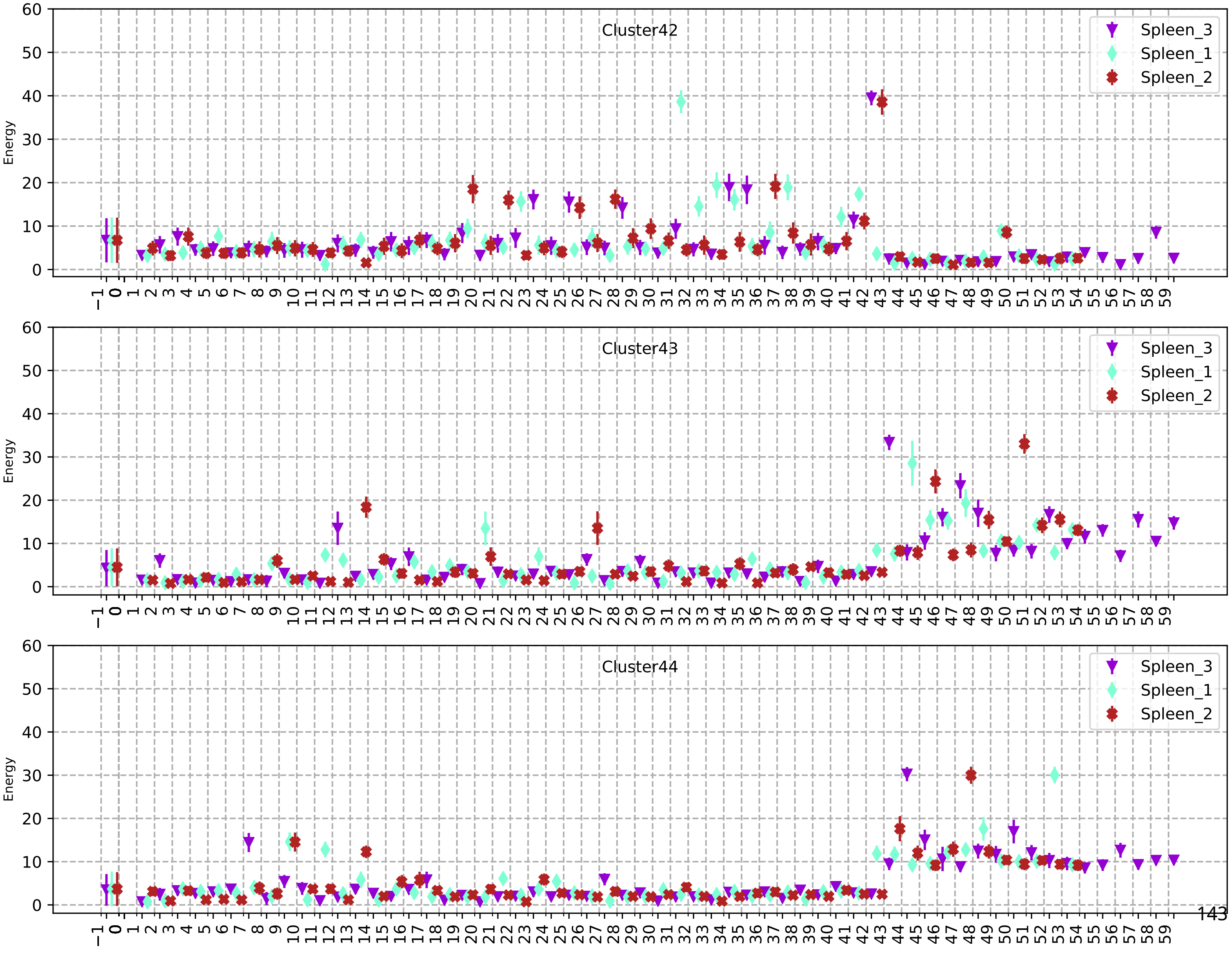
Cluster42
Cluster43
Cluster44
Spleen_3
Spleen_1
Spleen_2
Energy

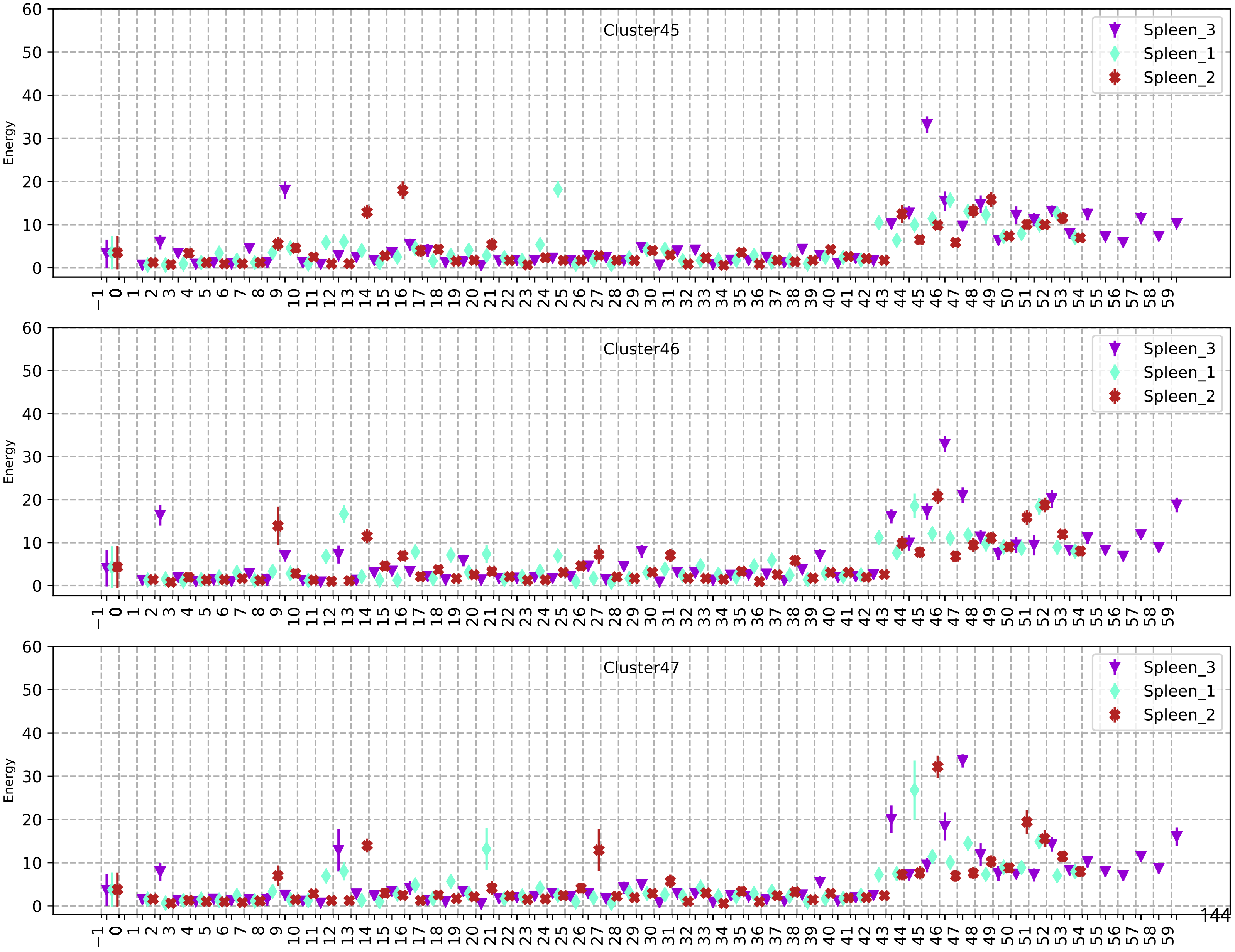
Cluster45
Cluster46
Cluster47
Energy
Spleen_3
Spleen_1
Spleen_2

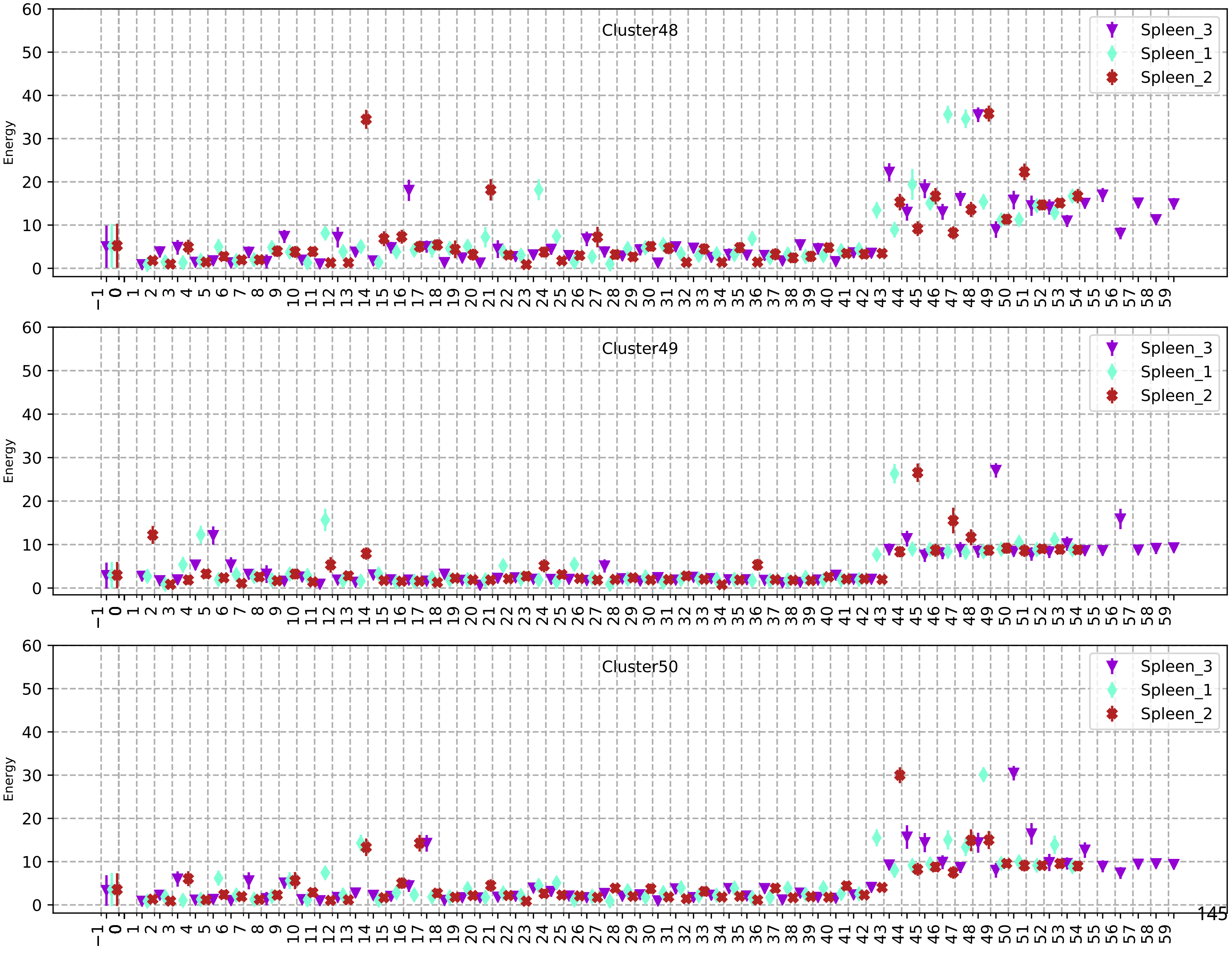
Cluster48
Spleen_3
Spleen_1
Spleen_2
Energy
Cluster49
Spleen_3
Spleen_1
Spleen_2
Energy
Cluster50
Spleen_3
Spleen_1
Spleen_2
Energy

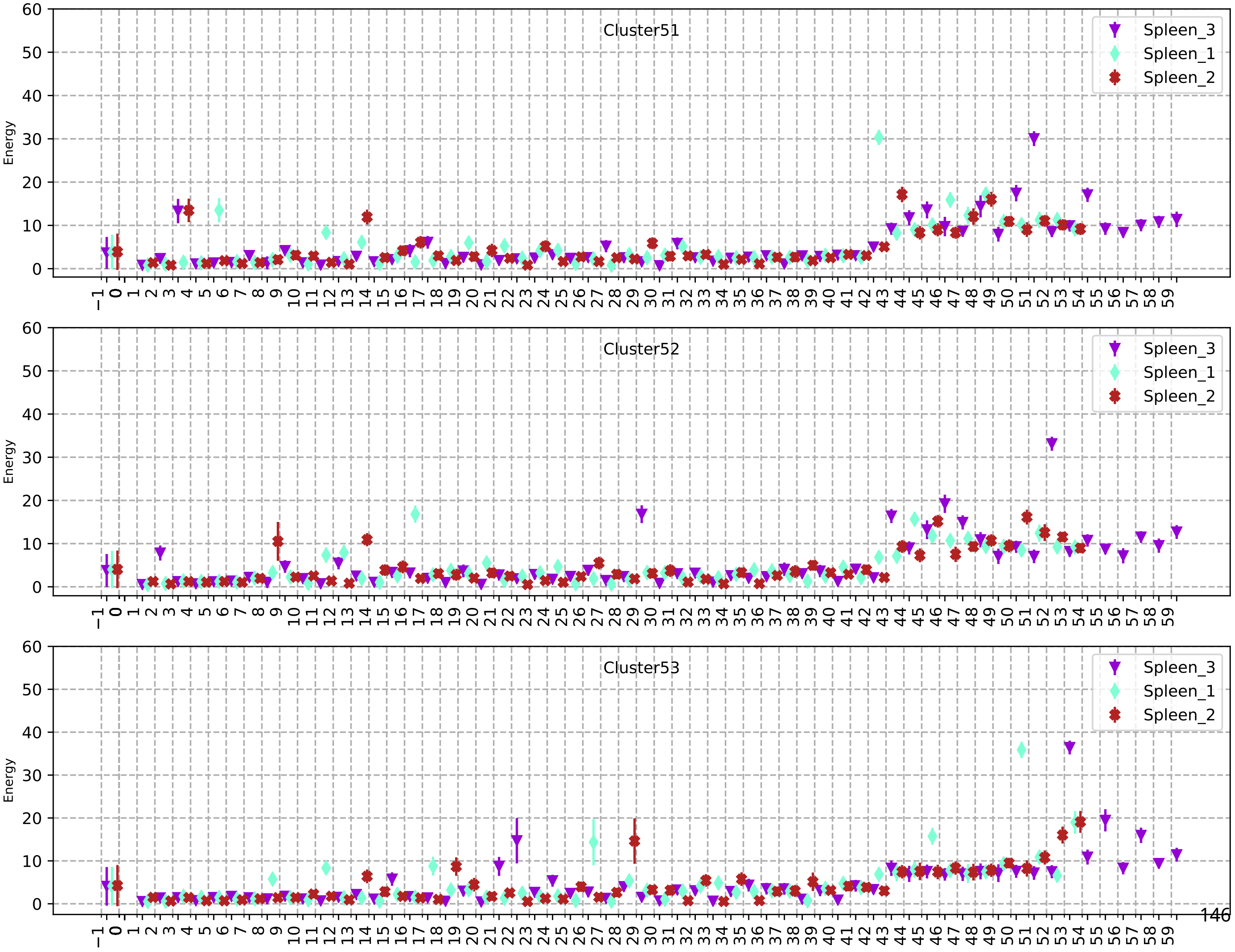
Cluster51
Cluster52
Cluster53
Energy
Spleen_3
Spleen_1
Spleen_2

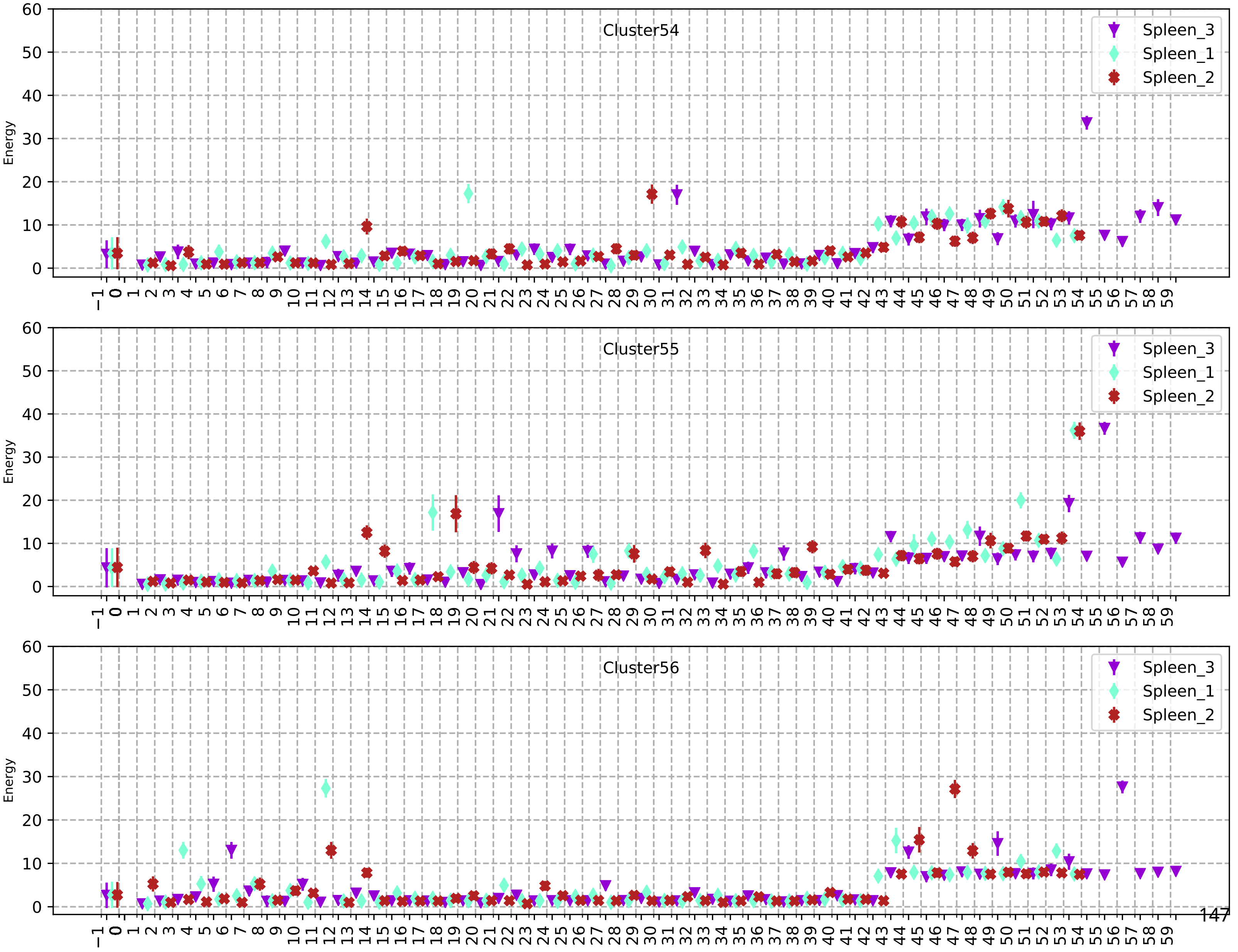
Cluster54
Cluster55
Cluster56
Spleen_3
Spleen_1
Spleen_2
Energy
0
10
20
30
40
50
60
−1 0 1 2 3 4 5 6 7 8 9 10 11 12 13 14 15 16 17 18 19 20 21 22 23 24 25 26 27 28 29 30 31 32 33 34 35 36 37 38 39 40 41 42 43 44 45 46 47 48 49 50 51 52 53 54 55 56 57 58 59

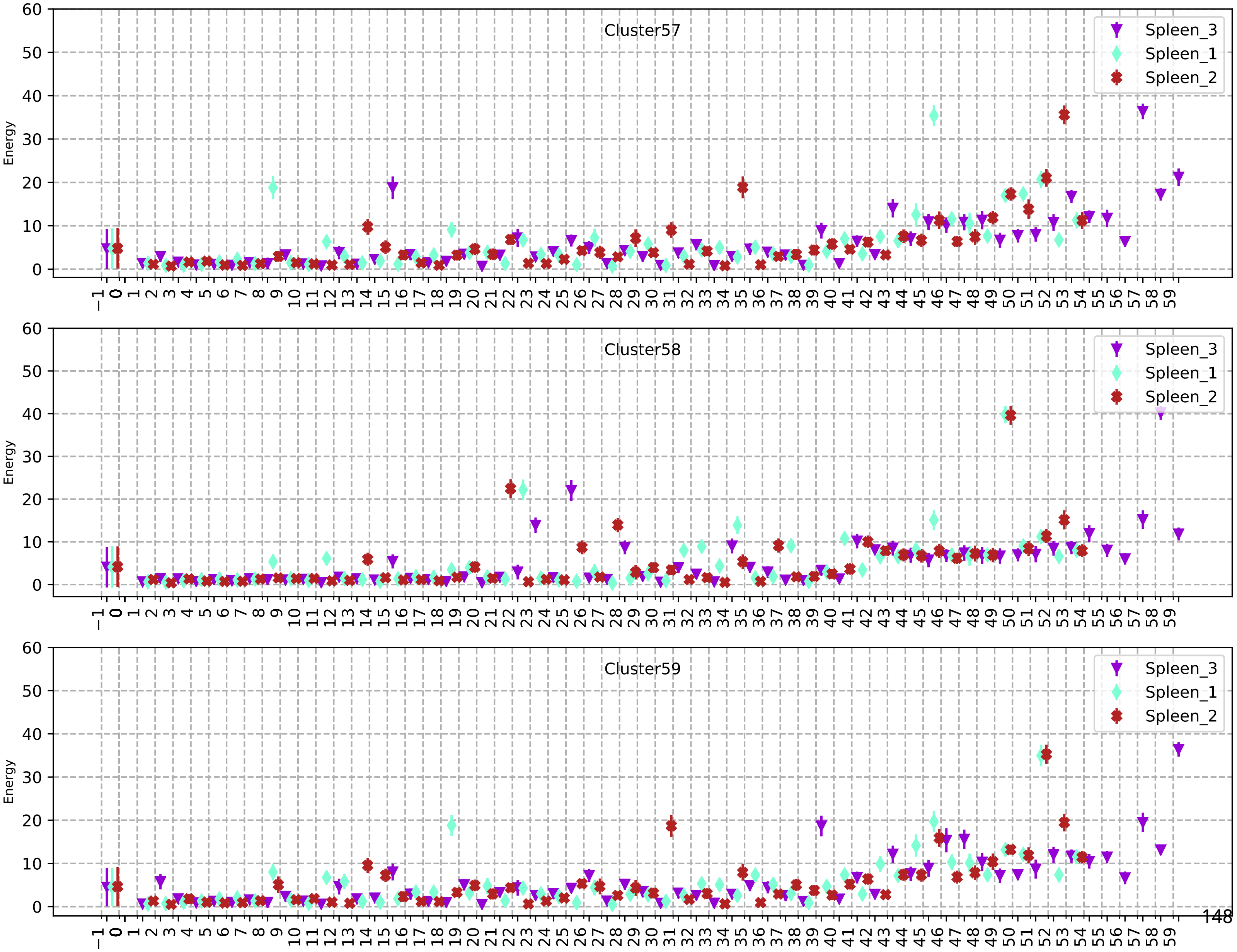
Cluster57
Cluster58
Cluster59
Energy
Spleen_3
Spleen_1
Spleen_2

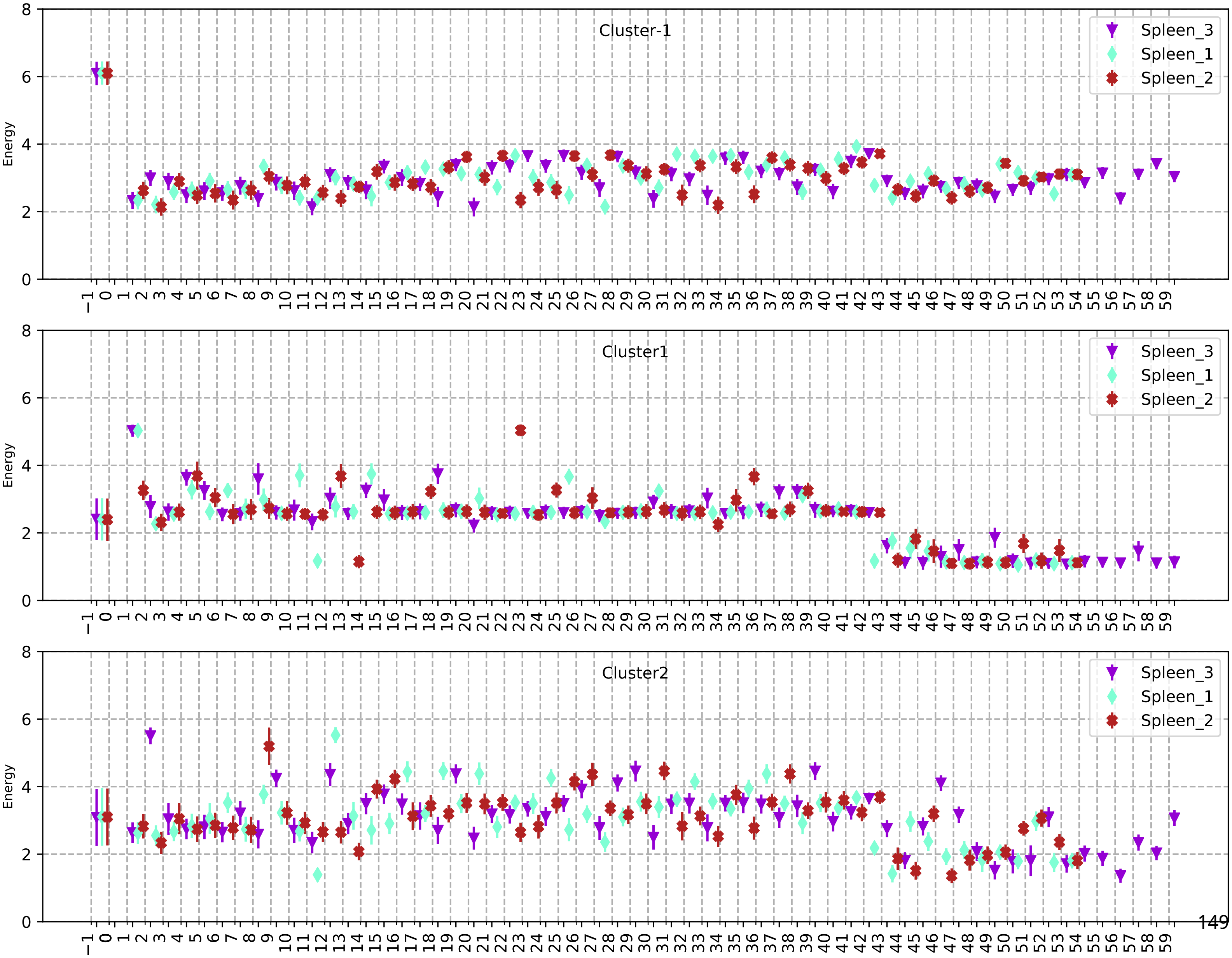
Cluster-1
Cluster1
Cluster2
Spleen_3
Spleen_1
Spleen_2
Energy

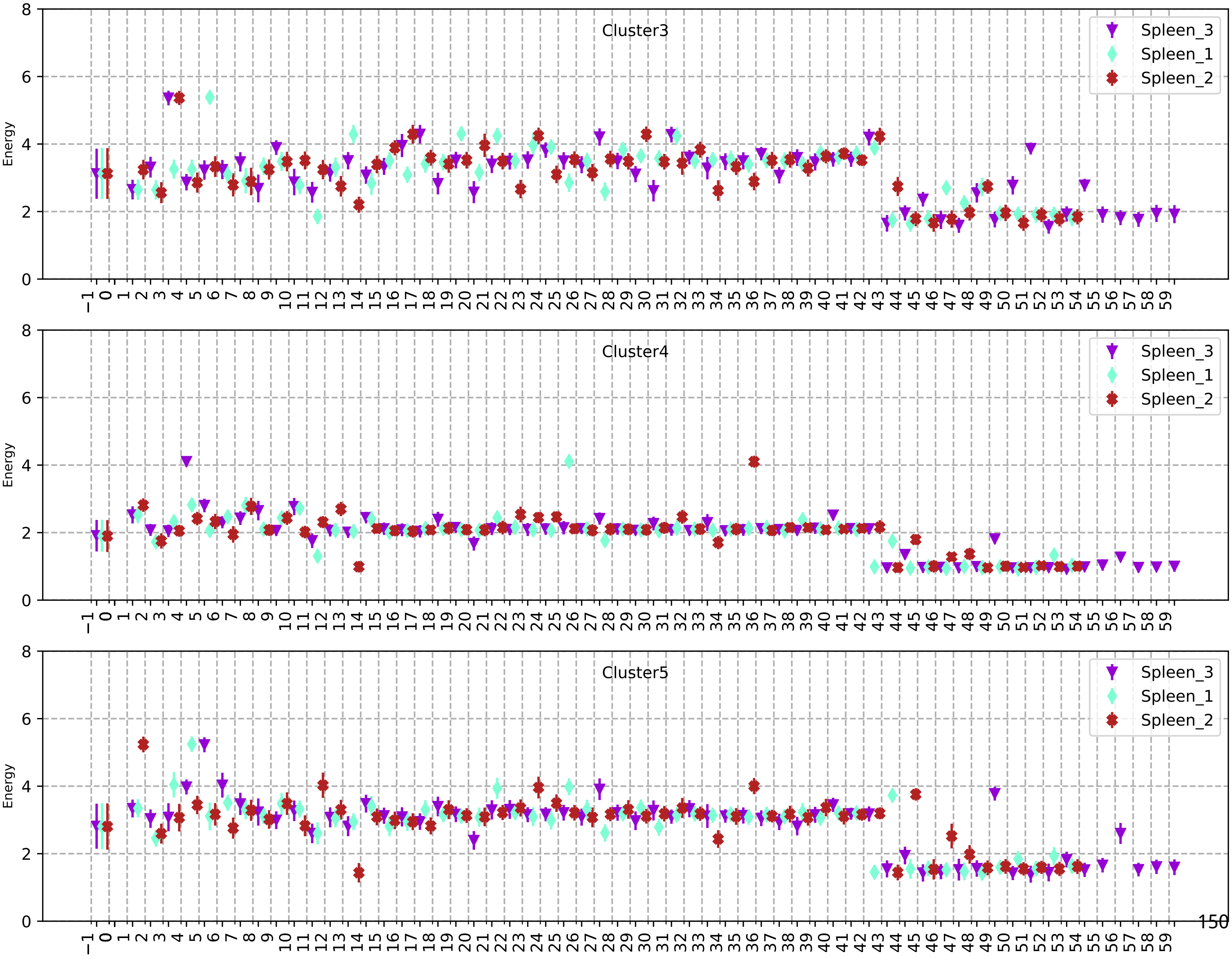
Cluster3
Spleen_3
Spleen_1
Spleen_2
Energy
Cluster4
Spleen_3
Spleen_1
Spleen_2
Energy
Cluster5
Spleen_3
Spleen_1
Spleen_2
Energy

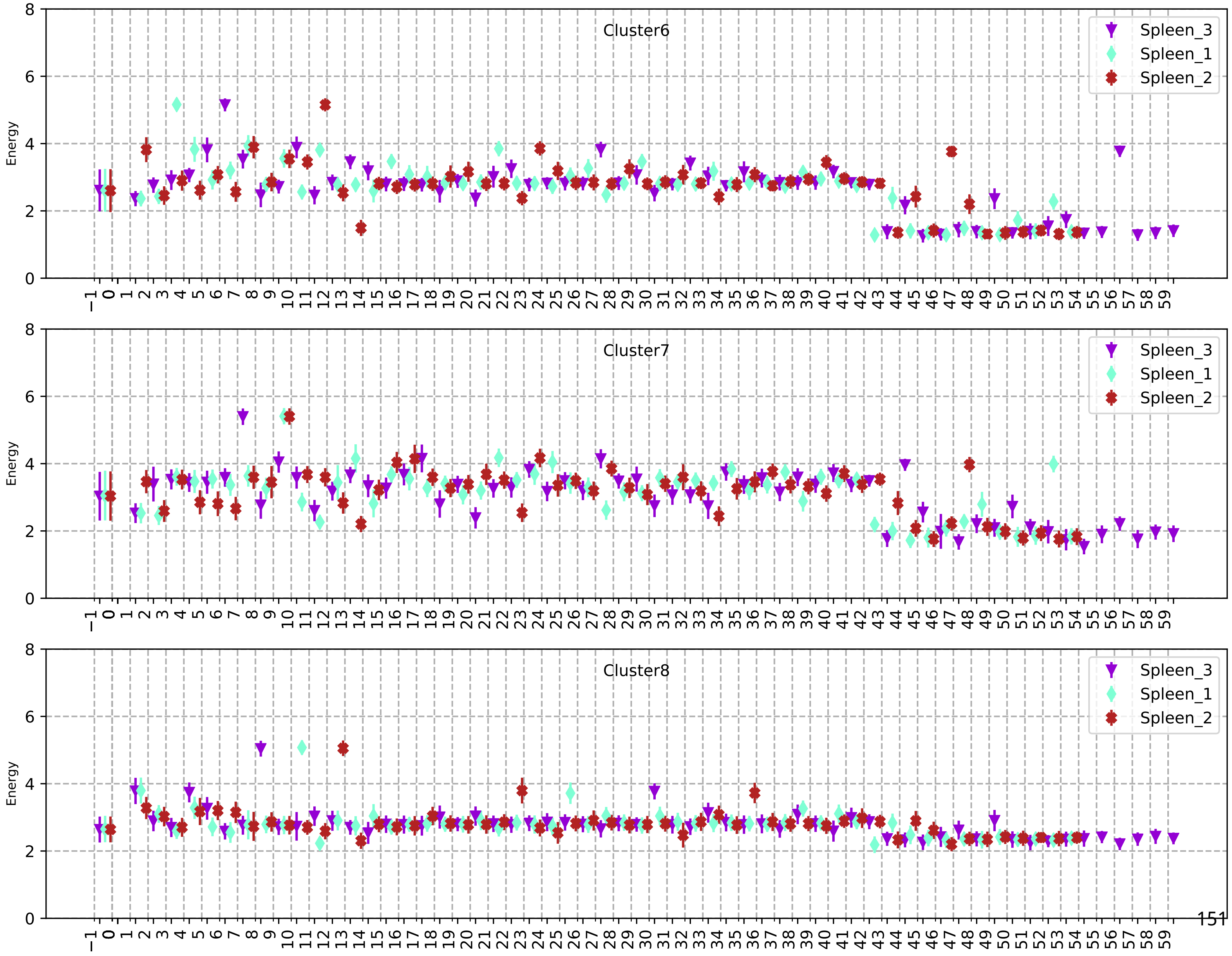
Cluster6
Spleen_3
Spleen_1
Spleen_2
Energy
Cluster7
Spleen_3
Spleen_1
Spleen_2
Energy
Cluster8
Spleen_3
Spleen_1
Spleen_2
Energy

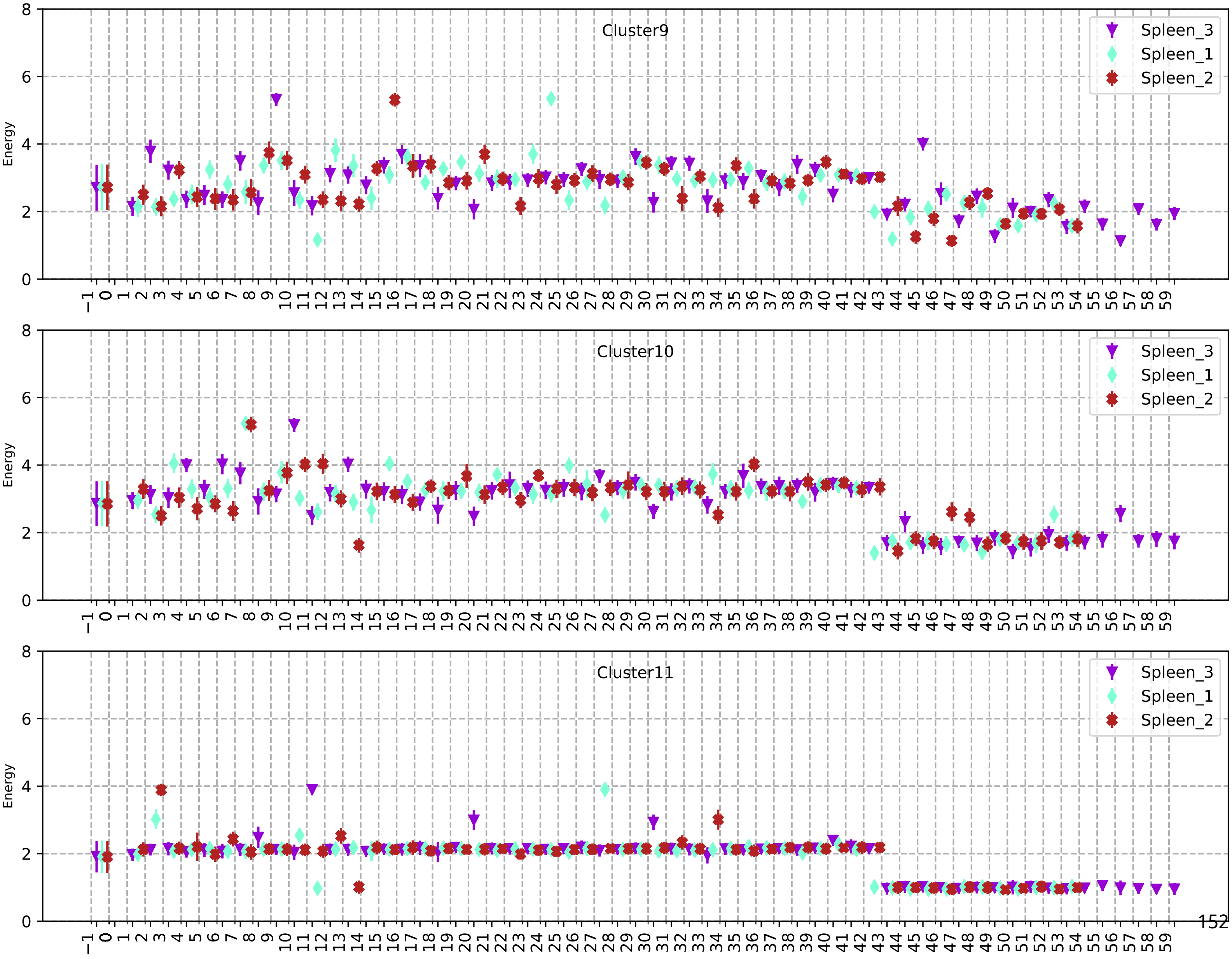
Cluster9
Energy
Spleen_3
Spleen_1
Spleen_2
Cluster10
Energy
Spleen_3
Spleen_1
Spleen_2
Cluster11
Energy
Spleen_3
Spleen_1
Spleen_2

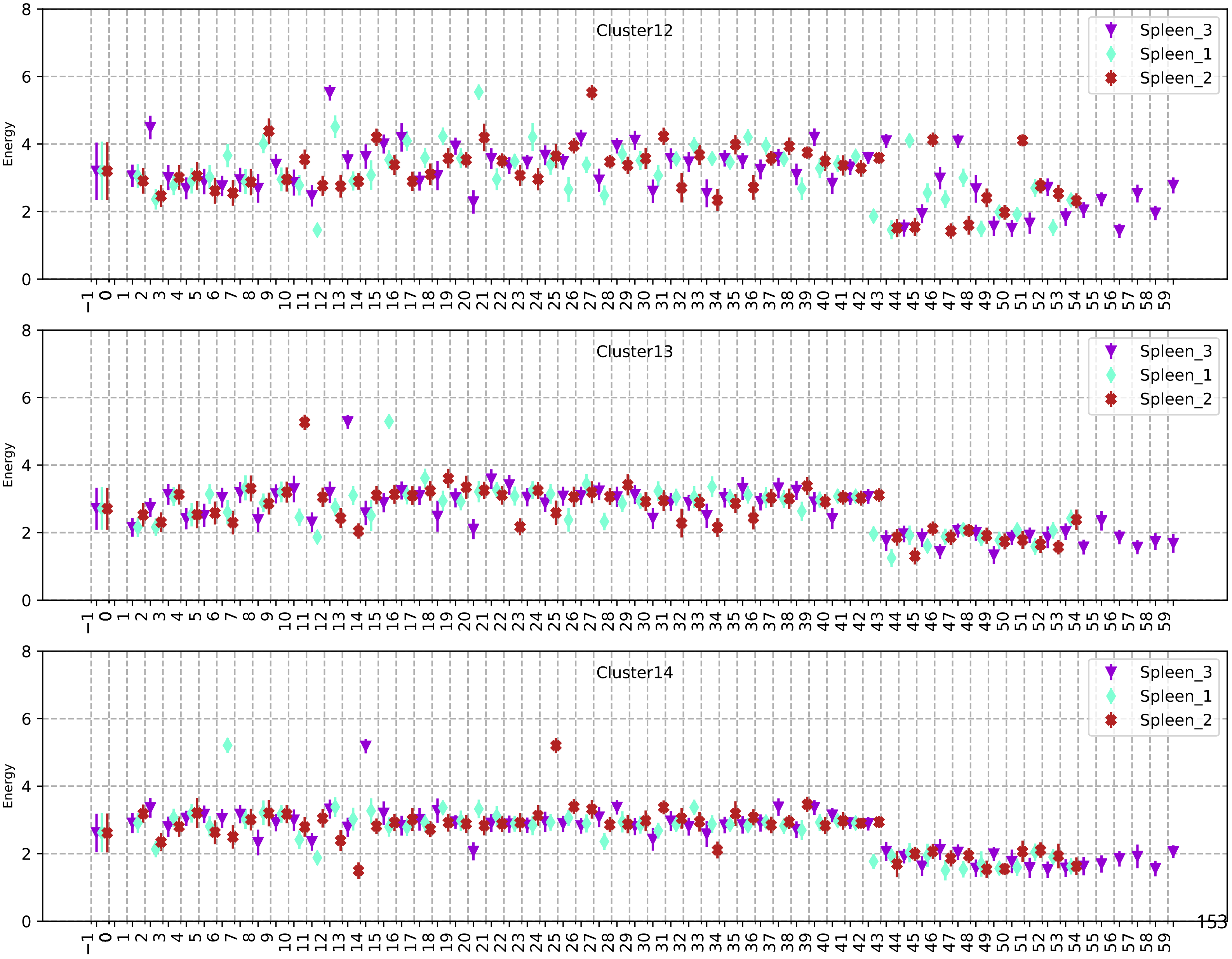
Cluster12
Cluster13
Cluster14
Energy
Spleen_3
Spleen_1
Spleen_2

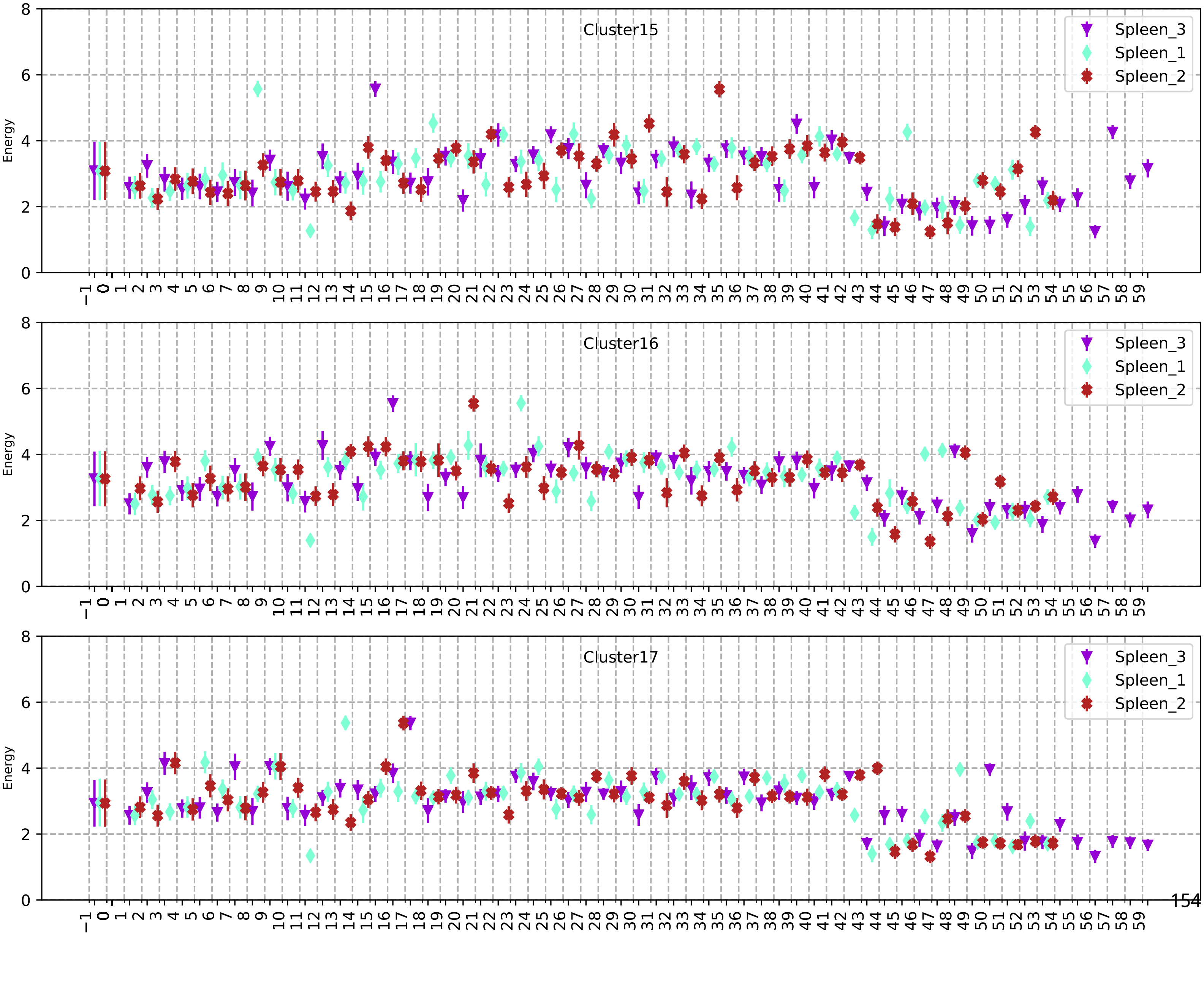
Cluster15
Spleen_3
Spleen_1
Spleen_2
Energy
Cluster16
Spleen_3
Spleen_1
Spleen_2
Energy
Cluster17
Spleen_3
Spleen_1
Spleen_2
Energy

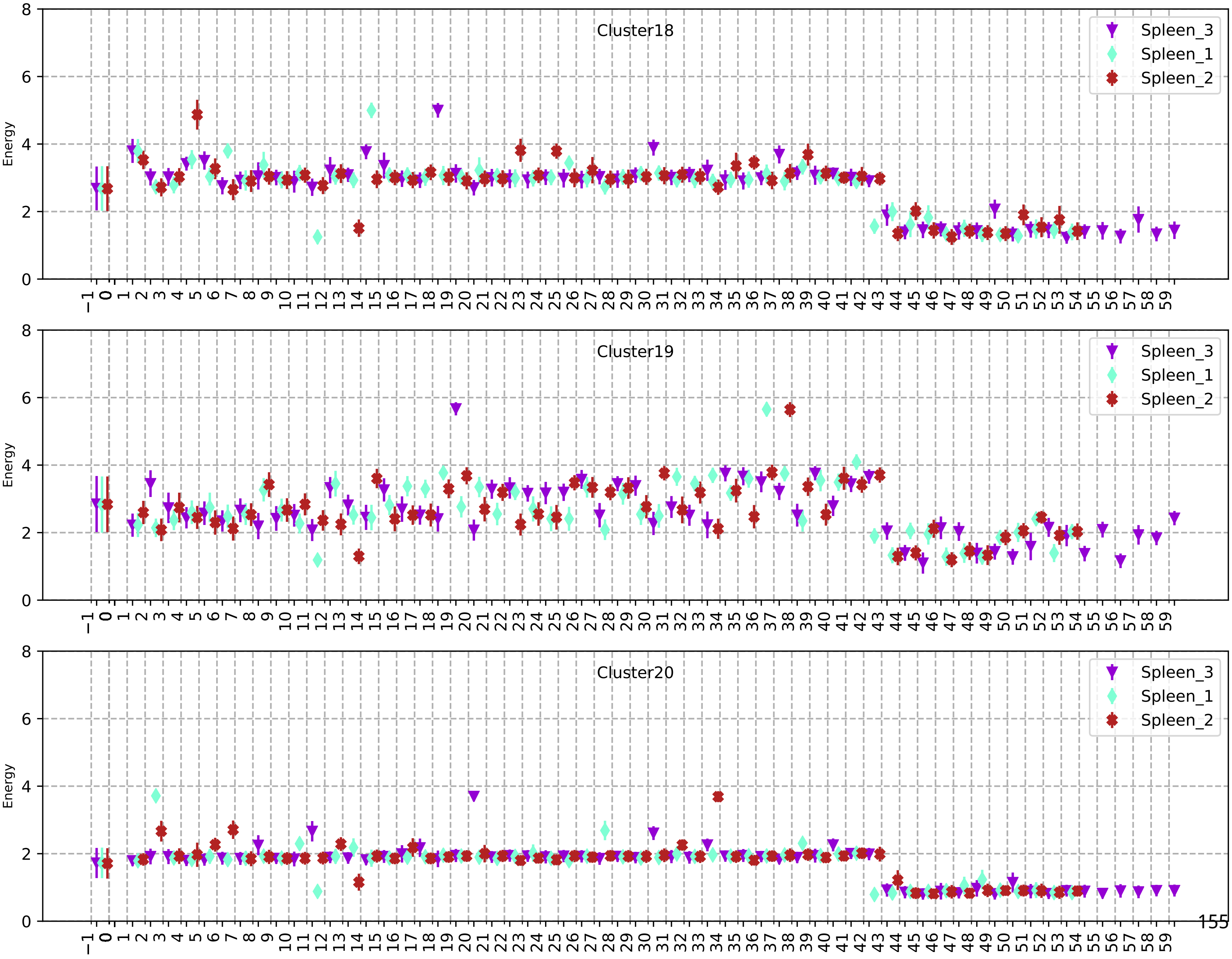
Cluster18
Cluster19
Cluster20
Energy
Spleen_3
Spleen_1
Spleen_2
0
2
4
6
8
−1 0 1 2 3 4 5 6 7 8 9 10 11 12 13 14 15 16 17 18 19 20 21 22 23 24 25 26 27 28 29 30 31 32 33 34 35 36 37 38 39 40 41 42 43 44 45 46 47 48 49 50 51 52 53 54 55 56 57 58 59

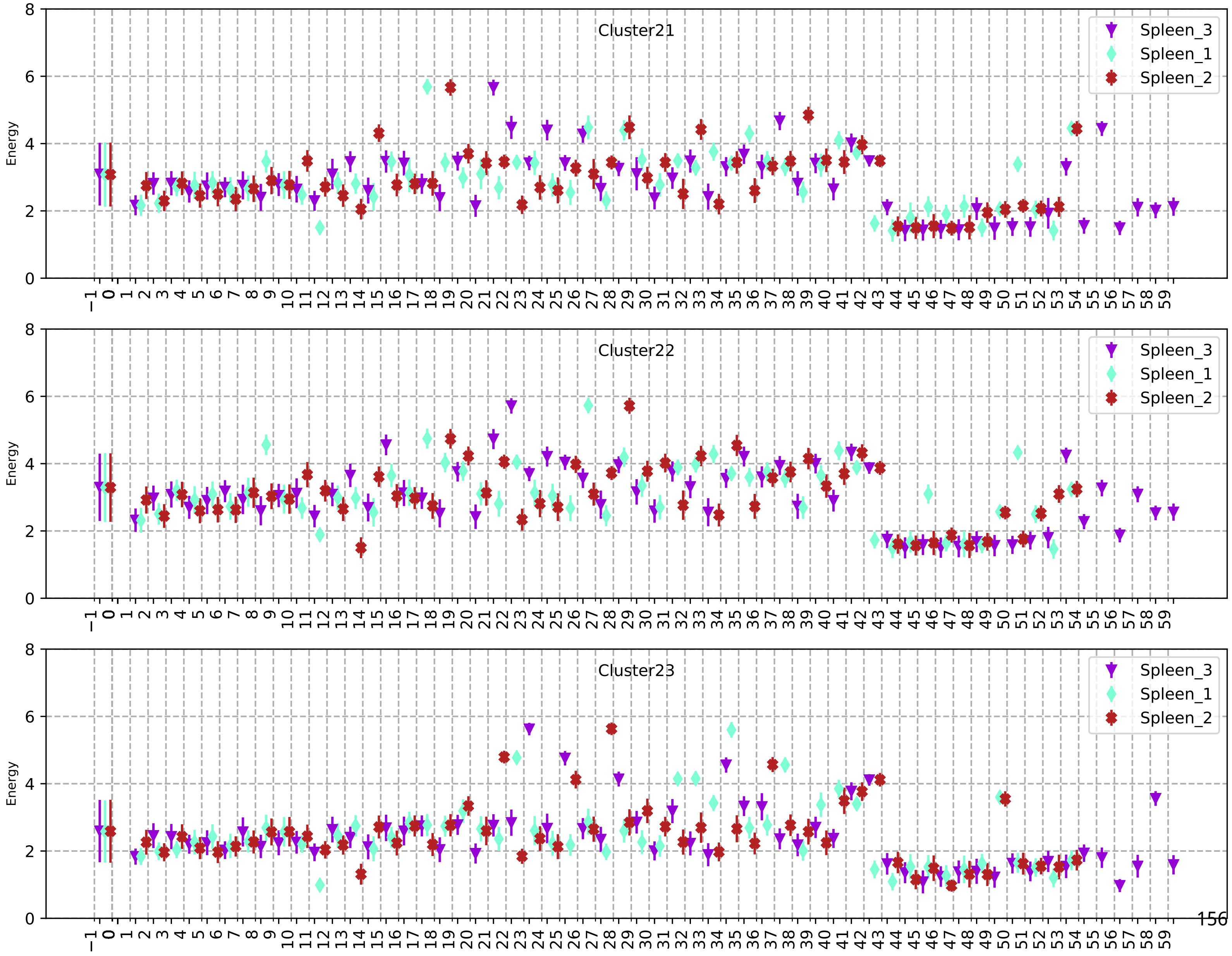
Cluster21
Cluster22
Cluster23
Spleen_3
Spleen_1
Spleen_2
Energy

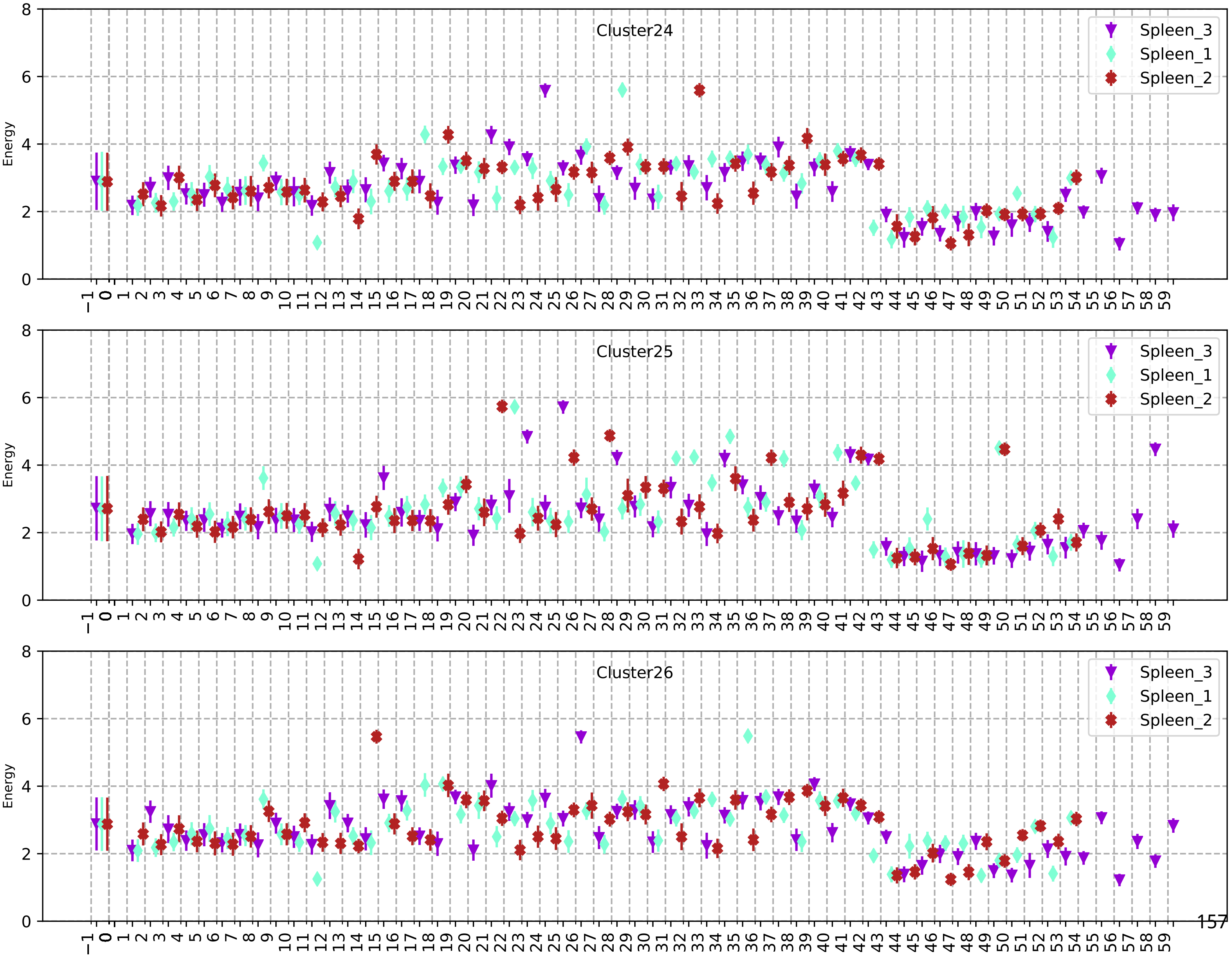
Cluster24
Cluster25
Cluster26
Energy
Spleen_3
Spleen_1
Spleen_2

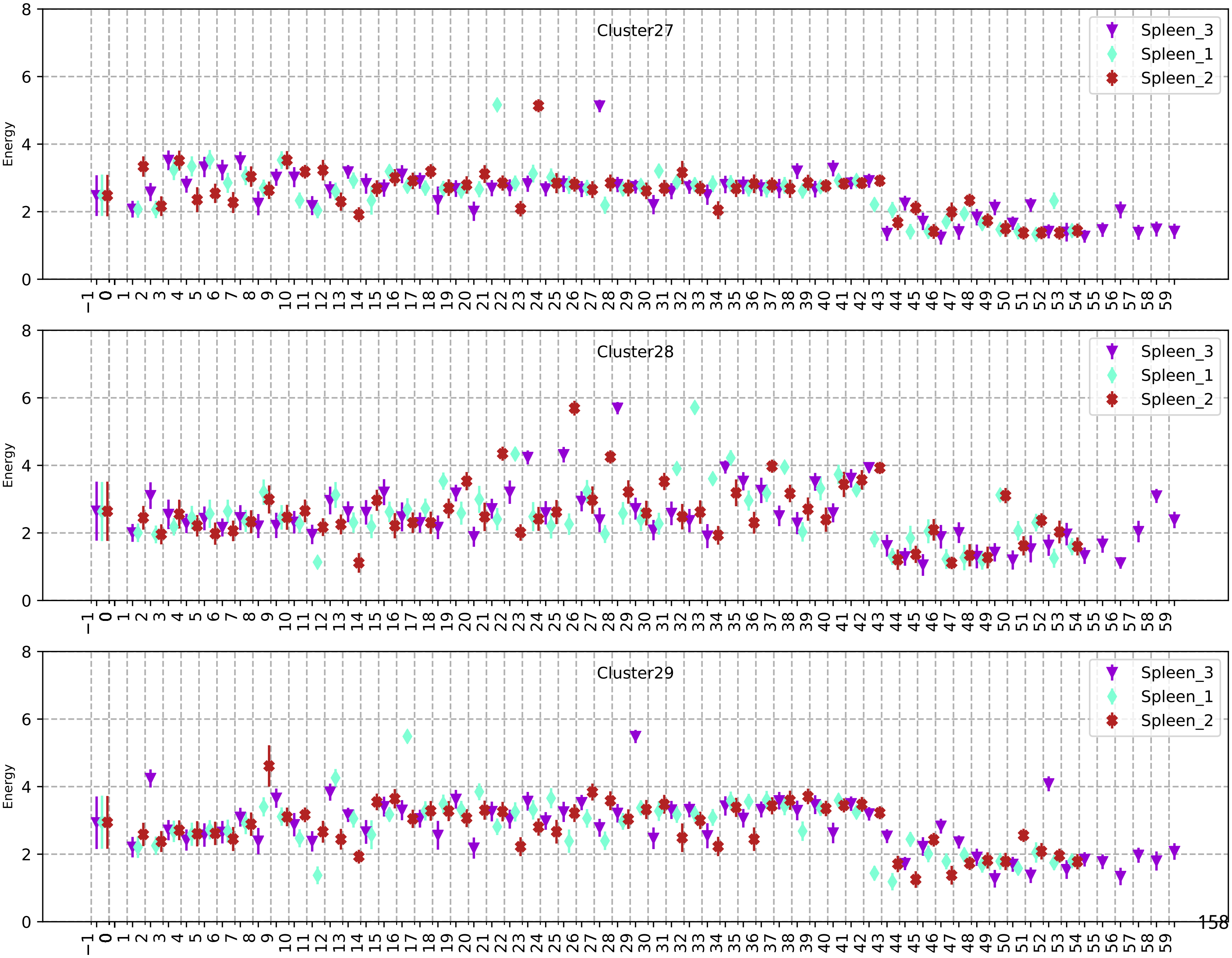
Cluster27
Spleen_3
Spleen_1
Spleen_2
Energy
Cluster28
Spleen_3
Spleen_1
Spleen_2
Energy
Cluster29
Spleen_3
Spleen_1
Spleen_2
Energy

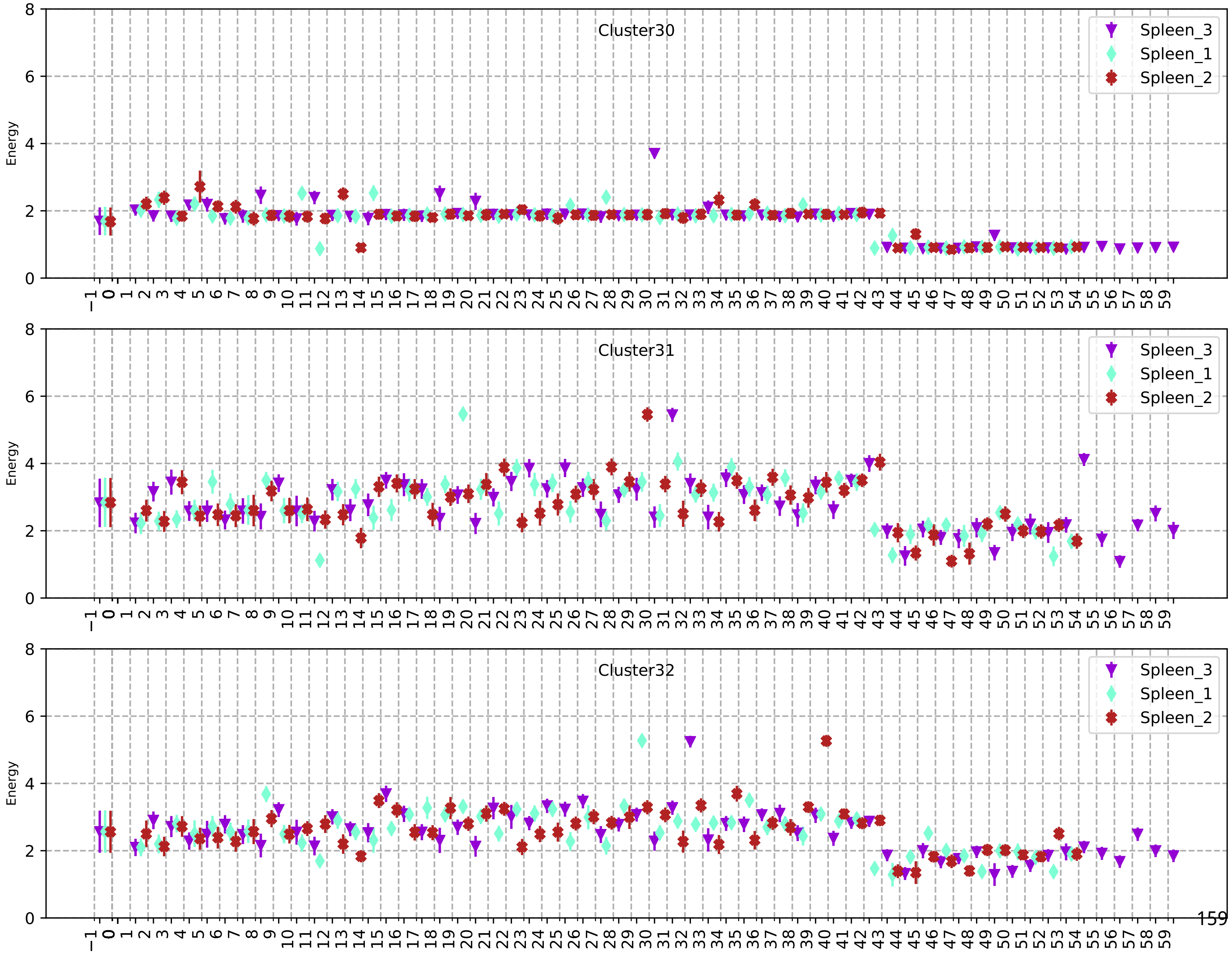
Cluster30
Cluster31
Cluster32
Spleen_3
Spleen_1
Spleen_2
Energy
0
2
4
6
8
−1 0 1 2 3 4 5 6 7 8 9 10 11 12 13 14 15 16 17 18 19 20 21 22 23 24 25 26 27 28 29 30 31 32 33 34 35 36 37 38 39 40 41 42 43 44 45 46 47 48 49 50 51 52 53 54 55 56 57 58 59

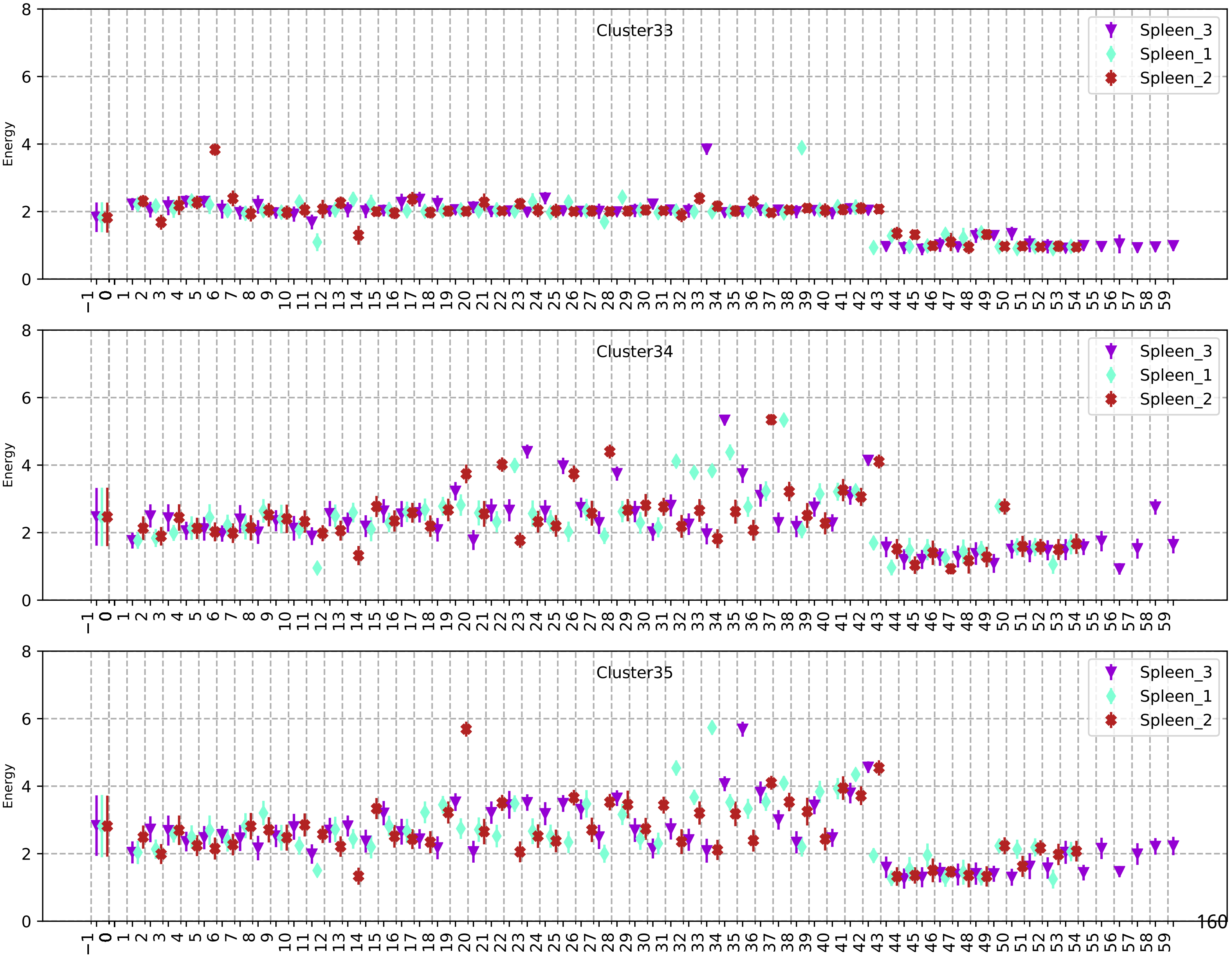
Cluster33
Energy
Spleen_3
Spleen_1
Spleen_2
Cluster34
Energy
Spleen_3
Spleen_1
Spleen_2
Cluster35
Energy
Spleen_3
Spleen_1
Spleen_2

207

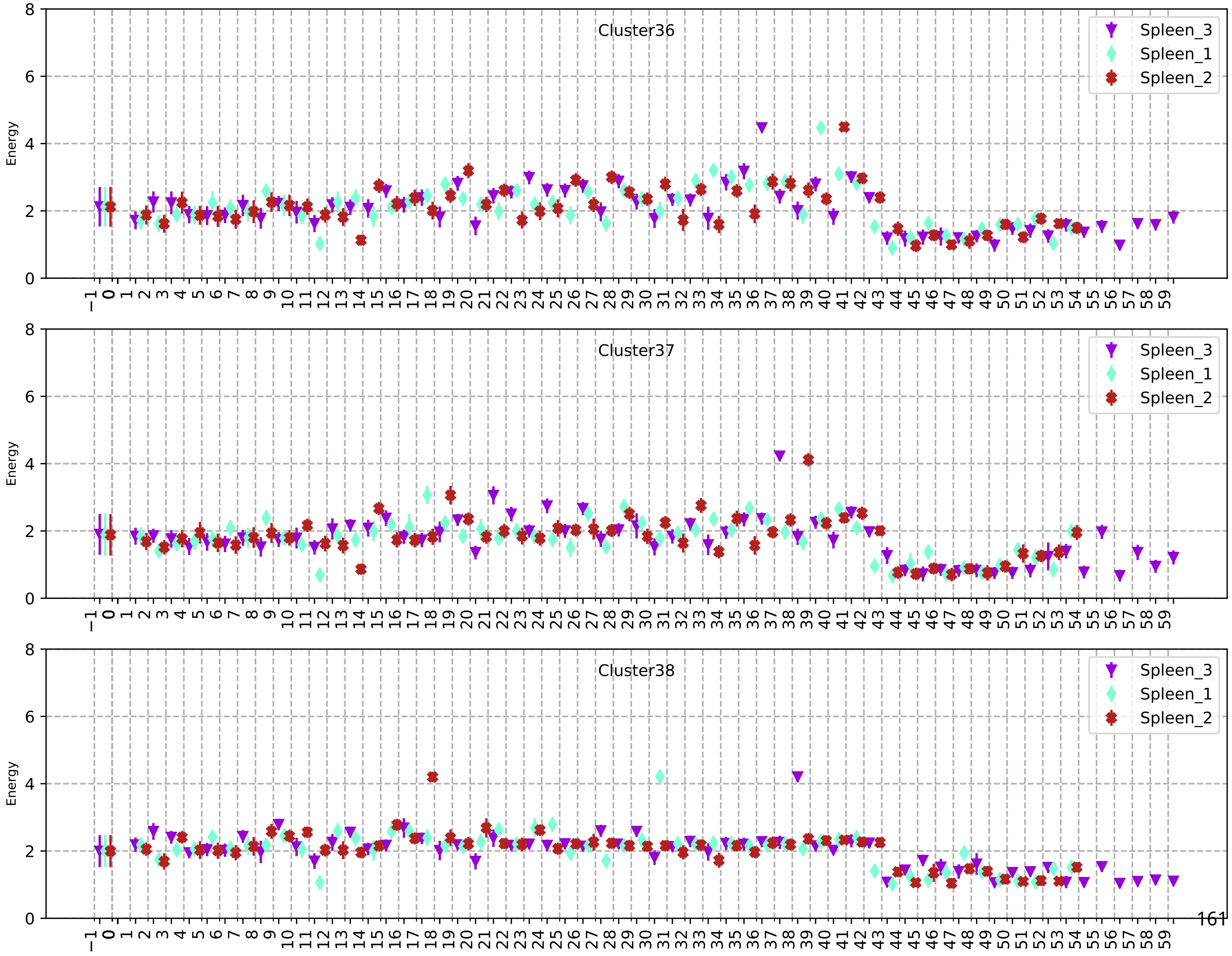
Cluster36
Cluster37
Cluster38
Energy
Spleen_3
Spleen_1
Spleen_2

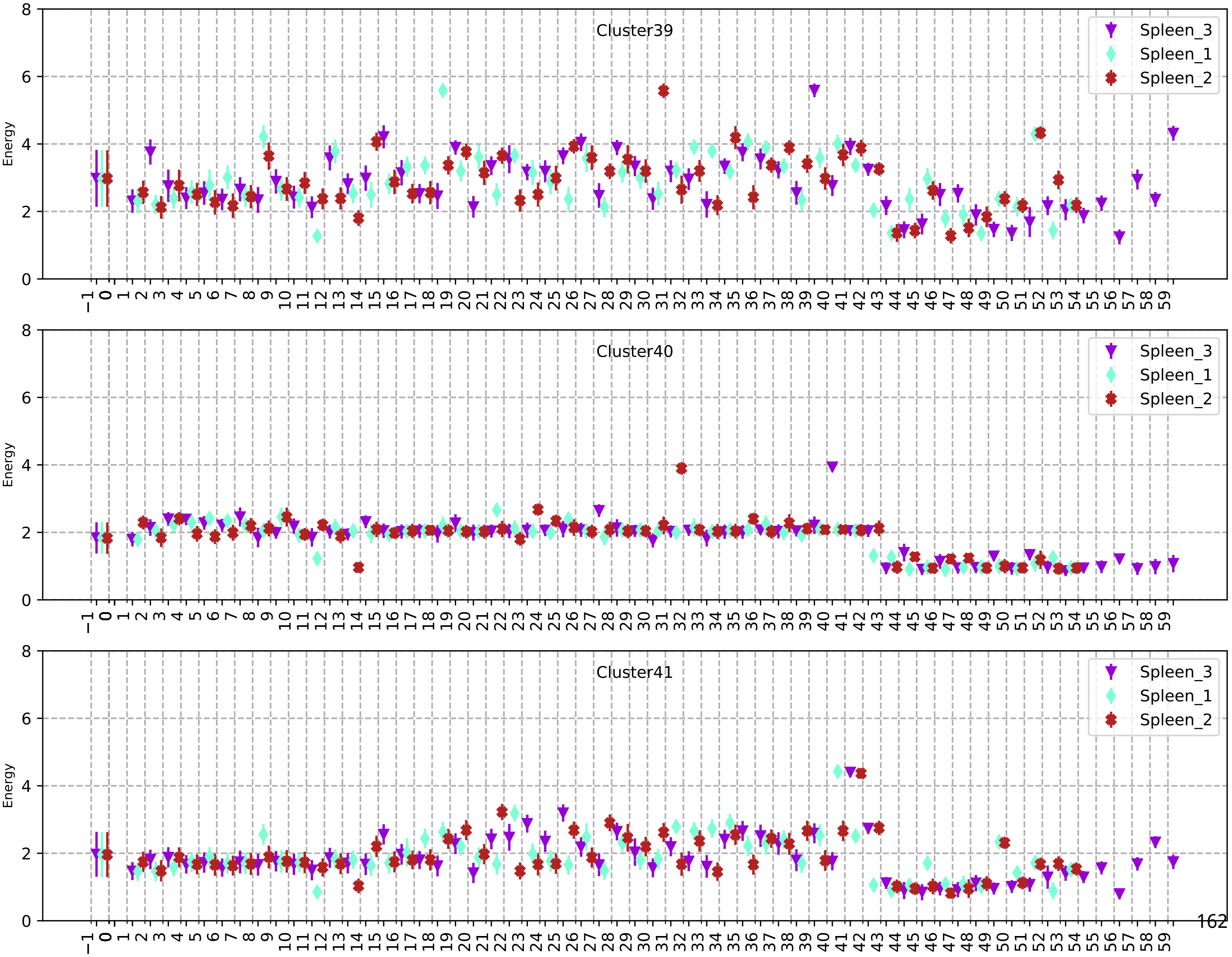
Cluster39
Cluster40
Cluster41
Spleen_3
Spleen_1
Spleen_2
Energy

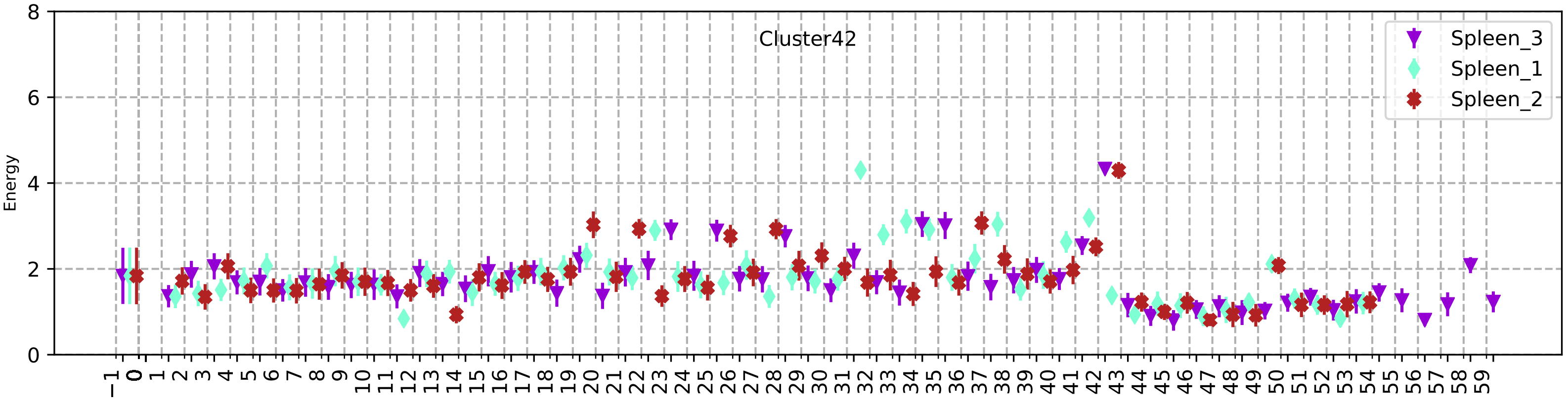
Cluster42
Spleen_3
Spleen_1
Spleen_2
Energy
0
2
4
6
8
−1 0 1 2 3 4 5 6 7 8 9 10 11 12 13 14 15 16 17 18 19 20 21 22 23 24 25 26 27 28 29 30 31 32 33 34 35 36 37 38 39 40 41 42 43 44 45 46 47 48 49 50 51 52 53 54 55 56 57 58 59

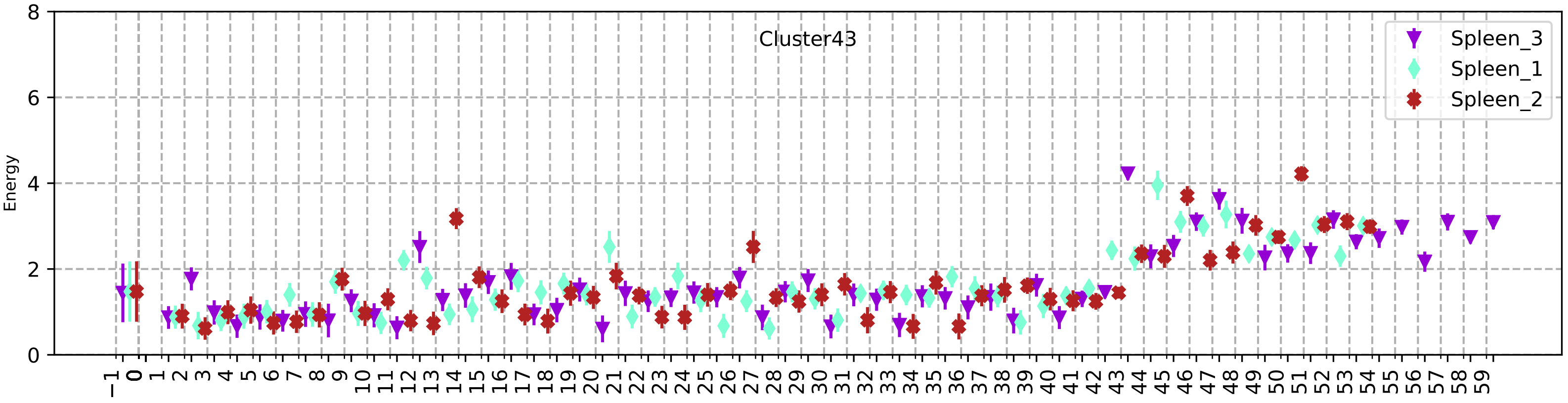
Cluster43
Spleen_3
Spleen_1
Spleen_2
Energy
0
2
4
6
8
−1 0 1 2 3 4 5 6 7 8 9 10 11 12 13 14 15 16 17 18 19 20 21 22 23 24 25 26 27 28 29 30 31 32 33 34 35 36 37 38 39 40 41 42 43 44 45 46 47 48 49 50 51 52 53 54 55 56 57 58 59

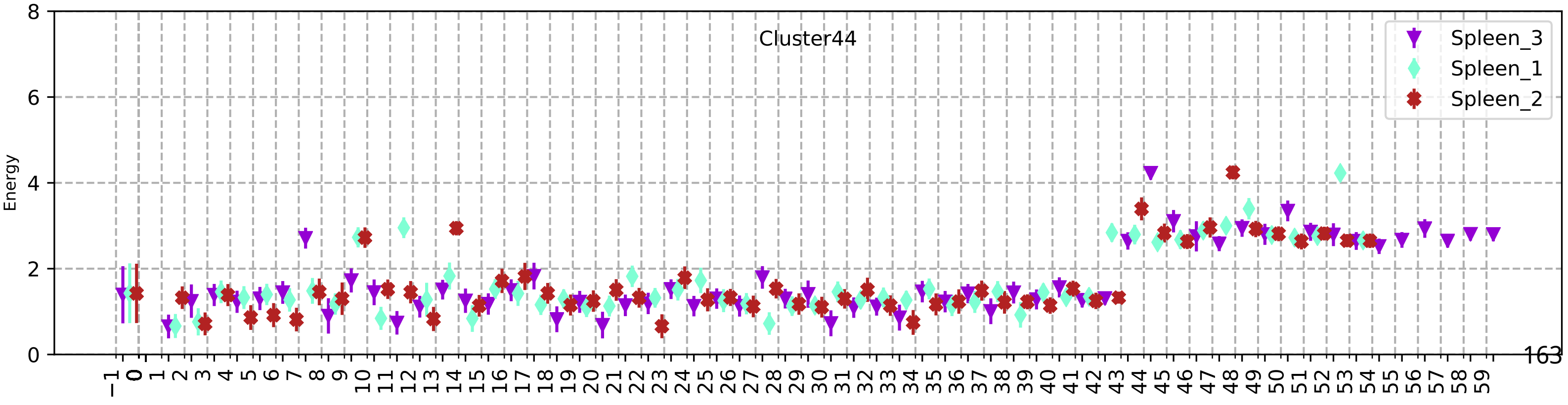
Cluster44
Spleen_3
Spleen_1
Spleen_2
Energy
0
2
4
6
8
−1 0 1 2 3 4 5 6 7 8 9 10 11 12 13 14 15 16 17 18 19 20 21 22 23 24 25 26 27 28 29 30 31 32 33 34 35 36 37 38 39 40 41 42 43 44 45 46 47 48 49 50 51 52 53 54 55 56 57 58 59

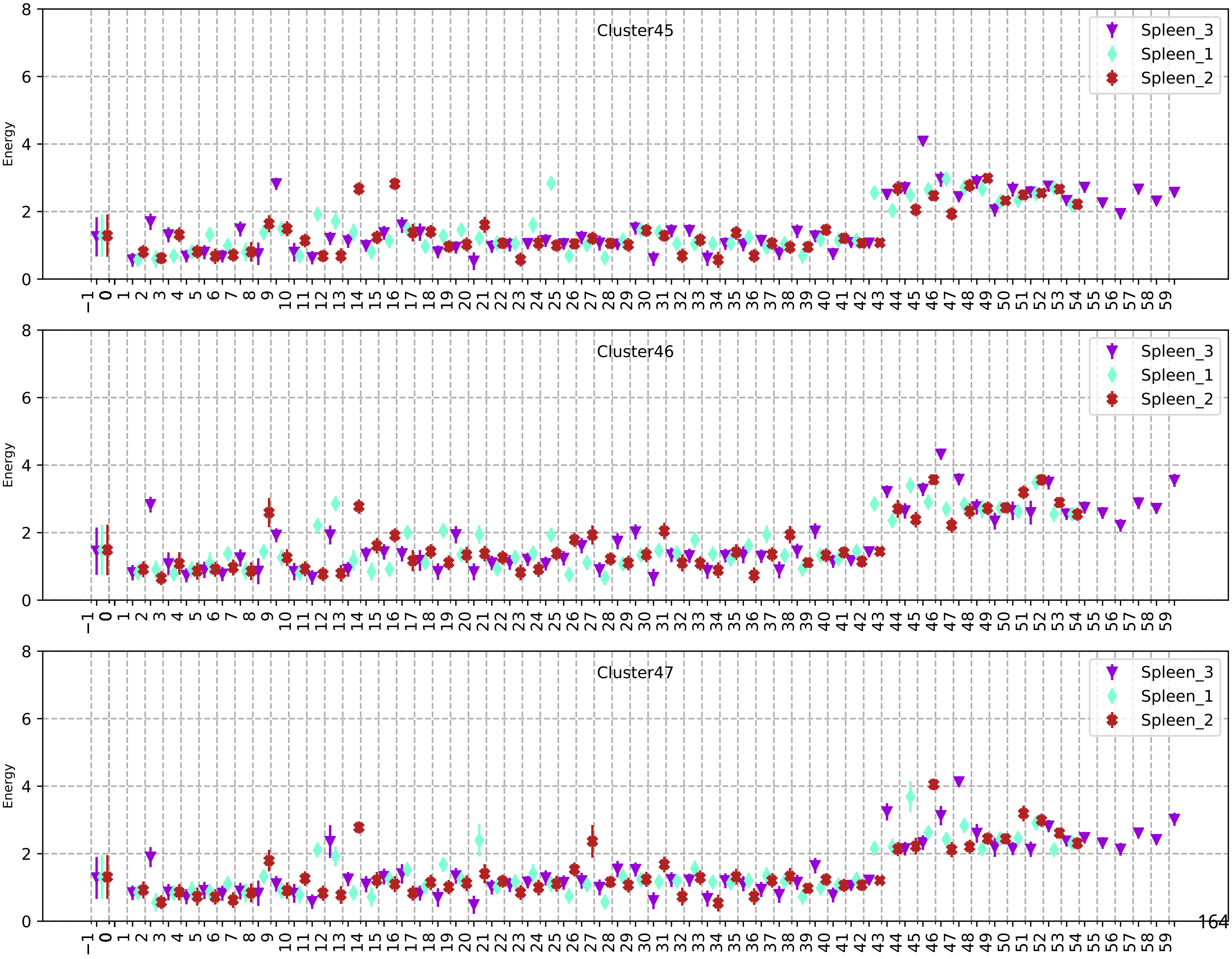
Cluster45
Spleen_3
Spleen_1
Spleen_2
Energy
Cluster46
Spleen_3
Spleen_1
Spleen_2
Energy
Cluster47
Spleen_3
Spleen_1
Spleen_2
Energy

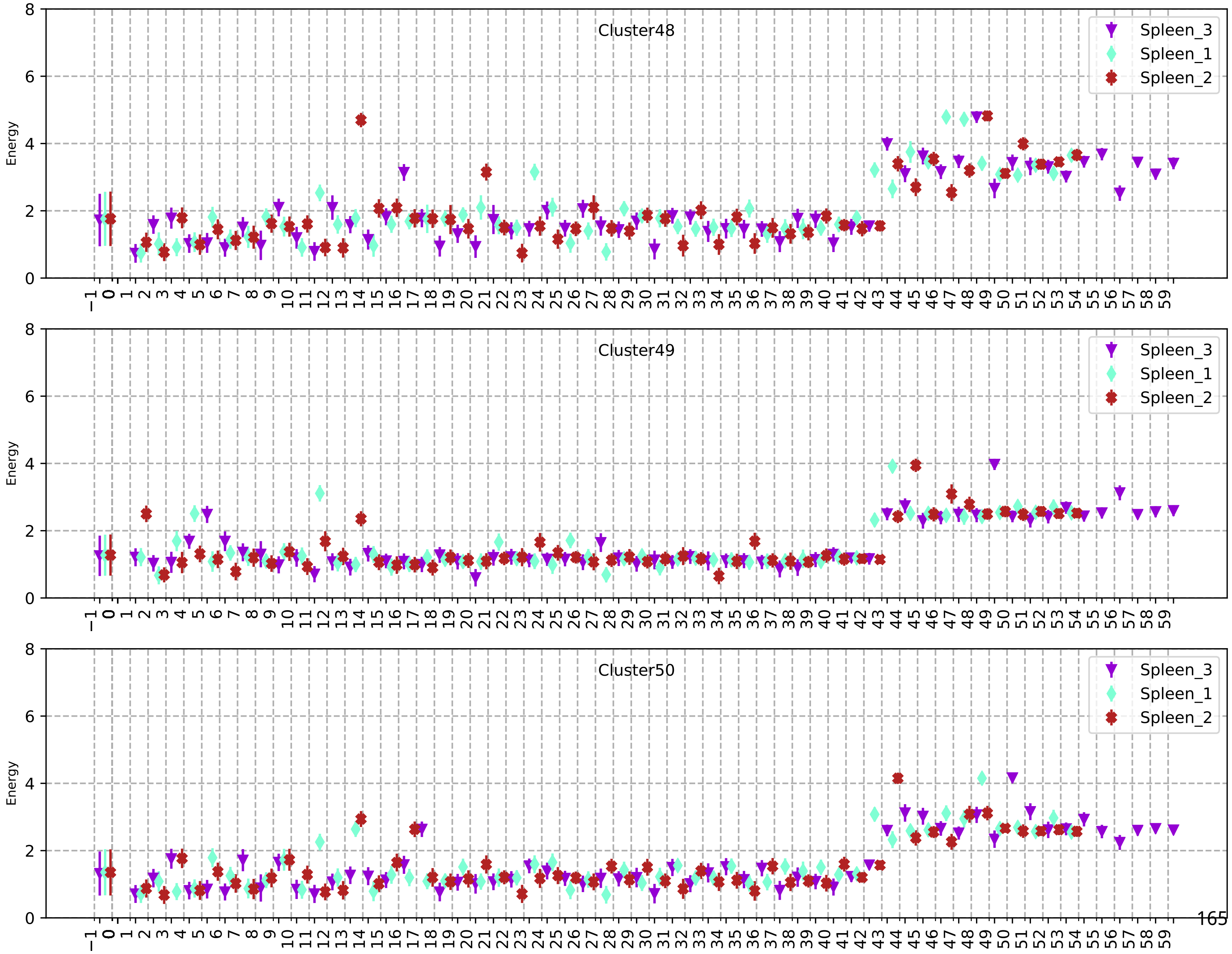
Cluster48
Spleen_3
Spleen_1
Spleen_2
Energy
Cluster49
Spleen_3
Spleen_1
Spleen_2
Energy
Cluster50
Spleen_3
Spleen_1
Spleen_2
Energy

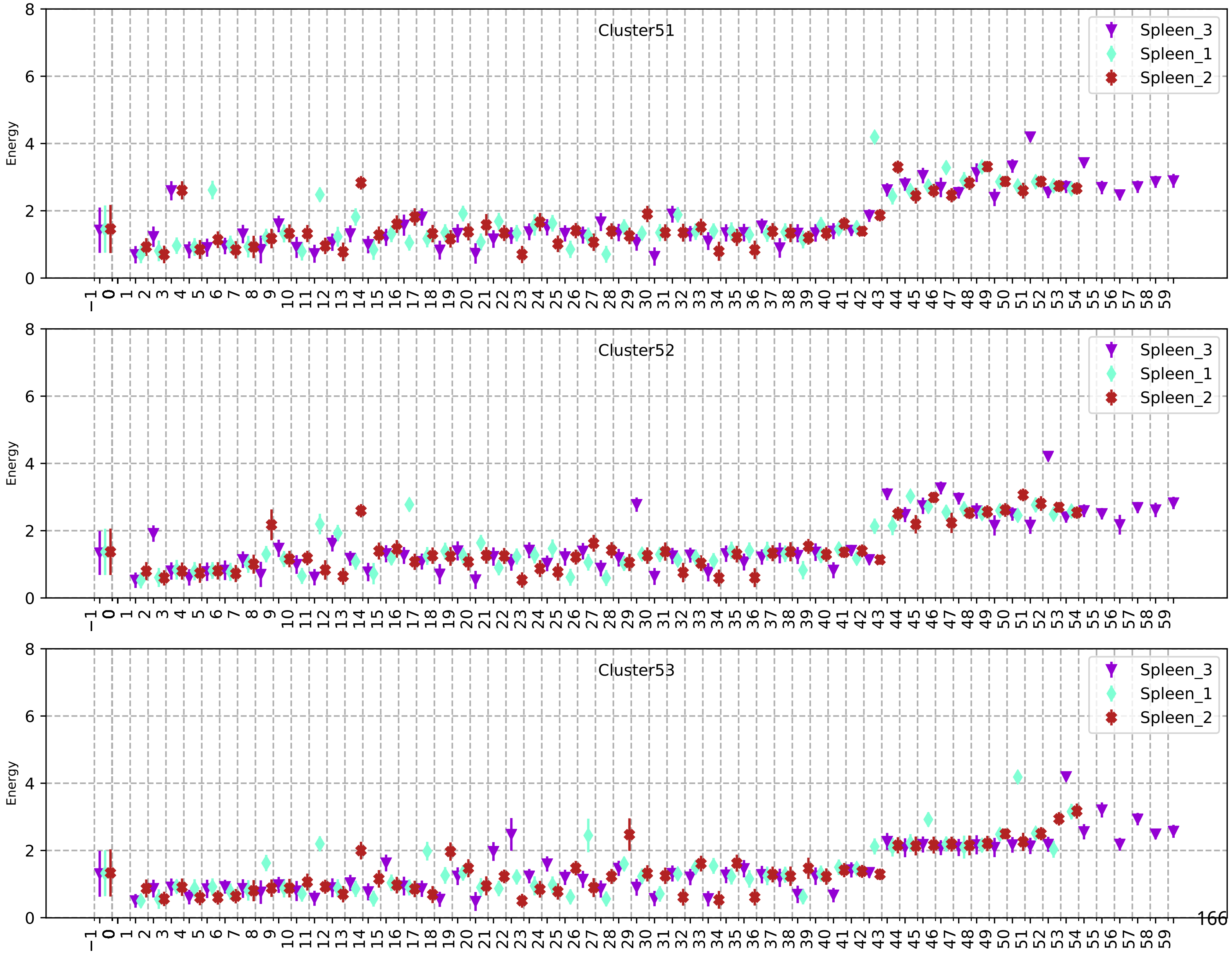
Cluster51
Spleen_3
Spleen_1
Spleen_2
Energy
Cluster52
Spleen_3
Spleen_1
Spleen_2
Energy
Cluster53
Spleen_3
Spleen_1
Spleen_2
Energy

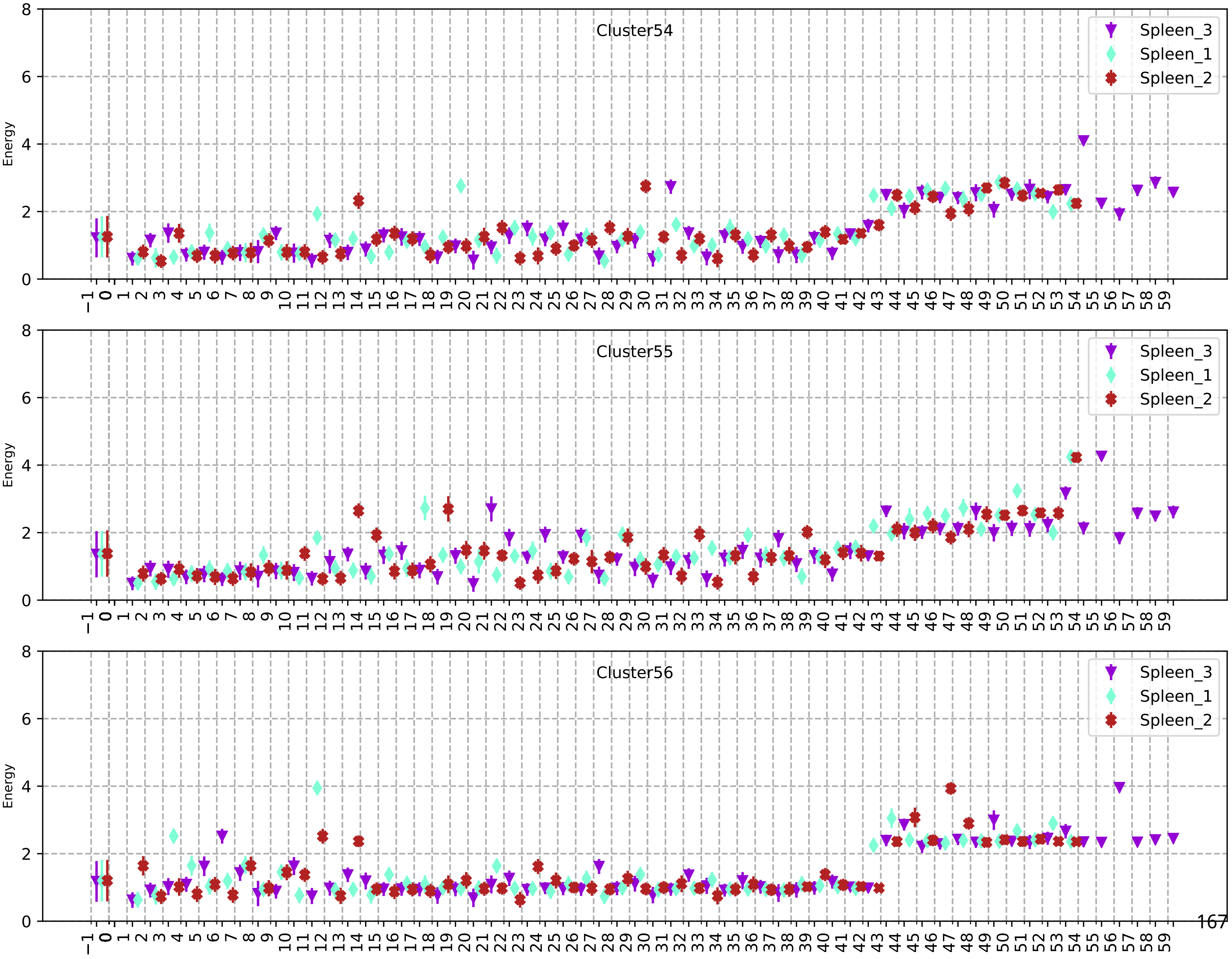
Cluster54
Cluster55
Cluster56
Spleen_3
Spleen_1
Spleen_2
Energy

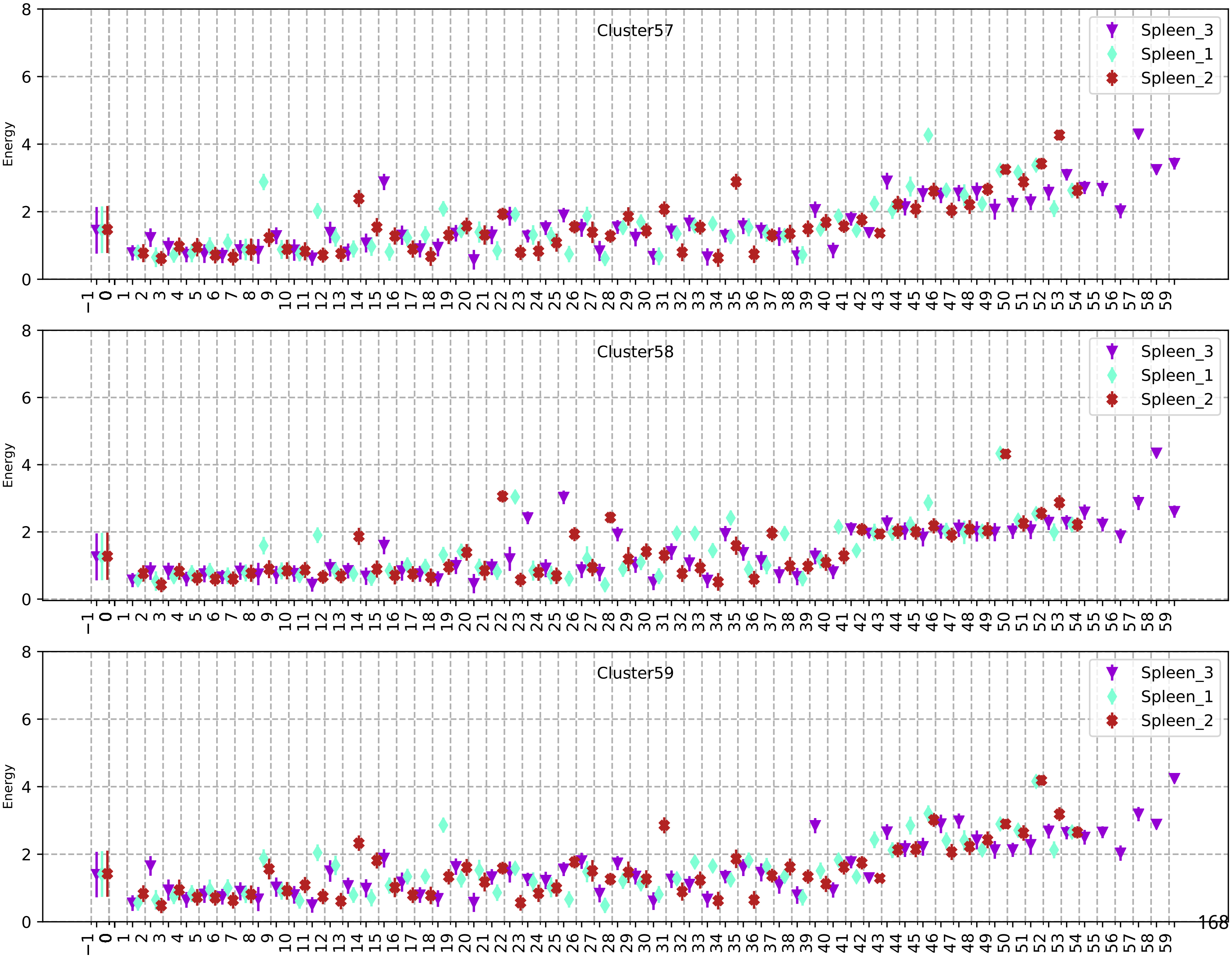
Cluster57
Cluster58
Cluster59
Energy
Spleen_3
Spleen_1
Spleen_2
